# Long COVID: G Protein-Coupled Receptors (GPCRs) responsible for persistent post-COVID symptoms

**DOI:** 10.1101/2022.12.12.520110

**Authors:** Sanisha Das, Suresh Kumar

**Affiliations:** Department of Diagnostic & Allied Health Science, Faculty of Health and Life Sciences, Management and Science University, Shah Alam, Selangor, Malaysia

**Keywords:** SARS-CoV-2, long COVID, COVID-19, biomarkers, therapeutic alternatives

## Abstract

As of early December 2022, COVID-19 had a significant impact on the lives of people all around the world, with over 630 million documented cases and over 6 million deaths. A recent clinical analysis revealed that under certain conditions, a patient’s disease symptoms are more likely to persist. Long COVID is characterised by many symptoms that continue long after the SARS-CoV-2 infection has resolved. This work utilised computational methods to analyse the persistence of COVID symptoms after recovery and to identify the relevant genes. Based on functional similarity, differentially expressed genes (DEGs) of SARS-CoV-2 infection and 255 symptoms of long covid were examined, and potential genes were identified based on the rank of functional similarity. Then, hub genes were identified by analysing the interactions between proteins. Using the identified key genes and the drug-gene interaction score, FDA drugs with potential for possible alternatives were identified. Also discovered were the gene ontology and pathways for 255 distinct symptoms. A website (https://longcovid.omicstutorials.com/) with a list of significant genes identified as biomarkers and potential treatments for each symptom was created. All of the hub genes associated with the symptoms, GNGT1, GNG12, GNB3, GNB4, GNG13, GNG8, GNG3, GNG7, GNG10, and GNAI1, were discovered to be associated with G-protein coupled receptors. This demonstrates that persistent COVID infection affects various organ systems and promotes chronic inflammation following infection. CTLA4, PTPN22, KIT, KRAS, NF1, RET, and CTNNB1 were identified as the common genes that regulate T-cell immunity via GPCR and cause a variety of symptoms, including autoimmunity, cardiovascular, dermatological, general symptoms, gastrointestinal, pulmonary, reproductive, genitourinary, and endocrine symptoms (RGEM). Among other functions, they were found to be involved in the positive regulation of protein localization to the cell cortex, the regulation of triglyceride metabolism, the binding of G protein-coupled receptors, the binding of G protein-coupled serotonin receptors, the heterotrimeric G-protein complex, and the cell cortex region. These biomarker data, together with the gene ontology and pathway information that accompanies them, are intended to aid in determining the cause and improving the efficacy of treatment.

## INTRODUCTION

The coronavirus disease 2019 (COVID-19) has had a huge impact on people’s lives globally, with over 630 million recorded cases and over 6 million deaths as of early December 2022 (Del Rio, Collins, & Malani, 2020). Severe Acute Respiratory Syndrome Coronavirus 2 (SARS-CoV-2) infects human cells through the angiotensin-converting enzyme 2 (ACE2) receptor on the cell membrane. Once within the body, the virus replicates and matures, triggering an inflammatory response defined by the activation and infiltration of immune cells by a cytokine storm. The presence of the ACE2 receptor in numerous cells across the human body suggests that SARS-CoV-2 can affect multiple organs. (Crook, Raza, Nowell, Young, & Edison, 2021). Recent clinical research has revealed some cases in which the symptoms of this condition tend to stay in the affected individual. Before these discoveries, it was considered that symptoms that persisted, much alone worsened, were exceptional. This disorder has been termed long COVID, post-acute sequelae of COVID (PASC), or post-acute COVID-19 syndrome (PACS) by researchers since it is defined by the existence of a variety of persistent symptoms long after the acute SARS-CoV-2 infection (Deer et al., 2021).

Long COVID has been shown to affect patients of all ages, ranging from those with very mild symptoms to those with chronic or severe symptoms. Similar to acute COVID-19, chronic COVID can affect multiple organs or organ systems, including the cardiovascular, respiratory, neurological, musculoskeletal, and nervous systems, among others. The most frequently reported symptoms are cardiac or pulmonary abnormalities, sleep disturbances, emotional and mental disorders, dyspnea, muscle and joint pains, and cognitive impairment **(Table 3)**.

**TABLE 1:** Identified hub genes and repurposed FDA drugs for all symptoms.

| INDEX | HUB GENES (RANKED) | FDA-APPROVED REPURPOSED DRUGS |
| --- | --- | --- |
| 1 | GNGT1 | - |
| 2 | GNG12 | - |
| 3 | GNB3 | CLONIDINE, SILDENAFIL, TORSEMIDE, NORTRIPTYLINE |
| 4 | GNB4 | - |
| 5 | GNG13 | - |
| 6 | GNG8 | - |
| 7 | GNG3 | - |
| 8 | GNG7 | - |
| 9 | GNG10 | - |
| 10 | GNAI1 | - |

**TABLE 2:**
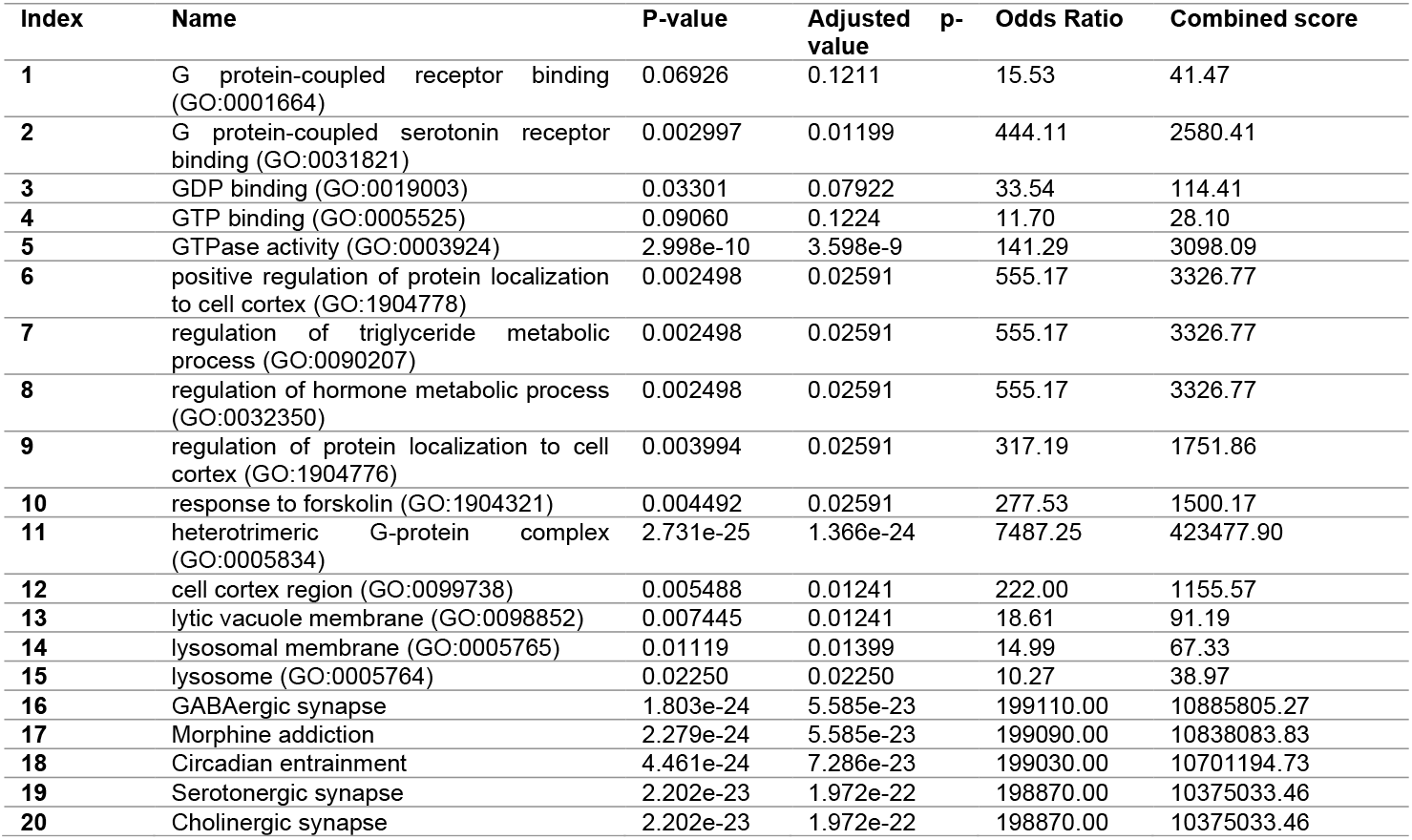
Gene enrichment for identified hub genes for all symptoms.

**TABLE 3:**
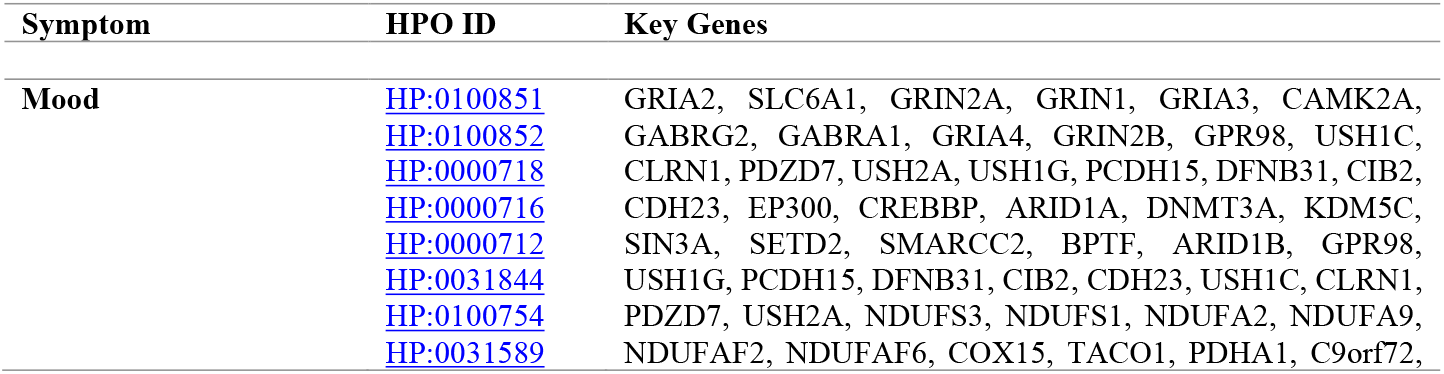
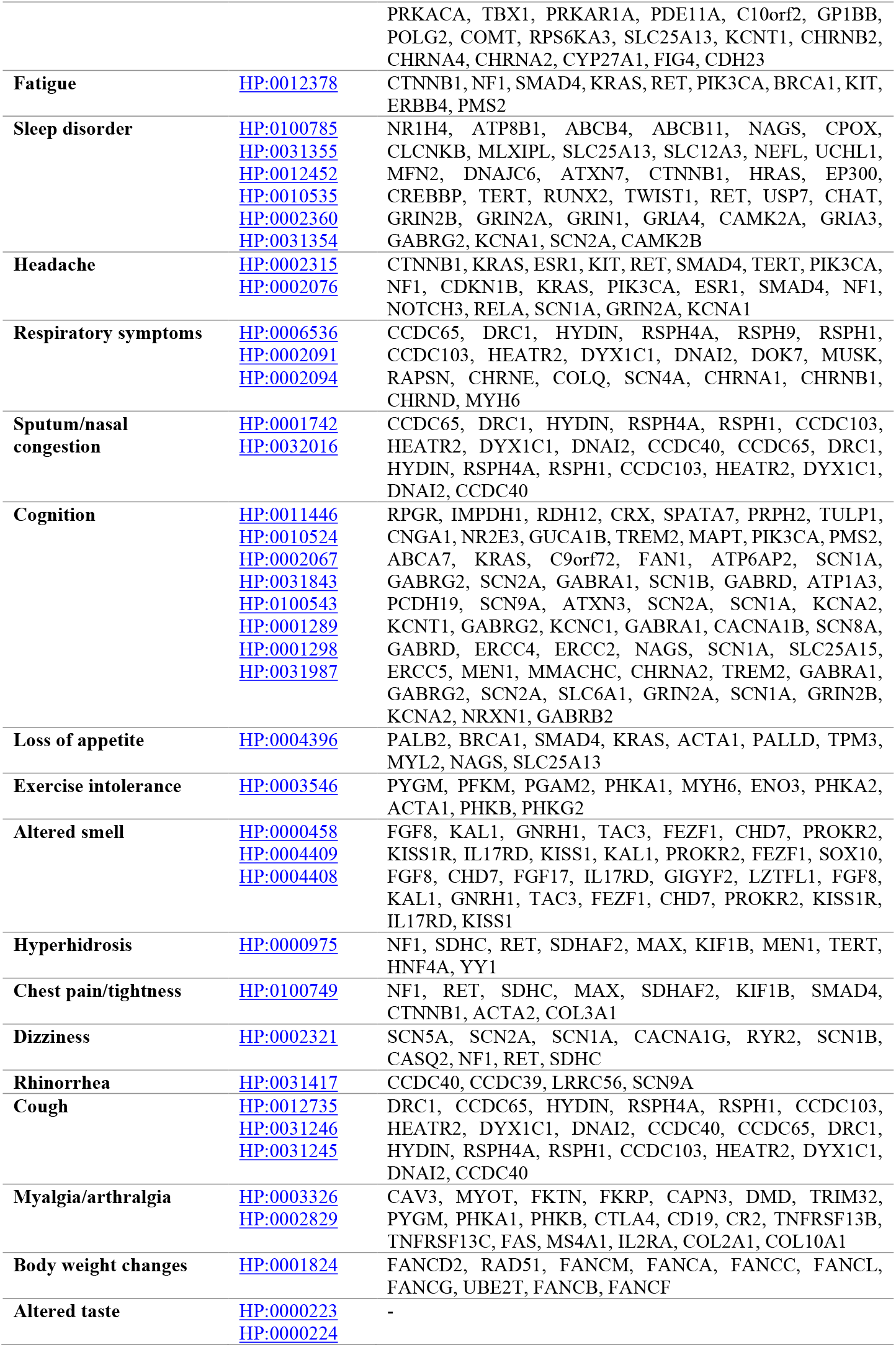
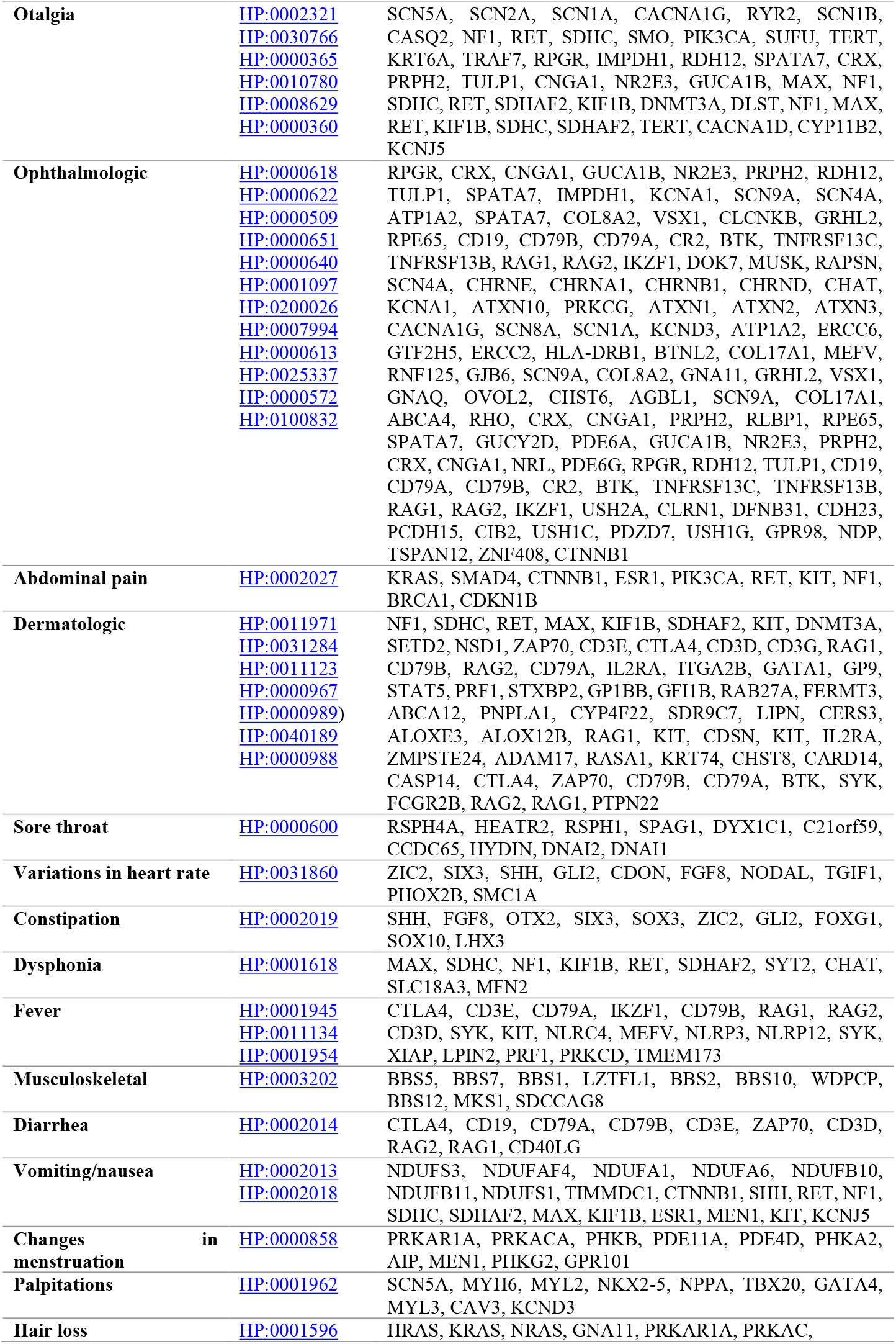
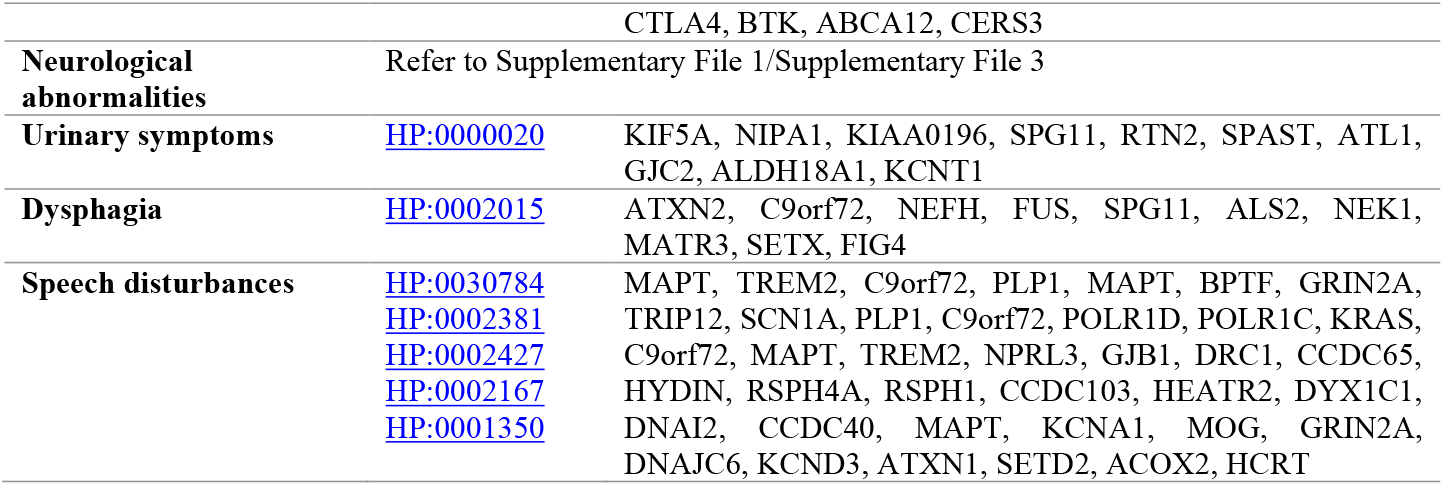
Identified common clinical manifestations in children and adolescents and respective hub genes.

Because the underlying causes and symptom patterns vary from individual to individual, it has become increasingly difficult for patients and medical professionals to identify and classify long COVID. To combat this, the definition of long COVID has been expanded to include the onset of new symptoms and chronic diseases (Deer et al., 2021), symptomatic infection (four to twelve weeks after initial acute infection), and symptoms developing twelve weeks after initial acute infection (Crook et al., 2021).

Several risk factors increase the likelihood that a patient who has recently recovered from acute COVID-19 will develop long-term COVID. To the COVID-19 rapid guideline written by the National Institute for Health and Care Excellence (NICE), the female gender, non-white and Asian ethnicities, poor pre-pandemic mental health, poor general health, the prevalence of asthma, overweight/obesity, smoking and/or vaping habits, and hospitalization history, to name a few. In addition, the number of acute symptoms not only determines the severity of the illness but also whether the patient will develop long-term COVID, the greater the number of symptoms, the greater the risk. Additionally, the presence of comorbidities preceding the progression of COVID-19 is a risk factor.

Although the advent of long-term symptoms following a SARS-CoV-2 infection may appear unexpected or unusual, it is a common occurrence of some viral infections. The persistent symptoms or chronic diseases of some infected people were linked to previously identified viral and bacterial infections (Proal & VanElzakker, 2021). For example, the Ebola virus has been associated with a severe illness that develops after the acute infection, with reservoirs still being discovered in human tissue years after viral clearance from the patient’s blood (Wilson et al., 2018). This example is analogous to long COVID remains long after the SARS-CoV-2 virus has been eliminated from the blood of a “recovered” patient.

After the acute stage of infection and risk factors that make a person more susceptible to developing long COVID symptoms, the future management of patients and evaluation of their treatment options will depend on additional research into the long-term impact of long COVID symptoms (Brown, Yahyouche, Haroon, Camaradou, & Turner, 2022).

## MATERIALS AND METHODS

### Dataset Retrieval and Construction

We have searched through all publications that reported for long COVID symptoms-systematic reviews, meta-analyses, and other publications (citing above all articles) from public repositories such as PubMed, LitCovid database, Embase database, etc. and have collected 255 symptoms of long COVID displayed in patients, published in peer-reviewed journals.

A gene list for the SARS-CoV-2 virus was created using data retrieved from the SARS-CoV-2 Infection Database (https://sarscovidb.org/), containing all differentially expressed genes (DEGs) identified after the SARS-CoV-2 infection, mined from various published articles in renowned scientific repositories, and H2V (http://www.datjar.com:40090/h2v/), which is a database containing all human proteins/genes that respond to SARS-CoV-2, SARS-CoV, and MERS-CoV. For this investigation, only the DEGs from each database were collected and merged.

Human Phenotype Ontology (HPO) concepts are increasingly being used to aid in the definition of patient phenotypes in diagnostic settings. Many HPO keywords are now mapped to putative causative genes with binary associations, and the HPO annotation database is routinely updated to offer precise phenotype information on a wide range of human diseases. Individual gene associations were extracted from the Human Phenotype Ontology (HPO) database (https://hpo.jas.org/app/) for each of the 255 symptoms included in this study. Each HPO ID corresponds to a phenotypic abnormality. After retrieving the individual gene lists, each symptom was categorized according to the organ system that was reported. The categories include autoimmunity, cardiovascular, dermatological, gastrointestinal, general symptoms, head, eyes, ears, nose, and throat (HEENT), lab, neuropsychiatric, pulmonary, and reproductive-genitourinary-endocrinological metabolism **(Figure 1)**.

**FIGURE 1:**
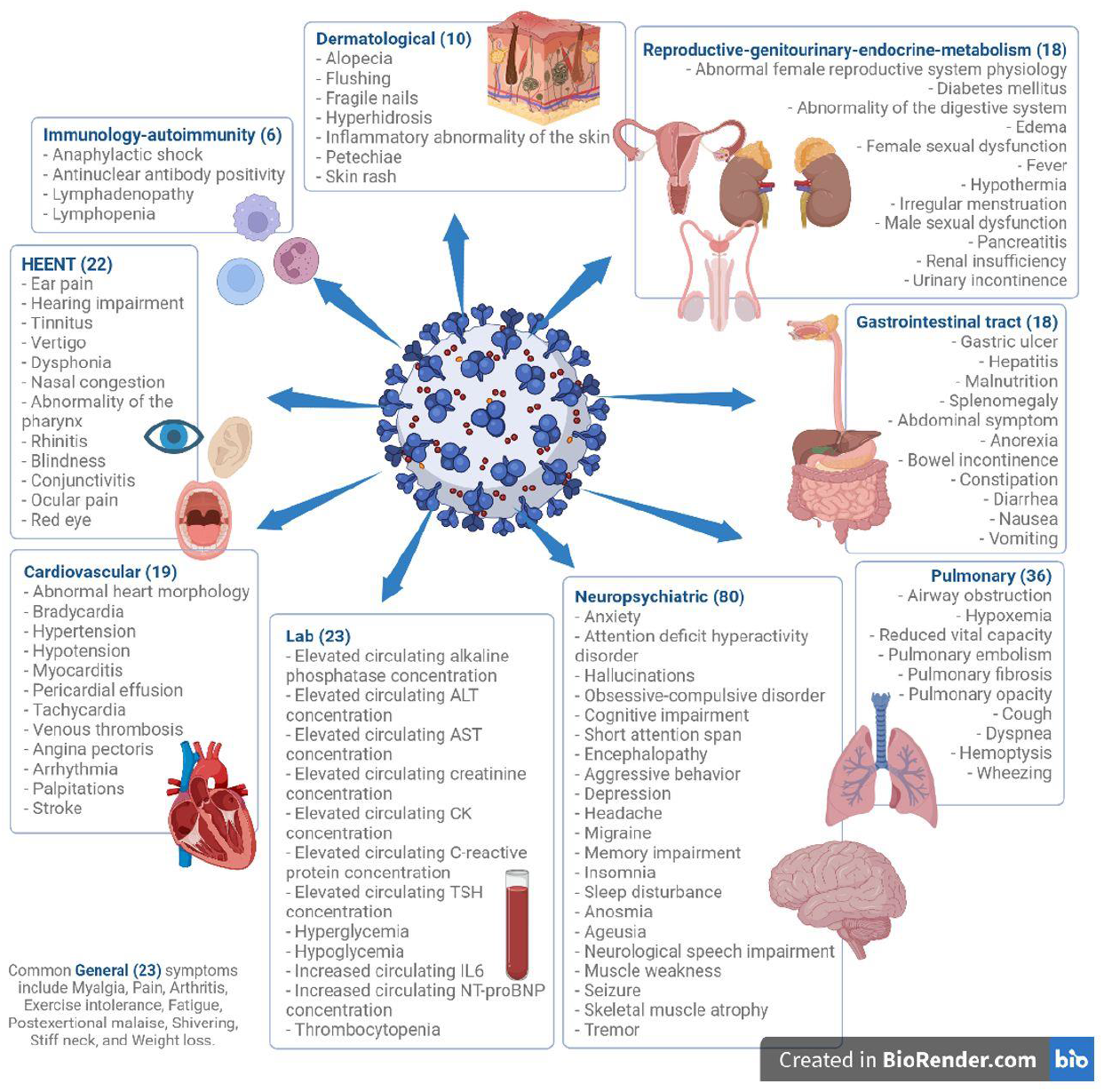
The schematic diagram of reported 255 symptoms of long covid.

### Gene Ranking

ToppGene Suite (https://toppgene.cchmc.org/), a portal for gene list enrichment analysis and candidate gene ranking based on functional annotations and protein interaction networks, was used to compare each symptom dataset to the SARS-CoV-2 dataset. The training gene set was comprised of SARS-CoV-2 DEG (differentially expressed genes), while the test gene set was comprised of symptom genes. GO: Molecular function, GO: Cellular component, GO: Biological process, Human phenotype, Mouse phenotype, Pathway, PubMed, Interaction, co-expression atlas (human protein atlas), ToppCell atlas, and Disease were selected as training parameters. The following sub-parameters were chosen for the ToppCell atlas:

Bronchoalveolar lavage atlas of COVID-19 patients, bronchoalveolar lavage atlas of severe obstructive pulmonary disease COVID-19 patients, COVID-19 patients’ CD8+ memory T cells, COVID-19 autopsy atlas (lung, liver, kidney, heart), COVID-19 B cell and plasma cell atlas in PBMC and BAL, COVID-19 autopsy atlas (lung, liver, kidney, heart), COVID-19 BAL atlas, COVID-19 leukocytes derived from cerebrospinal fluid, COVID-19 lung atlas, COVID-19 lung autopsy data, COVID-19 PBMC myeloid cell atlas, COVID-19 PBMC neutrophil cell atlas, and COVID-19 PBMC platelet cell atlas, COVID-19 cDC atlas, COVID-19 T cell atlas (PBMC), and COVID-19 T cell atlas (BAL).

Combining five COVID-19 peripheral blood mononuclear cell (PBMC) datasets, Integration of multiple COVID-19 patient sampling locations Large-scale integration of immune-mediated diseases (COVID-19 + Influenza + Sepsis + multiple sclerosis) COVID-19 single-cell data, PBMC atlas of patients with COVID-19, and PBMC atlas of patients with COVID-19 and influenza. Human cell lines infected with SARS-CoV-2, upper airway, and bronchi atlas of COVID-19 patients.

### Individual symptoms-Key genes identification using Protein-Protein Interaction analysis

The resulting prioritized genes were recorded from the Toppgene suite and further studied for hub genes using network analysis using Cytoscape.

The prioritized gene lists for each of the 255 long covid symptoms were, in turn, analyzed through Protein-Protein network analysis and the key genes were analyzed using CytoHubba (Chin et al., 2014) tool, which is a plugin in Cytoscape. It was used to rank the top 10 nodes for each symptom, from their respective STRING protein-protein interaction networks. CytoHubba uses 11 topological analysis methods, which cover Degree, Edge Percolated Component, Maximum Neighbourhood Component, Density of Maximum Neighbourhood Component, and Maximal Clique Centrality (six centralities). For this investigation, MCC was used.

### All 255 symptoms Gene prioritization based on phenotype and hub gene identification

It has been reported that Long Covid has multiple symptoms that last longer. For this aspect, we are hoping to analyze all reported symptoms that can be used to infer prioritized genes and to comprehend the underlying mechanism.

The HPO IDs retrieved from the HPO database were entered into the Phen2Gene tool (https://phen2gene.wglab.org/), which is a real-time phenotype-based gene prioritization tool using HPO IDs. Using the default weight model criterion, which weights HPO terms by skewness, Phen2Gene ranked all the genes; the top 1000 genes as test set genes were then prioritized using functional similarity analysis against the training gene set of SARS-CoV-2 DEG (differentially expressed genes) in ToppGene Suite. The resulting prioritized genes from the Toppgene suite were analyzed using protein-protein interaction analysis for the identification of hub genes using Cytoscape. Cytohubba was used to identify the key top 10 genes in the network.

### Identification of FDA of approved drugs for drug repurposing

The Drug Gene Interaction Database-DGIdb ((https://www.dgidb.org/), a web resource that provides information on drug-gene interactions and druggable genes from articles, databases, and other web-based sources, was then used to enter each set of hub genes for all 255 symptoms as a list. This was done to identify at least one drug approved by the US Food and Drug Administration for each gene. The DGIdb compiled information from 22 different databases, 43 different gene categories, and 31 different types of interactions. Only drugs with an interaction value of more than 0.8 with the genes were included in the results **(Supplementary File 1)**.

### Gene Enrichment Analysis

Gene enrichment or functional enrichment refers to identifying enriched or overrepresented genes in a list of ranked genes, that have an association with a particular disease and its phenotypes. For this stage, gene ontology tables on biological processes, molecular function, and cellular components and KEGG pathway tables for each set of hub genes were retrieved from Enrichr (https://maayanlab.cloud/Enrichr/), a suite of gene set enrichment analysis tools (**Supplementary File 2:** individual symptoms, Table 2: overall symptoms).

## RESULTS

From various online repositories, a collection of systematic reviews, metaanalyses, and cohort studies were obtained and studied. A total of 255 symptoms **(Figure 1)** were identified, compiled, and analyzed further. As a result, gene lists corresponding to each symptom were downloaded from the Human Phenotype Ontology (HPO) database, along with the SARS-CoV-2 gene list. These lists were divided into ten categories based on the phenotypes with which the symptoms were associated.

The overall workflow is shown in **Figure 2**. Gene prioritization was performed utilizing the ToppGene Suite to rank the candidate genes we had collected against all the parameters associated with the SARS-CoV-2 virus and the COVID-19 infection, followed by a gene enrichment analysis utilizing Enrichr to identify the most frequent genes on the list. This eliminated the genes that did not have a high frequency and therefore did not likely influence the phenotype of a symptom as strongly as the enriched genes. The enriched genes were ranked using Cytoscape and the 11 scoring methods developed by Cytohubba, a Cytoscape plugin. The Drug Gene Interaction Database-DGIdb was used to find therapeutic suggestions approved by the US Food and Drug Administration for each ranked gene. Only drugs with a score of 0.8 or higher with each gene are included.

**FIGURE 2:**
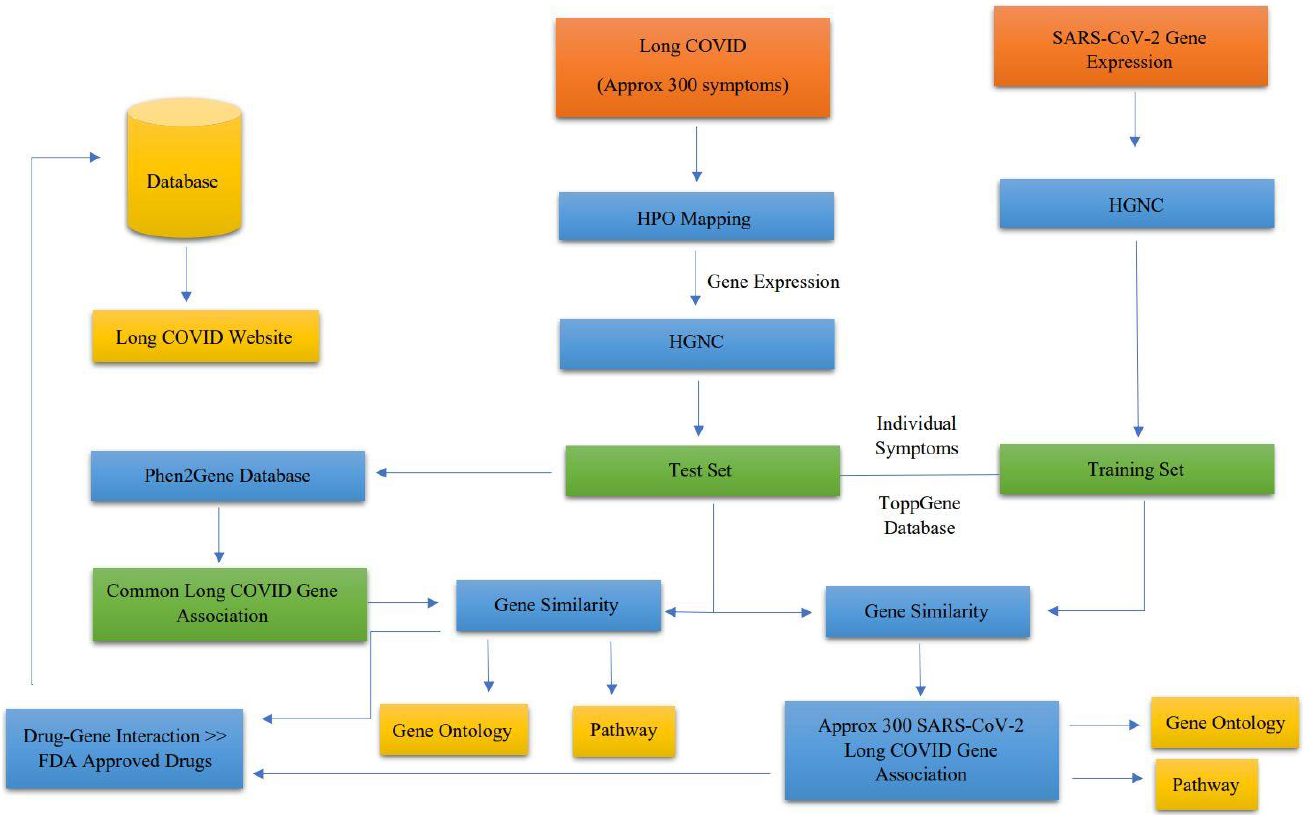
Overall workflow of identification of key genes of 255 reported long covid symptoms and identification of FDA-approved repurposed drugs.

## DISCUSSION

### Cardiovascular

Cardiovascular abnormalities such as abnormal morphology, abnormal heart rate, fluctuating blood pressure, and elevated troponin levels have been linked to an increased risk of death in patients hospitalized with SARS-CoV-2. As a result, cardiac issues that appear as a symptom of long COVID can be a serious problem for patients. Many people, including those who would be considered low risk, have ongoing myocardial inflammation, increased troponin levels, manifestations that lead to chest pain (angina pectoris), and other residual symptoms, according to cohort studies (Crook et al., 2021). The persistence could be attributed to residual viral remnants in the patient’s immune system, resulting in chronic inflammation or autoimmunity caused by an acute infection. The development of postural orthostatic tachycardia syndrome (POTS) is an example of autonomic dysfunction (Crook et al., 2021). As we will see later in this manuscript, autoimmunity is not limited to cardiovascular issues.

The European Society of Cardiology has published comprehensive guidelines for the assessment and treatment of the wide range of cardiovascular conditions that can manifest in a patient with long COVID. Several cohort studies and systematic reviews have highlighted the need to investigate specific cardiac biomarkers that can then be translated into potential therapeutic options; these biomarkers have been identified in our study. The National Institute for Health and Care Excellence (NICE) guidelines (NICE) et al., 2020) have recommended blockers for use against angina pectoris, cardiac arrhythmias, hypertension, and other immunotherapies to aid in recovery until more concrete treatment methods are determined with the help of the identified biomarkers.

SCN5A, MYBPC3, and MYH6 are the frequent key genes associated with cardiovascular symptoms in long covid patients. Brugada syndrome (BrS), progressive cardiac conduction disorder (Lev-Lenègre illness), dilated cardiomyopathy sick sinus syndrome, and atrial fibrillation have all been linked to SCN5A gene mutations that cause a loss of function. A common cause of familial hypertrophic cardiomyopathy is the MYBPC3 gene. This gene has been identified as the cause of left ventricular noncompaction, which occurs when the lower left chamber of the heart (left ventricle) fails to grow properly. The MYH6 gene has been linked to an increased incidence of sick sinus syndrome. The hub genes (**Supplementary File 1**: Tables 10 & 11) for cardiovascular symptoms for long covid patients were seen as most commonly involved in the pathways for African trypanosomiasis, Adrenergic signaling in cardiomyocytes, Cardiac muscle contraction, Circadian entrainment, Regulation of lipolysis in adipocytes, Viral myocarditis, and cGMP-PKG signaling pathway, which is responsible for the regulation of vascular smooth muscle cell relaxation and contraction, anti-cardiac hypertrophy, anti-atherosclerosis, and anti-vascular injury/restenosis. Adrenergic signaling works to control the contraction of the heart by working on adrenaline and noradrenaline receptors, therefore a malfunction in its receptors can lead to heart failure. Another important pathway is the regulation of lipolysis in adipocytes since if lipids are not adequately broken down in the adipocytes in cardiac muscle, this could lead to a host of other heart-related problems, some as severe as a heart attack.

### Dermatological

Even though they are less frequently reported, cutaneous symptoms can be a significant problem. From brittle nails and skin inflammation to severe conditions such as alopecia and petechiae, few studies have closely monitored each individual who has been diagnosed with the aforementioned conditions. One to two months after the acute infection, the most prevalent symptoms were increased hair loss and skin rashes (Thuangtong et al., 2021).

Excessive dermatological abnormalities can be caused by two statistically significant variables. First, the amount of post-COVID effects on skin lesions and hair loss is determined by the severity of the acute infection. The visibility of the symptom correlates with the severity of the acute infection. Second, gender, which has been identified as a general risk factor for long-term COVID, can influence which subset of patients is more likely to experience symptoms. As a result, medical personnel must thoroughly monitor females before, during, and after hospitalisation. (Thuangtong et al., 2021).

Autoimmune disorders may result in a rash because they induce inflammation in skin cells. CTLA4, KIT, RAG1, and ABCA12 are the most prevalent genes among all dermatological symptoms. Another severe skin ailment caused by mutations in the ABCA12 gene is lamellar ichthyosis type 2. The hub genes (**Supplementary File 1**: Table 2) for dermatological symptoms reported for long covid patients were seen as most commonly involved in the pathways for Acute myeloid leukemia, Central carbon metabolism in cancer, MAPK, and Ras signaling pathway, and Melanogenesis.

### Gastrointestinal

According to Kariyawasam et al., 11.4-61.1% of infected patients have reported gastrointestinal symptoms. Anorexia, diarrhea, nausea, vomiting, and abdominal pain are among the most prevalent COVID-19 symptoms (Kariyawasam, Jayarajah, Riza, Abeysuriya, & Seneviratne, 2021). Other conditions include gastric ulcers, malnutrition, and enlarged liver and/or spleen.

The onset of gastrointestinal symptoms can occur at any time. They appear at the beginning of the infection, before other clinical characteristics, in some people, but appear later in most. Patients who displayed gastrointestinal issues were admitted to hospitals considerably slower than those displaying respiratory issues. There was a corresponding delay in diagnosis and treatment management by the healthcare professionals, and subsequently, those patients had to experience a more prolonged hospital stay because their discharge was postponed until the viral infection was marked cleared. It can be relatively easy to jump to the conclusion that the late discharge of patients with GI symptoms is due to delayed treatments, however, studies have shown that this could be due to a combination of factors. For one, patients with GI manifestations have been proven to have an increased viral load and viral replication rate. This means that the transmission rate from these individuals would be on the higher end of the spectrum too (Kariyawasam et al., 2021). The timing and combination of symptoms in each individual can be attributed to a variety of factors, including ethnicities, geographical differences, and any prior comorbidities, making it difficult for medical professionals to rely on a single diagnostic pathway, such as routine endoscopies, to identify bleeding episodes. To recognise long COVID at an early stage, it is necessary to remain vigilant and keenly aware of the probability of any of the symptoms manifesting.

The common key genes associated with gastrointestinal symptoms of long covid patients are ACTG2, CD79A, and CD79B. Mutations in the ACTG2 gene have been identified as the cause of megacystis-microcolon-intestinal hypoperistalsis syndrome (MMIHS). People with intestinal pseudo-obstruction, a disease that disrupts the smooth muscle contractions that transport food through the digestive tract, have the ACTG2 gene (peristalsis). The hub genes (**Supplementary File 1:** Tables 12 & 13) of gastrointestinal symptoms reported for long covid patients were seen as most commonly involved in the pathways for AGE-RAGE signaling pathway in diabetic complications, Colorectal cancer, Epstein-Barr virus infection, Gastric cancer, PD-L1 expression and PD-1 checkpoint pathway in cancer, Primary immunodeficiency, Th1, Th2, and Th17 cell differentiation.

### General

Due to the subjective nature of general and constitutional symptoms, the reporting of these symptoms has varied considerably between investigations and studies. This indicates that the appearance of these symptoms can be easily misinterpreted by patients and misdiagnosed by physicians as any other illness/disease besides long COVID. This category contains some of the most common and widely reported symptoms (Groff et al., 2021). Among the symptoms are fatigue, joint inflammation, weight loss, xerostomia, myalgia, and chest pain. Persistent fever has also been a recurrent symptom; however, after approximately 60 days, the levels decrease. Groff et al. state that high fever rates may be caused by abnormally elevated anti-SARS-CoV-2-immunoglobulin G levels, which are commonly associated with disease severity, particularly in patients who are frontline medical personnel.

Fatigue has garnered a great deal of attention because infected populations around the world have realized that it is much more than simply being overtired; it is a state of perpetual exhaustion and weariness that negatively affects a person’s energy level, motivation, and ability to concentrate. 92.9 percent and 93.5 percent of admitted and nonadmitted patients, respectively, reported persistent fatigue 79 days after the onset of an acute infection, according to a cross-sectional study. Several additional studies have found associations between long-term COVID and fatigue as a common symptom, drawing parallels between long-term COVID fatigue and myalgic encephalomyelitis/chronic fatigue syndrome (ME/CFS) (Crook et al., 2021). Both conditions share similar manifestations, including fatigue, pain, neuropsychiatric, and autoimmune symptoms, with a significant number of patients suffering from impaired mobility and post-exertional malaise. The systemic review conducted by Wong et al. includes a quantitative analysis in which symptoms from 21 studies were compared to ME/CFS symptoms in general. One study on long-term COVID revealed that 25 of the 29 ME/CFS symptoms were present. This is a strong indication that additional research is required, given the examples of such preliminary studies that suggest overlapping symptoms, as well as the need for close monitoring and individualized treatment plans (Wong & Weitzer, 2021).

The common genes associated with general symptoms reported among long covid patients are SMAD4, COL2A1, COL9A3, and CTNNB1. The hub genes (**Supplementary File 1:** Tables 8 & 9) reported for general symptoms of long covid patients are associated most with pathways of cancer such as Breast cancer, Chemical carcinogenesis, Chronic myeloid leukemia, Endometrial cancer, Pancreatic cancer, Thyroid cancer, and signaling pathways such as Apelin signaling pathway, Calcium signaling pathway, ECM-receptor interaction, Glucagon signaling pathway, Insulin signaling pathway, Signaling pathways regulating pluripotency of stem cells, Thyroid hormone signaling pathway, and cGMP-PKG signaling pathway to name a few.

### HEENT (Head, Ears, Nose, Throat, Eyes)

In addition to being the primary sites for collecting samples for COVID-19 testing, nasal, nasopharyngeal, and/or oropharyngeal tissues are also the primary sites of infection. However, most of the literature and publications on COVID-19 focus primarily on the sequelae of the lower respiratory tract because these manifestations are relatively more lethal. This category of symptoms can be further subdivided into ears, ENT, and eyes (El Anwar, Eesa, Mansour, Zake, & Hendawy, 2021). Auditory and vestibular systems are predominantly located in the brainstem. Therefore, brainstem disruption caused by SARS-CoV-2-induced inflammation (Jafari, Kolb, & Mohajerani, 2022) can result in pharynx abnormalities, sensory impairments such as hearing impairment, hyperacusis, tinnitus, and vertigo, ocular involvement such as conjunctivitis and blurred vision, and motor deficits, but is not limited to these conditions.

The common key genes found in HEENT symptoms in long COVID patients are CNGA1, CRX, PRPH2, and SPATA7 are associated with both central and peripheral retinal degenerations. The hub genes (**Supplementary File 1:** Tables 4, 5 & 6) for head, ears, nose, throat, and eyes reported symptoms of long covid patients are observed most frequently in the pathways for Phototransduction, Retinol metabolism, Cortisol synthesis, and secretion, Cholinergic synapse, Basal cell carcinoma, cAMP signaling pathway, and cGMP-PKG signaling pathway.

### Lab

COVID-19, which is caused by SARS-CoV-2, is characterized by a broad spectrum of symptoms, a cytokine storm, and multi-organ failure. Cytokine storms have also been observed in SARS-CoV-infected patients, resulting in a poor prognosis. This suggests that a cytokine storm may play a role in the pathogenesis of COVID-19, as serum interleukin-6 levels are strongly correlated with a shorter life expectancy. In addition to elevated systemic cytokine levels and activated immune cells, COVID-19 exhibits several clinical and laboratory abnormalities, such as elevated CRP and D-dimer levels, renal failure, and pericardial effusions, as is typical of cytokine storm disorders (Fajgenbaum & June, 2020). SLC2A3, TTC26, and CASR are the main important genes detected in COVID patients with long-term symptoms. Fanconi-Bickel Syndrome and Type 2 Diabetes Mellitus are two diseases related to SLC2A2. Hydrocephalus and Biliary, Renal, Neurologic, and Skeletal Syndrome are diseases related to TTC26. The CASR gene is associated with type 1 autosomal dominant hypocalcemia, which is characterised by low blood calcium levels (hypocalcemia). The symptoms identified (**Supplementary File 1**: Table 7) are reported for lab-related symptoms which occur in long covid patients.

Interestingly, abnormal concentrations of serum cytokines and hormones have been reported as findings in patients with long COVID. In a meta-analysis done by Malik et al., it was found that specific biomarkers included lymphocytopenia, lower platelet count, and higher C-reactive protein (CRP), creatine kinase (CK), D-dimer, lactate dehydrogenase (LDH), aspartate transaminase (AST), aminotransferase (ALT), and creatinine levels. Lymphocytes play an important part in immunological homeostasis and the inflammatory response that protects the body against viral infections. Most of the papers included in this meta-analysis revealed that one of the hallmarks of SARS-CoV-2 infection is a decrease in the number of lymphocytes, which is associated with poor outcomes.

In long COVID patients, elevated D-dimer levels are related to a threefold increase in the likelihood of poor outcomes. The enhanced inflammatory response in COVID-19, combined with hypoxia caused by severe pneumonia, eventually activates coagulation and fibrinolysis, resulting in a hypercoagulable condition that causes multiorgan failure. When hepatocytes are damaged, aspartate transaminase (AST) and alanine aminotransferase (ALT) is released, resulting in elevated serum levels (abnormal liver function). In a study conducted by Cai et al, 76.3% of the 417 COVID-19 patients had unusual liver tests, while 21.5% acquired liver injury during hospitalization, characterized by ALT, AST, total bilirubin, and gamma-glutamyltransferase values rose to more than three times the upper limit of normal (Cai et al., 2020). Patients with abnormal liver tests had a considerably increased risk of acquiring severe pneumonia, and subsequently, long COVID. LDH is found in all tissues and is engaged in the interconversion of pyruvate and lactate via a nicotinamide adenine dinucleotide (NADH)-dependent process. Abnormal LDH levels can occur because of reduced oxygenation, which causes an upregulation of the glycolytic pathway and various organ damage. The enzyme CK is found in numerous tissues throughout the body, including the heart, brain, and skeletal muscle. Elevated CK levels in the blood are a sign of muscle injury. Malik et al. discovered an increase in poor outcomes in patients with increased CK levels (Malik et al., 2021).

### Immunology-autoimmunity

Pulmonary microangiopathy with signs of fibrin thrombi, active platelets, and neutrophil extracellular traps in arteries is a key histological characteristic of severe COVID-19. An influx of neutrophils, monocytes, and macrophages has also been found in various other organs other than the lungs, the heart, the central nervous system, and the liver (Knight et al., 2021). This serves as an indication that autoimmunity also contributes to multi-system organ failure in a person with developed long COVID. Some of the clinical characteristics displayed by patients suffering from moderate to severe COVID-19 are similar to those seen in autoimmune disorders. Furthermore, there have been multiple instances of patients developing well-defined autoimmune illnesses such as arthritis and Type 1 diabetes concurrently with or shortly following an acute SARS-CoV-2 infection. These findings have prompted researchers to wonder whether de novo autoimmunity may play a role in the etiology of long COVID (Knight et al., 2021).

At 12 months after the onset of COVID-19 symptoms, 43.6% of patients had antinuclear antibody titers in their blood, according to a recent study conducted with 96 individuals. The frequency of neurocognitive symptoms in the cohort was substantially higher in the group with antinuclear antibody titers than in the group without. Viruses that cause autoimmunity have various characteristics including the ability to tilt the host’s immune response to less tolerance through the creation of autoreactive lymphocytes, as seen in the cohort study. This association provides some evidence to support the claim that long COVID is due to viral residue in the individual’s body.

Although immune dysregulation has been linked to PASC, empirical evidence is lacking (Su et al., 2022). Existing questions include: Do pre-existing autoantibodies increase the likelihood that a person with COVID-19 will develop PASC? What is the prevalence of de novo autoantibodies during the acute phase of SARS-CoV-2 infection, and will these patients develop an autoimmune disease? Moreover, is virus-induced autoreactivity accountable for at least a portion of the diverse clinical manifestations associated with PASC? Our investigation uncovered identifiable biomarkers that can aid in answering at least one of the questions.

CTLA4, IL2R, and CD19 are the most common important genes implicated in autoimmune symptoms. CTLA-4 or CTLA4, also known as CD152, is a protein receptor that modulates immunological responses and operates as an immune checkpoint. Cytotoxic T-Lymphocyte Antigen 4 (CTLA-4) or CD152 is an inhibitory molecule essential for the maintenance of self-antigen tolerance. CTLA-4 has been linked to several autoimmune disorders, including systemic lupus erythematosus (SLE). SLE is an autoimmune disorder in which the immune system targets the body’s tissues, resulting in extensive inflammation and tissue destruction in the afflicted organs. It can impact the joints, the skin, the brain, the lungs, the kidneys, and the blood vessels.

Patients with severe Coronavirus Disease 2019 (COVID-19), caused by the new severe acute respiratory syndrome coronavirus 2 (SARS-CoV-2), had considerably raised plasma levels of IL2R. Dysfunction of regulatory T cells (Tregs) may be a main, causative event or the outcome of immune system perturbations during disease progression. Polymorphisms in genes related to Treg function, such as IL2RA, are associated with an increased risk of autoimmune illness. This pathogenic autoimmunity is the direct outcome of a breakdown in immune control in Type 1 diabetes. CD19 has also been linked to autoimmune illnesses, such as rheumatoid arthritis and multiple sclerosis. The hub genes (**Supplementary File 1**: Tables 1) for autoimmunity-related symptoms reported for long covid patients were seen as most involved in the pathways for Allograft rejection, Autoimmune thyroid disease, Graft-versus-host disease, Natural killer cell-mediated cytotoxicity, and T cell receptor signaling pathway. We can note from this that the genes play a role in causing the host immune system’s unique set of HLA proteins to build a response against the host’s cells which is recognized as foreign.

### Neuropsychiatric

It has been noted that COVID-19 patients have also frequently experienced neurological symptoms related to speech and language, cognitive dysfunction, headaches, smell and taste, emotions and mood, memory, behavior, and sleep. Neurologic abnormalities were found in 36.4% of 214 hospitalized COVID-19 patients in Wuhan, China, which had a higher percentage of patients with severe infections, at approximately 45.5%. In a cohort study of 236,379 patients that studied neurological and psychiatric outcomes after 6 months, 33.6% had average manifestations, 38.73% were hospitalized, 46.42% were admitted into intensive therapy, and 62.34% had encephalopathy. Likewise, neurologic results were found in 84.5% of infected patients who were hospitalized in France (Taquet et al., 2021).

The common key genes identified in Neuropsychiatric symptoms in long covid patients are SCN1A, C9orf72, GABRG2, and GRIN2A. The SCN1A gene is one of the most frequently mutated genes related to epilepsy in the human genome. Significantly, C9orf72 mutations are the first pathogenic mechanism discovered as a genetic connection between familial frontotemporal dementia (FTD) and amyotrophic lateral sclerosis (ALS) (ALS). This gene has been linked to epilepsy and febrile seizures due to mutations. Multiple neurological problems, including epilepsy, intellectual impairment, autism spectrum disorders, developmental delay, and schizophrenia, are associated with GRIN2A mutations. The symptoms reported related to neuropsychiatric symptoms for long covid patients are more than 60 related symptoms. (**Supplementary File 1:** Tables 17 to 25)

Zhang et. al. conducted an extensive study where expression of the viral proteins and viral particles were found in neurospheres and brain organoids infected with SARS-CoV-2, implying that SARS-CoV-2 can infect the brain effectively (Zhang et al., 2020). Fotuhi et. al. outlines the different ways the brain can get impacted during a wave of acute infection and how this impact varies based on the severity of the infection (Fotuhi, Hachinski, & Whitehouse, 2009). Looking into this categorization, it is possible to draw a connection to the neuropsychiatric impact of long COVID, based on how the brain fared during the acute SARS-CoV-2 infection.

In a mild infection, the brain can be infected via the olfactory nerve or the bloodstream. The binding of the SARS-CoV-2 to ACE2 receptors is connected to the nasal and gustatory cells. The virus-induced cytokine storm, created as an inflammatory response, is in a controlled manner. Smell and taste abnormalities are usually treated without the need for any complex procedures (Fotuhi et al., 2009). In a moderate infection, the brain can be infected via the brainstem or the respiratory brain center. It is associated with the triggering of a significant immune response with high amounts of cytokines, which raises ferritin, C-reactive protein, and D-dimer concentrations. If the individual were to experience a stroke at this point, there would be a production of blood clots, blockage of arteries, or venous thrombosis; all of which have been noted to be symptoms of long COVID as well. This massive immune response can cause further damage to the central nervous system and/or the peripheral nervous system (Fotuhi et al., 2009). In a severe infection, the brain can be directly infected via the nerves connecting the pulmonary system and the brain. The cytokine storm generated as an inflammatory response is no longer controlled and disrupts the blood-brain barrier, allowing inflammatory agents and other components of the blood to filter into the cerebrospinal fluid. Encephalopathy, seizures, and tremors, arise from edema and brain damage. Hypertension increases the risk of cerebral bleeding (Fotuhi et al., 2009).

Cognitive disorders are mental health conditions that impair the cognitive capacities of an individual such as memory, motor function, language, attention, and social cognition. Hypoxemia, inflammatory reaction infection, and anesthetic medications can all cause cognitive impairment. Additionally, demyelinating diseases such as transverse myelitis, multiple sclerosis, acute disseminated encephalomyelitis (ADEM), and neuromyelitis optica spectrum disorder have been reported post-COVID.

Current neuropsychological testing and biochemical or neuroimaging evaluation are thought to aid in estimating the amount and course of neurological symptoms post-COVID. These investigations raised awareness of the necessity for earlier monitoring of cognition in COVID-19 patients, as well as the importance of prompt medical intervention if any disorientation is evident upon hospital admission. It is crucial to note that different cultures have varying executive function requirements; thus, disease effects on this cognitive trait can vary (Hosseini, Nadjafi, & Ashtary, 2021).

### Pulmonary

Adeloye et. al used the Child Health and Nutrition Research Initiative (CHNRI) method to carry out a research exercise focused on tackling the long COVID burden in patients who have experienced and survived an acute SARS-CoV-2 infection. In the results, although the emphasis was on airway illnesses, there are certain aspects listed that are largely relevant to the investigation of long COVID and therapeutic options in patients who were not confined to hospitals or those that had comorbidities.

The research centered on comparing manifestations in COVID-19 survivors with and without any previous airway-related illness. Fatigue, sarcopenia, anxiety, depression, cardiovascular problems, and hospital readmission are all comorbidities and consequences. Investigation of the incidence and risk variables (such as smoking and ethnicity) linked with acquiring new symptomatic airway disease or structural airway abnormalities on radiological imaging was also reported (Adeloye et al., 2021).

CCDC103, CCDC40, and CCDC65 are the common key genes linked with pulmonary symptoms in long covid patients. CCDC103 is connected with Ciliary Dyskinesia, Primary, 17, and Dextrocardia With Situs Inversus.CCDC40 associated with Kartagener’s syndrome is a rare autosomal recessive ciliary condition characterised by situs inversus, recurrent sinusitis, and bronchiectasis. The hub genes (**Supplementary File 1:** Tables 14, 15 & 16) of pulmonary-related symptoms for long covid patients were seen as most commonly involved in the pathways for Arrhythmogenic right ventricular cardiomyopathy, Asthma, Cell adhesion molecules, Cytokine-cytokine receptor interaction, Fluid shear stress and atherosclerosis, Huntington disease, Neuroactive ligand-receptor interaction, cGMP-PKG signaling pathway.

### Reproductive, Genitourinary, Endocrine, and Metabolism (RGEM)

Bowe et al’s cohort study assessed post-acute kidney outcomes in a cohort of 89,216 COVID-19 30-day survivors. The findings demonstrate that those who survived COVID-19 had a higher risk (and incidence) of AKI (acute kidney injury), eGFR (estimated glomerular filtration rate) decrease, ESKD (end-stage kidney disease), and MAKE (major adverse kidney events) after the initial 30 days of infection. These findings imply that persons with COVID-19 have a higher risk of bad renal outcomes beyond the acute phase of infection. People with COVID-19 should receive post-acute care that includes attention and treatment for both acute and chronic kidney disease. The reason or mechanisms behind the increased risk of AKI, eGFR decrease, ESKD, and MAKE in the post-acute phase of COVID-19 infection are unknown. Other plausible causes include dysregulated immunological response or autoimmunity, chronic inflammation, and autonomic nervous system disturbances. Mechanisms connected to changes in broader economic and social situations in the context of the worldwide pandemic, which may have affected people with COVID-19 differently, may potentially be at work (Bowe, Xie, Xu, & Al-Aly, 2021).

Bowe et. al highlights the urgent requirement for a better understanding of the molecular and epidemiologic drivers of the post-acute renal sequelae of SARS-CoV-2 infection to assist influence care options. Hence, our findings on the role of biomarkers responsible for these sequelae in the various gene ontologies and pathways, coupled with the suggested therapeutic options may shed light on future investigations.

Numerous endocrine glands exhibit elevated concentrations of ACE2 and TMPRSS2. This, along with multiple case reports of thyroid and pituitary dysfunction in COVID-19-infected patients, has generated significant interest in the virus’ endocrine consequences. Possible risk factors for coronavirus disease 2019 (COVID-19) infection on fertility include the presence of angiotensin-converting enzyme2 (ACE2) on testes, a drop in significant sex hormone ratios, and fever linked with COVID-19 infection. Recent research has demonstrated a discrepancy between men and women in terms of COVID-19 prevalence and comorbidities. SARS-CoV-2 also induces a cytokine storm, which is an excessive immune response with a broad spectrum of cytokine release that establishes a systemic proinflammatory milieu that may facilitate insulin resistance and beta cell hyperstimulation, leading to beta cell dysfunction and death. Additionally, SARS-CoV-2 may exacerbate the preexisting proinflammatory state observed in type 2 diabetes (T2D), hence lowering patient lifespan and exacerbating comorbidities. These abnormalities continue post-COVID-19 in COVID-19-recovered patients. Smaller, single-center investigations are actively investigating the impact of COVID-19 on the genitourinary system and initiating the creation of theories concerning the mechanism behind urine symptoms in COVID-19 patients. Women are more vulnerable to SARS-CoV-2 infection. SARS-CoV-2 can infect the follicular membrane and granular cells of the ovaries, lowering egg quality and causing female infertility. Chronic inflammation caused by SARS-CoV-2, as well as following long COVID, can pose serious risks to the ovaries and the hypothalamus-pituitary ovarian axis, compromising female reproductive structure and function. This, for example, could result in irregular menstruation. Because of their extensive virus exposure and high ACE2 expression in their reproductive system, women are especially sensitive to SARS-CoV-2 (Liu et al., 2021).

Males in the COVID-19 pandemic had significantly higher contagiousness and mortality rates than females, according to multiple studies. According to reports, SARS-CoV-2, like SARS-CoV, has not been found in testicular tissues. Male COVID-19 patients, on the other hand, suffer testicular tissue damage and obvious clinical symptoms. In men with long COVID, male genitourinary (GU) damage was also observed. Inflammation, fever, and hypoxia can all affect the male hypothalamic-pituitary-gonadal axis, testicular function, and spermatogenesis. One of the most prevalent symptoms in patients is fever, and high body temperatures are harmful to the testis, limiting spermatogenesis through a variety of processes (Liu et al., 2021).

FGF8, CHD7, and FEZF1 are the most often identified important genes in RGEM-related symptoms of long covid. People with Kallmann syndrome, a sickness characterised by hypogonadotropic hypogonadism (a condition affecting the synthesis of hormones that regulate sexual development) and an impaired sense of smell, have been shown to carry the CHD7 gene. Hypogonadotropic Hypogonadism 22 with or without anosmia and Kallmann Syndrome are related to FEZF1. The hub genes (**Supplementary File 1:** Table 3) for reproductive, genitourinary, endocrine, and metabolism-related reported symptoms of long covid patients were seen as most commonly involved in the pathways for NF-kappa B signaling pathway (regulator of innate immunity), Neuroactive ligand-receptor interaction, ECM-receptor interaction, GnRH signaling pathway, Propanoate metabolism, Protein digestion and absorption, Regulation of lipolysis in adipocytes, Purine metabolism, Tyrosine metabolism, Rap1 signaling pathway, and Chemical carcinogenesis.

### Role of G-protein-coupled receptors (GPCRs) in Long COVID symptoms

GPCRs may play a role in the immune response to viral infections, including SARS-CoV-2, the virus that causes COVID-19, according to some research. G-protein-coupled receptors (GPCRs) allow viruses to enter and infect host cells. GPCRs are proteins found on the cell surface and are involved in a variety of physiological processes, such as the perception of hormones, neurotransmitters, and other signalling molecules. Our ability to respond to environmental stimuli, such as taste and smell, indicates the need for a mechanism with a high degree of sensitivity. G protein-coupled receptors (GPCRs) are crucial to these activities. In addition, they regulate our emotions, behaviour, and immune system. GPCRs have been demonstrated to interact with other important physiological molecules, such as growth factors and their receptors, by generating transactivation signals that regulate cell motility. When a virus infects a host cell, it can use GPCRs as receptors to gain access to the cell. COVID-19 is caused by the SARS-CoV-2 virus, which uses the ACE2 receptor. AT1R is a clinically significant GPCR that binds to therenin–angiotensin system hormone angiotensin II (AngII). Signaling by the activated AT1R involves stimulation of the Gq11/12, intracellular production of calcium, and lipid messengers, and activation of protein kinases and a GPCR to enter and infect respiratory tract cells. Once a virus enters a cell, it can replicate and cause disease. In this way, the virus “hijacks” the GPCR to enter the cell. Notably, GPCRs are not the only receptors that viruses can use to enter host cells. Therefore, it makes sense that the hub genes identified for all of the collective symptoms included in our study are associated with GPCRs **(Table 1)**. G protein-coupled receptors (GPCRs) are a class of proteins that participate in numerous physiological processes, including T-cell modulation. These proteins are located on the cell surface and play a role in the transmission of signals from the cell’s exterior to its interior, where they can initiate a response.

T-cells are an essential type of white blood cell for the immune system. They contribute to the recognition and defence against microbial and viral invaders. T-cell activity can be modulated by the activation of GPCRs on the surface of T-cells by various molecules, such as hormones or cytokines.

Specifically, cytokines, which are immune response-coordinating proteins, can activate GPCRs on T-cells. Upon activation of these GPCRs, T-cells can proliferate and produce more cytokines, thereby enhancing the immune response.

In conclusion, GPCRs play a crucial role in T-cell modulation by transmitting signals from the cell’s exterior to its interior, where they can trigger a response that aids in immune response coordination. Endogenous self-reactive autoantibodies (AAs) recognise a variety of G-protein-coupled receptors (GPCRs). They are frequently associated with cardiovascular, neurological, and autoimmune diseases, and in some cases, they directly affect the progression of the disease. Numerous GPCR AAs modulate receptor signalling, but the molecular details of their activity are unknown.

T cell signalling proteins CTLA4 and PD-1, as well as the GPCR A2AR, can block the T cell receptor (TCR) and its accelerators via inhibitory allosteric receptor-receptor interactions in heteroreceptor complexes. Cells communicate in a variety of ways to perform their normal functions. Because of their ability to bind guanine nucleotide, GTP-binding proteins, or G proteins, are a key component of a signalling cascade. G proteins interact with and control a variety of effector molecules, including calcium, potassium channels, adenylyl cyclase, phospholipases C and D, and protein kinases. It is now known that both subunits regulate a vast number of effectors. These genes are GNAI1, GNGT1, GNG12, GNB3, GNB4, GNG13, GNG8, GNG3, GNG7, and GNG10 **(Figure 3)**. The ability of GPCRs to bind diverse ligands, change conformation, up- or down-regulate membrane expression, and interact with other proteins and receptors to form homo-or heterodimers contributes to their adaptability to changing environmental conditions. GPCRs, therefore, interact with a vast array of extracellular and intracellular proteins, including signalling molecules and other membrane proteins. (Legler & Thelen, 2018; Riemekasten, Petersen, & Heidecke, 2020) Approximately half of the over 800 human GPCRs are sensory receptors that regulate olfaction, light perception, and pheromone signalling. More than 90% of the remaining non-sensory GPCRs, numbering over 370, are expressed in the brain and mediate communication from a variety of ligands, thereby regulating a variety of physiological processes throughout the human organism, primarily endocrine and neurological functions. GPCRs are the most used therapeutic drug targets (De Oliveira, Ramos, Amaro, Dias, & Vieira, 2019).

**FIGURE 3:**
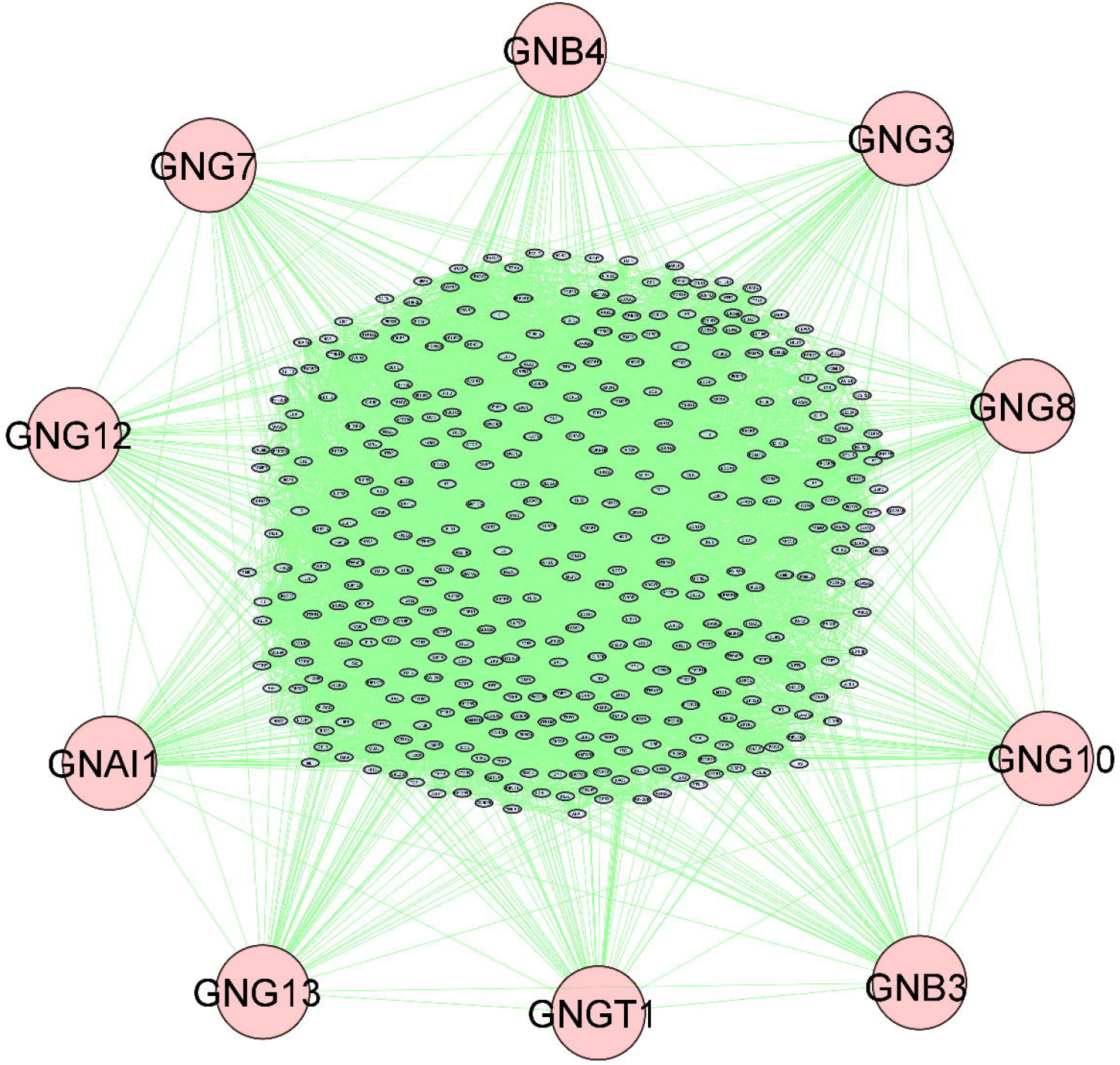
Protein-protein interaction network overview built using STRING in Cytoscape. Nodes represent proteins, edges represent the interaction between two nodes (proteins). The top 10 key hub genes of 255 symptoms of long covid are related to G-protein coupled receptors shown in circular nodes in pink.

At the subcellular level, Gi/o-coupled GPCRs can be detected and analysed for functions such as neurite outgrowth, neurotransmitter release, and synaptic plasticity, which are then processed into complex brain processes such as memory, learning, and cognition. Serotonin, dopamine, cannabinoid, and metabotropic glutamate receptors enhance short- and long-term memory, memory consolidation, and learning. The majority of GPCRs, including dopamine, serotonin, and opioid receptors, play an essential role in modulating and affecting behaviour and emotions. GPCR signalling is also crucial for sleep and circadian rhythm regulation (De Oliveira et al., 2019).

As shown in **Table 2**, we also investigated the pathways, biological processes, molecular functions, and cellular components that occur most frequently in each of the individual symptoms (Talla et al., 2022).

## CONCLUSION

Long COVID has been identified as a major worldwide health concern, but because research on this syndrome is still in its infancy, its prognosis for the future is impossible to predict. The identification of the 255 symptoms in the different organ systems, the biomarker discoveries, and the respective FDA-Approved repurposed drugs, as well as their associated gene ontology and pathway insights from this study, will assist in determining the cause of long COVID and personalising treatment by facilitating the diagnosis of long COVID in patients by healthcare professionals around the world.

## ACKNOWLEDGEMENT

The authors would like to acknowledge the Department of Diagnostic and Allied Health Sciences, Faculty of Health and Life Sciences of Management and Science University, Shah Alam, Selangor Darul Ehsan, Malaysia for giving us the opportunity and research facilities to conduct this research.

