## Supplementary Table 1-3 for "Long COVID: G Protein-Coupled Receptors (GPCRs) responsible for persistent post-COVID symptoms": Supplementary File 1- hub genes and FDA-approved drugs for 255 symptoms.docx

### TABLE 1: Identified hub genes and repurposed FDA drugs for 255 individual symptoms

### Table 1 Immunology-autoimmunity

| **Symptom (HPO ID)** | **Hub Genes (ranked)** | **FDA-Approved repurposed drugs** |
| --- | --- | --- |
| Anaphylactic shock ([HP:0100845](https://hpo.jax.org/app/browse/term/HP:0100845)) | CBL | - |
|  | KIT | NILOTINIB, IMATINIB |
|  | KITLG | - |
|  | GRB2 | - |
|  | EPOR | DARBEPOETIN ALFA, PEGINESATIDE ACETATE, EPOETIN BETA, CAPSAICIN |
|  | PTPN11 | - |
|  | NRAS | BINIMETINIB |
|  | PIK3R1 | - |
|  | HRAS | - |
|  | PTPN6 | TOFACITINIB |
| Antinuclear antibody positivity ([HP:0003493](https://hpo.jax.org/app/browse/term/HP:0003493)) | CTLA4 | IPILIMUMAB, ABATACEPT |
|  | IL2RA | BASILIXIMAB, DACLIZUMAB, ALDESLEUKIN, DENILEUKIN DIFTITOX, CETIRIZINE |
|  | FAS | TENIPOSIDE |
|  | FASLG | - |
|  | FCGR2B | - |
|  | RIPK1 | - |
|  | IL12RB1 | - |
|  | MS4A1 | TOSITUMOMAB, OFATUMUMAB, OBINUTUZUMAB, OCRELIZUMAB, IBRITUMOMAB TIUXETAN, YTTRIUM Y 90 IBRITUMOMAB TIUXETAN, RITUXIMAB |
|  | PTPN22 | - |
|  | ACP5 | FILGRASTIM, ALTEPLASE |
| Anti-thyroid peroxidase antibody positivity ([HP:0025379](https://hpo.jax.org/app/browse/term/HP:0025379)) | RASGRP1 | - |
|  | HRAS | - |
|  | KRAS | PANITUMUMAB, CETUXIMAB |
|  | NRAS | BINIMETINIB |
|  | RAP1B | - |
|  | RAP1A | - |
|  | PLCG1 | CEFIXIME, DICLOFENAC |
|  | PRKCQ | - |
|  | SKAP1 | - |
|  | DGKZ | - |
| Anti-thyroglobulin antibody positivity ([HP:0032069](https://hpo.jax.org/app/browse/term/HP:0032069)) | - | - |
| Lymphadenopathy ([HP:0002716](https://hpo.jax.org/app/browse/term/HP:0002716)) | CTLA4 | IPILIMUMAB, ABATACEPT |
|  | CD19 | BLINATUMOMAB |
|  | IL2RA | BASILIXIMAB, DACLIZUMAB, ALDESLEUKIN, DENILEUKIN DIFTITOX, CETIRIZINE |
|  | STAT5B | TOFACITINIB, DASATINIB, EPOETIN ALFA |
|  | PRF1 | EMAPALUMAB |
|  | FAS | TENIPOSIDE |
|  | ZAP70 | - |
|  | RAG1 | - |
|  | FASLG | - |
|  | SYK | FOSTAMATINIB |
| Lymphopenia ([HP:0001888](https://hpo.jax.org/app/browse/term/HP:0001888)) | CD19 | BLINATUMOMAB |
|  | CTLA4 | IPILIMUMAB, ABATACEPT |
|  | ZAP70 | - |
|  | CD3E | MUROMMONAB-CD3, CATUMAXOMAB, BLINATUMOMAB |
|  | RAG1 | - |
|  | RAG2 | - |
|  | CD3D | MUROMMONAB-CD3, CATUMAXOMAB, BLINATUMOMAB |
|  | IL2RA | BASILIXIMAB, DACLIZUMAB, ALDESLEUKIN, DENILEUKIN DIFTITOX, CETIRIZINE |
|  | CD3G | MUROMMONAB-CD3, CATUMAXOMAB, BLINATUMOMAB |
|  | IKZF1 | LENALIDOMIDE |

### Table 2 Dermatological

| **Symptom (HPO ID)** | **Hub Genes (ranked)** | **FDA-Approved repurposed drugs** |
| --- | --- | --- |
| Alopecia ([HP:0001596](https://hpo.jax.org/app/browse/term/HP:0001596)) | HRAS | - |
|  | KRAS | PANITUMUMAB, CETUXIMAB |
|  | NRAS | BINIMETINIB |
|  | GNA11 | BINIMETINIB, CABOZANTINIB |
|  | PRKAR1A | - |
|  | PRKACA | - |
|  | CTLA4 | IPILIMUMAB, ABATACEPT |
|  | BTK | IBRUTINIB |
|  | ABCA12 | - |
|  | CERS3 | - |
| Dermatographic urticaria ([HP:0011971](https://hpo.jax.org/app/browse/term/HP:0011971)) | - | - |
| Flushing ([HP:0031284](https://hpo.jax.org/app/browse/term/HP:0031284)) | NF1 | TRAMETINIB, BINIMETINIB, COBIMETINIB, DABRAFENIB, VEMURAFENIB |
|  | SDHC | - |
|  | RET | CABOZANTINIB, VANDETANIB |
|  | MAX | - |
|  | KIF1B | - |
|  | SDHAF2 | - |
|  | KIT | NILOTINIB, IMATINIB |
|  | DNMT3A | DECITABINE, IDARUBICIN, WARFARIN |
|  | SETD2 | - |
|  | NSD1 | - |
| Fragile nails ([HP:0001808](https://hpo.jax.org/app/browse/term/HP:0001808)) | COL17A1 | - |
|  | HRAS | - |
|  | ITGB4 | - |
|  | DSP | ENALAPRIL |
|  | EFNB1 | - |
|  | DLX3 | - |
|  | ERCC2 | OXALIPLATIN, LEUCOVORIN |
|  | GTF2H5 | - |
|  | CAMK2G | - |
|  | MSX1 | - |
| Hyperhidrosis ([HP:0000975](https://hpo.jax.org/app/browse/term/HP:0000975)) | NF1 | TRAMETINIB, BINIMETINIB, COBIMETINIB, DABRAFENIB, VEMURAFENIB |
|  | SDHC | - |
|  | RET | CABOZANTINIB, VANDETANIB |
|  | SDHAF2 | - |
|  | MAX | - |
|  | KIF1B | - |
|  | MEN1 | OLSALAZINE |
|  | TERT | OMACETAXINE MEPESUCCINATE |
|  | HNF4A | UREA |
|  | YY1 | - |
| Inflammatory abnormality of the skin ([HP:0011123](https://hpo.jax.org/app/browse/term/HP:0011123)) | ZAP70 | - |
|  | CD3E | MUROMMONAB-CD3, CATUMAXOMAB, BLINATUMOMAB |
|  | CTLA4 | IPILIMUMAB, ABATACEPT |
|  | CD3D | MUROMMONAB-CD3, CATUMAXOMAB, BLINATUMOMAB |
|  | CD3G | MUROMMONAB-CD3, CATUMAXOMAB, BLINATUMOMAB |
|  | RAG1 | - |
|  | CD79B | POLATUZUMAB VEDOTIN |
|  | RAG2 | - |
|  | CD79A | - |
|  | IL2RA | BASILIXIMAB, DACLIZUMAB, ALDESLEUKIN, DENILEUKIN DIFTITOX, CETIRIZINE |
| Petechiae ([HP:0000967](https://hpo.jax.org/app/browse/term/HP:0000967)) | ITGA2B | TIROFIBAN, ABCIXIMAB, EPTIFIBATIDE, THROMBIN |
|  | GATA1 | CYTARABINE, DAUNORUBICIN |
|  | GP9 | - |
|  | STAT5B | TOFACITINIB, DASATINIB, EPOETIN ALFA |
|  | PRF1 | EMAPALUMAB |
|  | STXBP2 | EMAPALUMAB |
|  | GP1BB | - |
|  | GFI1B | - |
|  | RAB27A | EMAPALUMAB |
|  | FERMT3 | - |
| Pruritus ([HP:0000989](https://hpo.jax.org/app/browse/term/HP:0000989)) | ABCA12 | - |
|  | PNPLA1 | - |
|  | CYP4F22 | - |
|  | SDR9C7 | - |
|  | LIPN | - |
|  | CERS3 | - |
|  | ALOXE3 | - |
|  | ALOX12B | - |
|  | RAG1 | - |
|  | KIT | NILOTINIB, IMATINIB |
| Scaling skin ([HP:0040189](https://hpo.jax.org/app/browse/term/HP:0040189)) | CDSN | CARBOPLATIN |
|  | KIT | NILOTINIB, IMATINIB |
|  | IL2RA | BASILIXIMAB, DACLIZUMAB, ALDESLEUKIN, DENILEUKIN DIFTITOX, CETIRIZINE |
|  | ZMPSTE24 | - |
|  | ADAM17 | - |
|  | RASA1 | TRAMETINIB |
|  | KRT74 | - |
|  | CHST8 | - |
|  | CARD14 | - |
|  | CASP14 | - |
| Skin rash ([HP:0000988](https://hpo.jax.org/app/browse/term/HP:0000988)) | CTLA4 | IPILIMUMAB, ABATACEPT |
|  | ZAP70 | - |
|  | CD79B | POLATUZUMAB VEDOTIN |
|  | CD79A | - |
|  | BTK | IBRUTINIB |
|  | SYK | FOSTAMATINIB |
|  | FCGR2B | - |
|  | RAG2 | - |
|  | RAG1 | - |
|  | PTPN22 | - |

### Table 3 Reproductive-Genitourinary-Endocrine-Metabolism

| **Symptom (HPO ID)** | **Hub Genes (ranked)** | **FDA-Approved repurposed drugs** |
| --- | --- | --- |
| Abnormal female reproductive system physiology ([HP:0030012](https://hpo.jax.org/app/browse/term/HP:0030012)) | GNRH1 | ZALCITABINE, GOSERELIN, AMINOGLUTETHIMIDE, DITIOCARB, DAPSONE, CAPTOPRIL |
|  | GNRHR | NAFARELIN, GONADORELIN, GANIRELIX ACETATE, DEGARELIX, CETRORELIX, ABARELIX, BUSERELIN, ELAGOLIX, LEUPROLIDE, TRIPTORELIN PAMOATE |
|  | KISS1R | - |
|  | KAL1 | - |
|  | TAC3 | LIOTHYRONINE SODIUM |
|  | FGF8 | - |
|  | FEZF1 | - |
|  | CHD7 | - |
|  | KISS1 | - |
|  | IL17RD | - |
| Decreased glomerular filtration rate ([HP:0012213](https://hpo.jax.org/app/browse/term/HP:0012213)) | BSND | - |
|  | CLCNKB | - |
|  | CLCNKA | - |
|  | CASR | CINACALCET, VELCALCETIDE, CINACALCET HYDROCHLORIDE, PAMIDRONIC ACID |
|  | SLC22A12 | LESINURAD, SULFINPYRAZONE |
|  | SLC2A9 | - |
|  | G6PC | - |
|  | ITGA3 | - |
|  | SLC7A7 | - |
|  | HGD | - |
| Diabetes mellitus ([HP:0000819](https://hpo.jax.org/app/browse/term/HP:0000819)) | CRX | - |
|  | GUCA1B | - |
|  | PRPH2 | - |
|  | RDH12 | - |
|  | RPGR | - |
|  | NR2E3 | - |
|  | TULP1 |  |
|  | SPATA7 | - |
|  | PRPF3 | - |
|  | PDE6G | PENTOXIFYLLINE |
| Abnormality of the digestive system ([HP:0025031](https://hpo.jax.org/app/browse/term/HP:0025031)) | DRC1 | - |
|  | CCDC65 | - |
|  | HYDIN | - |
|  | RSPH4A | - |
|  | RSPH1 | - |
|  | CCDC103 | - |
|  | HEATR2 | - |
|  | DYX1C1 | - |
|  | CCDC40 | - |
|  | CCDC39 | - |
| Edema ([HP:0000969](https://hpo.jax.org/app/browse/term/HP:0000969)) | NR2E3 | - |
|  | RHO | - |
|  | NRL | - |
|  | RLBP1 | - |
|  | PRPF8 | - |
|  | PRPH2 | - |
|  | PDE6G | PENTOXIFYLLINE |
|  | RP9 | - |
|  | KRAS | PANITUMUMAB, CETUXIMAB |
|  | NRAS | BINIMETINIB |
| Female sexual dysfunction ([HP:0030014](https://hpo.jax.org/app/browse/term/HP:0030014)) | KAL1 | - |
|  | FGF8 | - |
|  | GNRH1 | ZALCITABINE, GOSERELIN, AMINOGLUTETHIMIDE, DITIOCARB, DAPSONE, CAPTOPRIL |
|  | TAC3 | LIOTHYRONINE SODIUM |
|  | KISS1R | - |
|  | GNRHR | NAFARELIN, GONADORELIN, GANIRELIX ACETATE, DEGARELIX, CETRORELIX, ABARELIX, BUSERELIN, ELAGOLIX, LEUPROLIDE, TRIPTORELIN PAMOATE |
|  | FEZF1 | - |
|  | CHD7 | - |
|  | IL17RD | - |
|  | KISS1 | - |
| Fever ([HP:0001945](https://hpo.jax.org/app/browse/term/HP:0001945)) | CTLA4 | IPILIMUMAB, ABATACEPT |
|  | CD3E | MUROMMONAB-CD3, CATUMAXOMAB, BLINATUMOMAB |
|  | CD79A | - |
|  | IKZF1 | LENALIDOMIDE |
|  | CD79B | POLATUZUMAB VEDOTIN |
|  | RAG1 | - |
|  | RAG2 | - |
|  | CD3D | MUROMMONAB-CD3, CATUMAXOMAB, BLINATUMOMAB |
|  | SYK | FOSTAMATINIB |
|  | KIT | NILOTINIB, IMATINIB |
| Heat intolerance ([HP:0002046](https://hpo.jax.org/app/browse/term/HP:0002046)) | EDAR | - |
|  | OCA2 | - |
|  | EDA | - |
|  | EDARADD | - |
|  | UBE3A | - |
|  | POFUT1 | - |
|  | ITPR2 | - |
|  | CLDN10 | - |
|  | HNRNPK | - |
|  | LBX1 | - |
| Hypothermia ([HP:0002045](https://hpo.jax.org/app/browse/term/HP:0002045)) | TSHB | MESTRANOL, LEVOTHYROXINE, BACLOFEN |
|  | TG | ACEBUTOLOL, AMINOGLETETHIMIDE |
|  | TPO | CARBIMAZOLE, METHIMAZOLE, PROPYLTHIOURACIL |
|  | TSHR | THYROTROPIN ALFA |
|  | POU1F1 | - |
|  | PROP1 | - |
|  | LHX4 | - |
|  | SLC5A5 | - |
|  | LHX3 | - |
|  | SUCLG1 | - |
| Irregular menstruation ([HP:0000858](https://hpo.jax.org/app/browse/term/HP:0000858)) | PRKAR1A | - |
|  | PRKACA | - |
|  | PHKB | - |
|  | PDE11A | TADALAFIL |
|  | PDE4D | DYPHYLLINE, ROFLUMILAST, APREMILAST |
|  | PHKA2 | - |
|  | AIP | - |
|  | MEN1 | OLSALAZINE |
|  | PHKG2 | - |
|  | GPR101 | - |
| Low-grade fever ([HP:0011134](https://hpo.jax.org/app/browse/term/HP:0011134)) | - | - |
| Male sexual dysfunction ([HP:0040307](https://hpo.jax.org/app/browse/term/HP:0040307)) | KAL1 | - |
|  | FGF8 | - |
|  | GNRH1 | ZALCITABINE, GOSERELIN, AMINOGLUTETHIMIDE, DITIOCARB, DAPSONE, CAPTOPRIL |
|  | TAC3 | LIOTHYRONINE SODIUM |
|  | KISS1R | - |
|  | GNRHR | NAFARELIN, GONADORELIN, GANIRELIX ACETATE, DEGARELIX, CETRORELIX, ABARELIX, BUSERELIN, ELAGOLIX, LEUPROLIDE, TRIPTORELIN PAMOATE |
|  | FEZF1 | - |
|  | CHD7 | - |
|  | IL17RD | - |
|  | KISS1 | - |
| Menorrhagia ([HP:0000132](https://hpo.jax.org/app/browse/term/HP:0000132)) | GP9 | - |
|  | GP1BB | - |
|  | ITGA2B | TIROFIBAN, ABCIXIMAB, EPTIFIBATIDE, THROMBIN |
|  | F8 | SORBITOL, RANITIDINE |
|  | BLOC1S5 | - |
|  | SLFN14 | - |
|  | BLOC1S3 | - |
|  | PRKACG | - |
|  | PRLR | SOMATROPIN |
| Pancreatitis ([HP:0001733](https://hpo.jax.org/app/browse/term/HP:0001733)) | CASR | CINACALCET, VELCALCETIDE, CINACALCET HYDROCHLORIDE, PAMIDRONIC ACID |
|  | CTRC | - |
|  | PRSS1 | - |
|  | PRSS3P2 | - |
|  | CTLA4 | IPILIMUMAB, ABATACEPT |
|  | BCKDHB | - |
|  | PTPN22 | - |
|  | BCKDHA | - |
|  | G6PC | - |
|  | NFS1 | - |
| Recurrent fever ([HP:0001954](https://hpo.jax.org/app/browse/term/HP:0001954)) | NLRC4 | - |
|  | MEFV | - |
|  | NLRP3 | ANAKINRA, TRICLOCARBAN |
|  | NLRP12 | - |
|  | SYK | FOSTAMATINIB |
|  | XIAP | ETOPOSIDE PHOSPHATE |
|  | LPIN2 | - |
|  | PRF1 | EMAPALUMAB |
|  | PRKCD | - |
|  | TMEM173 | - |
| Renal insufficiency ([HP:0000083](https://hpo.jax.org/app/browse/term/HP:0000083)) | RAD51 | - |
|  | FANCM | - |
|  | FANCD2 | DOXORUBICIN HYDROCHLORIDE |
|  | FANCA | CISPLATIN |
|  | FANCB | - |
|  | SLX4 | - |
|  | FANCE | - |
|  | FANCC | MITOMYCIN, CHLORAMBUCIL, MELPHALAN |
|  | FANCF | - |
|  | FANCG | - |
| Temperature instability ([HP:0005968](https://hpo.jax.org/app/browse/term/HP:0005968)) | ZIC2 | - |
|  | SIX3 | - |
|  | SHH | VISMODEGIB, CAFFEINE |
|  | GLI2 | - |
|  | FGF8 | - |
|  | CDON | - |
|  | NODAL | - |
|  | SNRPN | - |
|  | NPAP1 | - |
|  | DDC | CARBIDOPA, BENSERAZIDE |
| Urinary incontinence ([HP:0000020](https://hpo.jax.org/app/browse/term/HP:0000020)) | KIF5A | - |
|  | NIPA1 | - |
|  | KIAA0196 | - |
|  | SPG11 | - |
|  | RTN2 | - |
|  | SPAST | - |
|  | ATL1 | - |
|  | GJC2 | CARBENOXOLONE |
|  | ALDH18A1 | - |
|  | KCNT1 | CLOFILIUM, LOXAPINE, QUINIDINE |

### Table 4 HEENT-ear

| **Symptom (HPO ID)** | **Hub Genes (ranked)** | **FDA-Approved repurposed drugs** |
| --- | --- | --- |
| Ear pain ([HP:0030766](https://hpo.jax.org/app/browse/term/HP:0030766)) | SMO | VISMODEGIB, SONIDEGIB, GLASDEGIB, POSACONAZOLE, SONIDEGIB PHOSPHATE |
|  | PIK3CA | - |
|  | SUFU | SONIDEGIB |
|  | TERT | OMACETAXINE MEPESUCCINATE |
|  | KRT6A | - |
|  | TRAF7 | - |
| Hearing impairment ([HP:0000365](https://hpo.jax.org/app/browse/term/HP:0000365)) | RPGR | - |
|  | IMPDH1 | MYCOPHENOLATE MOFETIL HYDROCHLORIDE, RIBAVIRIN, MYCOPHENOLATE MOFETIL, MYCOPHENOLIC ACID |
|  | RDH12 | - |
|  | SPATA7 | - |
|  | CRX | - |
|  | PRPH2 | - |
|  | TULP1 | - |
|  | CNGA1 | DEQUALINIUM |
|  | NR2E3 | - |
|  | GUCA1B | - |
| Hyperacusis ([HP:0010780](https://hpo.jax.org/app/browse/term/HP:0010780)) | - | - |
| Pulsatile tinnitus ([HP:0008629](https://hpo.jax.org/app/browse/term/HP:0008629)) | MAX | - |
|  | NF1 | TRAMETINIB, BINIMETINIB, COBIMETINIB, DABRAFENIB, VEMURAFENIB |
|  | SDHC | - |
|  | RET | CABOZANTINIB, VANDETANIB |
|  | SDHAF2 | - |
|  | KIF1B | - |
|  | DNMT3A | DECITABINE, IDARUBICIN, WARFARIN |
|  | DLST | - |
| Tinnitus ([HP:0000360](https://hpo.jax.org/app/browse/term/HP:0000360)) | NF1 | TRAMETINIB, BINIMETINIB, COBIMETINIB, DABRAFENIB, VEMURAFENIB |
|  | MAX | - |
|  | RET | CABOZANTINIB, VANDETANIB |
|  | KIF1B | - |
|  | SDHC | - |
|  | SDHAF2 | - |
|  | TERT | OMACETAXINE MEPESUCCINATE |
|  | CACNA1D | ISRADIPINE, CLEVIDIPINE, NIMODIPINE, NITRENDIPINE |
|  | CYP11B2 | IMADIPRIL, BENAZEPRIL, METOCLOPRAMIDE |
|  | KCNJ5 | - |
| Vertigo ([HP:0002321](https://hpo.jax.org/app/browse/term/HP:0002321)) | SCN5A | INDECAINIDE, BENZONATATE, FOSPHENYTOIN, MORICIZINE, HEXYLCAINE, PROCAINAMIDE, MEXILETINE, DISOPYRAMIDE |
|  | SCN2A | - |
|  | SCN1A | ETHOSUXIMIDE |
|  | CACNA1G | ETHOSUXIMIDE, ZONISAMIDE, TRIMETHADIONE, METHSUXIMIDE, PARAMETHADIONE, FLUNARIZINE, PHENSUXIMIDE |
|  | RYR2 | PROCAINE |
|  | SCN1B | ZONISAMIDE |
|  | CASQ2 | - |
|  | NF1 | TRAMETINIB, BINIMETINIB, COBIMETINIB, DABRAFENIB, VEMURAFENIB |
|  | RET | CABOZANTINIB, VANDETANIB |
|  | SDHC | - |

### Table 5 HEENT-ENT

| **Symptom (HPO ID)** | **Hub Genes (ranked)** | **FDA-Approved repurposed drugs** |
| --- | --- | --- |
| Abnormality of the pharynx ([HP:0000600](https://hpo.jax.org/app/browse/term/HP:0000600)) | RSPH4A | - |
|  | HEATR2 | - |
|  | RSPH1 | - |
|  | SPAG1 | - |
|  | DYX1C1 | - |
|  | C21orf59 | - |
|  | CCDC65 | - |
|  | HYDIN | - |
|  | DNAI2 | - |
|  | DNAI1 | - |
| Dysphonia ([HP:0001618](https://hpo.jax.org/app/browse/term/HP:0001618)) | MAX | - |
|  | SDHC | - |
|  | NF1 | TRAMETINIB, BINIMETINIB, COBIMETINIB, DABRAFENIB, VEMURAFENIB |
|  | KIF1B | - |
|  | RET | CABOZANTINIB, VANDETANIB |
|  | SDHAF2 | - |
|  | SYT2 | - |
|  | CHAT | RIVASTIGMINE, GALANTAMINE, DONEPEZILM PILOCARPINE |
|  | SLC18A3 | - |
|  | MFN2 | - |
| Nasal congestion ([HP:0001742](https://hpo.jax.org/app/browse/term/HP:0001742)) | CCDC65 | - |
|  | DRC1 | - |
|  | HYDIN | - |
|  | RSPH4A | - |
|  | RSPH1 | - |
|  | CCDC103 | - |
|  | HEATR2 | - |
|  | DYX1C1 | - |
|  | DNAI2 | - |
|  | CCDC40 | - |
| Rhinitis ([HP:0012384](https://hpo.jax.org/app/browse/term/HP:0012384)) | CCDC65 | - |
|  | DRC1 | - |
|  | HYDIN | - |
|  | RSPH4A | - |
|  | RSPH1 | - |
|  | CCDC103 | - |
|  | HEATR2 | - |
|  | DYX1C1 | - |
|  | DNAI2 | - |
|  | CCDC40 | - |

### Table 6 HEENT-eye

| **Symptom (HPO ID)** | **Hub Genes (ranked)** | **FDA-Approved repurposed drugs** |
| --- | --- | --- |
| Blindness ([HP:0000618](https://hpo.jax.org/app/browse/term/HP:0000618)) | RPGR | - |
|  | CRX | - |
|  | CNGA1 | DEQUALINIUM |
|  | GUCA1B | - |
|  | NR2E3 | - |
|  | PRPH2 | - |
|  | RDH12 | - |
|  | TULP1 | - |
|  | SPATA7 | - |
|  | IMPDH1 | MYCOPHENOLATE MOFETIL HYDROCHLORIDE, RIBAVIRIN, MYCOPHENOLATE MOFETIL, MYCOPHENOLIC ACID |
| Blurred vision ([HP:0000622](https://hpo.jax.org/app/browse/term/HP:0000622)) | KCNA1 | BUPIVACAINE, ISOFLURANE, DESFLURANE |
|  | SCN9A | - |
|  | SCN4A | LEVOBUPIVACAINE HYDROCHLORIDE, BUPIVACAINE HYDROCHLORIDE |
|  | ATP1A2 | ACETYLDIGITOXIN, DESLANOSIDE |
|  | SPATA7 | - |
|  | COL8A2 | - |
|  | VSX1 | - |
|  | CLCNKB | - |
|  | GRHL2 | - |
|  | RPE65 | - |
| Conjunctivitis ([HP:0000509](https://hpo.jax.org/app/browse/term/HP:0000509)) | CD19 | BLINATUMOMAB |
|  | CD79B | POLATUZUMAB VEDOTIN |
|  | CD79A | - |
|  | CR2 | PROGESTERONE |
|  | BTK | IBRUTINIB |
|  | TNFRSF13C | - |
|  | TNFRSF13B | - |
|  | RAG1 | - |
|  | RAG2 | - |
|  | IKZF1 | LENALIDOMIDE |
| Diplopia ([HP:0000651](https://hpo.jax.org/app/browse/term/HP:0000651)) | DOK7 | - |
|  | MUSK | - |
|  | RAPSN | - |
|  | SCN4A | LEVOBUPIVACAINE HYDROCHLORIDE, BUPIVACAINE HYDROCHLORIDE |
|  | CHRNE | SUCCINYLCHOLINE CHLORIDE, PIPECURONIUM BROMIDE, MIVACURIUM CHLORIDE, METOCURINE IODIDE, DOXACURIUM CHLORIDE, VECURONIUM BROMIDE, DECAMETHONIUM BROMIDE |
|  | CHRNA1 | - |
|  | CHRNB1 | SUCCINYLCHOLINE CHLORIDE, PIPECURONIUM BROMIDE, MIVACURIUM CHLORIDE, METOCURINE IODIDE, DOXACURIUM CHLORIDE, VECURONIUM BROMIDE, DECAMETHONIUM BROMIDE |
|  | CHRND | SUCCINYLCHOLINE CHLORIDE, PIPECURONIUM BROMIDE, MIVACURIUM CHLORIDE, METOCURINE IODIDE, DOXACURIUM CHLORIDE, VECURONIUM BROMIDE, DECAMETHONIUM BROMIDE |
|  | CHAT | RIVASTIGMINE, GALANTAMINE, DONEPEZILM PILOCARPINE |
|  | KCNA1 | BUPIVACAINE, ISOFLURANE, DESFLURANE |
| Gaze-evoked nystagmus ([HP:0000640](https://hpo.jax.org/app/browse/term/HP:0000640)) | ATXN10 | - |
|  | PRKCG | - |
|  | ATXN1 | - |
|  | ATXN2 | - |
|  | ATXN3 | - |
|  | CACNA1G | ETHOSUXIMIDE, ZONISAMIDE, TRIMETHADIONE, METHSUXIMIDE, PARAMETHADIONE, FLUNARIZINE, PHENSUXIMIDE |
|  | SCN8A | - |
|  | SCN1A | ETHOSUXIMIDE |
|  | KCND3 | - |
|  | ATP1A2 | ACETYLDIGITOXIN, DESLANOSIDE |
| Keratoconjunctivitis sicca ([HP:0001097](https://hpo.jax.org/app/browse/term/HP:0001097)) | ERCC6 | DEXTROMETHORPHAN, CYCLOSPORINE |
|  | GTF2H5 | - |
|  | ERCC2 | OXALIPLATIN, LEUCOVORIN |
|  | HLA-DRB1 | CLAVULANIC ACID, PITAVASTATIN |
|  | BTNL2 | - |
|  | COL17A1 | - |
|  | MEFV | - |
|  | RNF125 | - |
|  | GJB6 | CARBENOXOLONE |
|  | SCN9A | - |
| Ocular pain ([HP:0200026](https://hpo.jax.org/app/browse/term/HP:0200026)) | COL8A2 | - |
|  | GNA11 | BINIMETINIB, CABOZANTINIB |
|  | GRHL2 | - |
|  | VSX1 | - |
|  | GNAQ | VERTEPORFIN |
|  | OVOL2 | - |
|  | CHST6 | - |
|  | AGBL1 | - |
|  | SCN9A | - |
|  | COL17A1 | - |
| Peripheral visual field loss ([HP:0007994](https://hpo.jax.org/app/browse/term/HP:0007994)) | ABCA4 | - |
|  | RHO | - |
|  | CRX | - |
|  | CNGA1 | DEQUALINIUM |
|  | PRPH2 | - |
|  | RLBP1 | - |
|  | RPE65 | - |
|  | SPATA7 | - |
|  | GUCY2D | RIOCIGUAT |
|  | PDE6A | PENTOXIFYLLINE |
| Photophobia ([HP:0000613](https://hpo.jax.org/app/browse/term/HP:0000613)) | GUCA1B | - |
|  | NR2E3 | - |
|  | PRPH2 | - |
|  | CRX | - |
|  | CNGA1 | DEQUALINIUM |
|  | NRL | - |
|  | PDE6G | PENTOXIFYLLINE |
|  | RPGR | - |
|  | RDH12 | - |
|  | TULP1 | - |
| Red eye ([HP:0025337](https://hpo.jax.org/app/browse/term/HP:0025337)) | CD19 | BLINATUMOMAB |
|  | CD79A | - |
|  | CD79B | POLATUZUMAB VEDOTIN |
|  | CR2 | PROGESTERONE |
|  | BTK | IBRUTINIB |
|  | TNFRSF13C | - |
|  | TNFRSF13B | - |
|  | RAG1 | - |
|  | RAG2 | - |
|  | IKZF1 | LENALIDOMIDE |
| Visual loss ([HP:0000572](https://hpo.jax.org/app/browse/term/HP:0000572)) | USH2A | - |
|  | CLRN1 | - |
|  | DFNB31 | - |
|  | CDH23 | METHYLPHENIDATE |
|  | PCDH15 | - |
|  | CIB2 | - |
|  | USH1C | - |
|  | PDZD7 | - |
|  | USH1G | - |
|  | GPR98 | - |
| Vitreous floaters ([HP:0100832](https://hpo.jax.org/app/browse/term/HP:0100832)) | NDP | - |
|  | TSPAN12 | - |
|  | ZNF408 | - |
|  | CTNNB1 | FLUORESCEN SODIUM |

### Table 7 Lab

| **Symptom (HPO ID)** | **Hub Genes (ranked)** | **FDA-Approved repurposed drugs** |
| --- | --- | --- |
| Abnormal calcium-phosphate regulating hormone level ([HP:0100530](https://hpo.jax.org/app/browse/term/HP:0100530)) | PTH | IBANDRONIC ACID, CLOMIPHENE, CALCITONIN, TRILOSTANE, TERIPARATIDE, PENICILLAMINE, HYDROXYUREA, RANITIDINE |
|  | VDR | CALCIUM, DIHYDROTACHYSTEROL |
|  | KL | - |
|  | SLC34A3 | - |
|  | CASR | CINACALCET, VELCALCETIDE, CINACALCET HYDROCHLORIDE, PAMIDRONIC ACID |
|  | SLC34A1 | - |
|  | RET | CABOZANTINIB, VANDETANIB |
|  | NF1 | TRAMETINIB, BINIMETINIB, COBIMETINIB, DABRAFENIB, VEMURAFENIB |
|  | TRPV6 | - |
|  | SDHC | - |
| Abnormal circulating protein concentration ([HP:0010876](https://hpo.jax.org/app/browse/term/HP:0010876)) | TCAP | - |
|  | LDB3 | - |
|  | CSRP3 | - |
|  | MYH6 | - |
|  | MYL2 | - |
|  | DMD | ETEPLIRSEN, ATALUREN |
|  | ACTN2 | - |
|  | MYBPC3 | - |
|  | TNNI3 | - |
|  | MYH7 | - |
| Abnormality of fibrinolysis ([HP:0040224](https://hpo.jax.org/app/browse/term/HP:0040224)) | - | - |
| Decreased circulating calcifediol concentration ([HP:0012053](https://hpo.jax.org/app/browse/term/HP:0012053)) | CYP3A4 | - |
|  | CYP2C9 | - |
|  | CYP2C19 | - |
|  | ABCB1 | - |
|  | CYP3A7 | MICONAZOLE, QUININE, ETONOGESTREL, COBICISTAT |
|  | UGT2B7 | EFAVIRENZ, FLURBIPROFEN, APREPITANT |
|  | UGT1A8 | FEBUXOSTAT |
|  | POR | TACROLIMUS |
|  | CYB5A | - |
|  | CES1 | CLOPIDOGREL, TRANDOLAPRIL, DIACETYLMORPHINE |
| Elevated erythrocyte sedimentation rate ([HP:0003565](https://hpo.jax.org/app/browse/term/HP:0003565)) | PTPN22 | - |
|  | CTLA4 | IPILIMUMAB, ABATACEPT |
|  | HLA-DRB1 | CLAVULANIC ACID, PITAVASTATIN |
|  | CIITA | - |
|  | MEFV | - |
|  | IL2RA | BASILIXIMAB, DACLIZUMAB, ALDESLEUKIN, DENILEUKIN DIFTITOX, CETIRIZINE |
|  | LPIN2 | - |
|  | NLRP3 | ANAKINRA, TRICLOCARBAN |
|  | TMEM173 | - |
|  | NLRP12 | - |
| Elevated circulating alkaline phosphatase concentration ([HP:0003155](https://hpo.jax.org/app/browse/term/HP:0003155)) | PIGV | - |
|  | PIGA | - |
|  | PIGL | - |
|  | PIGB | - |
|  | PIGQ | - |
|  | PIGY | - |
|  | PGAP2 | - |
|  | PIGW | - |
|  | FGF23 | BUROSUMAB |
|  | PHEX | CALCITRIOL |
| Elevated circulating alanine aminotransferase concentration ([HP:0031964](https://hpo.jax.org/app/browse/term/HP:0031964)) | NR1H4 | OBETICHOLIC ACID, CHENODIOL |
|  | BAAT | - |
|  | SLC51B | - |
|  | SLC2A2 | METFORMIN |
|  | SLC25A13 | - |
|  | SCO1 | - |
|  | DAK | - |
|  | TTC26 | - |
|  | KIF12 | - |
|  | PKHD1 | - |
| Elevated circulating aspartate aminotransferase concentration ([HP:0031956](https://hpo.jax.org/app/browse/term/HP:0031956)) | NR1H4 | OBETICHOLIC ACID, CHENODIOL |
|  | BAAT | - |
|  | SLC51B | - |
|  | SLC2A2 | METFORMIN |
|  | COG8 | - |
|  | TTC26 | - |
|  | KIF12 | - |
|  | NFS1 | - |
|  | PGM1 | - |
|  | MARS | - |
| Elevated circulating creatine kinase concentration ([HP:0003236](https://hpo.jax.org/app/browse/term/HP:0003236)) | TCAP | - |
|  | LDB3 | - |
|  | CSRP3 | - |
|  | MYH6 | - |
|  | DMD | ETEPLIRSEN, ATALUREN |
|  | MYL2 | - |
|  | ACTN2 | - |
|  | MYBPC3 | - |
|  | TNNI3 | - |
|  | MYH7 | - |
| Elevated circulating creatinine concentration ([HP:0003259](https://hpo.jax.org/app/browse/term/HP:0003259)) | WDR19 | - |
|  | CC2D2A | - |
|  | TTC26 | - |
|  | INVS | DULOXETINE |
|  | ADAMTS13 | - |
|  | CFHR3 | - |
|  | TREX1 | - |
|  | FAN1 | - |
|  | IKBKAP | - |
|  | HNF1B | METFORMIN |
| Elevated circulating C-reactive protein concentration ([HP:0011227](https://hpo.jax.org/app/browse/term/HP:0011227)) | NLRC4 | - |
|  | MEFV | - |
|  | NLRP3 | ANAKINRA, TRICLOCARBAN |
|  | PSTPIP1 | - |
|  | PTPN22 | - |
|  | CTLA4 | IPILIMUMAB, ABATACEPT |
|  | CIITA | - |
|  | SYK | FOSTAMATINIB |
|  | PRSS1 | - |
|  | HLA-DRB1 | CLAVULANIC ACID, PITAVASTATIN |
| Increased circulating ferritin concentration ([HP:0003281](https://hpo.jax.org/app/browse/term/HP:0003281)) | XIAP | ETOPOSIDE PHOSPHATE |
|  | SLC25A38 | - |
|  | STXBP2 | EMAPALUMAB |
|  | PRF1 | EMAPALUMAB |
|  | GLRX5 | - |
|  | CPOX | - |
|  | NLRC4 | - |
|  | SLC19A1 | RALTITREXED, PEMETREXED, LEUCOVORIN |
|  | TTC26 | - |
|  | SLC7A7 | - |
| Elevated gamma-glutamyltransferase level ([HP:0030948](https://hpo.jax.org/app/browse/term/HP:0030948)) | - | - |
| Increased circulating interleukin 6 ([HP:0030783](https://hpo.jax.org/app/browse/term/HP:0030783)) | - | - |
| Increased circulating lactate dehydrogenase concentration ([HP:0025435](https://hpo.jax.org/app/browse/term/HP:0025435)) | ACAD9 | - |
|  | SLC7A7 | - |
|  | CPT2 | - |
|  | SLC25A13 | - |
|  | RHAG | - |
|  | SLC19A1 | RALTITREXED, PEMETREXED, LEUCOVORIN |
|  | RB1 | FOSTAMATINIB, RIBOCICLIB, LORLATINIB |
|  | HLA-DRB1 | CLAVULANIC ACID, PITAVASTATIN |
|  | RHCE | - |
|  | PIGA | - |
| Increased circulating NT-proBNP concentration ([HP:0031185](https://hpo.jax.org/app/browse/term/HP:0031185)) | - | - |
| Elevated circulating thyroid-stimulating hormone concentration ([HP:0002925](https://hpo.jax.org/app/browse/term/HP:0002925)) | SLC5A5 | - |
|  | TSHR | THYROTROPIN ALFA |
|  | TG | ACEBUTOLOL, AMINOGLETETHIMIDE |
|  | NKX2-1 | - |
|  | TPO | CARBIMAZOLE, METHIMAZOLE, PROPYLTHIOURACIL |
|  | SLC16A2 | - |
|  | SECISBP2 | - |
|  | SLC35A2 | - |
|  | PRKAR1A | - |
|  | CDH23 | METHYLPHENIDATE |
| Hyperglycemia ([HP:0003074](https://hpo.jax.org/app/browse/term/HP:0003074)) | ABCC8 | REPAGLINIDE, GLICLAZIDE, GLIQUIDONE, GLIPIZIDE, NATEGLINIDE, GLIMEPIRIDE, ACETOHEXAMIDE, GLYBURIDE, STERILE TOLBUTAMIDE SODIUM, TOLAZAMIDE |
|  | INS | INULIN, INSULIN GLARGINE |
|  | KCNJ11 | GLIMEPIRIDE, GLIQUIDONE, GLIPIZIDE, GLICLAZIDE, THIAMYLAL, GLYBURIDE, STERILE TOLBUTAMIDE SODIUM, DIAZOXIDE |
|  | GCK | - |
|  | PAX4 | REPAGLINIDE |
|  | PDX1 | - |
|  | HNF4A | UREA |
|  | HNF1A | - |
|  | SLC2A2 | METFORMIN |
|  | NEUROD1 | REPAGLINIDE, DEFEROXAMINE |
| Hypocalcemia ([HP:0002901](https://hpo.jax.org/app/browse/term/HP:0002901)) | PTH | IBANDRONIC ACID, CLOMIPHENE, CALCITONIN, TRILOSTANE, TERIPARATIDE, PENICILLAMINE, HYDROXYUREA, RANITIDINE |
|  | CASR | CINACALCET, VELCALCETIDE, CINACALCET HYDROCHLORIDE, PAMIDRONIC ACID |
|  | VDR | CALCIPOTRIOL, CALCIUM, DIHYDROTACHYSTEROL |
|  | FGF23 | BUROSUMAB |
|  | TNFSF11 | DENOSUMAB, LENALIDOMIDE, ANASTROZOLE, LETROZOLE |
|  | CYP2R1 | CALCIFEDIOL, RIBAVIRIN |
|  | TBX1 | - |
|  | DGCR2 | - |
|  | TNFRSF11A | DICLOFENAC, ACETAMINOPHEN |
|  | GCM2 | - |
| Hypofibrinogenemia ([HP:0011900](https://hpo.jax.org/app/browse/term/HP:0011900)) | XIAP | ETOPOSIDE PHOSPHATE |
|  | PRF1 | EMAPALUMAB |
|  | EPB42 | - |
|  | ANK1 | - |
|  | STXBP2 | EMAPALUMAB |
|  | SPTA1 | - |
|  | STAT5B | TOFACITINIB, DASATINIB, EPOETIN ALFA |
|  | AHCY | - |
|  | NLRC4 | - |
|  | PRKAR1A | - |
| Hypoglycemia ([HP:0001943](https://hpo.jax.org/app/browse/term/HP:0001943)) | INS | INULIN, INSULIN GLARGINE |
|  | GCK | - |
|  | ABCC8 | REPAGLINIDE, GLICLAZIDE, GLIQUIDONE, GLIPIZIDE, NATEGLINIDE, GLIMEPIRIDE, ACETOHEXAMIDE, GLYBURIDE, STERILE TOLBUTAMIDE SODIUM, TOLAZAMIDE |
|  | KCNJ11 | GLIMEPIRIDE, GLIQUIDONE, GLIPIZIDE, GLICLAZIDE, THIAMYLAL, GLYBURIDE, STERILE TOLBUTAMIDE SODIUM, DIAZOXIDE |
|  | PAX4 | REPAGLINIDE |
|  | HNF4A | UREA |
|  | HNF1A | - |
|  | PDX1 | - |
|  | SLC2A2 | METFORMIN |
|  | NEUROD1 | REPAGLINIDE, DEFEROXAMINE |
| Hypophosphatemia ([HP:0002148](https://hpo.jax.org/app/browse/term/HP:0002148)) | FGF23 | BUROSUMAB |
|  | SLC34A3 | - |
|  | PHEX | CALCITRIOL |
|  | SLC34A1 | - |
|  | VDR | CALCIPOTRIOL, CALCIUM, DIHYDROTACHYSTEROL |
|  | DMP1 | - |
|  | CASR | CINACALCET, VELCALCETIDE, CINACALCET HYDROCHLORIDE, PAMIDRONIC ACID |
|  | ENPP1 | - |
|  | TNFSF11 | DENOSUMAB, LENALIDOMIDE, ANASTROZOLE, LETROZOLE |
|  | CLCN5 | - |
| Thrombocytopenia ([HP:0001873](https://hpo.jax.org/app/browse/term/HP:0001873)) | BRCA1 | - |
|  | RAD51 | - |
|  | FANCM | - |
|  | PALB2 | MITOMYCIN, OLAPRIB |
|  | SLX4 | - |
|  | FANCA | CISPLATIN |
|  | FANCD2 | DOXORUBICIN HYDROCHLORIDE |
|  | FANCC | MITOMYCIN, CHLORAMBUCIL, MELPHALAN |
|  | FANCE | - |
|  | FANCL | - |

### Table 8 General-pain

| **Symptom (HPO ID)** | **Hub Genes (ranked)** | **FDA-Approved repurposed drugs** |
| --- | --- | --- |
| Arthralgia ([HP:0002829](https://hpo.jax.org/app/browse/term/HP:0002829)) | CTLA4 | IPILIMUMAB, ABATACEPT |
|  | CD19 | BLINATUMOMAB |
|  | CR2 | PROGESTERONE |
|  | TNFRSF13B | - |
|  | TNFRSF13C | - |
|  | FAS | TENIPOSIDE |
|  | MS4A1 | TOSITUMOMAB, OFATUMUMAB, OBINUTUZUMAB, OCRELIZUMAB, IBRITUMOMAB TIUXETAN, YTTRIUM Y 90 IBRITUMOMAB TIUXETAN, RITUXIMAB |
|  | IL2RA | BASILIXIMAB, DACLIZUMAB, ALDESLEUKIN, DENILEUKIN DIFTITOX, CETIRIZINE |
|  | COL2A1 | COLLAGENASE CLOSTRIDIUM HISTOLYTICUM, OCRIPLASMIN |
|  | COL10A1 | - |
| Back pain ([HP:0003418](https://hpo.jax.org/app/browse/term/HP:0003418)) | PIK3CA | - |
|  | KRAS | PANITUMUMAB, CETUXIMAB |
|  | SMAD4 | LYSINE, ALECTINIB |
|  | BRCA1 | - |
|  | PALB2 | OLAPARIB, MITOMYCIN |
|  | TERT | OMACETAXINE MEPESUCCINATE |
|  | TBX6 | - |
|  | ALDH18A1 | - |
|  | RASA1 | TRAMETINIB |
|  | MESP2 | - |
| Bone pain ([HP:0002653](https://hpo.jax.org/app/browse/term/HP:0002653)) | PHEX | CALCITRIOL |
|  | FGF23 | BUROSUMAB |
|  | SLC34A3 | - |
|  | DMP1 | - |
|  | SLC34A1 | - |
|  | VDR | CALCIPOTRIOL, CALCIUM, DIHYDROTACHYSTEROL |
|  | ENPP1 | - |
|  | TNFSF11 | DENOSUMAB, LENALIDOMIDE, ANASTROZOLE, LETROZOLE |
|  | CLCN5 | - |
|  | STAT5B | TOFACITINIB, DASATINIB, EPOETIN ALFA |
| Chest pain ([HP:0100749](https://hpo.jax.org/app/browse/term/HP:0100749)) | NF1 | TRAMETINIB, BINIMETINIB, COBIMETINIB, DABRAFENIB, VEMURAFENIB |
|  | RET | CABOZANTINIB, VANDETANIB |
|  | SDHC | - |
|  | MAX | - |
|  | SDHAF2 | - |
|  | KIF1B | - |
|  | SMAD4 | LYSINE, ALECTINIB |
|  | CTNNB1 | FLUORESCEN SODIUM |
|  | ACTA2 | - |
|  | COL3A1 | COLLAGENASE CLOSTRIDIUM HISTOLYTICUM, OCRIPLASMIN |
| Limb pain ([HP:0009763](https://hpo.jax.org/app/browse/term/HP:0009763)) | COL9A1 | - |
|  | COL9A3 | - |
|  | COL2A1 | COLLAGENASE CLOSTRIDIUM HISTOLYTICUM, OCRIPLASMIN |
|  | COL5A2 | COLLAGENASE CLOSTRIDIUM HISTOLYTICUM, OCRIPLASMIN |
|  | MATN3 | - |
|  | SPTLC1 | - |
|  | BSCL2 | - |
|  | MFN2 | - |
|  | CPOX | - |
|  | ATL1 | - |
| Myalgia ([HP:0003326](https://hpo.jax.org/app/browse/term/HP:0003326)) | CAV3 | - |
|  | MYOT | - |
|  | FKTN | - |
|  | FKRP | - |
|  | CAPN3 | - |
|  | DMD | ETEPLIRSEN, ATALUREN |
|  | TRIM32 |  |
|  | PYGM | - |
|  | PHKA1 | - |
|  | PHKB | - |
| Pain ([HP:0012531](https://hpo.jax.org/app/browse/term/HP:0012531)) | KRAS | PANITUMUMAB, CETUXIMAB |
|  | CTNNB1 | FLUORESCEN SODIUM |
|  | ESR1 | CLOMIPHENE, LEVONORGESTREL |
|  | PIK3CA | - |
|  | SMAD4 | LYSINE, ALECTINIB |
|  | NF1 | TRAMETINIB, BINIMETINIB, COBIMETINIB, DABRAFENIB, VEMURAFENIB |
|  | KIT | NILOTINIB, IMATINIB |
|  | RET | CABOZANTINIB, VANDETANIB |
|  | ERBB4 | DACOMITINIB |
|  | GNAQ | VERTEPORFIN |
| Pedal edema ([HP:0010741](https://hpo.jax.org/app/browse/term/HP:0010741)) | NKX2-5 | - |
|  | TBX20 | - |
|  | MYH6 | - |
|  | GATA4 | WARFARIN |
|  | GATA6 | - |
|  | SNRPN | - |
|  | SETD2 | - |
|  | MAGEL2 | - |
|  | OCA2 | - |
|  | RBM8A | - |

### Table 9 General-symptom

| **Symptom (HPO ID)** | **Hub Genes (ranked)** | **FDA-Approved repurposed drugs** |
| --- | --- | --- |
| Arthritis ([HP:0001369](https://hpo.jax.org/app/browse/term/HP:0001369)) | COL2A1 | COLLAGENASE CLOSTRIDIUM HISTOLYTICUM, OCRIPLASMIN |
|  | CTLA4 | IPILIMUMAB, ABATACEPT |
|  | SYK | FOSTAMATINIB |
|  | BTK | IBRUTINIB |
|  | CD79B | POLATUZUMAB VEDOTIN |
|  | ACAN | - |
|  | FCGR2B | - |
|  | COL9A3 | - |
|  | MATN3 | - |
|  | ASPN | - |
| Asthenia ([HP:0025406](https://hpo.jax.org/app/browse/term/HP:0025406)) | - | - |
| Chills ([HP:0025143](https://hpo.jax.org/app/browse/term/HP:0025143)) | EPB42 | - |
|  | ANK1 | - |
|  | SPTA1 | - |
|  | EPB41 | BUPROPION, NICOTINE |
|  | NLRP3 | ANAKINRA, TRICLOCARBAN |
|  | PKHD1 | - |
| Constitutional symptom ([HP:0025142](https://hpo.jax.org/app/browse/term/HP:0025142)) | MYH6 | - |
|  | MYL2 | - |
|  | MYBPC3 | - |
|  | TNNI3 | - |
|  | MYH7 | - |
|  | SCN5A | INDECAINIDE, BENZONATATE, FOSPHENYTOIN, MORICIZINE, HEXYLCAINE, PROCAINAMIDE, MEXILETINE, DISOPYRAMIDE |
|  | RYR2 | PROCAINE |
|  | MYL3 | - |
|  | LDB3 | - |
|  | CASQ2 | - |
| Difficulty walking ([HP:0002355](https://hpo.jax.org/app/browse/term/HP:0002355)) | REEP1 | - |
|  | RTN2 | - |
|  | SPG21 | - |
|  | AP4E1 | - |
|  | AP4B1 | - |
|  | SPG11 | - |
|  | AP4S1 | - |
|  | KIAA0196 | - |
|  | BSCL2 | - |
|  | GJC2 | CARBENOXOLONE |
| Exercise intolerance ([HP:0003546](https://hpo.jax.org/app/browse/term/HP:0003546)) | PYGM | - |
|  | PFKM | - |
|  | PGAM2 | - |
|  | PHKA1 | - |
|  | MYH6 | - |
|  | ENO3 | - |
|  | PHKA2 | - |
|  | ACTA1 | - |
|  | PHKB | - |
|  | PHKG2 | - |
| Fatigue ([HP:0012378)](https://hpo.jax.org/app/browse/term/HP:0012378) | CTNNB1 | FLUORESCEN SODIUM |
|  | NF1 | TRAMETINIB, BINIMETINIB, COBIMETINIB, DABRAFENIB, VEMURAFENIB |
|  | SMAD4 | LYSINE, ALECTINIB |
|  | KRAS | PANITUMUMAB, CETUXIMAB |
|  | RET | CABOZANTINIB, VANDETANIB |
|  | PIK3CA | - |
|  | BRCA1 | - |
|  | KIT | NILOTINIB, IMATINIB |
|  | ERBB4 | DACOMITINIB |
|  | PMS2 | NIVOLUMAB |
| Impaired ability to dress oneself ([HP:0031060](https://hpo.jax.org/app/browse/term/HP:0031060)) | - | - |
| Impairment of activities of daily living ([HP:0031058](https://hpo.jax.org/app/browse/term/HP:0031058)) | KIF5A | - |
|  | KIAA0196 | - |
|  | RTN2 | - |
|  | NIPA1 | - |
|  | SPG11 | - |
|  | SPAST | - |
|  | ATL1 | - |
|  | GJC2 | CARBENOXOLONE |
|  | PLP1 | - |
|  | GABRA1 | METHOHEXITAL, BARBITAL, THIAMYLAL, ZALEPLON, ZOPICLONE, ZOLPIDEM |
| Night sweats ([HP:0030166](https://hpo.jax.org/app/browse/term/HP:0030166)) | - | - |
| Postexertional malaise ([HP:0030973](https://hpo.jax.org/app/browse/term/HP:0030973)) | SLC2A9 | - |
|  | COL9A1 | - |
|  | SCN5A | INDECAINIDE, BENZONATATE, FOSPHENYTOIN, MORICIZINE, HEXYLCAINE, PROCAINAMIDE, MEXILETINE, DISOPYRAMIDE |
|  | NPPA | CLORTHALIDONE, AMLODIPINE |
|  | SLC22A12 | LESINURAD, SULFINPYRAZONE |
|  | COL9A3 | - |
|  | PYGM | - |
|  | SVIL | - |
| Shivering ([HP:0025144](https://hpo.jax.org/app/browse/term/HP:0025144)) | CHRNA2 | PIPECURONIUM, METOCURINE, CISATRACURIUM, PANCURONIUM, MECAMYLAMINE, DOXACURIUM, MIVACURIUM, DECAMETHONIUM, VECURONIUM, TUBOCURARINE |
|  | CHRNB2 | NICOTINE, VARENICLINE, VARENICLINE TARTRATE, NICOTINE POLACRILEX |
|  | CHRNA4 | VARENICLINE |
|  | CHRNB4 | PENTOLINIUM TARTRATE, TRIMETHAPHAN CAMSYLATE, MECAMYLAMINE HYDROCHLORIDE |
|  | CHRNA7 | GALANTAMINE |
|  | CHRNA9 | NICOTINE |
|  | CHRNA3 | VARENICLINE, PENTOLINIUM TARTRATE, TREMETHAPHAN CAMYSLATE, MECAMYLAMINE HYDROCHLORIDE |
|  | CHRNB3 | - |
|  | CHRNB1 | SUCCINYLCHOLINE CHLORIDE, PIPECURONIUM BROMIDE, MIVACURIUM CHLORIDE, METOCURINE IODIDE, DOXACURIUM CHLORIDE, VECURONIUM BROMIDE, DECAMETHONIUM BROMIDE |
|  | KCNT1 | CLOFILIUM, LOXAPINE, QUINIDINE |
| Stiff neck ([HP:0025258](https://hpo.jax.org/app/browse/term/HP:0025258)) | - | - |
| Weight loss ([HP:0001824](https://hpo.jax.org/app/browse/term/HP:0001824)) | FANCD2 | DOXORUBICIN HYDROCHLORIDE |
|  | RAD51 | - |
|  | FANCM | - |
|  | FANCA | CISPLATIN |
|  | FANCC | MITOMYCIN, CHLORAMBUCIL, MELPHALAN |
|  | FANCL | - |
|  | FANCG | - |
|  | UBE2T | - |
|  | FANCB | - |
|  | FANCF | - |
| Xerostomia ([HP:0000217](https://hpo.jax.org/app/browse/term/HP:0000217)) | C9orf72 | - |
|  | FUS | TRETINOIN |
|  | ATXN2 | - |
|  | MATR3 | - |
|  | NEK1 | - |
|  | NEFH | - |
|  | FIG4 | - |
|  | TAF15 | - |
|  | CCNF | - |
|  | GLE1 | - |

### Table 10 Cardiovascular-finding

| **Symptom (HPO ID)** | **Hub Genes (ranked)** | **FDA-Approved repurposed drugs** |
| --- | --- | --- |
| Abnormal heart morphology ([HP:0001627](https://hpo.jax.org/app/browse/term/HP:0001627)) | DRC1 | - |
|  | CCDC65 | - |
|  | HYDIN | - |
|  | RSPH4A | - |
|  | RSPH1 | - |
|  | CCDC103 | - |
|  | HEATR2 | - |
|  | DYX1C1 | - |
|  | DNAI2 | - |
|  | CCDC40 | - |
| Abnormal heart rate variability ([HP:0031860](https://hpo.jax.org/app/browse/term/HP:0031860)) | ZIC2 | - |
|  | SIX3 | - |
|  | SHH | VISMODEGIB, CAFFEINE |
|  | GLI2 | - |
|  | CDON | - |
|  | FGF8 | - |
|  | NODAL | - |
|  | TGIF1 | - |
|  | PHOX2B | - |
|  | SMC1A | - |
| Abnormal left ventricular function ([HP:0005162](https://hpo.jax.org/app/browse/term/HP:0005162)) | MYH6 | - |
|  | TNNI3 | - |
|  | NKX2-5 | - |
|  | GATA4 | WARFARIN |
|  | SCN5A | INDECAINIDE, BENZONATATE, FOSPHENYTOIN, MORICIZINE, HEXYLCAINE, PROCAINAMIDE, MEXILETINE, DISOPYRAMIDE |
|  | NPPA | CLORTHALIDONE, AMLODIPINE |
|  | MYH7 | - |
|  | TBX20 | - |
|  | ACTA2 | - |
|  | LDB3 | - |
| Abnormal pericardium morphology ([HP:0001697](https://hpo.jax.org/app/browse/term/HP:0001697)) | CTLA4 | IPILIMUMAB, ABATACEPT |
|  | PTPN22 | - |
|  | HLA-DRB1 | CLAVULANIC ACID, PITAVASTATIN |
|  | IL23R | - |
|  | MYBPC3 | - |
|  | CCBE1 | - |
|  | FAS | TENIPOSIDE |
|  | FCGR2B | - |
|  | ABCC9 | NICORANDIL, PINACIDIL, TOLBUTAMIDE, GLIMEPIRIDE |
|  | PTPN14 | - |
| Bradycardia ([HP:0001662](https://hpo.jax.org/app/browse/term/HP:0001662)) | SCN5A | INDECAINIDE, BENZONATATE, FOSPHENYTOIN, MORICIZINE, HEXYLCAINE, PROCAINAMIDE, MEXILETINE, DISOPYRAMIDE |
|  | ANK2 | - |
|  | CAV3 | - |
|  | CASQ2 | - |
|  | KCNE2 | - |
|  | HCN4 | IVABRADINE HYDROCHLORIDE, DRONEDARONE HYDROCHLORIDE, IVABRADINE |
|  | KCNJ5 | - |
|  | SCN10A | HEXYLCAINE, BENZOCAINE, PROPARACAINE, ROPIVACAINE, LEVOBUPIVACAINE, DIBUCAINE, PROCAINE, BENOXINATE, CHLOROPROCAINE |
|  | TRDN | - |
|  | CACNA1D | ISRADIPINE, CLEVIDIPINE, NIMODIPINE, NITRENDIPINE |
| Increased circulating troponin I concentration ([HP:0410173](https://hpo.jax.org/app/browse/term/HP:0410173)) | SVIL | - |
|  | ACTB | ETHINYL ESTRADIOL, CYCLOPHOSPHAMIDE |
|  | KDM1A | TRANYLCYPROMINE |
|  | ACTG1 | VINCRISTINE |
|  | FLOT2 | - |
|  | FLOT1 | - |
|  | AR | - |
|  | NEB | - |
|  | POTEF | - |
|  | KIF14 | - |
| Increased circulating troponin T concentration ([HP:0410174](https://hpo.jax.org/app/browse/term/HP:0410174)) | - | - |
| Hypertension ([HP:0000822](https://hpo.jax.org/app/browse/term/HP:0000822)) | BBS1 | - |
|  | BBS2 | - |
|  | MKS1 | - |
|  | SDCCAG8 | - |
|  | BBS5 | - |
|  | BBS7 | - |
|  | BBS12 | - |
|  | BBS10 | - |
|  | WDPCP | - |
|  | CCDC28B | - |
| Hypotension ([HP:0002615](https://hpo.jax.org/app/browse/term/HP:0002615)) | REN | ALISKIREN, ALISKIREN FUMARATE |
|  | KCNJ1 | GLYMIDINE, ACETOHEXAMIDE, MINOXIDIL, GLIMEPIRIDE, TOLBUTAMIDE, GLICLAZIDE, TOLAZAMIDE, GLIPIZIDE, CHLORPROPAMIDE |
|  | SCNN1A | AMILORIDE, TRIAMTERENE, AMILORIDE HYDROCHLORIDE |
|  | NR3C2 | EPLERENONE, DESOXYCORTICOSTERONE PIVALATE, DROSPIRENONE, SPIRONOLACTONE, DESOXYCORTICOSTERONE ACETATE, FLUDROCORTISONE ACETATE, ONAPRISTONE |
|  | SLC12A3 | POLYTHIAZIDE, BENDROFLUMETHIAZIDE, QUINETHAZONE, METOLAZONE, BENZTHIAZIDE, TRICHLORMETHIAZIDE, CHLOROTHIZIDE, CHLOROTHIAZIDE SODIUM, INDAPAMIDE, HYDROCHL OROTHIAZIDE |
|  | CLCNKB | - |
|  | SLC12A1 | HYDROFLUMETHIAZIDE, METHYCLOTHIAZIDE, CHLORMERODRIN, FUROSEMIDE, TRICHLORMETHIAZIDE, BUMETANIDE, CHLORTHALIDONE, TORSEMIDE, METOLAZONE, BENDROFLUMETHIAZIDE |
|  | ACE | CILAZAPRIL, SPIRAPRIL, MOEXIPRIL, FOSINOPRIL, LISINOPRIL, QUINAPRIL, PERINDOPRIL, BENAZEPRIL, TRANDOLAPRIL, CAPTOPRIL |
|  | LHX4 | - |
|  | POU1F1 | - |
| Myocarditis ([HP:0012819](https://hpo.jax.org/app/browse/term/HP:0012819)) | - | - |
| Pericardial effusion ([HP:0001698](https://hpo.jax.org/app/browse/term/HP:0001698)) | MYBPC3 | - |
|  | CCBE1 | - |
|  | HLA-DRB1 | CLAVULANIC ACID, PITAVASTATIN |
|  | CLCNKB | - |
|  | ABCC9 | NICORANDIL, PINACIDIL, TOLBUTAMIDE, GLIMEPIRIDE |
|  | ADAMTS3 | - |
|  | PTPN14 | - |
|  | ENPP1 | - |
|  | SLC12A3 | POLYTHIAZIDE, BENDROFLUMETHIAZIDE, QUINETHAZONE, METOLAZONE, BENZTHIAZIDE, TRICHLORMETHIAZIDE, CHLOROTHIZIDE, CHLOROTHIAZIDE SODIUM, INDAPAMIDE, HYDROCHL OROTHIAZIDE |
|  | ABCC6 | SODIUM CHLORIDE, THALIDOMIDE |
| Reduced ejection fraction ([HP:0012664](https://hpo.jax.org/app/browse/term/HP:0012664)) | GTPBP3 | - |
|  | POLG2 | - |
|  | NPPA | CLORTHALIDONE, AMLODIPINE |
|  | C10orf2 | - |
|  | RPL3L | - |
|  | SCN5A | INDECAINIDE, BENZONATATE, FOSPHENYTOIN, MORICIZINE, HEXYLCAINE, PROCAINAMIDE, MEXILETINE, DISOPYRAMIDE |
|  | FHOD3 | - |
|  | PPCS | - |
|  | COA6 | - |
| Tachycardia ([HP:0001649](https://hpo.jax.org/app/browse/term/HP:0001649)) | SCN5A | INDECAINIDE, BENZONATATE, FOSPHENYTOIN, MORICIZINE, HEXYLCAINE, PROCAINAMIDE, MEXILETINE, DISOPYRAMIDE |
|  | HCN4 | IVABRADINE HYDROCHLORIDE, DRONEDARONE HYDROCHLORIDE, IVABRADINE |
|  | KCND3 | - |
|  | CACNB2 | FELODIPINE, AMLODIPINE |
|  | KCNE3 | - |
|  | SCN1B | ZONISAMIDE |
|  | ABCC9 | NICORANDIL, PINACIDIL, TOLBUTAMIDE, GLIMEPIRIDE |
|  | SCN3B | ZONISAMIDE |
|  | RYR2 | PROCAINE |
|  | KCNJ5 | - |
| Venous thrombosis ([HP:0004936](https://hpo.jax.org/app/browse/term/HP:0004936)) | CTNNB1 | FLUORESCEN SODIUM |
|  | CASR | CINACALCET, VELCALCETIDE, CINACALCET HYDROCHLORIDE, PAMIDRONIC ACID |
|  | PRSS3P2 | - |
|  | SMAD4 | LYSINE, ALECTINIB |
|  | CTLA4 | IPILIMUMAB, ABATACEPT |
|  | PRSS1 | - |
|  | CTRC | - |
|  | GNAQ | VERTEPORFIN |
|  | MET | CRIZOTINIB, OSIMERTINIB |
|  | PTPN22 | - |

### Table 11 Cardiovascular-symptom

| **Symptom (HPO ID)** | **Hub Genes (ranked)** | **FDA-Approved repurposed drugs** |
| --- | --- | --- |
| Angina pectoris ([HP:0001681](https://hpo.jax.org/app/browse/term/HP:0001681)) | ABCG5 | - |
|  | ABCG8 | ATORVASTATIN |
|  | CYP27A1 | CHOLESTYRAMINE |
|  | ABCC6 | SODIUM CHLORIDE, THALIDOMIDE |
|  | LIPC | PENTONSAN POLYSULFATE SODIUM, FLUVASTATTIN |
|  | ENPP1 | - |
|  | PTPN22 | - |
|  | CTLA4 | IPILIMUMAB, ABATACEPT |
|  | ZMPSTE24 | - |
|  | GLA | MIGALASTAT, PINDOLOL |
| Arrhythmia ([HP:0011675](https://hpo.jax.org/app/browse/term/HP:0011675)) | MYH6 | - |
|  | ACTN2 | - |
|  | TNNI3 | - |
|  | MYL2 | - |
|  | MYBPC3 | - |
|  | CSRP3 | - |
|  | MYH7 | - |
|  | RYR2 | PROCAINE |
|  | LDB3 | - |
|  | MYL3 | - |
| Palpitations ([HP:0001962](https://hpo.jax.org/app/browse/term/HP:0001962)) | SCN5A | INDECAINIDE, BENZONATATE, FOSPHENYTOIN, MORICIZINE, HEXYLCAINE, PROCAINAMIDE, MEXILETINE, DISOPYRAMIDE |
|  | MYH6 | - |
|  | MYL2 | - |
|  | NKX2-5 | - |
|  | NPPA | CLORTHALIDONE, AMLODIPINE |
|  | TBX20 | - |
|  | GATA4 | WARFARIN |
|  | MYL3 | - |
|  | CAV3 | - |
|  | KCND3 | - |
| Stroke ([HP:0001297](https://hpo.jax.org/app/browse/term/HP:0001297)) | NKX2-5 | - |
|  | MYH6 | - |
|  | TNNI3 | - |
|  | SCN5A | INDECAINIDE, BENZONATATE, FOSPHENYTOIN, MORICIZINE, HEXYLCAINE, PROCAINAMIDE, MEXILETINE, DISOPYRAMIDE |
|  | NPPA | CLORTHALIDONE, AMLODIPINE |
|  | GATA4 | WARFARIN |
|  | TBX20 | - |
|  | MYBPC3 | - |
|  | ACTA2 | - |
|  | COL3A1 | COLLAGENASE CLOSTRIDIUM HISTOLYTICUM, OCRIPLASMIN |
| Syncope ([HP:0001279](https://hpo.jax.org/app/browse/term/HP:0001279)) | SCN5A | INDECAINIDE, BENZONATATE, FOSPHENYTOIN, MORICIZINE, HEXYLCAINE, PROCAINAMIDE, MEXILETINE, DISOPYRAMIDE |
|  | HCN4 | IVABRADINE HYDROCHLORIDE, DRONEDARONE HYDROCHLORIDE, IVABRADINE |
|  | KCND3 | - |
|  | CACNB2 | FELODIPINE, AMLODIPINE |
|  | KCNE2 |  |
|  | KCNJ5 | - |
|  | SCN1B | ZONISAMIDE |
|  | ANK2 | - |
|  | KCNE3 | - |
|  | RYR2 | PROCAINE |

### Table 12 gi-finding

| **Symptom (HPO ID)** | **Hub Genes (ranked)** | **FDA-Approved repurposed drugs** |
| --- | --- | --- |
| Abnormal pancreas morphology ([HP:0012090](https://hpo.jax.org/app/browse/term/HP:0012090)) | RPGRIP1L | - |
|  | TMEM216 | - |
|  | MKS1 | - |
|  | TCTN2 | - |
|  | TCTN3 | - |
|  | CC2D2A | - |
|  | B9D2 | - |
|  | B9D1 | - |
|  | TMEM231 | - |
|  | NPHP3 | - |
| Gastric ulcer ([HP:0002592](https://hpo.jax.org/app/browse/term/HP:0002592)) | ARID1B | - |
|  | SMARCA4 | ABEMACICLIB, RIBOCICLIB |
|  | SMARCC1 | - |
|  | SMARCA2 | - |
|  | SMARCB1 | PANOBINOSTAT |
|  | SMARCC2 | - |
|  | SMARCD1 | - |
|  | SMARCE1 | - |
|  | ACTL6A | - |
|  | SS18 | - |
| Gastroesophageal reflux ([HP:0002020](https://hpo.jax.org/app/browse/term/HP:0002020)) | GRIN1 | MEMANTINE, ORPHENADRINE |
|  | GRIN2B | MEMANTINE, FELBAMATE, ACAMPROSATE |
|  | CAMK2A | - |
|  | GABRG2 | - |
|  | GABRB2 | - |
|  | NRXN1 | DULOXETINE |
|  | CACNA1B | ZICONOTIDE, ZICONOTIDE ACETATE, AMLODIPINE, LEVETIRACETAM |
|  | CAMK2B | - |
|  | ERBB4 | DACOMITINIB |
|  | HRAS | - |
| Gastroparesis ([HP:0002578](https://hpo.jax.org/app/browse/term/HP:0002578)) | SNRPN | - |
|  | MAGEL2 | - |
|  | OCA2 | - |
|  | NDN | - |
|  | C10orf2 | - |
|  | POLG2 | - |
|  | DNAJC6 | - |
|  | ACTG2 | - |
|  | MGAT2 | - |
|  | TMEM70 | - |
| Hepatic steatosis ([HP:0001397](https://hpo.jax.org/app/browse/term/HP:0001397)) | ACADS | - |
|  | ETFDH | METHYLPHENIDATE |
|  | CPT2 | - |
|  | HADH | - |
|  | SLC25A20 | - |
|  | ACOX1 | - |
|  | ACAD9 | - |
|  | SLC22A5 | IMATINIB |
|  | PLIN1 | SUFENTANIL CITRATE, CITRIC ACID, ALVERINE CITRATE, TOREMIFENE CITRATE |
|  | CIDEC | - |
| Hepatitis ([HP:0012115](https://hpo.jax.org/app/browse/term/HP:0012115)) | SYK | FOSTAMATINIB |
|  | PIK3CA | - |
|  | CD40LG | - |
|  | CD79B | POLATUZUMAB VEDOTIN |
|  | CD79A | - |
|  | CD3D | MUROMMONAB-CD3, CATUMAXOMAB, BLINATUMOMAB |
|  | CD3E | MUROMMONAB-CD3, CATUMAXOMAB, BLINATUMOMAB |
|  | BTK | IBRUTINIB |
|  | FASLG | - |
|  | FAS | TENIPOSIDE |
| Hepatomegaly ([HP:0002240](https://hpo.jax.org/app/browse/term/HP:0002240)) | CTLA4 | IPILIMUMAB, ABATACEPT |
|  | CD40LG | - |
|  | CD3E | MUROMMONAB-CD3, CATUMAXOMAB, BLINATUMOMAB |
|  | PRF1 | EMAPALUMAB |
|  | ZAP70 | - |
|  | CD3D | MUROMMONAB-CD3, CATUMAXOMAB, BLINATUMOMAB |
|  | CD19 | BLINATUMOMAB |
|  | IL2RA | BASILIXIMAB, DACLIZUMAB, ALDESLEUKIN, DENILEUKIN DIFTITOX, CETIRIZINE |
|  | RAG1 | - |
|  | RAG2 | - |
| Malnutrition ([HP:0004395](https://hpo.jax.org/app/browse/term/HP:0004395)) | MSX1 | - |
|  | PVRL1 | - |
|  | IRF6 | - |
|  | SBDS | - |
|  | SLC5A1 | CANAGLIFLOZIN, EMPAGLIFLOZIN, DAPAGLIFLOZIN, ERTUGLIFLOZIN |
|  | SLC7A7 | - |
|  | EFTUD1 | - |
|  | HLA-DQA1 | LAPATINIB, AZATHIOPRINE, MERCAPTOPURINE |
|  | ACTG2 | - |
|  | CRLF1 | - |
| Splenomegaly ([HP:0001744](https://hpo.jax.org/app/browse/term/HP:0001744)) | CD40LG | - |
|  | CD19 | BLINATUMOMAB |
|  | CTLA4 | IPILIMUMAB, ABATACEPT |
|  | IL2RA | BASILIXIMAB, DACLIZUMAB, ALDESLEUKIN, DENILEUKIN DIFTITOX, CETIRIZINE |
|  | ZAP70 | - |
|  | CD3E | MUROMMONAB-CD3, CATUMAXOMAB, BLINATUMOMAB |
|  | CD3D | MUROMMONAB-CD3, CATUMAXOMAB, BLINATUMOMAB |
|  | PRF1 | EMAPALUMAB |
|  | RAG1 | - |
|  | RAG2 | - |

### Table 13 gi-symptom

| **Symptom (HPO ID)** | **Hub Genes (ranked)** | **FDA-Approved repurposed drugs** |
| --- | --- | --- |
| Abdominal pain ([HP:0002027](https://hpo.jax.org/app/browse/term/HP:0002027)) | KRAS | PANITUMUMAB, CETUXIMAB |
|  | SMAD4 | LYSINE, ALECTINIB |
|  | CTNNB1 | FLUORESCEN SODIUM |
|  | ESR1 | CLOMIPHENE, LEVONORGESTREL |
|  | PIK3CA | - |
|  | RET | CABOZANTINIB, VANDETANIB |
|  | KIT | NILOTINIB, IMATINIB |
|  | NF1 | TRAMETINIB, BINIMETINIB, COBIMETINIB, DABRAFENIB, VEMURAFENIB |
|  | BRCA1 | - |
|  | CDKN1B | RALTITREXED |
| Abdominal symptom ([HP:0011458](https://hpo.jax.org/app/browse/term/HP:0011458)) | SCN2A | - |
|  | SCN1A | ETHOSUXIMIDE |
|  | SCN8A | - |
|  | SCN9A | - |
|  | SCN5A | INDECAINIDE, BENZONATATE, FOSPHENYTOIN, MORICIZINE, HEXYLCAINE, PROCAINAMIDE, MEXILETINE, DISOPYRAMIDE |
|  | SCN11A | - |
|  | SCN10A | HEXYLCAINE, BENZOCAINE, PROPARACAINE, ROPIVACAINE, LEVOBUPIVACAINE, DIBUCAINE, PROCAINE, BENOXINATE, CHLOROPROCAINE |
|  | SCN1B | ZONISAMIDE |
|  | SCN4A | LEVOBUPIVACAINE HYDROCHLORIDE, BUPIVACAINE HYDROCHLORIDE |
|  | SCN3A | - |
| Anorexia ([HP:0002039](https://hpo.jax.org/app/browse/term/HP:0002039)) | BRCA1 | - |
|  | SMAD4 | LYSINE, ALECTINIB |
|  | KRAS | PANITUMUMAB, CETUXIMAB |
|  | CDKN1B | RALTITREXED |
|  | PTPN22 | - |
|  | CD3D | MUROMMONAB-CD3, CATUMAXOMAB, BLINATUMOMAB |
|  | CD3E | MUROMMONAB-CD3, CATUMAXOMAB, BLINATUMOMAB |
|  | SYK | FOSTAMATINIB |
|  | MEN1 | OLSALAZINE |
|  | PALB2 | OLAPARIB, MITOMYCIN |
| Bowel incontinence ([HP:0002607](https://hpo.jax.org/app/browse/term/HP:0002607)) | TBX1 | - |
|  | VANGL1 | - |
|  | RTN2 | - |
|  | FUZ | - |
|  | LRIG2 | - |
|  | PLP1 | - |
|  | HPSE2 | - |
|  | GP1BB | - |
|  | C19orf12 | - |
|  | COMT | ENTACAPONE, OPICAPONE, REMIFENTANIL, SUFENTANIL, MODAFINIL, MORPHINE, METHADONE |
| Constipation ([HP:0002019](https://hpo.jax.org/app/browse/term/HP:0002019)) | SHH | VISMODEGIB, CAFFEINE |
|  | FGF8 | - |
|  | OTX2 | - |
|  | SIX3 | - |
|  | SOX3 | - |
|  | ZIC2 | - |
|  | GLI2 | - |
|  | FOXG1 | - |
|  | SOX10 | VEMURAFENIB |
|  | LHX3 | - |
| Diarrhea ([HP:0002014](https://hpo.jax.org/app/browse/term/HP:0002014)) | CTLA4 | IPILIMUMAB, ABATACEPT |
|  | CD19 | BLINATUMOMAB |
|  | CD79A | - |
|  | CD79B | POLATUZUMAB VEDOTIN |
|  | CD3E | MUROMMONAB-CD3, CATUMAXOMAB, BLINATUMOMAB |
|  | ZAP70 | - |
|  | CD3D | MUROMMONAB-CD3, CATUMAXOMAB, BLINATUMOMAB |
|  | RAG2 | - |
|  | RAG1 | - |
|  | CD40LG | - |
| Nausea ([HP:0002018](https://hpo.jax.org/app/browse/term/HP:0002018)) | RET | CABOZANTINIB, VANDETANIB |
|  | NF1 | TRAMETINIB, BINIMETINIB, COBIMETINIB, DABRAFENIB, VEMURAFENIB |
|  | SDHC | - |
|  | SDHAF2 | - |
|  | MAX | - |
|  | KIF1B | - |
|  | ESR1 | CLOMIPHENE, LEVONORGESTREL |
|  | MEN1 | OLSALAZINE |
|  | KIT | NILOTINIB, IMATINIB |
|  | KCNJ5 | - |
| Poor appetite ([HP:0004396](https://hpo.jax.org/app/browse/term/HP:0004396)) | PALB2 | OLAPARIB, MITOMYCIN |
|  | BRCA1 | - |
|  | SMAD4 | LYSINE, ALECTINIB |
|  | KRAS | PANITUMUMAB, CETUXIMAB |
|  | ACTA1 | - |
|  | PALLD | - |
|  | TPM3 | CORTICOTROPIN |
|  | MYL2 | - |
|  | NAGS | - |
|  | SLC25A13 | - |
| Vomiting ([HP:0002013](https://hpo.jax.org/app/browse/term/HP:0002013)) | NDUFS3 | - |
|  | NDUFAF4 | - |
|  | NDUFA1 | - |
|  | NDUFA6 | - |
|  | NDUFB10 | - |
|  | NDUFB11 | - |
|  | NDUFS1 | - |
|  | TIMMDC1 | - |
|  | CTNNB1 | FLUORESCEN SODIUM |
|  | SHH | VISMODEGIB, CAFFEINE |

### Table 14 Pulmonary-finding

| **Symptom (HPO ID)** | **Hub Genes (ranked)** | **FDA-Approved repurposed drugs** |
| --- | --- | --- |
| Abnormality on pulmonary function testing ([HP:0030878](https://hpo.jax.org/app/browse/term/HP:0030878)) | RAPSN | - |
|  | CHRNA1 | - |
|  | DOK7 | - |
|  | MUSK | - |
|  | SCN4A | LEVOBUPIVACAINE HYDROCHLORIDE, BUPIVACAINE HYDROCHLORIDE |
|  | CHRND | SUCCINYLCHOLINE CHLORIDE, PIPECURONIUM BROMIDE, MIVACURIUM CHLORIDE, METOCURINE IODIDE, DOXACURIUM CHLORIDE, VECURONIUM BROMIDE, DECAMETHONIUM BROMIDE |
|  | CHRNB1 | SUCCINYLCHOLINE CHLORIDE, PIPECURONIUM BROMIDE, MIVACURIUM CHLORIDE, METOCURINE IODIDE, DOXACURIUM CHLORIDE, VECURONIUM BROMIDE, DECAMETHONIUM BROMIDE |
|  | CHRNE | SUCCINYLCHOLINE CHLORIDE, PIPECURONIUM BROMIDE, MIVACURIUM CHLORIDE, METOCURINE IODIDE, DOXACURIUM CHLORIDE, VECURONIUM BROMIDE, DECAMETHONIUM BROMIDE |
|  | CTLA4 | IPILIMUMAB, ABATACEPT |
|  | CD19 | BLINATUMOMAB |
| Airway obstruction ([HP:0006536](https://hpo.jax.org/app/browse/term/HP:0006536)) | CCDC65 | - |
|  | DRC1 | - |
|  | HYDIN | - |
|  | RSPH4A | - |
|  | RSPH9 | - |
|  | RSPH1 | - |
|  | CCDC103 | - |
|  | HEATR2 | - |
|  | DYX1C1 | - |
|  | DNAI2 | - |
| Decreased DLCO ([HP:0045051](https://hpo.jax.org/app/browse/term/HP:0045051)) | - | - |
| Ground-glass opacification ([HP:0025179](https://hpo.jax.org/app/browse/term/HP:0025179)) | TERT | OMACETAXINE MEPESUCCINATE |
|  | RTEL1 | - |
|  | OBFC1 | ATENOLOL |
|  | ABCA3 | - |
|  | DHX36 | - |
|  | NKX2-1 | - |
|  | DSP | ENALAPRIL |
|  | STON1 | - |
|  | BMPR2 | - |
| Hypoxemia ([HP:0012418](https://hpo.jax.org/app/browse/term/HP:0012418)) | NKX2-1 | - |
|  | GATA6 | - |
|  | TPM3 | CORTICOTROPIN |
|  | MYL2 | - |
|  | ACTA1 | - |
|  | HLA-DRB1 | CLAVULANIC ACID, PITAVASTATIN |
|  | ZFPM2 | - |
|  | ABCA3 | - |
|  | BTNL2 | - |
|  | RHAG | - |
| Oxygen desaturation on exertion ([HP:0030874](https://hpo.jax.org/app/browse/term/HP:0030874)) | NKX2-1 | - |
|  | PAX8 | - |
|  | NAPSA | - |
|  | KRT7 | - |
|  | TG | ACEBUTOLOL, AMINOGLETETHIMIDE |
|  | SMAD3 | -- |
|  | NCOA1 | - |
|  | FOXA2 | - |
|  | FOXA1 | - |
|  | SFTPB | STREPTOZOCIN |
| Pleuritis ([HP:0002102](https://hpo.jax.org/app/browse/term/HP:0002102)) | CTLA4 | IPILIMUMAB, ABATACEPT |
|  | PTPN22 | - |
|  | FCGR2B | - |
|  | IL23R | - |
|  | FAS | TENIPOSIDE |
|  | MEFV | - |
|  | TREX1 | - |
| Pulmonary embolism ([HP:0002204](https://hpo.jax.org/app/browse/term/HP:0002204)) | SMAD4 | LYSINE, ALECTINIB |
|  | GDF2 | - |
|  | GNAQ | VERTEPORFIN |
|  | CBSL | - |
|  | ENSP00000381231 | - |
|  | COIL | - |
|  | TGS1 | - |
|  | MMACHC | - |
|  | MEFV | - |
|  | KCNJ5 | - |
| Reduced forced expiratory volume in one second ([HP:0032342](https://hpo.jax.org/app/browse/term/HP:0032342)) | RTEL1 | - |
|  | DNA2 | - |
|  | WRN | - |
|  | RAD51 | - |
|  | PIF1 | - |
|  | BLM | - |
|  | EXO1 | CAPECITABINE |
|  | TERF1 | - |
|  | MMS19 | - |
|  | FAM96B | - |
| Reduced FEV1/FVC ratio ([HP:0030877](https://hpo.jax.org/app/browse/term/HP:0030877)) | - | - |
| Reduced forced vital capacity ([HP:0032341](https://hpo.jax.org/app/browse/term/HP:0032341)) | MYH7 | - |
|  | ACTA1 | - |
|  | UNC45B | - |
|  | TPM3 | CORTICOTROPIN |
|  | FKRP | - |
|  | SLC25A21 | - |
|  | MCIDAS | - |
|  | GGPS1 | ZOLEDRONIC ACID |
|  | RTEL1 | - |
|  | SYT2 | - |
| Reduced vital capacity ([HP:0002792](https://hpo.jax.org/app/browse/term/HP:0002792)) | RAPSN | - |
|  | CHRNA1 | - |
|  | CHRND | SUCCINYLCHOLINE CHLORIDE, PIPECURONIUM BROMIDE, MIVACURIUM CHLORIDE, METOCURINE IODIDE, DOXACURIUM CHLORIDE, VECURONIUM BROMIDE, DECAMETHONIUM BROMIDE |
|  | CHRNE | SUCCINYLCHOLINE CHLORIDE, PIPECURONIUM BROMIDE, MIVACURIUM CHLORIDE, METOCURINE IODIDE, DOXACURIUM CHLORIDE, VECURONIUM BROMIDE, DECAMETHONIUM BROMIDE |
|  | CHRNB1 | SUCCINYLCHOLINE CHLORIDE, PIPECURONIUM BROMIDE, MIVACURIUM CHLORIDE, METOCURINE IODIDE, DOXACURIUM CHLORIDE, VECURONIUM BROMIDE, DECAMETHONIUM BROMIDE |
|  | DOK7 | - |
|  | MUSK | - |
|  | SCN4A | LEVOBUPIVACAINE HYDROCHLORIDE, BUPIVACAINE HYDROCHLORIDE |
|  | ACTA1 | - |
|  | MYH7 | - |
| Restrictive ventilatory defect ([HP:0002091](https://hpo.jax.org/app/browse/term/HP:0002091)) | SCN4A | LEVOBUPIVACAINE HYDROCHLORIDE, BUPIVACAINE HYDROCHLORIDE |
|  | CHRNE | SUCCINYLCHOLINE CHLORIDE, PIPECURONIUM BROMIDE, MIVACURIUM CHLORIDE, METOCURINE IODIDE, DOXACURIUM CHLORIDE, VECURONIUM BROMIDE, DECAMETHONIUM BROMIDE |
|  | RAPSN | - |
|  | CHRND | SUCCINYLCHOLINE CHLORIDE, PIPECURONIUM BROMIDE, MIVACURIUM CHLORIDE, METOCURINE IODIDE, DOXACURIUM CHLORIDE, VECURONIUM BROMIDE, DECAMETHONIUM BROMIDE |
|  | CHRNA1 | SUCCINYLCHOLINE CHLORIDE, PIPECURONIUM BROMIDE, MIVACURIUM CHLORIDE, METOCURINE IODIDE, DOXACURIUM CHLORIDE, VECURONIUM BROMIDE, DECAMETHONIUM BROMIDE |
|  | CHRNB1 | SUCCINYLCHOLINE CHLORIDE, PIPECURONIUM CHLORIDE, MIVACURIUM CHLORIDE, METOCURINE IODIDE, DOXACURIUM CHLORIDE, VECURONIUM BROMIDE, DECAMETHONIUM BROMIDE |
|  | DOK7 | - |
|  | MUSK | - |
|  | CTLA4 | IPILIMUMAB, ABATACEPT |
|  | CD19 | BLINATUMOMAB |

### Table 15 Pulmonary-imaging

| **Symptom (HPO ID)** | **Hub Genes (ranked)** | **FDA-Approved repurposed drugs** |
| --- | --- | --- |
| Abnormal pulmonary thoracic imaging finding ([HP:0031983](https://hpo.jax.org/app/browse/term/HP:0031983)) | DRC1 | - |
|  | CCDC65 | - |
|  | HYDIN | - |
|  | RSPH4A | - |
|  | RSPH1 | - |
|  | CCDC103 | - |
|  | HEATR2 | - |
|  | DYX1C1 | - |
|  | DNAI2 | - |
|  | CCDC40 | - |
| Atelectasis ([HP:0100750](https://hpo.jax.org/app/browse/term/HP:0100750)) | CCDC65 | - |
|  | DRC1 | - |
|  | HYDIN | - |
|  | RSPH4A | - |
|  | RSPH1 | - |
|  | CCDC103 | - |
|  | HEATR2 | - |
|  | DYX1C1 | - |
|  | DNAI2 | - |
|  | CCDC40 | - |
| Bronchiectasis ([HP:0002110](https://hpo.jax.org/app/browse/term/HP:0002110)) | DRC1 | - |
|  | CCDC65 | - |
|  | HYDIN | - |
|  | RSPH4A | - |
|  | RSPH1 | - |
|  | CCDC103 | - |
|  | HEATR2 | - |
|  | DYX1C1 | - |
|  | DNAI2 | - |
|  | CCDC40 | - |
| Centrilobular ground-glass opacification on pulmonary HRCT ([HP:0025180](https://hpo.jax.org/app/browse/term/HP:0025180)) | BMPR2 | - |
|  | ACVR1 | - |
|  | SMAD6 | - |
|  | BMP4 | - |
|  | BMP2 | - |
|  | BMP10 | - |
|  | BMP7 | PEGASPARGASE |
|  | GDF5 | - |
|  | BMP6 | - |
|  | GDF2 | - |
| Interlobular septal thickening ([HP:0030879](https://hpo.jax.org/app/browse/term/HP:0030879)) | BMPR2 | - |
|  | ACVR1 | - |
|  | SMAD6 | - |
|  | BMP4 | - |
|  | BMP2 | - |
|  | BMP10 | - |
|  | BMP7 | PEGASPARGASE |
|  | GDF5 | - |
|  | BMP6 | - |
|  | GDF2 | - |
| Parenchymal consolidation ([HP:0032177](https://hpo.jax.org/app/browse/term/HP:0032177)) | NKX2-1 | - |
|  | NAPSA | - |
|  | PAX8 | - |
|  | KRT7 | - |
|  | TG | ACEBUTOLOL, AMINOGLETETHIMIDE |
|  | SMAD3 | - |
|  | NCOA1 | - |
|  | FOXA2 | - |
|  | FOXA1 | - |
|  | SFTPB | STREPTOZOCIN |
| Pleural thickening ([HP:0031944](https://hpo.jax.org/app/browse/term/HP:0031944)) | CARD10 | - |
|  | PRKCQ | - |
|  | BCL10 | - |
|  | MALT1 | - |
|  | CARD11 | - |
|  | PRKCB | - |
|  | TRAF6 | - |
|  | IKBKG | SULFADOXINE, ARTESUNATE, PRIMAQUINE, PYRIMETHAMINE |
|  | CARD14 | - |
|  | RRNAD1 | - |
| Pulmonary bulla ([HP:0032446](https://hpo.jax.org/app/browse/term/HP:0032446)) | - | - |
| Pulmonary fibrosis ([HP:0002206](https://hpo.jax.org/app/browse/term/HP:0002206)) | CTLA4 | IPILIMUMAB, ABATACEPT |
|  | PTPN22 | - |
|  | HLA-DRB1 | CLAVULANIC ACID, PITAVASTATIN |
|  | NR5A1 | CORTICOTROPIN, OXYTOCIN |
|  | TERT | OMACETAXINE MEPESUCCINATE |
|  | BMP15 | - |
|  | NKX2-1 | - |
|  | BTNL2 | - |
|  | OBFC1 | ATENOLOL |
|  | FASLG | - |
| Pulmonary interstitial high-resolution computed tomography abnormality ([HP:0025389](https://hpo.jax.org/app/browse/term/HP:0025389)) | TERT | OMACETAXINE MEPESUCCINATE |
|  | OBFC1 | ATENOLOL |
|  | RTEL1 | - |
|  | NKX2-1 | - |
|  | ABCA3 | - |
|  | DHX36 | - |
|  | COL3A1 | COLLAGENASE CLOSTRIDIUM HISTOLYTICUM, OCRIPLASMIN |
|  | HLA-DRB1 | CLAVULANIC ACID, PITAVASTATIN |
|  | BMPR2 | - |
|  | DSP | ENALAPRIL |
| Pulmonary opacity ([HP:0031457](https://hpo.jax.org/app/browse/term/HP:0031457)) | TERT | OMACETAXINE MEPESUCCINATE |
|  | OBFC1 | ATENOLOL |
|  | RTEL1 | - |
|  | NKX2-1 | - |
|  | ABCA3 | - |
|  | DHX36 | - |
|  | DSP | ENALAPRIL |
|  | COL3A1 | COLLAGENASE CLOSTRIDIUM HISTOLYTICUM, OCRIPLASMIN |
|  | BMPR2 | - |
| Reticular pattern on pulmonary HRCT ([HP:0025390](https://hpo.jax.org/app/browse/term/HP:0025390)) | TERT | OMACETAXINE MEPESUCCINATE |
|  | OBFC1 | ATENOLOL |
|  | RTEL1 | - |
|  | DHX36 | - |
|  | ABCA3 | - |
|  | DSP | ENALAPRIL |

### Table 16 Pulmonary-symptom

| **Symptom (HPO ID)** | **Hub Genes (ranked)** | **FDA-Approved repurposed drugs** |
| --- | --- | --- |
| Abnormal breath sound ([HP:0030829](https://hpo.jax.org/app/browse/term/HP:0030829)) | DRC1 | - |
|  | CCDC65 | - |
|  | HYDIN | - |
|  | RSPH4A | - |
|  | RSPH1 | - |
|  | CCDC103 | - |
|  | HEATR2 | - |
|  | DYX1C1 | - |
|  | DNAI2 | - |
|  | CCDC40 | - |
| Abnormal sputum ([HP:0032016](https://hpo.jax.org/app/browse/term/HP:0032016)) | CCDC65 | - |
|  | DRC1 | - |
|  | HYDIN | - |
|  | RSPH4A | - |
|  | RSPH1 | - |
|  | CCDC103 | - |
|  | HEATR2 | - |
|  | DYX1C1 | - |
|  | DNAI2 | - |
|  | CCDC40 | - |
| Cough ([HP:0012735](https://hpo.jax.org/app/browse/term/HP:0012735)) | DRC1 | - |
|  | CCDC65 | - |
|  | HYDIN | - |
|  | RSPH4A | - |
|  | RSPH1 | - |
|  | CCDC103 | - |
|  | HEATR2 | - |
|  | DYX1C1 | - |
|  | DNAI2 | - |
|  | CCDC40 | - |
| Dyspnea ([HP:0002094](https://hpo.jax.org/app/browse/term/HP:0002094)) | DOK7 | - |
|  | MUSK | - |
|  | RAPSN | - |
|  | CHRNE | SUCCINYLCHOLINE CHLORIDE, PIPECURONIUM BROMIDE, MIVACURIUM CHLORIDE, METOCURINE IODIDE, DOXACURIUM CHLORIDE, VECURONIUM BROMIDE, DECAMETHONIUM BROMIDE |
|  | COLQ | - |
|  | SCN4A | LEVOBUPIVACAINE HYDROCHLORIDE, BUPIVACAINE HYDROCHLORIDE |
|  | CHRNA1 | - |
|  | CHRNB1 | SUCCINYLCHOLINE CHLORIDE, PIPECURONIUM BROMIDE, MIVACURIUM CHLORIDE, METOCURINE IODIDE, DOXACURIUM CHLORIDE, VECURONIUM BROMIDE, DECAMETHONIUM BROMIDE |
|  | CHRND | SUCCINYLCHOLINE CHLORIDE, PIPECURONIUM BROMIDE, MIVACURIUM CHLORIDE, METOCURINE IODIDE, DOXACURIUM CHLORIDE, VECURONIUM BROMIDE, DECAMETHONIUM BROMIDE |
|  | MYH6 | - |
| Exertional dyspnea ([HP:0002875](https://hpo.jax.org/app/browse/term/HP:0002875)) | RAPSN | - |
|  | CHRNA1 | - |
|  | SCN4A | LEVOBUPIVACAINE HYDROCHLORIDE, BUPIVACAINE HYDROCHLORIDE |
|  | CHRNE | SUCCINYLCHOLINE CHLORIDE, PIPECURONIUM BROMIDE, MIVACURIUM CHLORIDE, METOCURINE IODIDE, DOXACURIUM CHLORIDE, VECURONIUM BROMIDE, DECAMETHONIUM BROMIDE |
|  | CHRND | SUCCINYLCHOLINE CHLORIDE, PIPECURONIUM BROMIDE, MIVACURIUM CHLORIDE, METOCURINE IODIDE, DOXACURIUM CHLORIDE, VECURONIUM BROMIDE, DECAMETHONIUM BROMIDE |
|  | COLQ | - |
|  | CHRNB1 | SUCCINYLCHOLINE CHLORIDE, PIPECURONIUM BROMIDE, MIVACURIUM CHLORIDE, METOCURINE IODIDE, DOXACURIUM CHLORIDE, VECURONIUM BROMIDE, DECAMETHONIUM BROMIDE |
|  | DOK7 | - |
|  | MUSK | - |
|  | NKX2-5 | - |
| Hemoptysis ([HP:0002105](https://hpo.jax.org/app/browse/term/HP:0002105)) | ACTA2 | - |
|  | COL3A1 | COLLAGENASE CLOSTRIDIUM HISTOLYTICUM, OCRIPLASMIN |
|  | MFAP5 | - |
|  | COL5A2 | COLLAGENASE CLOSTRIDIUM HISTOLYTICUM, OCRIPLASMIN |
|  | ELN | - |
|  | IL23R | - |
|  | IL17F | - |
|  | IL17RA | BRODALUMAB |
|  | PRKG1 | - |
|  | CTLA4 | IPILIMUMAB, ABATACEPT |
| Nonproductive cough ([HP:0031246](https://hpo.jax.org/app/browse/term/HP:0031246)) | - | - |
| Productive cough ([HP:0031245](https://hpo.jax.org/app/browse/term/HP:0031245)) | CCDC65 | - |
|  | DRC1 | - |
|  | HYDIN | - |
|  | RSPH4A | - |
|  | RSPH1 | - |
|  | CCDC103 | - |
|  | HEATR2 | - |
|  | DYX1C1 | - |
|  | DNAI2 | - |
|  | CCDC40 | - |
| Rhinorrhea ([HP:0031417](https://hpo.jax.org/app/browse/term/HP:0031417)) | CCDC40 | - |
|  | CCDC39 | - |
|  | LRRC56 | - |
|  | SCN9A | - |
| Tachypnea ([HP:0002789](https://hpo.jax.org/app/browse/term/HP:0002789)) | TMEM231 | - |
|  | TMEM138 | - |
|  | B9D1 | - |
|  | CC2D2A | - |
|  | CEP41 | - |
|  | TCTN3 | - |
|  | TCTN2 | - |
|  | RPGRIP1L | - |
|  | MKS1 | - |
|  | TMEM216 | - |
| Wheezing ([HP:0030828](https://hpo.jax.org/app/browse/term/HP:0030828)) | CCDC65 | - |
|  | DRC1 | - |
|  | HYDIN | - |
|  | RSPH4A | - |
|  | RSPH1 | - |
|  | CCDC103 | - |
|  | HEATR2 | - |
|  | DYX1C1 | - |
|  | DNAI2 | - |
|  | CCDC40 | - |

### Table 17 Neuropsychiatric-behavioral

| **Symptom (HPO ID)** | **Hub Genes (ranked)** | **FDA-Approved repurposed drugs** |
| --- | --- | --- |
| Anxiety ([HP:0000739](https://hpo.jax.org/app/browse/term/HP:0000739)) | GPR98 | - |
|  | USH1C | - |
|  | USH2A | - |
|  | USH1G | - |
|  | PCDH15 | - |
|  | DFNB31 | - |
|  | CIB2 | - |
|  | CDH23 | METHYLPHENIDATE |
|  | CLRN1 | - |
|  | PDZD7 | - |
| Apathy ([HP:0000741](https://hpo.jax.org/app/browse/term/HP:0000741)) | ZIC2 | - |
|  | SIX3 | - |
|  | SHH | VISMODEGIB, CAFFEINE |
|  | GLI2 | - |
|  | FGF8 | - |
|  | CDON | - |
|  | MAPT | - |
|  | NODAL | - |
|  | C9orf72 | - |
|  | ATXN10 | - |
| Attention deficit hyperactivity disorder ([HP:0007018](https://hpo.jax.org/app/browse/term/HP:0007018)) | GABRG2 | - |
|  | GABRA1 | METHOHEXITAL, BARBITAL, THIAMYLAL, ZALEPLON, ZOPICLONE, ZOLPIDEM |
|  | SCN1A | ETHOSUXIMIDE |
|  | KCNA2 | - |
|  | KCNT1 | CLOFILIUM, LOXAPINE, QUINIDINE |
|  | SCN8A | - |
|  | GRIN2A | MEMANTINE, GLYCINE, DEXTROMETHORPHAN POLISTIREX |
|  | SLC6A1 | TIAGABINE, TIAGABINE HYDROCHLORIDE |
|  | CACNA1B | ZICONOTIDE, ZICONOTIDE ACETATE, AMLODIPINE, LEVETIRACETAM |
|  | ATP1A3 | ACETYLDIGITOXIN, DESLANOSIDE, DIGITOXIN |
| Auditory hallucinations ([HP:0008765](https://hpo.jax.org/app/browse/term/HP:0008765)) | - | - |
| Behavioral abnormalities ([HP:0000708](https://hpo.jax.org/app/browse/term/HP:0000708)) | GUCA1B | - |
|  | PRPH2 | - |
|  | NR2E3 | - |
|  | CRX | - |
|  | CNGA1 | DEQUALINIUM |
|  | NRL | - |
|  | PDE6G | PENTOXIFYLLINE |
|  | RPGR | - |
|  | RDH12 | - |
|  | TULP1 | - |
| Delusions ([HP:0000746](https://hpo.jax.org/app/browse/term/HP:0000746)) | COMT | ENTACAPONE, OPICAPONE, REMIFENTANIL, SUFENTANIL, MODAFINIL, MORPHINE, METHADONE |
|  | DGCR2 | - |
|  | ENSP00000331681 | - |
|  | TBX1 | - |
|  | EIF2B2 | - |
|  | DAOA | - |
|  | EIF2B3 | - |
|  | JPH3 | - |
|  | EIF2B1 | - |
|  | DGCR14 | - |
| Hallucinations ([HP:0000738](https://hpo.jax.org/app/browse/term/HP:0000738)) | GPR98 | - |
|  | USH1C | - |
|  | USH2A | - |
|  | DFNB31 | - |
|  | CIB2 | - |
|  | CDH23 | METHYLPHENIDATE |
|  | USH1G | - |
|  | PCDH15 | - |
|  | CLRN1 | - |
|  | PDZD7 | - |
| Impulsivity ([HP:0100710](https://hpo.jax.org/app/browse/term/HP:0100710)) | SCN1A | ETHOSUXIMIDE |
|  | SCN2A | - |
|  | KCNA2 | - |
|  | GABRG2 | - |
|  | GABRA1 | METHOHEXITAL, BARBITAL, THIAMYLAL, ZALEPLON, ZOPICLONE, ZOLPIDEM |
|  | SCN8A | - |
|  | PCDH19 | - |
|  | GRIN2A | MEMANTINE, GLYCINE, DEXTROMETHORPHAN POLISTIREX |
|  | SCN1B | ZONISAMIDE |
|  | CACNA1B | ZICONOTIDE, ZICONOTIDE ACETATE, AMLODIPINE, LEVETIRACETAM |
| Irritability ([HP:0000737](https://hpo.jax.org/app/browse/term/HP:0000737)) | SHH | VISMODEGIB, CAFFEINE |
|  | SIX3 | - |
|  | FGF8 | - |
|  | ZIC2 | - |
|  | PAX2 | PROGESTERONE |
|  | GLI2 | - |
|  | FOXG1 | - |
|  | NPHS1 | LOSARTAN |
|  | TH | TYROSINE, METYROSINE, IOBENGUANE |
|  | AQP2 | - |
| Obsessive-compulsive disorder ([HP:0000722](https://hpo.jax.org/app/browse/term/HP:0000722)) | GABRG2 | - |
|  | SCN1A | ETHOSUXIMIDE |
|  | SCN2A | - |
|  | GABRA1 | METHOHEXITAL, BARBITAL, THIAMYLAL, ZALEPLON, ZOPICLONE, ZOLPIDEM |
|  | GABRD | - |
|  | SCN1B | ZONISAMIDE |
|  | GRIA2 | PERAMPANEL |
|  | SCN9A | - |
|  | PCDH19 | - |
|  | EP300 | - |
| Panic attack ([HP:0025269](https://hpo.jax.org/app/browse/term/HP:0025269)) | MAX | - |
|  | NF1 | TRAMETINIB, BINIMETINIB, COBIMETINIB, DABRAFENIB, VEMURAFENIB |
|  | SDHC | - |
|  | KIF1B | - |
|  | SDHAF2 | - |
|  | RET | CABOZANTINIB, VANDETANIB |
|  | EP300 | - |
|  | BPTF | - |
|  | CREBBP | - |
|  | TRIP12 | - |
| Paranoia ([HP:0011999](https://hpo.jax.org/app/browse/term/HP:0011999)) | ENSP00000331681 | - |
|  | DGCR2 | - |
|  | TBX1 | - |
|  | DGCR14 | - |
|  | CPOX | - |
|  | DGCR8 | - |
|  | HMBS | - |
|  | TIMM8A | - |
|  | VPS13A | - |
|  | IMPA1 | LITHIUM CARBONATE, LITHIUM CITRATE |
| Personality disorders ([HP:0012075](https://hpo.jax.org/app/browse/term/HP:0012075)) | - | - |
| Phonophobia ([HP:0002183](https://hpo.jax.org/app/browse/term/HP:0002183)) | SGCE | - |
|  | SCN1A | ETHOSUXIMIDE |
|  | KCNT1 | CLOFILIUM, LOXAPINE, QUINIDINE |
|  | KCTD17 | - |
|  | TOR1A | HALOPERIDOL |
|  | ENSP00000381231 | - |
|  | CYP27A1 | CHOLESTYRAMINE |
|  | CUX2 | RIFAMPIN |
|  | XK | - |
|  | SPG21 | - |
| Polydipsia ([HP:0001959](https://hpo.jax.org/app/browse/term/HP:0001959)) | EP300 | - |
|  | CREBBP | - |
|  | ESR1 | CLOMIPHENE, LEVONORGESTREL |
|  | MLXIPL | - |
|  | ELN | - |
|  | NCF1 | - |
|  | DNAJC30 | - |
| Short attention span ([HP:0000736](https://hpo.jax.org/app/browse/term/HP:0000736)) | SCN1A | ETHOSUXIMIDE |
|  | SCN2A | - |
|  | GABRG2 | - |
|  | KCNT1 | CLOFILIUM, LOXAPINE, QUINIDINE |
|  | GABRA1 | METHOHEXITAL, BARBITAL, THIAMYLAL, ZALEPLON, ZOPICLONE, ZOLPIDEM |
|  | SCN8A | - |
|  | KCNA2 | - |
|  | PCDH19 | - |
|  | SLC6A1 | TIAGABINE, TIAGABINE HYDROCHLORIDE |
|  | GRIN2A | MEMANTINE, GLYCINE, DEXTROMETHORPHAN POLISTIREX |
| Visual hallucinations ([HP:0002367](https://hpo.jax.org/app/browse/term/HP:0002367)) | DNAJC5 | - |
|  | FBXO7 | - |
|  | GIGYF2 | - |
|  | SNCB | - |
|  | NPRL3 | - |
|  | FIG4 | - |
|  | LGI1 | - |
|  | RELN | - |
|  | EPM2A | - |

### Table 18 Neuropsychiatric-cognitivedysfunction

| **Symptom (HPO ID)** | **Hub Genes (ranked)** | **FDA-Approved repurposed drugs** |
| --- | --- | --- |
| Abnormality of higher mental function ([HP:0011446](https://hpo.jax.org/app/browse/term/HP:0011446)) | RPGR | - |
|  | IMPDH1 | MYCOPHENOLATE MOFETIL HYDROCHLORIDE, RIBAVIRIN, MYCOPHENOLATE MOFETIL, MYCOPHENOLIC ACID |
|  | RDH12 | - |
|  | CRX | - |
|  | SPATA7 | - |
|  | PRPH2 | - |
|  | TULP1 | - |
|  | CNGA1 | DEQUALINIUM |
|  | NR2E3 | - |
|  | GUCA1B | - |
| Agnosia ([HP:0010524](https://hpo.jax.org/app/browse/term/HP:0010524)) | TREM2 | - |
|  | MAPT | - |
|  | PIK3CA | - |
|  | PMS2 | NIVOLUMAB |
|  | ABCA7 | - |
|  | KRAS | PANITUMUMAB, CETUXIMAB |
|  | C9orf72 | - |
|  | FAN1 | - |
|  | ATP6AP2 | - |
| Bradykinesia ([HP:0002067](https://hpo.jax.org/app/browse/term/HP:0002067)) | SCN1A | ETHOSUXIMIDE |
|  | GABRG2 | - |
|  | SCN2A | - |
|  | GABRA1 | METHOHEXITAL, BARBITAL, THIAMYLAL, ZALEPLON, ZOPICLONE, ZOLPIDEM |
|  | SCN1B | ZONISAMIDE |
|  | GABRD | - |
|  | ATP1A3 | ACETYLDIGITOXIN, DESLANOSIDE, DIGITOXIN |
|  | PCDH19 | - |
|  | SCN9A | - |
|  | ATXN3 | - |
| Bradyphrenia ([HP:0031843](https://hpo.jax.org/app/browse/term/HP:0031843)) | - | - |
| Cognitive impairment ([HP:0100543](https://hpo.jax.org/app/browse/term/HP:0100543)) | SCN2A | - |
|  | SCN1A | ETHOSUXIMIDE |
|  | KCNA2 | - |
|  | KCNT1 | CLOFILIUM, LOXAPINE, QUINIDINE |
|  | GABRG2 | - |
|  | KCNC1 | - |
|  | GABRA1 | METHOHEXITAL, BARBITAL, THIAMYLAL, ZALEPLON, ZOPICLONE, ZOLPIDEM |
|  | CACNA1B | ZICONOTIDE, ZICONOTIDE ACETATE, AMLODIPINE, LEVETIRACETAM |
|  | SCN8A | - |
|  | GABRD | - |
| Confusion ([HP:0001289](https://hpo.jax.org/app/browse/term/HP:0001289)) | ERCC4 | - |
|  | ERCC2 | OXALIPLATIN, LEUCOVORIN |
|  | NAGS | - |
|  | SCN1A | ETHOSUXIMIDE |
|  | SLC25A15 | - |
|  | ERCC5 | - |
|  | MEN1 | OLSALAZINE |
|  | MMACHC | - |
|  | CHRNA2 | PIPECURONIUM, METOCURINE, CISATRACURIUM, PANCURONIUM, MECAMYLAMINE, DOXACURIUM, MIVACURIUM, DECAMETHONIUM, VECURONIUM, TUBOCURARINE |
|  | TREM2 | - |
| Encephalopathy ([HP:0001298](https://hpo.jax.org/app/browse/term/HP:0001298)) | GABRA1 | METHOHEXITAL, BARBITAL, THIAMYLAL, ZALEPLON, ZOPICLONE, ZOLPIDEM |
|  | GABRG2 | - |
|  | SCN2A | - |
|  | SLC6A1 | TIAGABINE, TIAGABINE HYDROCHLORIDE |
|  | GRIN2A | MEMANTINE, GLYCINE, DEXTROMETHORPHAN POLISTIREX |
|  | SCN1A | ETHOSUXIMIDE |
|  | GRIN2B | MEMANTINE, FELBAMATE, ACAMPROSATE |
|  | KCNA2 | - |
|  | NRXN1 | DULOXETINE |
|  | GABRB2 | - |
| Diminished ability to concentrate ([HP:0031987](https://hpo.jax.org/app/browse/term/HP:0031987)) | - | - |

### Table 19 Neuropsychiatric-emotion-mood

| **Symptom (HPO ID)** | **Hub Genes (ranked)** | **FDA-Approved repurposed drugs** |
| --- | --- | --- |
| Abnormal emotion/affect behavior ([HP:0100851](https://hpo.jax.org/app/browse/term/HP:0100851)) | GRIA2 | PERAMPANEL |
|  | SLC6A1 | TIAGABINE, TIAGABINE HYDROCHLORIDE |
|  | GRIN2A | MEMANTINE, GLYCINE, DEXTROMETHORPHAN POLISTIREX |
|  | GRIN1 | MEMANTINE, ORPHENADRINE |
|  | GRIA3 | PERAMPANEL |
|  | CAMK2A | - |
|  | GABRG2 | - |
|  | GABRA1 | METHOHEXITAL, BARBITAL, THIAMYLAL, ZALEPLON, ZOPICLONE, ZOLPIDEM |
|  | GRIA4 | PERAMPANEL, PIRACETAM |
|  | GRIN2B | MEMANTINE, FELBAMATE, ACAMPROSATE |
| Abnormal fear/anxiety-related behavior ([HP:0100852](https://hpo.jax.org/app/browse/term/HP:0100852)) | GPR98 | - |
|  | USH1C | - |
|  | CLRN1 | - |
|  | PDZD7 | - |
|  | USH2A | - |
|  | USH1G | - |
|  | PCDH15 | - |
|  | DFNB31 | - |
|  | CIB2 | - |
|  | CDH23 | METHYLPHENIDATE |
| Aggressive behavior ([HP:0000718](https://hpo.jax.org/app/browse/term/HP:0000718)) | EP300 | - |
|  | CREBBP | - |
|  | ARID1A | ATEZOLIZUMAB, NIVOLUMAB, PEMBROLIZUMAB |
|  | DNMT3A | DECITABINE, IDARUBICIN, WARFARIN |
|  | KDM5C | EVEROLIMUS, SUNTINIB |
|  | SIN3A | - |
|  | SETD2 | - |
|  | SMARCC2 | - |
|  | BPTF | - |
|  | ARID1B | - |
| Depression ([HP:0000716](https://hpo.jax.org/app/browse/term/HP:0000716)) | GPR98 | - |
|  | USH1G | - |
|  | PCDH15 | - |
|  | DFNB31 | - |
|  | CIB2 | - |
|  | CDH23 | METHYLPHENIDATE |
|  | USH1C | - |
|  | CLRN1 | - |
|  | PDZD7 | - |
|  | USH2A | - |
| Emotional lability ([HP:0000712](https://hpo.jax.org/app/browse/term/HP:0000712)) | NDUFS3 | - |
|  | NDUFS1 | - |
|  | NDUFA2 | - |
|  | NDUFA9 | - |
|  | NDUFAF2 | - |
|  | NDUFAF6 | - |
|  | COX15 | - |
|  | TACO1 | - |
|  | PDHA1 | - |
|  | C9orf72 | - |
| Euphoria ([HP:0031844](https://hpo.jax.org/app/browse/term/HP:0031844)) | - | - |
| Mania ([HP:0100754](https://hpo.jax.org/app/browse/term/HP:0100754)) | PRKACA | - |
|  | TBX1 | - |
|  | PRKAR1A | - |
|  | PDE11A | TADALAFIL |
|  | C10orf2 | - |
|  | GP1BB | - |
|  | POLG2 | - |
|  | COMT | ENTACAPONE, OPICAPONE, REMIFENTANIL, SUFENTANIL, MODAFINIL, MORPHINE, METHADONE |
|  | RPS6KA3 | - |
|  | SLC25A13 | - |
| Suicidal ideation ([HP:0031589](https://hpo.jax.org/app/browse/term/HP:0031589)) | KCNT1 | CLOFILIUM, LOXAPINE, QUINIDINE |
|  | CHRNB2 | NICOTINE, VARENICLINE, VARENICLINE TARTRATE, NICOTINE POLACRILEX |
|  | CHRNA4 | VARENICLINE |
|  | CHRNA2 | PIPECURONIUM, METOCURINE, CISATRACURIUM, PANCURONIUM, MECAMYLAMINE, DOXACURIUM, MIVACURIUM, DECAMETHONIUM, VECURONIUM, TUBOCURARINE |
|  | CYP27A1 | CHOLESTYRAMINE |
|  | FIG4 | - |
|  | CDH23 | METHYLPHENIDATE |

### Table 20 Neuropsychiatric-headache

| **Symptom (HPO ID)** | **Hub Genes (ranked)** | **FDA-Approved repurposed drugs** |
| --- | --- | --- |
| Headache ([HP:0002315](https://hpo.jax.org/app/browse/term/HP:0002315)) | CTNNB1 | FLUORESCEN SODIUM |
|  | KRAS | PANITUMUMAB, CETUXIMAB |
|  | ESR1 | CLOMIPHENE, LEVONORGESTREL |
|  | KIT | NILOTINIB, IMATINIB |
|  | RET | CABOZANTINIB, VANDETANIB |
|  | SMAD4 | LYSINE, ALECTINIB |
|  | TERT | OMACETAXINE MEPESUCCINATE |
|  | PIK3CA | - |
|  | NF1 | TRAMETINIB, BINIMETINIB, COBIMETINIB, DABRAFENIB, VEMURAFENIB |
|  | CDKN1B | RALTITREXED |
| Migraine ([HP:0002076](https://hpo.jax.org/app/browse/term/HP:0002076)) | KRAS | PANITUMUMAB, CETUXIMAB |
|  | PIK3CA | - |
|  | ESR1 | CLOMIPHENE, LEVONORGESTREL |
|  | SMAD4 | LYSINE, ALECTINIB |
|  | NF1 | TRAMETINIB, BINIMETINIB, COBIMETINIB, DABRAFENIB, VEMURAFENIB |
|  | NOTCH3 | - |
|  | RELA | - |
|  | SCN1A | ETHOSUXIMIDE |
|  | GRIN2A | MEMANTINE, GLYCINE, DEXTROMETHORPHAN POLISTIREX |
|  | KCNA1 | BUPIVACAINE, ISOFLURANE, DESFLURANE |

### Table 21 Neuropsychiatric-memory

| **Symptom (HPO ID)** | **Hub Genes (ranked)** | **FDA-Approved repurposed drugs** |
| --- | --- | --- |
| Memory impairment ([HP:0002354](https://hpo.jax.org/app/browse/term/HP:0002354)) | MAPT | - |
|  | NF1 | TRAMETINIB, BINIMETINIB, COBIMETINIB, DABRAFENIB, VEMURAFENIB |
|  | KRAS | PANITUMUMAB, CETUXIMAB |
|  | ATXN3 | - |
|  | PIK3CA | - |
|  | PRKACA | - |
|  | ATXN1 | - |
|  | C9orf72 | - |
|  | PRKAR1A | - |
|  | PRKCG | - |

### Table 22 Neuropsychiatric-sleep

| **Symptom (HPO ID)** | **Hub Genes (ranked)** | **FDA-Approved repurposed drugs** |
| --- | --- | --- |
| Insomnia ([HP:0100785](https://hpo.jax.org/app/browse/term/HP:0100785)) | NR1H4 | OBETICHOLIC ACID, CHENODIOL |
|  | ATP8B1 | - |
|  | ABCB4 | BENZQUINAMIDE |
|  | ABCB11 | CHOLIC ACID, CHENODIOL, HYDROXYPROGESTERONE CAPROATE |
|  | NAGS | - |
|  | CPOX | - |
|  | CLCNKB | - |
|  | MLXIPL | - |
|  | SLC25A13 | - |
|  | SLC12A3 | POLYTHIAZIDE, BENDROFLUMETHIAZIDE, QUINETHAZONE, METOLAZONE, BENZTHIAZIDE, TRICHLORMETHIAZIDE, CHLOROTHIZIDE, CHLOROTHIAZIDE SODIUM, INDAPAMIDE, HYDROCHL OROTHIAZIDE |
| Maintenance insomnia ([HP:0031355](https://hpo.jax.org/app/browse/term/HP:0031355)) | - | - |
| Restless legs ([HP:0012452](https://hpo.jax.org/app/browse/term/HP:0012452)) | NEFL | - |
|  | UCHL1 | - |
|  | MFN2 | - |
|  | DNAJC6 | - |
|  | ATXN7 | - |
| Sleep apnea ([HP:0010535](https://hpo.jax.org/app/browse/term/HP:0010535)) | CTNNB1 | FLUORESCEN SODIUM |
|  | HRAS | - |
|  | EP300 | - |
|  | CREBBP | - |
|  | TERT | OMACETAXINE MEPESUCCINATE |
|  | RUNX2 | - |
|  | TWIST1 | - |
|  | RET | CABOZANTINIB, VANDETANIB |
|  | USP7 | - |
|  | CHAT | RIVASTIGMINE, GALANTAMINE, DONEPEZILM PILOCARPINE |
| Sleep disturbance ([HP:0002360](https://hpo.jax.org/app/browse/term/HP:0002360)) | GRIN2B | MEMANTINE, FELBAMATE, ACAMPROSATE |
|  | GRIN2A | MEMANTINE, GLYCINE, DEXTROMETHORPHAN POLISTIREX |
|  | GRIN1 | MEMANTINE, ORPHENADRINE |
|  | GRIA4 | PERAMPANEL, PIRACETAM |
|  | CAMK2A | - |
|  | GRIA3 | METHOHEXITAL, BARBITAL, THIAMYLAL, ZALEPLON, ZOPICLONE, ZOLPIDEM |
|  | GABRG2 | - |
|  | KCNA1 | BUPIVACAINE, ISOFLURANE, DESFLURANE |
|  | SCN2A | - |
|  | CAMK2B | - |
| Sleep onset insomnia ([HP:0031354](https://hpo.jax.org/app/browse/term/HP:0031354)) | - | - |

### Table 23 Neuropsychiatric-smell-taste

| **Symptom (HPO ID)** | **Hub Genes (ranked)** | **FDA-Approved repurposed drugs** |
| --- | --- | --- |
| Abnormality of taste sensation ([HP:0000223](https://hpo.jax.org/app/browse/term/HP:0000223)) | - | - |
| Anosmia ([HP:0000458](https://hpo.jax.org/app/browse/term/HP:0000458)) | FGF8 | - |
|  | KAL1 | - |
|  | GNRH1 | ZALCITABINE, GOSERELIN, AMINOGLUTETHIMIDE, DITIOCARB, DAPSONE, CAPTOPRIL |
|  | TAC3 | LIOTHYRONINE SODIUM |
|  | FEZF1 | - |
|  | CHD7 | - |
|  | PROKR2 | - |
|  | KISS1R | - |
|  | IL17RD | - |
|  | KISS1 | - |
| Hypogeusia ([HP:0000224](https://hpo.jax.org/app/browse/term/HP:0000224)) | - | - |
| Hyposmia ([HP:0004409](https://hpo.jax.org/app/browse/term/HP:0004409)) | KAL1 | - |
|  | PROKR2 | - |
|  | FEZF1 | - |
|  | SOX10 | VEMURAFENIB |
|  | FGF8 | - |
|  | CHD7 | - |
|  | FGF17 | - |
|  | IL17RD | - |
|  | GIGYF2 | - |
|  | LZTFL1 | - |
| Abnormality of the sense of smell ([HP:0004408](https://hpo.jax.org/app/browse/term/HP:0004408)) | FGF8 | - |
|  | KAL1 | - |
|  | GNRH1 | ZALCITABINE, GOSERELIN, AMINOGLUTETHIMIDE, DITIOCARB, DAPSONE, CAPTOPRIL |
|  | TAC3 | LIOTHYRONINE SODIUM |
|  | FEZF1 | - |
|  | CHD7 | - |
|  | PROKR2 | - |
|  | KISS1R | - |
|  | IL17RD | - |
|  | KISS1 | - |

### Table 24 Neuropsychiatric-speech-language

| **Symptom (HPO ID)** | **Hub Genes (ranked)** | **FDA-Approved repurposed drugs** |
| --- | --- | --- |
| Anomic aphasia ([HP:0030784](https://hpo.jax.org/app/browse/term/HP:0030784)) | MAPT | - |
|  | TREM2 | - |
|  | C9orf72 | - |
|  | PLP1 | - |
| Aphasia ([HP:0002381](https://hpo.jax.org/app/browse/term/HP:0002381)) | MAPT | - |
|  | BPTF | - |
|  | GRIN2A | MEMANTINE, GLYCINE, DEXTROMETHORPHAN POLISTIREX |
|  | TRIP12 | - |
|  | SCN1A | ETHOSUXIMIDE |
|  | PLP1 | - |
|  | C9orf72 | - |
|  | POLR1D | - |
|  | POLR1C | - |
|  | KRAS | PANITUMUMAB, CETUXIMAB |
| Expressive aphasia ([HP:0002427](https://hpo.jax.org/app/browse/term/HP:0002427)) | C9orf72 | - |
|  | MAPT | - |
|  | TREM2 | - |
|  | NPRL3 | - |
|  | GJB1 | CARBENOXOLONE |
| Neurological speech impairment ([HP:0002167](https://hpo.jax.org/app/browse/term/HP:0002167)) | DRC1 | - |
|  | CCDC65 | - |
|  | HYDIN | - |
|  | RSPH4A | - |
|  | RSPH1 | - |
|  | CCDC103 | - |
|  | HEATR2 | - |
|  | DYX1C1 | - |
|  | DNAI2 | - |
|  | CCDC40 | - |
| Slurred speech ([HP:0001350](https://hpo.jax.org/app/browse/term/HP:0001350)) | MAPT | - |
|  | KCNA1 | BUPIVACAINE, ISOFLURANE, DESFLURANE |
|  | MOG | - |
|  | GRIN2A | MEMANTINE, GLYCINE, DEXTROMETHORPHAN POLISTIREX |
|  | DNAJC6 | - |
|  | KCND3 | - |
|  | ATXN1 | - |
|  | SETD2 | - |
|  | ACOX2 | - |
|  | HCRT | - |

### Table 25 Neuropsychiatric-finding

| **Symptom (HPO ID)** | **Hub Genes (ranked)** | **FDA-Approved repurposed drugs** |
| --- | --- | --- |
| Abnormal reflex ([HP:0031826](https://hpo.jax.org/app/browse/term/HP:0031826)) | RDH12 | - |
|  | NR2E3 | - |
|  | CRX | - |
|  | GUCA1B | - |
|  | PRPH2 | - |
|  | RPGR | - |
|  | TULP1 | - |
|  | PRPF3 | - |
|  | PDE6G | PENTOXIFYLLINE |
|  | CNGA1 | DEQUALINIUM |
| Abnormality of movement ([HP:0100022](https://hpo.jax.org/app/browse/term/HP:0100022)) | RDH12 | - |
|  | NR2E3 | - |
|  | CRX | - |
|  | GUCA1B | - |
|  | PRPH2 | - |
|  | RPGR | - |
|  | TULP1 | - |
|  | CNGA1 | DEQUALINIUM |
|  | IMPDH1 | MYCOPHENOLATE MOFETIL HYDROCHLORIDE, RIBAVIRIN, MYCOPHENOLATE MOFETIL, MYCOPHENOLIC ACID |
|  | NRL | - |
| Ataxia ([HP:0001251](https://hpo.jax.org/app/browse/term/HP:0001251)) | B9D1 | - |
|  | CC2D2A | - |
|  | MKS1 | - |
|  | TMEM216 | - |
|  | RPGRIP1L | - |
|  | TCTN2 | - |
|  | NPHP1 | - |
|  | TMEM231 | - |
|  | TCTN3 | - |
|  | TCTN1 | - |
| Babinski sign ([HP:0003487](https://hpo.jax.org/app/browse/term/HP:0003487)) | REEP1 | - |
|  | RTN2 | - |
|  | SPG21 | - |
|  | SPG11 | - |
|  | KIAA0196 | - |
|  | NIPA1 | - |
|  | AP4B1 | - |
|  | AP4S1 | - |
|  | AP4E1 | - |
|  | SPAST | - |
| Dysarthria ([HP:0001260](https://hpo.jax.org/app/browse/term/HP:0001260)) | ATXN2 | - |
|  | SPG11 | - |
|  | C9orf72 | - |
|  | FUS | TRETINOIN |
|  | MATR3 | - |
|  | NEFH | - |
|  | SETX | - |
|  | NEK1 | - |
|  | FIG4 | - |
|  | ALS2 | - |
| Dysmetria ([HP:0001310](https://hpo.jax.org/app/browse/term/HP:0001310)) | ATXN7 | - |
|  | ATXN10 | - |
|  | PPP2R2B | - |
|  | ATXN2 | - |
|  | KCNC3 | - |
|  | ATXN1 | - |
|  | ATN1 | - |
|  | TGM6 | - |
|  | PRKCG | - |
|  | FXN | EPOETIN BETA |
| Dysphagia ([HP:0002015](https://hpo.jax.org/app/browse/term/HP:0002015)) | ATXN2 | - |
|  | C9orf72 | - |
|  | NEFH | - |
|  | FUS | TRETINOIN |
|  | SPG11 | - |
|  | ALS2 | - |
|  | NEK1 | - |
|  | MATR3 | - |
|  | SETX | - |
|  | FIG4 | - |
| Dystonia ([HP:0001332](https://hpo.jax.org/app/browse/term/HP:0001332)) | GRIN2B | MEMANTINE, FELBAMATE, ACAMPROSATE |
|  | GRIA2 | PERAMPANEL |
|  | SCN2A | - |
|  | CACNA1B | ZICONOTIDE, ZICONOTIDE ACETATE, AMLODIPINE, LEVETIRACETAM |
|  | KCNA1 | BUPIVACAINE, ISOFLURANE, DESFLURANE |
|  | GRIN1 | MEMANTINE, ORPHENADRINE |
|  | GRIN2A | MEMANTINE, GLYCINE, DEXTROMETHORPHAN POLISTIREX |
|  | CAMK2A | - |
|  | GABRB2 | - |
|  | GRIK2 | BUTABARBITAL, BUTETHAL, THIOPENTAL, MEPHOBARBITAL, METHARBITAL, TALBUTAL, BUTALBITAL |
| Facial paralysis ([HP:0007209](https://hpo.jax.org/app/browse/term/HP:0007209)) | ATP1A2 | ACETYLDIGITOXIN, DESLANOSIDE |
|  | TCIRG1 | - |
|  | SCN1A | ETHOSUXIMIDE |
|  | TNFSF11 | DENOSUMAB, LENALIDOMIDE, ANASTROZOLE, LETROZOLE |
|  | SH3TC2 | - |
| Frontal release signs ([HP:0000743](https://hpo.jax.org/app/browse/term/HP:0000743)) | - | - |
| Gait disturbance ([HP:0001288](https://hpo.jax.org/app/browse/term/HP:0001288)) | GRIA2 | PERAMPANEL |
|  | GABRA1 | METHOHEXITAL, BARBITAL, THIAMYLAL, ZALEPLON, ZOPICLONE, ZOLPIDEM |
|  | GABRG2 | - |
|  | GABRB2 | - |
|  | GRIN1 | MEMANTINE, ORPHENADRINE |
|  | CAMK2A | - |
|  | KCNA1 | BUPIVACAINE, ISOFLURANE, DESFLURANE |
|  | GABRD | - |
|  | SLC12A5 | BUMETANIDE |
|  | SLC6A1 | TIAGABINE, TIAGABINE HYDROCHLORIDE |
| Hand muscle weakness ([HP:0030237](https://hpo.jax.org/app/browse/term/HP:0030237)) | RAPSN | - |
|  | CHRNA1 | - |
|  | CHRNE | SUCCINYLCHOLINE CHLORIDE, PIPECURONIUM BROMIDE, MIVACURIUM CHLORIDE, METOCURINE IODIDE, DOXACURIUM CHLORIDE, VECURONIUM BROMIDE, DECAMETHONIUM BROMIDE |
|  | CHRNB1 | SUCCINYLCHOLINE CHLORIDE, PIPECURONIUM BROMIDE, MIVACURIUM CHLORIDE, METOCURINE IODIDE, DOXACURIUM CHLORIDE, VECURONIUM BROMIDE, DECAMETHONIUM BROMIDE |
|  | CHRND | SUCCINYLCHOLINE CHLORIDE, PIPECURONIUM BROMIDE, MIVACURIUM CHLORIDE, METOCURINE IODIDE, DOXACURIUM CHLORIDE, VECURONIUM BROMIDE, DECAMETHONIUM BROMIDE |
|  | DOK7 | - |
|  | MUSK | - |
|  | SCN4A | LEVOBUPIVACAINE HYDROCHLORIDE, BUPIVACAINE HYDROCHLORIDE |
|  | COLQ | - |
|  | MFN2 | - |
| Hyperesthesia ([HP:0100963](https://hpo.jax.org/app/browse/term/HP:0100963)) | - | - |
| Hyperkinetic movements ([HP:0002487](https://hpo.jax.org/app/browse/term/HP:0002487)) | GRIN2A | MEMANTINE, GLYCINE, DEXTROMETHORPHAN POLISTIREX |
|  | GAD1 | METHADONE |
|  | CACNA1B | ZICONOTIDE, ZICONOTIDE ACETATE, AMLODIPINE, LEVETIRACETAM |
|  | GRIN1 | MEMANTINE, ORPHENADRINE |
|  | SCN1A | ETHOSUXIMIDE |
|  | GNAO1 | - |
|  | SLC6A3 | DEXMETHYLPHENIDATE, PHENMETRAZINE, MAZINDOL, DIETHYLPROPION, ARMODAFNIL, DEXTROAMPHETAMINE, PHENTERMINE, BUPROPION, MODAFINIL, LISDEXAMFETAMINE |
|  | IQSEC2 | - |
|  | ALDH5A1 | CHLORMERODRIN, VALPROATE SODIUM, DIVALPROEX SODIUM, VALPROIC ACID |
|  | PNKD | - |
| Hypotonia ([HP:0001252](https://hpo.jax.org/app/browse/term/HP:0001252)) | NDUFS3 | - |
|  | NDUFA9 | - |
|  | NDUFA2 | - |
|  | NDUFC2 | - |
|  | NDUFA6 | - |
|  | NDUFA12 | - |
|  | NDUFA1 | - |
|  | NDUFA8 | - |
|  | NDUFB10 | - |
|  | NDUFB11 | - |
| Muscle spasm ([HP:0003394](https://hpo.jax.org/app/browse/term/HP:0003394)) | C9orf72 | - |
|  | FUS | TRETINOIN |
|  | ATXN2 | - |
|  | MATR3 | - |
|  | NEK1 | - |
|  | FIG4 | - |
|  | NEFH | - |
|  | TAF15 | - |
|  | CCNF | - |
|  | GLE1 | - |
| Muscle weakness ([HP:0001324](https://hpo.jax.org/app/browse/term/HP:0001324)) | FKTN | - |
|  | FKRP | - |
|  | POMGNT1 | - |
|  | SCN4A | LEVOBUPIVACAINE HYDROCHLORIDE, BUPIVACAINE HYDROCHLORIDE |
|  | SCN1A | ETHOSUXIMIDE |
|  | SCN2A | - |
|  | SCN8A | - |
|  | MYOT | - |
|  | SCN11A | - |
|  | SGCB | - |
| Orthostatic hypotension ([HP:0001278](https://hpo.jax.org/app/browse/term/HP:0001278)) | GIGYF2 | - |
|  | CHCHD2 | - |
|  | SPG11 | - |
|  | IL12RB1 | - |
|  | LEP | SOYBEAN OIL, GEMFIBROZIL |
|  | CYB561 | - |
|  | AAAS | - |
|  | IKBKAP | - |
|  | ARSA | DECAMETHONIUM |
|  | CHRNA3 | VARENICLINE, PENTOLINIUM TARTRATE, TREMETHAPHAN CAMYSLATE, MECAMYLAMINE HYDROCHLORIDE |
| Paresthesia ([HP:0003401](https://hpo.jax.org/app/browse/term/HP:0003401)) | CASR | CINACALCET, VELCALCETIDE, CINACALCET HYDROCHLORIDE, PAMIDRONIC ACID |
|  | CLCNKB | - |
|  | KCNJ1 | GLYMIDINE, ACETOHEXAMIDE, MINOXIDIL, GLIMEPIRIDE, TOLBUTAMIDE, GLICLAZIDE, TOLAZAMIDE, GLIPIZIDE, CHLORPROPAMIDE |
|  | PIK3CA | - |
|  | KRAS | PANITUMUMAB, CETUXIMAB |
|  | SLC12A3 | POLYTHIAZIDE, BENDROFLUMETHIAZIDE, QUINETHAZONE, METOLAZONE, BENZTHIAZIDE, TRICHLORMETHIAZIDE, CHLOROTHIZIDE, CHLOROTHIAZIDE SODIUM, INDAPAMIDE, HYDROCHL OROTHIAZIDE |
|  | THPO | - |
|  | SLC12A1 | HYDROFLUMETHIAZIDE, METHYCLOTHIAZIDE, CHLORMERODRIN, FUROSEMIDE, TRICHLORMETHIAZIDE, BUMETANIDE, CHLORTHALIDONE, TORSEMIDE, METOLAZONE, BENDROFLUMETHIAZIDE |
|  | PTPN22 | - |
|  | GATA1 | CYTARABINE, DAUNORUBICIN |
| Parkinsonism ([HP:0001300](https://hpo.jax.org/app/browse/term/HP:0001300)) | GABRG2 | - |
|  | SCN2A | - |
|  | GABRA1 | METHOHEXITAL, BARBITAL, THIAMYLAL, ZALEPLON, ZOPICLONE, ZOLPIDEM |
|  | SCN1A | ETHOSUXIMIDE |
|  | PCDH19 | - |
|  | ATP1A3 | ACETYLDIGITOXIN, DESLANOSIDE, DIGITOXIN |
|  | SCN1B | ZONISAMIDE |
|  | KIF5A | - |
|  | C9orf72 | - |
|  | ATXN2 | - |
| Polyneuropathy ([HP:0001271](https://hpo.jax.org/app/browse/term/HP:0001271)) | PEX11B | - |
|  | PEX7 | - |
|  | PEX12 | - |
|  | SETX | - |
|  | GRM1 | MORPHINE, CAFFEINE |
|  | MYOT | - |
|  | COQ7 | - |
|  | PDYN | MORPHINE |
|  | ERCC6 | DEXTROMETHORPHAN, CYCLOSPORINE |
|  | LDB3 | - |
| Rigidity ([HP:0002063](https://hpo.jax.org/app/browse/term/HP:0002063)) | SCN1A | ETHOSUXIMIDE |
|  | SCN2A | - |
|  | KCNA2 | - |
|  | SCN8A | - |
|  | SCN9A | - |
|  | KCND3 | - |
|  | KCNA4 | - |
|  | SCN1B | ZONISAMIDE |
|  | KCNB1 | - |
|  | CACNA1B | ZICONOTIDE, ZICONOTIDE ACETATE, AMLODIPINE, LEVETIRACETAM |
| Seizure ([HP:0001250](https://hpo.jax.org/app/browse/term/HP:0001250)) | CC2D2A | - |
|  | B9D1 | - |
|  | TCTN2 | - |
|  | TMEM231 | - |
|  | TMEM216 | - |
|  | B9D2 | - |
|  | MKS1 | - |
|  | RPGRIP1L | - |
|  | TCTN1 | - |
|  | TMEM138 | - |
| Skeletal muscle atrophy ([HP:0003202](https://hpo.jax.org/app/browse/term/HP:0003202)) | BBS5 | - |
|  | BBS7 | - |
|  | BBS1 | - |
|  | LZTFL1 | - |
|  | BBS2 | - |
|  | BBS10 | - |
|  | WDPCP | - |
|  | BBS12 | - |
|  | MKS1 | - |
|  | SDCCAG8 | - |
| Somatic sensory dysfunction ([HP:0003474](https://hpo.jax.org/app/browse/term/HP:0003474)) | BSCL2 | - |
|  | REEP1 | - |
|  | KIF5A | - |
|  | SPG11 | - |
|  | NIPA1 | - |
|  | KIAA0196 | - |
|  | RTN2 | - |
|  | SPAST | - |
|  | ATL1 | - |
|  | MPZ | - |
| Spasticity ([HP:0001257](https://hpo.jax.org/app/browse/term/HP:0001257)) | SPG11 | - |
|  | REEP1 | - |
|  | RTN2 | - |
|  | SPG21 | - |
|  | KIAA0196 | - |
|  | NIPA1 | - |
|  | AP4E1 | - |
|  | AP4B1 | - |
|  | AP4S1 | - |
|  | KIF5A | - |
| Tremor ([HP:0001337](https://hpo.jax.org/app/browse/term/HP:0001337)) | B9D1 | - |
|  | CC2D2A | - |
|  | TCTN2 | - |
|  | CEP41 | - |
|  | TCTN3 | - |
|  | MKS1 | - |
|  | TMEM216 | - |
|  | RPGRIP1L | - |
|  | TCTN1 | - |
|  | NPHP1 | - |
| Unilateral facial palsy ([HP:0012799](https://hpo.jax.org/app/browse/term/HP:0012799)) | - | - |
