## Supplementary Table 1-3 for "Long COVID: G Protein-Coupled Receptors (GPCRs) responsible for persistent post-COVID symptoms": Supplementary File 2.1-Immunology-autoimmunity.docx

Anaphylactic shock ([HP:0100845](https://hpo.jax.org/app/browse/term/HP:0100845))

KEGG Pathway

| **Index** | **Name** | **P-value** | **Adjusted p-value** | **Odds Ratio** | **Combined score** |
| --- | --- | --- | --- | --- | --- |
| 1 | Chronic myeloid leukemia | 5.105e-13 | 3.140e-11 | 426.86 | 12081.50 |
| 2 | Phospholipase D signaling pathway | 1.240e-13 | 1.525e-11 | 328.47 | 9761.61 |
| 3 | Acute myeloid leukemia | 9.011e-11 | 1.232e-9 | 321.42 | 7434.41 |
| 4 | Renal cell carcinoma | 1.048e-10 | 1.289e-9 | 311.34 | 7154.30 |
| 5 | Proteoglycans in cancer | 1.254e-12 | 5.141e-11 | 233.24 | 6391.87 |

GO: Biological Process

| **Index** | **Name** | **P-value** | **Adjusted p-value** | **Odds Ratio** | **Combined score** |
| --- | --- | --- | --- | --- | --- |
| 1 | entry of bacterium into host cell (GO:0035635) | 0.000006290 | 0.0001510 | 832.67 | 9972.50 |
| 2 | positive regulation of intracellular signal transduction (GO:1902533) | 7.703e-14 | 2.403e-11 | 326.03 | 9844.29 |
| 3 | positive regulation of phosphatidylinositol 3-kinase signaling (GO:0014068) | 1.840e-10 | 2.870e-8 | 276.64 | 6201.20 |
| 4 | epidermal growth factor receptor signaling pathway (GO:0007173) | 5.561e-9 | 3.470e-7 | 309.26 | 5878.15 |
| 5 | embryonic hemopoiesis (GO:0035162) | 0.00001750 | 0.0003211 | 454.07 | 4973.59 |

GO: Molecular Function

| **Index** | **Name** | **P-value** | **Adjusted p-value** | **Odds Ratio** | **Combined score** |
| --- | --- | --- | --- | --- | --- |
| 1 | neurotrophin TRK receptor binding (GO:0005167) | 0.000003371 | 0.00003624 | 1249.13 | 15739.23 |
| 2 | neurotrophin TRKA receptor binding (GO:0005168) | 0.000003371 | 0.00003624 | 1249.13 | 15739.23 |
| 3 | phosphotyrosine residue binding (GO:0001784) | 2.064e-9 | 7.512e-8 | 403.17 | 8062.84 |
| 4 | protein phosphorylated amino acid binding (GO:0045309) | 3.494e-9 | 7.512e-8 | 350.04 | 6815.94 |
| 5 | ErbB-3 class receptor binding (GO:0043125) | 0.002498 | 0.007672 | 555.17 | 3326.77 |

GO: Cellular Component

| **Index** | **Name** | **P-value** | **Adjusted p-value** | **Odds Ratio** | **Combined score** |
| --- | --- | --- | --- | --- | --- |
| 1 | phosphatidylinositol 3-kinase complex, class I (GO:0097651) | 0.002498 | 0.03312 | 555.17 | 3326.77 |
| 2 | tertiary granule (GO:0070820) | 0.002880 | 0.03312 | 30.60 | 179.00 |
| 3 | tertiary granule lumen (GO:1904724) | 0.02717 | 0.09534 | 41.02 | 147.91 |
| 4 | filopodium (GO:0030175) | 0.02863 | 0.09534 | 38.86 | 138.07 |
| 5 | specific granule lumen (GO:0035580) | 0.03058 | 0.09534 | 36.30 | 126.60 |

Antinuclear antibody positivity ([HP:0003493](https://hpo.jax.org/app/browse/term/HP:0003493))

KEGG Pathway

| **Index** | **Name** | **P-value** | **Adjusted p-value** | **Odds Ratio** | **Combined score** |
| --- | --- | --- | --- | --- | --- |
| 1 | Autoimmune thyroid disease | 0.000002081 | 0.00006243 | 170.91 | 2236.02 |
| 2 | Riboflavin metabolism | 0.003994 | 0.008875 | 317.19 | 1751.86 |
| 3 | Measles | 4.542e-7 | 0.00002725 | 98.05 | 1431.99 |
| 4 | African trypanosomiasis | 0.0001485 | 0.0007952 | 142.54 | 1256.48 |
| 5 | Allograft rejection | 0.0001567 | 0.0007952 | 138.57 | 1214.06 |

GO: Biological Process

| **Index** | **Name** | **P-value** | **Adjusted p-value** | **Odds Ratio** | **Combined score** |
| --- | --- | --- | --- | --- | --- |
| 1 | T cell apoptotic process (GO:0070231) | 3.147e-9 | 0.000001026 | 2141.36 | 41921.23 |
| 2 | lymphocyte apoptotic process (GO:0070227) | 0.000003371 | 0.0002005 | 1249.13 | 15739.23 |
| 3 | negative regulation of B cell proliferation (GO:0030889) | 0.00001481 | 0.0004023 | 499.50 | 5554.53 |
| 4 | negative regulation of extrinsic apoptotic signaling pathway via death domain receptors (GO:1902042) | 5.342e-7 | 0.00008708 | 275.93 | 3985.12 |
| 5 | necroptotic process (GO:0070266) | 0.00002690 | 0.0006264 | 356.71 | 3753.85 |

GO: Molecular Function

| **Index** | **Name** | **P-value** | **Adjusted p-value** | **Odds Ratio** | **Combined score** |
| --- | --- | --- | --- | --- | --- |
| 1 | IgG binding (GO:0019864) | 0.002498 | 0.01298 | 555.17 | 3326.77 |
| 2 | acid phosphatase activity (GO:0003993) | 0.002997 | 0.01298 | 444.11 | 2580.41 |
| 3 | interleukin-12 receptor binding (GO:0005143) | 0.002997 | 0.01298 | 444.11 | 2580.41 |
| 4 | non-membrane spanning protein tyrosine phosphatase activity (GO:0004726) | 0.002997 | 0.01298 | 444.11 | 2580.41 |
| 5 | ferric iron binding (GO:0008199) | 0.003495 | 0.01298 | 370.07 | 2093.27 |

GO: Cellular Component

| **Index** | **Name** | **P-value** | **Adjusted p-value** | **Odds Ratio** | **Combined score** |
| --- | --- | --- | --- | --- | --- |
| 1 | death-inducing signaling complex (GO:0031264) | 0.000006290 | 0.00008806 | 832.67 | 9972.50 |
| 2 | CD95 death-inducing signaling complex (GO:0031265) | 0.002498 | 0.009961 | 555.17 | 3326.77 |
| 3 | integral component of plasma membrane (GO:0005887) | 0.00002378 | 0.0001664 | 19.21 | 204.50 |
| 4 | membrane raft (GO:0045121) | 0.002846 | 0.009961 | 30.79 | 180.49 |
| 5 | cytoplasmic side of plasma membrane (GO:0009898) | 0.02717 | 0.07298 | 41.02 | 147.91 |

Anti-thyroid peroxidase antibody positivity ([HP:0025379](https://hpo.jax.org/app/browse/term/HP:0025379))

KEGG Pathway

| **Index** | **Name** | **P-value** | **Adjusted p-value** | **Odds Ratio** | **Combined score** |
| --- | --- | --- | --- | --- | --- |
| 1 | PD-L1 expression and PD-1 checkpoint pathway in cancer | 1.354e-12 | 8.178e-11 | 359.77 | 9831.65 |
| 2 | T cell receptor signaling pathway | 3.526e-12 | 9.698e-11 | 304.47 | 8029.08 |
| 3 | Long-term potentiation | 9.011e-11 | 1.647e-9 | 321.42 | 7434.41 |
| 4 | Renal cell carcinoma | 1.048e-10 | 1.647e-9 | 311.34 | 7154.30 |
| 5 | Neurotrophin signaling pathway | 8.044e-12 | 1.770e-10 | 263.85 | 6740.44 |

GO: Biological Process

| **Index** | **Name** | **P-value** | **Adjusted p-value** | **Odds Ratio** | **Combined score** |
| --- | --- | --- | --- | --- | --- |
| 1 | negative regulation of neurotransmitter secretion (GO:0046929) | 0.000004719 | 0.0002463 | 999.25 | 12254.80 |
| 2 | regulation of phospholipase C activity (GO:1900274) | 4.082e-8 | 0.000005327 | 713.50 | 12139.53 |
| 3 | negative regulation of regulated secretory pathway (GO:1903306) | 0.000006290 | 0.0002736 | 832.67 | 9972.50 |
| 4 | Rap protein signal transduction (GO:0032486) | 0.00001010 | 0.0003767 | 624.44 | 7182.69 |
| 5 | positive regulation of integrin activation (GO:0033625) | 0.00001235 | 0.0004028 | 555.03 | 6273.06 |

GO: Molecular Function

| **Index** | **Name** | **P-value** | **Adjusted p-value** | **Odds Ratio** | **Combined score** |
| --- | --- | --- | --- | --- | --- |
| 1 | GDP binding (GO:0019003) | 9.011e-11 | 2.433e-9 | 321.42 | 7434.41 |
| 2 | neurotrophin TRK receptor binding (GO:0005167) | 0.002997 | 0.008990 | 444.11 | 2580.41 |
| 3 | neurotrophin TRKA receptor binding (GO:0005168) | 0.002997 | 0.008990 | 444.11 | 2580.41 |
| 4 | GTP binding (GO:0005525) | 1.733e-8 | 2.283e-7 | 107.64 | 1923.63 |
| 5 | guanyl ribonucleotide binding (GO:0032561) | 3.229e-8 | 2.283e-7 | 94.65 | 1632.51 |

GO: Cellular Component

| **Index** | **Name** | **P-value** | **Adjusted p-value** | **Odds Ratio** | **Combined score** |
| --- | --- | --- | --- | --- | --- |
| 1 | glutamatergic synapse (GO:0098978) | 0.0005185 | 0.005012 | 74.34 | 562.35 |
| 2 | aggresome (GO:0016235) | 0.01737 | 0.07195 | 65.22 | 264.33 |
| 3 | secretory granule membrane (GO:0030667) | 0.0002842 | 0.005012 | 31.18 | 254.65 |
| 4 | membrane raft (GO:0045121) | 0.002846 | 0.01651 | 30.79 | 180.49 |
| 5 | azurophil granule membrane (GO:0035577) | 0.02863 | 0.1038 | 38.86 | 138.07 |

Lymphadenopathy ([HP:0002716](https://hpo.jax.org/app/browse/term/HP:0002716))

KEGG Pathway

| **Index** | **Name** | **P-value** | **Adjusted p-value** | **Odds Ratio** | **Combined score** |
| --- | --- | --- | --- | --- | --- |
| 1 | Autoimmune thyroid disease | 9.117e-9 | 3.145e-7 | 271.31 | 5022.72 |
| 2 | Allograft rejection | 7.523e-7 | 0.00001038 | 244.35 | 3445.31 |
| 3 | Primary immunodeficiency | 7.523e-7 | 0.00001038 | 244.35 | 3445.31 |
| 4 | Natural killer cell mediated cytotoxicity | 2.740e-9 | 1.891e-7 | 157.65 | 3108.13 |
| 5 | Graft-versus-host disease | 0.000001023 | 0.00001083 | 219.24 | 3024.01 |

GO: Biological Process

| **Index** | **Name** | **P-value** | **Adjusted p-value** | **Odds Ratio** | **Combined score** |
| --- | --- | --- | --- | --- | --- |
| 1 | cellular response to interleukin-2 (GO:0071352) | 1.482e-8 | 0.000001912 | 1070.46 | 19297.66 |
| 2 | interleukin-2-mediated signaling pathway (GO:0038110) | 1.482e-8 | 0.000001912 | 1070.46 | 19297.66 |
| 3 | positive regulation of killing of cells of other organism (GO:0051712) | 0.000003371 | 0.0001112 | 1249.13 | 15739.23 |
| 4 | T cell apoptotic process (GO:0070231) | 0.000004719 | 0.0001353 | 999.25 | 12254.80 |
| 5 | positive regulation of cell killing (GO:0031343) | 0.00002354 | 0.0005607 | 384.17 | 4094.01 |

GO: Molecular Function

| **Index** | **Name** | **P-value** | **Adjusted p-value** | **Odds Ratio** | **Combined score** |
| --- | --- | --- | --- | --- | --- |
| 1 | glucocorticoid receptor binding (GO:0035259) | 0.004492 | 0.02545 | 277.53 | 1500.17 |
| 2 | tumor necrosis factor-activated receptor activity (GO:0005031) | 0.004492 | 0.02545 | 277.53 | 1500.17 |
| 3 | non-membrane spanning protein tyrosine kinase activity (GO:0004715) | 0.0001327 | 0.002172 | 151.19 | 1349.73 |
| 4 | phosphotyrosine residue binding (GO:0001784) | 0.0001485 | 0.002172 | 142.54 | 1256.48 |
| 5 | protein phosphorylated amino acid binding (GO:0045309) | 0.0001917 | 0.002172 | 124.69 | 1067.29 |

GO: Cellular Component

| **Index** | **Name** | **P-value** | **Adjusted p-value** | **Odds Ratio** | **Combined score** |
| --- | --- | --- | --- | --- | --- |
| 1 | T cell receptor complex (GO:0042101) | 0.00001481 | 0.0002073 | 499.50 | 5554.53 |
| 2 | CD95 death-inducing signaling complex (GO:0031265) | 0.002498 | 0.009549 | 555.17 | 3326.77 |
| 3 | cytolytic granule (GO:0044194) | 0.003495 | 0.009549 | 370.07 | 2093.27 |
| 4 | death-inducing signaling complex (GO:0031264) | 0.003994 | 0.009549 | 317.19 | 1751.86 |
| 5 | early phagosome (GO:0032009) | 0.005985 | 0.01197 | 201.81 | 1032.95 |

Lymphopenia ([HP:0001888](https://hpo.jax.org/app/browse/term/HP:0001888))

KEGG Pathway

| **Index** | **Name** | **P-value** | **Adjusted p-value** | **Odds Ratio** | **Combined score** |
| --- | --- | --- | --- | --- | --- |
| 1 | Primary immunodeficiency | 6.488e-15 | 1.687e-13 | 935.53 | 30562.71 |
| 2 | Th1 and Th2 cell differentiation | 4.565e-10 | 5.115e-9 | 228.77 | 4920.26 |
| 3 | Hematopoietic cell lineage | 6.629e-10 | 5.115e-9 | 211.66 | 4473.28 |
| 4 | T cell receptor signaling pathway | 8.515e-10 | 5.115e-9 | 200.92 | 4196.00 |
| 5 | Th17 cell differentiation | 9.837e-10 | 5.115e-9 | 194.98 | 4043.83 |

GO: Biological Process

| **Index** | **Name** | **P-value** | **Adjusted p-value** | **Odds Ratio** | **Combined score** |
| --- | --- | --- | --- | --- | --- |
| 1 | positive thymic T cell selection (GO:0045059) | 1.574e-13 | 1.858e-11 | 13326.00 | 392846.55 |
| 2 | V(D)J recombination (GO:0033151) | 0.00001235 | 0.0001457 | 555.03 | 6273.06 |
| 3 | T cell differentiation in thymus (GO:0033077) | 0.00002041 | 0.0002189 | 416.21 | 4494.85 |
| 4 | antigen receptor-mediated signaling pathway (GO:0050851) | 1.176e-10 | 6.937e-9 | 166.01 | 3795.72 |
| 5 | regulation of T cell differentiation (GO:0045580) | 8.148e-7 | 0.00001868 | 237.55 | 3330.49 |

GO: Molecular Function

| **Index** | **Name** | **P-value** | **Adjusted p-value** | **Odds Ratio** | **Combined score** |
| --- | --- | --- | --- | --- | --- |
| 1 | signaling receptor complex adaptor activity (GO:0030159) | 0.0001651 | 0.006108 | 134.82 | 1174.13 |
| 2 | endodeoxyribonuclease activity (GO:0004520) | 0.009957 | 0.06999 | 116.79 | 538.34 |
| 3 | phosphatidylinositol-3,5-bisphosphate binding (GO:0080025) | 0.01194 | 0.06999 | 96.46 | 427.12 |
| 4 | phosphatidylinositol-3,4-bisphosphate binding (GO:0043325) | 0.01293 | 0.06999 | 88.73 | 385.85 |
| 5 | phosphatidylinositol-3,4,5-trisphosphate binding (GO:0005547) | 0.01737 | 0.06999 | 65.22 | 264.33 |

GO: Cellular Component

| **Index** | **Name** | **P-value** | **Adjusted p-value** | **Odds Ratio** | **Combined score** |
| --- | --- | --- | --- | --- | --- |
| 1 | alpha-beta T cell receptor complex (GO:0042105) | 1.799e-9 | 1.169e-8 | 2855.29 | 57494.92 |
| 2 | T cell receptor complex (GO:0042101) | 1.556e-11 | 2.023e-10 | 1665.17 | 41439.58 |
| 3 | clathrin-coated endocytic vesicle (GO:0045334) | 0.000008701 | 0.00003770 | 104.05 | 1212.39 |
| 4 | pericentric heterochromatin (GO:0005721) | 0.006979 | 0.01134 | 170.74 | 847.70 |
| 5 | clathrin-coated endocytic vesicle membrane (GO:0030669) | 0.0005185 | 0.001348 | 74.34 | 562.35 |
