## Supplementary Table 1-3 for "Long COVID: G Protein-Coupled Receptors (GPCRs) responsible for persistent post-COVID symptoms": Supplementary File 2.2-Dermatological-finding.docx

Dermatographical-finding

Alopecia ([HP:0001596](https://hpo.jax.org/app/browse/term/HP:0001596))

KEGG Pathway

| **Index** | **Name** | **P-value** | **Adjusted p-value** | **Odds Ratio** | **Combined score** |
| --- | --- | --- | --- | --- | --- |
| 1 | Gap junction | 3.640e-10 | 3.232e-8 | 239.84 | 5212.76 |
| 2 | GnRH signaling pathway | 4.823e-10 | 3.232e-8 | 226.16 | 4851.66 |
| 3 | Long-term depression | 1.516e-8 | 3.385e-7 | 237.31 | 4272.71 |
| 4 | GnRH secretion | 1.973e-8 | 3.762e-7 | 221.44 | 3928.66 |
| 5 | Cholinergic synapse | 1.297e-9 | 5.648e-8 | 184.09 | 3767.09 |

GO: Biological Process

| **Index** | **Name** | **P-value** | **Adjusted p-value** | **Odds Ratio** | **Combined score** |
| --- | --- | --- | --- | --- | --- |
| 1 | activation of protein kinase A activity (GO:0034199) | 0.00003048 | 0.001338 | 332.92 | 3461.84 |
| 2 | regulation of miRNA metabolic process (GO:2000628) | 0.002498 | 0.02623 | 555.17 | 3326.77 |
| 3 | negative regulation of regulatory T cell differentiation (GO:0045590) | 0.002498 | 0.02623 | 555.17 | 3326.77 |
| 4 | cellular response to glucagon stimulus (GO:0071377) | 0.00003830 | 0.001338 | 293.72 | 2987.15 |
| 5 | response to glucagon (GO:0033762) | 0.00004701 | 0.001357 | 262.78 | 2618.60 |

GO: Molecular Function

| **Index** | **Name** | **P-value** | **Adjusted p-value** | **Odds Ratio** | **Combined score** |
| --- | --- | --- | --- | --- | --- |
| 1 | cAMP-dependent protein kinase activity (GO:0004691) | 0.002498 | 0.01049 | 555.17 | 3326.77 |
| 2 | cyclic nucleotide-dependent protein kinase activity (GO:0004690) | 0.002997 | 0.01049 | 444.11 | 2580.41 |
| 3 | G protein-coupled serotonin receptor binding (GO:0031821) | 0.002997 | 0.01049 | 444.11 | 2580.41 |
| 4 | apolipoprotein receptor binding (GO:0034190) | 0.002997 | 0.01049 | 444.11 | 2580.41 |
| 5 | sphingosine N-acyltransferase activity (GO:0050291) | 0.003495 | 0.01112 | 370.07 | 2093.27 |

GO: Cellular Component

| **Index** | **Name** | **P-value** | **Adjusted p-value** | **Odds Ratio** | **Combined score** |
| --- | --- | --- | --- | --- | --- |
| 1 | lamellar body (GO:0042599) | 0.006482 | 0.03025 | 184.98 | 932.06 |
| 2 | membrane raft (GO:0045121) | 0.00006115 | 0.0008561 | 53.12 | 515.34 |
| 3 | plasma membrane raft (GO:0044853) | 0.0007315 | 0.004096 | 62.22 | 449.25 |
| 4 | heterotrimeric G-protein complex (GO:0005834) | 0.01638 | 0.06553 | 69.30 | 284.93 |
| 5 | Golgi membrane (GO:0000139) | 0.00005745 | 0.0008561 | 27.81 | 271.55 |

Flushing ([HP:0031284](https://hpo.jax.org/app/browse/term/HP:0031284))

KEGG Pathway

| **Index** | **Name** | **P-value** | **Adjusted p-value** | **Odds Ratio** | **Combined score** |
| --- | --- | --- | --- | --- | --- |
| 1 | Lysine degradation | 0.0004323 | 0.005336 | 81.68 | 632.69 |
| 2 | Central carbon metabolism in cancer | 0.0005336 | 0.005336 | 73.24 | 551.94 |
| 3 | Citrate cycle (TCA cycle) | 0.01490 | 0.07451 | 76.48 | 321.69 |
| 4 | Thyroid cancer | 0.01835 | 0.07865 | 61.59 | 246.23 |
| 5 | MAPK signaling pathway | 0.0003495 | 0.005336 | 29.01 | 230.90 |

GO: Biological Process

| **Index** | **Name** | **P-value** | **Adjusted p-value** | **Odds Ratio** | **Combined score** |
| --- | --- | --- | --- | --- | --- |
| 1 | glial cell-derived neurotrophic factor receptor signaling pathway (GO:0035860) | 0.002498 | 0.02897 | 555.17 | 3326.77 |
| 2 | positive regulation of smooth muscle cell differentiation (GO:0051152) | 0.002498 | 0.02897 | 555.17 | 3326.77 |
| 3 | positive regulation of leukocyte mediated immunity (GO:0002705) | 0.002498 | 0.02897 | 555.17 | 3326.77 |
| 4 | immature B cell differentiation (GO:0002327) | 0.002498 | 0.02897 | 555.17 | 3326.77 |
| 5 | regulation of dendritic cell cytokine production (GO:0002730) | 0.002498 | 0.02897 | 555.17 | 3326.77 |

GO: Molecular Function

| **Index** | **Name** | **P-value** | **Adjusted p-value** | **Odds Ratio** | **Combined score** |
| --- | --- | --- | --- | --- | --- |
| 1 | histone methyltransferase activity (H3-K36 specific) (GO:0046975) | 0.00001235 | 0.0005308 | 555.03 | 6273.06 |
| 2 | histone methyltransferase activity (H4-K20 specific) (GO:0042799) | 0.002498 | 0.01790 | 555.17 | 3326.77 |
| 3 | DNA-methyltransferase activity (GO:0009008) | 0.002997 | 0.01841 | 444.11 | 2580.41 |
| 4 | S-methyltransferase activity (GO:0008172) | 0.003495 | 0.01879 | 370.07 | 2093.27 |
| 5 | phosphatidylethanolamine binding (GO:0008429) | 0.004492 | 0.02146 | 277.53 | 1500.17 |

GO: Cellular Component

| **Index** | **Name** | **P-value** | **Adjusted p-value** | **Odds Ratio** | **Combined score** |
| --- | --- | --- | --- | --- | --- |
| 1 | MLL1 complex (GO:0071339) | 0.01293 | 0.05371 | 88.73 | 385.85 |
| 2 | MLL1/2 complex (GO:0044665) | 0.01293 | 0.05371 | 88.73 | 385.85 |
| 3 | axon (GO:0030424) | 0.0001190 | 0.002619 | 42.19 | 381.27 |
| 4 | vesicle membrane (GO:0012506) | 0.03591 | 0.1129 | 30.74 | 102.25 |
| 5 | neuron projection (GO:0043005) | 0.002216 | 0.02438 | 15.06 | 92.07 |

Fragile nails ([HP:0001808](https://hpo.jax.org/app/browse/term/HP:0001808))

KEGG Pathway

| **Index** | **Name** | **P-value** | **Adjusted p-value** | **Odds Ratio** | **Combined score** |
| --- | --- | --- | --- | --- | --- |
| 1 | Basal transcription factors | 0.0002202 | 0.008572 | 115.97 | 976.58 |
| 2 | Nucleotide excision repair | 0.0002403 | 0.008572 | 110.81 | 923.40 |
| 3 | Long-term potentiation | 0.0004889 | 0.01151 | 76.63 | 584.21 |
| 4 | Glioma | 0.0006123 | 0.01151 | 68.21 | 504.62 |
| 5 | Arrhythmogenic right ventricular cardiomyopathy | 0.0006453 | 0.01151 | 66.38 | 487.64 |

GO: Biological Process

| **Index** | **Name** | **P-value** | **Adjusted p-value** | **Odds Ratio** | **Combined score** |
| --- | --- | --- | --- | --- | --- |
| 1 | hemidesmosome assembly (GO:0031581) | 0.00001481 | 0.001888 | 499.50 | 5554.53 |
| 2 | regulation of miRNA metabolic process (GO:2000628) | 0.002498 | 0.01370 | 555.17 | 3326.77 |
| 3 | positive regulation of intrinsic apoptotic signaling pathway by p53 class mediator (GO:1902255) | 0.002498 | 0.01370 | 555.17 | 3326.77 |
| 4 | nucleotide-excision repair, DNA incision, 3'-to lesion (GO:0006295) | 0.00004701 | 0.001888 | 262.78 | 2618.60 |
| 5 | nucleotide-excision repair, preincision complex stabilization (GO:0006293) | 0.00004701 | 0.001888 | 262.78 | 2618.60 |

GO: Molecular Function

| **Index** | **Name** | **P-value** | **Adjusted p-value** | **Odds Ratio** | **Combined score** |
| --- | --- | --- | --- | --- | --- |
| 1 | neuregulin binding (GO:0038132) | 0.002498 | 0.02709 | 555.17 | 3326.77 |
| 2 | cell adhesive protein binding involved in bundle of His cell-Purkinje myocyte communication (GO:0086083) | 0.002498 | 0.02709 | 555.17 | 3326.77 |
| 3 | calcium-dependent protein serine/threonine phosphatase activity (GO:0004723) | 0.003495 | 0.02709 | 370.07 | 2093.27 |
| 4 | 5'-3' DNA helicase activity (GO:0043139) | 0.003495 | 0.02709 | 370.07 | 2093.27 |
| 5 | protein binding involved in heterotypic cell-cell adhesion (GO:0086080) | 0.004990 | 0.03094 | 246.68 | 1307.49 |

GO: Cellular Component

| **Index** | **Name** | **P-value** | **Adjusted p-value** | **Odds Ratio** | **Combined score** |
| --- | --- | --- | --- | --- | --- |
| 1 | hemidesmosome (GO:0030056) | 0.000004719 | 0.0001746 | 999.25 | 12254.80 |
| 2 | transcription factor TFIIH core complex (GO:0000439) | 0.00001010 | 0.0001827 | 624.44 | 7182.69 |
| 3 | transcription factor TFIIH holo complex (GO:0005675) | 0.00001481 | 0.0001827 | 499.50 | 5554.53 |
| 4 | transcription factor TFIID complex (GO:0005669) | 0.0001038 | 0.0009603 | 172.08 | 1578.44 |
| 5 | intercalated disc (GO:0014704) | 0.01540 | 0.09034 | 73.93 | 308.54 |

Hyperhidrosis ([HP:0000975](https://hpo.jax.org/app/browse/term/HP:0000975))

KEGG Pathway

| **Index** | **Name** | **P-value** | **Adjusted p-value** | **Odds Ratio** | **Combined score** |
| --- | --- | --- | --- | --- | --- |
| 1 | Maturity onset diabetes of the young | 0.01293 | 0.08047 | 88.73 | 385.85 |
| 2 | Citrate cycle (TCA cycle) | 0.01490 | 0.08047 | 76.48 | 321.69 |
| 3 | Thyroid cancer | 0.01835 | 0.08258 | 61.59 | 246.23 |
| 4 | Transcriptional misregulation in cancer | 0.003922 | 0.05294 | 26.05 | 144.36 |
| 5 | Central carbon metabolism in cancer | 0.03446 | 0.1196 | 32.08 | 108.04 |

GO: Biological Process

| **Index** | **Name** | **P-value** | **Adjusted p-value** | **Odds Ratio** | **Combined score** |
| --- | --- | --- | --- | --- | --- |
| 1 | establishment of protein localization to telomere (GO:0070200) | 0.002498 | 0.03493 | 555.17 | 3326.77 |
| 2 | glial cell-derived neurotrophic factor receptor signaling pathway (GO:0035860) | 0.002498 | 0.03493 | 555.17 | 3326.77 |
| 3 | Schwann cell differentiation (GO:0014037) | 0.002498 | 0.03493 | 555.17 | 3326.77 |
| 4 | astrocyte development (GO:0014002) | 0.002997 | 0.03493 | 444.11 | 2580.41 |
| 5 | positive regulation of adenylate cyclase activity (GO:0045762) | 0.003495 | 0.03493 | 370.07 | 2093.27 |

GO: Molecular Function

| **Index** | **Name** | **P-value** | **Adjusted p-value** | **Odds Ratio** | **Combined score** |
| --- | --- | --- | --- | --- | --- |
| 1 | four-way junction DNA binding (GO:0000400) | 0.00003048 | 0.0007162 | 332.92 | 3461.84 |
| 2 | Y-form DNA binding (GO:0000403) | 0.002498 | 0.01280 | 555.17 | 3326.77 |
| 3 | RNA-directed DNA polymerase activity (GO:0003964) | 0.002997 | 0.01280 | 444.11 | 2580.41 |
| 4 | telomerase activity (GO:0003720) | 0.002997 | 0.01280 | 444.11 | 2580.41 |
| 5 | phosphatidylethanolamine binding (GO:0008429) | 0.004492 | 0.01759 | 277.53 | 1500.17 |

GO: Cellular Component

| **Index** | **Name** | **P-value** | **Adjusted p-value** | **Odds Ratio** | **Combined score** |
| --- | --- | --- | --- | --- | --- |
| 1 | transferase complex, transferring phosphorus-containing groups (GO:0061695) | 0.004492 | 0.03878 | 277.53 | 1500.17 |
| 2 | Ino80 complex (GO:0031011) | 0.006979 | 0.03878 | 170.74 | 847.70 |
| 3 | INO80-type complex (GO:0097346) | 0.01194 | 0.03878 | 96.46 | 427.12 |
| 4 | MLL1 complex (GO:0071339) | 0.01293 | 0.03878 | 88.73 | 385.85 |
| 5 | MLL1/2 complex (GO:0044665) | 0.01293 | 0.03878 | 88.73 | 385.85 |

Inflammatory abnormality of the skin ([HP:0011123](https://hpo.jax.org/app/browse/term/HP:0011123))

KEGG Pathway

| **Index** | **Name** | **P-value** | **Adjusted p-value** | **Odds Ratio** | **Combined score** |
| --- | --- | --- | --- | --- | --- |
| 1 | Primary immunodeficiency | 6.488e-15 | 1.687e-13 | 935.53 | 30562.71 |
| 2 | Th1 and Th2 cell differentiation | 4.565e-10 | 5.934e-9 | 228.77 | 4920.26 |
| 3 | T cell receptor signaling pathway | 8.515e-10 | 6.394e-9 | 200.92 | 4196.00 |
| 4 | Th17 cell differentiation | 9.837e-10 | 6.394e-9 | 194.98 | 4043.83 |
| 5 | PD-L1 expression and PD-1 checkpoint pathway in cancer | 7.537e-8 | 3.919e-7 | 156.12 | 2560.47 |

GO: Biological Process

| **Index** | **Name** | **P-value** | **Adjusted p-value** | **Odds Ratio** | **Combined score** |
| --- | --- | --- | --- | --- | --- |
| 1 | positive thymic T cell selection (GO:0045059) | 1.574e-13 | 1.401e-11 | 13326.00 | 392846.55 |
| 2 | antigen receptor-mediated signaling pathway (GO:0050851) | 6.059e-13 | 2.696e-11 | 259.71 | 7306.12 |
| 3 | V(D)J recombination (GO:0033151) | 0.00001235 | 0.00009988 | 555.03 | 6273.06 |
| 4 | lymphocyte differentiation (GO:0030098) | 2.870e-10 | 6.785e-9 | 252.04 | 5537.62 |
| 5 | B cell activation (GO:0042113) | 3.049e-10 | 6.785e-9 | 248.88 | 5453.09 |

GO: Molecular Function

| **Index** | **Name** | **P-value** | **Adjusted p-value** | **Odds Ratio** | **Combined score** |
| --- | --- | --- | --- | --- | --- |
| 1 | signaling receptor complex adaptor activity (GO:0030159) | 0.0001651 | 0.005613 | 134.82 | 1174.13 |
| 2 | endodeoxyribonuclease activity (GO:0004520) | 0.009957 | 0.07074 | 116.79 | 538.34 |
| 3 | phosphatidylinositol-3,5-bisphosphate binding (GO:0080025) | 0.01194 | 0.07074 | 96.46 | 427.12 |
| 4 | phosphatidylinositol-3,4-bisphosphate binding (GO:0043325) | 0.01293 | 0.07074 | 88.73 | 385.85 |
| 5 | phosphatidylinositol-3,4,5-trisphosphate binding (GO:0005547) | 0.01737 | 0.07074 | 65.22 | 264.33 |

GO: Cellular Component

| **Index** | **Name** | **P-value** | **Adjusted p-value** | **Odds Ratio** | **Combined score** |
| --- | --- | --- | --- | --- | --- |
| 1 | alpha-beta T cell receptor complex (GO:0042105) | 1.799e-9 | 1.259e-8 | 2855.29 | 57494.92 |
| 2 | T cell receptor complex (GO:0042101) | 1.556e-11 | 2.179e-10 | 1665.17 | 41439.58 |
| 3 | clathrin-coated endocytic vesicle (GO:0045334) | 0.000008701 | 0.00004060 | 104.05 | 1212.39 |
| 4 | clathrin-coated endocytic vesicle membrane (GO:0030669) | 0.0005185 | 0.001452 | 74.34 | 562.35 |
| 5 | clathrin-coated vesicle membrane (GO:0030665) | 0.0008802 | 0.002054 | 56.54 | 397.78 |

Petechiae ([HP:0000967](https://hpo.jax.org/app/browse/term/HP:0000967))

KEGG Pathway

| **Index** | **Name** | **P-value** | **Adjusted p-value** | **Odds Ratio** | **Combined score** |
| --- | --- | --- | --- | --- | --- |
| 1 | Platelet activation | 2.872e-7 | 0.00001120 | 110.39 | 1662.81 |
| 2 | ECM-receptor interaction | 0.000009659 | 0.0001790 | 100.36 | 1158.94 |
| 3 | Hematopoietic cell lineage | 0.00001377 | 0.0001790 | 88.81 | 994.11 |
| 4 | Allograft rejection | 0.01884 | 0.09536 | 59.92 | 237.98 |
| 5 | Graft-versus-host disease | 0.02081 | 0.09536 | 54.06 | 209.35 |

GO: Biological Process

| **Index** | **Name** | **P-value** | **Adjusted p-value** | **Odds Ratio** | **Combined score** |
| --- | --- | --- | --- | --- | --- |
| 1 | regulation of glycoprotein metabolic process (GO:1903018) | 0.002498 | 0.02593 | 555.17 | 3326.77 |
| 2 | extracellular exosome biogenesis (GO:0097734) | 0.002498 | 0.02593 | 555.17 | 3326.77 |
| 3 | positive regulation of killing of cells of other organism (GO:0051712) | 0.002997 | 0.02593 | 444.11 | 2580.41 |
| 4 | taurine metabolic process (GO:0019530) | 0.002997 | 0.02593 | 444.11 | 2580.41 |
| 5 | positive regulation of erythrocyte differentiation (GO:0045648) | 0.00007845 | 0.003662 | 199.65 | 1887.30 |

GO: Molecular Function

| **Index** | **Name** | **P-value** | **Adjusted p-value** | **Odds Ratio** | **Combined score** |
| --- | --- | --- | --- | --- | --- |
| 1 | glucocorticoid receptor binding (GO:0035259) | 0.004492 | 0.04040 | 277.53 | 1500.17 |
| 2 | C2H2 zinc finger domain binding (GO:0070742) | 0.005985 | 0.04040 | 201.81 | 1032.95 |
| 3 | syntaxin-1 binding (GO:0017075) | 0.01095 | 0.04795 | 105.66 | 476.99 |
| 4 | wide pore channel activity (GO:0022829) | 0.01243 | 0.04795 | 92.44 | 405.55 |
| 5 | channel activity (GO:0015267) | 0.02277 | 0.06147 | 49.25 | 186.27 |

GO: Cellular Component

| **Index** | **Name** | **P-value** | **Adjusted p-value** | **Odds Ratio** | **Combined score** |
| --- | --- | --- | --- | --- | --- |
| 1 | cytolytic granule (GO:0044194) | 0.000004719 | 0.0001557 | 999.25 | 12254.80 |
| 2 | multivesicular body membrane (GO:0032585) | 0.004492 | 0.02139 | 277.53 | 1500.17 |
| 3 | primary lysosome (GO:0005766) | 0.005488 | 0.02139 | 222.00 | 1155.57 |
| 4 | melanosome membrane (GO:0033162) | 0.005985 | 0.02139 | 201.81 | 1032.95 |
| 5 | pigment granule membrane (GO:0090741) | 0.005985 | 0.02139 | 201.81 | 1032.95 |

Pruritus ([HP:0000989](https://hpo.jax.org/app/browse/term/HP:0000989))

KEGG Pathway

| **Index** | **Name** | **P-value** | **Adjusted p-value** | **Odds Ratio** | **Combined score** |
| --- | --- | --- | --- | --- | --- |
| 1 | Primary immunodeficiency | 0.01884 | 0.09914 | 59.92 | 237.98 |
| 2 | ABC transporters | 0.02228 | 0.09914 | 50.37 | 191.61 |
| 3 | Sphingolipid metabolism | 0.02424 | 0.09914 | 46.16 | 171.72 |
| 4 | Arachidonic acid metabolism | 0.03009 | 0.09914 | 36.91 | 129.31 |
| 5 | Acute myeloid leukemia | 0.03301 | 0.09914 | 33.54 | 114.41 |

GO: Biological Process

| **Index** | **Name** | **P-value** | **Adjusted p-value** | **Odds Ratio** | **Combined score** |
| --- | --- | --- | --- | --- | --- |
| 1 | ceramide metabolic process (GO:0006672) | 6.794e-14 | 1.175e-11 | 610.44 | 18508.64 |
| 2 | ceramide biosynthetic process (GO:0046513) | 7.979e-12 | 6.901e-10 | 539.27 | 13780.66 |
| 3 | amide biosynthetic process (GO:0043604) | 2.433e-11 | 1.403e-9 | 424.32 | 10370.14 |
| 4 | regulation of water loss via skin (GO:0033561) | 8.685e-8 | 0.000002146 | 535.02 | 8698.92 |
| 5 | establishment of skin barrier (GO:0061436) | 8.685e-8 | 0.000002146 | 535.02 | 8698.92 |

GO: Molecular Function

| **Index** | **Name** | **P-value** | **Adjusted p-value** | **Odds Ratio** | **Combined score** |
| --- | --- | --- | --- | --- | --- |
| 1 | oxidoreductase activity, acting on single donors with incorporation of molecular oxygen, incorporation of two atoms of oxygen (GO:0016702) | 0.00003428 | 0.0009598 | 312.09 | 3208.63 |
| 2 | apolipoprotein receptor binding (GO:0034190) | 0.002997 | 0.02447 | 444.11 | 2580.41 |
| 3 | sphingosine N-acyltransferase activity (GO:0050291) | 0.003495 | 0.02447 | 370.07 | 2093.27 |
| 4 | NAD-retinol dehydrogenase activity (GO:0004745) | 0.009462 | 0.04577 | 123.28 | 574.57 |
| 5 | endodeoxyribonuclease activity (GO:0004520) | 0.009957 | 0.04577 | 116.79 | 538.34 |

GO: Cellular Component

| **Index** | **Name** | **P-value** | **Adjusted p-value** | **Odds Ratio** | **Combined score** |
| --- | --- | --- | --- | --- | --- |
| 1 | lamellar body (GO:0042599) | 0.006482 | 0.05834 | 184.98 | 932.06 |
| 2 | lipid droplet (GO:0005811) | 0.03785 | 0.1414 | 29.11 | 95.32 |
| 3 | endoplasmic reticulum membrane (GO:0005789) | 0.04712 | 0.1414 | 6.79 | 20.74 |
| 4 | Golgi membrane (GO:0000139) | 0.2125 | 0.4781 | 4.60 | 7.13 |
| 5 | bounding membrane of organelle (GO:0098588) | 0.3237 | 0.5827 | 2.79 | 3.15 |

Scaling skin ([HP:0040189](https://hpo.jax.org/app/browse/term/HP:0040189))

KEGG Pathway

| **Index** | **Name** | **P-value** | **Adjusted p-value** | **Odds Ratio** | **Combined score** |
| --- | --- | --- | --- | --- | --- |
| 1 | Terpenoid backbone biosynthesis | 0.01095 | 0.06911 | 105.66 | 476.99 |
| 2 | Hematopoietic cell lineage | 0.001064 | 0.02872 | 51.27 | 351.00 |
| 3 | Various types of N-glycan biosynthesis | 0.01933 | 0.08459 | 58.34 | 230.20 |
| 4 | Notch signaling pathway | 0.02912 | 0.08459 | 38.18 | 135.03 |
| 5 | Acute myeloid leukemia | 0.03301 | 0.08459 | 33.54 | 114.41 |

GO: Biological Process

| **Index** | **Name** | **P-value** | **Adjusted p-value** | **Odds Ratio** | **Combined score** |
| --- | --- | --- | --- | --- | --- |
| 1 | receptor transactivation (GO:0035624) | 0.002498 | 0.02820 | 555.17 | 3326.77 |
| 2 | positive regulation of smooth muscle cell differentiation (GO:0051152) | 0.002498 | 0.02820 | 555.17 | 3326.77 |
| 3 | positive regulation of leukocyte mediated immunity (GO:0002705) | 0.002498 | 0.02820 | 555.17 | 3326.77 |
| 4 | immature B cell differentiation (GO:0002327) | 0.002498 | 0.02820 | 555.17 | 3326.77 |
| 5 | regulation of dendritic cell cytokine production (GO:0002730) | 0.002498 | 0.02820 | 555.17 | 3326.77 |

GO: Molecular Function

| **Index** | **Name** | **P-value** | **Adjusted p-value** | **Odds Ratio** | **Combined score** |
| --- | --- | --- | --- | --- | --- |
| 1 | N-acetylgalactosamine 4-O-sulfotransferase activity (GO:0001537) | 0.002498 | 0.01806 | 555.17 | 3326.77 |
| 2 | keratin filament binding (GO:1990254) | 0.002997 | 0.01806 | 444.11 | 2580.41 |
| 3 | interleukin-6 receptor binding (GO:0005138) | 0.003495 | 0.01806 | 370.07 | 2093.27 |
| 4 | potassium channel inhibitor activity (GO:0019870) | 0.004492 | 0.01989 | 277.53 | 1500.17 |
| 5 | cysteine-type endopeptidase activity involved in execution phase of apoptosis (GO:0097200) | 0.006482 | 0.02472 | 184.98 | 932.06 |

GO: Cellular Component

| **Index** | **Name** | **P-value** | **Adjusted p-value** | **Odds Ratio** | **Combined score** |
| --- | --- | --- | --- | --- | --- |
| 1 | desmosome (GO:0030057) | 0.008469 | 0.08046 | 138.71 | 661.82 |
| 2 | aggresome (GO:0016235) | 0.01737 | 0.08384 | 65.22 | 264.33 |
| 3 | cornified envelope (GO:0001533) | 0.02130 | 0.08384 | 52.77 | 203.13 |
| 4 | cell-cell junction (GO:0005911) | 0.007662 | 0.08046 | 18.33 | 89.29 |
| 5 | intrinsic component of endoplasmic reticulum membrane (GO:0031227) | 0.05889 | 0.1399 | 18.40 | 52.10 |

Skin rash ([HP:0000988](https://hpo.jax.org/app/browse/term/HP:0000988))

KEGG Pathway

| **Index** | **Name** | **P-value** | **Adjusted p-value** | **Odds Ratio** | **Combined score** |
| --- | --- | --- | --- | --- | --- |
| 1 | Primary immunodeficiency | 4.712e-12 | 1.461e-10 | 604.76 | 15772.67 |
| 2 | B cell receptor signaling pathway | 2.384e-10 | 3.695e-9 | 262.03 | 5805.76 |
| 3 | NF-kappa B signaling pathway | 0.00001596 | 0.0001649 | 84.39 | 932.17 |
| 4 | Osteoclast differentiation | 0.00002904 | 0.0002251 | 68.66 | 717.28 |
| 5 | Fc epsilon RI signaling pathway | 0.0005036 | 0.003122 | 75.47 | 573.09 |

GO: Biological Process

| **Index** | **Name** | **P-value** | **Adjusted p-value** | **Odds Ratio** | **Combined score** |
| --- | --- | --- | --- | --- | --- |
| 1 | regulation of neutrophil activation (GO:1902563) | 0.000002248 | 0.00007819 | 1665.58 | 21661.60 |
| 2 | B cell receptor signaling pathway (GO:0050853) | 2.614e-12 | 1.869e-10 | 688.31 | 18357.31 |
| 3 | lymphocyte differentiation (GO:0030098) | 9.479e-13 | 9.721e-11 | 382.92 | 10601.05 |
| 4 | B cell activation (GO:0042113) | 1.020e-12 | 9.721e-11 | 378.06 | 10438.72 |
| 5 | positive regulation of B cell differentiation (GO:0045579) | 0.000008085 | 0.0002102 | 713.68 | 8368.26 |

GO: Molecular Function

| **Index** | **Name** | **P-value** | **Adjusted p-value** | **Odds Ratio** | **Combined score** |
| --- | --- | --- | --- | --- | --- |
| 1 | phospholipase binding (GO:0043274) | 0.00002041 | 0.0002925 | 416.21 | 4494.85 |
| 2 | non-membrane spanning protein tyrosine kinase activity (GO:0004715) | 5.841e-7 | 0.00002512 | 267.29 | 3836.51 |
| 3 | IgG binding (GO:0019864) | 0.002498 | 0.01343 | 555.17 | 3326.77 |
| 4 | non-membrane spanning protein tyrosine phosphatase activity (GO:0004726) | 0.002997 | 0.01432 | 444.11 | 2580.41 |
| 5 | phosphatidylinositol-3,4,5-trisphosphate binding (GO:0005547) | 0.0001327 | 0.001277 | 151.19 | 1349.73 |

GO: Cellular Component

| **Index** | **Name** | **P-value** | **Adjusted p-value** | **Odds Ratio** | **Combined score** |
| --- | --- | --- | --- | --- | --- |
| 1 | T cell receptor complex (GO:0042101) | 0.00001481 | 0.0001925 | 499.50 | 5554.53 |
| 2 | early phagosome (GO:0032009) | 0.005985 | 0.02594 | 201.81 | 1032.95 |
| 3 | membrane raft (GO:0045121) | 0.00006115 | 0.0003975 | 53.12 | 515.34 |
| 4 | multivesicular body (GO:0005771) | 0.02326 | 0.06771 | 48.17 | 181.19 |
| 5 | cytoplasmic side of plasma membrane (GO:0009898) | 0.02717 | 0.06771 | 41.02 | 147.91 |
