## Supplementary Table 1-3 for "Long COVID: G Protein-Coupled Receptors (GPCRs) responsible for persistent post-COVID symptoms": Supplementary File 2.3-RGEM.docx

Reproductive-Genitourinary-Endocrine-Metabolism

Abnormal female reproductive system physiology ([HP:0030012](https://hpo.jax.org/app/browse/term/HP:0030012))

KEGG Pathway

| **Index** | **Name** | **P-value** | **Adjusted p-value** | **Odds Ratio** | **Combined score** |
| --- | --- | --- | --- | --- | --- |
| 1 | GnRH secretion | 0.000003690 | 0.00002583 | 140.02 | 1751.57 |
| 2 | Neuroactive ligand-receptor interaction | 3.287e-7 | 0.000004602 | 58.49 | 873.21 |
| 3 | GnRH signaling pathway | 0.0009395 | 0.004384 | 54.67 | 381.04 |
| 4 | Melanoma | 0.03543 | 0.1240 | 31.17 | 104.12 |
| 5 | Breast cancer | 0.07113 | 0.1448 | 15.10 | 39.92 |

GO: Biological Process

| **Index** | **Name** | **P-value** | **Adjusted p-value** | **Odds Ratio** | **Combined score** |
| --- | --- | --- | --- | --- | --- |
| 1 | neuroepithelial cell differentiation (GO:0060563) | 0.002997 | 0.03670 | 444.11 | 2580.41 |
| 2 | negative regulation of striated muscle tissue development (GO:0045843) | 0.003495 | 0.03670 | 370.07 | 2093.27 |
| 3 | regulation of growth hormone secretion (GO:0060123) | 0.003495 | 0.03670 | 370.07 | 2093.27 |
| 4 | nose development (GO:0043584) | 0.003495 | 0.03670 | 370.07 | 2093.27 |
| 5 | limb development (GO:0060173) | 0.00009070 | 0.007618 | 184.84 | 1720.52 |

GO: Molecular Function

| **Index** | **Name** | **P-value** | **Adjusted p-value** | **Odds Ratio** | **Combined score** |
| --- | --- | --- | --- | --- | --- |
| 1 | type 1 fibroblast growth factor receptor binding (GO:0005105) | 0.002498 | 0.02123 | 555.17 | 3326.77 |
| 2 | type 2 fibroblast growth factor receptor binding (GO:0005111) | 0.002498 | 0.02123 | 555.17 | 3326.77 |
| 3 | interleukin-17 receptor activity (GO:0030368) | 0.003994 | 0.02263 | 317.19 | 1751.86 |
| 4 | peptide hormone receptor binding (GO:0051428) | 0.006979 | 0.02366 | 170.74 | 847.70 |
| 5 | neuropeptide receptor binding (GO:0071855) | 0.008966 | 0.02366 | 130.54 | 615.43 |

GO: Cellular Component

| **Index** | **Name** | **P-value** | **Adjusted p-value** | **Odds Ratio** | **Combined score** |
| --- | --- | --- | --- | --- | --- |
| 1 | cilium (GO:0005929) | 0.1124 | 0.4624 | 9.30 | 20.33 |
| 2 | Golgi membrane (GO:0000139) | 0.2125 | 0.4624 | 4.60 | 7.13 |
| 3 | integral component of plasma membrane (GO:0005887) | 0.1613 | 0.4624 | 3.19 | 5.82 |
| 4 | neuron projection (GO:0043005) | 0.2457 | 0.4624 | 3.89 | 5.46 |
| 5 | nucleolus (GO:0005730) | 0.3117 | 0.4624 | 2.92 | 3.41 |

Decreased glomerular filtration rate ([HP:0012213](https://hpo.jax.org/app/browse/term/HP:0012213))

KEGG Pathway

| **Index** | **Name** | **P-value** | **Adjusted p-value** | **Odds Ratio** | **Combined score** |
| --- | --- | --- | --- | --- | --- |
| 1 | Collecting duct acid secretion | 0.01342 | 0.08282 | 85.32 | 367.79 |
| 2 | Tyrosine metabolism | 0.01786 | 0.08282 | 63.35 | 255.00 |
| 3 | Arrhythmogenic right ventricular cardiomyopathy | 0.03785 | 0.08282 | 29.11 | 95.32 |
| 4 | ECM-receptor interaction | 0.04315 | 0.08282 | 25.42 | 79.89 |
| 5 | Hypertrophic cardiomyopathy | 0.04411 | 0.08282 | 24.85 | 77.54 |

GO: Biological Process

| **Index** | **Name** | **P-value** | **Adjusted p-value** | **Odds Ratio** | **Combined score** |
| --- | --- | --- | --- | --- | --- |
| 1 | urate metabolic process (GO:0046415) | 0.00001235 | 0.0002551 | 555.03 | 6273.06 |
| 2 | erythrose 4-phosphate/phosphoenolpyruvate family amino acid catabolic process (GO:1902222) | 0.002498 | 0.02532 | 555.17 | 3326.77 |
| 3 | L-phenylalanine catabolic process (GO:0006559) | 0.002498 | 0.02532 | 555.17 | 3326.77 |
| 4 | urate transport (GO:0015747) | 0.002498 | 0.02532 | 555.17 | 3326.77 |
| 5 | tyrosine catabolic process (GO:0006572) | 0.002997 | 0.02532 | 444.11 | 2580.41 |

GO: Molecular Function

| **Index** | **Name** | **P-value** | **Adjusted p-value** | **Odds Ratio** | **Combined score** |
| --- | --- | --- | --- | --- | --- |
| 1 | urate transmembrane transporter activity (GO:0015143) | 0.000002248 | 0.00004244 | 1665.58 | 21661.60 |
| 2 | voltage-gated chloride channel activity (GO:0005247) | 0.00001010 | 0.00007746 | 624.44 | 7182.69 |
| 3 | voltage-gated anion channel activity (GO:0008308) | 0.00002690 | 0.0001547 | 356.71 | 3753.85 |
| 4 | carbohydrate:proton symporter activity (GO:0005351) | 0.003495 | 0.01608 | 370.07 | 2093.27 |
| 5 | chloride channel activity (GO:0005254) | 0.000003690 | 0.00004244 | 140.02 | 1751.57 |

GO: Cellular Component

| **Index** | **Name** | **P-value** | **Adjusted p-value** | **Odds Ratio** | **Combined score** |
| --- | --- | --- | --- | --- | --- |
| 1 | filopodium membrane (GO:0031527) | 0.005985 | 0.01995 | 201.81 | 1032.95 |
| 2 | excitatory synapse (GO:0060076) | 0.01243 | 0.02585 | 92.44 | 405.55 |
| 3 | cell projection membrane (GO:0031253) | 0.0009195 | 0.004598 | 55.28 | 386.48 |
| 4 | synaptic membrane (GO:0097060) | 0.01293 | 0.02585 | 88.73 | 385.85 |
| 5 | brush border membrane (GO:0031526) | 0.01835 | 0.03058 | 61.59 | 246.23 |

Diabetes mellitus ([HP:0000819](https://hpo.jax.org/app/browse/term/HP:0000819))

KEGG Pathway

| **Index** | **Name** | **P-value** | **Adjusted p-value** | **Odds Ratio** | **Combined score** |
| --- | --- | --- | --- | --- | --- |
| 1 | Phototransduction | 0.00008446 | 0.0003379 | 191.96 | 1800.45 |
| 2 | Retinol metabolism | 0.03349 | 0.06698 | 33.04 | 112.22 |
| 3 | Purine metabolism | 0.06267 | 0.07253 | 17.24 | 47.76 |
| 4 | Spliceosome | 0.07253 | 0.07253 | 14.80 | 38.82 |

GO: Biological Process

| **Index** | **Name** | **P-value** | **Adjusted p-value** | **Odds Ratio** | **Combined score** |
| --- | --- | --- | --- | --- | --- |
| 1 | protein localization to ciliary transition zone (GO:1904491) | 0.002997 | 0.02825 | 444.11 | 2580.41 |
| 2 | protein localization to non-motile cilium (GO:0097499) | 0.002997 | 0.02825 | 444.11 | 2580.41 |
| 3 | visual perception (GO:0007601) | 9.004e-8 | 0.000003238 | 149.07 | 2418.38 |
| 4 | sensory perception of light stimulus (GO:0050953) | 9.813e-8 | 0.000003238 | 145.78 | 2352.45 |
| 5 | regulation of rhodopsin mediated signaling pathway (GO:0022400) | 0.00006709 | 0.001476 | 217.03 | 2085.58 |

GO: Molecular Function

| **Index** | **Name** | **P-value** | **Adjusted p-value** | **Odds Ratio** | **Combined score** |
| --- | --- | --- | --- | --- | --- |
| 1 | guanylate cyclase activator activity (GO:0030250) | 0.002498 | 0.03296 | 555.17 | 3326.77 |
| 2 | NADP-retinol dehydrogenase activity (GO:0052650) | 0.002997 | 0.03296 | 444.11 | 2580.41 |
| 3 | leucine zipper domain binding (GO:0043522) | 0.004990 | 0.03469 | 246.68 | 1307.49 |
| 4 | LRR domain binding (GO:0030275) | 0.006979 | 0.03469 | 170.74 | 847.70 |
| 5 | alcohol dehydrogenase (NADP+) activity (GO:0008106) | 0.008469 | 0.03469 | 138.71 | 661.82 |

GO: Cellular Component

| **Index** | **Name** | **P-value** | **Adjusted p-value** | **Odds Ratio** | **Combined score** |
| --- | --- | --- | --- | --- | --- |
| 1 | spliceosomal tri-snRNP complex (GO:0097526) | 0.01589 | 0.06098 | 71.54 | 296.32 |
| 2 | U4/U6 x U5 tri-snRNP complex (GO:0046540) | 0.01589 | 0.06098 | 71.54 | 296.32 |
| 3 | U2-type precatalytic spliceosome (GO:0071005) | 0.02473 | 0.06098 | 45.22 | 167.30 |
| 4 | precatalytic spliceosome (GO:0071011) | 0.02570 | 0.06098 | 43.44 | 159.04 |
| 5 | sperm flagellum (GO:0036126) | 0.02570 | 0.06098 | 43.44 | 159.04 |

Abnormality of the digestive system ([HP:0025031](https://hpo.jax.org/app/browse/term/HP:0025031))

GO: Biological Process

| **Index** | **Name** | **P-value** | **Adjusted p-value** | **Odds Ratio** | **Combined score** |
| --- | --- | --- | --- | --- | --- |
| 1 | axoneme assembly (GO:0035082) | 2.534e-18 | 9.374e-17 | 1725.20 | 69899.68 |
| 2 | regulation of cilium movement (GO:0003352) | 6.605e-12 | 8.147e-11 | 2220.44 | 57161.21 |
| 3 | determination of digestive tract left/right asymmetry (GO:0071907) | 3.147e-9 | 2.911e-8 | 2141.36 | 41921.23 |
| 4 | epithelial cilium movement involved in determination of left/right asymmetry (GO:0060287) | 7.548e-9 | 5.072e-8 | 1427.43 | 26695.75 |
| 5 | determination of pancreatic left/right asymmetry (GO:0035469) | 0.000002248 | 0.000004621 | 1665.58 | 21661.60 |

GO: Molecular Function

| **Index** | **Name** | **P-value** | **Adjusted p-value** | **Odds Ratio** | **Combined score** |
| --- | --- | --- | --- | --- | --- |
| 1 | protein homodimerization activity (GO:0042803) | 0.2762 | 0.2762 | 3.39 | 4.36 |

GO: Cellular Conponent

| **Index** | **Name** | **P-value** | **Adjusted p-value** | **Odds Ratio** | **Combined score** |
| --- | --- | --- | --- | --- | --- |
| 1 | cilium (GO:0005929) | 0.000003817 | 0.00002672 | 56.53 | 705.26 |
| 2 | motile cilium (GO:0031514) | 0.0005036 | 0.001763 | 75.47 | 573.09 |
| 3 | nucleolus (GO:0005730) | 0.3117 | 0.5529 | 2.92 | 3.41 |
| 4 | nuclear lumen (GO:0031981) | 0.3159 | 0.5529 | 2.87 | 3.31 |
| 5 | intracellular non-membrane-bounded organelle (GO:0043232) | 0.4493 | 0.6290 | 1.81 | 1.45 |

Edema ([HP:0000969](https://hpo.jax.org/app/browse/term/HP:0000969))

KEGG Pathway

| **Index** | **Name** | **P-value** | **Adjusted p-value** | **Odds Ratio** | **Combined score** |
| --- | --- | --- | --- | --- | --- |
| 1 | Phototransduction | 0.00008446 | 0.002864 | 191.96 | 1800.45 |
| 2 | Thyroid cancer | 0.0001485 | 0.002864 | 142.54 | 1256.48 |
| 3 | Bladder cancer | 0.0001826 | 0.002864 | 127.89 | 1100.92 |
| 4 | Endometrial cancer | 0.0003664 | 0.002864 | 88.99 | 704.07 |
| 5 | VEGF signaling pathway | 0.0003792 | 0.002864 | 87.43 | 688.70 |

GO: Biological Process

| **Index** | **Name** | **P-value** | **Adjusted p-value** | **Odds Ratio** | **Combined score** |
| --- | --- | --- | --- | --- | --- |
| 1 | rhodopsin mediated signaling pathway (GO:0016056) | 0.00001235 | 0.0008240 | 555.03 | 6273.06 |
| 2 | phototransduction, visible light (GO:0007603) | 0.00002354 | 0.0008240 | 384.17 | 4094.01 |
| 3 | retinal rod cell development (GO:0046548) | 0.003495 | 0.01439 | 370.07 | 2093.27 |
| 4 | vitamin A metabolic process (GO:0006776) | 0.003495 | 0.01439 | 370.07 | 2093.27 |
| 5 | regulation of rhodopsin mediated signaling pathway (GO:0022400) | 0.00006709 | 0.001565 | 217.03 | 2085.58 |

GO: Molecular Function

| **Index** | **Name** | **P-value** | **Adjusted p-value** | **Odds Ratio** | **Combined score** |
| --- | --- | --- | --- | --- | --- |
| 1 | U2 snRNA binding (GO:0030620) | 0.002498 | 0.01597 | 555.17 | 3326.77 |
| 2 | 11-cis retinal binding (GO:0005502) | 0.002997 | 0.01597 | 444.11 | 2580.41 |
| 3 | pre-mRNA intronic binding (GO:0097157) | 0.003994 | 0.01597 | 317.19 | 1751.86 |
| 4 | U1 snRNA binding (GO:0030619) | 0.003994 | 0.01597 | 317.19 | 1751.86 |
| 5 | retinal binding (GO:0016918) | 0.004492 | 0.01597 | 277.53 | 1500.17 |

GO: Cellular Component

| **Index** | **Name** | **P-value** | **Adjusted p-value** | **Odds Ratio** | **Combined score** |
| --- | --- | --- | --- | --- | --- |
| 1 | U2-type catalytic step 1 spliceosome (GO:0071006) | 0.005985 | 0.05120 | 201.81 | 1032.95 |
| 2 | U5 snRNP (GO:0005682) | 0.008469 | 0.05120 | 138.71 | 661.82 |
| 3 | Golgi-associated vesicle membrane (GO:0030660) | 0.01194 | 0.05120 | 96.46 | 427.12 |
| 4 | U2-type catalytic step 2 spliceosome (GO:0071007) | 0.01490 | 0.05120 | 76.48 | 321.69 |
| 5 | ciliary membrane (GO:0060170) | 0.01540 | 0.05120 | 73.93 | 308.54 |

Female sexual dysfunction ([HP:0030014](https://hpo.jax.org/app/browse/term/HP:0030014))

KEGG Pathway

| **Index** | **Name** | **P-value** | **Adjusted p-value** | **Odds Ratio** | **Combined score** |
| --- | --- | --- | --- | --- | --- |
| 1 | GnRH secretion | 0.000003690 | 0.00002583 | 140.02 | 1751.57 |
| 2 | Neuroactive ligand-receptor interaction | 3.287e-7 | 0.000004602 | 58.49 | 873.21 |
| 3 | GnRH signaling pathway | 0.0009395 | 0.004384 | 54.67 | 381.04 |
| 4 | Melanoma | 0.03543 | 0.1240 | 31.17 | 104.12 |
| 5 | Breast cancer | 0.07113 | 0.1448 | 15.10 | 39.92 |

GO: Biological Process

| **Index** | **Name** | **P-value** | **Adjusted p-value** | **Odds Ratio** | **Combined score** |
| --- | --- | --- | --- | --- | --- |
| 1 | neuroepithelial cell differentiation (GO:0060563) | 0.002997 | 0.03670 | 444.11 | 2580.41 |
| 2 | negative regulation of striated muscle tissue development (GO:0045843) | 0.003495 | 0.03670 | 370.07 | 2093.27 |
| 3 | regulation of growth hormone secretion (GO:0060123) | 0.003495 | 0.03670 | 370.07 | 2093.27 |
| 4 | nose development (GO:0043584) | 0.003495 | 0.03670 | 370.07 | 2093.27 |
| 5 | limb development (GO:0060173) | 0.00009070 | 0.007618 | 184.84 | 1720.52 |

GO: Molecular Function

| **Index** | **Name** | **P-value** | **Adjusted p-value** | **Odds Ratio** | **Combined score** |
| --- | --- | --- | --- | --- | --- |
| 1 | type 1 fibroblast growth factor receptor binding (GO:0005105) | 0.002498 | 0.02123 | 555.17 | 3326.77 |
| 2 | type 2 fibroblast growth factor receptor binding (GO:0005111) | 0.002498 | 0.02123 | 555.17 | 3326.77 |
| 3 | interleukin-17 receptor activity (GO:0030368) | 0.003994 | 0.02263 | 317.19 | 1751.86 |
| 4 | peptide hormone receptor binding (GO:0051428) | 0.006979 | 0.02366 | 170.74 | 847.70 |
| 5 | neuropeptide receptor binding (GO:0071855) | 0.008966 | 0.02366 | 130.54 | 615.43 |

GO: Cellular Component

| **Index** | **Name** | **P-value** | **Adjusted p-value** | **Odds Ratio** | **Combined score** |
| --- | --- | --- | --- | --- | --- |
| 1 | cilium (GO:0005929) | 0.1124 | 0.4624 | 9.30 | 20.33 |
| 2 | Golgi membrane (GO:0000139) | 0.2125 | 0.4624 | 4.60 | 7.13 |
| 3 | integral component of plasma membrane (GO:0005887) | 0.1613 | 0.4624 | 3.19 | 5.82 |
| 4 | neuron projection (GO:0043005) | 0.2457 | 0.4624 | 3.89 | 5.46 |
| 5 | nucleolus (GO:0005730) | 0.3117 | 0.4624 | 2.92 | 3.41 |

Fever ([HP:0001945](https://hpo.jax.org/app/browse/term/HP:0001945)

KEGG Pathway

| **Index** | **Name** | **P-value** | **Adjusted p-value** | **Odds Ratio** | **Combined score** |
| --- | --- | --- | --- | --- | --- |
| 1 | Primary immunodeficiency | 4.712e-12 | 1.838e-10 | 604.76 | 15772.67 |
| 2 | B cell receptor signaling pathway | 0.000007524 | 0.0001467 | 109.41 | 1290.72 |
| 3 | Hematopoietic cell lineage | 0.00001377 | 0.0001556 | 88.81 | 994.11 |
| 4 | T cell receptor signaling pathway | 0.00001596 | 0.0001556 | 84.39 | 932.17 |
| 5 | PD-L1 expression and PD-1 checkpoint pathway in cancer | 0.0008609 | 0.005123 | 57.19 | 403.64 |

GO: Biological Process

| **Index** | **Name** | **P-value** | **Adjusted p-value** | **Odds Ratio** | **Combined score** |
| --- | --- | --- | --- | --- | --- |
| 1 | positive thymic T cell selection (GO:0045059) | 0.000002248 | 0.00007997 | 1665.58 | 21661.60 |
| 2 | lymphocyte differentiation (GO:0030098) | 9.479e-13 | 2.360e-10 | 382.92 | 10601.05 |
| 3 | B cell receptor signaling pathway (GO:0050853) | 1.451e-9 | 7.224e-8 | 443.56 | 9026.96 |
| 4 | B cell differentiation (GO:0030183) | 5.558e-11 | 6.920e-9 | 355.96 | 8405.46 |
| 5 | V(D)J recombination (GO:0033151) | 0.00001235 | 0.0003842 | 555.03 | 6273.06 |

GO: Molecular Function

| **Index** | **Name** | **P-value** | **Adjusted p-value** | **Odds Ratio** | **Combined score** |
| --- | --- | --- | --- | --- | --- |
| 1 | phospholipase binding (GO:0043274) | 0.006979 | 0.06082 | 170.74 | 847.70 |
| 2 | endodeoxyribonuclease activity (GO:0004520) | 0.009957 | 0.06082 | 116.79 | 538.34 |
| 3 | phosphatidylinositol-3,5-bisphosphate binding (GO:0080025) | 0.01194 | 0.06082 | 96.46 | 427.12 |
| 4 | phosphatidylinositol-3,4-bisphosphate binding (GO:0043325) | 0.01293 | 0.06082 | 88.73 | 385.85 |
| 5 | protein tyrosine kinase activity (GO:0004713) | 0.001264 | 0.04803 | 46.90 | 312.97 |

GO: Cellular Component

| **Index** | **Name** | **P-value** | **Adjusted p-value** | **Odds Ratio** | **Combined score** |
| --- | --- | --- | --- | --- | --- |
| 1 | T cell receptor complex (GO:0042101) | 1.975e-8 | 3.358e-7 | 951.48 | 16879.14 |
| 2 | alpha-beta T cell receptor complex (GO:0042105) | 0.000003371 | 0.00002866 | 1249.13 | 15739.23 |
| 3 | early phagosome (GO:0032009) | 0.005985 | 0.01978 | 201.81 | 1032.95 |
| 4 | pericentric heterochromatin (GO:0005721) | 0.006979 | 0.01978 | 170.74 | 847.70 |
| 5 | clathrin-coated endocytic vesicle (GO:0045334) | 0.0007857 | 0.004452 | 59.96 | 428.66 |

Heat intolerance ([HP:0002046](https://hpo.jax.org/app/browse/term/HP:0002046))

KEGG Pathway

| **Index** | **Name** | **P-value** | **Adjusted p-value** | **Odds Ratio** | **Combined score** |
| --- | --- | --- | --- | --- | --- |
| 1 | NF-kappa B signaling pathway | 0.00001596 | 0.0009258 | 84.39 | 932.17 |
| 2 | Other types of O-glycan biosynthesis | 0.02326 | 0.1047 | 48.17 | 181.19 |
| 3 | Viral carcinogenesis | 0.004372 | 0.1047 | 24.61 | 133.71 |
| 4 | Long-term depression | 0.02960 | 0.1047 | 37.53 | 132.12 |
| 5 | GnRH secretion | 0.03155 | 0.1047 | 35.14 | 121.47 |

GO: Biological Process

| **Index** | **Name** | **P-value** | **Adjusted p-value** | **Odds Ratio** | **Combined score** |
| --- | --- | --- | --- | --- | --- |
| 1 | positive regulation of low-density lipoprotein receptor activity (GO:1905599) | 0.002498 | 0.05769 | 555.17 | 3326.77 |
| 2 | progesterone receptor signaling pathway (GO:0050847) | 0.003994 | 0.06070 | 317.19 | 1751.86 |
| 3 | regulation of low-density lipoprotein particle clearance (GO:0010988) | 0.004990 | 0.06320 | 246.68 | 1307.49 |
| 4 | tumor necrosis factor-mediated signaling pathway (GO:0033209) | 0.00002215 | 0.001683 | 75.39 | 807.99 |
| 5 | odontogenesis of dentin-containing tooth (GO:0042475) | 0.007973 | 0.07527 | 147.96 | 714.91 |

GO: Molecular Function

| **Index** | **Name** | **P-value** | **Adjusted p-value** | **Odds Ratio** | **Combined score** |
| --- | --- | --- | --- | --- | --- |
| 1 | aromatic amino acid transmembrane transporter activity (GO:0015173) | 0.004492 | 0.04235 | 277.53 | 1500.17 |
| 2 | inositol 1,4,5 trisphosphate binding (GO:0070679) | 0.005488 | 0.04235 | 222.00 | 1155.57 |
| 3 | fucosyltransferase activity (GO:0008417) | 0.005985 | 0.04235 | 201.81 | 1032.95 |
| 4 | death receptor binding (GO:0005123) | 0.007476 | 0.04235 | 158.54 | 776.21 |
| 5 | calcium-release channel activity (GO:0015278) | 0.008469 | 0.04235 | 138.71 | 661.82 |

GO: Cellular Component

| **Index** | **Name** | **P-value** | **Adjusted p-value** | **Odds Ratio** | **Combined score** |
| --- | --- | --- | --- | --- | --- |
| 1 | platelet dense tubular network membrane (GO:0031095) | 0.004492 | 0.03352 | 277.53 | 1500.17 |
| 2 | platelet dense tubular network (GO:0031094) | 0.005488 | 0.03352 | 222.00 | 1155.57 |
| 3 | melanosome membrane (GO:0033162) | 0.005985 | 0.03352 | 201.81 | 1032.95 |
| 4 | pigment granule membrane (GO:0090741) | 0.005985 | 0.03352 | 201.81 | 1032.95 |
| 5 | chitosome (GO:0045009) | 0.005985 | 0.03352 | 201.81 | 1032.95 |

Hypothermia ([HP:0002045](https://hpo.jax.org/app/browse/term/HP:0002045))

KEGG Pathway

| **Index** | **Name** | **P-value** | **Adjusted p-value** | **Odds Ratio** | **Combined score** |
| --- | --- | --- | --- | --- | --- |
| 1 | Thyroid hormone synthesis | 1.608e-10 | 1.447e-9 | 284.57 | 6417.37 |
| 2 | Autoimmune thyroid disease | 9.117e-9 | 4.103e-8 | 271.31 | 5022.72 |
| 3 | Regulation of lipolysis in adipocytes | 0.0003294 | 0.0009883 | 94.04 | 754.04 |
| 4 | Citrate cycle (TCA cycle) | 0.01490 | 0.02009 | 76.48 | 321.69 |
| 5 | Propanoate metabolism | 0.01687 | 0.02009 | 67.20 | 274.29 |

GO: Biological Process

| **Index** | **Name** | **P-value** | **Adjusted p-value** | **Odds Ratio** | **Combined score** |
| --- | --- | --- | --- | --- | --- |
| 1 | thyroid hormone generation (GO:0006590) | 1.975e-8 | 0.000001007 | 951.48 | 16879.14 |
| 2 | response to gonadotropin (GO:0034698) | 0.002997 | 0.02206 | 444.11 | 2580.41 |
| 3 | cellular response to gonadotropin stimulus (GO:0071371) | 0.003495 | 0.02206 | 370.07 | 2093.27 |
| 4 | thyroid gland development (GO:0030878) | 0.003994 | 0.02206 | 317.19 | 1751.86 |
| 5 | embryonic hemopoiesis (GO:0035162) | 0.006482 | 0.02755 | 184.98 | 932.06 |

GO: Molecular Function

| **Index** | **Name** | **P-value** | **Adjusted p-value** | **Odds Ratio** | **Combined score** |
| --- | --- | --- | --- | --- | --- |
| 1 | anion:sodium symporter activity (GO:0015373) | 0.002498 | 0.01310 | 555.17 | 3326.77 |
| 2 | symporter activity (GO:0015293) | 0.004990 | 0.01310 | 246.68 | 1307.49 |
| 3 | methyl-CpG binding (GO:0008327) | 0.01194 | 0.02507 | 96.46 | 427.12 |
| 4 | inorganic anion transmembrane transporter activity (GO:0015103) | 0.01441 | 0.02751 | 79.21 | 335.86 |
| 5 | sequence-specific DNA binding (GO:0043565) | 0.0002742 | 0.005758 | 18.29 | 150.01 |

GO: Cellular Component

| **Index** | **Name** | **P-value** | **Adjusted p-value** | **Odds Ratio** | **Combined score** |
| --- | --- | --- | --- | --- | --- |
| 1 | extracellular membrane-bounded organelle (GO:0065010) | 0.02766 | 0.1310 | 40.27 | 144.50 |
| 2 | extracellular vesicle (GO:1903561) | 0.02912 | 0.1310 | 38.18 | 135.03 |
| 3 | basolateral plasma membrane (GO:0016323) | 0.07300 | 0.2150 | 14.70 | 38.46 |
| 4 | vesicle (GO:0031982) | 0.1074 | 0.2150 | 9.76 | 21.77 |
| 5 | mitochondrial matrix (GO:0005759) | 0.1610 | 0.2150 | 6.29 | 11.49 |

Irregular menstruation ([HP:0000858](https://hpo.jax.org/app/browse/term/HP:0000858))

KEGG Pathway

| **Index** | **Name** | **P-value** | **Adjusted p-value** | **Odds Ratio** | **Combined score** |
| --- | --- | --- | --- | --- | --- |
| 1 | Insulin signaling pathway | 3.435e-9 | 2.405e-7 | 150.44 | 2931.94 |
| 2 | Glucagon signaling pathway | 1.586e-7 | 0.000005551 | 128.72 | 2015.32 |
| 3 | Cushing syndrome | 7.027e-7 | 0.00001640 | 87.59 | 1240.99 |
| 4 | Morphine addiction | 0.00001068 | 0.0001496 | 96.93 | 1109.48 |
| 5 | Calcium signaling pathway | 0.000004013 | 0.00007022 | 55.80 | 693.40 |

GO: Biological Process

| **Index** | **Name** | **P-value** | **Adjusted p-value** | **Odds Ratio** | **Combined score** |
| --- | --- | --- | --- | --- | --- |
| 1 | glycogen catabolic process (GO:0005980) | 4.082e-8 | 0.000002964 | 713.50 | 12139.53 |
| 2 | glucan catabolic process (GO:0009251) | 4.082e-8 | 0.000002964 | 713.50 | 12139.53 |
| 3 | cellular polysaccharide catabolic process (GO:0044247) | 5.023e-8 | 0.000002964 | 658.58 | 11068.58 |
| 4 | negative regulation of cAMP-mediated signaling (GO:0043951) | 0.00001010 | 0.0002980 | 624.44 | 7182.69 |
| 5 | cellular glucan metabolic process (GO:0006073) | 0.00001235 | 0.0003122 | 555.03 | 6273.06 |

GO: Molecular Function

| **Index** | **Name** | **P-value** | **Adjusted p-value** | **Odds Ratio** | **Combined score** |
| --- | --- | --- | --- | --- | --- |
| 1 | cAMP-dependent protein kinase activity (GO:0004691) | 0.002498 | 0.01463 | 555.17 | 3326.77 |
| 2 | beta-2 adrenergic receptor binding (GO:0031698) | 0.002498 | 0.01463 | 555.17 | 3326.77 |
| 3 | Y-form DNA binding (GO:0000403) | 0.002498 | 0.01463 | 555.17 | 3326.77 |
| 4 | 3',5'-cyclic-nucleotide phosphodiesterase activity (GO:0004114) | 0.00004701 | 0.0006328 | 262.78 | 2618.60 |
| 5 | cyclic nucleotide-dependent protein kinase activity (GO:0004690) | 0.002997 | 0.01536 | 444.11 | 2580.41 |

GO: Cellular Component

| **Index** | **Name** | **P-value** | **Adjusted p-value** | **Odds Ratio** | **Combined score** |
| --- | --- | --- | --- | --- | --- |
| 1 | calcium channel complex (GO:0034704) | 0.0002202 | 0.002422 | 115.97 | 976.58 |
| 2 | cation channel complex (GO:0034703) | 0.0005802 | 0.002682 | 70.14 | 522.67 |
| 3 | plasma membrane raft (GO:0044853) | 0.0007315 | 0.002682 | 62.22 | 449.25 |
| 4 | voltage-gated calcium channel complex (GO:0005891) | 0.01540 | 0.03387 | 73.93 | 308.54 |
| 5 | membrane raft (GO:0045121) | 0.002846 | 0.007826 | 30.79 | 180.49 |

Male sexual dysfunction ([HP:0040307](https://hpo.jax.org/app/browse/term/HP:0040307))

KEGG Pathway

| **Index** | **Name** | **P-value** | **Adjusted p-value** | **Odds Ratio** | **Combined score** |
| --- | --- | --- | --- | --- | --- |
| 1 | GnRH secretion | 0.000003690 | 0.00002583 | 140.02 | 1751.57 |
| 2 | Neuroactive ligand-receptor interaction | 3.287e-7 | 0.000004602 | 58.49 | 873.21 |
| 3 | GnRH signaling pathway | 0.0009395 | 0.004384 | 54.67 | 381.04 |
| 4 | Melanoma | 0.03543 | 0.1240 | 31.17 | 104.12 |
| 5 | Breast cancer | 0.07113 | 0.1448 | 15.10 | 39.92 |

GO: Biological Process

| **Index** | **Name** | **P-value** | **Adjusted p-value** | **Odds Ratio** | **Combined score** |
| --- | --- | --- | --- | --- | --- |
| 1 | neuroepithelial cell differentiation (GO:0060563) | 0.002997 | 0.03670 | 444.11 | 2580.41 |
| 2 | negative regulation of striated muscle tissue development (GO:0045843) | 0.003495 | 0.03670 | 370.07 | 2093.27 |
| 3 | regulation of growth hormone secretion (GO:0060123) | 0.003495 | 0.03670 | 370.07 | 2093.27 |
| 4 | nose development (GO:0043584) | 0.003495 | 0.03670 | 370.07 | 2093.27 |
| 5 | limb development (GO:0060173) | 0.00009070 | 0.007618 | 184.84 | 1720.52 |

GO: Molecular Function

| **Index** | **Name** | **P-value** | **Adjusted p-value** | **Odds Ratio** | **Combined score** |
| --- | --- | --- | --- | --- | --- |
| 1 | type 1 fibroblast growth factor receptor binding (GO:0005105) | 0.002498 | 0.02123 | 555.17 | 3326.77 |
| 2 | type 2 fibroblast growth factor receptor binding (GO:0005111) | 0.002498 | 0.02123 | 555.17 | 3326.77 |
| 3 | interleukin-17 receptor activity (GO:0030368) | 0.003994 | 0.02263 | 317.19 | 1751.86 |
| 4 | peptide hormone receptor binding (GO:0051428) | 0.006979 | 0.02366 | 170.74 | 847.70 |
| 5 | neuropeptide receptor binding (GO:0071855) | 0.008966 | 0.02366 | 130.54 | 615.43 |

GO: Cellular Component

| **Index** | **Name** | **P-value** | **Adjusted p-value** | **Odds Ratio** | **Combined score** |
| --- | --- | --- | --- | --- | --- |
| 1 | cilium (GO:0005929) | 0.1124 | 0.4624 | 9.30 | 20.33 |
| 2 | Golgi membrane (GO:0000139) | 0.2125 | 0.4624 | 4.60 | 7.13 |
| 3 | integral component of plasma membrane (GO:0005887) | 0.1613 | 0.4624 | 3.19 | 5.82 |
| 4 | neuron projection (GO:0043005) | 0.2457 | 0.4624 | 3.89 | 5.46 |
| 5 | nucleolus (GO:0005730) | 0.3117 | 0.4624 | 2.92 | 3.41 |

Menorrhagia ([HP:0000132](https://hpo.jax.org/app/browse/term/HP:0000132))

KEGG Pathway

| **Index** | **Name** | **P-value** | **Adjusted p-value** | **Odds Ratio** | **Combined score** |
| --- | --- | --- | --- | --- | --- |
| 1 | Platelet activation | 1.731e-7 | 0.00001454 | 132.47 | 2062.51 |
| 2 | ECM-receptor interaction | 0.000006783 | 0.0002708 | 117.09 | 1393.55 |
| 3 | Hematopoietic cell lineage | 0.000009671 | 0.0002708 | 103.62 | 1196.43 |
| 4 | Dilated cardiomyopathy | 0.0008030 | 0.01686 | 60.48 | 431.03 |
| 5 | Vasopressin-regulated water reabsorption | 0.01963 | 0.09128 | 57.99 | 227.93 |

GO: Biological Process

| **Index** | **Name** | **P-value** | **Adjusted p-value** | **Odds Ratio** | **Combined score** |
| --- | --- | --- | --- | --- | --- |
| 1 | endosome to melanosome transport (GO:0035646) | 0.000006469 | 0.0003170 | 815.67 | 9746.03 |
| 2 | endosome to pigment granule transport (GO:0043485) | 0.000006469 | 0.0003170 | 815.67 | 9746.03 |
| 3 | anterograde synaptic vesicle transport (GO:0048490) | 0.00001884 | 0.0004292 | 439.08 | 4776.89 |
| 4 | synaptic vesicle transport along microtubule (GO:0099517) | 0.00001884 | 0.0004292 | 439.08 | 4776.89 |
| 5 | negative regulation of blood coagulation, intrinsic pathway (GO:2000267) | 0.002248 | 0.01185 | 624.59 | 3808.55 |

GO: Molecular Function

| **Index** | **Name** | **P-value** | **Adjusted p-value** | **Odds Ratio** | **Combined score** |
| --- | --- | --- | --- | --- | --- |
| 1 | leukemia inhibitory factor receptor activity (GO:0004923) | 0.002697 | 0.01112 | 499.65 | 2955.69 |
| 2 | oncostatin-M receptor activity (GO:0004924) | 0.002697 | 0.01112 | 499.65 | 2955.69 |
| 3 | ciliary neurotrophic factor receptor activity (GO:0004897) | 0.003595 | 0.01112 | 356.86 | 2008.48 |
| 4 | ciliary neurotrophic factor receptor binding (GO:0005127) | 0.004043 | 0.01112 | 312.23 | 1720.62 |
| 5 | protein kinase A regulatory subunit binding (GO:0034237) | 0.009858 | 0.02169 | 118.87 | 549.11 |

GO: Cellular Component

| **Index** | **Name** | **P-value** | **Adjusted p-value** | **Odds Ratio** | **Combined score** |
| --- | --- | --- | --- | --- | --- |
| 1 | platelet alpha granule membrane (GO:0031092) | 0.007625 | 0.06100 | 156.05 | 760.96 |
| 2 | platelet alpha granule (GO:0031091) | 0.0007063 | 0.01130 | 64.62 | 468.85 |
| 3 | endosome lumen (GO:0031904) | 0.01164 | 0.06209 | 99.83 | 444.56 |
| 4 | endoplasmic reticulum-Golgi intermediate compartment membrane (GO:0033116) | 0.02184 | 0.08498 | 51.93 | 198.60 |
| 5 | platelet alpha granule lumen (GO:0031093) | 0.02975 | 0.08498 | 37.74 | 132.64 |

Pancreatitis ([HP:0001733](https://hpo.jax.org/app/browse/term/HP:0001733))

KEGG Pathway

| **Index** | **Name** | **P-value** | **Adjusted p-value** | **Odds Ratio** | **Combined score** |
| --- | --- | --- | --- | --- | --- |
| 1 | Sulfur relay system | 0.003994 | 0.01864 | 317.19 | 1751.86 |
| 2 | Propanoate metabolism | 0.0001252 | 0.001752 | 155.92 | 1401.11 |
| 3 | Valine, leucine and isoleucine degradation | 0.0002507 | 0.001755 | 108.39 | 898.70 |
| 4 | Thiamine metabolism | 0.007476 | 0.02617 | 158.54 | 776.21 |
| 5 | Autoimmune thyroid disease | 0.02619 | 0.07247 | 42.60 | 155.17 |

GO: Biological Process

| **Index** | **Name** | **P-value** | **Adjusted p-value** | **Odds Ratio** | **Combined score** |
| --- | --- | --- | --- | --- | --- |
| 1 | positive regulation of toll-like receptor 9 signaling pathway (GO:0034165) | 0.002498 | 0.02080 | 555.17 | 3326.77 |
| 2 | negative regulation of regulatory T cell differentiation (GO:0045590) | 0.002498 | 0.02080 | 555.17 | 3326.77 |
| 3 | branched-chain amino acid catabolic process (GO:0009083) | 0.00004255 | 0.001616 | 277.39 | 2791.90 |
| 4 | branched-chain amino acid metabolic process (GO:0009081) | 0.00004255 | 0.001616 | 277.39 | 2791.90 |
| 5 | cobalamin metabolic process (GO:0009235) | 0.00004701 | 0.001616 | 262.78 | 2618.60 |

GO: Molecular Function

| **Index** | **Name** | **P-value** | **Adjusted p-value** | **Odds Ratio** | **Combined score** |
| --- | --- | --- | --- | --- | --- |
| 1 | oxidoreductase activity, acting on the aldehyde or oxo group of donors, disulfide as acceptor (GO:0016624) | 0.000004719 | 0.00009909 | 999.25 | 12254.80 |
| 2 | non-membrane spanning protein tyrosine phosphatase activity (GO:0004726) | 0.002997 | 0.01573 | 444.11 | 2580.41 |
| 3 | sulfurtransferase activity (GO:0016783) | 0.003994 | 0.01677 | 317.19 | 1751.86 |
| 4 | carboxy-lyase activity (GO:0016831) | 0.01144 | 0.02901 | 100.85 | 450.83 |
| 5 | phosphatidylinositol phospholipase C activity (GO:0004435) | 0.01144 | 0.02901 | 100.85 | 450.83 |

GO: Cellular Component

| **Index** | **Name** | **P-value** | **Adjusted p-value** | **Odds Ratio** | **Combined score** |
| --- | --- | --- | --- | --- | --- |
| 1 | mitochondrial alpha-ketoglutarate dehydrogenase complex (GO:0005947) | 0.000002248 | 0.00003147 | 1665.58 | 21661.60 |
| 2 | mitochondrial matrix (GO:0005759) | 0.0005723 | 0.004006 | 24.40 | 182.19 |
| 3 | cytoplasmic side of plasma membrane (GO:0009898) | 0.02717 | 0.09509 | 41.02 | 147.91 |
| 4 | clathrin-coated vesicle (GO:0030136) | 0.03930 | 0.09731 | 28.00 | 90.64 |
| 5 | clathrin-coated endocytic vesicle (GO:0045334) | 0.04171 | 0.09731 | 26.33 | 83.66 |

Recurrent fever ([HP:0001954](https://hpo.jax.org/app/browse/term/HP:0001954))

KEGG Pathway

| **Index** | **Name** | **P-value** | **Adjusted p-value** | **Odds Ratio** | **Combined score** |
| --- | --- | --- | --- | --- | --- |
| 1 | NOD-like receptor signaling pathway | 1.030e-10 | 5.666e-9 | 169.84 | 3905.74 |
| 2 | C-type lectin receptor signaling pathway | 0.00001596 | 0.0004389 | 84.39 | 932.17 |
| 3 | Yersinia infection | 0.00003643 | 0.0006678 | 63.51 | 649.04 |
| 4 | Fc gamma R-mediated phagocytosis | 0.001021 | 0.01075 | 52.36 | 360.55 |
| 5 | NF-kappa B signaling pathway | 0.001173 | 0.01075 | 48.75 | 328.95 |

GO: Biological Process

| **Index** | **Name** | **P-value** | **Adjusted p-value** | **Odds Ratio** | **Combined score** |
| --- | --- | --- | --- | --- | --- |
| 1 | positive regulation of killing of cells of other organism (GO:0051712) | 0.000003371 | 0.00009468 | 1249.13 | 15739.23 |
| 2 | negative regulation of interleukin-1 production (GO:0032692) | 3.628e-7 | 0.00001359 | 316.87 | 4699.01 |
| 3 | positive regulation of superoxide anion generation (GO:0032930) | 0.00002041 | 0.0005289 | 416.21 | 4494.85 |
| 4 | positive regulation of cysteine-type endopeptidase activity (GO:2001056) | 1.733e-8 | 0.000003461 | 229.10 | 4094.25 |
| 5 | positive regulation of cell killing (GO:0031343) | 0.00002354 | 0.0005289 | 384.17 | 4094.01 |

GO: Molecular Function

| **Index** | **Name** | **P-value** | **Adjusted p-value** | **Odds Ratio** | **Combined score** |
| --- | --- | --- | --- | --- | --- |
| 1 | endopeptidase activator activity (GO:0061133) | 0.004990 | 0.03381 | 246.68 | 1307.49 |
| 2 | caspase binding (GO:0089720) | 0.005985 | 0.03381 | 201.81 | 1032.95 |
| 3 | phosphatidate phosphatase activity (GO:0008195) | 0.006482 | 0.03381 | 184.98 | 932.06 |
| 4 | phospholipase binding (GO:0043274) | 0.006979 | 0.03381 | 170.74 | 847.70 |
| 5 | protein kinase C activity (GO:0004697) | 0.006979 | 0.03381 | 170.74 | 847.70 |

GO: Cellular Component

| **Index** | **Name** | **P-value** | **Adjusted p-value** | **Odds Ratio** | **Combined score** |
| --- | --- | --- | --- | --- | --- |
| 1 | cytolytic granule (GO:0044194) | 0.003495 | 0.02155 | 370.07 | 2093.27 |
| 2 | early phagosome (GO:0032009) | 0.005985 | 0.02155 | 201.81 | 1032.95 |
| 3 | T cell receptor complex (GO:0042101) | 0.005985 | 0.02155 | 201.81 | 1032.95 |
| 4 | endolysosome (GO:0036019) | 0.01243 | 0.03730 | 92.44 | 405.55 |
| 5 | spindle microtubule (GO:0005876) | 0.03009 | 0.06771 | 36.91 | 129.31 |

Renal insufficiency ([HP:0000083](https://hpo.jax.org/app/browse/term/HP:0000083))

KEGG Pathway

| **Index** | **Name** | **P-value** | **Adjusted p-value** | **Odds Ratio** | **Combined score** |
| --- | --- | --- | --- | --- | --- |
| 1 | Fanconi anemia pathway | 8.485e-27 | 3.394e-26 | 199460.00 | 11973883.48 |
| 2 | Homologous recombination | 0.02032 | 0.04063 | 55.42 | 215.92 |
| 3 | Pancreatic cancer | 0.03736 | 0.04982 | 29.50 | 96.98 |
| 4 | Pathways in cancer | 0.2360 | 0.2360 | 4.08 | 5.89 |

GO: Biological Process

| **Index** | **Name** | **P-value** | **Adjusted p-value** | **Odds Ratio** | **Combined score** |
| --- | --- | --- | --- | --- | --- |
| 1 | interstrand cross-link repair (GO:0036297) | 1.037e-26 | 7.674e-25 | 199450.00 | 11933257.10 |
| 2 | DNA repair (GO:0006281) | 4.636e-19 | 1.715e-17 | 197020.00 | 8317234.26 |
| 3 | double-strand break repair via synthesis-dependent strand annealing (GO:0045003) | 0.000002248 | 0.00002773 | 1665.58 | 21661.60 |
| 4 | resolution of meiotic recombination intermediates (GO:0000712) | 0.00002690 | 0.0002844 | 356.71 | 3753.85 |
| 5 | positive regulation of protein monoubiquitination (GO:1902527) | 0.002498 | 0.009641 | 555.17 | 3326.77 |

GO: Molecular Function

| **Index** | **Name** | **P-value** | **Adjusted p-value** | **Odds Ratio** | **Combined score** |
| --- | --- | --- | --- | --- | --- |
| 1 | DNA polymerase binding (GO:0070182) | 0.00003428 | 0.0006170 | 312.09 | 3208.63 |
| 2 | four-way junction helicase activity (GO:0009378) | 0.002997 | 0.01348 | 444.11 | 2580.41 |
| 3 | crossover junction endodeoxyribonuclease activity (GO:0008821) | 0.002997 | 0.01348 | 444.11 | 2580.41 |
| 4 | 5'-flap endonuclease activity (GO:0017108) | 0.003994 | 0.01348 | 317.19 | 1751.86 |
| 5 | endodeoxyribonuclease activity, producing 3'-phosphomonoesters (GO:0016889) | 0.004492 | 0.01348 | 277.53 | 1500.17 |

GO: Cellular Component

| **Index** | **Name** | **P-value** | **Adjusted p-value** | **Odds Ratio** | **Combined score** |
| --- | --- | --- | --- | --- | --- |
| 1 | condensed chromosome (GO:0000793) | 0.0003175 | 0.002858 | 95.86 | 772.11 |
| 2 | condensed nuclear chromosome (GO:0000794) | 0.009957 | 0.02240 | 116.79 | 538.34 |
| 3 | chromosome (GO:0005694) | 0.002744 | 0.01235 | 31.38 | 185.09 |
| 4 | nuclear chromosome (GO:0000228) | 0.04074 | 0.06623 | 26.98 | 86.34 |
| 5 | nuclear lumen (GO:0031981) | 0.005076 | 0.01523 | 11.12 | 58.74 |

Temperature instability ([HP:0005968](https://hpo.jax.org/app/browse/term/HP:0005968))

KEGG Pathway

| **Index** | **Name** | **P-value** | **Adjusted p-value** | **Odds Ratio** | **Combined score** |
| --- | --- | --- | --- | --- | --- |
| 1 | Hedgehog signaling pathway | 0.000002460 | 0.00006643 | 161.22 | 2082.13 |
| 2 | Phenylalanine metabolism | 0.008469 | 0.04573 | 138.71 | 661.82 |
| 3 | Basal cell carcinoma | 0.0004323 | 0.005837 | 81.68 | 632.69 |
| 4 | Tyrosine metabolism | 0.01786 | 0.08026 | 63.35 | 255.00 |
| 5 | Tryptophan metabolism | 0.02081 | 0.08026 | 54.06 | 209.35 |

GO: Biological Process

| **Index** | **Name** | **P-value** | **Adjusted p-value** | **Odds Ratio** | **Combined score** |
| --- | --- | --- | --- | --- | --- |
| 1 | regulation of nodal signaling pathway involved in determination of lateral mesoderm left/right asymmetry (GO:1900175) | 0.000002248 | 0.0002038 | 1665.58 | 21661.60 |
| 2 | positive regulation of T cell differentiation in thymus (GO:0033089) | 0.000003371 | 0.0002038 | 1249.13 | 15739.23 |
| 3 | dorsal/ventral pattern formation (GO:0009953) | 3.331e-10 | 8.062e-8 | 665.67 | 14526.48 |
| 4 | pituitary gland development (GO:0021983) | 0.000004719 | 0.0002038 | 999.25 | 12254.80 |
| 5 | proximal/distal pattern formation (GO:0009954) | 0.000006290 | 0.0002174 | 832.67 | 9972.50 |

GO: Molecular Function

| **Index** | **Name** | **P-value** | **Adjusted p-value** | **Odds Ratio** | **Combined score** |
| --- | --- | --- | --- | --- | --- |
| 1 | morphogen activity (GO:0016015) | 0.000004719 | 0.0001274 | 999.25 | 12254.80 |
| 2 | type 1 fibroblast growth factor receptor binding (GO:0005105) | 0.002498 | 0.01686 | 555.17 | 3326.77 |
| 3 | type 2 fibroblast growth factor receptor binding (GO:0005111) | 0.002498 | 0.01686 | 555.17 | 3326.77 |
| 4 | patched binding (GO:0005113) | 0.003495 | 0.01887 | 370.07 | 2093.27 |
| 5 | activin receptor binding (GO:0070697) | 0.004990 | 0.02245 | 246.68 | 1307.49 |

GO: Cellular Component

| **Index** | **Name** | **P-value** | **Adjusted p-value** | **Odds Ratio** | **Combined score** |
| --- | --- | --- | --- | --- | --- |
| 1 | U4 snRNP (GO:0005687) | 0.004492 | 0.03319 | 277.53 | 1500.17 |
| 2 | prespliceosome (GO:0071010) | 0.007476 | 0.03319 | 158.54 | 776.21 |
| 3 | U2-type prespliceosome (GO:0071004) | 0.007476 | 0.03319 | 158.54 | 776.21 |
| 4 | U5 snRNP (GO:0005682) | 0.008469 | 0.03319 | 138.71 | 661.82 |
| 5 | U1 snRNP (GO:0005685) | 0.008966 | 0.03319 | 130.54 | 615.43 |

Urinary incontinence ([HP:0000020](https://hpo.jax.org/app/browse/term/HP:0000020))

KEGG Pathway

| **Index** | **Name** | **P-value** | **Adjusted p-value** | **Odds Ratio** | **Combined score** |
| --- | --- | --- | --- | --- | --- |
| 1 | Arginine and proline metabolism | 0.02473 | 0.09066 | 45.22 | 167.30 |
| 2 | Non-small cell lung cancer | 0.03543 | 0.09743 | 31.17 | 104.12 |
| 3 | Amyotrophic lateral sclerosis | 0.01350 | 0.09066 | 13.56 | 58.36 |
| 4 | Dopaminergic synapse | 0.06409 | 0.1410 | 16.84 | 46.28 |
| 5 | Pathways of neurodegeneration | 0.02233 | 0.09066 | 10.32 | 39.22 |

GO: Biological Process

| **Index** | **Name** | **P-value** | **Adjusted p-value** | **Odds Ratio** | **Combined score** |
| --- | --- | --- | --- | --- | --- |
| 1 | synaptic vesicle transport (GO:0048489) | 0.00001235 | 0.0006999 | 555.03 | 6273.06 |
| 2 | synaptic vesicle localization (GO:0097479) | 0.00001750 | 0.0006999 | 454.07 | 4973.59 |
| 3 | proline biosynthetic process (GO:0006561) | 0.002498 | 0.01711 | 555.17 | 3326.77 |
| 4 | anterograde dendritic transport (GO:0098937) | 0.002498 | 0.01711 | 555.17 | 3326.77 |
| 5 | anterograde dendritic transport of neurotransmitter receptor complex (GO:0098971) | 0.002498 | 0.01711 | 555.17 | 3326.77 |

GO: Molecular Function

| **Index** | **Name** | **P-value** | **Adjusted p-value** | **Odds Ratio** | **Combined score** |
| --- | --- | --- | --- | --- | --- |
| 1 | gap junction channel activity involved in cell communication by electrical coupling (GO:1903763) | 0.002997 | 0.02873 | 444.11 | 2580.41 |
| 2 | outward rectifier potassium channel activity (GO:0015271) | 0.005488 | 0.02873 | 222.00 | 1155.57 |
| 3 | oxidoreductase activity, acting on the aldehyde or oxo group of donors, NAD or NADP as acceptor (GO:0016620) | 0.009957 | 0.02873 | 116.79 | 538.34 |
| 4 | gap junction channel activity (GO:0005243) | 0.01194 | 0.02873 | 96.46 | 427.12 |
| 5 | wide pore channel activity (GO:0022829) | 0.01243 | 0.02873 | 92.44 | 405.55 |

GO: Cellular Component

| **Index** | **Name** | **P-value** | **Adjusted p-value** | **Odds Ratio** | **Combined score** |
| --- | --- | --- | --- | --- | --- |
| 1 | endoplasmic reticulum tubular network membrane (GO:0098826) | 0.002498 | 0.05861 | 555.17 | 3326.77 |
| 2 | connexin complex (GO:0005922) | 0.01045 | 0.05861 | 110.94 | 506.01 |
| 3 | endoplasmic reticulum tubular network (GO:0071782) | 0.01144 | 0.05861 | 100.85 | 450.83 |
| 4 | gap junction (GO:0005921) | 0.01194 | 0.05861 | 96.46 | 427.12 |
| 5 | Golgi cis cisterna (GO:0000137) | 0.01243 | 0.05861 | 92.44 | 405.55 |
