## Supplementary Table 1-3 for "Long COVID: G Protein-Coupled Receptors (GPCRs) responsible for persistent post-COVID symptoms": Supplementary File 2.4-HEENT.docx

HEENT-ear

Ear pain ([HP:0030766](https://hpo.jax.org/app/browse/term/HP:0030766))

KEGG Pathway

| **Index** | **Name** | **P-value** | **Adjusted p-value** | **Odds Ratio** | **Combined score** |
| --- | --- | --- | --- | --- | --- |
| 1 | Hedgehog signaling pathway | 0.0001147 | 0.004940 | 184.63 | 1675.22 |
| 2 | Basal cell carcinoma | 0.0001453 | 0.004940 | 163.39 | 1443.80 |
| 3 | Pathways in cancer | 0.000007064 | 0.0007205 | 73.88 | 876.24 |
| 4 | Aldosterone-regulated sodium reabsorption | 0.01105 | 0.06141 | 110.88 | 499.54 |
| 5 | Gastric cancer | 0.0008109 | 0.02050 | 67.51 | 480.47 |

GO: Biological Process

| **Index** | **Name** | **P-value** | **Adjusted p-value** | **Odds Ratio** | **Combined score** |
| --- | --- | --- | --- | --- | --- |
| 1 | establishment of protein localization to telomere (GO:0070200) | 0.001499 | 0.01866 | 999.50 | 6499.56 |
| 2 | determination of left/right asymmetry in lateral mesoderm (GO:0003140) | 0.001499 | 0.01866 | 999.50 | 6499.56 |
| 3 | contact inhibition (GO:0060242) | 0.001799 | 0.01866 | 799.56 | 5053.71 |
| 4 | mesenchymal to epithelial transition involved in metanephros morphogenesis (GO:0003337) | 0.001799 | 0.01866 | 799.56 | 5053.71 |
| 5 | negative regulation of actin filament depolymerization (GO:0030835) | 0.001799 | 0.01866 | 799.56 | 5053.71 |

GO: Molecular Function

| **Index** | **Name** | **P-value** | **Adjusted p-value** | **Odds Ratio** | **Combined score** |
| --- | --- | --- | --- | --- | --- |
| 1 | RNA-directed DNA polymerase activity (GO:0003964) | 0.001799 | 0.01416 | 799.56 | 5053.71 |
| 2 | telomerase activity (GO:0003720) | 0.001799 | 0.01416 | 799.56 | 5053.71 |
| 3 | patched binding (GO:0005113) | 0.002098 | 0.01416 | 666.27 | 4108.59 |
| 4 | 1-phosphatidylinositol-4-phosphate 3-kinase activity (GO:0035005) | 0.002098 | 0.01416 | 666.27 | 4108.59 |
| 5 | 1-phosphatidylinositol-3-kinase activity (GO:0016303) | 0.002997 | 0.01481 | 444.11 | 2580.41 |

GO: Cellular Component

| **Index** | **Name** | **P-value** | **Adjusted p-value** | **Odds Ratio** | **Combined score** |
| --- | --- | --- | --- | --- | --- |
| 1 | phosphatidylinositol 3-kinase complex, class I (GO:0097651) | 0.001499 | 0.02697 | 999.50 | 6499.56 |
| 2 | transferase complex, transferring phosphorus-containing groups (GO:0061695) | 0.002697 | 0.02697 | 499.65 | 2955.69 |
| 3 | intercalated disc (GO:0014704) | 0.009265 | 0.03706 | 133.09 | 623.08 |
| 4 | ciliary membrane (GO:0060170) | 0.009265 | 0.03706 | 133.09 | 623.08 |
| 5 | cell-cell contact zone (GO:0044291) | 0.01402 | 0.04180 | 86.73 | 370.11 |

Hearing impairment ([HP:0000365](https://hpo.jax.org/app/browse/term/HP:0000365))

KEGG Pathway

| **Index** | **Name** | **P-value** | **Adjusted p-value** | **Odds Ratio** | **Combined score** |
| --- | --- | --- | --- | --- | --- |
| 1 | Phototransduction | 0.00008446 | 0.0005068 | 191.96 | 1800.45 |
| 2 | Retinol metabolism | 0.03349 | 0.09401 | 33.04 | 112.22 |
| 3 | Drug metabolism | 0.05272 | 0.09401 | 20.65 | 60.76 |
| 4 | Purine metabolism | 0.06267 | 0.09401 | 17.24 | 47.76 |
| 5 | cGMP-PKG signaling pathway | 0.08045 | 0.09654 | 13.27 | 33.44 |

GO: Biological Process

| **Index** | **Name** | **P-value** | **Adjusted p-value** | **Odds Ratio** | **Combined score** |
| --- | --- | --- | --- | --- | --- |
| 1 | protein localization to ciliary transition zone (GO:1904491) | 0.002997 | 0.02240 | 444.11 | 2580.41 |
| 2 | protein localization to non-motile cilium (GO:0097499) | 0.002997 | 0.02240 | 444.11 | 2580.41 |
| 3 | visual perception (GO:0007601) | 9.004e-8 | 0.000002895 | 149.07 | 2418.38 |
| 4 | sensory perception of light stimulus (GO:0050953) | 9.813e-8 | 0.000002895 | 145.78 | 2352.45 |
| 5 | regulation of rhodopsin mediated signaling pathway (GO:0022400) | 0.00006709 | 0.001319 | 217.03 | 2085.58 |

GO: Molecular Function

| **Index** | **Name** | **P-value** | **Adjusted p-value** | **Odds Ratio** | **Combined score** |
| --- | --- | --- | --- | --- | --- |
| 1 | guanylate cyclase activator activity (GO:0030250) | 0.002498 | 0.01921 | 555.17 | 3326.77 |
| 2 | NADP-retinol dehydrogenase activity (GO:0052650) | 0.002997 | 0.01921 | 444.11 | 2580.41 |
| 3 | intracellular cGMP-activated cation channel activity (GO:0005223) | 0.003495 | 0.01921 | 370.07 | 2093.27 |
| 4 | intracellular cAMP-activated cation channel activity (GO:0005222) | 0.004492 | 0.01921 | 277.53 | 1500.17 |
| 5 | leucine zipper domain binding (GO:0043522) | 0.004990 | 0.01921 | 246.68 | 1307.49 |

GO: Cellular Component

| **Index** | **Name** | **P-value** | **Adjusted p-value** | **Odds Ratio** | **Combined score** |
| --- | --- | --- | --- | --- | --- |
| 1 | sperm flagellum (GO:0036126) | 0.02570 | 0.1766 | 43.44 | 159.04 |
| 2 | 9+2 motile cilium (GO:0097729) | 0.02814 | 0.1766 | 39.55 | 141.22 |
| 3 | azurophil granule lumen (GO:0035578) | 0.04411 | 0.1766 | 24.85 | 77.54 |
| 4 | cytoplasmic vesicle lumen (GO:0060205) | 0.05605 | 0.1766 | 19.37 | 55.82 |
| 5 | ficolin-1-rich granule lumen (GO:1904813) | 0.05984 | 0.1766 | 18.09 | 50.96 |

Pulsatile tinnitus ([HP:0008629](https://hpo.jax.org/app/browse/term/HP:0008629))

KEGG Pathway

| **Index** | **Name** | **P-value** | **Adjusted p-value** | **Odds Ratio** | **Combined score** |
| --- | --- | --- | --- | --- | --- |
| 1 | Citrate cycle (TCA cycle) | 0.00006056 | 0.001453 | 237.67 | 2308.19 |
| 2 | Thyroid cancer | 0.01471 | 0.07585 | 79.19 | 334.14 |
| 3 | Tryptophan metabolism | 0.01668 | 0.07585 | 69.52 | 284.57 |
| 4 | Cysteine and methionine metabolism | 0.01983 | 0.07585 | 58.14 | 227.95 |
| 5 | Lysine degradation | 0.02493 | 0.07585 | 45.92 | 169.53 |

GO: Biological Process

| **Index** | **Name** | **P-value** | **Adjusted p-value** | **Odds Ratio** | **Combined score** |
| --- | --- | --- | --- | --- | --- |
| 1 | succinyl-CoA metabolic process (GO:0006104) | 0.001999 | 0.03464 | 713.86 | 4436.86 |
| 2 | glial cell-derived neurotrophic factor receptor signaling pathway (GO:0035860) | 0.001999 | 0.03464 | 713.86 | 4436.86 |
| 3 | Schwann cell differentiation (GO:0014037) | 0.001999 | 0.03464 | 713.86 | 4436.86 |
| 4 | astrocyte development (GO:0014002) | 0.002398 | 0.03464 | 571.06 | 3445.29 |
| 5 | positive regulation of adenylate cyclase activity (GO:0045762) | 0.002797 | 0.03464 | 475.86 | 2797.66 |

GO: Molecular Function

| **Index** | **Name** | **P-value** | **Adjusted p-value** | **Odds Ratio** | **Combined score** |
| --- | --- | --- | --- | --- | --- |
| 1 | DNA-methyltransferase activity (GO:0009008) | 0.002398 | 0.03235 | 571.06 | 3445.29 |
| 2 | S-methyltransferase activity (GO:0008172) | 0.002797 | 0.03235 | 475.86 | 2797.66 |
| 3 | phosphatidylethanolamine binding (GO:0008429) | 0.003595 | 0.03235 | 356.86 | 2008.48 |
| 4 | kinesin binding (GO:0019894) | 0.01154 | 0.07047 | 101.86 | 454.45 |
| 5 | E-box binding (GO:0070888) | 0.01629 | 0.07047 | 71.26 | 293.40 |

GO: Cellular Component

| **Index** | **Name** | **P-value** | **Adjusted p-value** | **Odds Ratio** | **Combined score** |
| --- | --- | --- | --- | --- | --- |
| 1 | oxoglutarate dehydrogenase complex (GO:0045252) | 0.001999 | 0.01599 | 713.86 | 4436.86 |
| 2 | axon (GO:0030424) | 0.00005639 | 0.001353 | 59.08 | 577.97 |
| 3 | MLL1 complex (GO:0071339) | 0.01035 | 0.03550 | 114.10 | 521.46 |
| 4 | MLL1/2 complex (GO:0044665) | 0.01035 | 0.03550 | 114.10 | 521.46 |
| 5 | neuron projection (GO:0043005) | 0.001078 | 0.01294 | 21.09 | 144.11 |

Tinnitus ([HP:0000360](https://hpo.jax.org/app/browse/term/HP:0000360))

KEGG Pathway

| **Index** | **Name** | **P-value** | **Adjusted p-value** | **Odds Ratio** | **Combined score** |
| --- | --- | --- | --- | --- | --- |
| 1 | Aldosterone synthesis and secretion | 0.00001335 | 0.0007611 | 89.75 | 1007.36 |
| 2 | GnRH secretion | 0.0004462 | 0.008477 | 80.35 | 619.92 |
| 3 | Circadian entrainment | 0.001021 | 0.01456 | 52.36 | 360.55 |
| 4 | Citrate cycle (TCA cycle) | 0.01490 | 0.06534 | 76.48 | 321.69 |
| 5 | Serotonergic synapse | 0.001382 | 0.01576 | 44.77 | 294.78 |

GO: Biological Process

| **Index** | **Name** | **P-value** | **Adjusted p-value** | **Odds Ratio** | **Combined score** |
| --- | --- | --- | --- | --- | --- |
| 1 | positive regulation of adenylate cyclase activity (GO:0045762) | 0.000004719 | 0.001368 | 999.25 | 12254.80 |
| 2 | positive regulation of cyclase activity (GO:0031281) | 0.00002041 | 0.002276 | 416.21 | 4494.85 |
| 3 | positive regulation of lyase activity (GO:0051349) | 0.00002354 | 0.002276 | 384.17 | 4094.01 |
| 4 | establishment of protein localization to telomere (GO:0070200) | 0.002498 | 0.03750 | 555.17 | 3326.77 |
| 5 | glial cell-derived neurotrophic factor receptor signaling pathway (GO:0035860) | 0.002498 | 0.03750 | 555.17 | 3326.77 |

GO: Molecular Function

| **Index** | **Name** | **P-value** | **Adjusted p-value** | **Odds Ratio** | **Combined score** |
| --- | --- | --- | --- | --- | --- |
| 1 | voltage-gated calcium channel activity involved in cardiac muscle cell action potential (GO:0086007) | 0.002498 | 0.03707 | 555.17 | 3326.77 |
| 2 | RNA-directed DNA polymerase activity (GO:0003964) | 0.002997 | 0.03707 | 444.11 | 2580.41 |
| 3 | telomerase activity (GO:0003720) | 0.002997 | 0.03707 | 444.11 | 2580.41 |
| 4 | oxidoreductase activity, acting on paired donors, with incorporation or reduction of molecular oxygen, reduced iron-sulfur protein as one donor, and incorporation of one atom of oxygen (GO:0016713) | 0.004492 | 0.03707 | 277.53 | 1500.17 |
| 5 | phosphatidylethanolamine binding (GO:0008429) | 0.004492 | 0.03707 | 277.53 | 1500.17 |

GO: Cellular Component

| **Index** | **Name** | **P-value** | **Adjusted p-value** | **Odds Ratio** | **Combined score** |
| --- | --- | --- | --- | --- | --- |
| 1 | transferase complex, transferring phosphorus-containing groups (GO:0061695) | 0.004492 | 0.03893 | 277.53 | 1500.17 |
| 2 | MLL1 complex (GO:0071339) | 0.01293 | 0.04003 | 88.73 | 385.85 |
| 3 | MLL1/2 complex (GO:0044665) | 0.01293 | 0.04003 | 88.73 | 385.85 |
| 4 | axon (GO:0030424) | 0.0001190 | 0.003095 | 42.19 | 381.27 |
| 5 | voltage-gated calcium channel complex (GO:0005891) | 0.01540 | 0.04003 | 73.93 | 308.54 |

Vertigo ([HP:0002321](https://hpo.jax.org/app/browse/term/HP:0002321))

KEGG Pathway

| **Index** | **Name** | **P-value** | **Adjusted p-value** | **Odds Ratio** | **Combined score** |
| --- | --- | --- | --- | --- | --- |
| 1 | Calcium signaling pathway | 0.000004013 | 0.0001445 | 55.80 | 693.40 |
| 2 | Adrenergic signaling in cardiomyocytes | 0.00004774 | 0.0008593 | 57.85 | 575.61 |
| 3 | Cardiac muscle contraction | 0.0008229 | 0.009193 | 58.54 | 415.82 |
| 4 | Circadian entrainment | 0.001021 | 0.009193 | 52.36 | 360.55 |
| 5 | Citrate cycle (TCA cycle) | 0.01490 | 0.06706 | 76.48 | 321.69 |

GO: Biological Process

| **Index** | **Name** | **P-value** | **Adjusted p-value** | **Odds Ratio** | **Combined score** |
| --- | --- | --- | --- | --- | --- |
| 1 | membrane depolarization during action potential (GO:0086010) | 6.190e-13 | 1.752e-10 | 950.90 | 26730.60 |
| 2 | membrane depolarization during AV node cell action potential (GO:0086045) | 0.000002248 | 0.00003535 | 1665.58 | 21661.60 |
| 3 | membrane depolarization during SA node cell action potential (GO:0086046) | 0.000002248 | 0.00003535 | 1665.58 | 21661.60 |
| 4 | SA node cell to atrial cardiac muscle cell signaling (GO:0086018) | 0.000002248 | 0.00003535 | 1665.58 | 21661.60 |
| 5 | cardiac muscle cell action potential involved in contraction (GO:0086002) | 1.597e-12 | 1.785e-10 | 767.85 | 20856.82 |

GO: Molecular Function

| **Index** | **Name** | **P-value** | **Adjusted p-value** | **Odds Ratio** | **Combined score** |
| --- | --- | --- | --- | --- | --- |
| 1 | voltage-gated sodium channel activity (GO:0005248) | 2.480e-13 | 8.184e-12 | 1174.88 | 34101.37 |
| 2 | voltage-gated sodium channel activity involved in cardiac muscle cell action potential (GO:0086006) | 0.000002248 | 0.00002473 | 1665.58 | 21661.60 |
| 3 | sodium channel activity (GO:0005272) | 4.712e-12 | 7.774e-11 | 604.76 | 15772.67 |
| 4 | voltage-gated calcium channel activity involved in cardiac muscle cell action potential (GO:0086007) | 0.002498 | 0.01374 | 555.17 | 3326.77 |
| 5 | nitric-oxide synthase binding (GO:0050998) | 0.003495 | 0.01647 | 370.07 | 2093.27 |

GO: Cellular Component

| **Index** | **Name** | **P-value** | **Adjusted p-value** | **Odds Ratio** | **Combined score** |
| --- | --- | --- | --- | --- | --- |
| 1 | voltage-gated sodium channel complex (GO:0001518) | 5.833e-14 | 1.867e-12 | 1664.83 | 50731.79 |
| 2 | sodium channel complex (GO:0034706) | 5.000e-13 | 8.001e-12 | 998.50 | 28281.59 |
| 3 | junctional sarcoplasmic reticulum membrane (GO:0014701) | 0.000008085 | 0.00003696 | 713.68 | 8368.26 |
| 4 | calcium channel complex (GO:0034704) | 0.000001263 | 0.00001347 | 203.55 | 2764.61 |
| 5 | sarcoplasmic reticulum membrane (GO:0033017) | 0.00008446 | 0.0003003 | 191.96 | 1800.45 |

HEENT-ENT

Abnormality of the pharynx ([HP:0000600](https://hpo.jax.org/app/browse/term/HP:0000600))

KEGG Pathway

| **Index** | **Name** | **P-value** | **Adjusted p-value** | **Odds Ratio** | **Combined score** |
| --- | --- | --- | --- | --- | --- |
| 1 | Huntington disease | 0.009682 | 0.02024 | 16.19 | 75.08 |
| 2 | Amyotrophic lateral sclerosis | 0.01350 | 0.02024 | 13.56 | 58.36 |
| 3 | Pathways of neurodegeneration | 0.02233 | 0.02233 | 10.32 | 39.22 |

GO: Biological Process

| **Index** | **Name** | **P-value** | **Adjusted p-value** | **Odds Ratio** | **Combined score** |
| --- | --- | --- | --- | --- | --- |
| 1 | axonemal dynein complex assembly (GO:0070286) | 1.125e-9 | 1.305e-8 | 475.29 | 9793.38 |
| 2 | axoneme assembly (GO:0035082) | 1.451e-9 | 1.305e-8 | 443.56 | 9026.96 |
| 3 | outer dynein arm assembly (GO:0036158) | 0.00003428 | 0.0001234 | 312.09 | 3208.63 |
| 4 | cilium movement (GO:0003341) | 0.000001964 | 0.00001178 | 174.41 | 2291.88 |
| 5 | determination of bilateral symmetry (GO:0009855) | 0.00009070 | 0.0002721 | 184.84 | 1720.52 |

GO: Molecular Function

| **Index** | **Name** | **P-value** | **Adjusted p-value** | **Odds Ratio** | **Combined score** |
| --- | --- | --- | --- | --- | --- |
| 1 | motor activity (GO:0003774) | 0.0004745 | 0.001423 | 77.84 | 595.70 |
| 2 | microtubule motor activity (GO:0003777) | 0.02766 | 0.04148 | 40.27 | 144.50 |
| 3 | nucleoside-triphosphatase activity (GO:0017111) | 0.1306 | 0.1306 | 7.91 | 16.09 |

GO: Cellular Component

| **Index** | **Name** | **P-value** | **Adjusted p-value** | **Odds Ratio** | **Combined score** |
| --- | --- | --- | --- | --- | --- |
| 1 | outer dynein arm (GO:0036157) | 0.000008085 | 0.00008893 | 713.68 | 8368.26 |
| 2 | motile cilium (GO:0031514) | 0.0005036 | 0.001847 | 75.47 | 573.09 |
| 3 | cilium (GO:0005929) | 0.0001854 | 0.001020 | 36.18 | 310.92 |
| 4 | sperm flagellum (GO:0036126) | 0.02570 | 0.06192 | 43.44 | 159.04 |
| 5 | 9+2 motile cilium (GO:0097729) | 0.02814 | 0.06192 | 39.55 | 141.22 |

Dysphonia ([HP:0001618](https://hpo.jax.org/app/browse/term/HP:0001618))

KEGG Pathway

| **Index** | **Name** | **P-value** | **Adjusted p-value** | **Odds Ratio** | **Combined score** |
| --- | --- | --- | --- | --- | --- |
| 1 | Citrate cycle (TCA cycle) | 0.01490 | 0.08712 | 76.48 | 321.69 |
| 2 | Cholinergic synapse | 0.001382 | 0.03456 | 44.77 | 294.78 |
| 3 | Thyroid cancer | 0.01835 | 0.08712 | 61.59 | 246.23 |
| 4 | Mitophagy | 0.03349 | 0.08712 | 33.04 | 112.22 |
| 5 | Central carbon metabolism in cancer | 0.03446 | 0.08712 | 32.08 | 108.04 |

GO: Biological Process

| **Index** | **Name** | **P-value** | **Adjusted p-value** | **Odds Ratio** | **Combined score** |
| --- | --- | --- | --- | --- | --- |
| 1 | glial cell-derived neurotrophic factor receptor signaling pathway (GO:0035860) | 0.002498 | 0.04559 | 555.17 | 3326.77 |
| 2 | positive regulation of vascular associated smooth muscle cell apoptotic process (GO:1905461) | 0.002498 | 0.04559 | 555.17 | 3326.77 |
| 3 | Schwann cell differentiation (GO:0014037) | 0.002498 | 0.04559 | 555.17 | 3326.77 |
| 4 | neuromuscular synaptic transmission (GO:0007274) | 0.00004255 | 0.009105 | 277.39 | 2791.90 |
| 5 | astrocyte development (GO:0014002) | 0.002997 | 0.04559 | 444.11 | 2580.41 |

GO: Molecular Function

| **Index** | **Name** | **P-value** | **Adjusted p-value** | **Odds Ratio** | **Combined score** |
| --- | --- | --- | --- | --- | --- |
| 1 | inositol 1,3,4,5 tetrakisphosphate binding (GO:0043533) | 0.002498 | 0.04791 | 555.17 | 3326.77 |
| 2 | O-acetyltransferase activity (GO:0016413) | 0.003994 | 0.04791 | 317.19 | 1751.86 |
| 3 | phosphatidylethanolamine binding (GO:0008429) | 0.004492 | 0.04791 | 277.53 | 1500.17 |
| 4 | organic cation transmembrane transporter activity (GO:0015101) | 0.007973 | 0.06378 | 147.96 | 714.91 |
| 5 | kinesin binding (GO:0019894) | 0.01441 | 0.07433 | 79.21 | 335.86 |

GO: Cellular Component

| **Index** | **Name** | **P-value** | **Adjusted p-value** | **Odds Ratio** | **Combined score** |
| --- | --- | --- | --- | --- | --- |
| 1 | clathrin coat of endocytic vesicle (GO:0030128) | 0.003495 | 0.01592 | 370.07 | 2093.27 |
| 2 | AP-1 adaptor complex (GO:0030121) | 0.003495 | 0.01592 | 370.07 | 2093.27 |
| 3 | AP-2 adaptor complex (GO:0030122) | 0.003495 | 0.01592 | 370.07 | 2093.27 |
| 4 | clathrin coat of trans-Golgi network vesicle (GO:0030130) | 0.004492 | 0.01842 | 277.53 | 1500.17 |
| 5 | clathrin coat of coated pit (GO:0030132) | 0.004990 | 0.01860 | 246.68 | 1307.49 |

Nasal congestion ([HP:0001742](https://hpo.jax.org/app/browse/term/HP:0001742))

KEGG Pathway

| **Index** | **Name** | **P-value** | **Adjusted p-value** | **Odds Ratio** | **Combined score** |
| --- | --- | --- | --- | --- | --- |
| 1 | Huntington disease | 0.1429 | 0.2137 | 7.17 | 13.95 |
| 2 | Amyotrophic lateral sclerosis | 0.1678 | 0.2137 | 6.01 | 10.72 |
| 3 | Pathways of neurodegeneration | 0.2137 | 0.2137 | 4.57 | 7.06 |

GO: Biological Process

| **Index** | **Name** | **P-value** | **Adjusted p-value** | **Odds Ratio** | **Combined score** |
| --- | --- | --- | --- | --- | --- |
| 1 | axoneme assembly (GO:0035082) | 3.163e-15 | 1.170e-13 | 1069.39 | 35704.16 |
| 2 | regulation of cilium movement (GO:0003352) | 1.078e-8 | 9.972e-8 | 1223.45 | 22444.87 |
| 3 | axonemal dynein complex assembly (GO:0070286) | 1.893e-12 | 3.501e-11 | 739.37 | 19957.87 |
| 4 | determination of digestive tract left/right asymmetry (GO:0071907) | 0.000004719 | 0.00001940 | 999.25 | 12254.80 |
| 5 | epithelial cilium movement involved in determination of left/right asymmetry (GO:0060287) | 0.000008085 | 0.00002991 | 713.68 | 8368.26 |

GO: Molecular Function

| **Index** | **Name** | **P-value** | **Adjusted p-value** | **Odds Ratio** | **Combined score** |
| --- | --- | --- | --- | --- | --- |
| 1 | microtubule motor activity (GO:0003777) | 0.02766 | 0.04878 | 40.27 | 144.50 |
| 2 | motor activity (GO:0003774) | 0.03252 | 0.04878 | 34.06 | 116.68 |
| 3 | protein homodimerization activity (GO:0042803) | 0.2762 | 0.2762 | 3.39 | 4.36 |

GO: Cellular Component

| **Index** | **Name** | **P-value** | **Adjusted p-value** | **Odds Ratio** | **Combined score** |
| --- | --- | --- | --- | --- | --- |
| 1 | outer dynein arm (GO:0036157) | 0.004492 | 0.01497 | 277.53 | 1500.17 |
| 2 | motile cilium (GO:0031514) | 0.0005036 | 0.002518 | 75.47 | 573.09 |
| 3 | cilium (GO:0005929) | 0.0001854 | 0.001854 | 36.18 | 310.92 |
| 4 | sperm flagellum (GO:0036126) | 0.02570 | 0.05629 | 43.44 | 159.04 |
| 5 | 9+2 motile cilium (GO:0097729) | 0.02814 | 0.05629 | 39.55 | 141.22 |

Rhinitis ([HP:0012384](https://hpo.jax.org/app/browse/term/HP:0012384))

KEGG Pathway

| **Index** | **Name** | **P-value** | **Adjusted p-value** | **Odds Ratio** | **Combined score** |
| --- | --- | --- | --- | --- | --- |
| 1 | Huntington disease | 0.1429 | 0.2137 | 7.17 | 13.95 |
| 2 | Amyotrophic lateral sclerosis | 0.1678 | 0.2137 | 6.01 | 10.72 |
| 3 | Pathways of neurodegeneration | 0.2137 | 0.2137 | 4.57 | 7.06 |

GO: Biological Process

| **Index** | **Name** | **P-value** | **Adjusted p-value** | **Odds Ratio** | **Combined score** |
| --- | --- | --- | --- | --- | --- |
| 1 | axoneme assembly (GO:0035082) | 3.163e-15 | 1.170e-13 | 1069.39 | 35704.16 |
| 2 | regulation of cilium movement (GO:0003352) | 1.078e-8 | 9.972e-8 | 1223.45 | 22444.87 |
| 3 | axonemal dynein complex assembly (GO:0070286) | 1.893e-12 | 3.501e-11 | 739.37 | 19957.87 |
| 4 | determination of digestive tract left/right asymmetry (GO:0071907) | 0.000004719 | 0.00001940 | 999.25 | 12254.80 |
| 5 | epithelial cilium movement involved in determination of left/right asymmetry (GO:0060287) | 0.000008085 | 0.00002991 | 713.68 | 8368.26 |

GO: Molecular Function

| **Index** | **Name** | **P-value** | **Adjusted p-value** | **Odds Ratio** | **Combined score** |
| --- | --- | --- | --- | --- | --- |
| 1 | microtubule motor activity (GO:0003777) | 0.02766 | 0.04878 | 40.27 | 144.50 |
| 2 | motor activity (GO:0003774) | 0.03252 | 0.04878 | 34.06 | 116.68 |
| 3 | protein homodimerization activity (GO:0042803) | 0.2762 | 0.2762 | 3.39 | 4.36 |

GO: Cellular Component

| **Index** | **Name** | **P-value** | **Adjusted p-value** | **Odds Ratio** | **Combined score** |
| --- | --- | --- | --- | --- | --- |
| 1 | outer dynein arm (GO:0036157) | 0.004492 | 0.01497 | 277.53 | 1500.17 |
| 2 | motile cilium (GO:0031514) | 0.0005036 | 0.002518 | 75.47 | 573.09 |
| 3 | cilium (GO:0005929) | 0.0001854 | 0.001854 | 36.18 | 310.92 |
| 4 | sperm flagellum (GO:0036126) | 0.02570 | 0.05629 | 43.44 | 159.04 |
| 5 | 9+2 motile cilium (GO:0097729) | 0.02814 | 0.05629 | 39.55 | 141.22 |

HEENT-eye

Blindness ([HP:0000618](https://hpo.jax.org/app/browse/term/HP:0000618))

KEGG Pathway

| **Index** | **Name** | **P-value** | **Adjusted p-value** | **Odds Ratio** | **Combined score** |
| --- | --- | --- | --- | --- | --- |
| 1 | Phototransduction | 0.00008446 | 0.0005068 | 191.96 | 1800.45 |
| 2 | Retinol metabolism | 0.03349 | 0.09401 | 33.04 | 112.22 |
| 3 | Drug metabolism | 0.05272 | 0.09401 | 20.65 | 60.76 |
| 4 | Purine metabolism | 0.06267 | 0.09401 | 17.24 | 47.76 |
| 5 | cGMP-PKG signaling pathway | 0.08045 | 0.09654 | 13.27 | 33.44 |

GO: Biological Process

| **Index** | **Name** | **P-value** | **Adjusted p-value** | **Odds Ratio** | **Combined score** |
| --- | --- | --- | --- | --- | --- |
| 1 | protein localization to ciliary transition zone (GO:1904491) | 0.002997 | 0.02240 | 444.11 | 2580.41 |
| 2 | protein localization to non-motile cilium (GO:0097499) | 0.002997 | 0.02240 | 444.11 | 2580.41 |
| 3 | visual perception (GO:0007601) | 9.004e-8 | 0.000002895 | 149.07 | 2418.38 |
| 4 | sensory perception of light stimulus (GO:0050953) | 9.813e-8 | 0.000002895 | 145.78 | 2352.45 |
| 5 | regulation of rhodopsin mediated signaling pathway (GO:0022400) | 0.00006709 | 0.001319 | 217.03 | 2085.58 |

GO: Molecular Function

| **Index** | **Name** | **P-value** | **Adjusted p-value** | **Odds Ratio** | **Combined score** |
| --- | --- | --- | --- | --- | --- |
| 1 | guanylate cyclase activator activity (GO:0030250) | 0.002498 | 0.01921 | 555.17 | 3326.77 |
| 2 | NADP-retinol dehydrogenase activity (GO:0052650) | 0.002997 | 0.01921 | 444.11 | 2580.41 |
| 3 | intracellular cGMP-activated cation channel activity (GO:0005223) | 0.003495 | 0.01921 | 370.07 | 2093.27 |
| 4 | intracellular cAMP-activated cation channel activity (GO:0005222) | 0.004492 | 0.01921 | 277.53 | 1500.17 |
| 5 | leucine zipper domain binding (GO:0043522) | 0.004990 | 0.01921 | 246.68 | 1307.49 |

GO: Cellular Component

| **Index** | **Name** | **P-value** | **Adjusted p-value** | **Odds Ratio** | **Combined score** |
| --- | --- | --- | --- | --- | --- |
| 1 | sperm flagellum (GO:0036126) | 0.02570 | 0.1766 | 43.44 | 159.04 |
| 2 | 9+2 motile cilium (GO:0097729) | 0.02814 | 0.1766 | 39.55 | 141.22 |
| 3 | azurophil granule lumen (GO:0035578) | 0.04411 | 0.1766 | 24.85 | 77.54 |
| 4 | cytoplasmic vesicle lumen (GO:0060205) | 0.05605 | 0.1766 | 19.37 | 55.82 |
| 5 | ficolin-1-rich granule lumen (GO:1904813) | 0.05984 | 0.1766 | 18.09 | 50.96 |

Blurred vision ([HP:0000622](https://hpo.jax.org/app/browse/term/HP:0000622))

KEGG Pathway

| **Index** | **Name** | **P-value** | **Adjusted p-value** | **Odds Ratio** | **Combined score** |
| --- | --- | --- | --- | --- | --- |
| 1 | Proximal tubule bicarbonate reclamation | 0.01144 | 0.06159 | 100.85 | 450.83 |
| 2 | Collecting duct acid secretion | 0.01342 | 0.06159 | 85.32 | 367.79 |
| 3 | Protein digestion and absorption | 0.001151 | 0.02416 | 49.23 | 333.17 |
| 4 | Aldosterone-regulated sodium reabsorption | 0.01835 | 0.06159 | 61.59 | 246.23 |
| 5 | Carbohydrate digestion and absorption | 0.02326 | 0.06159 | 48.17 | 181.19 |

GO: Biological Process

| **Index** | **Name** | **P-value** | **Adjusted p-value** | **Odds Ratio** | **Combined score** |
| --- | --- | --- | --- | --- | --- |
| 1 | neuronal action potential (GO:0019228) | 1.379e-7 | 0.00001605 | 450.47 | 7115.97 |
| 2 | membrane depolarization during action potential (GO:0086010) | 2.326e-7 | 0.00001605 | 372.06 | 5682.76 |
| 3 | regulation of muscle system process (GO:0090257) | 0.00001481 | 0.0002555 | 499.50 | 5554.53 |
| 4 | regulation of respiratory gaseous exchange by nervous system process (GO:0002087) | 0.002498 | 0.01929 | 555.17 | 3326.77 |
| 5 | negative regulation of keratinocyte differentiation (GO:0045617) | 0.002498 | 0.01929 | 555.17 | 3326.77 |

GO: Molecular Function

| **Index** | **Name** | **P-value** | **Adjusted p-value** | **Odds Ratio** | **Combined score** |
| --- | --- | --- | --- | --- | --- |
| 1 | intronic transcription regulatory region sequence-specific DNA binding (GO:0001161) | 0.002498 | 0.01946 | 555.17 | 3326.77 |
| 2 | P-type sodium transporter activity (GO:0008554) | 0.002997 | 0.01946 | 444.11 | 2580.41 |
| 3 | P-type sodium:potassium-exchanging transporter activity (GO:0005391) | 0.002997 | 0.01946 | 444.11 | 2580.41 |
| 4 | sodium ion binding (GO:0031402) | 0.002997 | 0.01946 | 444.11 | 2580.41 |
| 5 | voltage-gated sodium channel activity (GO:0005248) | 0.00005170 | 0.002016 | 249.63 | 2463.82 |

GO: Cellular Component

| **Index** | **Name** | **P-value** | **Adjusted p-value** | **Odds Ratio** | **Combined score** |
| --- | --- | --- | --- | --- | --- |
| 1 | voltage-gated sodium channel complex (GO:0001518) | 0.00003048 | 0.0007010 | 332.92 | 3461.84 |
| 2 | sodium channel complex (GO:0034706) | 0.00006709 | 0.0007715 | 217.03 | 2085.58 |
| 3 | sodium:potassium-exchanging ATPase complex (GO:0005890) | 0.004990 | 0.01913 | 246.68 | 1307.49 |
| 4 | cation-transporting ATPase complex (GO:0090533) | 0.007973 | 0.02620 | 147.96 | 714.91 |
| 5 | basement membrane (GO:0005604) | 0.02570 | 0.06697 | 43.44 | 159.04 |

Conjunctivitis ([HP:0000509](https://hpo.jax.org/app/browse/term/HP:0000509))

KEGG Pathway

| **Index** | **Name** | **P-value** | **Adjusted p-value** | **Odds Ratio** | **Combined score** |
| --- | --- | --- | --- | --- | --- |
| 1 | Primary immunodeficiency | 5.941e-18 | 8.317e-17 | 1502.29 | 59587.92 |
| 2 | B cell receptor signaling pathway | 2.384e-10 | 1.669e-9 | 262.03 | 5805.76 |
| 3 | Intestinal immune network for IgA production | 0.0002507 | 0.0008775 | 108.39 | 898.70 |
| 4 | Epstein-Barr virus infection | 0.0001156 | 0.0005395 | 42.62 | 386.39 |
| 5 | Hematopoietic cell lineage | 0.001064 | 0.002736 | 51.27 | 351.00 |

GO: Biological Process

| **Index** | **Name** | **P-value** | **Adjusted p-value** | **Odds Ratio** | **Combined score** |
| --- | --- | --- | --- | --- | --- |
| 1 | lymphocyte differentiation (GO:0030098) | 9.479e-13 | 4.589e-11 | 382.92 | 10601.05 |
| 2 | B cell activation (GO:0042113) | 1.020e-12 | 4.589e-11 | 378.06 | 10438.72 |
| 3 | B cell receptor signaling pathway (GO:0050853) | 1.451e-9 | 3.264e-8 | 443.56 | 9026.96 |
| 4 | B cell differentiation (GO:0030183) | 5.558e-11 | 1.667e-9 | 355.96 | 8405.46 |
| 5 | V(D)J recombination (GO:0033151) | 0.00001235 | 0.0001587 | 555.03 | 6273.06 |

GO: Molecular Function

| **Index** | **Name** | **P-value** | **Adjusted p-value** | **Odds Ratio** | **Combined score** |
| --- | --- | --- | --- | --- | --- |
| 1 | phosphatidylinositol-3,4,5-trisphosphate binding (GO:0005547) | 0.0001327 | 0.004379 | 151.19 | 1349.73 |
| 2 | phospholipase activator activity (GO:0016004) | 0.005488 | 0.03290 | 222.00 | 1155.57 |
| 3 | complement receptor activity (GO:0004875) | 0.005488 | 0.03290 | 222.00 | 1155.57 |
| 4 | lipase activator activity (GO:0060229) | 0.005985 | 0.03290 | 201.81 | 1032.95 |
| 5 | phospholipase binding (GO:0043274) | 0.006979 | 0.03290 | 170.74 | 847.70 |

GO: Cellular Component

| **Index** | **Name** | **P-value** | **Adjusted p-value** | **Odds Ratio** | **Combined score** |
| --- | --- | --- | --- | --- | --- |
| 1 | pericentric heterochromatin (GO:0005721) | 0.006979 | 0.02443 | 170.74 | 847.70 |
| 2 | multivesicular body (GO:0005771) | 0.02326 | 0.05427 | 48.17 | 181.19 |
| 3 | membrane raft (GO:0045121) | 0.002846 | 0.01992 | 30.79 | 180.49 |
| 4 | late endosome (GO:0005770) | 0.09060 | 0.1586 | 11.70 | 28.10 |
| 5 | integral component of plasma membrane (GO:0005887) | 0.1613 | 0.2258 | 3.19 | 5.82 |

Diplopia ([HP:0000651](https://hpo.jax.org/app/browse/term/HP:0000651))

KEGG Pathway

| **Index** | **Name** | **P-value** | **Adjusted p-value** | **Odds Ratio** | **Combined score** |
| --- | --- | --- | --- | --- | --- |
| 1 | Neuroactive ligand-receptor interaction | 0.00001608 | 0.00004823 | 38.88 | 429.14 |
| 2 | Glycerophospholipid metabolism | 0.04794 | 0.05510 | 22.79 | 69.22 |
| 3 | Cholinergic synapse | 0.05510 | 0.05510 | 19.72 | 57.16 |

GO: Biological Process

| **Index** | **Name** | **P-value** | **Adjusted p-value** | **Odds Ratio** | **Combined score** |
| --- | --- | --- | --- | --- | --- |
| 1 | neuromuscular process (GO:0050905) | 8.685e-8 | 0.000001307 | 535.02 | 8698.92 |
| 2 | synaptic transmission, cholinergic (GO:0007271) | 8.685e-8 | 0.000001307 | 535.02 | 8698.92 |
| 3 | neuromuscular synaptic transmission (GO:0007274) | 1.021e-7 | 0.000001307 | 503.52 | 8105.11 |
| 4 | neuronal action potential (GO:0019228) | 1.379e-7 | 0.000001471 | 450.47 | 7115.97 |
| 5 | neuromuscular junction development (GO:0007528) | 1.812e-7 | 0.000001656 | 407.53 | 6326.45 |

GO: Molecular Function

| **Index** | **Name** | **P-value** | **Adjusted p-value** | **Odds Ratio** | **Combined score** |
| --- | --- | --- | --- | --- | --- |
| 1 | neurotransmitter receptor activity involved in regulation of postsynaptic membrane potential (GO:0099529) | 1.878e-10 | 5.821e-9 | 783.25 | 17541.62 |
| 2 | transmitter-gated ion channel activity involved in regulation of postsynaptic membrane potential (GO:1904315) | 8.580e-10 | 1.330e-8 | 511.90 | 10686.61 |
| 3 | transmitter-gated ion channel activity (GO:0022824) | 1.451e-9 | 1.499e-8 | 443.56 | 9026.96 |
| 4 | postsynaptic neurotransmitter receptor activity (GO:0098960) | 1.191e-7 | 8.551e-7 | 475.52 | 7581.26 |
| 5 | acetylcholine receptor activity (GO:0015464) | 1.379e-7 | 8.551e-7 | 450.47 | 7115.97 |

GO: Cellular Component

| **Index** | **Name** | **P-value** | **Adjusted p-value** | **Odds Ratio** | **Combined score** |
| --- | --- | --- | --- | --- | --- |
| 1 | neuromuscular junction (GO:0031594) | 4.712e-12 | 6.125e-11 | 604.76 | 15772.67 |
| 2 | acetylcholine-gated channel complex (GO:0005892) | 3.267e-8 | 1.416e-7 | 778.40 | 13417.28 |
| 3 | ion channel complex (GO:0034702) | 5.841e-7 | 0.000001898 | 267.29 | 3836.51 |
| 4 | neuron projection (GO:0043005) | 1.379e-9 | 8.967e-9 | 82.63 | 1685.72 |
| 5 | voltage-gated sodium channel complex (GO:0001518) | 0.008469 | 0.01835 | 138.71 | 661.82 |

Gaze-evoked nystagmus ([HP:0000640](https://hpo.jax.org/app/browse/term/HP:0000640))

KEGG Pathway

| **Index** | **Name** | **P-value** | **Adjusted p-value** | **Odds Ratio** | **Combined score** |
| --- | --- | --- | --- | --- | --- |
| 1 | Spinocerebellar ataxia | 2.465e-11 | 1.874e-9 | 217.37 | 5309.47 |
| 2 | Aldosterone-regulated sodium reabsorption | 0.0001485 | 0.002821 | 142.54 | 1256.48 |
| 3 | Aldosterone synthesis and secretion | 0.00001335 | 0.0005074 | 89.75 | 1007.36 |
| 4 | Endocrine and other factor-regulated calcium reabsorption | 0.0003059 | 0.004649 | 97.74 | 790.95 |
| 5 | GnRH secretion | 0.0004462 | 0.005651 | 80.35 | 619.92 |

GO: Biological Process

| **Index** | **Name** | **P-value** | **Adjusted p-value** | **Odds Ratio** | **Combined score** |
| --- | --- | --- | --- | --- | --- |
| 1 | membrane depolarization during action potential (GO:0086010) | 4.685e-10 | 7.550e-8 | 605.09 | 12998.28 |
| 2 | cardiac muscle cell action potential (GO:0086001) | 8.580e-10 | 7.550e-8 | 511.90 | 10686.61 |
| 3 | neuronal action potential (GO:0019228) | 1.379e-7 | 0.000006068 | 450.47 | 7115.97 |
| 4 | membrane depolarization (GO:0051899) | 4.430e-7 | 0.00001559 | 294.99 | 4315.59 |
| 5 | membrane repolarization (GO:0086009) | 0.00002690 | 0.0004734 | 356.71 | 3753.85 |

GO: Molecular Function

| **Index** | **Name** | **P-value** | **Adjusted p-value** | **Odds Ratio** | **Combined score** |
| --- | --- | --- | --- | --- | --- |
| 1 | voltage-gated sodium channel activity (GO:0005248) | 1.379e-7 | 0.000005517 | 450.47 | 7115.97 |
| 2 | sodium channel activity (GO:0005272) | 7.523e-7 | 0.00001505 | 244.35 | 3445.31 |
| 3 | voltage-gated calcium channel activity involved in cardiac muscle cell action potential (GO:0086007) | 0.002498 | 0.01869 | 555.17 | 3326.77 |
| 4 | P-type sodium transporter activity (GO:0008554) | 0.002997 | 0.01869 | 444.11 | 2580.41 |
| 5 | P-type sodium:potassium-exchanging transporter activity (GO:0005391) | 0.002997 | 0.01869 | 444.11 | 2580.41 |

GO: Cellular Component

| **Index** | **Name** | **P-value** | **Adjusted p-value** | **Odds Ratio** | **Combined score** |
| --- | --- | --- | --- | --- | --- |
| 1 | voltage-gated sodium channel complex (GO:0001518) | 6.098e-8 | 0.000001829 | 611.51 | 10158.88 |
| 2 | nuclear inclusion body (GO:0042405) | 0.00001235 | 0.00009259 | 555.03 | 6273.06 |
| 3 | sodium channel complex (GO:0034706) | 2.058e-7 | 0.000003087 | 388.99 | 5988.96 |
| 4 | sodium:potassium-exchanging ATPase complex (GO:0005890) | 0.004990 | 0.02138 | 246.68 | 1307.49 |
| 5 | cation-transporting ATPase complex (GO:0090533) | 0.007973 | 0.02658 | 147.96 | 714.91 |

Keratoconjunctivitis sicca ([HP:0001097](https://hpo.jax.org/app/browse/term/HP:0001097))

KEGG Pathway

| **Index** | **Name** | **P-value** | **Adjusted p-value** | **Odds Ratio** | **Combined score** |
| --- | --- | --- | --- | --- | --- |
| 1 | Nucleotide excision repair | 0.000001443 | 0.00004472 | 194.28 | 2612.86 |
| 2 | Basal transcription factors | 0.0002202 | 0.003413 | 115.97 | 976.58 |
| 3 | Asthma | 0.01540 | 0.08063 | 73.93 | 308.54 |
| 4 | Allograft rejection | 0.01884 | 0.08063 | 59.92 | 237.98 |
| 5 | Graft-versus-host disease | 0.02081 | 0.08063 | 54.06 | 209.35 |

GO: Biological Process

| **Index** | **Name** | **P-value** | **Adjusted p-value** | **Odds Ratio** | **Combined score** |
| --- | --- | --- | --- | --- | --- |
| 1 | positive regulation of CD4-positive, CD25-positive, alpha-beta regulatory T cell differentiation (GO:0032831) | 0.002498 | 0.01599 | 555.17 | 3326.77 |
| 2 | nucleotide-excision repair, DNA incision, 3'-to lesion (GO:0006295) | 0.00004701 | 0.001831 | 262.78 | 2618.60 |
| 3 | nucleotide-excision repair, preincision complex stabilization (GO:0006293) | 0.00004701 | 0.001831 | 262.78 | 2618.60 |
| 4 | regulation of T-helper cell differentiation (GO:0045622) | 0.002997 | 0.01599 | 444.11 | 2580.41 |
| 5 | positive regulation of T cell mediated immune response to tumor cell (GO:0002842) | 0.002997 | 0.01599 | 444.11 | 2580.41 |

GO: Molecular Function

| **Index** | **Name** | **P-value** | **Adjusted p-value** | **Odds Ratio** | **Combined score** |
| --- | --- | --- | --- | --- | --- |
| 1 | gap junction channel activity involved in cell communication by electrical coupling (GO:1903763) | 0.002997 | 0.02797 | 444.11 | 2580.41 |
| 2 | 5'-3' DNA helicase activity (GO:0043139) | 0.003495 | 0.02797 | 370.07 | 2093.27 |
| 3 | CD4 receptor binding (GO:0042609) | 0.003994 | 0.02797 | 317.19 | 1751.86 |
| 4 | MHC class II receptor activity (GO:0032395) | 0.004990 | 0.02797 | 246.68 | 1307.49 |
| 5 | MHC class II protein complex binding (GO:0023026) | 0.008469 | 0.02797 | 138.71 | 661.82 |

GO: Cellular Component

| **Index** | **Name** | **P-value** | **Adjusted p-value** | **Odds Ratio** | **Combined score** |
| --- | --- | --- | --- | --- | --- |
| 1 | transcription factor TFIIH core complex (GO:0000439) | 0.00001010 | 0.0003925 | 624.44 | 7182.69 |
| 2 | transcription factor TFIIH holo complex (GO:0005675) | 0.00001481 | 0.0003925 | 499.50 | 5554.53 |
| 3 | hemidesmosome (GO:0030056) | 0.003495 | 0.04631 | 370.07 | 2093.27 |
| 4 | transcription factor TFIID complex (GO:0005669) | 0.0001038 | 0.001834 | 172.08 | 1578.44 |
| 5 | MHC class II protein complex (GO:0042613) | 0.006482 | 0.05268 | 184.98 | 932.06 |

Ocular pain ([HP:0200026](https://hpo.jax.org/app/browse/term/HP:0200026))

KEGG Pathway

| **Index** | **Name** | **P-value** | **Adjusted p-value** | **Odds Ratio** | **Combined score** |
| --- | --- | --- | --- | --- | --- |
| 1 | Long-term depression | 0.0003921 | 0.005647 | 85.91 | 673.90 |
| 2 | GnRH secretion | 0.0004462 | 0.005647 | 80.35 | 619.92 |
| 3 | Cortisol synthesis and secretion | 0.0004602 | 0.005647 | 79.08 | 607.60 |
| 4 | Insulin secretion | 0.0008042 | 0.005647 | 59.24 | 422.16 |
| 5 | Gap junction | 0.0008418 | 0.005647 | 57.86 | 409.65 |

GO: Biological Process

| **Index** | **Name** | **P-value** | **Adjusted p-value** | **Odds Ratio** | **Combined score** |
| --- | --- | --- | --- | --- | --- |
| 1 | phototransduction, visible light (GO:0007603) | 0.00002354 | 0.0006787 | 384.17 | 4094.01 |
| 2 | detection of visible light (GO:0009584) | 0.00003048 | 0.0006787 | 332.92 | 3461.84 |
| 3 | negative regulation of keratinocyte differentiation (GO:0045617) | 0.002498 | 0.02248 | 555.17 | 3326.77 |
| 4 | protein side chain deglutamylation (GO:0035610) | 0.002498 | 0.02248 | 555.17 | 3326.77 |
| 5 | C-terminal protein deglutamylation (GO:0035609) | 0.002498 | 0.02248 | 555.17 | 3326.77 |

GO: Molecular Function

| **Index** | **Name** | **P-value** | **Adjusted p-value** | **Odds Ratio** | **Combined score** |
| --- | --- | --- | --- | --- | --- |
| 1 | G protein-coupled serotonin receptor binding (GO:0031821) | 0.000003371 | 0.00008765 | 1249.13 | 15739.23 |
| 2 | intronic transcription regulatory region sequence-specific DNA binding (GO:0001161) | 0.002498 | 0.02138 | 555.17 | 3326.77 |
| 3 | N-acetylglucosamine 6-O-sulfotransferase activity (GO:0001517) | 0.003495 | 0.02138 | 370.07 | 2093.27 |
| 4 | voltage-gated sodium channel activity (GO:0005248) | 0.01095 | 0.03558 | 105.66 | 476.99 |
| 5 | metallocarboxypeptidase activity (GO:0004181) | 0.01441 | 0.04163 | 79.21 | 335.86 |

GO: Cellular Component

| **Index** | **Name** | **P-value** | **Adjusted p-value** | **Odds Ratio** | **Combined score** |
| --- | --- | --- | --- | --- | --- |
| 1 | hemidesmosome (GO:0030056) | 0.003495 | 0.02964 | 370.07 | 2093.27 |
| 2 | heterotrimeric G-protein complex (GO:0005834) | 0.0001178 | 0.002474 | 160.96 | 1456.09 |
| 3 | voltage-gated sodium channel complex (GO:0001518) | 0.008469 | 0.02964 | 138.71 | 661.82 |
| 4 | sodium channel complex (GO:0034706) | 0.01243 | 0.03264 | 92.44 | 405.55 |
| 5 | basement membrane (GO:0005604) | 0.02570 | 0.04907 | 43.44 | 159.04 |

Peripheral visual field loss ([HP:0007994](https://hpo.jax.org/app/browse/term/HP:0007994))

KEGG Pathway

| **Index** | **Name** | **P-value** | **Adjusted p-value** | **Odds Ratio** | **Combined score** |
| --- | --- | --- | --- | --- | --- |
| 1 | Phototransduction | 6.413e-10 | 4.489e-9 | 554.61 | 11739.73 |
| 2 | Purine metabolism | 0.001796 | 0.006285 | 39.10 | 247.21 |
| 3 | ABC transporters | 0.02228 | 0.05198 | 50.37 | 191.61 |
| 4 | Retinol metabolism | 0.03349 | 0.05861 | 33.04 | 112.22 |
| 5 | cGMP-PKG signaling pathway | 0.08045 | 0.1126 | 13.27 | 33.44 |

GO: Biological Process

| **Index** | **Name** | **P-value** | **Adjusted p-value** | **Odds Ratio** | **Combined score** |
| --- | --- | --- | --- | --- | --- |
| 1 | phototransduction, visible light (GO:0007603) | 4.289e-11 | 1.930e-9 | 1210.85 | 28905.97 |
| 2 | rhodopsin mediated signaling pathway (GO:0016056) | 1.482e-8 | 2.220e-7 | 1070.46 | 19297.66 |
| 3 | regulation of rhodopsin mediated signaling pathway (GO:0022400) | 3.965e-10 | 8.921e-9 | 633.94 | 13723.68 |
| 4 | vitamin A metabolic process (GO:0006776) | 0.000004719 | 0.00002178 | 999.25 | 12254.80 |
| 5 | detection of visible light (GO:0009584) | 6.098e-8 | 4.573e-7 | 611.51 | 10158.88 |

GO: Molecular Function

| **Index** | **Name** | **P-value** | **Adjusted p-value** | **Odds Ratio** | **Combined score** |
| --- | --- | --- | --- | --- | --- |
| 1 | 11-cis retinal binding (GO:0005502) | 0.000003371 | 0.0001281 | 1249.13 | 15739.23 |
| 2 | retinal binding (GO:0016918) | 0.000008085 | 0.0001536 | 713.68 | 8368.26 |
| 3 | all-trans retinal binding (GO:0005503) | 0.002498 | 0.02029 | 555.17 | 3326.77 |
| 4 | phosphatidylethanolamine flippase activity (GO:0090555) | 0.002997 | 0.02029 | 444.11 | 2580.41 |
| 5 | intracellular cGMP-activated cation channel activity (GO:0005223) | 0.003495 | 0.02029 | 370.07 | 2093.27 |

GO: Cellular Component

| **Index** | **Name** | **P-value** | **Adjusted p-value** | **Odds Ratio** | **Combined score** |
| --- | --- | --- | --- | --- | --- |
| 1 | nuclear outer membrane (GO:0005640) | 0.007973 | 0.05235 | 147.96 | 714.91 |
| 2 | Golgi-associated vesicle membrane (GO:0030660) | 0.01194 | 0.05235 | 96.46 | 427.12 |
| 3 | ciliary membrane (GO:0060170) | 0.01540 | 0.05235 | 73.93 | 308.54 |
| 4 | Golgi-associated vesicle (GO:0005798) | 0.02326 | 0.06590 | 48.17 | 181.19 |
| 5 | cell projection membrane (GO:0031253) | 0.04507 | 0.1001 | 24.30 | 75.31 |

Photophobia ([HP:0000613](https://hpo.jax.org/app/browse/term/HP:0000613))

KEGG Pathway

| **Index** | **Name** | **P-value** | **Adjusted p-value** | **Odds Ratio** | **Combined score** |
| --- | --- | --- | --- | --- | --- |
| 1 | Phototransduction | 2.929e-7 | 0.000001465 | 342.26 | 5148.70 |
| 2 | Retinol metabolism | 0.03349 | 0.08373 | 33.04 | 112.22 |
| 3 | Purine metabolism | 0.06267 | 0.1006 | 17.24 | 47.76 |
| 4 | cGMP-PKG signaling pathway | 0.08045 | 0.1006 | 13.27 | 33.44 |
| 5 | cAMP signaling pathway | 0.1029 | 0.1029 | 10.22 | 23.24 |

GO: Biological Process

| **Index** | **Name** | **P-value** | **Adjusted p-value** | **Odds Ratio** | **Combined score** |
| --- | --- | --- | --- | --- | --- |
| 1 | rhodopsin mediated signaling pathway (GO:0016056) | 0.00001235 | 0.0001407 | 555.03 | 6273.06 |
| 2 | regulation of rhodopsin mediated signaling pathway (GO:0022400) | 2.058e-7 | 0.000003910 | 388.99 | 5988.96 |
| 3 | phototransduction, visible light (GO:0007603) | 0.00002354 | 0.0001917 | 384.17 | 4094.01 |
| 4 | eye photoreceptor cell development (GO:0042462) | 0.00002354 | 0.0001917 | 384.17 | 4094.01 |
| 5 | visual perception (GO:0007601) | 9.004e-8 | 0.000002797 | 149.07 | 2418.38 |

GO: Molecular Function

| **Index** | **Name** | **P-value** | **Adjusted p-value** | **Odds Ratio** | **Combined score** |
| --- | --- | --- | --- | --- | --- |
| 1 | leucine zipper domain binding (GO:0043522) | 0.00001010 | 0.0002829 | 624.44 | 7182.69 |
| 2 | LRR domain binding (GO:0030275) | 0.00002041 | 0.0002857 | 416.21 | 4494.85 |
| 3 | guanylate cyclase activator activity (GO:0030250) | 0.002498 | 0.01537 | 555.17 | 3326.77 |
| 4 | NADP-retinol dehydrogenase activity (GO:0052650) | 0.002997 | 0.01537 | 444.11 | 2580.41 |
| 5 | intracellular cGMP-activated cation channel activity (GO:0005223) | 0.003495 | 0.01537 | 370.07 | 2093.27 |

GO: Cellular Component

| **Index** | **Name** | **P-value** | **Adjusted p-value** | **Odds Ratio** | **Combined score** |
| --- | --- | --- | --- | --- | --- |
| 1 | sperm flagellum (GO:0036126) | 0.02570 | 0.08443 | 43.44 | 159.04 |
| 2 | 9+2 motile cilium (GO:0097729) | 0.02814 | 0.08443 | 39.55 | 141.22 |
| 3 | cilium (GO:0005929) | 0.1124 | 0.2248 | 9.30 | 20.33 |
| 4 | integral component of plasma membrane (GO:0005887) | 0.1613 | 0.2419 | 3.19 | 5.82 |
| 5 | nucleus (GO:0005634) | 0.3960 | 0.4752 | 1.48 | 1.37 |

Red eye ([HP:0025337](https://hpo.jax.org/app/browse/term/HP:0025337))

KEGG Pathway

| **Index** | **Name** | **P-value** | **Adjusted p-value** | **Odds Ratio** | **Combined score** |
| --- | --- | --- | --- | --- | --- |
| 1 | Primary immunodeficiency | 5.941e-18 | 8.317e-17 | 1502.29 | 59587.92 |
| 2 | B cell receptor signaling pathway | 2.384e-10 | 1.669e-9 | 262.03 | 5805.76 |
| 3 | Intestinal immune network for IgA production | 0.0002507 | 0.0008775 | 108.39 | 898.70 |
| 4 | Epstein-Barr virus infection | 0.0001156 | 0.0005395 | 42.62 | 386.39 |
| 5 | Hematopoietic cell lineage | 0.001064 | 0.002736 | 51.27 | 351.00 |

GO: Biological Process

| **Index** | **Name** | **P-value** | **Adjusted p-value** | **Odds Ratio** | **Combined score** |
| --- | --- | --- | --- | --- | --- |
| 1 | lymphocyte differentiation (GO:0030098) | 9.479e-13 | 4.589e-11 | 382.92 | 10601.05 |
| 2 | B cell activation (GO:0042113) | 1.020e-12 | 4.589e-11 | 378.06 | 10438.72 |
| 3 | B cell receptor signaling pathway (GO:0050853) | 1.451e-9 | 3.264e-8 | 443.56 | 9026.96 |
| 4 | B cell differentiation (GO:0030183) | 5.558e-11 | 1.667e-9 | 355.96 | 8405.46 |
| 5 | V(D)J recombination (GO:0033151) | 0.00001235 | 0.0001587 | 555.03 | 6273.06 |

GO: Molecular Function

| **Index** | **Name** | **P-value** | **Adjusted p-value** | **Odds Ratio** | **Combined score** |
| --- | --- | --- | --- | --- | --- |
| 1 | phosphatidylinositol-3,4,5-trisphosphate binding (GO:0005547) | 0.0001327 | 0.004379 | 151.19 | 1349.73 |
| 2 | phospholipase activator activity (GO:0016004) | 0.005488 | 0.03290 | 222.00 | 1155.57 |
| 3 | complement receptor activity (GO:0004875) | 0.005488 | 0.03290 | 222.00 | 1155.57 |
| 4 | lipase activator activity (GO:0060229) | 0.005985 | 0.03290 | 201.81 | 1032.95 |
| 5 | phospholipase binding (GO:0043274) | 0.006979 | 0.03290 | 170.74 | 847.70 |

GO: Cellular Component

| **Index** | **Name** | **P-value** | **Adjusted p-value** | **Odds Ratio** | **Combined score** |
| --- | --- | --- | --- | --- | --- |
| 1 | pericentric heterochromatin (GO:0005721) | 0.006979 | 0.02443 | 170.74 | 847.70 |
| 2 | multivesicular body (GO:0005771) | 0.02326 | 0.05427 | 48.17 | 181.19 |
| 3 | membrane raft (GO:0045121) | 0.002846 | 0.01992 | 30.79 | 180.49 |
| 4 | late endosome (GO:0005770) | 0.09060 | 0.1586 | 11.70 | 28.10 |
| 5 | integral component of plasma membrane (GO:0005887) | 0.1613 | 0.2258 | 3.19 | 5.82 |

Visual loss ([HP:0000572](https://hpo.jax.org/app/browse/term/HP:0000572))

GO: Biological Process

| **Index** | **Name** | **P-value** | **Adjusted p-value** | **Odds Ratio** | **Combined score** |
| --- | --- | --- | --- | --- | --- |
| 1 | equilibrioception (GO:0050957) | 5.668e-17 | 3.287e-15 | 19989.00 | 747771.16 |
| 2 | neuromuscular process controlling balance (GO:0050885) | 1.888e-14 | 2.738e-13 | 2220.11 | 70156.55 |
| 3 | sensory perception (GO:0007600) | 4.223e-14 | 4.899e-13 | 664.83 | 20473.93 |
| 4 | sensory perception of mechanical stimulus (GO:0050954) | 2.969e-15 | 7.316e-14 | 573.51 | 19184.17 |
| 5 | sensory perception of sound (GO:0007605) | 3.784e-15 | 7.316e-14 | 552.94 | 18362.12 |

GO: Molecular Function

| **Index** | **Name** | **P-value** | **Adjusted p-value** | **Odds Ratio** | **Combined score** |
| --- | --- | --- | --- | --- | --- |
| 1 | myosin binding (GO:0017022) | 0.02766 | 0.05531 | 40.27 | 144.50 |
| 2 | calcium ion binding (GO:0005509) | 0.01239 | 0.05531 | 14.19 | 62.33 |
| 3 | magnesium ion binding (GO:0000287) | 0.07066 | 0.1060 | 15.21 | 40.30 |
| 4 | metal ion binding (GO:0046872) | 0.02616 | 0.05531 | 9.45 | 34.45 |
| 5 | cadherin binding (GO:0045296) | 0.1499 | 0.1798 | 6.81 | 12.92 |

GO: Cellular Component

| **Index** | **Name** | **P-value** | **Adjusted p-value** | **Odds Ratio** | **Combined score** |
| --- | --- | --- | --- | --- | --- |
| 1 | microvillus (GO:0005902) | 0.0003539 | 0.003893 | 90.61 | 720.07 |
| 2 | actin-based cell projection (GO:0098858) | 0.0007493 | 0.004121 | 61.45 | 442.20 |
| 3 | catenin complex (GO:0016342) | 0.01540 | 0.05645 | 73.93 | 308.54 |
| 4 | basement membrane (GO:0005604) | 0.02570 | 0.07068 | 43.44 | 159.04 |
| 5 | cilium (GO:0005929) | 0.1124 | 0.2473 | 9.30 | 20.33 |

Vitreous floaters ([HP:0100832](https://hpo.jax.org/app/browse/term/HP:0100832))

KEGG Pathway

| **Index** | **Name** | **P-value** | **Adjusted p-value** | **Odds Ratio** | **Combined score** |
| --- | --- | --- | --- | --- | --- |
| 1 | Thyroid cancer | 0.007380 | 0.04977 | 184.81 | 907.25 |
| 2 | Endometrial cancer | 0.01155 | 0.04977 | 116.60 | 520.17 |
| 3 | Basal cell carcinoma | 0.01254 | 0.04977 | 107.17 | 469.28 |
| 4 | Adherens junction | 0.01413 | 0.04977 | 94.89 | 404.19 |
| 5 | Arrhythmogenic right ventricular cardiomyopathy | 0.01531 | 0.04977 | 87.37 | 365.12 |

GO: Biological Process

| **Index** | **Name** | **P-value** | **Adjusted p-value** | **Odds Ratio** | **Combined score** |
| --- | --- | --- | --- | --- | --- |
| 1 | cellular response to indole-3-methanol (GO:0071681) | 0.0009997 | 0.01417 | 1666.00 | 11508.86 |
| 2 | response to indole-3-methanol (GO:0071680) | 0.0009997 | 0.01417 | 1666.00 | 11508.86 |
| 3 | regulation of cellular response to vascular endothelial growth factor stimulus (GO:1902547) | 0.0009997 | 0.01417 | 1666.00 | 11508.86 |
| 4 | embryonic skeletal joint morphogenesis (GO:0060272) | 0.0009997 | 0.01417 | 1666.00 | 11508.86 |
| 5 | regulation of vasculature development (GO:1901342) | 0.00001051 | 0.001367 | 798.84 | 9157.07 |

GO: Molecular Function

| **Index** | **Name** | **P-value** | **Adjusted p-value** | **Odds Ratio** | **Combined score** |
| --- | --- | --- | --- | --- | --- |
| 1 | histone methyltransferase binding (GO:1990226) | 0.002198 | 0.01998 | 666.20 | 4077.19 |
| 2 | I-SMAD binding (GO:0070411) | 0.002598 | 0.01998 | 555.11 | 3304.67 |
| 3 | Wnt-activated receptor activity (GO:0042813) | 0.002997 | 0.01998 | 475.76 | 2764.28 |
| 4 | frizzled binding (GO:0005109) | 0.006783 | 0.02793 | 201.65 | 1006.89 |
| 5 | estrogen receptor binding (GO:0030331) | 0.006982 | 0.02793 | 195.71 | 971.56 |

GO: Cellular Component

| **Index** | **Name** | **P-value** | **Adjusted p-value** | **Odds Ratio** | **Combined score** |
| --- | --- | --- | --- | --- | --- |
| 1 | beta-catenin-TCF complex (GO:1990907) | 0.002598 | 0.02857 | 555.11 | 3304.67 |
| 2 | catenin complex (GO:0016342) | 0.006186 | 0.03402 | 221.84 | 1128.18 |
| 3 | adherens junction (GO:0005912) | 0.02614 | 0.08212 | 50.55 | 184.21 |
| 4 | basolateral plasma membrane (GO:0016323) | 0.02986 | 0.08212 | 44.10 | 154.85 |
| 5 | cell-cell junction (GO:0005911) | 0.05311 | 0.1052 | 24.35 | 71.48 |
