## Supplementary Table 1-3 for "Long COVID: G Protein-Coupled Receptors (GPCRs) responsible for persistent post-COVID symptoms": Supplementary File 2.5-Lab.docx

Abnormal calcium-phosphate regulating hormone level ([HP:0100530](https://hpo.jax.org/app/browse/term/HP:0100530))

KEGG Pathway

| **Index** | **Name** | **P-value** | **Adjusted p-value** | **Odds Ratio** | **Combined score** |
| --- | --- | --- | --- | --- | --- |
| 1 | Parathyroid hormone synthesis, secretion and action | 3.963e-12 | 1.189e-10 | 298.35 | 7832.87 |
| 2 | Mineral absorption | 1.516e-8 | 2.274e-7 | 237.31 | 4272.71 |
| 3 | Endocrine and other factor-regulated calcium reabsorption | 0.000002081 | 0.00002081 | 170.91 | 2236.02 |
| 4 | Ascorbate and aldarate metabolism | 0.01490 | 0.07865 | 76.48 | 321.69 |
| 5 | Citrate cycle (TCA cycle) | 0.01490 | 0.07865 | 76.48 | 321.69 |

GO: Biological Process

| **Index** | **Name** | **P-value** | **Adjusted p-value** | **Odds Ratio** | **Combined score** |
| --- | --- | --- | --- | --- | --- |
| 1 | cellular trivalent inorganic anion homeostasis (GO:0072502) | 0.000003371 | 0.0003860 | 1249.13 | 15739.23 |
| 2 | cellular phosphate ion homeostasis (GO:0030643) | 0.000003371 | 0.0003860 | 1249.13 | 15739.23 |
| 3 | phosphate ion homeostasis (GO:0055062) | 0.00001010 | 0.0007712 | 624.44 | 7182.69 |
| 4 | phosphate ion transport (GO:0006817) | 0.00003048 | 0.001745 | 332.92 | 3461.84 |
| 5 | cellular response to vitamin D (GO:0071305) | 0.002498 | 0.02737 | 555.17 | 3326.77 |

GO: Molecular Function

| **ndex** | **Name** | **P-value** | **Adjusted p-value** | **Odds Ratio** | **Combined score** |
| --- | --- | --- | --- | --- | --- |
| 1 | sodium:phosphate symporter activity (GO:0005436) | 0.00001235 | 0.0004691 | 555.03 | 6273.06 |
| 2 | beta-glucosidase activity (GO:0008422) | 0.002997 | 0.02845 | 444.11 | 2580.41 |
| 3 | glucosidase activity (GO:0015926) | 0.003994 | 0.02845 | 317.19 | 1751.86 |
| 4 | solute:sodium symporter activity (GO:0015370) | 0.00009715 | 0.001846 | 178.23 | 1646.74 |
| 5 | phosphatidylethanolamine binding (GO:0008429) | 0.004492 | 0.02845 | 277.53 | 1500.17 |

GO: Cellular Component

| **Index** | **Name** | **P-value** | **Adjusted p-value** | **Odds Ratio** | **Combined score** |
| --- | --- | --- | --- | --- | --- |
| 1 | brush border membrane (GO:0031526) | 0.01835 | 0.07799 | 61.59 | 246.23 |
| 2 | axon (GO:0030424) | 0.004415 | 0.03054 | 24.49 | 132.81 |
| 3 | vesicle (GO:0031982) | 0.005389 | 0.03054 | 22.06 | 115.23 |
| 4 | integral component of plasma membrane (GO:0005887) | 0.0003728 | 0.006338 | 12.80 | 101.01 |
| 5 | cell projection membrane (GO:0031253) | 0.04507 | 0.1277 | 24.30 | 75.31 |

Abnormal circulating protein concentration ([HP:0010876](https://hpo.jax.org/app/browse/term/HP:0010876))

KEGG Pathway

| **Index** | **Name** | **P-value** | **Adjusted p-value** | **Odds Ratio** | **Combined score** |
| --- | --- | --- | --- | --- | --- |
| 1 | Hypertrophic cardiomyopathy | 1.450e-12 | 1.834e-11 | 355.46 | 9689.66 |
| 2 | Dilated cardiomyopathy | 2.157e-12 | 1.834e-11 | 331.67 | 8909.27 |
| 3 | Cardiac muscle contraction | 6.874e-8 | 3.895e-7 | 159.90 | 2637.14 |
| 4 | Viral myocarditis | 0.000003034 | 0.00001032 | 149.87 | 1904.21 |
| 5 | Adrenergic signaling in cardiomyocytes | 6.163e-7 | 0.000002619 | 90.61 | 1295.71 |

GO: Biological Process

| **Index** | **Name** | **P-value** | **Adjusted p-value** | **Odds Ratio** | **Combined score** |
| --- | --- | --- | --- | --- | --- |
| 1 | actin-myosin filament sliding (GO:0033275) | 3.458e-21 | 2.403e-19 | 2661.33 | 125385.00 |
| 2 | muscle filament sliding (GO:0030049) | 3.458e-21 | 2.403e-19 | 2661.33 | 125385.00 |
| 3 | heart contraction (GO:0060047) | 6.755e-21 | 3.130e-19 | 2419.03 | 112349.58 |
| 4 | cardiac muscle contraction (GO:0060048) | 2.534e-18 | 8.804e-17 | 1725.20 | 69899.68 |
| 5 | cardiac muscle tissue morphogenesis (GO:0055008) | 2.132e-15 | 3.704e-14 | 1151.77 | 38908.86 |

GO: Molecular Function

| **Index** | **Name** | **P-value** | **Adjusted p-value** | **Odds Ratio** | **Combined score** |
| --- | --- | --- | --- | --- | --- |
| 1 | FATZ binding (GO:0051373) | 0.000002248 | 0.00001476 | 1665.58 | 21661.60 |
| 2 | titin binding (GO:0031432) | 2.567e-8 | 6.161e-7 | 856.29 | 14966.03 |
| 3 | myosin heavy chain binding (GO:0032036) | 0.000004719 | 0.00002265 | 999.25 | 12254.80 |
| 4 | nitric-oxide synthase binding (GO:0050998) | 0.003495 | 0.01198 | 370.07 | 2093.27 |
| 5 | telethonin binding (GO:0031433) | 0.003495 | 0.01198 | 370.07 | 2093.27 |

GO: Cellular Component

| **Index** | **Name** | **P-value** | **Adjusted p-value** | **Odds Ratio** | **Combined score** |
| --- | --- | --- | --- | --- | --- |
| 1 | cardiac myofibril (GO:0097512) | 8.995e-10 | 1.169e-8 | 4283.14 | 89214.48 |
| 2 | myofibril (GO:0030016) | 7.595e-13 | 1.975e-11 | 907.64 | 25328.60 |
| 3 | muscle myosin complex (GO:0005859) | 0.00002354 | 0.0002040 | 384.17 | 4094.01 |
| 4 | myosin filament (GO:0032982) | 0.002997 | 0.01113 | 444.11 | 2580.41 |
| 5 | pseudopodium (GO:0031143) | 0.004492 | 0.01460 | 277.53 | 1500.17 |

Decreased circulating calcifediol concentration ([HP:0012053](https://hpo.jax.org/app/browse/term/HP:0012053))

KEGG Pathway

| **Index** | **Name** | **P-value** | **Adjusted p-value** | **Odds Ratio** | **Combined score** |
| --- | --- | --- | --- | --- | --- |
| 1 | Drug metabolism | 4.444e-12 | 7.111e-11 | 292.47 | 7645.02 |
| 2 | Retinol metabolism | 9.725e-11 | 7.780e-10 | 316.30 | 7291.95 |
| 3 | Linoleic acid metabolism | 3.266e-7 | 7.466e-7 | 329.08 | 4914.57 |
| 4 | Steroid hormone biosynthesis | 1.622e-8 | 6.487e-8 | 233.13 | 4181.78 |
| 5 | Metabolism of xenobiotics by cytochrome P450 | 3.973e-8 | 1.271e-7 | 184.43 | 3142.85 |

GO: Biological Process

| **Index** | **Name** | **P-value** | **Adjusted p-value** | **Odds Ratio** | **Combined score** |
| --- | --- | --- | --- | --- | --- |
| 1 | monoterpenoid metabolic process (GO:0016098) | 1.799e-9 | 6.954e-8 | 2855.29 | 57494.92 |
| 2 | terpenoid metabolic process (GO:0006721) | 3.147e-9 | 9.125e-8 | 2141.36 | 41921.23 |
| 3 | lipid hydroxylation (GO:0002933) | 0.000004719 | 0.00006082 | 999.25 | 12254.80 |
| 4 | estrogen metabolic process (GO:0008210) | 8.580e-10 | 4.976e-8 | 511.90 | 10686.61 |
| 5 | regulation of anion transport (GO:0044070) | 0.000006290 | 0.00007296 | 832.67 | 9972.50 |

GO: Molecular Function

| **Index** | **Name** | **P-value** | **Adjusted p-value** | **Odds Ratio** | **Combined score** |
| --- | --- | --- | --- | --- | --- |
| 1 | estrogen 2-hydroxylase activity (GO:0101021) | 0.000002248 | 0.00004271 | 1665.58 | 21661.60 |
| 2 | estrogen 16-alpha-hydroxylase activity (GO:0101020) | 0.000006290 | 0.00007681 | 832.67 | 9972.50 |
| 3 | retinoic acid 4-hydroxylase activity (GO:0008401) | 0.000008085 | 0.00007681 | 713.68 | 8368.26 |
| 4 | steroid hydroxylase activity (GO:0008395) | 1.841e-9 | 6.998e-8 | 415.79 | 8362.69 |
| 5 | arachidonic acid epoxygenase activity (GO:0008392) | 0.00002690 | 0.0001704 | 356.71 | 3753.85 |

GO: Cellular Component

| **Index** | **Name** | **P-value** | **Adjusted p-value** | **Odds Ratio** | **Combined score** |
| --- | --- | --- | --- | --- | --- |
| 1 | external side of apical plasma membrane (GO:0098591) | 0.002498 | 0.01249 | 555.17 | 3326.77 |
| 2 | endoplasmic reticulum membrane (GO:0005789) | 3.704e-7 | 0.000003704 | 40.97 | 606.74 |
| 3 | lipid droplet (GO:0005811) | 0.03785 | 0.09462 | 29.11 | 95.32 |
| 4 | mitochondrial outer membrane (GO:0005741) | 0.06126 | 0.1146 | 17.66 | 49.31 |
| 5 | organelle outer membrane (GO:0031968) | 0.06879 | 0.1146 | 15.64 | 41.87 |

Elevated erythrocyte sedimentation rate ([HP:0003565](https://hpo.jax.org/app/browse/term/HP:0003565))

KEGG Pathway

| **Index** | **Name** | **P-value** | **Adjusted p-value** | **Odds Ratio** | **Combined score** |
| --- | --- | --- | --- | --- | --- |
| 1 | Autoimmune thyroid disease | 0.0003059 | 0.004690 | 97.74 | 790.95 |
| 2 | Influenza A | 0.00007175 | 0.001920 | 50.26 | 479.64 |
| 3 | Antigen processing and presentation | 0.0006621 | 0.006942 | 65.51 | 479.51 |
| 4 | NOD-like receptor signaling pathway | 0.00008348 | 0.001920 | 47.70 | 447.96 |
| 5 | Th1 and Th2 cell differentiation | 0.0009195 | 0.006942 | 55.28 | 386.48 |

GO: Biological Process

| **Index** | **Name** | **P-value** | **Adjusted p-value** | **Odds Ratio** | **Combined score** |
| --- | --- | --- | --- | --- | --- |
| 1 | positive regulation of MHC class I biosynthetic process (GO:0045345) | 0.000002248 | 0.0001704 | 1665.58 | 21661.60 |
| 2 | regulation of MHC class I biosynthetic process (GO:0045343) | 0.000004719 | 0.0002186 | 999.25 | 12254.80 |
| 3 | negative regulation of interleukin-1 production (GO:0032692) | 3.628e-7 | 0.0001009 | 316.87 | 4699.01 |
| 4 | regulation of interleukin-1 production (GO:0032652) | 0.00002690 | 0.0006798 | 356.71 | 3753.85 |
| 5 | positive regulation of CD4-positive, CD25-positive, alpha-beta regulatory T cell differentiation (GO:0032831) | 0.002498 | 0.01472 | 555.17 | 3326.77 |

GO: Molecular Function

| **Index** | **Name** | **P-value** | **Adjusted p-value** | **Odds Ratio** | **Combined score** |
| --- | --- | --- | --- | --- | --- |
| 1 | non-membrane spanning protein tyrosine phosphatase activity (GO:0004726) | 0.002997 | 0.02371 | 444.11 | 2580.41 |
| 2 | CD4 receptor binding (GO:0042609) | 0.003994 | 0.02371 | 317.19 | 1751.86 |
| 3 | MHC class II receptor activity (GO:0032395) | 0.004990 | 0.02371 | 246.68 | 1307.49 |
| 4 | phosphatidate phosphatase activity (GO:0008195) | 0.006482 | 0.02371 | 184.98 | 932.06 |
| 5 | peptidoglycan binding (GO:0042834) | 0.007476 | 0.02371 | 158.54 | 776.21 |

GO: Cellular Component

| **Index** | **Name** | **P-value** | **Adjusted p-value** | **Odds Ratio** | **Combined score** |
| --- | --- | --- | --- | --- | --- |
| 1 | MHC class II protein complex (GO:0042613) | 0.006482 | 0.07347 | 184.98 | 932.06 |
| 2 | MHC protein complex (GO:0042611) | 0.009957 | 0.07701 | 116.79 | 538.34 |
| 3 | clathrin-coated endocytic vesicle (GO:0045334) | 0.0007857 | 0.02671 | 59.96 | 428.66 |
| 4 | lumenal side of endoplasmic reticulum membrane (GO:0098553) | 0.01392 | 0.07701 | 82.15 | 351.18 |
| 5 | integral component of lumenal side of endoplasmic reticulum membrane (GO:0071556) | 0.01392 | 0.07701 | 82.15 | 351.18 |

Elevated circulating alkaline phosphatase concentration ([HP:0003155](https://hpo.jax.org/app/browse/term/HP:0003155))

KEGG Pathway

| **Index** | **Name** | **P-value** | **Adjusted p-value** | **Odds Ratio** | **Combined score** |
| --- | --- | --- | --- | --- | --- |
| 1 | Glycosylphosphatidylinositol (GPI)-anchor biosynthesis | 3.101e-19 | 3.721e-18 | 2452.58 | 104522.58 |
| 2 | Melanoma | 0.03543 | 0.1517 | 31.17 | 104.12 |
| 3 | Parathyroid hormone synthesis, secretion and action | 0.05176 | 0.1517 | 21.04 | 62.31 |
| 4 | Breast cancer | 0.07113 | 0.1517 | 15.10 | 39.92 |
| 5 | Gastric cancer | 0.07207 | 0.1517 | 14.90 | 39.18 |

GO: Biological Process

| **Index** | **Name** | **P-value** | **Adjusted p-value** | **Odds Ratio** | **Combined score** |
| --- | --- | --- | --- | --- | --- |
| 1 | preassembly of GPI anchor in ER membrane (GO:0016254) | 9.177e-21 | 1.377e-19 | 4662.00 | 215093.20 |
| 2 | GPI anchor metabolic process (GO:0006505) | 2.199e-22 | 1.320e-20 | 3994.00 | 199175.49 |
| 3 | GPI anchor biosynthetic process (GO:0006506) | 7.441e-22 | 2.232e-20 | 3327.67 | 161890.56 |
| 4 | glycolipid biosynthetic process (GO:0009247) | 5.437e-21 | 1.087e-19 | 2494.75 | 116407.64 |
| 5 | protein lipidation (GO:0006497) | 2.078e-19 | 2.494e-18 | 1504.68 | 64727.57 |

GO: Molecular Function

| **Index** | **Name** | **P-value** | **Adjusted p-value** | **Odds Ratio** | **Combined score** |
| --- | --- | --- | --- | --- | --- |
| 1 | type 1 fibroblast growth factor receptor binding (GO:0005105) | 0.002498 | 0.009366 | 555.17 | 3326.77 |
| 2 | acetylglucosaminyltransferase activity (GO:0008375) | 0.000001538 | 0.00002308 | 189.95 | 2542.46 |
| 3 | mannosyltransferase activity (GO:0000030) | 0.00005170 | 0.0003877 | 249.63 | 2463.82 |
| 4 | fibroblast growth factor receptor binding (GO:0005104) | 0.01144 | 0.03433 | 100.85 | 450.83 |
| 5 | O-acyltransferase activity (GO:0008374) | 0.01490 | 0.03726 | 76.48 | 321.69 |

GO: Cellular Component

| **Index** | **Name** | **P-value** | **Adjusted p-value** | **Odds Ratio** | **Combined score** |
| --- | --- | --- | --- | --- | --- |
| 1 | endoplasmic reticulum membrane (GO:0005789) | 1.048e-10 | 7.339e-10 | 109.58 | 2517.98 |
| 2 | Golgi lumen (GO:0005796) | 0.04890 | 0.1712 | 22.32 | 67.37 |
| 3 | endoplasmic reticulum lumen (GO:0005788) | 0.1337 | 0.3120 | 7.71 | 15.51 |
| 4 | Golgi membrane (GO:0000139) | 0.2125 | 0.3719 | 4.60 | 7.13 |
| 5 | bounding membrane of organelle (GO:0098588) | 0.3237 | 0.4103 | 2.79 | 3.15 |

Elevated circulating alanine aminotransferase concentration ([HP:0031964](https://hpo.jax.org/app/browse/term/HP:0031964))

KEGG Pathway

| **Index** | **Name** | **P-value** | **Adjusted p-value** | **Odds Ratio** | **Combined score** |
| --- | --- | --- | --- | --- | --- |
| 1 | Taurine and hypotaurine metabolism | 0.005488 | 0.03490 | 222.00 | 1155.57 |
| 2 | Bile secretion | 0.00001034 | 0.0001344 | 98.04 | 1125.55 |
| 3 | Primary bile acid biosynthesis | 0.008469 | 0.03490 | 138.71 | 661.82 |
| 4 | Maturity onset diabetes of the young | 0.01293 | 0.03490 | 88.73 | 385.85 |
| 5 | Biosynthesis of unsaturated fatty acids | 0.01342 | 0.03490 | 85.32 | 367.79 |

GO: Biological Process

| **Index** | **Name** | **P-value** | **Adjusted p-value** | **Odds Ratio** | **Combined score** |
| --- | --- | --- | --- | --- | --- |
| 1 | bile acid secretion (GO:0032782) | 0.002498 | 0.03435 | 555.17 | 3326.77 |
| 2 | fructose transmembrane transport (GO:0015755) | 0.002498 | 0.03435 | 555.17 | 3326.77 |
| 3 | dehydroascorbic acid transport (GO:0070837) | 0.002997 | 0.03435 | 444.11 | 2580.41 |
| 4 | cellular triglyceride homeostasis (GO:0035356) | 0.002997 | 0.03435 | 444.11 | 2580.41 |
| 5 | positive regulation of protein glycosylation (GO:0060050) | 0.002997 | 0.03435 | 444.11 | 2580.41 |

GO: Molecular Function

| **Index** | **Name** | **P-value** | **Adjusted p-value** | **Odds Ratio** | **Combined score** |
| --- | --- | --- | --- | --- | --- |
| 1 | fructose transmembrane transporter activity (GO:0005353) | 0.002498 | 0.02631 | 555.17 | 3326.77 |
| 2 | dehydroascorbic acid transmembrane transporter activity (GO:0033300) | 0.002997 | 0.02631 | 444.11 | 2580.41 |
| 3 | L-aspartate transmembrane transporter activity (GO:0015183) | 0.004492 | 0.02631 | 277.53 | 1500.17 |
| 4 | bile acid binding (GO:0032052) | 0.004492 | 0.02631 | 277.53 | 1500.17 |
| 5 | L-glutamate transmembrane transporter activity (GO:0005313) | 0.006979 | 0.02631 | 170.74 | 847.70 |

GO: Cellular Component

| **Index** | **Name** | **P-value** | **Adjusted p-value** | **Odds Ratio** | **Combined score** |
| --- | --- | --- | --- | --- | --- |
| 1 | anchored component of external side of plasma membrane (GO:0031362) | 0.009957 | 0.05883 | 116.79 | 538.34 |
| 2 | intrinsic component of external side of plasma membrane (GO:0031233) | 0.01194 | 0.05883 | 96.46 | 427.12 |
| 3 | myofibril (GO:0030016) | 0.01342 | 0.05883 | 85.32 | 367.79 |
| 4 | intrinsic component of mitochondrial inner membrane (GO:0031304) | 0.01933 | 0.05883 | 58.34 | 230.20 |
| 5 | anchored component of plasma membrane (GO:0046658) | 0.02277 | 0.05883 | 49.25 | 186.27 |

Elevated circulating aspartate aminotransferase concentration ([HP:0031956](https://hpo.jax.org/app/browse/term/HP:0031956))

KEGG Pathway

| **Index** | **Name** | **P-value** | **Adjusted p-value** | **Odds Ratio** | **Combined score** |
| --- | --- | --- | --- | --- | --- |
| 1 | Sulfur relay system | 0.003994 | 0.03557 | 317.19 | 1751.86 |
| 2 | Taurine and hypotaurine metabolism | 0.005488 | 0.03557 | 222.00 | 1155.57 |
| 3 | Bile secretion | 0.00001034 | 0.0002170 | 98.04 | 1125.55 |
| 4 | Thiamine metabolism | 0.007476 | 0.03557 | 158.54 | 776.21 |
| 5 | Primary bile acid biosynthesis | 0.008469 | 0.03557 | 138.71 | 661.82 |

GO: Biological Process

| **Index** | **Name** | **P-value** | **Adjusted p-value** | **Odds Ratio** | **Combined score** |
| --- | --- | --- | --- | --- | --- |
| 1 | bile acid secretion (GO:0032782) | 0.002498 | 0.02932 | 555.17 | 3326.77 |
| 2 | fructose transmembrane transport (GO:0015755) | 0.002498 | 0.02932 | 555.17 | 3326.77 |
| 3 | dehydroascorbic acid transport (GO:0070837) | 0.002997 | 0.02932 | 444.11 | 2580.41 |
| 4 | cellular triglyceride homeostasis (GO:0035356) | 0.002997 | 0.02932 | 444.11 | 2580.41 |
| 5 | positive regulation of protein glycosylation (GO:0060050) | 0.002997 | 0.02932 | 444.11 | 2580.41 |

GO: Molecular Function

| **Index** | **Name** | **P-value** | **Adjusted p-value** | **Odds Ratio** | **Combined score** |
| --- | --- | --- | --- | --- | --- |
| 1 | fructose transmembrane transporter activity (GO:0005353) | 0.002498 | 0.02695 | 555.17 | 3326.77 |
| 2 | dehydroascorbic acid transmembrane transporter activity (GO:0033300) | 0.002997 | 0.02695 | 444.11 | 2580.41 |
| 3 | sulfurtransferase activity (GO:0016783) | 0.003994 | 0.02695 | 317.19 | 1751.86 |
| 4 | intramolecular transferase activity, phosphotransferases (GO:0016868) | 0.004492 | 0.02695 | 277.53 | 1500.17 |
| 5 | bile acid binding (GO:0032052) | 0.004492 | 0.02695 | 277.53 | 1500.17 |

GO: Cellular Component

| **Index** | **Name** | **P-value** | **Adjusted p-value** | **Odds Ratio** | **Combined score** |
| --- | --- | --- | --- | --- | --- |
| 1 | microbody lumen (GO:0031907) | 0.02424 | 0.1686 | 46.16 | 171.72 |
| 2 | peroxisomal matrix (GO:0005782) | 0.02424 | 0.1686 | 46.16 | 171.72 |
| 3 | tertiary granule lumen (GO:1904724) | 0.02717 | 0.1686 | 41.02 | 147.91 |
| 4 | trans-Golgi network membrane (GO:0032588) | 0.04842 | 0.1686 | 22.55 | 68.29 |
| 5 | ficolin-1-rich granule lumen (GO:1904813) | 0.05984 | 0.1686 | 18.09 | 50.96 |

Elevated circulating creatine kinase concentration ([HP:0003236](https://hpo.jax.org/app/browse/term/HP:0003236))

KEGG Pathway

| **Index** | **Name** | **P-value** | **Adjusted p-value** | **Odds Ratio** | **Combined score** |
| --- | --- | --- | --- | --- | --- |
| 1 | Hypertrophic cardiomyopathy | 1.450e-12 | 1.834e-11 | 355.46 | 9689.66 |
| 2 | Dilated cardiomyopathy | 2.157e-12 | 1.834e-11 | 331.67 | 8909.27 |
| 3 | Cardiac muscle contraction | 6.874e-8 | 3.895e-7 | 159.90 | 2637.14 |
| 4 | Viral myocarditis | 0.000003034 | 0.00001032 | 149.87 | 1904.21 |
| 5 | Adrenergic signaling in cardiomyocytes | 6.163e-7 | 0.000002619 | 90.61 | 1295.71 |

GO: Biological Process

| **Index** | **Name** | **P-value** | **Adjusted p-value** | **Odds Ratio** | **Combined score** |
| --- | --- | --- | --- | --- | --- |
| 1 | actin-myosin filament sliding (GO:0033275) | 3.458e-21 | 2.403e-19 | 2661.33 | 125385.00 |
| 2 | muscle filament sliding (GO:0030049) | 3.458e-21 | 2.403e-19 | 2661.33 | 125385.00 |
| 3 | heart contraction (GO:0060047) | 6.755e-21 | 3.130e-19 | 2419.03 | 112349.58 |
| 4 | cardiac muscle contraction (GO:0060048) | 2.534e-18 | 8.804e-17 | 1725.20 | 69899.68 |
| 5 | cardiac muscle tissue morphogenesis (GO:0055008) | 2.132e-15 | 3.704e-14 | 1151.77 | 38908.86 |

GO: Molecular Function

| **Index** | **Name** | **P-value** | **Adjusted p-value** | **Odds Ratio** | **Combined score** |
| --- | --- | --- | --- | --- | --- |
| 1 | FATZ binding (GO:0051373) | 0.000002248 | 0.00001476 | 1665.58 | 21661.60 |
| 2 | titin binding (GO:0031432) | 2.567e-8 | 6.161e-7 | 856.29 | 14966.03 |
| 3 | myosin heavy chain binding (GO:0032036) | 0.000004719 | 0.00002265 | 999.25 | 12254.80 |
| 4 | nitric-oxide synthase binding (GO:0050998) | 0.003495 | 0.01198 | 370.07 | 2093.27 |
| 5 | telethonin binding (GO:0031433) | 0.003495 | 0.01198 | 370.07 | 2093.27 |

GO: Cellular Component

| **Index** | **Name** | **P-value** | **Adjusted p-value** | **Odds Ratio** | **Combined score** |
| --- | --- | --- | --- | --- | --- |
| 1 | cardiac myofibril (GO:0097512) | 8.995e-10 | 1.169e-8 | 4283.14 | 89214.48 |
| 2 | myofibril (GO:0030016) | 7.595e-13 | 1.975e-11 | 907.64 | 25328.60 |
| 3 | muscle myosin complex (GO:0005859) | 0.00002354 | 0.0002040 | 384.17 | 4094.01 |
| 4 | myosin filament (GO:0032982) | 0.002997 | 0.01113 | 444.11 | 2580.41 |
| 5 | pseudopodium (GO:0031143) | 0.004492 | 0.01460 | 277.53 | 1500.17 |

Elevated circulating creatinine concentration ([HP:0003259](https://hpo.jax.org/app/browse/term/HP:0003259))

KEGG Pathway

| **Index** | **Name** | **P-value** | **Adjusted p-value** | **Odds Ratio** | **Combined score** |
| --- | --- | --- | --- | --- | --- |
| 1 | Maturity onset diabetes of the young | 0.01293 | 0.05177 | 88.73 | 385.85 |
| 2 | Fanconi anemia pathway | 0.02668 | 0.05177 | 41.80 | 151.46 |
| 3 | Cytosolic DNA-sensing pathway | 0.03106 | 0.05177 | 35.71 | 123.99 |
| 4 | Complement and coagulation cascades | 0.04171 | 0.05213 | 26.33 | 83.66 |
| 5 | Wnt signaling pathway | 0.07998 | 0.07998 | 13.35 | 33.72 |

GO: Biological Process

| **Index** | **Name** | **P-value** | **Adjusted p-value** | **Odds Ratio** | **Combined score** |
| --- | --- | --- | --- | --- | --- |
| 1 | protein localization to cilium (GO:0061512) | 5.841e-7 | 0.00003739 | 267.29 | 3836.51 |
| 2 | DNA catabolic process, exonucleolytic (GO:0000738) | 0.002498 | 0.01743 | 555.17 | 3326.77 |
| 3 | nephron tubule development (GO:0072080) | 0.002997 | 0.01743 | 444.11 | 2580.41 |
| 4 | pronephros development (GO:0048793) | 0.002997 | 0.01743 | 444.11 | 2580.41 |
| 5 | protein localization to ciliary membrane (GO:1903441) | 0.003495 | 0.01864 | 370.07 | 2093.27 |

GO: Molecular Function

| **Index** | **Name** | **P-value** | **Adjusted p-value** | **Odds Ratio** | **Combined score** |
| --- | --- | --- | --- | --- | --- |
| 1 | MutLalpha complex binding (GO:0032405) | 0.003495 | 0.01454 | 370.07 | 2093.27 |
| 2 | MutSalpha complex binding (GO:0032407) | 0.003495 | 0.01454 | 370.07 | 2093.27 |
| 3 | 5'-flap endonuclease activity (GO:0017108) | 0.003994 | 0.01454 | 317.19 | 1751.86 |
| 4 | flap endonuclease activity (GO:0048256) | 0.004492 | 0.01454 | 277.53 | 1500.17 |
| 5 | ubiquitination-like modification-dependent protein binding (GO:0140035) | 0.004990 | 0.01454 | 246.68 | 1307.49 |

GO: Cellular Component

| **Index** | **Name** | **P-value** | **Adjusted p-value** | **Odds Ratio** | **Combined score** |
| --- | --- | --- | --- | --- | --- |
| 1 | non-motile cilium (GO:0097730) | 0.01589 | 0.06355 | 71.54 | 296.32 |
| 2 | motile cilium (GO:0031514) | 0.03349 | 0.08931 | 33.04 | 112.22 |
| 3 | cilium (GO:0005929) | 0.005910 | 0.04728 | 21.02 | 107.83 |
| 4 | endoplasmic reticulum lumen (GO:0005788) | 0.1337 | 0.2675 | 7.71 | 15.51 |
| 5 | endoplasmic reticulum membrane (GO:0005789) | 0.3041 | 0.4689 | 3.01 | 3.59 |

Elevated circulating C-reactive protein concentration ([HP:0011227](https://hpo.jax.org/app/browse/term/HP:0011227))

KEGG Pathway

| **Index** | **Name** | **P-value** | **Adjusted p-value** | **Odds Ratio** | **Combined score** |
| --- | --- | --- | --- | --- | --- |
| 1 | Influenza A | 0.000001065 | 0.00003395 | 78.66 | 1081.73 |
| 2 | NOD-like receptor signaling pathway | 0.000001306 | 0.00003395 | 74.63 | 1011.07 |
| 3 | Autoimmune thyroid disease | 0.0003059 | 0.003181 | 97.74 | 790.95 |
| 4 | Yersinia infection | 0.00003643 | 0.0006314 | 63.51 | 649.04 |
| 5 | Antigen processing and presentation | 0.0006621 | 0.005738 | 65.51 | 479.51 |

GO: Biological Process

| **Index** | **Name** | **P-value** | **Adjusted p-value** | **Odds Ratio** | **Combined score** |
| --- | --- | --- | --- | --- | --- |
| 1 | purinergic nucleotide receptor signaling pathway (GO:0035590) | 1.191e-7 | 0.00001834 | 475.52 | 7581.26 |
| 2 | regulation of interleukin-4 production (GO:0032673) | 1.586e-7 | 0.00001834 | 427.93 | 6700.13 |
| 3 | detection of bacterium (GO:0016045) | 0.00002041 | 0.0008853 | 416.21 | 4494.85 |
| 4 | positive regulation of cysteine-type endopeptidase activity (GO:2001056) | 1.733e-8 | 0.000006014 | 229.10 | 4094.25 |
| 5 | positive regulation of alpha-beta T cell proliferation (GO:0046641) | 0.00002690 | 0.0009334 | 356.71 | 3753.85 |

GO: Molecular Function

| **Index** | **Name** | **P-value** | **Adjusted p-value** | **Odds Ratio** | **Combined score** |
| --- | --- | --- | --- | --- | --- |
| 1 | non-membrane spanning protein tyrosine phosphatase activity (GO:0004726) | 0.002997 | 0.04205 | 444.11 | 2580.41 |
| 2 | CD4 receptor binding (GO:0042609) | 0.003994 | 0.04205 | 317.19 | 1751.86 |
| 3 | MHC class II receptor activity (GO:0032395) | 0.004990 | 0.04205 | 246.68 | 1307.49 |
| 4 | endopeptidase activator activity (GO:0061133) | 0.004990 | 0.04205 | 246.68 | 1307.49 |
| 5 | caspase binding (GO:0089720) | 0.005985 | 0.04205 | 201.81 | 1032.95 |

GO: Cellular Component

| **Index** | **Name** | **P-value** | **Adjusted p-value** | **Odds Ratio** | **Combined score** |
| --- | --- | --- | --- | --- | --- |
| 1 | early phagosome (GO:0032009) | 0.005985 | 0.04461 | 201.81 | 1032.95 |
| 2 | T cell receptor complex (GO:0042101) | 0.005985 | 0.04461 | 201.81 | 1032.95 |
| 3 | MHC class II protein complex (GO:0042613) | 0.006482 | 0.04461 | 184.98 | 932.06 |
| 4 | MHC protein complex (GO:0042611) | 0.009957 | 0.05548 | 116.79 | 538.34 |
| 5 | clathrin-coated endocytic vesicle (GO:0045334) | 0.0007857 | 0.03064 | 59.96 | 428.66 |

Increased circulating ferritin concentration ([HP:0003281](https://hpo.jax.org/app/browse/term/HP:0003281))

KEGG Pathway

| **Index** | **Name** | **P-value** | **Adjusted p-value** | **Odds Ratio** | **Combined score** |
| --- | --- | --- | --- | --- | --- |
| 1 | Vitamin digestion and absorption | 0.01194 | 0.07105 | 96.46 | 427.12 |
| 2 | Allograft rejection | 0.01884 | 0.07105 | 59.92 | 237.98 |
| 3 | Apoptosis | 0.002170 | 0.04193 | 35.45 | 217.39 |
| 4 | Graft-versus-host disease | 0.02081 | 0.07105 | 54.06 | 209.35 |
| 5 | Type I diabetes mellitus | 0.02130 | 0.07105 | 52.77 | 203.13 |

GO: Biological Process

| **Index** | **Name** | **P-value** | **Adjusted p-value** | **Odds Ratio** | **Combined score** |
| --- | --- | --- | --- | --- | --- |
| 1 | regulation of nucleotide-binding oligomerization domain containing signaling pathway (GO:0070424) | 0.002498 | 0.02906 | 555.17 | 3326.77 |
| 2 | folate transmembrane transport (GO:0098838) | 0.002498 | 0.02906 | 555.17 | 3326.77 |
| 3 | drug transport (GO:0015893) | 0.002498 | 0.02906 | 555.17 | 3326.77 |
| 4 | cyclic-GMP-AMP transmembrane import across plasma membrane (GO:0140361) | 0.002498 | 0.02906 | 555.17 | 3326.77 |
| 5 | guanine nucleotide transmembrane transport (GO:1903790) | 0.002997 | 0.02906 | 444.11 | 2580.41 |

GO: Molecular Function

| **Index** | **Name** | **P-value** | **Adjusted p-value** | **Odds Ratio** | **Combined score** |
| --- | --- | --- | --- | --- | --- |
| 1 | folic acid transmembrane transporter activity (GO:0008517) | 0.002498 | 0.02463 | 555.17 | 3326.77 |
| 2 | oxidoreductase activity, acting on the CH-CH group of donors, oxygen as acceptor (GO:0016634) | 0.003495 | 0.02463 | 370.07 | 2093.27 |
| 3 | glycine transmembrane transporter activity (GO:0015187) | 0.003994 | 0.02463 | 317.19 | 1751.86 |
| 4 | endopeptidase activator activity (GO:0061133) | 0.004990 | 0.02463 | 246.68 | 1307.49 |
| 5 | purine ribonucleotide transmembrane transporter activity (GO:0005346) | 0.005488 | 0.02463 | 222.00 | 1155.57 |

GO: Celullar Component

| **Index** | **Name** | **P-value** | **Adjusted p-value** | **Odds Ratio** | **Combined score** |
| --- | --- | --- | --- | --- | --- |
| 1 | cytolytic granule (GO:0044194) | 0.000004719 | 0.00008965 | 999.25 | 12254.80 |
| 2 | primary lysosome (GO:0005766) | 0.005488 | 0.05213 | 222.00 | 1155.57 |
| 3 | mitochondrial intermembrane space (GO:0005758) | 0.02863 | 0.09991 | 38.86 | 138.07 |
| 4 | spindle microtubule (GO:0005876) | 0.03009 | 0.09991 | 36.91 | 129.31 |
| 5 | organelle envelope lumen (GO:0031970) | 0.03155 | 0.09991 | 35.14 | 121.47 |

Increased circulating lactate dehydrogenase concentration ([HP:0025435](https://hpo.jax.org/app/browse/term/HP:0025435))

KEGG Pathway

| **Index** | **Name** | **P-value** | **Adjusted p-value** | **Odds Ratio** | **Combined score** |
| --- | --- | --- | --- | --- | --- |
| 1 | Vitamin digestion and absorption | 0.01194 | 0.09166 | 96.46 | 427.12 |
| 2 | Glycosylphosphatidylinositol (GPI)-anchor biosynthesis | 0.01293 | 0.09166 | 88.73 | 385.85 |
| 3 | Asthma | 0.01540 | 0.09166 | 73.93 | 308.54 |
| 4 | Allograft rejection | 0.01884 | 0.09166 | 59.92 | 237.98 |
| 5 | Bladder cancer | 0.02032 | 0.09166 | 55.42 | 215.92 |

GO: Biological Process

| **Index** | **Name** | **P-value** | **Adjusted p-value** | **Odds Ratio** | **Combined score** |
| --- | --- | --- | --- | --- | --- |
| 1 | ammonium transport (GO:0015696) | 0.000004719 | 0.0007957 | 999.25 | 12254.80 |
| 2 | ammonium transmembrane transport (GO:0072488) | 0.000006290 | 0.0007957 | 832.67 | 9972.50 |
| 3 | positive regulation of CD4-positive, CD25-positive, alpha-beta regulatory T cell differentiation (GO:0032831) | 0.002498 | 0.02804 | 555.17 | 3326.77 |
| 4 | folate transmembrane transport (GO:0098838) | 0.002498 | 0.02804 | 555.17 | 3326.77 |
| 5 | cyclic-GMP-AMP transmembrane import across plasma membrane (GO:0140361) | 0.002498 | 0.02804 | 555.17 | 3326.77 |

GO: Molecular Function

| **Index** | **Name** | **P-value** | **Adjusted p-value** | **Odds Ratio** | **Combined score** |
| --- | --- | --- | --- | --- | --- |
| 1 | ammonium transmembrane transporter activity (GO:0008519) | 0.00001010 | 0.0003839 | 624.44 | 7182.69 |
| 2 | folic acid transmembrane transporter activity (GO:0008517) | 0.002498 | 0.01582 | 555.17 | 3326.77 |
| 3 | carnitine O-acyltransferase activity (GO:0016406) | 0.002997 | 0.01627 | 444.11 | 2580.41 |
| 4 | CD4 receptor binding (GO:0042609) | 0.003994 | 0.01634 | 317.19 | 1751.86 |
| 5 | dicarboxylic acid transmembrane transporter activity (GO:0005310) | 0.00009070 | 0.001723 | 184.84 | 1720.52 |

GO: Cellular Component

| **Index** | **Name** | **P-value** | **Adjusted p-value** | **Odds Ratio** | **Combined score** |
| --- | --- | --- | --- | --- | --- |
| 1 | MHC class II protein complex (GO:0042613) | 0.006482 | 0.05575 | 184.98 | 932.06 |
| 2 | SWI/SNF complex (GO:0016514) | 0.009462 | 0.06117 | 123.28 | 574.57 |
| 3 | MHC protein complex (GO:0042611) | 0.009957 | 0.06117 | 116.79 | 538.34 |
| 4 | lumenal side of endoplasmic reticulum membrane (GO:0098553) | 0.01392 | 0.06648 | 82.15 | 351.18 |
| 5 | integral component of lumenal side of endoplasmic reticulum membrane (GO:0071556) | 0.01392 | 0.06648 | 82.15 | 351.18 |

Elevated circulating thyroid-stimulating hormone concentration ([HP:0002925](https://hpo.jax.org/app/browse/term/HP:0002925))

KEGG Pathway

| **Index** | **Name** | **P-value** | **Adjusted p-value** | **Odds Ratio** | **Combined score** |
| --- | --- | --- | --- | --- | --- |
| 1 | Thyroid hormone synthesis | 3.764e-8 | 3.012e-7 | 187.03 | 3197.34 |
| 2 | Autoimmune thyroid disease | 0.000002081 | 0.000008324 | 170.91 | 2236.02 |
| 3 | Tyrosine metabolism | 0.01786 | 0.04762 | 63.35 | 255.00 |
| 4 | Regulation of lipolysis in adipocytes | 0.02717 | 0.05434 | 41.02 | 147.91 |
| 5 | Thyroid hormone signaling pathway | 0.05889 | 0.08859 | 18.40 | 52.10 |

GO: Biological Process

| **Index** | **Name** | **P-value** | **Adjusted p-value** | **Odds Ratio** | **Combined score** |
| --- | --- | --- | --- | --- | --- |
| 1 | thyroid hormone generation (GO:0006590) | 1.975e-8 | 0.000002272 | 951.48 | 16879.14 |
| 2 | thyroid gland development (GO:0030878) | 0.000006290 | 0.0003617 | 832.67 | 9972.50 |
| 3 | lung morphogenesis (GO:0060425) | 0.002498 | 0.01914 | 555.17 | 3326.77 |
| 4 | epithelial tube branching involved in lung morphogenesis (GO:0060441) | 0.002997 | 0.01914 | 444.11 | 2580.41 |
| 5 | equilibrioception (GO:0050957) | 0.002997 | 0.01914 | 444.11 | 2580.41 |

GO: Molecular Function

| **Index** | **Name** | **P-value** | **Adjusted p-value** | **Odds Ratio** | **Combined score** |
| --- | --- | --- | --- | --- | --- |
| 1 | intronic transcription regulatory region sequence-specific DNA binding (GO:0001161) | 0.002498 | 0.02058 | 555.17 | 3326.77 |
| 2 | anion:sodium symporter activity (GO:0015373) | 0.002498 | 0.02058 | 555.17 | 3326.77 |
| 3 | thyroid hormone transmembrane transporter activity (GO:0015349) | 0.002997 | 0.02058 | 444.11 | 2580.41 |
| 4 | cAMP-dependent protein kinase inhibitor activity (GO:0004862) | 0.003994 | 0.02058 | 317.19 | 1751.86 |
| 5 | pyrimidine nucleotide-sugar transmembrane transporter activity (GO:0015165) | 0.003994 | 0.02058 | 317.19 | 1751.86 |

GO: Cellular Component

| **Index** | **Name** | **P-value** | **Adjusted p-value** | **Odds Ratio** | **Combined score** |
| --- | --- | --- | --- | --- | --- |
| 1 | catenin complex (GO:0016342) | 0.01540 | 0.07813 | 73.93 | 308.54 |
| 2 | integral component of Golgi membrane (GO:0030173) | 0.02717 | 0.07813 | 41.02 | 147.91 |
| 3 | extracellular membrane-bounded organelle (GO:0065010) | 0.02766 | 0.07813 | 40.27 | 144.50 |
| 4 | extracellular vesicle (GO:1903561) | 0.02912 | 0.07813 | 38.18 | 135.03 |
| 5 | intrinsic component of Golgi membrane (GO:0031228) | 0.02960 | 0.07813 | 37.53 | 132.12 |

Hyperglycemia ([HP:0003074](https://hpo.jax.org/app/browse/term/HP:0003074))

KEGG Pathway

| **Index** | **Name** | **P-value** | **Adjusted p-value** | **Odds Ratio** | **Combined score** |
| --- | --- | --- | --- | --- | --- |
| 1 | Maturity onset diabetes of the young | 1.106e-22 | 4.312e-21 | 4438.22 | 224380.54 |
| 2 | Type II diabetes mellitus | 2.198e-14 | 4.287e-13 | 748.13 | 23527.39 |
| 3 | Insulin secretion | 1.096e-12 | 1.425e-11 | 373.31 | 10280.78 |
| 4 | Neomycin, kanamycin and gentamicin biosynthesis | 0.002498 | 0.009741 | 555.17 | 3326.77 |
| 5 | Prolactin signaling pathway | 0.000004841 | 0.00004720 | 127.44 | 1559.65 |

GO: Biological Process

| **Index** | **Name** | **P-value** | **Adjusted p-value** | **Odds Ratio** | **Combined score** |
| --- | --- | --- | --- | --- | --- |
| 1 | regulation of peptide hormone secretion (GO:0090276) | 8.432e-16 | 2.268e-13 | 693.84 | 24082.54 |
| 2 | enteroendocrine cell differentiation (GO:0035883) | 0.000002248 | 0.00007559 | 1665.58 | 21661.60 |
| 3 | regulation of protein secretion (GO:0050708) | 3.712e-14 | 4.992e-12 | 392.95 | 12151.82 |
| 4 | nitric oxide mediated signal transduction (GO:0007263) | 7.315e-8 | 0.000002811 | 570.71 | 9377.24 |
| 5 | regulation of insulin secretion (GO:0050796) | 3.526e-12 | 3.162e-10 | 304.47 | 8029.08 |

GO: Molecular Function

| **Index** | **Name** | **P-value** | **Adjusted p-value** | **Odds Ratio** | **Combined score** |
| --- | --- | --- | --- | --- | --- |
| 1 | ATP-activated inward rectifier potassium channel activity (GO:0015272) | 0.000003371 | 0.0001416 | 1249.13 | 15739.23 |
| 2 | glucokinase activity (GO:0004340) | 0.002498 | 0.006556 | 555.17 | 3326.77 |
| 3 | mannokinase activity (GO:0019158) | 0.002498 | 0.006556 | 555.17 | 3326.77 |
| 4 | fructokinase activity (GO:0008865) | 0.002498 | 0.006556 | 555.17 | 3326.77 |
| 5 | fructose transmembrane transporter activity (GO:0005353) | 0.002498 | 0.006556 | 555.17 | 3326.77 |

GO: Cellular Component

| **Index** | **Name** | **P-value** | **Adjusted p-value** | **Odds Ratio** | **Combined score** |
| --- | --- | --- | --- | --- | --- |
| 1 | cation-transporting ATPase complex (GO:0090533) | 0.007973 | 0.05581 | 147.96 | 714.91 |
| 2 | potassium channel complex (GO:0034705) | 0.0006964 | 0.009749 | 63.82 | 463.95 |
| 3 | endosome lumen (GO:0031904) | 0.01293 | 0.06033 | 88.73 | 385.85 |
| 4 | endoplasmic reticulum-Golgi intermediate compartment membrane (GO:0033116) | 0.02424 | 0.08483 | 46.16 | 171.72 |
| 5 | Golgi lumen (GO:0005796) | 0.04890 | 0.1121 | 22.32 | 67.37 |

Hypocalcemia ([HP:0002901](https://hpo.jax.org/app/browse/term/HP:0002901))

KEGG Pathway

| **Index** | **Name** | **P-value** | **Adjusted p-value** | **Odds Ratio** | **Combined score** |
| --- | --- | --- | --- | --- | --- |
| 1 | Parathyroid hormone synthesis, secretion and action | 3.963e-12 | 9.115e-11 | 298.35 | 7832.87 |
| 2 | Rheumatoid arthritis | 0.00001141 | 0.0001312 | 94.76 | 1078.51 |
| 3 | Endocrine and other factor-regulated calcium reabsorption | 0.0003059 | 0.002345 | 97.74 | 790.95 |
| 4 | Prolactin signaling pathway | 0.0005336 | 0.003068 | 73.24 | 551.94 |
| 5 | Steroid biosynthesis | 0.009957 | 0.02290 | 116.79 | 538.34 |

GO: Biological Process

| **Index** | **Name** | **P-value** | **Adjusted p-value** | **Odds Ratio** | **Combined score** |
| --- | --- | --- | --- | --- | --- |
| 1 | positive regulation of prostaglandin secretion (GO:0032308) | 0.000002248 | 0.0002540 | 1665.58 | 21661.60 |
| 2 | positive regulation of fever generation (GO:0031622) | 0.000003371 | 0.0002540 | 1249.13 | 15739.23 |
| 3 | vitamin D metabolic process (GO:0042359) | 4.082e-8 | 0.000009226 | 713.50 | 12139.53 |
| 4 | phosphate ion homeostasis (GO:0055062) | 0.00001010 | 0.0005580 | 624.44 | 7182.69 |
| 5 | positive regulation of bone resorption (GO:0045780) | 0.00001235 | 0.0005580 | 555.03 | 6273.06 |

GO: Molecular Function

| **Index** | **Name** | **P-value** | **Adjusted p-value** | **Odds Ratio** | **Combined score** |
| --- | --- | --- | --- | --- | --- |
| 1 | type 1 fibroblast growth factor receptor binding (GO:0005105) | 0.002498 | 0.02582 | 555.17 | 3326.77 |
| 2 | bile acid binding (GO:0032052) | 0.004492 | 0.02582 | 277.53 | 1500.17 |
| 3 | tumor necrosis factor-activated receptor activity (GO:0005031) | 0.004492 | 0.02582 | 277.53 | 1500.17 |
| 4 | retinoid X receptor binding (GO:0046965) | 0.005488 | 0.02582 | 222.00 | 1155.57 |
| 5 | death receptor activity (GO:0005035) | 0.005985 | 0.02582 | 201.81 | 1032.95 |

GO: Cellular Component

| **Index** | **Name** | **P-value** | **Adjusted p-value** | **Odds Ratio** | **Combined score** |
| --- | --- | --- | --- | --- | --- |
| 1 | Golgi lumen (GO:0005796) | 0.04890 | 0.3423 | 22.32 | 67.37 |
| 2 | endoplasmic reticulum lumen (GO:0005788) | 0.1337 | 0.3763 | 7.71 | 15.51 |
| 3 | integral component of plasma membrane (GO:0005887) | 0.1613 | 0.3763 | 3.19 | 5.82 |
| 4 | endoplasmic reticulum membrane (GO:0005789) | 0.3041 | 0.3960 | 3.01 | 3.59 |
| 5 | intracellular membrane-bounded organelle (GO:0043231) | 0.2469 | 0.3960 | 1.90 | 2.66 |

Hypofibrinogenemia ([HP:0011900](https://hpo.jax.org/app/browse/term/HP:0011900))

KEGG Pathway

| **Index** | **Name** | **P-value** | **Adjusted p-value** | **Odds Ratio** | **Combined score** |
| --- | --- | --- | --- | --- | --- |
| 1 | Apoptosis | 0.00004054 | 0.001541 | 61.21 | 618.98 |
| 2 | Allograft rejection | 0.01884 | 0.08352 | 59.92 | 237.98 |
| 3 | Graft-versus-host disease | 0.02081 | 0.08352 | 54.06 | 209.35 |
| 4 | Type I diabetes mellitus | 0.02130 | 0.08352 | 52.77 | 203.13 |
| 5 | Necroptosis | 0.002710 | 0.04426 | 31.58 | 186.67 |

GO: Biological Process

| **Index** | **Name** | **P-value** | **Adjusted p-value** | **Odds Ratio** | **Combined score** |
| --- | --- | --- | --- | --- | --- |
| 1 | regulation of nucleotide-binding oligomerization domain containing signaling pathway (GO:0070424) | 0.002498 | 0.03427 | 555.17 | 3326.77 |
| 2 | positive regulation of killing of cells of other organism (GO:0051712) | 0.002997 | 0.03427 | 444.11 | 2580.41 |
| 3 | taurine metabolic process (GO:0019530) | 0.002997 | 0.03427 | 444.11 | 2580.41 |
| 4 | negative regulation of cAMP-dependent protein kinase activity (GO:2000480) | 0.003994 | 0.03427 | 317.19 | 1751.86 |
| 5 | regulation of pattern recognition receptor signaling pathway (GO:0062207) | 0.003994 | 0.03427 | 317.19 | 1751.86 |

GO: Molecular Function

| **Index** | **Name** | **P-value** | **Adjusted p-value** | **Odds Ratio** | **Combined score** |
| --- | --- | --- | --- | --- | --- |
| 1 | cAMP-dependent protein kinase inhibitor activity (GO:0004862) | 0.003994 | 0.03467 | 317.19 | 1751.86 |
| 2 | glucocorticoid receptor binding (GO:0035259) | 0.004492 | 0.03467 | 277.53 | 1500.17 |
| 3 | protein-glutamine gamma-glutamyltransferase activity (GO:0003810) | 0.004492 | 0.03467 | 277.53 | 1500.17 |
| 4 | cAMP-dependent protein kinase regulator activity (GO:0008603) | 0.004990 | 0.03467 | 246.68 | 1307.49 |
| 5 | endopeptidase activator activity (GO:0061133) | 0.004990 | 0.03467 | 246.68 | 1307.49 |

GO: Cellular Component

| **Index** | **Name** | **P-value** | **Adjusted p-value** | **Odds Ratio** | **Combined score** |
| --- | --- | --- | --- | --- | --- |
| 1 | cytolytic granule (GO:0044194) | 0.000004719 | 0.00006604 | 999.25 | 12254.80 |
| 2 | spectrin-associated cytoskeleton (GO:0014731) | 0.000006290 | 0.00006604 | 832.67 | 9972.50 |
| 3 | intrinsic component of the cytoplasmic side of the plasma membrane (GO:0031235) | 0.003994 | 0.01677 | 317.19 | 1751.86 |
| 4 | primary lysosome (GO:0005766) | 0.005488 | 0.01921 | 222.00 | 1155.57 |
| 5 | cytoplasmic side of plasma membrane (GO:0009898) | 0.0003294 | 0.002306 | 94.04 | 754.04 |

Hypoglycemia ([HP:0001943](https://hpo.jax.org/app/browse/term/HP:0001943))

KEGG Pathway

| **Index** | **Name** | **P-value** | **Adjusted p-value** | **Odds Ratio** | **Combined score** |
| --- | --- | --- | --- | --- | --- |
| 1 | Maturity onset diabetes of the young | 1.106e-22 | 4.312e-21 | 4438.22 | 224380.54 |
| 2 | Type II diabetes mellitus | 2.198e-14 | 4.287e-13 | 748.13 | 23527.39 |
| 3 | Insulin secretion | 1.096e-12 | 1.425e-11 | 373.31 | 10280.78 |
| 4 | Neomycin, kanamycin and gentamicin biosynthesis | 0.002498 | 0.009741 | 555.17 | 3326.77 |
| 5 | Prolactin signaling pathway | 0.000004841 | 0.00004720 | 127.44 | 1559.65 |

GO: Biological Process

| **Index** | **Name** | **P-value** | **Adjusted p-value** | **Odds Ratio** | **Combined score** |
| --- | --- | --- | --- | --- | --- |
| 1 | regulation of peptide hormone secretion (GO:0090276) | 8.432e-16 | 2.268e-13 | 693.84 | 24082.54 |
| 2 | enteroendocrine cell differentiation (GO:0035883) | 0.000002248 | 0.00007559 | 1665.58 | 21661.60 |
| 3 | regulation of protein secretion (GO:0050708) | 3.712e-14 | 4.992e-12 | 392.95 | 12151.82 |
| 4 | nitric oxide mediated signal transduction (GO:0007263) | 7.315e-8 | 0.000002811 | 570.71 | 9377.24 |
| 5 | regulation of insulin secretion (GO:0050796) | 3.526e-12 | 3.162e-10 | 304.47 | 8029.08 |

GO: Molecular Function

| **Index** | **Name** | **P-value** | **Adjusted p-value** | **Odds Ratio** | **Combined score** |
| --- | --- | --- | --- | --- | --- |
| 1 | ATP-activated inward rectifier potassium channel activity (GO:0015272) | 0.000003371 | 0.0001416 | 1249.13 | 15739.23 |
| 2 | glucokinase activity (GO:0004340) | 0.002498 | 0.006556 | 555.17 | 3326.77 |
| 3 | mannokinase activity (GO:0019158) | 0.002498 | 0.006556 | 555.17 | 3326.77 |
| 4 | fructokinase activity (GO:0008865) | 0.002498 | 0.006556 | 555.17 | 3326.77 |
| 5 | fructose transmembrane transporter activity (GO:0005353) | 0.002498 | 0.006556 | 555.17 | 3326.77 |

GO: Cellular Component

| **Index** | **Name** | **P-value** | **Adjusted p-value** | **Odds Ratio** | **Combined score** |
| --- | --- | --- | --- | --- | --- |
| 1 | cation-transporting ATPase complex (GO:0090533) | 0.007973 | 0.05581 | 147.96 | 714.91 |
| 2 | potassium channel complex (GO:0034705) | 0.0006964 | 0.009749 | 63.82 | 463.95 |
| 3 | endosome lumen (GO:0031904) | 0.01293 | 0.06033 | 88.73 | 385.85 |
| 4 | endoplasmic reticulum-Golgi intermediate compartment membrane (GO:0033116) | 0.02424 | 0.08483 | 46.16 | 171.72 |
| 5 | Golgi lumen (GO:0005796) | 0.04890 | 0.1121 | 22.32 | 67.37 |

Hypophosphatemia ([HP:0002148](https://hpo.jax.org/app/browse/term/HP:0002148))

KEGG Pathway

| **Index** | **Name** | **P-value** | **Adjusted p-value** | **Odds Ratio** | **Combined score** |
| --- | --- | --- | --- | --- | --- |
| 1 | Parathyroid hormone synthesis, secretion and action | 3.963e-12 | 1.149e-10 | 298.35 | 7832.87 |
| 2 | Mineral absorption | 0.000003034 | 0.00004400 | 149.87 | 1904.21 |
| 3 | Riboflavin metabolism | 0.003994 | 0.02895 | 317.19 | 1751.86 |
| 4 | Pantothenate and CoA biosynthesis | 0.01045 | 0.05052 | 110.94 | 506.01 |
| 5 | Nicotinate and nicotinamide metabolism | 0.01737 | 0.06474 | 65.22 | 264.33 |

GO: Biological Process

| **Index** | **Name** | **P-value** | **Adjusted p-value** | **Odds Ratio** | **Combined score** |
| --- | --- | --- | --- | --- | --- |
| 1 | cellular trivalent inorganic anion homeostasis (GO:0072502) | 1.799e-9 | 9.952e-8 | 2855.29 | 57494.92 |
| 2 | cellular phosphate ion homeostasis (GO:0030643) | 1.799e-9 | 9.952e-8 | 2855.29 | 57494.92 |
| 3 | phosphate ion homeostasis (GO:0055062) | 6.605e-12 | 1.096e-9 | 2220.44 | 57161.21 |
| 4 | trivalent inorganic anion homeostasis (GO:0072506) | 0.000002248 | 0.00007464 | 1665.58 | 21661.60 |
| 5 | negative regulation of bone mineralization (GO:0030502) | 0.00002041 | 0.0005583 | 416.21 | 4494.85 |

GO: Molecular Function

| **Index** | **Name** | **P-value** | **Adjusted p-value** | **Odds Ratio** | **Combined score** |
| --- | --- | --- | --- | --- | --- |
| 1 | sodium:phosphate symporter activity (GO:0005436) | 0.00001235 | 0.0005926 | 555.03 | 6273.06 |
| 2 | solute:proton antiporter activity (GO:0015299) | 0.002498 | 0.01713 | 555.17 | 3326.77 |
| 3 | type 1 fibroblast growth factor receptor binding (GO:0005105) | 0.002498 | 0.01713 | 555.17 | 3326.77 |
| 4 | phosphodiesterase I activity (GO:0004528) | 0.002498 | 0.01713 | 555.17 | 3326.77 |
| 5 | hydrolase activity, acting on acid anhydrides, in phosphorus-containing anhydrides (GO:0016818) | 0.003495 | 0.01996 | 370.07 | 2093.27 |

GO: Cellular Component

| **Index** | **Name** | **P-value** | **Adjusted p-value** | **Odds Ratio** | **Combined score** |
| --- | --- | --- | --- | --- | --- |
| 1 | brush border membrane (GO:0031526) | 0.01835 | 0.05811 | 61.59 | 246.23 |
| 2 | integral component of plasma membrane (GO:0005887) | 0.00002378 | 0.0004517 | 19.21 | 204.50 |
| 3 | vesicle (GO:0031982) | 0.005389 | 0.04011 | 22.06 | 115.23 |
| 4 | lytic vacuole membrane (GO:0098852) | 0.007445 | 0.04011 | 18.61 | 91.19 |
| 5 | endoplasmic reticulum lumen (GO:0005788) | 0.008444 | 0.04011 | 17.41 | 83.12 |

Thrombocytopenia ([HP:0001873](https://hpo.jax.org/app/browse/term/HP:0001873))

KEGG Pathway

| **Index** | **Name** | **P-value** | **Adjusted p-value** | **Odds Ratio** | **Combined score** |
| --- | --- | --- | --- | --- | --- |
| 1 | Fanconi anemia pathway | 8.485e-27 | 6.788e-26 | 199460.00 | 11973883.48 |
| 2 | Homologous recombination | 9.499e-7 | 0.000003800 | 225.02 | 3120.36 |
| 3 | Ubiquitin mediated proteolysis | 0.002110 | 0.005627 | 35.96 | 221.57 |
| 4 | Pancreatic cancer | 0.03736 | 0.07473 | 29.50 | 96.98 |
| 5 | Breast cancer | 0.07113 | 0.1138 | 15.10 | 39.92 |

GO: Biological Process

| **Index** | **Name** | **P-value** | **Adjusted p-value** | **Odds Ratio** | **Combined score** |
| --- | --- | --- | --- | --- | --- |
| 1 | interstrand cross-link repair (GO:0036297) | 8.598e-20 | 1.307e-17 | 1697.28 | 74510.81 |
| 2 | DNA repair (GO:0006281) | 3.166e-16 | 2.406e-14 | 613.53 | 21896.03 |
| 3 | double-strand break repair via synthesis-dependent strand annealing (GO:0045003) | 0.000002248 | 0.00004271 | 1665.58 | 21661.60 |
| 4 | double-strand break repair via homologous recombination (GO:0000724) | 5.976e-10 | 3.028e-8 | 216.28 | 4593.43 |
| 5 | resolution of meiotic recombination intermediates (GO:0000712) | 0.00002690 | 0.0004089 | 356.71 | 3753.85 |

GO: Molecular Function

| **Index** | **Name** | **P-value** | **Adjusted p-value** | **Odds Ratio** | **Combined score** |
| --- | --- | --- | --- | --- | --- |
| 1 | DNA polymerase binding (GO:0070182) | 0.00003428 | 0.0009598 | 312.09 | 3208.63 |
| 2 | four-way junction helicase activity (GO:0009378) | 0.002997 | 0.01892 | 444.11 | 2580.41 |
| 3 | crossover junction endodeoxyribonuclease activity (GO:0008821) | 0.002997 | 0.01892 | 444.11 | 2580.41 |
| 4 | 5'-flap endonuclease activity (GO:0017108) | 0.003994 | 0.01892 | 317.19 | 1751.86 |
| 5 | flap endonuclease activity (GO:0048256) | 0.004492 | 0.01892 | 277.53 | 1500.17 |

GO: Cellular Component

| **Index** | **Name** | **P-value** | **Adjusted p-value** | **Odds Ratio** | **Combined score** |
| --- | --- | --- | --- | --- | --- |
| 1 | condensed chromosome (GO:0000793) | 0.0003175 | 0.001429 | 95.86 | 772.11 |
| 2 | condensed nuclear chromosome (GO:0000794) | 0.009957 | 0.01792 | 116.79 | 538.34 |
| 3 | chromosome (GO:0005694) | 0.00005786 | 0.0005207 | 54.14 | 528.26 |
| 4 | nuclear chromosome (GO:0000228) | 0.04074 | 0.06111 | 26.98 | 86.34 |
| 5 | nucleus (GO:0005634) | 0.001775 | 0.005324 | 8.09 | 51.21 |
