## Supplementary Table 1-3 for "Long COVID: G Protein-Coupled Receptors (GPCRs) responsible for persistent post-COVID symptoms": Supplementary File 2.6-General.docx

General-pain

Arthralgia ([HP:0002829](https://hpo.jax.org/app/browse/term/HP:0002829))

KEGG Pathway

| **Index** | **Name** | **P-value** | **Adjusted p-value** | **Odds Ratio** | **Combined score** |
| --- | --- | --- | --- | --- | --- |
| 1 | Primary immunodeficiency | 7.523e-7 | 0.00001843 | 244.35 | 3445.31 |
| 2 | Hematopoietic cell lineage | 1.159e-7 | 0.000005680 | 139.61 | 2229.69 |
| 3 | Intestinal immune network for IgA production | 0.0002507 | 0.002457 | 108.39 | 898.70 |
| 4 | Autoimmune thyroid disease | 0.0003059 | 0.002498 | 97.74 | 790.95 |
| 5 | Cytokine-cytokine receptor interaction | 0.000009081 | 0.0001483 | 45.13 | 523.92 |

GO: Biological Process

| **Index** | **Name** | **P-value** | **Adjusted p-value** | **Odds Ratio** | **Combined score** |
| --- | --- | --- | --- | --- | --- |
| 1 | negative regulation of B cell proliferation (GO:0030889) | 0.00001481 | 0.0004628 | 499.50 | 5554.53 |
| 2 | B cell proliferation (GO:0042100) | 4.430e-7 | 0.00003339 | 294.99 | 4315.59 |
| 3 | B cell receptor signaling pathway (GO:0050853) | 5.342e-7 | 0.00003339 | 275.93 | 3985.12 |
| 4 | embryonic skeletal joint morphogenesis (GO:0060272) | 0.002498 | 0.01559 | 555.17 | 3326.77 |
| 5 | negative regulation of regulatory T cell differentiation (GO:0045590) | 0.002498 | 0.01559 | 555.17 | 3326.77 |

GO: Molecular Function

| **Index** | **Name** | **P-value** | **Adjusted p-value** | **Odds Ratio** | **Combined score** |
| --- | --- | --- | --- | --- | --- |
| 1 | MHC class II protein binding (GO:0042289) | 0.002997 | 0.01317 | 444.11 | 2580.41 |
| 2 | tumor necrosis factor-activated receptor activity (GO:0005031) | 0.004492 | 0.01317 | 277.53 | 1500.17 |
| 3 | platelet-derived growth factor binding (GO:0048407) | 0.005488 | 0.01317 | 222.00 | 1155.57 |
| 4 | complement receptor activity (GO:0004875) | 0.005488 | 0.01317 | 222.00 | 1155.57 |
| 5 | death receptor activity (GO:0005035) | 0.005985 | 0.01317 | 201.81 | 1032.95 |

GO: Cellular Component

| **Index** | **Name** | **P-value** | **Adjusted p-value** | **Odds Ratio** | **Combined score** |
| --- | --- | --- | --- | --- | --- |
| 1 | CD95 death-inducing signaling complex (GO:0031265) | 0.002498 | 0.01125 | 555.17 | 3326.77 |
| 2 | death-inducing signaling complex (GO:0031264) | 0.003994 | 0.01125 | 317.19 | 1751.86 |
| 3 | membrane raft (GO:0045121) | 0.002846 | 0.01125 | 30.79 | 180.49 |
| 4 | clathrin-coated vesicle (GO:0030136) | 0.03930 | 0.05097 | 28.00 | 90.64 |
| 5 | plasma membrane raft (GO:0044853) | 0.04026 | 0.05097 | 27.31 | 87.73 |

Back pain ([HP:0003418](https://hpo.jax.org/app/browse/term/HP:0003418))

KEGG Pathway

| **Index** | **Name** | **P-value** | **Adjusted p-value** | **Odds Ratio** | **Combined score** |
| --- | --- | --- | --- | --- | --- |
| 1 | Chronic myeloid leukemia | 0.000006207 | 0.0001552 | 116.93 | 1401.96 |
| 2 | Pancreatic cancer | 0.000006207 | 0.0001552 | 116.93 | 1401.96 |
| 3 | Gastric cancer | 6.000e-7 | 0.00006061 | 91.24 | 1307.15 |
| 4 | Aldosterone-regulated sodium reabsorption | 0.0001485 | 0.001326 | 142.54 | 1256.48 |
| 5 | Colorectal cancer | 0.000009013 | 0.0001878 | 102.79 | 1194.10 |

GO: Biological Process

| **Index** | **Name** | **P-value** | **Adjusted p-value** | **Odds Ratio** | **Combined score** |
| --- | --- | --- | --- | --- | --- |
| 1 | positive regulation of histone H3-K9 acetylation (GO:2000617) | 0.000002248 | 0.0007621 | 1665.58 | 21661.60 |
| 2 | regulation of histone H3-K9 acetylation (GO:2000615) | 0.00001235 | 0.002093 | 555.03 | 6273.06 |
| 3 | positive regulation of histone H3-K4 methylation (GO:0051571) | 0.00002354 | 0.002660 | 384.17 | 4094.01 |
| 4 | cellular response to indole-3-methanol (GO:0071681) | 0.002498 | 0.03309 | 555.17 | 3326.77 |
| 5 | establishment of protein localization to telomere (GO:0070200) | 0.002498 | 0.03309 | 555.17 | 3326.77 |

GO: Molecular Function

| **Index** | **Name** | **P-value** | **Adjusted p-value** | **Odds Ratio** | **Combined score** |
| --- | --- | --- | --- | --- | --- |
| 1 | transcription coactivator binding (GO:0001223) | 0.00004255 | 0.001127 | 277.39 | 2791.90 |
| 2 | RNA-directed DNA polymerase activity (GO:0003964) | 0.002997 | 0.02269 | 444.11 | 2580.41 |
| 3 | telomerase activity (GO:0003720) | 0.002997 | 0.02269 | 444.11 | 2580.41 |
| 4 | 1-phosphatidylinositol-4-phosphate 3-kinase activity (GO:0035005) | 0.003495 | 0.02316 | 370.07 | 2093.27 |
| 5 | potassium channel inhibitor activity (GO:0019870) | 0.004492 | 0.02404 | 277.53 | 1500.17 |

GO: Cellular Component

| **Index** | **Name** | **P-value** | **Adjusted p-value** | **Odds Ratio** | **Combined score** |
| --- | --- | --- | --- | --- | --- |
| 1 | phosphatidylinositol 3-kinase complex, class I (GO:0097651) | 0.002498 | 0.04492 | 555.17 | 3326.77 |
| 2 | transferase complex, transferring phosphorus-containing groups (GO:0061695) | 0.004492 | 0.04492 | 277.53 | 1500.17 |
| 3 | intercalated disc (GO:0014704) | 0.01540 | 0.09303 | 73.93 | 308.54 |
| 4 | cell-cell contact zone (GO:0044291) | 0.02326 | 0.09303 | 48.17 | 181.19 |
| 5 | mitochondrial outer membrane (GO:0005741) | 0.06126 | 0.1930 | 17.66 | 49.31 |

Bone pain ([HP:0002653](https://hpo.jax.org/app/browse/term/HP:0002653))

KEGG Pathway

| **Index** | **Name** | **P-value** | **Adjusted p-value** | **Odds Ratio** | **Combined score** |
| --- | --- | --- | --- | --- | --- |
| 1 | Parathyroid hormone synthesis, secretion and action | 9.380e-10 | 4.033e-8 | 196.92 | 4093.45 |
| 2 | Mineral absorption | 0.000003034 | 0.00006524 | 149.87 | 1904.21 |
| 3 | Riboflavin metabolism | 0.003994 | 0.02862 | 317.19 | 1751.86 |
| 4 | Prolactin signaling pathway | 0.0005336 | 0.005736 | 73.24 | 551.94 |
| 5 | Pantothenate and CoA biosynthesis | 0.01045 | 0.06421 | 110.94 | 506.01 |

GO: Biological Process

| **Index** | **Name** | **P-value** | **Adjusted p-value** | **Odds Ratio** | **Combined score** |
| --- | --- | --- | --- | --- | --- |
| 1 | cellular trivalent inorganic anion homeostasis (GO:0072502) | 1.799e-9 | 1.103e-7 | 2855.29 | 57494.92 |
| 2 | cellular phosphate ion homeostasis (GO:0030643) | 1.799e-9 | 1.103e-7 | 2855.29 | 57494.92 |
| 3 | phosphate ion homeostasis (GO:0055062) | 6.605e-12 | 1.215e-9 | 2220.44 | 57161.21 |
| 4 | trivalent inorganic anion homeostasis (GO:0072506) | 0.000002248 | 0.00008273 | 1665.58 | 21661.60 |
| 5 | negative regulation of bone mineralization (GO:0030502) | 0.00002041 | 0.0006188 | 416.21 | 4494.85 |

GO: Molecular Function

| **Index** | **Name** | **P-value** | **Adjusted p-value** | **Odds Ratio** | **Combined score** |
| --- | --- | --- | --- | --- | --- |
| 1 | sodium:phosphate symporter activity (GO:0005436) | 0.00001235 | 0.0005679 | 555.03 | 6273.06 |
| 2 | solute:proton antiporter activity (GO:0015299) | 0.002498 | 0.02087 | 555.17 | 3326.77 |
| 3 | type 1 fibroblast growth factor receptor binding (GO:0005105) | 0.002498 | 0.02087 | 555.17 | 3326.77 |
| 4 | phosphodiesterase I activity (GO:0004528) | 0.002498 | 0.02087 | 555.17 | 3326.77 |
| 5 | hydrolase activity, acting on acid anhydrides, in phosphorus-containing anhydrides (GO:0016818) | 0.003495 | 0.02087 | 370.07 | 2093.27 |

GO: Cellular Component

| **Index** | **Name** | **P-value** | **Adjusted p-value** | **Odds Ratio** | **Combined score** |
| --- | --- | --- | --- | --- | --- |
| 1 | brush border membrane (GO:0031526) | 0.01835 | 0.05811 | 61.59 | 246.23 |
| 2 | vesicle (GO:0031982) | 0.005389 | 0.04011 | 22.06 | 115.23 |
| 3 | integral component of plasma membrane (GO:0005887) | 0.0003728 | 0.007084 | 12.80 | 101.01 |
| 4 | lytic vacuole membrane (GO:0098852) | 0.007445 | 0.04011 | 18.61 | 91.19 |
| 5 | endoplasmic reticulum lumen (GO:0005788) | 0.008444 | 0.04011 | 17.41 | 83.12 |

Chest pain ([HP:0100749](https://hpo.jax.org/app/browse/term/HP:0100749))

KEGG Pathway

| **Index** | **Name** | **P-value** | **Adjusted p-value** | **Odds Ratio** | **Combined score** |
| --- | --- | --- | --- | --- | --- |
| 1 | Thyroid cancer | 0.0001485 | 0.004528 | 142.54 | 1256.48 |
| 2 | Adherens junction | 0.0005489 | 0.01116 | 72.18 | 541.87 |
| 3 | Colorectal cancer | 0.0008042 | 0.01226 | 59.24 | 422.16 |
| 4 | AGE-RAGE signaling pathway in diabetic complications | 0.001085 | 0.01324 | 50.74 | 346.39 |
| 5 | Citrate cycle (TCA cycle) | 0.01490 | 0.05682 | 76.48 | 321.69 |

GO: Biological Process

| **Index** | **Name** | **P-value** | **Adjusted p-value** | **Odds Ratio** | **Combined score** |
| --- | --- | --- | --- | --- | --- |
| 1 | positive regulation of histone H3-K4 methylation (GO:0051571) | 0.00002354 | 0.002621 | 384.17 | 4094.01 |
| 2 | sympathetic nervous system development (GO:0048485) | 0.00002690 | 0.002621 | 356.71 | 3753.85 |
| 3 | regulation of neuroblast proliferation (GO:1902692) | 0.00003048 | 0.002621 | 332.92 | 3461.84 |
| 4 | cellular response to indole-3-methanol (GO:0071681) | 0.002498 | 0.02358 | 555.17 | 3326.77 |
| 5 | positive regulation of cell proliferation involved in heart morphogenesis (GO:2000138) | 0.002498 | 0.02358 | 555.17 | 3326.77 |

GO: Molecular Function

| **Index** | **Name** | **P-value** | **Adjusted p-value** | **Odds Ratio** | **Combined score** |
| --- | --- | --- | --- | --- | --- |
| 1 | I-SMAD binding (GO:0070411) | 0.00001750 | 0.0007524 | 454.07 | 4973.59 |
| 2 | phosphatidylethanolamine binding (GO:0008429) | 0.004492 | 0.04719 | 277.53 | 1500.17 |
| 3 | histone methyltransferase binding (GO:1990226) | 0.005488 | 0.04719 | 222.00 | 1155.57 |
| 4 | platelet-derived growth factor binding (GO:0048407) | 0.005488 | 0.04719 | 222.00 | 1155.57 |
| 5 | transcription coregulator binding (GO:0001221) | 0.0003059 | 0.006576 | 97.74 | 790.95 |

GO: Cellular Component

| **Index** | **Name** | **P-value** | **Adjusted p-value** | **Odds Ratio** | **Combined score** |
| --- | --- | --- | --- | --- | --- |
| 1 | beta-catenin-TCF complex (GO:1990907) | 0.006482 | 0.06276 | 184.98 | 932.06 |
| 2 | MLL1 complex (GO:0071339) | 0.01293 | 0.06351 | 88.73 | 385.85 |
| 3 | MLL1/2 complex (GO:0044665) | 0.01293 | 0.06351 | 88.73 | 385.85 |
| 4 | axon (GO:0030424) | 0.0001190 | 0.003928 | 42.19 | 381.27 |
| 5 | catenin complex (GO:0016342) | 0.01540 | 0.06351 | 73.93 | 308.54 |

Limb pain ([HP:0009763](https://hpo.jax.org/app/browse/term/HP:0009763))

KEGG Pathway

| **Index** | **Name** | **P-value** | **Adjusted p-value** | **Odds Ratio** | **Combined score** |
| --- | --- | --- | --- | --- | --- |
| 1 | Protein digestion and absorption | 1.360e-7 | 0.000001632 | 133.95 | 2117.75 |
| 2 | ECM-receptor interaction | 0.000009659 | 0.00005795 | 100.36 | 1158.94 |
| 3 | Focal adhesion | 0.0001139 | 0.0004557 | 42.84 | 388.99 |
| 4 | Porphyrin and chlorophyll metabolism | 0.02130 | 0.04155 | 52.77 | 203.13 |
| 5 | Human papillomavirus infection | 0.0004945 | 0.001444 | 25.69 | 195.56 |

GO: Biological Process

| **Index** | **Name** | **P-value** | **Adjusted p-value** | **Odds Ratio** | **Combined score** |
| --- | --- | --- | --- | --- | --- |
| 1 | positive regulation of vascular associated smooth muscle cell apoptotic process (GO:1905461) | 0.002498 | 0.02597 | 555.17 | 3326.77 |
| 2 | embryonic skeletal joint morphogenesis (GO:0060272) | 0.002498 | 0.02597 | 555.17 | 3326.77 |
| 3 | embryonic skeletal joint development (GO:0072498) | 0.002997 | 0.02608 | 444.11 | 2580.41 |
| 4 | collagen fibril organization (GO:0030199) | 7.537e-8 | 0.000002437 | 156.12 | 2560.47 |
| 5 | regulation of vascular associated smooth muscle cell apoptotic process (GO:1905459) | 0.003495 | 0.02608 | 370.07 | 2093.27 |

GO: Molecular Function

| **Index** | **Name** | **P-value** | **Adjusted p-value** | **Odds Ratio** | **Combined score** |
| --- | --- | --- | --- | --- | --- |
| 1 | serine C-palmitoyltransferase activity (GO:0004758) | 0.002498 | 0.01223 | 555.17 | 3326.77 |
| 2 | C-palmitoyltransferase activity (GO:0016454) | 0.002498 | 0.01223 | 555.17 | 3326.77 |
| 3 | MHC class II protein binding (GO:0042289) | 0.002997 | 0.01223 | 444.11 | 2580.41 |
| 4 | oxidoreductase activity, acting on the CH-CH group of donors, oxygen as acceptor (GO:0016634) | 0.003495 | 0.01223 | 370.07 | 2093.27 |
| 5 | platelet-derived growth factor binding (GO:0048407) | 0.005488 | 0.01280 | 222.00 | 1155.57 |

GO: Cellular Component

| **Index** | **Name** | **P-value** | **Adjusted p-value** | **Odds Ratio** | **Combined score** |
| --- | --- | --- | --- | --- | --- |
| 1 | endoplasmic reticulum tubular network membrane (GO:0098826) | 0.002498 | 0.01149 | 555.17 | 3326.77 |
| 2 | serine C-palmitoyltransferase complex (GO:0017059) | 0.003994 | 0.01469 | 317.19 | 1751.86 |
| 3 | endoplasmic reticulum lumen (GO:0005788) | 1.348e-7 | 0.000003102 | 70.39 | 1113.55 |
| 4 | intrinsic component of mitochondrial membrane (GO:0098573) | 0.005985 | 0.01721 | 201.81 | 1032.95 |
| 5 | basement membrane (GO:0005604) | 0.0002944 | 0.001693 | 99.70 | 810.62 |

Myalgia ([HP:0003326](https://hpo.jax.org/app/browse/term/HP:0003326))

KEGG Pathway

| **Index** | **Name** | **P-value** | **Adjusted p-value** | **Odds Ratio** | **Combined score** |
| --- | --- | --- | --- | --- | --- |
| 1 | Mannose type O-glycan biosynthesis | 0.00005661 | 0.0003396 | 237.73 | 2324.81 |
| 2 | Glucagon signaling pathway | 0.00001738 | 0.0003129 | 81.95 | 898.15 |
| 3 | Insulin signaling pathway | 0.00003643 | 0.0003278 | 63.51 | 649.04 |
| 4 | Starch and sucrose metabolism | 0.01786 | 0.06429 | 63.35 | 255.00 |
| 5 | Viral myocarditis | 0.02960 | 0.08458 | 37.53 | 132.12 |

GO: Biological Process

| **Index** | **Name** | **P-value** | **Adjusted p-value** | **Odds Ratio** | **Combined score** |
| --- | --- | --- | --- | --- | --- |
| 1 | cellular glucan metabolic process (GO:0006073) | 1.482e-8 | 0.000001771 | 1070.46 | 19297.66 |
| 2 | regulation of skeletal muscle contraction (GO:0014819) | 0.000004719 | 0.0001410 | 999.25 | 12254.80 |
| 3 | glucan catabolic process (GO:0009251) | 4.082e-8 | 0.000002401 | 713.50 | 12139.53 |
| 4 | glycogen catabolic process (GO:0005980) | 4.082e-8 | 0.000002401 | 713.50 | 12139.53 |
| 5 | cellular polysaccharide catabolic process (GO:0044247) | 5.023e-8 | 0.000002401 | 658.58 | 11068.58 |

GO: Molecular Function

| **Index** | **Name** | **P-value** | **Adjusted p-value** | **Odds Ratio** | **Combined score** |
| --- | --- | --- | --- | --- | --- |
| 1 | 1,4-alpha-oligoglucan phosphorylase activity (GO:0004645) | 0.002498 | 0.02089 | 555.17 | 3326.77 |
| 2 | sodium ion binding (GO:0031402) | 0.002997 | 0.02089 | 444.11 | 2580.41 |
| 3 | nitric-oxide synthase binding (GO:0050998) | 0.003495 | 0.02089 | 370.07 | 2093.27 |
| 4 | potassium channel inhibitor activity (GO:0019870) | 0.004492 | 0.02089 | 277.53 | 1500.17 |
| 5 | Tat protein binding (GO:0030957) | 0.004990 | 0.02089 | 246.68 | 1307.49 |

GO: Cellular Component

| **Index** | **Name** | **P-value** | **Adjusted p-value** | **Odds Ratio** | **Combined score** |
| --- | --- | --- | --- | --- | --- |
| 1 | filopodium membrane (GO:0031527) | 0.005985 | 0.05387 | 201.81 | 1032.95 |
| 2 | sarcolemma (GO:0042383) | 0.0002944 | 0.007949 | 99.70 | 810.62 |
| 3 | myofibril (GO:0030016) | 0.01342 | 0.06661 | 85.32 | 367.79 |
| 4 | rough endoplasmic reticulum (GO:0005791) | 0.01441 | 0.06661 | 79.21 | 335.86 |
| 5 | neuromuscular junction (GO:0031594) | 0.01884 | 0.06661 | 59.92 | 237.98 |

Pain ([HP:0012531](https://hpo.jax.org/app/browse/term/HP:0012531))

KEGG Pathway

| **Index** | **Name** | **P-value** | **Adjusted p-value** | **Odds Ratio** | **Combined score** |
| --- | --- | --- | --- | --- | --- |
| 1 | Pathways in cancer | 1.006e-11 | 1.479e-9 | 148.89 | 3770.13 |
| 2 | Thyroid cancer | 6.931e-7 | 0.000008491 | 251.55 | 3567.44 |
| 3 | Central carbon metabolism in cancer | 2.843e-8 | 8.638e-7 | 201.25 | 3496.91 |
| 4 | Colorectal cancer | 6.560e-8 | 0.000001607 | 161.85 | 2677.02 |
| 5 | Breast cancer | 4.901e-9 | 3.602e-7 | 139.77 | 2674.44 |

GO: Biological Process

| **Index** | **Name** | **P-value** | **Adjusted p-value** | **Odds Ratio** | **Combined score** |
| --- | --- | --- | --- | --- | --- |
| 1 | positive regulation of histone H3-K4 methylation (GO:0051571) | 0.00002354 | 0.0008257 | 384.17 | 4094.01 |
| 2 | response to muscle stretch (GO:0035994) | 0.00002354 | 0.0008257 | 384.17 | 4094.01 |
| 3 | regulation of phospholipase C activity (GO:1900274) | 0.00002354 | 0.0008257 | 384.17 | 4094.01 |
| 4 | sympathetic nervous system development (GO:0048485) | 0.00002690 | 0.0008402 | 356.71 | 3753.85 |
| 5 | regulation of neuroblast proliferation (GO:1902692) | 0.00003048 | 0.0008402 | 332.92 | 3461.84 |

GO: Molecular Function

| **Index** | **Name** | **P-value** | **Adjusted p-value** | **Odds Ratio** | **Combined score** |
| --- | --- | --- | --- | --- | --- |
| 1 | I-SMAD binding (GO:0070411) | 0.00001750 | 0.0001966 | 454.07 | 4973.59 |
| 2 | RNA polymerase II general transcription initiation factor binding (GO:0001091) | 0.002498 | 0.01526 | 555.17 | 3326.77 |
| 3 | transcription coactivator binding (GO:0001223) | 0.00004255 | 0.0003900 | 277.39 | 2791.90 |
| 4 | G protein-coupled serotonin receptor binding (GO:0031821) | 0.002997 | 0.01602 | 444.11 | 2580.41 |
| 5 | transcription coregulator binding (GO:0001221) | 0.000002081 | 0.00005563 | 170.91 | 2236.02 |

GO: Cellular Component

| **Index** | **Name** | **P-value** | **Adjusted p-value** | **Odds Ratio** | **Combined score** |
| --- | --- | --- | --- | --- | --- |
| 1 | phosphatidylinositol 3-kinase complex, class I (GO:0097651) | 0.002498 | 0.03747 | 555.17 | 3326.77 |
| 2 | beta-catenin-TCF complex (GO:1990907) | 0.006482 | 0.04862 | 184.98 | 932.06 |
| 3 | intercalated disc (GO:0014704) | 0.01540 | 0.05461 | 73.93 | 308.54 |
| 4 | catenin complex (GO:0016342) | 0.01540 | 0.05461 | 73.93 | 308.54 |
| 5 | heterotrimeric G-protein complex (GO:0005834) | 0.01638 | 0.05461 | 69.30 | 284.93 |

Pedal edema ([HP:0010741](https://hpo.jax.org/app/browse/term/HP:0010741))

KEGG Pathway

| **Index** | **Name** | **P-value** | **Adjusted p-value** | **Odds Ratio** | **Combined score** |
| --- | --- | --- | --- | --- | --- |
| 1 | Thyroid hormone signaling pathway | 0.001583 | 0.01940 | 41.75 | 269.21 |
| 2 | cGMP-PKG signaling pathway | 0.002985 | 0.01940 | 30.04 | 174.65 |
| 3 | Viral myocarditis | 0.02960 | 0.07791 | 37.53 | 132.12 |
| 4 | Lysine degradation | 0.03106 | 0.07791 | 35.71 | 123.99 |
| 5 | Cardiac muscle contraction | 0.04267 | 0.07791 | 25.72 | 81.12 |

GO: Biological Process

| **Index** | **Name** | **P-value** | **Adjusted p-value** | **Odds Ratio** | **Combined score** |
| --- | --- | --- | --- | --- | --- |
| 1 | positive regulation of cardioblast differentiation (GO:0051891) | 0.000002248 | 0.00003414 | 1665.58 | 21661.60 |
| 2 | regulation of cardioblast differentiation (GO:0051890) | 0.000002248 | 0.00003414 | 1665.58 | 21661.60 |
| 3 | cardiac atrium morphogenesis (GO:0003209) | 1.217e-10 | 2.956e-8 | 887.78 | 20267.77 |
| 4 | atrial septum morphogenesis (GO:0060413) | 1.482e-8 | 0.000001200 | 1070.46 | 19297.66 |
| 5 | negative regulation of cardiac muscle cell apoptotic process (GO:0010667) | 0.000003371 | 0.00004819 | 1249.13 | 15739.23 |

GO: Molecular Function

| **Index** | **Name** | **P-value** | **Adjusted p-value** | **Odds Ratio** | **Combined score** |
| --- | --- | --- | --- | --- | --- |
| 1 | aromatic amino acid transmembrane transporter activity (GO:0015173) | 0.004492 | 0.01206 | 277.53 | 1500.17 |
| 2 | co-SMAD binding (GO:0070410) | 0.004990 | 0.01206 | 246.68 | 1307.49 |
| 3 | histone methyltransferase activity (H3-K36 specific) (GO:0046975) | 0.005488 | 0.01224 | 222.00 | 1155.57 |
| 4 | lysine N-methyltransferase activity (GO:0016278) | 0.01045 | 0.02165 | 110.94 | 506.01 |
| 5 | transcription regulatory region nucleic acid binding (GO:0001067) | 0.0001334 | 0.001634 | 40.56 | 361.91 |

GO: Cellular Component

| **Index** | **Name** | **P-value** | **Adjusted p-value** | **Odds Ratio** | **Combined score** |
| --- | --- | --- | --- | --- | --- |
| 1 | U4 snRNP (GO:0005687) | 0.004492 | 0.01971 | 277.53 | 1500.17 |
| 2 | melanosome membrane (GO:0033162) | 0.005985 | 0.01971 | 201.81 | 1032.95 |
| 3 | U2-type catalytic step 1 spliceosome (GO:0071006) | 0.005985 | 0.01971 | 201.81 | 1032.95 |
| 4 | pigment granule membrane (GO:0090741) | 0.005985 | 0.01971 | 201.81 | 1032.95 |
| 5 | chitosome (GO:0045009) | 0.005985 | 0.01971 | 201.81 | 1032.95 |

General-symptom

Arthritis ([HP:0001369](https://hpo.jax.org/app/browse/term/HP:0001369))

KEGG Pathway

| **Index** | **Name** | **P-value** | **Adjusted p-value** | **Odds Ratio** | **Combined score** |
| --- | --- | --- | --- | --- | --- |
| 1 | B cell receptor signaling pathway | 5.146e-8 | 0.000001492 | 172.41 | 2893.43 |
| 2 | Osteoclast differentiation | 0.00002904 | 0.0004211 | 68.66 | 717.28 |
| 3 | Fc epsilon RI signaling pathway | 0.0005036 | 0.004251 | 75.47 | 573.09 |
| 4 | ECM-receptor interaction | 0.0008418 | 0.004251 | 57.86 | 409.65 |
| 5 | Fc gamma R-mediated phagocytosis | 0.001021 | 0.004251 | 52.36 | 360.55 |

GO: Biological Process

| **Index** | **Name** | **P-value** | **Adjusted p-value** | **Odds Ratio** | **Combined score** |
| --- | --- | --- | --- | --- | --- |
| 1 | regulation of neutrophil activation (GO:1902563) | 0.000002248 | 0.0001323 | 1665.58 | 21661.60 |
| 2 | B cell receptor signaling pathway (GO:0050853) | 1.451e-9 | 3.568e-7 | 443.56 | 9026.96 |
| 3 | positive regulation of B cell differentiation (GO:0045579) | 0.000008085 | 0.0003315 | 713.68 | 8368.26 |
| 4 | negative regulation of B cell proliferation (GO:0030889) | 0.00001481 | 0.0004554 | 499.50 | 5554.53 |
| 5 | negative regulation of dendritic cell differentiation (GO:2001199) | 0.002498 | 0.01707 | 555.17 | 3326.77 |

GO: Molecular Function

| **Index** | **Name** | **P-value** | **Adjusted p-value** | **Odds Ratio** | **Combined score** |
| --- | --- | --- | --- | --- | --- |
| 1 | phospholipase binding (GO:0043274) | 0.00002041 | 0.0004286 | 416.21 | 4494.85 |
| 2 | IgG binding (GO:0019864) | 0.002498 | 0.01259 | 555.17 | 3326.77 |
| 3 | MHC class II protein binding (GO:0042289) | 0.002997 | 0.01259 | 444.11 | 2580.41 |
| 4 | non-membrane spanning protein tyrosine kinase activity (GO:0004715) | 0.0001327 | 0.001393 | 151.19 | 1349.73 |
| 5 | phospholipase activator activity (GO:0016004) | 0.005488 | 0.01571 | 222.00 | 1155.57 |

GO: Cellular Component

| **Index** | **Name** | **P-value** | **Adjusted p-value** | **Odds Ratio** | **Combined score** |
| --- | --- | --- | --- | --- | --- |
| 1 | early phagosome (GO:0032009) | 0.005985 | 0.02155 | 201.81 | 1032.95 |
| 2 | T cell receptor complex (GO:0042101) | 0.005985 | 0.02155 | 201.81 | 1032.95 |
| 3 | collagen-containing extracellular matrix (GO:0062023) | 5.619e-7 | 0.00001011 | 52.31 | 752.79 |
| 4 | endoplasmic reticulum lumen (GO:0005788) | 0.0003190 | 0.002871 | 29.95 | 241.12 |
| 5 | basement membrane (GO:0005604) | 0.02570 | 0.07335 | 43.44 | 159.04 |

Chills ([HP:0025143](https://hpo.jax.org/app/browse/term/HP:0025143))

KEGG Pathway

| **Index** | **Name** | **P-value** | **Adjusted p-value** | **Odds Ratio** | **Combined score** |
| --- | --- | --- | --- | --- | --- |
| 1 | Pertussis | 0.02259 | 0.07242 | 53.12 | 201.33 |
| 2 | C-type lectin receptor signaling pathway | 0.03080 | 0.07242 | 38.62 | 134.42 |
| 3 | Yersinia infection | 0.04041 | 0.07242 | 29.20 | 93.70 |
| 4 | Apoptosis | 0.04186 | 0.07242 | 28.16 | 89.37 |
| 5 | Necroptosis | 0.04677 | 0.07242 | 25.11 | 76.90 |

GO: Biological Process

| **Index** | **Name** | **P-value** | **Adjusted p-value** | **Odds Ratio** | **Combined score** |
| --- | --- | --- | --- | --- | --- |
| 1 | positive regulation of protein localization to cell cortex (GO:1904778) | 0.001499 | 0.02006 | 999.50 | 6499.56 |
| 2 | positive regulation of T-helper 2 cell differentiation (GO:0045630) | 0.001799 | 0.02006 | 799.56 | 5053.71 |
| 3 | regulation of digestive system process (GO:0044058) | 0.001799 | 0.02006 | 799.56 | 5053.71 |
| 4 | regulation of type 2 immune response (GO:0002828) | 0.001799 | 0.02006 | 799.56 | 5053.71 |
| 5 | establishment of centrosome localization (GO:0051660) | 0.002098 | 0.02006 | 666.27 | 4108.59 |

GO: Molecular Function

| **Index** | **Name** | **P-value** | **Adjusted p-value** | **Odds Ratio** | **Combined score** |
| --- | --- | --- | --- | --- | --- |
| 1 | protein-glutamine gamma-glutamyltransferase activity (GO:0003810) | 0.002697 | 0.01947 | 499.65 | 2955.69 |
| 2 | 1-phosphatidylinositol binding (GO:0005545) | 0.004492 | 0.01947 | 285.43 | 1542.87 |
| 3 | peptidoglycan binding (GO:0042834) | 0.004492 | 0.01947 | 285.43 | 1542.87 |
| 4 | ATPase binding (GO:0051117) | 0.02170 | 0.06749 | 55.34 | 211.96 |
| 5 | phosphatidylinositol binding (GO:0035091) | 0.02729 | 0.06749 | 43.74 | 157.53 |

GO: Cellular Component

| **Index** | **Name** | **P-value** | **Adjusted p-value** | **Odds Ratio** | **Combined score** |
| --- | --- | --- | --- | --- | --- |
| 1 | spectrin-associated cytoskeleton (GO:0014731) | 8.395e-10 | 1.679e-8 | 3997.80 | 83546.98 |
| 2 | intrinsic component of the cytoplasmic side of the plasma membrane (GO:0031235) | 0.002398 | 0.006851 | 571.06 | 3445.29 |
| 3 | cell cortex region (GO:0099738) | 0.003296 | 0.007538 | 399.68 | 2284.21 |
| 4 | cytoplasmic side of plasma membrane (GO:0009898) | 0.0001106 | 0.0007373 | 188.12 | 1713.74 |
| 5 | anchored component of external side of plasma membrane (GO:0031362) | 0.005986 | 0.01197 | 210.26 | 1076.21 |

Constitutional symptom ([HP:0025142](https://hpo.jax.org/app/browse/term/HP:0025142))

KEGG Pathway

| **Index** | **Name** | **P-value** | **Adjusted p-value** | **Odds Ratio** | **Combined score** |
| --- | --- | --- | --- | --- | --- |
| 1 | Cardiac muscle contraction | 2.733e-15 | 4.192e-14 | 580.71 | 19473.01 |
| 2 | Hypertrophic cardiomyopathy | 3.494e-15 | 4.192e-14 | 559.63 | 18629.01 |
| 3 | Dilated cardiomyopathy | 5.569e-15 | 4.455e-14 | 521.75 | 17124.60 |
| 4 | Adrenergic signaling in cardiomyocytes | 1.365e-13 | 8.188e-13 | 323.84 | 9593.14 |
| 5 | Viral myocarditis | 0.0003921 | 0.001569 | 85.91 | 673.90 |

GO: Biological Process

| **Index** | **Name** | **P-value** | **Adjusted p-value** | **Odds Ratio** | **Combined score** |
| --- | --- | --- | --- | --- | --- |
| 1 | heart contraction (GO:0060047) | 2.480e-24 | 4.242e-22 | 5613.19 | 305096.95 |
| 2 | cardiac muscle contraction (GO:0060048) | 1.284e-21 | 1.098e-19 | 3071.38 | 147746.28 |
| 3 | striated muscle contraction (GO:0006941) | 7.348e-20 | 4.188e-18 | 1734.26 | 76406.89 |
| 4 | ventricular cardiac muscle tissue morphogenesis (GO:0055010) | 2.534e-18 | 1.083e-16 | 1725.20 | 69899.68 |
| 5 | ventricular cardiac muscle tissue development (GO:0003229) | 4.171e-16 | 1.426e-14 | 1576.66 | 55834.51 |

GO: Molecular Function

| **Index** | **Name** | **P-value** | **Adjusted p-value** | **Odds Ratio** | **Combined score** |
| --- | --- | --- | --- | --- | --- |
| 1 | myosin heavy chain binding (GO:0032036) | 3.147e-9 | 1.070e-7 | 2141.36 | 41921.23 |
| 2 | voltage-gated sodium channel activity involved in cardiac muscle cell action potential (GO:0086006) | 0.002498 | 0.009436 | 555.17 | 3326.77 |
| 3 | nitric-oxide synthase binding (GO:0050998) | 0.003495 | 0.01188 | 370.07 | 2093.27 |
| 4 | actin monomer binding (GO:0003785) | 0.00006709 | 0.0006402 | 217.03 | 2085.58 |
| 5 | calcium channel inhibitor activity (GO:0019855) | 0.005488 | 0.01435 | 222.00 | 1155.57 |

GO: Cellular Component

| **Index** | **Name** | **P-value** | **Adjusted p-value** | **Odds Ratio** | **Combined score** |
| --- | --- | --- | --- | --- | --- |
| 1 | cardiac myofibril (GO:0097512) | 8.995e-10 | 1.394e-8 | 4283.14 | 89214.48 |
| 2 | myofibril (GO:0030016) | 7.595e-13 | 2.354e-11 | 907.64 | 25328.60 |
| 3 | muscle myosin complex (GO:0005859) | 4.082e-8 | 4.218e-7 | 713.50 | 12139.53 |
| 4 | junctional sarcoplasmic reticulum membrane (GO:0014701) | 0.000008085 | 0.00006266 | 713.68 | 8368.26 |
| 5 | myosin filament (GO:0032982) | 0.002997 | 0.009289 | 444.11 | 2580.41 |

Difficulty walking ([HP:0002355](https://hpo.jax.org/app/browse/term/HP:0002355))

KEGG Pathway

| **Index** | **Name** | **P-value** | **Adjusted p-value** | **Odds Ratio** | **Combined score** |
| --- | --- | --- | --- | --- | --- |
| 1 | Lysosome | 0.00002973 | 0.0001189 | 68.11 | 709.91 |
| 2 | Endocytosis | 0.1191 | 0.2137 | 8.74 | 18.59 |
| 3 | Amyotrophic lateral sclerosis | 0.1678 | 0.2137 | 6.01 | 10.72 |
| 4 | Pathways of neurodegeneration | 0.2137 | 0.2137 | 4.57 | 7.06 |

GO: Biological Process

| **Index** | **Name** | **P-value** | **Adjusted p-value** | **Odds Ratio** | **Combined score** |
| --- | --- | --- | --- | --- | --- |
| 1 | axo-dendritic transport (GO:0008088) | 0.002997 | 0.02576 | 444.11 | 2580.41 |
| 2 | lipid droplet formation (GO:0140042) | 0.004990 | 0.02576 | 246.68 | 1307.49 |
| 3 | synaptic vesicle transport (GO:0048489) | 0.005488 | 0.02576 | 222.00 | 1155.57 |
| 4 | post-Golgi vesicle-mediated transport (GO:0006892) | 0.000009993 | 0.0003110 | 99.19 | 1142.03 |
| 5 | transport along microtubule (GO:0010970) | 0.006482 | 0.02576 | 184.98 | 932.06 |

GO: Molecular Function

| **Index** | **Name** | **P-value** | **Adjusted p-value** | **Odds Ratio** | **Combined score** |
| --- | --- | --- | --- | --- | --- |
| 1 | gap junction channel activity involved in cell communication by electrical coupling (GO:1903763) | 0.002997 | 0.01065 | 444.11 | 2580.41 |
| 2 | olfactory receptor binding (GO:0031849) | 0.002997 | 0.01065 | 444.11 | 2580.41 |
| 3 | CD4 receptor binding (GO:0042609) | 0.003994 | 0.01065 | 317.19 | 1751.86 |
| 4 | gap junction channel activity (GO:0005243) | 0.01194 | 0.01989 | 96.46 | 427.12 |
| 5 | wide pore channel activity (GO:0022829) | 0.01243 | 0.01989 | 92.44 | 405.55 |

GO: Cellular Component

| **Index** | **Name** | **P-value** | **Adjusted p-value** | **Odds Ratio** | **Combined score** |
| --- | --- | --- | --- | --- | --- |
| 1 | endosome lumen (GO:0031904) | 2.326e-7 | 0.000007211 | 372.06 | 5682.76 |
| 2 | trans-Golgi network membrane (GO:0032588) | 0.00001377 | 0.0002134 | 88.81 | 994.11 |
| 3 | trans-Golgi network transport vesicle (GO:0030140) | 0.007476 | 0.02575 | 158.54 | 776.21 |
| 4 | connexin complex (GO:0005922) | 0.01045 | 0.03084 | 110.94 | 506.01 |
| 5 | endoplasmic reticulum tubular network (GO:0071782) | 0.01144 | 0.03084 | 100.85 | 450.83 |

Exercise intolerance ([HP:0003546](https://hpo.jax.org/app/browse/term/HP:0003546))

KEGG Pathway

| **Index** | **Name** | **P-value** | **Adjusted p-value** | **Odds Ratio** | **Combined score** |
| --- | --- | --- | --- | --- | --- |
| 1 | Glucagon signaling pathway | 1.217e-14 | 2.677e-13 | 464.10 | 14869.80 |
| 2 | Insulin signaling pathway | 3.435e-9 | 3.779e-8 | 150.44 | 2931.94 |
| 3 | Glycolysis / Gluconeogenesis | 0.000004240 | 0.00002332 | 133.43 | 1650.70 |
| 4 | Calcium signaling pathway | 0.000004013 | 0.00002332 | 55.80 | 693.40 |
| 5 | Central carbon metabolism in cancer | 0.0005336 | 0.002348 | 73.24 | 551.94 |

GO: Biological Process

| **Index** | **Name** | **P-value** | **Adjusted p-value** | **Odds Ratio** | **Combined score** |
| --- | --- | --- | --- | --- | --- |
| 1 | glycogen catabolic process (GO:0005980) | 2.832e-14 | 8.923e-13 | 1998.00 | 62328.01 |
| 2 | glucan catabolic process (GO:0009251) | 2.832e-14 | 8.923e-13 | 1998.00 | 62328.01 |
| 3 | cellular polysaccharide catabolic process (GO:0044247) | 4.118e-14 | 8.923e-13 | 1816.27 | 55978.80 |
| 4 | cellular glucan metabolic process (GO:0006073) | 1.038e-11 | 1.349e-10 | 1903.14 | 48133.11 |
| 5 | glycogen metabolic process (GO:0005977) | 9.244e-13 | 1.502e-11 | 868.13 | 24055.55 |

GO: Molecular Function

| **Index** | **Name** | **P-value** | **Adjusted p-value** | **Odds Ratio** | **Combined score** |
| --- | --- | --- | --- | --- | --- |
| 1 | calmodulin-dependent protein kinase activity (GO:0004683) | 1.812e-7 | 0.000003080 | 407.53 | 6326.45 |
| 2 | 1,4-alpha-oligoglucan phosphorylase activity (GO:0004645) | 0.002498 | 0.01485 | 555.17 | 3326.77 |
| 3 | fructose binding (GO:0070061) | 0.002997 | 0.01485 | 444.11 | 2580.41 |
| 4 | phosphofructokinase activity (GO:0008443) | 0.003495 | 0.01485 | 370.07 | 2093.27 |
| 5 | intramolecular transferase activity, phosphotransferases (GO:0016868) | 0.004492 | 0.01527 | 277.53 | 1500.17 |

GO: Cellular Component

| **Index** | **Name** | **P-value** | **Adjusted p-value** | **Odds Ratio** | **Combined score** |
| --- | --- | --- | --- | --- | --- |
| 1 | muscle myosin complex (GO:0005859) | 0.007476 | 0.06711 | 158.54 | 776.21 |
| 2 | myofibril (GO:0030016) | 0.01342 | 0.06711 | 85.32 | 367.79 |
| 3 | filopodium (GO:0030175) | 0.02863 | 0.08148 | 38.86 | 138.07 |
| 4 | actin filament (GO:0005884) | 0.03543 | 0.08148 | 31.17 | 104.12 |
| 5 | actin-based cell projection (GO:0098858) | 0.04074 | 0.08148 | 26.98 | 86.34 |

Fatigue ([HP:0012378)](https://hpo.jax.org/app/browse/term/HP:0012378)

KEGG Pathway

| **Index** | **Name** | **P-value** | **Adjusted p-value** | **Odds Ratio** | **Combined score** |
| --- | --- | --- | --- | --- | --- |
| 1 | Thyroid cancer | 6.931e-7 | 0.00001135 | 251.55 | 3567.44 |
| 2 | Central carbon metabolism in cancer | 2.843e-8 | 0.000001862 | 201.25 | 3496.91 |
| 3 | Colorectal cancer | 6.560e-8 | 0.000002148 | 161.85 | 2677.02 |
| 4 | Breast cancer | 4.901e-9 | 6.420e-7 | 139.77 | 2674.44 |
| 5 | Endometrial cancer | 0.000002737 | 0.00002988 | 155.34 | 1989.64 |

GO: Biological Process

| **Index** | **Name** | **P-value** | **Adjusted p-value** | **Odds Ratio** | **Combined score** |
| --- | --- | --- | --- | --- | --- |
| 1 | positive regulation of histone H3-K9 acetylation (GO:2000617) | 0.000002248 | 0.0001148 | 1665.58 | 21661.60 |
| 2 | response to indole-3-methanol (GO:0071680) | 0.000002248 | 0.0001148 | 1665.58 | 21661.60 |
| 3 | cellular response to indole-3-methanol (GO:0071681) | 0.000002248 | 0.0001148 | 1665.58 | 21661.60 |
| 4 | positive regulation of histone H3-K4 methylation (GO:0051571) | 4.082e-8 | 0.00001065 | 713.50 | 12139.53 |
| 5 | regulation of histone H3-K4 methylation (GO:0051569) | 8.685e-8 | 0.00001511 | 535.02 | 8698.92 |

GO: Molecular Function

| **Index** | **Name** | **P-value** | **Adjusted p-value** | **Odds Ratio** | **Combined score** |
| --- | --- | --- | --- | --- | --- |
| 1 | I-SMAD binding (GO:0070411) | 0.00001750 | 0.0002458 | 454.07 | 4973.59 |
| 2 | DNA insertion or deletion binding (GO:0032135) | 0.002498 | 0.01717 | 555.17 | 3326.77 |
| 3 | 1-phosphatidylinositol-4-phosphate 3-kinase activity (GO:0035005) | 0.003495 | 0.01748 | 370.07 | 2093.27 |
| 4 | MutSalpha complex binding (GO:0032407) | 0.003495 | 0.01748 | 370.07 | 2093.27 |
| 5 | transmembrane receptor protein kinase activity (GO:0019199) | 0.000003034 | 0.00008344 | 149.87 | 1904.21 |

GO: Cellular Component

| **Index** | **Name** | **P-value** | **Adjusted p-value** | **Odds Ratio** | **Combined score** |
| --- | --- | --- | --- | --- | --- |
| 1 | phosphatidylinositol 3-kinase complex, class I (GO:0097651) | 0.002498 | 0.03497 | 555.17 | 3326.77 |
| 2 | beta-catenin-TCF complex (GO:1990907) | 0.006482 | 0.04538 | 184.98 | 932.06 |
| 3 | intercalated disc (GO:0014704) | 0.01540 | 0.04881 | 73.93 | 308.54 |
| 4 | catenin complex (GO:0016342) | 0.01540 | 0.04881 | 73.93 | 308.54 |
| 5 | basolateral plasma membrane (GO:0016323) | 0.002449 | 0.03497 | 33.29 | 200.15 |

Impairment of activities of daily living ([HP:0031058](https://hpo.jax.org/app/browse/term/HP:0031058))

KEGG Pathway

| **Index** | **Name** | **P-value** | **Adjusted p-value** | **Odds Ratio** | **Combined score** |
| --- | --- | --- | --- | --- | --- |
| 1 | Nicotine addiction | 0.01983 | 0.1019 | 56.84 | 222.86 |
| 2 | Non-small cell lung cancer | 0.03543 | 0.1019 | 31.17 | 104.12 |
| 3 | Taste transduction | 0.04219 | 0.1019 | 26.02 | 82.37 |
| 4 | GABAergic synapse | 0.04363 | 0.1019 | 25.13 | 78.70 |
| 5 | Morphine addiction | 0.04459 | 0.1019 | 24.57 | 76.41 |

GO: Biological Process

| **Index** | **Name** | **P-value** | **Adjusted p-value** | **Odds Ratio** | **Combined score** |
| --- | --- | --- | --- | --- | --- |
| 1 | synaptic vesicle transport (GO:0048489) | 0.00001235 | 0.0003498 | 555.03 | 6273.06 |
| 2 | synaptic vesicle localization (GO:0097479) | 0.00001750 | 0.0003718 | 454.07 | 4973.59 |
| 3 | anterograde dendritic transport (GO:0098937) | 0.002498 | 0.01769 | 555.17 | 3326.77 |
| 4 | anterograde dendritic transport of neurotransmitter receptor complex (GO:0098971) | 0.002498 | 0.01769 | 555.17 | 3326.77 |
| 5 | establishment of vesicle localization (GO:0051650) | 0.00003428 | 0.0005827 | 312.09 | 3208.63 |

GO: Molecular Function

| **Index** | **Name** | **P-value** | **Adjusted p-value** | **Odds Ratio** | **Combined score** |
| --- | --- | --- | --- | --- | --- |
| 1 | gap junction channel activity involved in cell communication by electrical coupling (GO:1903763) | 0.002997 | 0.02309 | 444.11 | 2580.41 |
| 2 | benzodiazepine receptor activity (GO:0008503) | 0.004990 | 0.02309 | 246.68 | 1307.49 |
| 3 | extracellular ligand-gated ion channel activity (GO:0005230) | 0.005488 | 0.02309 | 222.00 | 1155.57 |
| 4 | GABA-gated chloride ion channel activity (GO:0022851) | 0.006482 | 0.02309 | 184.98 | 932.06 |
| 5 | inhibitory extracellular ligand-gated ion channel activity (GO:0005237) | 0.006979 | 0.02309 | 170.74 | 847.70 |

GO: Cellular Component

| **Index** | **Name** | **P-value** | **Adjusted p-value** | **Odds Ratio** | **Combined score** |
| --- | --- | --- | --- | --- | --- |
| 1 | endoplasmic reticulum tubular network membrane (GO:0098826) | 0.002498 | 0.04559 | 555.17 | 3326.77 |
| 2 | GABA-A receptor complex (GO:1902711) | 0.009462 | 0.04559 | 123.28 | 574.57 |
| 3 | connexin complex (GO:0005922) | 0.01045 | 0.04559 | 110.94 | 506.01 |
| 4 | endoplasmic reticulum tubular network (GO:0071782) | 0.01144 | 0.04559 | 100.85 | 450.83 |
| 5 | gap junction (GO:0005921) | 0.01194 | 0.04559 | 96.46 | 427.12 |

Postexertional malaise ([HP:0030973](https://hpo.jax.org/app/browse/term/HP:0030973))

KEGG Pathway

| **Index** | **Name** | **P-value** | **Adjusted p-value** | **Odds Ratio** | **Combined score** |
| --- | --- | --- | --- | --- | --- |
| 1 | ECM-receptor interaction | 0.0005268 | 0.007568 | 77.16 | 582.42 |
| 2 | Protein digestion and absorption | 0.0007207 | 0.007568 | 65.65 | 474.97 |
| 3 | Starch and sucrose metabolism | 0.01431 | 0.04412 | 81.46 | 345.92 |
| 4 | African trypanosomiasis | 0.01471 | 0.04412 | 79.19 | 334.14 |
| 5 | Regulation of lipolysis in adipocytes | 0.02179 | 0.05721 | 52.75 | 201.82 |

GO: Biological Process

| **Index** | **Name** | **P-value** | **Adjusted p-value** | **Odds Ratio** | **Combined score** |
| --- | --- | --- | --- | --- | --- |
| 1 | regulation of atrial cardiac muscle cell membrane repolarization (GO:0060372) | 0.000002937 | 0.0003671 | 1332.47 | 16973.13 |
| 2 | urate metabolic process (GO:0046415) | 0.000007686 | 0.0004804 | 740.11 | 8715.63 |
| 3 | positive regulation of action potential (GO:0045760) | 0.001999 | 0.01520 | 713.86 | 4436.86 |
| 4 | bundle of His cell action potential (GO:0086043) | 0.001999 | 0.01520 | 713.86 | 4436.86 |
| 5 | response to denervation involved in regulation of muscle adaptation (GO:0014894) | 0.001999 | 0.01520 | 713.86 | 4436.86 |

GO: Molecular Function

| **Index** | **Name** | **P-value** | **Adjusted p-value** | **Odds Ratio** | **Combined score** |
| --- | --- | --- | --- | --- | --- |
| 1 | urate transmembrane transporter activity (GO:0015143) | 0.000001399 | 0.00003498 | 2221.00 | 29938.36 |
| 2 | 1,4-alpha-oligoglucan phosphorylase activity (GO:0004645) | 0.001999 | 0.01399 | 713.86 | 4436.86 |
| 3 | voltage-gated sodium channel activity involved in cardiac muscle cell action potential (GO:0086006) | 0.001999 | 0.01399 | 713.86 | 4436.86 |
| 4 | carbohydrate:proton symporter activity (GO:0005351) | 0.002797 | 0.01399 | 475.86 | 2797.66 |
| 5 | nitric-oxide synthase binding (GO:0050998) | 0.002797 | 0.01399 | 475.86 | 2797.66 |

GO: Cellular Component

| **Index** | **Name** | **P-value** | **Adjusted p-value** | **Odds Ratio** | **Combined score** |
| --- | --- | --- | --- | --- | --- |
| 1 | basement membrane (GO:0005604) | 0.0001838 | 0.004082 | 132.95 | 1143.57 |
| 2 | voltage-gated sodium channel complex (GO:0001518) | 0.006781 | 0.03899 | 178.36 | 890.65 |
| 3 | sodium channel complex (GO:0034706) | 0.009958 | 0.04581 | 118.86 | 547.86 |
| 4 | intercalated disc (GO:0014704) | 0.01233 | 0.04728 | 95.06 | 417.81 |
| 5 | brush border membrane (GO:0031526) | 0.01471 | 0.04832 | 79.19 | 334.14 |

Shivering ([HP:0025144](https://hpo.jax.org/app/browse/term/HP:0025144))

KEGG Pathway

| **Index** | **Name** | **P-value** | **Adjusted p-value** | **Odds Ratio** | **Combined score** |
| --- | --- | --- | --- | --- | --- |
| 1 | Neuroactive ligand-receptor interaction | 1.080e-15 | 7.557e-15 | 532.90 | 18364.79 |
| 2 | Cholinergic synapse | 1.297e-9 | 3.027e-9 | 184.09 | 3767.09 |
| 3 | Nicotine addiction | 8.806e-7 | 0.000001541 | 231.12 | 3222.36 |
| 4 | Chemical carcinogenesis | 5.518e-10 | 1.931e-9 | 127.19 | 2711.44 |
| 5 | Calcium signaling pathway | 0.1137 | 0.1592 | 9.18 | 19.96 |

GO: Biological Process

| **Index** | **Name** | **P-value** | **Adjusted p-value** | **Odds Ratio** | **Combined score** |
| --- | --- | --- | --- | --- | --- |
| 1 | synaptic transmission, cholinergic (GO:0007271) | 5.352e-24 | 7.011e-22 | 7265.09 | 389297.06 |
| 2 | response to acetylcholine (GO:1905144) | 8.995e-10 | 1.964e-8 | 4283.14 | 89214.48 |
| 3 | anterograde trans-synaptic signaling (GO:0098916) | 5.108e-17 | 3.346e-15 | 756.57 | 28381.44 |
| 4 | cellular response to acetylcholine (GO:1905145) | 1.078e-8 | 2.017e-7 | 1223.45 | 22444.87 |
| 5 | chemical synaptic transmission (GO:0007268) | 4.030e-16 | 1.760e-14 | 596.76 | 21153.59 |

GO: Molecular Function

| **Index** | **Name** | **P-value** | **Adjusted p-value** | **Odds Ratio** | **Combined score** |
| --- | --- | --- | --- | --- | --- |
| 1 | acetylcholine-gated cation-selective channel activity (GO:0022848) | 1.169e-26 | 2.571e-25 | 26649.33 | 1591265.17 |
| 2 | excitatory extracellular ligand-gated ion channel activity (GO:0005231) | 1.441e-23 | 1.057e-22 | 6146.77 | 323285.26 |
| 3 | postsynaptic neurotransmitter receptor activity (GO:0098960) | 1.441e-23 | 1.057e-22 | 6146.77 | 323285.26 |
| 4 | acetylcholine receptor activity (GO:0015464) | 2.264e-23 | 1.245e-22 | 5707.43 | 297599.13 |
| 5 | transmitter-gated ion channel activity involved in regulation of postsynaptic membrane potential (GO:1904315) | 4.141e-22 | 1.822e-21 | 3630.55 | 178753.16 |

GO: Cellular Component

| **Index** | **Name** | **P-value** | **Adjusted p-value** | **Odds Ratio** | **Combined score** |
| --- | --- | --- | --- | --- | --- |
| 1 | acetylcholine-gated channel complex (GO:0005892) | 2.127e-25 | 3.190e-24 | 13322.67 | 756859.90 |
| 2 | ion channel complex (GO:0034702) | 1.665e-21 | 1.248e-20 | 2957.48 | 141499.80 |
| 3 | neuron projection (GO:0043005) | 9.076e-14 | 4.538e-13 | 319.90 | 9606.86 |
| 4 | integral component of plasma membrane (GO:0005887) | 5.182e-10 | 1.943e-9 | 115.51 | 2469.57 |
| 5 | plasma membrane raft (GO:0044853) | 0.000007807 | 0.00002342 | 108.02 | 1270.32 |

Weight loss ([HP:0001824](https://hpo.jax.org/app/browse/term/HP:0001824))

KEGG Pathway

| **Index** | **Name** | **P-value** | **Adjusted p-value** | **Odds Ratio** | **Combined score** |
| --- | --- | --- | --- | --- | --- |
| 1 | Fanconi anemia pathway | 8.485e-27 | 4.242e-26 | 199460.00 | 11973883.48 |
| 2 | Homologous recombination | 0.02032 | 0.05079 | 55.42 | 215.92 |
| 3 | Pancreatic cancer | 0.03736 | 0.06227 | 29.50 | 96.98 |
| 4 | Ubiquitin mediated proteolysis | 0.06785 | 0.08481 | 15.87 | 42.69 |
| 5 | Pathways in cancer | 0.2360 | 0.2360 | 4.08 | 5.89 |

GO: Biological Process

| **Index** | **Name** | **P-value** | **Adjusted p-value** | **Odds Ratio** | **Combined score** |
| --- | --- | --- | --- | --- | --- |
| 1 | interstrand cross-link repair (GO:0036297) | 1.037e-26 | 7.259e-25 | 199450.00 | 11933257.10 |
| 2 | DNA repair (GO:0006281) | 4.636e-19 | 1.623e-17 | 197020.00 | 8317234.26 |
| 3 | double-strand break repair via synthesis-dependent strand annealing (GO:0045003) | 0.000002248 | 0.00003147 | 1665.58 | 21661.60 |
| 4 | positive regulation of protein monoubiquitination (GO:1902527) | 0.002498 | 0.01165 | 555.17 | 3326.77 |
| 5 | positive regulation of DNA ligation (GO:0051106) | 0.002498 | 0.01165 | 555.17 | 3326.77 |

GO: Molecular Function

| **Index** | **Name** | **P-value** | **Adjusted p-value** | **Odds Ratio** | **Combined score** |
| --- | --- | --- | --- | --- | --- |
| 1 | DNA polymerase binding (GO:0070182) | 0.00003428 | 0.0006856 | 312.09 | 3208.63 |
| 2 | four-way junction helicase activity (GO:0009378) | 0.002997 | 0.02703 | 444.11 | 2580.41 |
| 3 | 3'-5' DNA helicase activity (GO:0043138) | 0.007973 | 0.02703 | 147.96 | 714.91 |
| 4 | four-way junction DNA binding (GO:0000400) | 0.008469 | 0.02703 | 138.71 | 661.82 |
| 5 | single-stranded DNA helicase activity (GO:0017116) | 0.009462 | 0.02703 | 123.28 | 574.57 |

GO: Cellular Component

| **Index** | **Name** | **P-value** | **Adjusted p-value** | **Odds Ratio** | **Combined score** |
| --- | --- | --- | --- | --- | --- |
| 1 | condensed chromosome (GO:0000793) | 0.0003175 | 0.001508 | 95.86 | 772.11 |
| 2 | condensed nuclear chromosome (GO:0000794) | 0.009957 | 0.01713 | 116.79 | 538.34 |
| 3 | chromosome (GO:0005694) | 0.002744 | 0.008232 | 31.38 | 185.09 |
| 4 | nuclear lumen (GO:0031981) | 0.0003351 | 0.001508 | 17.32 | 138.56 |
| 5 | nuclear chromosome (GO:0000228) | 0.04074 | 0.04074 | 26.98 | 86.34 |

Xerostomia ([HP:0000217](https://hpo.jax.org/app/browse/term/HP:0000217))

KEGG Pathway

| **Index** | **Name** | **P-value** | **Adjusted p-value** | **Odds Ratio** | **Combined score** |
| --- | --- | --- | --- | --- | --- |
| 1 | Amyotrophic lateral sclerosis | 6.887e-9 | 6.199e-8 | 82.26 | 1545.90 |
| 2 | mRNA surveillance pathway | 0.001042 | 0.003127 | 51.81 | 355.72 |
| 3 | Pathways of neurodegeneration | 0.00005889 | 0.0002650 | 27.63 | 269.09 |
| 4 | Basal transcription factors | 0.02228 | 0.03342 | 50.37 | 191.61 |
| 5 | RNA transport | 0.003686 | 0.007059 | 26.91 | 150.79 |

GO: Biological Process

| **Index** | **Name** | **P-value** | **Adjusted p-value** | **Odds Ratio** | **Combined score** |
| --- | --- | --- | --- | --- | --- |
| 1 | regulation of cytoplasmic mRNA processing body assembly (GO:0010603) | 0.003495 | 0.03848 | 370.07 | 2093.27 |
| 2 | intermediate filament bundle assembly (GO:0045110) | 0.003495 | 0.03848 | 370.07 | 2093.27 |
| 3 | regulation of translational termination (GO:0006449) | 0.003994 | 0.03848 | 317.19 | 1751.86 |
| 4 | negative regulation of mRNA catabolic process (GO:1902373) | 0.00009070 | 0.005179 | 184.84 | 1720.52 |
| 5 | RNA stabilization (GO:0043489) | 0.0001178 | 0.005179 | 160.96 | 1456.09 |

GO: Molecular Function

| **Index** | **Name** | **P-value** | **Adjusted p-value** | **Odds Ratio** | **Combined score** |
| --- | --- | --- | --- | --- | --- |
| 1 | epidermal growth factor receptor binding (GO:0005154) | 0.01293 | 0.04721 | 88.73 | 385.85 |
| 2 | kinesin binding (GO:0019894) | 0.01441 | 0.04721 | 79.21 | 335.86 |
| 3 | miRNA binding (GO:0035198) | 0.01490 | 0.04721 | 76.48 | 321.69 |
| 4 | regulatory RNA binding (GO:0061980) | 0.01983 | 0.04721 | 56.84 | 222.86 |
| 5 | cyclin-dependent protein serine/threonine kinase regulator activity (GO:0016538) | 0.02179 | 0.04721 | 51.54 | 197.22 |

GO: Cellular Component

| **Index** | **Name** | **P-value** | **Adjusted p-value** | **Odds Ratio** | **Combined score** |
| --- | --- | --- | --- | --- | --- |
| 1 | neurofibrillary tangle (GO:0097418) | 0.002498 | 0.03372 | 555.17 | 3326.77 |
| 2 | nuclear inner membrane (GO:0005637) | 0.01392 | 0.08047 | 82.15 | 351.18 |
| 3 | cyclin-dependent protein kinase holoenzyme complex (GO:0000307) | 0.01490 | 0.08047 | 76.48 | 321.69 |
| 4 | serine/threonine protein kinase complex (GO:1902554) | 0.01835 | 0.08258 | 61.59 | 246.23 |
| 5 | SCF ubiquitin ligase complex (GO:0019005) | 0.02863 | 0.09043 | 38.86 | 138.07 |
