## Supplementary Table 1-3 for "Long COVID: G Protein-Coupled Receptors (GPCRs) responsible for persistent post-COVID symptoms": Supplementary File 2.7-Cardiovascular.docx

Abnormal heart rate variability ([HP:0031860](https://hpo.jax.org/app/browse/term/HP:0031860))

KEGG Pathway

| **Index** | **Name** | **P-value** | **Adjusted p-value** | **Odds Ratio** | **Combined score** |
| --- | --- | --- | --- | --- | --- |
| 1 | Hedgehog signaling pathway | 0.000002460 | 0.00005167 | 161.22 | 2082.13 |
| 2 | Basal cell carcinoma | 0.0004323 | 0.004540 | 81.68 | 632.69 |
| 3 | TGF-beta signaling pathway | 0.0009597 | 0.006718 | 54.07 | 375.73 |
| 4 | Gastric cancer | 0.002386 | 0.01002 | 33.75 | 203.77 |
| 5 | Melanoma | 0.03543 | 0.1240 | 31.17 | 104.12 |

GO: Biological Process

| **Index** | **Name** | **P-value** | **Adjusted p-value** | **Odds Ratio** | **Combined score** |
| --- | --- | --- | --- | --- | --- |
| 1 | regulation of nodal signaling pathway involved in determination of lateral mesoderm left/right asymmetry (GO:1900175) | 0.000002248 | 0.00009367 | 1665.58 | 21661.60 |
| 2 | positive regulation of T cell differentiation in thymus (GO:0033089) | 0.000003371 | 0.0001204 | 1249.13 | 15739.23 |
| 3 | dorsal/ventral pattern formation (GO:0009953) | 3.331e-10 | 8.328e-8 | 665.67 | 14526.48 |
| 4 | pituitary gland development (GO:0021983) | 0.000004719 | 0.0001263 | 999.25 | 12254.80 |
| 5 | proximal/distal pattern formation (GO:0009954) | 0.000006290 | 0.0001430 | 832.67 | 9972.50 |

GO: Molecular Function

| **Index** | **Name** | **P-value** | **Adjusted p-value** | **Odds Ratio** | **Combined score** |
| --- | --- | --- | --- | --- | --- |
| 1 | morphogen activity (GO:0016015) | 0.000004719 | 0.0001321 | 999.25 | 12254.80 |
| 2 | type 1 fibroblast growth factor receptor binding (GO:0005105) | 0.002498 | 0.006994 | 555.17 | 3326.77 |
| 3 | type 2 fibroblast growth factor receptor binding (GO:0005111) | 0.002498 | 0.006994 | 555.17 | 3326.77 |
| 4 | patched binding (GO:0005113) | 0.003495 | 0.008897 | 370.07 | 2093.27 |
| 5 | activin receptor binding (GO:0070697) | 0.004990 | 0.01164 | 246.68 | 1307.49 |

GO: Cellular Component

| **Index** | **Name** | **P-value** | **Adjusted p-value** | **Odds Ratio** | **Combined score** |
| --- | --- | --- | --- | --- | --- |
| 1 | condensed nuclear chromosome (GO:0000794) | 0.009957 | 0.1025 | 116.79 | 538.34 |
| 2 | condensed chromosome (GO:0000793) | 0.02668 | 0.1245 | 41.80 | 151.46 |
| 3 | nuclear chromosome (GO:0000228) | 0.04074 | 0.1426 | 26.98 | 86.34 |
| 4 | collagen-containing extracellular matrix (GO:0062023) | 0.01465 | 0.1025 | 12.97 | 54.78 |
| 5 | chromosome (GO:0005694) | 0.07720 | 0.1834 | 13.86 | 35.50 |

Abnormal left ventricular function ([HP:0005162](https://hpo.jax.org/app/browse/term/HP:0005162))

KEGG Pathway

| **Index** | **Name** | **P-value** | **Adjusted p-value** | **Odds Ratio** | **Combined score** |
| --- | --- | --- | --- | --- | --- |
| 1 | Adrenergic signaling in cardiomyocytes | 6.163e-7 | 0.000009942 | 90.61 | 1295.71 |
| 2 | Cardiac muscle contraction | 0.000009332 | 0.00005271 | 101.56 | 1176.29 |
| 3 | Hypertrophic cardiomyopathy | 0.00001034 | 0.00005271 | 98.04 | 1125.55 |
| 4 | cGMP-PKG signaling pathway | 9.469e-7 | 0.000009942 | 81.09 | 1124.75 |
| 5 | Dilated cardiomyopathy | 0.00001255 | 0.00005271 | 91.69 | 1034.81 |

GO: Biological Process

| **Index** | **Name** | **P-value** | **Adjusted p-value** | **Odds Ratio** | **Combined score** |
| --- | --- | --- | --- | --- | --- |
| 1 | adult heart development (GO:0007512) | 7.548e-9 | 1.446e-7 | 1427.43 | 26695.75 |
| 2 | positive regulation of cardioblast differentiation (GO:0051891) | 0.000002248 | 0.00002073 | 1665.58 | 21661.60 |
| 3 | regulation of cardioblast differentiation (GO:0051890) | 0.000002248 | 0.00002073 | 1665.58 | 21661.60 |
| 4 | cardiac atrium morphogenesis (GO:0003209) | 1.217e-10 | 6.059e-9 | 887.78 | 20267.77 |
| 5 | cardiac muscle tissue morphogenesis (GO:0055008) | 1.893e-12 | 4.713e-10 | 739.37 | 19957.87 |

GO: Molecular Function

| **Index** | **Name** | **P-value** | **Adjusted p-value** | **Odds Ratio** | **Combined score** |
| --- | --- | --- | --- | --- | --- |
| 1 | voltage-gated sodium channel activity involved in cardiac muscle cell action potential (GO:0086006) | 0.002498 | 0.01343 | 555.17 | 3326.77 |
| 2 | nitric-oxide synthase binding (GO:0050998) | 0.003495 | 0.01343 | 370.07 | 2093.27 |
| 3 | co-SMAD binding (GO:0070410) | 0.004990 | 0.01343 | 246.68 | 1307.49 |
| 4 | calcium channel inhibitor activity (GO:0019855) | 0.005488 | 0.01372 | 222.00 | 1155.57 |
| 5 | muscle alpha-actinin binding (GO:0051371) | 0.007476 | 0.01744 | 158.54 | 776.21 |

GO: Cellular Component

| **Index** | **Name** | **P-value** | **Adjusted p-value** | **Odds Ratio** | **Combined score** |
| --- | --- | --- | --- | --- | --- |
| 1 | myofibril (GO:0030016) | 2.616e-7 | 0.000006279 | 356.54 | 5403.81 |
| 2 | muscle myosin complex (GO:0005859) | 0.00002354 | 0.0002825 | 384.17 | 4094.01 |
| 3 | cardiac myofibril (GO:0097512) | 0.002498 | 0.01798 | 555.17 | 3326.77 |
| 4 | myosin filament (GO:0032982) | 0.002997 | 0.01798 | 444.11 | 2580.41 |
| 5 | voltage-gated sodium channel complex (GO:0001518) | 0.008469 | 0.03785 | 138.71 | 661.82 |

Abnormal pericardium morphology ([HP:0001697](https://hpo.jax.org/app/browse/term/HP:0001697))

KEGG Pathway

| **Index** | **Name** | **P-value** | **Adjusted p-value** | **Odds Ratio** | **Combined score** |
| --- | --- | --- | --- | --- | --- |
| 1 | Autoimmune thyroid disease | 0.000002081 | 0.0001145 | 170.91 | 2236.02 |
| 2 | Allograft rejection | 0.0001567 | 0.002763 | 138.57 | 1214.06 |
| 3 | Graft-versus-host disease | 0.0001917 | 0.002763 | 124.69 | 1067.29 |
| 4 | Type I diabetes mellitus | 0.0002010 | 0.002763 | 121.64 | 1035.45 |
| 5 | Inflammatory bowel disease | 0.0004602 | 0.005062 | 79.08 | 607.60 |

GO: Biological Process

| **Index** | **Name** | **P-value** | **Adjusted p-value** | **Odds Ratio** | **Combined score** |
| --- | --- | --- | --- | --- | --- |
| 1 | regulation of natural killer cell proliferation (GO:0032817) | 0.000004719 | 0.0002982 | 999.25 | 12254.80 |
| 2 | positive regulation of memory T cell differentiation (GO:0043382) | 0.000008085 | 0.0003505 | 713.68 | 8368.26 |
| 3 | regulation of memory T cell differentiation (GO:0043380) | 0.00001010 | 0.0003536 | 624.44 | 7182.69 |
| 4 | negative regulation of B cell proliferation (GO:0030889) | 0.00001481 | 0.0004712 | 499.50 | 5554.53 |
| 5 | negative regulation of interleukin-10 production (GO:0032693) | 0.00003048 | 0.0008889 | 332.92 | 3461.84 |

GO: Molecular Function

| **Index** | **Name** | **P-value** | **Adjusted p-value** | **Odds Ratio** | **Combined score** |
| --- | --- | --- | --- | --- | --- |
| 1 | IgG binding (GO:0019864) | 0.002498 | 0.01940 | 555.17 | 3326.77 |
| 2 | ATP-activated inward rectifier potassium channel activity (GO:0015272) | 0.002997 | 0.01940 | 444.11 | 2580.41 |
| 3 | interleukin-12 receptor binding (GO:0005143) | 0.002997 | 0.01940 | 444.11 | 2580.41 |
| 4 | non-membrane spanning protein tyrosine phosphatase activity (GO:0004726) | 0.002997 | 0.01940 | 444.11 | 2580.41 |
| 5 | myosin heavy chain binding (GO:0032036) | 0.003495 | 0.01940 | 370.07 | 2093.27 |

GO: Cellular Component

| **Index** | **Name** | **P-value** | **Adjusted p-value** | **Odds Ratio** | **Combined score** |
| --- | --- | --- | --- | --- | --- |
| 1 | cardiac myofibril (GO:0097512) | 0.002498 | 0.02498 | 555.17 | 3326.77 |
| 2 | CD95 death-inducing signaling complex (GO:0031265) | 0.002498 | 0.02498 | 555.17 | 3326.77 |
| 3 | death-inducing signaling complex (GO:0031264) | 0.003994 | 0.02662 | 317.19 | 1751.86 |
| 4 | MHC class II protein complex (GO:0042613) | 0.006482 | 0.03704 | 184.98 | 932.06 |
| 5 | cation-transporting ATPase complex (GO:0090533) | 0.007973 | 0.03986 | 147.96 | 714.91 |

Bradycardia ([HP:0001662](https://hpo.jax.org/app/browse/term/HP:0001662))

KEGG Pathway

| **Index** | **Name** | **P-value** | **Adjusted p-value** | **Odds Ratio** | **Combined score** |
| --- | --- | --- | --- | --- | --- |
| 1 | Cardiac muscle contraction | 0.000009332 | 0.0004106 | 101.56 | 1176.29 |
| 2 | GnRH secretion | 0.0004462 | 0.006544 | 80.35 | 619.92 |
| 3 | Circadian entrainment | 0.001021 | 0.009173 | 52.36 | 360.55 |
| 4 | Aldosterone synthesis and secretion | 0.001042 | 0.009173 | 51.81 | 355.72 |
| 5 | Calcium signaling pathway | 0.0001924 | 0.004233 | 35.72 | 305.61 |

GO: Biological Process

| **Index** | **Name** | **P-value** | **Adjusted p-value** | **Odds Ratio** | **Combined score** |
| --- | --- | --- | --- | --- | --- |
| 1 | membrane depolarization during SA node cell action potential (GO:0086046) | 1.574e-13 | 1.286e-11 | 13326.00 | 392846.55 |
| 2 | SA node cell action potential (GO:0086015) | 1.102e-12 | 4.498e-11 | 4441.56 | 122294.78 |
| 3 | SA node cell to atrial cardiac muscle cell signaling (GO:0086018) | 8.995e-10 | 1.224e-8 | 4283.14 | 89214.48 |
| 4 | cardiac muscle cell action potential (GO:0086001) | 1.397e-15 | 3.423e-13 | 1247.88 | 42682.61 |
| 5 | regulation of cardiac muscle cell membrane repolarization (GO:0099623) | 2.480e-13 | 1.519e-11 | 1174.88 | 34101.37 |

GO: Molecular Function

| **Index** | **Name** | **P-value** | **Adjusted p-value** | **Odds Ratio** | **Combined score** |
| --- | --- | --- | --- | --- | --- |
| 1 | voltage-gated sodium channel activity (GO:0005248) | 1.379e-7 | 0.000005103 | 450.47 | 7115.97 |
| 2 | voltage-gated potassium channel activity involved in ventricular cardiac muscle cell action potential repolarization (GO:1902282) | 0.00001481 | 0.0001370 | 499.50 | 5554.53 |
| 3 | voltage-gated potassium channel activity involved in cardiac muscle cell action potential repolarization (GO:0086008) | 0.00002354 | 0.0001742 | 384.17 | 4094.01 |
| 4 | sodium channel activity (GO:0005272) | 7.523e-7 | 0.00001392 | 244.35 | 3445.31 |
| 5 | voltage-gated calcium channel activity involved in cardiac muscle cell action potential (GO:0086007) | 0.002498 | 0.007701 | 555.17 | 3326.77 |

GO: Cellular Component

| **Index** | **Name** | **P-value** | **Adjusted p-value** | **Odds Ratio** | **Combined score** |
| --- | --- | --- | --- | --- | --- |
| 1 | junctional sarcoplasmic reticulum membrane (GO:0014701) | 0.000008085 | 0.00008624 | 713.68 | 8368.26 |
| 2 | voltage-gated sodium channel complex (GO:0001518) | 0.00003048 | 0.0002438 | 332.92 | 3461.84 |
| 3 | sarcolemma (GO:0042383) | 0.000001964 | 0.00006284 | 174.41 | 2291.88 |
| 4 | sodium channel complex (GO:0034706) | 0.00006709 | 0.0004294 | 217.03 | 2085.58 |
| 5 | sarcoplasmic reticulum membrane (GO:0033017) | 0.00008446 | 0.0004505 | 191.96 | 1800.45 |

Increased circulating troponin I concentration ([HP:0410173](https://hpo.jax.org/app/browse/term/HP:0410173))

KEGG Pathway

| **Index** | **Name** | **P-value** | **Adjusted p-value** | **Odds Ratio** | **Combined score** |
| --- | --- | --- | --- | --- | --- |
| 1 | Vibrio cholerae infection | 0.0002721 | 0.003227 | 103.86 | 852.65 |
| 2 | Viral myocarditis | 0.0003921 | 0.003227 | 85.91 | 673.90 |
| 3 | Adherens junction | 0.0005489 | 0.003227 | 72.18 | 541.87 |
| 4 | Gastric acid secretion | 0.0006287 | 0.003227 | 67.28 | 496.00 |
| 5 | Arrhythmogenic right ventricular cardiomyopathy | 0.0006453 | 0.003227 | 66.38 | 487.64 |

GO: Biological Process

| **Index** | **Name** | **P-value** | **Adjusted p-value** | **Odds Ratio** | **Combined score** |
| --- | --- | --- | --- | --- | --- |
| 1 | protein localization to membrane raft (GO:1903044) | 0.000002248 | 0.0003131 | 1665.58 | 21661.60 |
| 2 | membrane raft assembly (GO:0001765) | 0.000003371 | 0.0003131 | 1249.13 | 15739.23 |
| 3 | regulation of toll-like receptor 3 signaling pathway (GO:0034139) | 0.000006290 | 0.0003560 | 832.67 | 9972.50 |
| 4 | morphogenesis of a polarized epithelium (GO:0001738) | 0.00001481 | 0.0006986 | 499.50 | 5554.53 |
| 5 | membrane raft organization (GO:0031579) | 0.00002354 | 0.0007403 | 384.17 | 4094.01 |

GO: Molecular Function

| **Index** | **Name** | **P-value** | **Adjusted p-value** | **Odds Ratio** | **Combined score** |
| --- | --- | --- | --- | --- | --- |
| 1 | RNA polymerase II general transcription initiation factor binding (GO:0001091) | 0.002498 | 0.03992 | 555.17 | 3326.77 |
| 2 | nitric-oxide synthase binding (GO:0050998) | 0.003495 | 0.03992 | 370.07 | 2093.27 |
| 3 | histone demethylase activity (H3-K4 specific) (GO:0032453) | 0.004492 | 0.03992 | 277.53 | 1500.17 |
| 4 | Tat protein binding (GO:0030957) | 0.004990 | 0.03992 | 246.68 | 1307.49 |
| 5 | histone demethylase activity (H3-K9 specific) (GO:0032454) | 0.006482 | 0.04445 | 184.98 | 932.06 |

GO: Cellular Component

| **Index** | **Name** | **P-value** | **Adjusted p-value** | **Odds Ratio** | **Combined score** |
| --- | --- | --- | --- | --- | --- |
| 1 | flotillin complex (GO:0016600) | 0.000002248 | 0.00001911 | 1665.58 | 21661.60 |
| 2 | actin cytoskeleton (GO:0015629) | 2.954e-9 | 1.004e-7 | 95.23 | 1870.24 |
| 3 | cortical actin cytoskeleton (GO:0030864) | 0.0001917 | 0.0008146 | 124.69 | 1067.29 |
| 4 | cell-cell contact zone (GO:0044291) | 0.0002403 | 0.0008171 | 110.81 | 923.40 |
| 5 | focal adhesion (GO:0005925) | 6.150e-7 | 0.000007616 | 51.33 | 734.10 |

Hypertension ([HP:0000822](https://hpo.jax.org/app/browse/term/HP:0000822))

GO: Biological Process

| **Index** | **Name** | **P-value** | **Adjusted p-value** | **Odds Ratio** | **Combined score** |
| --- | --- | --- | --- | --- | --- |
| 1 | cilium assembly (GO:0060271) | 5.098e-16 | 3.568e-14 | 580.87 | 20453.89 |
| 2 | pigment granule transport (GO:0051904) | 7.315e-8 | 8.534e-7 | 570.71 | 9377.24 |
| 3 | melanosome transport (GO:0032402) | 7.315e-8 | 8.534e-7 | 570.71 | 9377.24 |
| 4 | establishment of melanosome localization (GO:0032401) | 8.685e-8 | 8.685e-7 | 535.02 | 8698.92 |
| 5 | melanosome localization (GO:0032400) | 1.191e-7 | 0.000001042 | 475.52 | 7581.26 |

GO: Molecular Function

| **Index** | **Name** | **P-value** | **Adjusted p-value** | **Odds Ratio** | **Combined score** |
| --- | --- | --- | --- | --- | --- |
| 1 | patched binding (GO:0005113) | 0.003495 | 0.006990 | 370.07 | 2093.27 |
| 2 | RNA polymerase II-specific DNA-binding transcription factor binding (GO:0061629) | 1.780e-8 | 8.402e-8 | 107.05 | 1910.30 |
| 3 | DNA-binding transcription factor binding (GO:0140297) | 2.801e-8 | 8.402e-8 | 97.47 | 1695.14 |
| 4 | phosphatidylinositol-3-phosphate binding (GO:0032266) | 0.01933 | 0.02900 | 58.34 | 230.20 |
| 5 | phosphatidylinositol phosphate binding (GO:1901981) | 0.04986 | 0.05983 | 21.88 | 65.61 |

GO: Cellular Component

| **Index** | **Name** | **P-value** | **Adjusted p-value** | **Odds Ratio** | **Combined score** |
| --- | --- | --- | --- | --- | --- |
| 1 | motile cilium (GO:0031514) | 0.03349 | 0.06698 | 33.04 | 112.22 |
| 2 | neuron projection (GO:0043005) | 0.02994 | 0.06698 | 8.77 | 30.77 |
| 3 | cilium (GO:0005929) | 0.1124 | 0.1276 | 9.30 | 20.33 |
| 4 | cell-cell junction (GO:0005911) | 0.1276 | 0.1276 | 8.12 | 16.71 |

Hypotension ([HP:0002615](https://hpo.jax.org/app/browse/term/HP:0002615))

KEGG Pathway

| **Index** | **Name** | **P-value** | **Adjusted p-value** | **Odds Ratio** | **Combined score** |
| --- | --- | --- | --- | --- | --- |
| 1 | Aldosterone-regulated sodium reabsorption | 6.931e-7 | 0.000007624 | 251.55 | 3567.44 |
| 2 | Renin-angiotensin system | 0.00005661 | 0.0003113 | 237.73 | 2324.81 |
| 3 | Renin secretion | 0.0005185 | 0.001901 | 74.34 | 562.35 |
| 4 | Collecting duct acid secretion | 0.01342 | 0.02953 | 85.32 | 367.79 |
| 5 | Diabetic cardiomyopathy | 0.004372 | 0.01202 | 24.61 | 133.71 |

GO: Biological Process

| **Index** | **Name** | **P-value** | **Adjusted p-value** | **Odds Ratio** | **Combined score** |
| --- | --- | --- | --- | --- | --- |
| 1 | monovalent inorganic anion homeostasis (GO:0055083) | 0.00001010 | 0.00008268 | 624.44 | 7182.69 |
| 2 | chloride ion homeostasis (GO:0055064) | 0.00001010 | 0.00008268 | 624.44 | 7182.69 |
| 3 | sodium ion homeostasis (GO:0055078) | 2.058e-7 | 0.00001151 | 388.99 | 5988.96 |
| 4 | angiotensin maturation (GO:0002003) | 0.00001481 | 0.0001003 | 499.50 | 5554.53 |
| 5 | regulation of angiotensin levels in blood (GO:0002002) | 0.00001481 | 0.0001003 | 499.50 | 5554.53 |

GO: Molecular Function

| **Index** | **Name** | **P-value** | **Adjusted p-value** | **Odds Ratio** | **Combined score** |
| --- | --- | --- | --- | --- | --- |
| 1 | potassium:chloride symporter activity (GO:0015379) | 0.000008085 | 0.0001650 | 713.68 | 8368.26 |
| 2 | sodium:chloride symporter activity (GO:0015378) | 0.000008085 | 0.0001650 | 713.68 | 8368.26 |
| 3 | cation:chloride symporter activity (GO:0015377) | 0.00001010 | 0.0001650 | 624.44 | 7182.69 |
| 4 | anion:sodium symporter activity (GO:0015373) | 0.002498 | 0.01630 | 555.17 | 3326.77 |
| 5 | ATP-activated inward rectifier potassium channel activity (GO:0015272) | 0.002997 | 0.01630 | 444.11 | 2580.41 |

GO: Cellular Component

| **Index** | **Name** | **P-value** | **Adjusted p-value** | **Odds Ratio** | **Combined score** |
| --- | --- | --- | --- | --- | --- |
| 1 | sodium channel complex (GO:0034706) | 0.01243 | 0.07368 | 92.44 | 405.55 |
| 2 | ciliary membrane (GO:0060170) | 0.01540 | 0.07368 | 73.93 | 308.54 |
| 3 | acrosomal vesicle (GO:0001669) | 0.02375 | 0.07368 | 47.15 | 176.34 |
| 4 | motile cilium (GO:0031514) | 0.03349 | 0.07368 | 33.04 | 112.22 |
| 5 | cation channel complex (GO:0034703) | 0.03591 | 0.07368 | 30.74 | 102.25 |

Pericardial effusion ([HP:0001698](https://hpo.jax.org/app/browse/term/HP:0001698))

KEGG Pathway

| **Index** | **Name** | **P-value** | **Adjusted p-value** | **Odds Ratio** | **Combined score** |
| --- | --- | --- | --- | --- | --- |
| 1 | Riboflavin metabolism | 0.003994 | 0.06789 | 317.19 | 1751.86 |
| 2 | ABC transporters | 0.0002202 | 0.007487 | 115.97 | 976.58 |
| 3 | Pantothenate and CoA biosynthesis | 0.01045 | 0.07158 | 110.94 | 506.01 |
| 4 | Collecting duct acid secretion | 0.01342 | 0.07158 | 85.32 | 367.79 |
| 5 | Asthma | 0.01540 | 0.07158 | 73.93 | 308.54 |

GO: Biological Process

| **Index** | **Name** | **P-value** | **Adjusted p-value** | **Odds Ratio** | **Combined score** |
| --- | --- | --- | --- | --- | --- |
| 1 | positive regulation of vascular endothelial growth factor signaling pathway (GO:1900748) | 0.000003371 | 0.0007248 | 1249.13 | 15739.23 |
| 2 | positive regulation of CD4-positive, CD25-positive, alpha-beta regulatory T cell differentiation (GO:0032831) | 0.002498 | 0.02728 | 555.17 | 3326.77 |
| 3 | nucleoside triphosphate catabolic process (GO:0009143) | 0.002498 | 0.02728 | 555.17 | 3326.77 |
| 4 | regulation of vascular endothelial growth factor signaling pathway (GO:1900746) | 0.00003830 | 0.004117 | 293.72 | 2987.15 |
| 5 | cellular trivalent inorganic anion homeostasis (GO:0072502) | 0.002997 | 0.02728 | 444.11 | 2580.41 |

GO: Molecular Function

| **Index** | **Name** | **P-value** | **Adjusted p-value** | **Odds Ratio** | **Combined score** |
| --- | --- | --- | --- | --- | --- |
| 1 | anion:sodium symporter activity (GO:0015373) | 0.002498 | 0.01746 | 555.17 | 3326.77 |
| 2 | phosphodiesterase I activity (GO:0004528) | 0.002498 | 0.01746 | 555.17 | 3326.77 |
| 3 | ATP-activated inward rectifier potassium channel activity (GO:0015272) | 0.002997 | 0.01746 | 444.11 | 2580.41 |
| 4 | hydrolase activity, acting on acid anhydrides, in phosphorus-containing anhydrides (GO:0016818) | 0.003495 | 0.01746 | 370.07 | 2093.27 |
| 5 | ABC-type glutathione S-conjugate transporter activity (GO:0015431) | 0.003495 | 0.01746 | 370.07 | 2093.27 |

GO: Cellular Component

| **Index** | **Name** | **P-value** | **Adjusted p-value** | **Odds Ratio** | **Combined score** |
| --- | --- | --- | --- | --- | --- |
| 1 | cardiac myofibril (GO:0097512) | 0.002498 | 0.04621 | 555.17 | 3326.77 |
| 2 | MHC class II protein complex (GO:0042613) | 0.006482 | 0.05149 | 184.98 | 932.06 |
| 3 | cation-transporting ATPase complex (GO:0090533) | 0.007973 | 0.05149 | 147.96 | 714.91 |
| 4 | MHC protein complex (GO:0042611) | 0.009957 | 0.05149 | 116.79 | 538.34 |
| 5 | myofibril (GO:0030016) | 0.01342 | 0.05149 | 85.32 | 367.79 |

Reduced ejection fraction ([HP:0012664](https://hpo.jax.org/app/browse/term/HP:0012664))

KEGG Pathway

| **Index** | **Name** | **P-value** | **Adjusted p-value** | **Odds Ratio** | **Combined score** |
| --- | --- | --- | --- | --- | --- |
| 1 | Pantothenate and CoA biosynthesis | 0.009412 | 0.06589 | 124.82 | 582.37 |
| 2 | African trypanosomiasis | 0.01653 | 0.07714 | 69.29 | 284.26 |
| 3 | Regulation of lipolysis in adipocytes | 0.02448 | 0.08482 | 46.15 | 171.21 |
| 4 | Thermogenesis | 0.004571 | 0.06400 | 24.55 | 132.26 |
| 5 | Renin secretion | 0.03063 | 0.08482 | 36.62 | 127.66 |

GO: Biological Process

| **Index** | **Name** | **P-value** | **Adjusted p-value** | **Odds Ratio** | **Combined score** |
| --- | --- | --- | --- | --- | --- |
| 1 | regulation of atrial cardiac muscle cell membrane repolarization (GO:0060372) | 0.000003775 | 0.0005097 | 1142.06 | 14260.84 |
| 2 | positive regulation of action potential (GO:0045760) | 0.002248 | 0.01755 | 624.59 | 3808.55 |
| 3 | bundle of His cell action potential (GO:0086043) | 0.002248 | 0.01755 | 624.59 | 3808.55 |
| 4 | response to denervation involved in regulation of muscle adaptation (GO:0014894) | 0.002248 | 0.01755 | 624.59 | 3808.55 |
| 5 | positive regulation of cardiac muscle contraction (GO:0060452) | 0.002248 | 0.01755 | 624.59 | 3808.55 |

GO: Molecular Function

| **Index** | **Name** | **P-value** | **Adjusted p-value** | **Odds Ratio** | **Combined score** |
| --- | --- | --- | --- | --- | --- |
| 1 | voltage-gated sodium channel activity involved in cardiac muscle cell action potential (GO:0086006) | 0.002248 | 0.02517 | 624.59 | 3808.55 |
| 2 | acid-amino acid ligase activity (GO:0016881) | 0.003146 | 0.02517 | 416.35 | 2398.85 |
| 3 | nitric-oxide synthase binding (GO:0050998) | 0.003146 | 0.02517 | 416.35 | 2398.85 |
| 4 | neuropeptide receptor binding (GO:0071855) | 0.008072 | 0.02629 | 146.87 | 707.80 |
| 5 | DNA polymerase binding (GO:0070182) | 0.008072 | 0.02629 | 146.87 | 707.80 |

GO: Cellular Component

| **Index** | **Name** | **P-value** | **Adjusted p-value** | **Odds Ratio** | **Combined score** |
| --- | --- | --- | --- | --- | --- |
| 1 | voltage-gated sodium channel complex (GO:0001518) | 0.007625 | 0.05451 | 156.05 | 760.96 |
| 2 | sodium channel complex (GO:0034706) | 0.01120 | 0.05451 | 103.99 | 467.16 |
| 3 | intercalated disc (GO:0014704) | 0.01387 | 0.05451 | 83.17 | 355.83 |
| 4 | cell-cell contact zone (GO:0044291) | 0.02096 | 0.05451 | 54.20 | 209.49 |
| 5 | sarcolemma (GO:0042383) | 0.02316 | 0.05451 | 48.87 | 184.02 |
| 6 | cytosolic large ribosomal subunit (GO:0022625) | 0.02448 | 0.05451 | 46.15 | 171.21 |

Tachycardia ([HP:0001649](https://hpo.jax.org/app/browse/term/HP:0001649))

KEGG Pathway

| **Index** | **Name** | **P-value** | **Adjusted p-value** | **Odds Ratio** | **Combined score** |
| --- | --- | --- | --- | --- | --- |
| 1 | Adrenergic signaling in cardiomyocytes | 6.163e-7 | 0.00001664 | 90.61 | 1295.71 |
| 2 | Oxytocin signaling pathway | 0.00005163 | 0.0006971 | 56.31 | 555.83 |
| 3 | Arrhythmogenic right ventricular cardiomyopathy | 0.0006453 | 0.003940 | 66.38 | 487.64 |
| 4 | Cardiac muscle contraction | 0.0008229 | 0.003940 | 58.54 | 415.82 |
| 5 | Hypertrophic cardiomyopathy | 0.0008802 | 0.003940 | 56.54 | 397.78 |

GO: Biological Process

| **Index** | **Name** | **P-value** | **Adjusted p-value** | **Odds Ratio** | **Combined score** |
| --- | --- | --- | --- | --- | --- |
| 1 | regulation of heart rate by cardiac conduction (GO:0086091) | 4.350e-21 | 8.135e-19 | 2575.35 | 120743.21 |
| 2 | SA node cell to atrial cardiac muscle cell signaling (GO:0086018) | 8.995e-10 | 1.682e-8 | 4283.14 | 89214.48 |
| 3 | membrane depolarization during cardiac muscle cell action potential (GO:0086012) | 5.833e-14 | 5.454e-12 | 1664.83 | 50731.79 |
| 4 | regulation of atrial cardiac muscle cell membrane depolarization (GO:0060371) | 3.147e-9 | 3.942e-8 | 2141.36 | 41921.23 |
| 5 | SA node cell action potential (GO:0086015) | 3.147e-9 | 3.942e-8 | 2141.36 | 41921.23 |

GO: Molecular Function

| **Index** | **Name** | **P-value** | **Adjusted p-value** | **Odds Ratio** | **Combined score** |
| --- | --- | --- | --- | --- | --- |
| 1 | voltage-gated sodium channel activity involved in cardiac muscle cell action potential (GO:0086006) | 8.995e-10 | 1.889e-8 | 4283.14 | 89214.48 |
| 2 | voltage-gated potassium channel activity involved in ventricular cardiac muscle cell action potential repolarization (GO:1902282) | 1.975e-8 | 2.765e-7 | 951.48 | 16879.14 |
| 3 | voltage-gated sodium channel activity (GO:0005248) | 2.294e-10 | 9.637e-9 | 739.70 | 16417.99 |
| 4 | voltage-gated potassium channel activity involved in cardiac muscle cell action potential repolarization (GO:0086008) | 4.082e-8 | 4.286e-7 | 713.50 | 12139.53 |
| 5 | sodium channel inhibitor activity (GO:0019871) | 0.000006290 | 0.00003774 | 832.67 | 9972.50 |

GO: Cellular Component

| **Index** | **Name** | **P-value** | **Adjusted p-value** | **Odds Ratio** | **Combined score** |
| --- | --- | --- | --- | --- | --- |
| 1 | voltage-gated sodium channel complex (GO:0001518) | 6.098e-8 | 0.000001890 | 611.51 | 10158.88 |
| 2 | sodium channel complex (GO:0034706) | 2.058e-7 | 0.000003190 | 388.99 | 5988.96 |
| 3 | intercalated disc (GO:0014704) | 0.0001038 | 0.0005364 | 172.08 | 1578.44 |
| 4 | junctional sarcoplasmic reticulum membrane (GO:0014701) | 0.004492 | 0.01392 | 277.53 | 1500.17 |
| 5 | voltage-gated potassium channel complex (GO:0008076) | 0.000005496 | 0.00004259 | 121.96 | 1477.11 |

Venous thrombosis ([HP:0004936](https://hpo.jax.org/app/browse/term/HP:0004936))

KEGG Pathway

| **Index** | **Name** | **P-value** | **Adjusted p-value** | **Odds Ratio** | **Combined score** |
| --- | --- | --- | --- | --- | --- |
| 1 | Adherens junction | 0.000005053 | 0.0005053 | 125.56 | 1531.25 |
| 2 | Gastric cancer | 0.00004680 | 0.002230 | 58.25 | 580.74 |
| 3 | Hepatocellular carcinoma | 0.00006690 | 0.002230 | 51.49 | 494.97 |
| 4 | Bacterial invasion of epithelial cells | 0.0006453 | 0.01076 | 66.38 | 487.64 |
| 5 | Colorectal cancer | 0.0008042 | 0.01149 | 59.24 | 422.16 |

GO: Biological Process

| **Index** | **Name** | **P-value** | **Adjusted p-value** | **Odds Ratio** | **Combined score** |
| --- | --- | --- | --- | --- | --- |
| 1 | entry of bacterium into host cell (GO:0035635) | 0.000006290 | 0.002201 | 832.67 | 9972.50 |
| 2 | entry into host (GO:0044409) | 0.00001750 | 0.002747 | 454.07 | 4973.59 |
| 3 | positive regulation of histone H3-K4 methylation (GO:0051571) | 0.00002354 | 0.002747 | 384.17 | 4094.01 |
| 4 | cellular response to indole-3-methanol (GO:0071681) | 0.002498 | 0.02399 | 555.17 | 3326.77 |
| 5 | regulation of guanyl-nucleotide exchange factor activity (GO:1905097) | 0.002498 | 0.02399 | 555.17 | 3326.77 |

GO: Molecular Function

| **Index** | **Name** | **P-value** | **Adjusted p-value** | **Odds Ratio** | **Combined score** |
| --- | --- | --- | --- | --- | --- |
| 1 | I-SMAD binding (GO:0070411) | 0.00001750 | 0.0008399 | 454.07 | 4973.59 |
| 2 | G protein-coupled serotonin receptor binding (GO:0031821) | 0.002997 | 0.01798 | 444.11 | 2580.41 |
| 3 | non-membrane spanning protein tyrosine phosphatase activity (GO:0004726) | 0.002997 | 0.01798 | 444.11 | 2580.41 |
| 4 | histone methyltransferase binding (GO:1990226) | 0.005488 | 0.02927 | 222.00 | 1155.57 |
| 5 | transcription coregulator binding (GO:0001221) | 0.0003059 | 0.007341 | 97.74 | 790.95 |

GO: Cellular Component

| **Index** | **Name** | **P-value** | **Adjusted p-value** | **Odds Ratio** | **Combined score** |
| --- | --- | --- | --- | --- | --- |
| 1 | beta-catenin-TCF complex (GO:1990907) | 0.006482 | 0.1037 | 184.98 | 932.06 |
| 2 | catenin complex (GO:0016342) | 0.01540 | 0.1037 | 73.93 | 308.54 |
| 3 | heterotrimeric G-protein complex (GO:0005834) | 0.01638 | 0.1037 | 69.30 | 284.93 |
| 4 | cytoplasmic side of plasma membrane (GO:0009898) | 0.02717 | 0.1132 | 41.02 | 147.91 |
| 5 | clathrin-coated vesicle (GO:0030136) | 0.03930 | 0.1132 | 28.00 | 90.64 |

Cardiovascular-symptom

Angina pectoris ([HP:0001681](https://hpo.jax.org/app/browse/term/HP:0001681))

KEGG Pathway

| **Index** | **Name** | **P-value** | **Adjusted p-value** | **Odds Ratio** | **Combined score** |
| --- | --- | --- | --- | --- | --- |
| 1 | Cholesterol metabolism | 7.176e-9 | 1.579e-7 | 289.04 | 5420.31 |
| 2 | ABC transporters | 0.000001263 | 0.00001389 | 203.55 | 2764.61 |
| 3 | Riboflavin metabolism | 0.003994 | 0.01464 | 317.19 | 1751.86 |
| 4 | Fat digestion and absorption | 0.0002010 | 0.001474 | 121.64 | 1035.45 |
| 5 | Primary bile acid biosynthesis | 0.008469 | 0.02662 | 138.71 | 661.82 |

GO: Biological Process

| **Index** | **Name** | **P-value** | **Adjusted p-value** | **Odds Ratio** | **Combined score** |
| --- | --- | --- | --- | --- | --- |
| 1 | regulation of intestinal cholesterol absorption (GO:0030300) | 0.000006290 | 0.0003522 | 832.67 | 9972.50 |
| 2 | intestinal cholesterol absorption (GO:0030299) | 0.000008085 | 0.0003622 | 713.68 | 8368.26 |
| 3 | negative regulation of cholesterol transport (GO:0032375) | 0.00001235 | 0.0003950 | 555.03 | 6273.06 |
| 4 | intestinal lipid absorption (GO:0098856) | 0.00001235 | 0.0003950 | 555.03 | 6273.06 |
| 5 | nucleoside triphosphate catabolic process (GO:0009143) | 0.002498 | 0.01797 | 555.17 | 3326.77 |

GO: Molecular Function

| **Index** | **Name** | **P-value** | **Adjusted p-value** | **Odds Ratio** | **Combined score** |
| --- | --- | --- | --- | --- | --- |
| 1 | lipoprotein lipase activity (GO:0004465) | 0.002498 | 0.01443 | 555.17 | 3326.77 |
| 2 | phosphodiesterase I activity (GO:0004528) | 0.002498 | 0.01443 | 555.17 | 3326.77 |
| 3 | cholesterol transfer activity (GO:0120020) | 0.00003428 | 0.0004405 | 312.09 | 3208.63 |
| 4 | sterol transfer activity (GO:0120015) | 0.00003830 | 0.0004405 | 293.72 | 2987.15 |
| 5 | non-membrane spanning protein tyrosine phosphatase activity (GO:0004726) | 0.002997 | 0.01443 | 444.11 | 2580.41 |

GO: Cellular Component

| **Index** | **Name** | **P-value** | **Adjusted p-value** | **Odds Ratio** | **Combined score** |
| --- | --- | --- | --- | --- | --- |
| 1 | ATP-binding cassette (ABC) transporter complex (GO:0043190) | 0.000003371 | 0.00008091 | 1249.13 | 15739.23 |
| 2 | cytoplasmic side of plasma membrane (GO:0009898) | 0.02717 | 0.1323 | 41.02 | 147.91 |
| 3 | clathrin-coated vesicle (GO:0030136) | 0.03930 | 0.1323 | 28.00 | 90.64 |
| 4 | clathrin-coated endocytic vesicle (GO:0045334) | 0.04171 | 0.1323 | 26.33 | 83.66 |
| 5 | lysosomal lumen (GO:0043202) | 0.04219 | 0.1323 | 26.02 | 82.37 |

Arrhythmia ([HP:0011675](https://hpo.jax.org/app/browse/term/HP:0011675))

KEGG Pathway

| **Index** | **Name** | **P-value** | **Adjusted p-value** | **Odds Ratio** | **Combined score** |
| --- | --- | --- | --- | --- | --- |
| 1 | Hypertrophic cardiomyopathy | 3.494e-15 | 6.683e-14 | 559.63 | 18629.01 |
| 2 | Dilated cardiomyopathy | 5.569e-15 | 6.683e-14 | 521.75 | 17124.60 |
| 3 | Cardiac muscle contraction | 1.177e-12 | 9.416e-12 | 368.69 | 10127.07 |
| 4 | Adrenergic signaling in cardiomyocytes | 3.297e-11 | 1.978e-10 | 206.73 | 4989.52 |
| 5 | Viral myocarditis | 0.0003921 | 0.001569 | 85.91 | 673.90 |

GO: Biological Process

| **Index** | **Name** | **P-value** | **Adjusted p-value** | **Odds Ratio** | **Combined score** |
| --- | --- | --- | --- | --- | --- |
| 1 | heart contraction (GO:0060047) | 6.755e-21 | 1.067e-18 | 2419.03 | 112349.58 |
| 2 | ventricular cardiac muscle tissue morphogenesis (GO:0055010) | 2.534e-18 | 1.334e-16 | 1725.20 | 69899.68 |
| 3 | cardiac muscle contraction (GO:0060048) | 2.534e-18 | 1.334e-16 | 1725.20 | 69899.68 |
| 4 | actin-myosin filament sliding (GO:0033275) | 5.941e-18 | 1.877e-16 | 1502.29 | 59587.92 |
| 5 | muscle filament sliding (GO:0030049) | 5.941e-18 | 1.877e-16 | 1502.29 | 59587.92 |

GO: Molecular Function

| **Index** | **Name** | **P-value** | **Adjusted p-value** | **Odds Ratio** | **Combined score** |
| --- | --- | --- | --- | --- | --- |
| 1 | myosin heavy chain binding (GO:0032036) | 3.147e-9 | 9.440e-8 | 2141.36 | 41921.23 |
| 2 | titin binding (GO:0031432) | 0.00001750 | 0.0001750 | 454.07 | 4973.59 |
| 3 | FATZ binding (GO:0051373) | 0.002498 | 0.01070 | 555.17 | 3326.77 |
| 4 | telethonin binding (GO:0031433) | 0.003495 | 0.01311 | 370.07 | 2093.27 |
| 5 | actin monomer binding (GO:0003785) | 0.00006709 | 0.0005032 | 217.03 | 2085.58 |

GO: Cellular Component

| **Index** | **Name** | **P-value** | **Adjusted p-value** | **Odds Ratio** | **Combined score** |
| --- | --- | --- | --- | --- | --- |
| 1 | cardiac myofibril (GO:0097512) | 8.995e-10 | 1.484e-8 | 4283.14 | 89214.48 |
| 2 | myofibril (GO:0030016) | 7.595e-13 | 2.506e-11 | 907.64 | 25328.60 |
| 3 | muscle myosin complex (GO:0005859) | 4.082e-8 | 4.490e-7 | 713.50 | 12139.53 |
| 4 | myosin filament (GO:0032982) | 0.002997 | 0.01978 | 444.11 | 2580.41 |
| 5 | pseudopodium (GO:0031143) | 0.004492 | 0.02118 | 277.53 | 1500.17 |

Palpitations ([HP:0001962](https://hpo.jax.org/app/browse/term/HP:0001962))

KEGG Pathway

| **Index** | **Name** | **P-value** | **Adjusted p-value** | **Odds Ratio** | **Combined score** |
| --- | --- | --- | --- | --- | --- |
| 1 | Adrenergic signaling in cardiomyocytes | 6.163e-7 | 0.00001849 | 90.61 | 1295.71 |
| 2 | Cardiac muscle contraction | 0.000009332 | 0.00009412 | 101.56 | 1176.29 |
| 3 | Hypertrophic cardiomyopathy | 0.00001034 | 0.00009412 | 98.04 | 1125.55 |
| 4 | Dilated cardiomyopathy | 0.00001255 | 0.00009412 | 91.69 | 1034.81 |
| 5 | cGMP-PKG signaling pathway | 0.00006572 | 0.0003943 | 51.81 | 498.94 |

GO: Biological Process

| **Index** | **Name** | **P-value** | **Adjusted p-value** | **Odds Ratio** | **Combined score** |
| --- | --- | --- | --- | --- | --- |
| 1 | positive regulation of cardioblast differentiation (GO:0051891) | 0.000002248 | 0.00002428 | 1665.58 | 21661.60 |
| 2 | regulation of cardioblast differentiation (GO:0051890) | 0.000002248 | 0.00002428 | 1665.58 | 21661.60 |
| 3 | cardiac atrium morphogenesis (GO:0003209) | 1.217e-10 | 9.855e-9 | 887.78 | 20267.77 |
| 4 | cardiac muscle tissue morphogenesis (GO:0055008) | 1.893e-12 | 6.132e-10 | 739.37 | 19957.87 |
| 5 | atrial septum morphogenesis (GO:0060413) | 1.482e-8 | 4.001e-7 | 1070.46 | 19297.66 |

GO: Molecular Function

| **Index** | **Name** | **P-value** | **Adjusted p-value** | **Odds Ratio** | **Combined score** |
| --- | --- | --- | --- | --- | --- |
| 1 | myosin heavy chain binding (GO:0032036) | 0.000004719 | 0.0002218 | 999.25 | 12254.80 |
| 2 | voltage-gated sodium channel activity involved in cardiac muscle cell action potential (GO:0086006) | 0.002498 | 0.01612 | 555.17 | 3326.77 |
| 3 | nitric-oxide synthase binding (GO:0050998) | 0.003495 | 0.01612 | 370.07 | 2093.27 |
| 4 | actin monomer binding (GO:0003785) | 0.00006709 | 0.001577 | 217.03 | 2085.58 |
| 5 | potassium channel inhibitor activity (GO:0019870) | 0.004492 | 0.01612 | 277.53 | 1500.17 |

GO: Cellular Component

| **Index** | **Name** | **P-value** | **Adjusted p-value** | **Odds Ratio** | **Combined score** |
| --- | --- | --- | --- | --- | --- |
| 1 | muscle myosin complex (GO:0005859) | 0.00002354 | 0.0006357 | 384.17 | 4094.01 |
| 2 | cardiac myofibril (GO:0097512) | 0.002498 | 0.01124 | 555.17 | 3326.77 |
| 3 | myofibril (GO:0030016) | 0.00007845 | 0.001059 | 199.65 | 1887.30 |
| 4 | sarcolemma (GO:0042383) | 0.0002944 | 0.002647 | 99.70 | 810.62 |
| 5 | caveola (GO:0005901) | 0.0003921 | 0.002647 | 85.91 | 673.90 |

Stroke ([HP:0001297](https://hpo.jax.org/app/browse/term/HP:0001297))

KEGG Pathway

| **Index** | **Name** | **P-value** | **Adjusted p-value** | **Odds Ratio** | **Combined score** |
| --- | --- | --- | --- | --- | --- |
| 1 | Hypertrophic cardiomyopathy | 0.00001034 | 0.0001569 | 98.04 | 1125.55 |
| 2 | Dilated cardiomyopathy | 0.00001255 | 0.0001569 | 91.69 | 1034.81 |
| 3 | Adrenergic signaling in cardiomyocytes | 0.00004774 | 0.0003978 | 57.85 | 575.61 |
| 4 | cGMP-PKG signaling pathway | 0.00006572 | 0.0004108 | 51.81 | 498.94 |
| 5 | Cardiac muscle contraction | 0.0008229 | 0.004114 | 58.54 | 415.82 |

GO: Biological Process

| **Index** | **Name** | **P-value** | **Adjusted p-value** | **Odds Ratio** | **Combined score** |
| --- | --- | --- | --- | --- | --- |
| 1 | positive regulation of cardioblast differentiation (GO:0051891) | 0.000002248 | 0.00002290 | 1665.58 | 21661.60 |
| 2 | regulation of cardioblast differentiation (GO:0051890) | 0.000002248 | 0.00002290 | 1665.58 | 21661.60 |
| 3 | cardiac atrium morphogenesis (GO:0003209) | 1.217e-10 | 6.691e-9 | 887.78 | 20267.77 |
| 4 | cardiac muscle tissue morphogenesis (GO:0055008) | 1.893e-12 | 5.205e-10 | 739.37 | 19957.87 |
| 5 | atrial septum morphogenesis (GO:0060413) | 1.482e-8 | 2.911e-7 | 1070.46 | 19297.66 |

GO: Molecular Function

| **Index** | **Name** | **P-value** | **Adjusted p-value** | **Odds Ratio** | **Combined score** |
| --- | --- | --- | --- | --- | --- |
| 1 | voltage-gated sodium channel activity involved in cardiac muscle cell action potential (GO:0086006) | 0.002498 | 0.01463 | 555.17 | 3326.77 |
| 2 | myosin heavy chain binding (GO:0032036) | 0.003495 | 0.01463 | 370.07 | 2093.27 |
| 3 | nitric-oxide synthase binding (GO:0050998) | 0.003495 | 0.01463 | 370.07 | 2093.27 |
| 4 | co-SMAD binding (GO:0070410) | 0.004990 | 0.01463 | 246.68 | 1307.49 |
| 5 | calcium channel inhibitor activity (GO:0019855) | 0.005488 | 0.01463 | 222.00 | 1155.57 |

GO: Cellular Component

| **Index** | **Name** | **P-value** | **Adjusted p-value** | **Odds Ratio** | **Combined score** |
| --- | --- | --- | --- | --- | --- |
| 1 | cardiac myofibril (GO:0097512) | 0.000002248 | 0.00002361 | 1665.58 | 21661.60 |
| 2 | myofibril (GO:0030016) | 2.616e-7 | 0.000005494 | 356.54 | 5403.81 |
| 3 | muscle myosin complex (GO:0005859) | 0.007476 | 0.04446 | 158.54 | 776.21 |
| 4 | voltage-gated sodium channel complex (GO:0001518) | 0.008469 | 0.04446 | 138.71 | 661.82 |
| 5 | sodium channel complex (GO:0034706) | 0.01243 | 0.04619 | 92.44 | 405.55 |

Syncope ([HP:0001279](https://hpo.jax.org/app/browse/term/HP:0001279))

KEGG Pathway

| **Index** | **Name** | **P-value** | **Adjusted p-value** | **Odds Ratio** | **Combined score** |
| --- | --- | --- | --- | --- | --- |
| 1 | Adrenergic signaling in cardiomyocytes | 6.163e-7 | 0.00001726 | 90.61 | 1295.71 |
| 2 | Oxytocin signaling pathway | 0.00005163 | 0.0007229 | 56.31 | 555.83 |
| 3 | Arrhythmogenic right ventricular cardiomyopathy | 0.0006453 | 0.004086 | 66.38 | 487.64 |
| 4 | Cardiac muscle contraction | 0.0008229 | 0.004086 | 58.54 | 415.82 |
| 5 | Hypertrophic cardiomyopathy | 0.0008802 | 0.004086 | 56.54 | 397.78 |

GO: Biological Process

| **Index** | **Name** | **P-value** | **Adjusted p-value** | **Odds Ratio** | **Combined score** |
| --- | --- | --- | --- | --- | --- |
| 1 | regulation of heart rate by cardiac conduction (GO:0086091) | 1.500e-24 | 3.121e-22 | 5988.00 | 328479.32 |
| 2 | membrane depolarization during SA node cell action potential (GO:0086046) | 8.995e-10 | 1.701e-8 | 4283.14 | 89214.48 |
| 3 | SA node cell to atrial cardiac muscle cell signaling (GO:0086018) | 8.995e-10 | 1.701e-8 | 4283.14 | 89214.48 |
| 4 | membrane depolarization during cardiac muscle cell action potential (GO:0086012) | 5.833e-14 | 2.799e-12 | 1664.83 | 50731.79 |
| 5 | regulation of ventricular cardiac muscle cell membrane repolarization (GO:0060307) | 8.075e-14 | 2.799e-12 | 1536.69 | 46327.23 |

GO: Molecular Function

| **Index** | **Name** | **P-value** | **Adjusted p-value** | **Odds Ratio** | **Combined score** |
| --- | --- | --- | --- | --- | --- |
| 1 | voltage-gated potassium channel activity involved in ventricular cardiac muscle cell action potential repolarization (GO:1902282) | 1.556e-11 | 6.069e-10 | 1665.17 | 41439.58 |
| 2 | voltage-gated potassium channel activity involved in cardiac muscle cell action potential repolarization (GO:0086008) | 4.289e-11 | 8.363e-10 | 1210.85 | 28905.97 |
| 3 | voltage-gated sodium channel activity involved in cardiac muscle cell action potential (GO:0086006) | 0.000002248 | 0.00001461 | 1665.58 | 21661.60 |
| 4 | voltage-gated sodium channel activity (GO:0005248) | 1.379e-7 | 0.000001345 | 450.47 | 7115.97 |
| 5 | voltage-gated potassium channel activity (GO:0005249) | 2.538e-10 | 3.299e-9 | 258.61 | 5713.86 |

GO: Cellular Component

| **Index** | **Name** | **P-value** | **Adjusted p-value** | **Odds Ratio** | **Combined score** |
| --- | --- | --- | --- | --- | --- |
| 1 | intercalated disc (GO:0014704) | 4.016e-7 | 0.000006626 | 305.54 | 4499.94 |
| 2 | voltage-gated sodium channel complex (GO:0001518) | 0.00003048 | 0.0001437 | 332.92 | 3461.84 |
| 3 | voltage-gated potassium channel complex (GO:0008076) | 3.373e-8 | 0.000001113 | 192.47 | 3311.50 |
| 4 | cell-cell contact zone (GO:0044291) | 0.000001443 | 0.00001587 | 194.28 | 2612.86 |
| 5 | sarcolemma (GO:0042383) | 0.000001964 | 0.00001620 | 174.41 | 2291.88 |
