## Supplementary Table 1-3 for "Long COVID: G Protein-Coupled Receptors (GPCRs) responsible for persistent post-COVID symptoms": Supplementary File 2.8-Gastrointestinal.docx

gi-finding

Abnormal pancreas morphology ([HP:0012090](https://hpo.jax.org/app/browse/term/HP:0012090))

GO: Biological Process

| **Index** | **Name** | **P-value** | **Adjusted p-value** | **Odds Ratio** | **Combined score** |
| --- | --- | --- | --- | --- | --- |
| 1 | cilium assembly (GO:0060271) | 7.884e-19 | 4.178e-17 | 196860.00 | 8205971.98 |
| 2 | ciliary basal body-plasma membrane docking (GO:0097711) | 8.554e-18 | 2.267e-16 | 915.08 | 35962.78 |
| 3 | cilium organization (GO:0044782) | 2.747e-17 | 4.853e-16 | 812.51 | 30983.64 |
| 4 | plasma membrane bounded cell projection assembly (GO:0120031) | 1.681e-16 | 2.227e-15 | 659.81 | 23965.68 |
| 5 | smoothened signaling pathway (GO:0007224) | 3.049e-12 | 2.693e-11 | 665.33 | 17642.09 |

GO: Molecular Function

| **Index** | **Name** | **P-value** | **Adjusted p-value** | **Odds Ratio** | **Combined score** |
| --- | --- | --- | --- | --- | --- |
| 1 | gamma-tubulin binding (GO:0043015) | 0.01144 | 0.02289 | 100.85 | 450.83 |
| 2 | tubulin binding (GO:0015631) | 0.1433 | 0.1433 | 7.15 | 13.88 |

GO: Cellular Component

| **Index** | **Name** | **P-value** | **Adjusted p-value** | **Odds Ratio** | **Combined score** |
| --- | --- | --- | --- | --- | --- |
| 1 | ciliary membrane (GO:0060170) | 4.016e-7 | 0.000001406 | 305.54 | 4499.94 |
| 2 | cilium (GO:0005929) | 5.245e-10 | 3.672e-9 | 128.31 | 2741.69 |
| 3 | cell projection membrane (GO:0031253) | 0.00001104 | 0.00002576 | 95.83 | 1093.80 |
| 4 | bounding membrane of organelle (GO:0098588) | 0.005508 | 0.009639 | 10.78 | 56.10 |
| 5 | cell-cell junction (GO:0005911) | 0.1276 | 0.1786 | 8.12 | 16.71 |

Gastric ulcer ([HP:0002592](https://hpo.jax.org/app/browse/term/HP:0002592))

KEGG Pathway

| **Index** | **Name** | **P-value** | **Adjusted p-value** | **Odds Ratio** | **Combined score** |
| --- | --- | --- | --- | --- | --- |
| 1 | Hepatocellular carcinoma | 1.664e-18 | 4.992e-18 | 1122.51 | 45952.51 |
| 2 | Thermogenesis | 3.221e-17 | 4.832e-17 | 797.77 | 30294.76 |
| 3 | Transcriptional misregulation in cancer | 0.09198 | 0.09198 | 11.52 | 27.48 |

GO: Biological Process

| **Index** | **Name** | **P-value** | **Adjusted p-value** | **Odds Ratio** | **Combined score** |
| --- | --- | --- | --- | --- | --- |
| 1 | regulation of transcription by RNA polymerase II (GO:0006357) | 2.617e-10 | 2.061e-9 | 177940.00 | 3926054.56 |
| 2 | ATP-dependent chromatin remodeling (GO:0043044) | 8.343e-21 | 5.256e-19 | 2347.76 | 108543.80 |
| 3 | nucleosome disassembly (GO:0006337) | 4.377e-17 | 6.894e-16 | 2497.25 | 94065.34 |
| 4 | chromatin disassembly (GO:0031498) | 4.377e-17 | 6.894e-16 | 2497.25 | 94065.34 |
| 5 | chromatin remodeling (GO:0006338) | 1.773e-20 | 5.584e-19 | 1904.94 | 86635.00 |

GO: Molecular Function

| **Index** | **Name** | **P-value** | **Adjusted p-value** | **Odds Ratio** | **Combined score** |
| --- | --- | --- | --- | --- | --- |
| 1 | nucleosomal DNA binding (GO:0031492) | 2.605e-15 | 5.210e-14 | 1109.06 | 37243.54 |
| 2 | RNA polymerase I core promoter sequence-specific DNA binding (GO:0001164) | 0.000006290 | 0.00004193 | 832.67 | 9972.50 |
| 3 | RNA polymerase I transcription regulatory region sequence-specific DNA binding (GO:0001163) | 0.000006290 | 0.00004193 | 832.67 | 9972.50 |
| 4 | Tat protein binding (GO:0030957) | 0.00001010 | 0.00005052 | 624.44 | 7182.69 |
| 5 | core promoter sequence-specific DNA binding (GO:0001046) | 0.0001485 | 0.0004949 | 142.54 | 1256.48 |

GO: Cellular Component

| **Index** | **Name** | **P-value** | **Adjusted p-value** | **Odds Ratio** | **Combined score** |
| --- | --- | --- | --- | --- | --- |
| 1 | SWI/SNF complex (GO:0016514) | 3.273e-32 | 5.236e-31 | 199810.00 | 14485640.47 |
| 2 | npBAF complex (GO:0071564) | 1.169e-26 | 9.350e-26 | 26649.33 | 1591265.17 |
| 3 | nBAF complex (GO:0071565) | 2.127e-25 | 1.134e-24 | 13322.67 | 756859.90 |
| 4 | Ino80 complex (GO:0031011) | 0.006979 | 0.01396 | 170.74 | 847.70 |
| 5 | H4/H2A histone acetyltransferase complex (GO:0043189) | 0.009462 | 0.01593 | 123.28 | 574.57 |

Gastroesophageal reflux ([HP:0002020](https://hpo.jax.org/app/browse/term/HP:0002020))

KEGG Pathway

| **Index** | **Name** | **P-value** | **Adjusted p-value** | **Odds Ratio** | **Combined score** |
| --- | --- | --- | --- | --- | --- |
| 1 | Nicotine addiction | 6.174e-12 | 7.100e-10 | 570.14 | 14715.76 |
| 2 | Long-term potentiation | 9.011e-11 | 5.182e-9 | 321.42 | 7434.41 |
| 3 | Amphetamine addiction | 2.681e-8 | 9.684e-7 | 204.36 | 3562.87 |
| 4 | ErbB signaling pathway | 6.256e-8 | 0.000001199 | 163.86 | 2717.97 |
| 5 | Circadian entrainment | 1.067e-7 | 0.000001754 | 142.63 | 2289.62 |

GO: Biological Process

| **Index** | **Name** | **P-value** | **Adjusted p-value** | **Odds Ratio** | **Combined score** |
| --- | --- | --- | --- | --- | --- |
| 1 | cellular response to histamine (GO:0071420) | 0.000004719 | 0.0001184 | 999.25 | 12254.80 |
| 2 | excitatory chemical synaptic transmission (GO:0098976) | 0.000004719 | 0.0001184 | 999.25 | 12254.80 |
| 3 | regulation of NMDA receptor activity (GO:2000310) | 1.125e-9 | 1.412e-7 | 475.29 | 9793.38 |
| 4 | positive regulation of synapse maturation (GO:0090129) | 0.000008085 | 0.0001691 | 713.68 | 8368.26 |
| 5 | response to histamine (GO:0034776) | 0.00001010 | 0.0001937 | 624.44 | 7182.69 |

GO: Molecular Function

| **Index** | **Name** | **P-value** | **Adjusted p-value** | **Odds Ratio** | **Combined score** |
| --- | --- | --- | --- | --- | --- |
| 1 | glutamate-gated calcium ion channel activity (GO:0022849) | 0.000002248 | 0.0001304 | 1665.58 | 21661.60 |
| 2 | NMDA glutamate receptor activity (GO:0004972) | 0.000006290 | 0.0001401 | 832.67 | 9972.50 |
| 3 | glycine binding (GO:0016594) | 0.00001235 | 0.0001502 | 555.03 | 6273.06 |
| 4 | GABA-gated chloride ion channel activity (GO:0022851) | 0.00001750 | 0.0001692 | 454.07 | 4973.59 |
| 5 | ionotropic glutamate receptor activity (GO:0004970) | 0.00003048 | 0.0002221 | 332.92 | 3461.84 |

GO: Cellular Component

| **Index** | **Name** | **P-value** | **Adjusted p-value** | **Odds Ratio** | **Combined score** |
| --- | --- | --- | --- | --- | --- |
| 1 | NMDA selective glutamate receptor complex (GO:0017146) | 0.000006290 | 0.0001069 | 832.67 | 9972.50 |
| 2 | GABA-A receptor complex (GO:1902711) | 0.00003830 | 0.0003256 | 293.72 | 2987.15 |
| 3 | synaptic membrane (GO:0097060) | 0.00007266 | 0.0004941 | 207.98 | 1981.99 |
| 4 | ionotropic glutamate receptor complex (GO:0008328) | 0.0001107 | 0.0006274 | 166.33 | 1515.06 |
| 5 | neuron projection (GO:0043005) | 8.582e-8 | 0.000002918 | 53.02 | 862.66 |

Gastroparesis ([HP:0002578](https://hpo.jax.org/app/browse/term/HP:0002578))

KEGG Pathway

| **Index** | **Name** | **P-value** | **Adjusted p-value** | **Odds Ratio** | **Combined score** |
| --- | --- | --- | --- | --- | --- |
| 1 | Various types of N-glycan biosynthesis | 0.01933 | 0.04945 | 58.34 | 230.20 |
| 2 | N-Glycan biosynthesis | 0.02473 | 0.04945 | 45.22 | 167.30 |
| 3 | Vascular smooth muscle contraction | 0.06456 | 0.08608 | 16.72 | 45.80 |
| 4 | Endocytosis | 0.1191 | 0.1191 | 8.74 | 18.59 |

GO: Biological Process

| **Index** | **Name** | **P-value** | **Adjusted p-value** | **Odds Ratio** | **Combined score** |
| --- | --- | --- | --- | --- | --- |
| 1 | mitochondrial proton-transporting ATP synthase complex assembly (GO:0033615) | 0.002997 | 0.03069 | 444.11 | 2580.41 |
| 2 | proton-transporting ATP synthase complex assembly (GO:0043461) | 0.003495 | 0.03069 | 370.07 | 2093.27 |
| 3 | mitochondrial DNA replication (GO:0006264) | 0.003994 | 0.03069 | 317.19 | 1751.86 |
| 4 | regulation of DNA-directed DNA polymerase activity (GO:1900262) | 0.004492 | 0.03069 | 277.53 | 1500.17 |
| 5 | positive regulation of DNA-directed DNA polymerase activity (GO:1900264) | 0.004492 | 0.03069 | 277.53 | 1500.17 |

GO: Molecular Function

| **Index** | **Name** | **P-value** | **Adjusted p-value** | **Odds Ratio** | **Combined score** |
| --- | --- | --- | --- | --- | --- |
| 1 | aromatic amino acid transmembrane transporter activity (GO:0015173) | 0.004492 | 0.05238 | 277.53 | 1500.17 |
| 2 | DNA polymerase binding (GO:0070182) | 0.008966 | 0.05238 | 130.54 | 615.43 |
| 3 | acetylglucosaminyltransferase activity (GO:0008375) | 0.02375 | 0.05238 | 47.15 | 176.34 |
| 4 | cation transmembrane transporter activity (GO:0008324) | 0.02375 | 0.05238 | 47.15 | 176.34 |
| 5 | manganese ion binding (GO:0030145) | 0.02375 | 0.05238 | 47.15 | 176.34 |

GO: Cellukar Component

| **Index** | **Name** | **P-value** | **Adjusted p-value** | **Odds Ratio** | **Combined score** |
| --- | --- | --- | --- | --- | --- |
| 1 | myosin filament (GO:0032982) | 0.002997 | 0.02987 | 444.11 | 2580.41 |
| 2 | U4 snRNP (GO:0005687) | 0.004492 | 0.02987 | 277.53 | 1500.17 |
| 3 | melanosome membrane (GO:0033162) | 0.005985 | 0.02987 | 201.81 | 1032.95 |
| 4 | pigment granule membrane (GO:0090741) | 0.005985 | 0.02987 | 201.81 | 1032.95 |
| 5 | intrinsic component of mitochondrial membrane (GO:0098573) | 0.005985 | 0.02987 | 201.81 | 1032.95 |

Hepatic steatosis ([HP:0001397](https://hpo.jax.org/app/browse/term/HP:0001397))

KEGG Pathway

| **Index** | **Name** | **P-value** | **Adjusted p-value** | **Odds Ratio** | **Combined score** |
| --- | --- | --- | --- | --- | --- |
| 1 | Fatty acid degradation | 3.852e-9 | 6.933e-8 | 341.04 | 6607.62 |
| 2 | Butanoate metabolism | 0.00008446 | 0.0004372 | 191.96 | 1800.45 |
| 3 | beta-Alanine metabolism | 0.00009715 | 0.0004372 | 178.23 | 1646.74 |
| 4 | PPAR signaling pathway | 0.000005727 | 0.00005154 | 120.24 | 1451.28 |
| 5 | Propanoate metabolism | 0.0001252 | 0.0004506 | 155.92 | 1401.11 |

GO: Biological Process

| **Index** | **Name** | **P-value** | **Adjusted p-value** | **Odds Ratio** | **Combined score** |
| --- | --- | --- | --- | --- | --- |
| 1 | carnitine transport (GO:0015879) | 0.000003371 | 0.00004551 | 1249.13 | 15739.23 |
| 2 | fatty acid beta-oxidation (GO:0006635) | 2.433e-11 | 1.314e-9 | 424.32 | 10370.14 |
| 3 | fatty acid beta-oxidation using acyl-CoA dehydrogenase (GO:0033539) | 0.00001010 | 0.00009093 | 624.44 | 7182.69 |
| 4 | carnitine shuttle (GO:0006853) | 0.00001235 | 0.00009523 | 555.03 | 6273.06 |
| 5 | fatty acid oxidation (GO:0019395) | 1.415e-8 | 3.821e-7 | 241.64 | 4367.22 |

GO: Molecular Function

| **Index** | **Name** | **P-value** | **Adjusted p-value** | **Odds Ratio** | **Combined score** |
| --- | --- | --- | --- | --- | --- |
| 1 | acyl-CoA dehydrogenase activity (GO:0003995) | 0.000008085 | 0.00009259 | 713.68 | 8368.26 |
| 2 | quaternary ammonium group transmembrane transporter activity (GO:0015651) | 0.00001235 | 0.00009259 | 555.03 | 6273.06 |
| 3 | acyl carnitine transmembrane transporter activity (GO:0015227) | 0.002498 | 0.005825 | 555.17 | 3326.77 |
| 4 | carnitine O-acyltransferase activity (GO:0016406) | 0.002997 | 0.005825 | 444.11 | 2580.41 |
| 5 | 3-hydroxyacyl-CoA dehydrogenase activity (GO:0003857) | 0.003495 | 0.005825 | 370.07 | 2093.27 |

GO: Cellular Component

| **Index** | **Name** | **P-value** | **Adjusted p-value** | **Odds Ratio** | **Combined score** |
| --- | --- | --- | --- | --- | --- |
| 1 | mitochondrial membrane (GO:0031966) | 0.00005604 | 0.001233 | 27.99 | 274.03 |
| 2 | mitochondrial envelope (GO:0005740) | 0.001741 | 0.007661 | 39.73 | 252.41 |
| 3 | brush border membrane (GO:0031526) | 0.01835 | 0.05043 | 61.59 | 246.23 |
| 4 | intrinsic component of mitochondrial inner membrane (GO:0031304) | 0.01933 | 0.05043 | 58.34 | 230.20 |
| 5 | mitochondrial inner membrane (GO:0005743) | 0.0004815 | 0.003148 | 25.93 | 198.08 |

Hepatitis ([HP:0012115](https://hpo.jax.org/app/browse/term/HP:0012115))

KEGG Pathway

| **Index** | **Name** | **P-value** | **Adjusted p-value** | **Odds Ratio** | **Combined score** |
| --- | --- | --- | --- | --- | --- |
| 1 | Primary immunodeficiency | 4.712e-12 | 5.842e-10 | 604.76 | 15772.67 |
| 2 | B cell receptor signaling pathway | 2.384e-10 | 9.853e-9 | 262.03 | 5805.76 |
| 3 | Chagas disease | 7.715e-10 | 2.392e-8 | 205.08 | 4303.17 |
| 4 | Allograft rejection | 7.523e-7 | 0.00001037 | 244.35 | 3445.31 |
| 5 | Epstein-Barr virus infection | 2.001e-10 | 9.853e-9 | 151.48 | 3383.01 |

GO: Biological Process

| **Index** | **Name** | **P-value** | **Adjusted p-value** | **Odds Ratio** | **Combined score** |
| --- | --- | --- | --- | --- | --- |
| 1 | positive thymic T cell selection (GO:0045059) | 0.000002248 | 0.0001192 | 1665.58 | 21661.60 |
| 2 | T cell apoptotic process (GO:0070231) | 0.000004719 | 0.0001786 | 999.25 | 12254.80 |
| 3 | B cell receptor signaling pathway (GO:0050853) | 1.451e-9 | 1.922e-7 | 443.56 | 9026.96 |
| 4 | positive regulation of B cell differentiation (GO:0045579) | 0.000008085 | 0.0002678 | 713.68 | 8368.26 |
| 5 | antigen receptor-mediated signaling pathway (GO:0050851) | 6.059e-13 | 1.606e-10 | 259.71 | 7306.12 |

GO: Molecular Function

| **Index** | **Name** | **P-value** | **Adjusted p-value** | **Odds Ratio** | **Combined score** |
| --- | --- | --- | --- | --- | --- |
| 1 | phospholipase binding (GO:0043274) | 0.00002041 | 0.0005919 | 416.21 | 4494.85 |
| 2 | 1-phosphatidylinositol-4-phosphate 3-kinase activity (GO:0035005) | 0.003495 | 0.01709 | 370.07 | 2093.27 |
| 3 | tumor necrosis factor-activated receptor activity (GO:0005031) | 0.004492 | 0.01709 | 277.53 | 1500.17 |
| 4 | non-membrane spanning protein tyrosine kinase activity (GO:0004715) | 0.0001327 | 0.001924 | 151.19 | 1349.73 |
| 5 | 1-phosphatidylinositol-3-kinase activity (GO:0016303) | 0.004990 | 0.01709 | 246.68 | 1307.49 |

GO: Cellular Component

| **Index** | **Name** | **P-value** | **Adjusted p-value** | **Odds Ratio** | **Combined score** |
| --- | --- | --- | --- | --- | --- |
| 1 | T cell receptor complex (GO:0042101) | 1.975e-8 | 3.753e-7 | 951.48 | 16879.14 |
| 2 | alpha-beta T cell receptor complex (GO:0042105) | 0.000003371 | 0.00003203 | 1249.13 | 15739.23 |
| 3 | phosphatidylinositol 3-kinase complex, class I (GO:0097651) | 0.002498 | 0.009491 | 555.17 | 3326.77 |
| 4 | CD95 death-inducing signaling complex (GO:0031265) | 0.002498 | 0.009491 | 555.17 | 3326.77 |
| 5 | death-inducing signaling complex (GO:0031264) | 0.003994 | 0.01111 | 317.19 | 1751.86 |

Hepatomegaly ([HP:0002240](https://hpo.jax.org/app/browse/term/HP:0002240))

KEGG Pathway

| **Index** | **Name** | **P-value** | **Adjusted p-value** | **Odds Ratio** | **Combined score** |
| --- | --- | --- | --- | --- | --- |
| 1 | Primary immunodeficiency | 5.941e-18 | 2.198e-16 | 1502.29 | 59587.92 |
| 2 | T cell receptor signaling pathway | 8.515e-10 | 1.575e-8 | 200.92 | 4196.00 |
| 3 | Th1 and Th2 cell differentiation | 8.619e-8 | 0.000001063 | 150.77 | 2452.58 |
| 4 | Autoimmune thyroid disease | 0.000002081 | 0.00001283 | 170.91 | 2236.02 |
| 5 | Hematopoietic cell lineage | 1.159e-7 | 0.000001072 | 139.61 | 2229.69 |

GO: Biological Process

| **Index** | **Name** | **P-value** | **Adjusted p-value** | **Odds Ratio** | **Combined score** |
| --- | --- | --- | --- | --- | --- |
| 1 | positive thymic T cell selection (GO:0045059) | 8.995e-10 | 1.151e-7 | 4283.14 | 89214.48 |
| 2 | V(D)J recombination (GO:0033151) | 0.00001235 | 0.0001580 | 555.03 | 6273.06 |
| 3 | T cell differentiation in thymus (GO:0033077) | 0.00002041 | 0.0002375 | 416.21 | 4494.85 |
| 4 | B cell activation involved in immune response (GO:0002312) | 0.00002690 | 0.0002869 | 356.71 | 3753.85 |
| 5 | regulation of T cell differentiation (GO:0045580) | 8.148e-7 | 0.00002027 | 237.55 | 3330.49 |

GO: Molecular Function

| **Index** | **Name** | **P-value** | **Adjusted p-value** | **Odds Ratio** | **Combined score** |
| --- | --- | --- | --- | --- | --- |
| 1 | endodeoxyribonuclease activity (GO:0004520) | 0.009957 | 0.06558 | 116.79 | 538.34 |
| 2 | phosphatidylinositol-3,5-bisphosphate binding (GO:0080025) | 0.01194 | 0.06558 | 96.46 | 427.12 |
| 3 | wide pore channel activity (GO:0022829) | 0.01243 | 0.06558 | 92.44 | 405.55 |
| 4 | phosphatidylinositol-3,4-bisphosphate binding (GO:0043325) | 0.01293 | 0.06558 | 88.73 | 385.85 |
| 5 | tumor necrosis factor receptor superfamily binding (GO:0032813) | 0.01392 | 0.06558 | 82.15 | 351.18 |

GO: Cellular Component

| **Index** | **Name** | **P-value** | **Adjusted p-value** | **Odds Ratio** | **Combined score** |
| --- | --- | --- | --- | --- | --- |
| 1 | T cell receptor complex (GO:0042101) | 1.975e-8 | 2.765e-7 | 951.48 | 16879.14 |
| 2 | alpha-beta T cell receptor complex (GO:0042105) | 0.000003371 | 0.00002360 | 1249.13 | 15739.23 |
| 3 | cytolytic granule (GO:0044194) | 0.003495 | 0.009787 | 370.07 | 2093.27 |
| 4 | clathrin-coated endocytic vesicle (GO:0045334) | 0.0007857 | 0.002750 | 59.96 | 428.66 |
| 5 | clathrin-coated endocytic vesicle membrane (GO:0030669) | 0.03398 | 0.07719 | 32.55 | 110.09 |

Malnutrition ([HP:0004395](https://hpo.jax.org/app/browse/term/HP:0004395))

KEGG Pathway

| **Index** | **Name** | **P-value** | **Adjusted p-value** | **Odds Ratio** | **Combined score** |
| --- | --- | --- | --- | --- | --- |
| 1 | Asthma | 0.01540 | 0.07448 | 73.93 | 308.54 |
| 2 | Allograft rejection | 0.01884 | 0.07448 | 59.92 | 237.98 |
| 3 | Graft-versus-host disease | 0.02081 | 0.07448 | 54.06 | 209.35 |
| 4 | Type I diabetes mellitus | 0.02130 | 0.07448 | 52.77 | 203.13 |
| 5 | Carbohydrate digestion and absorption | 0.02326 | 0.07448 | 48.17 | 181.19 |

GO: Biological Process

| **Index** | **Name** | **P-value** | **Adjusted p-value** | **Odds Ratio** | **Combined score** |
| --- | --- | --- | --- | --- | --- |
| 1 | bone marrow development (GO:0048539) | 0.002498 | 0.03376 | 555.17 | 3326.77 |
| 2 | mature ribosome assembly (GO:0042256) | 0.002498 | 0.03376 | 555.17 | 3326.77 |
| 3 | glucose import across plasma membrane (GO:0098708) | 0.002498 | 0.03376 | 555.17 | 3326.77 |
| 4 | positive regulation of intrinsic apoptotic signaling pathway by p53 class mediator (GO:1902255) | 0.002498 | 0.03376 | 555.17 | 3326.77 |
| 5 | hexose import across plasma membrane (GO:0140271) | 0.002997 | 0.03376 | 444.11 | 2580.41 |

GO: Molecular Function

| **Index** | **Name** | **P-value** | **Adjusted p-value** | **Odds Ratio** | **Combined score** |
| --- | --- | --- | --- | --- | --- |
| 1 | glucose:sodium symporter activity (GO:0005412) | 0.003495 | 0.02991 | 370.07 | 2093.27 |
| 2 | ciliary neurotrophic factor receptor binding (GO:0005127) | 0.004492 | 0.02991 | 277.53 | 1500.17 |
| 3 | MHC class II receptor activity (GO:0032395) | 0.004990 | 0.02991 | 246.68 | 1307.49 |
| 4 | carbohydrate:cation symporter activity (GO:0005402) | 0.005985 | 0.02991 | 201.81 | 1032.95 |
| 5 | basic amino acid transmembrane transporter activity (GO:0015174) | 0.006482 | 0.02991 | 184.98 | 932.06 |

GO: Cellular Component

| **Index** | **Name** | **P-value** | **Adjusted p-value** | **Odds Ratio** | **Combined score** |
| --- | --- | --- | --- | --- | --- |
| 1 | myosin filament (GO:0032982) | 0.002997 | 0.07355 | 444.11 | 2580.41 |
| 2 | intracellular vesicle (GO:0097708) | 0.005488 | 0.07355 | 222.00 | 1155.57 |
| 3 | MHC class II protein complex (GO:0042613) | 0.006482 | 0.07355 | 184.98 | 932.06 |
| 4 | supramolecular fiber (GO:0099512) | 0.009462 | 0.07355 | 123.28 | 574.57 |
| 5 | MHC protein complex (GO:0042611) | 0.009957 | 0.07355 | 116.79 | 538.34 |

Splenomegaly ([HP:0001744](https://hpo.jax.org/app/browse/term/HP:0001744))

KEGG Pathway

| **Index** | **Name** | **P-value** | **Adjusted p-value** | **Odds Ratio** | **Combined score** |
| --- | --- | --- | --- | --- | --- |
| 1 | Primary immunodeficiency | 5.941e-18 | 2.198e-16 | 1502.29 | 59587.92 |
| 2 | T cell receptor signaling pathway | 8.515e-10 | 1.575e-8 | 200.92 | 4196.00 |
| 3 | Th1 and Th2 cell differentiation | 8.619e-8 | 0.000001063 | 150.77 | 2452.58 |
| 4 | Autoimmune thyroid disease | 0.000002081 | 0.00001283 | 170.91 | 2236.02 |
| 5 | Hematopoietic cell lineage | 1.159e-7 | 0.000001072 | 139.61 | 2229.69 |

GO: Biological Process

| **Index** | **Name** | **P-value** | **Adjusted p-value** | **Odds Ratio** | **Combined score** |
| --- | --- | --- | --- | --- | --- |
| 1 | positive thymic T cell selection (GO:0045059) | 8.995e-10 | 1.151e-7 | 4283.14 | 89214.48 |
| 2 | V(D)J recombination (GO:0033151) | 0.00001235 | 0.0001580 | 555.03 | 6273.06 |
| 3 | T cell differentiation in thymus (GO:0033077) | 0.00002041 | 0.0002375 | 416.21 | 4494.85 |
| 4 | B cell activation involved in immune response (GO:0002312) | 0.00002690 | 0.0002869 | 356.71 | 3753.85 |
| 5 | regulation of T cell differentiation (GO:0045580) | 8.148e-7 | 0.00002027 | 237.55 | 3330.49 |

GO: Molecular Function

| **Index** | **Name** | **P-value** | **Adjusted p-value** | **Odds Ratio** | **Combined score** |
| --- | --- | --- | --- | --- | --- |
| 1 | endodeoxyribonuclease activity (GO:0004520) | 0.009957 | 0.06558 | 116.79 | 538.34 |
| 2 | phosphatidylinositol-3,5-bisphosphate binding (GO:0080025) | 0.01194 | 0.06558 | 96.46 | 427.12 |
| 3 | wide pore channel activity (GO:0022829) | 0.01243 | 0.06558 | 92.44 | 405.55 |
| 4 | phosphatidylinositol-3,4-bisphosphate binding (GO:0043325) | 0.01293 | 0.06558 | 88.73 | 385.85 |
| 5 | tumor necrosis factor receptor superfamily binding (GO:0032813) | 0.01392 | 0.06558 | 82.15 | 351.18 |

GO: Cellular Component

| **Index** | **Name** | **P-value** | **Adjusted p-value** | **Odds Ratio** | **Combined score** |
| --- | --- | --- | --- | --- | --- |
| 1 | T cell receptor complex (GO:0042101) | 1.975e-8 | 2.765e-7 | 951.48 | 16879.14 |
| 2 | alpha-beta T cell receptor complex (GO:0042105) | 0.000003371 | 0.00002360 | 1249.13 | 15739.23 |
| 3 | cytolytic granule (GO:0044194) | 0.003495 | 0.009787 | 370.07 | 2093.27 |
| 4 | clathrin-coated endocytic vesicle (GO:0045334) | 0.0007857 | 0.002750 | 59.96 | 428.66 |
| 5 | clathrin-coated endocytic vesicle membrane (GO:0030669) | 0.03398 | 0.07719 | 32.55 | 110.09 |

gi-symptom

Abdominal pain ([HP:0002027](https://hpo.jax.org/app/browse/term/HP:0002027))

KEGG Pathway

| **Index** | **Name** | **P-value** | **Adjusted p-value** | **Odds Ratio** | **Combined score** |
| --- | --- | --- | --- | --- | --- |
| 1 | Breast cancer | 2.916e-11 | 1.910e-9 | 211.16 | 5122.38 |
| 2 | Pathways in cancer | 1.006e-11 | 1.318e-9 | 148.89 | 3770.13 |
| 3 | Thyroid cancer | 6.931e-7 | 0.000006985 | 251.55 | 3567.44 |
| 4 | Central carbon metabolism in cancer | 2.843e-8 | 9.312e-7 | 201.25 | 3496.91 |
| 5 | Chronic myeloid leukemia | 3.973e-8 | 0.000001041 | 184.43 | 3142.85 |

GO: Biological Process

| **Index** | **Name** | **P-value** | **Adjusted p-value** | **Odds Ratio** | **Combined score** |
| --- | --- | --- | --- | --- | --- |
| 1 | positive regulation of histone H3-K9 acetylation (GO:2000617) | 0.000002248 | 0.0001365 | 1665.58 | 21661.60 |
| 2 | response to indole-3-methanol (GO:0071680) | 0.000002248 | 0.0001365 | 1665.58 | 21661.60 |
| 3 | cellular response to indole-3-methanol (GO:0071681) | 0.000002248 | 0.0001365 | 1665.58 | 21661.60 |
| 4 | positive regulation of histone H3-K4 methylation (GO:0051571) | 4.082e-8 | 0.00002302 | 713.50 | 12139.53 |
| 5 | regulation of histone H3-K4 methylation (GO:0051569) | 8.685e-8 | 0.00002449 | 535.02 | 8698.92 |

GO: Molecular Function

| **Index** | **Name** | **P-value** | **Adjusted p-value** | **Odds Ratio** | **Combined score** |
| --- | --- | --- | --- | --- | --- |
| 1 | I-SMAD binding (GO:0070411) | 0.00001750 | 0.0005512 | 454.07 | 4973.59 |
| 2 | RNA polymerase II general transcription initiation factor binding (GO:0001091) | 0.002498 | 0.01311 | 555.17 | 3326.77 |
| 3 | transcription coactivator binding (GO:0001223) | 0.00004255 | 0.0008935 | 277.39 | 2791.90 |
| 4 | transcription coregulator binding (GO:0001221) | 0.000002081 | 0.0001311 | 170.91 | 2236.02 |
| 5 | 1-phosphatidylinositol-4-phosphate 3-kinase activity (GO:0035005) | 0.003495 | 0.01694 | 370.07 | 2093.27 |

GO: Cellular Component

| **Index** | **Name** | **P-value** | **Adjusted p-value** | **Odds Ratio** | **Combined score** |
| --- | --- | --- | --- | --- | --- |
| 1 | phosphatidylinositol 3-kinase complex, class I (GO:0097651) | 0.002498 | 0.04538 | 555.17 | 3326.77 |
| 2 | Cul4A-RING E3 ubiquitin ligase complex (GO:0031464) | 0.005488 | 0.04538 | 222.00 | 1155.57 |
| 3 | beta-catenin-TCF complex (GO:1990907) | 0.006482 | 0.04538 | 184.98 | 932.06 |
| 4 | intercalated disc (GO:0014704) | 0.01540 | 0.04725 | 73.93 | 308.54 |
| 5 | catenin complex (GO:0016342) | 0.01540 | 0.04725 | 73.93 | 308.54 |

Abdominal symptom ([HP:0011458](https://hpo.jax.org/app/browse/term/HP:0011458))

KEGG Pathway

| **Index** | **Name** | **P-value** | **Adjusted p-value** | **Odds Ratio** | **Combined score** |
| --- | --- | --- | --- | --- | --- |
| 1 | Taste transduction | 0.000009013 | 0.00002704 | 102.79 | 1194.10 |
| 2 | Adrenergic signaling in cardiomyocytes | 0.002417 | 0.003626 | 33.52 | 201.95 |
| 3 | Dopaminergic synapse | 0.06409 | 0.06409 | 16.84 | 46.28 |

GO: Biological Process

| **Index** | **Name** | **P-value** | **Adjusted p-value** | **Odds Ratio** | **Combined score** |
| --- | --- | --- | --- | --- | --- |
| 1 | membrane depolarization during action potential (GO:0086010) | 1.882e-30 | 1.544e-28 | 199740.00 | 13671207.76 |
| 2 | membrane depolarization (GO:0051899) | 2.287e-29 | 9.375e-28 | 199680.00 | 13168480.83 |
| 3 | sodium ion transmembrane transport (GO:0035725) | 1.419e-24 | 2.909e-23 | 199130.00 | 10934655.87 |
| 4 | sodium ion transport (GO:0006814) | 2.029e-24 | 3.327e-23 | 199100.00 | 10861806.72 |
| 5 | inorganic cation transmembrane transport (GO:0098662) | 1.975e-19 | 2.314e-18 | 197260.00 | 8495671.35 |

GO: Molecular Function

| **Index** | **Name** | **P-value** | **Adjusted p-value** | **Odds Ratio** | **Combined score** |
| --- | --- | --- | --- | --- | --- |
| 1 | voltage-gated sodium channel activity (GO:0005248) | 2.291e-31 | 2.979e-30 | 199780.00 | 14094685.11 |
| 2 | sodium channel activity (GO:0005272) | 1.676e-28 | 1.089e-27 | 199620.00 | 12766920.69 |
| 3 | voltage-gated sodium channel activity involved in cardiac muscle cell action potential (GO:0086006) | 0.000002248 | 0.000009742 | 1665.58 | 21661.60 |
| 4 | nitric-oxide synthase binding (GO:0050998) | 0.003495 | 0.01038 | 370.07 | 2093.27 |
| 5 | sodium channel inhibitor activity (GO:0019871) | 0.003994 | 0.01038 | 317.19 | 1751.86 |

GO: Cellular Component

| **Index** | **Name** | **P-value** | **Adjusted p-value** | **Odds Ratio** | **Combined score** |
| --- | --- | --- | --- | --- | --- |
| 1 | voltage-gated sodium channel complex (GO:0001518) | 6.889e-33 | 9.645e-32 | 199830.00 | 14798476.21 |
| 2 | sodium channel complex (GO:0034706) | 1.158e-30 | 8.108e-30 | 199750.00 | 13768878.75 |
| 3 | integral component of plasma membrane (GO:0005887) | 4.006e-12 | 1.402e-11 | 185460.00 | 4867070.13 |
| 4 | axon (GO:0030424) | 4.513e-15 | 2.106e-14 | 403.96 | 13343.53 |
| 5 | neuron projection (GO:0043005) | 1.454e-11 | 4.072e-11 | 141.91 | 3541.27 |

Anorexia ([HP:0002039](https://hpo.jax.org/app/browse/term/HP:0002039))

KEGG Pathway

| **Index** | **Name** | **P-value** | **Adjusted p-value** | **Odds Ratio** | **Combined score** |
| --- | --- | --- | --- | --- | --- |
| 1 | Chronic myeloid leukemia | 0.000006207 | 0.0002193 | 116.93 | 1401.96 |
| 2 | Primary immunodeficiency | 0.0001567 | 0.001186 | 138.57 | 1214.06 |
| 3 | PD-L1 expression and PD-1 checkpoint pathway in cancer | 0.000009993 | 0.0002468 | 99.19 | 1142.03 |
| 4 | Homologous recombination | 0.0001826 | 0.001290 | 127.89 | 1100.92 |
| 5 | AGE-RAGE signaling pathway in diabetic complications | 0.00001419 | 0.0002468 | 87.89 | 981.15 |

GO: Biological Process

| **Index** | **Name** | **P-value** | **Adjusted p-value** | **Odds Ratio** | **Combined score** |
| --- | --- | --- | --- | --- | --- |
| 1 | positive regulation of histone H3-K9 acetylation (GO:2000617) | 0.000002248 | 0.0002439 | 1665.58 | 21661.60 |
| 2 | positive thymic T cell selection (GO:0045059) | 0.000002248 | 0.0002439 | 1665.58 | 21661.60 |
| 3 | regulation of histone H3-K9 acetylation (GO:2000615) | 0.00001235 | 0.001072 | 555.03 | 6273.06 |
| 4 | positive regulation of histone H3-K4 methylation (GO:0051571) | 0.00002354 | 0.001459 | 384.17 | 4094.01 |
| 5 | positive regulation of alpha-beta T cell proliferation (GO:0046641) | 0.00002690 | 0.001459 | 356.71 | 3753.85 |

GO: Molecular Function

| **Index** | **Name** | **P-value** | **Adjusted p-value** | **Odds Ratio** | **Combined score** |
| --- | --- | --- | --- | --- | --- |
| 1 | Y-form DNA binding (GO:0000403) | 0.002498 | 0.02034 | 555.17 | 3326.77 |
| 2 | R-SMAD binding (GO:0070412) | 0.00003830 | 0.002183 | 293.72 | 2987.15 |
| 3 | non-membrane spanning protein tyrosine phosphatase activity (GO:0004726) | 0.002997 | 0.02135 | 444.11 | 2580.41 |
| 4 | cyclin-dependent protein serine/threonine kinase inhibitor activity (GO:0004861) | 0.004990 | 0.02844 | 246.68 | 1307.49 |
| 5 | I-SMAD binding (GO:0070411) | 0.006482 | 0.03194 | 184.98 | 932.06 |

GO: Cellular Component

| **Index** | **Name** | **P-value** | **Adjusted p-value** | **Odds Ratio** | **Combined score** |
| --- | --- | --- | --- | --- | --- |
| 1 | T cell receptor complex (GO:0042101) | 1.975e-8 | 5.136e-7 | 951.48 | 16879.14 |
| 2 | alpha-beta T cell receptor complex (GO:0042105) | 0.000003371 | 0.00004383 | 1249.13 | 15739.23 |
| 3 | Cul4A-RING E3 ubiquitin ligase complex (GO:0031464) | 0.005488 | 0.03890 | 222.00 | 1155.57 |
| 4 | early phagosome (GO:0032009) | 0.005985 | 0.03890 | 201.81 | 1032.95 |
| 5 | Cul4-RING E3 ubiquitin ligase complex (GO:0080008) | 0.01687 | 0.07312 | 67.20 | 274.29 |

Bowel incontinence ([HP:0002607](https://hpo.jax.org/app/browse/term/HP:0002607))

KEGG Pathway

| **Index** | **Name** | **P-value** | **Adjusted p-value** | **Odds Ratio** | **Combined score** |
| --- | --- | --- | --- | --- | --- |
| 1 | Glycosaminoglycan degradation | 0.009462 | 0.08929 | 123.28 | 574.57 |
| 2 | Tyrosine metabolism | 0.01786 | 0.08929 | 63.35 | 255.00 |
| 3 | Steroid hormone biosynthesis | 0.03009 | 0.09155 | 36.91 | 129.31 |
| 4 | ECM-receptor interaction | 0.04315 | 0.09155 | 25.42 | 79.89 |
| 5 | Hematopoietic cell lineage | 0.04842 | 0.09155 | 22.55 | 68.29 |

GO: Biological Process

| **Index** | **Name** | **P-value** | **Adjusted p-value** | **Odds Ratio** | **Combined score** |
| --- | --- | --- | --- | --- | --- |
| 1 | middle ear morphogenesis (GO:0042474) | 0.002498 | 0.03291 | 555.17 | 3326.77 |
| 2 | embryonic viscerocranium morphogenesis (GO:0048703) | 0.002498 | 0.03291 | 555.17 | 3326.77 |
| 3 | enamel mineralization (GO:0070166) | 0.002498 | 0.03291 | 555.17 | 3326.77 |
| 4 | coronary artery morphogenesis (GO:0060982) | 0.002498 | 0.03291 | 555.17 | 3326.77 |
| 5 | establishment of planar polarity (GO:0001736) | 0.002997 | 0.03291 | 444.11 | 2580.41 |

GO: Molecular Function

| **Index** | **Name** | **P-value** | **Adjusted p-value** | **Odds Ratio** | **Combined score** |
| --- | --- | --- | --- | --- | --- |
| 1 | O-methyltransferase activity (GO:0008171) | 0.004990 | 0.05988 | 246.68 | 1307.49 |
| 2 | hydrolase activity, hydrolyzing O-glycosyl compounds (GO:0004553) | 0.01835 | 0.07418 | 61.59 | 246.23 |
| 3 | S-adenosylmethionine-dependent methyltransferase activity (GO:0008757) | 0.02130 | 0.07418 | 52.77 | 203.13 |
| 4 | methyltransferase activity (GO:0008168) | 0.02473 | 0.07418 | 45.22 | 167.30 |
| 5 | protein homodimerization activity (GO:0042803) | 0.2762 | 0.4520 | 3.39 | 4.36 |

GO: Cellular Component

| **Index** | **Name** | **P-value** | **Adjusted p-value** | **Odds Ratio** | **Combined score** |
| --- | --- | --- | --- | --- | --- |
| 1 | sarcoplasmic reticulum (GO:0016529) | 0.02228 | 0.1607 | 50.37 | 191.61 |
| 2 | intermediate filament (GO:0005882) | 0.02473 | 0.1607 | 45.22 | 167.30 |
| 3 | intermediate filament cytoskeleton (GO:0045111) | 0.04122 | 0.1786 | 26.65 | 84.98 |
| 4 | intrinsic component of endoplasmic reticulum membrane (GO:0031227) | 0.05889 | 0.1789 | 18.40 | 52.10 |
| 5 | integral component of endoplasmic reticulum membrane (GO:0030176) | 0.06879 | 0.1789 | 15.64 | 41.87 |

Constipation ([HP:0002019](https://hpo.jax.org/app/browse/term/HP:0002019))

KEGG Pathway

| **Index** | **Name** | **P-value** | **Adjusted p-value** | **Odds Ratio** | **Combined score** |
| --- | --- | --- | --- | --- | --- |
| 1 | Hedgehog signaling pathway | 0.0003416 | 0.003675 | 92.30 | 736.71 |
| 2 | Basal cell carcinoma | 0.0004323 | 0.003675 | 81.68 | 632.69 |
| 3 | Gastric cancer | 0.002386 | 0.01014 | 33.75 | 203.77 |
| 4 | Melanoma | 0.03543 | 0.1205 | 31.17 | 104.12 |
| 5 | Pathways in cancer | 0.001943 | 0.01014 | 15.80 | 98.63 |

GO: Biological Process

| **Index** | **Name** | **P-value** | **Adjusted p-value** | **Odds Ratio** | **Combined score** |
| --- | --- | --- | --- | --- | --- |
| 1 | pituitary gland development (GO:0021983) | 3.147e-9 | 2.087e-7 | 2141.36 | 41921.23 |
| 2 | positive regulation of T cell differentiation in thymus (GO:0033089) | 0.000003371 | 0.00004792 | 1249.13 | 15739.23 |
| 3 | dorsal/ventral pattern formation (GO:0009953) | 3.331e-10 | 6.629e-8 | 665.67 | 14526.48 |
| 4 | proximal/distal pattern formation (GO:0009954) | 0.000006290 | 0.00007823 | 832.67 | 9972.50 |
| 5 | dopaminergic neuron differentiation (GO:0071542) | 1.021e-7 | 0.000002033 | 503.52 | 8105.11 |

GO: Molecular Function

| **Index** | **Name** | **P-value** | **Adjusted p-value** | **Odds Ratio** | **Combined score** |
| --- | --- | --- | --- | --- | --- |
| 1 | type 1 fibroblast growth factor receptor binding (GO:0005105) | 0.002498 | 0.006744 | 555.17 | 3326.77 |
| 2 | type 2 fibroblast growth factor receptor binding (GO:0005111) | 0.002498 | 0.006744 | 555.17 | 3326.77 |
| 3 | morphogen activity (GO:0016015) | 0.003495 | 0.007864 | 370.07 | 2093.27 |
| 4 | patched binding (GO:0005113) | 0.003495 | 0.007864 | 370.07 | 2093.27 |
| 5 | eukaryotic initiation factor 4E binding (GO:0008190) | 0.004990 | 0.01036 | 246.68 | 1307.49 |

GO: Cellular Component

| **Index** | **Name** | **P-value** | **Adjusted p-value** | **Odds Ratio** | **Combined score** |
| --- | --- | --- | --- | --- | --- |
| 1 | nucleus (GO:0005634) | 0.001775 | 0.01775 | 8.09 | 51.21 |
| 2 | intracellular membrane-bounded organelle (GO:0043231) | 0.004410 | 0.02205 | 6.66 | 36.14 |
| 3 | membrane raft (GO:0045121) | 0.07859 | 0.2620 | 13.60 | 34.59 |
| 4 | cilium (GO:0005929) | 0.1124 | 0.2675 | 9.30 | 20.33 |
| 5 | endoplasmic reticulum lumen (GO:0005788) | 0.1337 | 0.2675 | 7.71 | 15.51 |

Diarrhea ([HP:0002014](https://hpo.jax.org/app/browse/term/HP:0002014))

KEGG Pathway

| **Index** | **Name** | **P-value** | **Adjusted p-value** | **Odds Ratio** | **Combined score** |
| --- | --- | --- | --- | --- | --- |
| 1 | Primary immunodeficiency | 3.458e-21 | 1.037e-19 | 2661.33 | 125385.00 |
| 2 | T cell receptor signaling pathway | 8.515e-10 | 1.277e-8 | 200.92 | 4196.00 |
| 3 | B cell receptor signaling pathway | 0.000007524 | 0.00006625 | 109.41 | 1290.72 |
| 4 | PD-L1 expression and PD-1 checkpoint pathway in cancer | 0.000009993 | 0.00006625 | 99.19 | 1142.03 |
| 5 | Th1 and Th2 cell differentiation | 0.00001104 | 0.00006625 | 95.83 | 1093.80 |

GO: Biological Process

| **Index** | **Name** | **P-value** | **Adjusted p-value** | **Odds Ratio** | **Combined score** |
| --- | --- | --- | --- | --- | --- |
| 1 | positive thymic T cell selection (GO:0045059) | 8.995e-10 | 2.541e-8 | 4283.14 | 89214.48 |
| 2 | B cell activation (GO:0042113) | 1.020e-12 | 5.761e-11 | 378.06 | 10438.72 |
| 3 | B cell receptor signaling pathway (GO:0050853) | 1.451e-9 | 3.278e-8 | 443.56 | 9026.96 |
| 4 | antigen receptor-mediated signaling pathway (GO:0050851) | 6.059e-13 | 5.761e-11 | 259.71 | 7306.12 |
| 5 | V(D)J recombination (GO:0033151) | 0.00001235 | 0.0001163 | 555.03 | 6273.06 |

GO: Molecular Function

| **Index** | **Name** | **P-value** | **Adjusted p-value** | **Odds Ratio** | **Combined score** |
| --- | --- | --- | --- | --- | --- |
| 1 | endodeoxyribonuclease activity (GO:0004520) | 0.009957 | 0.06431 | 116.79 | 538.34 |
| 2 | phosphatidylinositol-3,5-bisphosphate binding (GO:0080025) | 0.01194 | 0.06431 | 96.46 | 427.12 |
| 3 | phosphatidylinositol-3,4-bisphosphate binding (GO:0043325) | 0.01293 | 0.06431 | 88.73 | 385.85 |
| 4 | tumor necrosis factor receptor superfamily binding (GO:0032813) | 0.01392 | 0.06431 | 82.15 | 351.18 |
| 5 | phosphatidylinositol-3,4,5-trisphosphate binding (GO:0005547) | 0.01737 | 0.06431 | 65.22 | 264.33 |

GO: Cellular Component

| **Index** | **Name** | **P-value** | **Adjusted p-value** | **Odds Ratio** | **Combined score** |
| --- | --- | --- | --- | --- | --- |
| 1 | T cell receptor complex (GO:0042101) | 1.975e-8 | 2.765e-7 | 951.48 | 16879.14 |
| 2 | alpha-beta T cell receptor complex (GO:0042105) | 0.000003371 | 0.00002360 | 1249.13 | 15739.23 |
| 3 | clathrin-coated endocytic vesicle (GO:0045334) | 0.0007857 | 0.002750 | 59.96 | 428.66 |
| 4 | multivesicular body (GO:0005771) | 0.02326 | 0.05427 | 48.17 | 181.19 |
| 5 | membrane raft (GO:0045121) | 0.002846 | 0.007968 | 30.79 | 180.49 |

Nausea ([HP:0002018](https://hpo.jax.org/app/browse/term/HP:0002018))

KEGG Pathway

| **Index** | **Name** | **P-value** | **Adjusted p-value** | **Odds Ratio** | **Combined score** |
| --- | --- | --- | --- | --- | --- |
| 1 | Central carbon metabolism in cancer | 0.0005336 | 0.007471 | 73.24 | 551.94 |
| 2 | Citrate cycle (TCA cycle) | 0.01490 | 0.07824 | 76.48 | 321.69 |
| 3 | Thyroid cancer | 0.01835 | 0.08564 | 61.59 | 246.23 |
| 4 | MAPK signaling pathway | 0.0003495 | 0.007340 | 29.01 | 230.90 |
| 5 | Pathways in cancer | 0.00009084 | 0.003815 | 24.62 | 229.14 |

GO: Biological Process

| **Index** | **Name** | **P-value** | **Adjusted p-value** | **Odds Ratio** | **Combined score** |
| --- | --- | --- | --- | --- | --- |
| 1 | regulation of phospholipase C activity (GO:1900274) | 0.00002354 | 0.003932 | 384.17 | 4094.01 |
| 2 | glial cell-derived neurotrophic factor receptor signaling pathway (GO:0035860) | 0.002498 | 0.02831 | 555.17 | 3326.77 |
| 3 | atrial cardiac muscle cell membrane repolarization (GO:0099624) | 0.002498 | 0.02831 | 555.17 | 3326.77 |
| 4 | positive regulation of smooth muscle cell differentiation (GO:0051152) | 0.002498 | 0.02831 | 555.17 | 3326.77 |
| 5 | membrane repolarization during atrial cardiac muscle cell action potential (GO:0098914) | 0.002498 | 0.02831 | 555.17 | 3326.77 |

GO: Molecular Function

| **Index** | **Name** | **P-value** | **Adjusted p-value** | **Odds Ratio** | **Combined score** |
| --- | --- | --- | --- | --- | --- |
| 1 | RNA polymerase II general transcription initiation factor binding (GO:0001091) | 0.002498 | 0.02498 | 555.17 | 3326.77 |
| 2 | Y-form DNA binding (GO:0000403) | 0.002498 | 0.02498 | 555.17 | 3326.77 |
| 3 | phosphatidylethanolamine binding (GO:0008429) | 0.004492 | 0.02807 | 277.53 | 1500.17 |
| 4 | voltage-gated potassium channel activity involved in ventricular cardiac muscle cell action potential repolarization (GO:1902282) | 0.005985 | 0.03325 | 201.81 | 1032.95 |
| 5 | voltage-gated potassium channel activity involved in cardiac muscle cell action potential repolarization (GO:0086008) | 0.007476 | 0.03738 | 158.54 | 776.21 |

GO: Cellular Component

| **Index** | **Name** | **P-value** | **Adjusted p-value** | **Odds Ratio** | **Combined score** |
| --- | --- | --- | --- | --- | --- |
| 1 | MLL1 complex (GO:0071339) | 0.01293 | 0.06103 | 88.73 | 385.85 |
| 2 | MLL1/2 complex (GO:0044665) | 0.01293 | 0.06103 | 88.73 | 385.85 |
| 3 | axon (GO:0030424) | 0.0001190 | 0.002976 | 42.19 | 381.27 |
| 4 | vesicle membrane (GO:0012506) | 0.03591 | 0.09824 | 30.74 | 102.25 |
| 5 | voltage-gated potassium channel complex (GO:0008076) | 0.03591 | 0.09824 | 30.74 | 102.25 |

Poor appetite ([HP:0004396](https://hpo.jax.org/app/browse/term/HP:0004396))

KEGG Pathway

| **Index** | **Name** | **P-value** | **Adjusted p-value** | **Odds Ratio** | **Combined score** |
| --- | --- | --- | --- | --- | --- |
| 1 | Thyroid cancer | 0.0001485 | 0.005965 | 142.54 | 1256.48 |
| 2 | Homologous recombination | 0.0001826 | 0.005965 | 127.89 | 1100.92 |
| 3 | Fanconi anemia pathway | 0.0003175 | 0.007780 | 95.86 | 772.11 |
| 4 | Apelin signaling pathway | 0.00003643 | 0.003570 | 63.51 | 649.04 |
| 5 | Chronic myeloid leukemia | 0.0006287 | 0.009585 | 67.28 | 496.00 |

GO: Biological Process

| **Index** | **Name** | **P-value** | **Adjusted p-value** | **Odds Ratio** | **Combined score** |
| --- | --- | --- | --- | --- | --- |
| 1 | positive regulation of histone H3-K9 acetylation (GO:2000617) | 0.000002248 | 0.0001776 | 1665.58 | 21661.60 |
| 2 | regulation of histone H3-K9 acetylation (GO:2000615) | 0.00001235 | 0.0007315 | 555.03 | 6273.06 |
| 3 | positive regulation of histone H3-K4 methylation (GO:0051571) | 0.00002354 | 0.001116 | 384.17 | 4094.01 |
| 4 | actin-myosin filament sliding (GO:0033275) | 7.523e-7 | 0.00008915 | 244.35 | 3445.31 |
| 5 | muscle filament sliding (GO:0030049) | 7.523e-7 | 0.00008915 | 244.35 | 3445.31 |

GO: Molecular Function

| **Index** | **Name** | **P-value** | **Adjusted p-value** | **Odds Ratio** | **Combined score** |
| --- | --- | --- | --- | --- | --- |
| 1 | myosin heavy chain binding (GO:0032036) | 0.003495 | 0.03337 | 370.07 | 2093.27 |
| 2 | arginine binding (GO:0034618) | 0.003994 | 0.03337 | 317.19 | 1751.86 |
| 3 | L-aspartate transmembrane transporter activity (GO:0015183) | 0.004492 | 0.03337 | 277.53 | 1500.17 |
| 4 | I-SMAD binding (GO:0070411) | 0.006482 | 0.03337 | 184.98 | 932.06 |
| 5 | L-glutamate transmembrane transporter activity (GO:0005313) | 0.006979 | 0.03337 | 170.74 | 847.70 |

GO: Cellular Component

| **Index** | **Name** | **P-value** | **Adjusted p-value** | **Odds Ratio** | **Combined score** |
| --- | --- | --- | --- | --- | --- |
| 1 | cardiac myofibril (GO:0097512) | 0.002498 | 0.01399 | 555.17 | 3326.77 |
| 2 | actin filament (GO:0005884) | 0.000005272 | 0.0001476 | 123.73 | 1503.75 |
| 3 | actin cytoskeleton (GO:0015629) | 0.00001191 | 0.0001668 | 42.05 | 476.73 |
| 4 | excitatory synapse (GO:0060076) | 0.01243 | 0.04786 | 92.44 | 405.55 |
| 5 | myofibril (GO:0030016) | 0.01342 | 0.04786 | 85.32 | 367.79 |

Vomiting ([HP:0002013](https://hpo.jax.org/app/browse/term/HP:0002013))

KEGG Pathway

| **Index** | **Name** | **P-value** | **Adjusted p-value** | **Odds Ratio** | **Combined score** |
| --- | --- | --- | --- | --- | --- |
| 1 | Oxidative phosphorylation | 1.586e-11 | 3.250e-10 | 234.60 | 5833.97 |
| 2 | Thermogenesis | 3.007e-12 | 1.233e-10 | 204.97 | 5437.89 |
| 3 | Retrograde endocannabinoid signaling | 3.038e-11 | 4.124e-10 | 209.66 | 5077.41 |
| 4 | Non-alcoholic fatty liver disease | 4.023e-11 | 4.124e-10 | 199.74 | 4781.09 |
| 5 | Diabetic cardiomyopathy | 2.061e-10 | 1.408e-9 | 150.71 | 3361.18 |

GO: Biological Process

| **Index** | **Name** | **P-value** | **Adjusted p-value** | **Odds Ratio** | **Combined score** |
| --- | --- | --- | --- | --- | --- |
| 1 | mitochondrial respiratory chain complex I assembly (GO:0032981) | 1.353e-19 | 1.583e-17 | 1595.20 | 69306.12 |
| 2 | NADH dehydrogenase complex assembly (GO:0010257) | 1.353e-19 | 1.583e-17 | 1595.20 | 69306.12 |
| 3 | mitochondrial respiratory chain complex assembly (GO:0033108) | 5.457e-18 | 4.257e-16 | 971.12 | 38601.66 |
| 4 | mitochondrial electron transport, NADH to ubiquinone (GO:0006120) | 7.666e-15 | 4.485e-13 | 907.14 | 29483.69 |
| 5 | aerobic electron transport chain (GO:0019646) | 3.065e-13 | 1.305e-11 | 467.02 | 13456.41 |

GO: Molecular Function

| **Index** | **Name** | **P-value** | **Adjusted p-value** | **Odds Ratio** | **Combined score** |
| --- | --- | --- | --- | --- | --- |
| 1 | NADH dehydrogenase (quinone) activity (GO:0050136) | 3.049e-12 | 3.507e-11 | 665.33 | 17642.09 |
| 2 | NADH dehydrogenase (ubiquinone) activity (GO:0008137) | 3.049e-12 | 3.507e-11 | 665.33 | 17642.09 |
| 3 | oxidoreduction-driven active transmembrane transporter activity (GO:0015453) | 4.283e-11 | 3.284e-10 | 376.17 | 8980.56 |
| 4 | morphogen activity (GO:0016015) | 0.003495 | 0.01340 | 370.07 | 2093.27 |
| 5 | NADH dehydrogenase activity (GO:0003954) | 0.003495 | 0.01340 | 370.07 | 2093.27 |

GO: Cellular Component

| **Index** | **Name** | **P-value** | **Adjusted p-value** | **Odds Ratio** | **Combined score** |
| --- | --- | --- | --- | --- | --- |
| 1 | mitochondrial respiratory chain complex I (GO:0005747) | 1.232e-14 | 1.355e-13 | 831.42 | 26628.24 |
| 2 | respiratory chain complex I (GO:0045271) | 1.232e-14 | 1.355e-13 | 831.42 | 26628.24 |
| 3 | mitochondrial inner membrane (GO:0005743) | 3.444e-11 | 2.525e-10 | 142.97 | 3444.50 |
| 4 | mitochondrial envelope (GO:0005740) | 2.343e-9 | 8.590e-9 | 162.85 | 3236.19 |
| 5 | organelle inner membrane (GO:0019866) | 5.010e-11 | 2.756e-10 | 135.26 | 3207.90 |
