## Supplementary Table 1-3 for "Long COVID: G Protein-Coupled Receptors (GPCRs) responsible for persistent post-COVID symptoms": Supplementary File 2.9-Pulmonary.docx

Pulmonary-finding

Abnormality on pulmonary function testing ([HP:0030878](https://hpo.jax.org/app/browse/term/HP:0030878))

KEGG Pathway

| **Index** | **Name** | **P-value** | **Adjusted p-value** | **Odds Ratio** | **Combined score** |
| --- | --- | --- | --- | --- | --- |
| 1 | Neuroactive ligand-receptor interaction | 0.00001608 | 0.0001608 | 38.88 | 429.14 |
| 2 | Primary immunodeficiency | 0.01884 | 0.07259 | 59.92 | 237.98 |
| 3 | Autoimmune thyroid disease | 0.02619 | 0.07259 | 42.60 | 155.17 |
| 4 | B cell receptor signaling pathway | 0.03978 | 0.07259 | 27.65 | 89.16 |
| 5 | Rheumatoid arthritis | 0.04555 | 0.07259 | 24.03 | 74.23 |

GO: Biological Process

| **Index** | **Name** | **P-value** | **Adjusted p-value** | **Odds Ratio** | **Combined score** |
| --- | --- | --- | --- | --- | --- |
| 1 | synaptic transmission, cholinergic (GO:0007271) | 8.685e-8 | 0.000003648 | 535.02 | 8698.92 |
| 2 | neuromuscular junction development (GO:0007528) | 1.812e-7 | 0.000004033 | 407.53 | 6326.45 |
| 3 | negative regulation of regulatory T cell differentiation (GO:0045590) | 0.002498 | 0.01614 | 555.17 | 3326.77 |
| 4 | neuromuscular process (GO:0050905) | 0.00003830 | 0.0003971 | 293.72 | 2987.15 |
| 5 | postsynaptic membrane organization (GO:0001941) | 0.00003830 | 0.0003971 | 293.72 | 2987.15 |

GO: Molecular Function

| **Index** | **Name** | **P-value** | **Adjusted p-value** | **Odds Ratio** | **Combined score** |
| --- | --- | --- | --- | --- | --- |
| 1 | neurotransmitter receptor activity involved in regulation of postsynaptic membrane potential (GO:0099529) | 1.878e-10 | 4.131e-9 | 783.25 | 17541.62 |
| 2 | transmitter-gated ion channel activity involved in regulation of postsynaptic membrane potential (GO:1904315) | 8.580e-10 | 9.438e-9 | 511.90 | 10686.61 |
| 3 | transmitter-gated ion channel activity (GO:0022824) | 1.451e-9 | 1.064e-8 | 443.56 | 9026.96 |
| 4 | postsynaptic neurotransmitter receptor activity (GO:0098960) | 1.191e-7 | 6.068e-7 | 475.52 | 7581.26 |
| 5 | acetylcholine receptor activity (GO:0015464) | 1.379e-7 | 6.068e-7 | 450.47 | 7115.97 |

GO: Cellular Component

| **Index** | **Name** | **P-value** | **Adjusted p-value** | **Odds Ratio** | **Combined score** |
| --- | --- | --- | --- | --- | --- |
| 1 | neuromuscular junction (GO:0031594) | 4.712e-12 | 5.183e-11 | 604.76 | 15772.67 |
| 2 | acetylcholine-gated channel complex (GO:0005892) | 3.267e-8 | 1.198e-7 | 778.40 | 13417.28 |
| 3 | ion channel complex (GO:0034702) | 5.841e-7 | 0.000001606 | 267.29 | 3836.51 |
| 4 | integral component of plasma membrane (GO:0005887) | 3.020e-8 | 1.198e-7 | 51.30 | 888.24 |
| 5 | voltage-gated sodium channel complex (GO:0001518) | 0.008469 | 0.01553 | 138.71 | 661.82 |

Airway obstruction ([HP:0006536](https://hpo.jax.org/app/browse/term/HP:0006536))

KEGG Pathway

| **Index** | **Name** | **P-value** | **Adjusted p-value** | **Odds Ratio** | **Combined score** |
| --- | --- | --- | --- | --- | --- |
| 1 | Huntington disease | 0.1429 | 0.2137 | 7.17 | 13.95 |
| 2 | Amyotrophic lateral sclerosis | 0.1678 | 0.2137 | 6.01 | 10.72 |
| 3 | Pathways of neurodegeneration | 0.2137 | 0.2137 | 4.57 | 7.06 |

GO:Biological Process

| **Index** | **Name** | **P-value** | **Adjusted p-value** | **Odds Ratio** | **Combined score** |
| --- | --- | --- | --- | --- | --- |
| 1 | axoneme assembly (GO:0035082) | 3.163e-15 | 9.488e-14 | 1069.39 | 35704.16 |
| 2 | axonemal dynein complex assembly (GO:0070286) | 1.125e-9 | 1.688e-8 | 475.29 | 9793.38 |
| 3 | regulation of cilium movement (GO:0003352) | 0.00001010 | 0.00004330 | 624.44 | 7182.69 |
| 4 | cilium-dependent cell motility (GO:0060285) | 1.586e-7 | 0.000001189 | 427.93 | 6700.13 |
| 5 | regulation of microtubule-based movement (GO:0060632) | 0.00001235 | 0.00004629 | 555.03 | 6273.06 |

GO: Molecular Function

| **Index** | **Name** | **P-value** | **Adjusted p-value** | **Odds Ratio** | **Combined score** |
| --- | --- | --- | --- | --- | --- |
| 1 | microtubule motor activity (GO:0003777) | 0.02766 | 0.04878 | 40.27 | 144.50 |
| 2 | motor activity (GO:0003774) | 0.03252 | 0.04878 | 34.06 | 116.68 |
| 3 | protein homodimerization activity (GO:0042803) | 0.2762 | 0.2762 | 3.39 | 4.36 |

GO: Cellular Component

| **Index** | **Name** | **P-value** | **Adjusted p-value** | **Odds Ratio** | **Combined score** |
| --- | --- | --- | --- | --- | --- |
| 1 | motile cilium (GO:0031514) | 0.000004434 | 0.00004434 | 131.37 | 1619.33 |
| 2 | outer dynein arm (GO:0036157) | 0.004492 | 0.01123 | 277.53 | 1500.17 |
| 3 | 9+2 motile cilium (GO:0097729) | 0.0003539 | 0.001180 | 90.61 | 720.07 |
| 4 | cilium (GO:0005929) | 0.0001854 | 0.0009270 | 36.18 | 310.92 |
| 5 | sperm flagellum (GO:0036126) | 0.02570 | 0.05141 | 43.44 | 159.04 |

Ground-glass opacification ([HP:0025179](https://hpo.jax.org/app/browse/term/HP:0025179))

KEGG Pathway

| **Index** | **Name** | **P-value** | **Adjusted p-value** | **Odds Ratio** | **Combined score** |
| --- | --- | --- | --- | --- | --- |
| 1 | ABC transporters | 0.02007 | 0.1185 | 56.67 | 221.48 |
| 2 | Arrhythmogenic right ventricular cardiomyopathy | 0.03413 | 0.1185 | 32.75 | 110.63 |
| 3 | RNA degradation | 0.03500 | 0.1185 | 31.91 | 106.98 |
| 4 | TGF-beta signaling pathway | 0.04152 | 0.1185 | 26.74 | 85.09 |
| 5 | Fluid shear stress and atherosclerosis | 0.06085 | 0.1185 | 17.98 | 50.34 |

GO: Biological Process

| **Index** | **Name** | **P-value** | **Adjusted p-value** | **Odds Ratio** | **Combined score** |
| --- | --- | --- | --- | --- | --- |
| 1 | positive regulation of chromosome organization (GO:2001252) | 0.00001884 | 0.002717 | 439.08 | 4776.89 |
| 2 | establishment of protein localization to telomere (GO:0070200) | 0.002248 | 0.02312 | 624.59 | 3808.55 |
| 3 | positive regulation of cardioblast differentiation (GO:0051891) | 0.002248 | 0.02312 | 624.59 | 3808.55 |
| 4 | lung morphogenesis (GO:0060425) | 0.002248 | 0.02312 | 624.59 | 3808.55 |
| 5 | tricuspid valve development (GO:0003175) | 0.002248 | 0.02312 | 624.59 | 3808.55 |

GO: Molecular Function

| **Index** | **Name** | **P-value** | **Adjusted p-value** | **Odds Ratio** | **Combined score** |
| --- | --- | --- | --- | --- | --- |
| 1 | intronic transcription regulatory region sequence-specific DNA binding (GO:0001161) | 0.002248 | 0.01537 | 624.59 | 3808.55 |
| 2 | phosphatidylcholine flippase activity (GO:0140345) | 0.002248 | 0.01537 | 624.59 | 3808.55 |
| 3 | phosphatidylcholine transfer activity (GO:0120019) | 0.002248 | 0.01537 | 624.59 | 3808.55 |
| 4 | cell adhesive protein binding involved in bundle of His cell-Purkinje myocyte communication (GO:0086083) | 0.002248 | 0.01537 | 624.59 | 3808.55 |
| 5 | RNA-directed DNA polymerase activity (GO:0003964) | 0.002697 | 0.01537 | 499.65 | 2955.69 |

GO: Cellular Component

| **Index** | **Name** | **P-value** | **Adjusted p-value** | **Odds Ratio** | **Combined score** |
| --- | --- | --- | --- | --- | --- |
| 1 | alveolar lamellar body (GO:0097208) | 0.002697 | 0.03580 | 499.65 | 2955.69 |
| 2 | transferase complex, transferring phosphorus-containing groups (GO:0061695) | 0.004043 | 0.03580 | 312.23 | 1720.62 |
| 3 | spanning component of plasma membrane (GO:0044214) | 0.005836 | 0.03580 | 208.11 | 1070.48 |
| 4 | lamellar body (GO:0042599) | 0.005836 | 0.03580 | 208.11 | 1070.48 |
| 5 | spanning component of membrane (GO:0089717) | 0.008966 | 0.04355 | 131.39 | 619.44 |

Hypoxemia ([HP:0012418](https://hpo.jax.org/app/browse/term/HP:0012418))

KEGG Pathway

| **Index** | **Name** | **P-value** | **Adjusted p-value** | **Odds Ratio** | **Combined score** |
| --- | --- | --- | --- | --- | --- |
| 1 | Cardiac muscle contraction | 0.0008229 | 0.01301 | 58.54 | 415.82 |
| 2 | Hypertrophic cardiomyopathy | 0.0008802 | 0.01301 | 56.54 | 397.78 |
| 3 | Dilated cardiomyopathy | 0.001001 | 0.01301 | 52.91 | 365.49 |
| 4 | Asthma | 0.01540 | 0.08420 | 73.93 | 308.54 |
| 5 | Thyroid cancer | 0.01835 | 0.08420 | 61.59 | 246.23 |

GO: Biological Process

| **Index** | **Name** | **P-value** | **Adjusted p-value** | **Odds Ratio** | **Combined score** |
| --- | --- | --- | --- | --- | --- |
| 1 | actin-myosin filament sliding (GO:0033275) | 7.523e-7 | 0.00008577 | 244.35 | 3445.31 |
| 2 | muscle filament sliding (GO:0030049) | 7.523e-7 | 0.00008577 | 244.35 | 3445.31 |
| 3 | positive regulation of myoblast proliferation (GO:2000288) | 0.002498 | 0.01785 | 555.17 | 3326.77 |
| 4 | negative regulation of transforming growth factor beta1 production (GO:0032911) | 0.002498 | 0.01785 | 555.17 | 3326.77 |
| 5 | positive regulation of CD4-positive, CD25-positive, alpha-beta regulatory T cell differentiation (GO:0032831) | 0.002498 | 0.01785 | 555.17 | 3326.77 |

GO: Molecular Function

| **Index** | **Name** | **P-value** | **Adjusted p-value** | **Odds Ratio** | **Combined score** |
| --- | --- | --- | --- | --- | --- |
| 1 | intronic transcription regulatory region sequence-specific DNA binding (GO:0001161) | 0.002498 | 0.02231 | 555.17 | 3326.77 |
| 2 | phosphatidylcholine flippase activity (GO:0140345) | 0.002498 | 0.02231 | 555.17 | 3326.77 |
| 3 | phosphatidylcholine transfer activity (GO:0120019) | 0.002498 | 0.02231 | 555.17 | 3326.77 |
| 4 | myosin heavy chain binding (GO:0032036) | 0.003495 | 0.02231 | 370.07 | 2093.27 |
| 5 | CD4 receptor binding (GO:0042609) | 0.003994 | 0.02231 | 317.19 | 1751.86 |

GO: Cellular Component

| **Index** | **Name** | **P-value** | **Adjusted p-value** | **Odds Ratio** | **Combined score** |
| --- | --- | --- | --- | --- | --- |
| 1 | cardiac myofibril (GO:0097512) | 0.002498 | 0.02197 | 555.17 | 3326.77 |
| 2 | alveolar lamellar body (GO:0097208) | 0.002997 | 0.02197 | 444.11 | 2580.41 |
| 3 | MHC class II protein complex (GO:0042613) | 0.006482 | 0.03169 | 184.98 | 932.06 |
| 4 | lamellar body (GO:0042599) | 0.006482 | 0.03169 | 184.98 | 932.06 |
| 5 | MHC protein complex (GO:0042611) | 0.009957 | 0.04381 | 116.79 | 538.34 |

Oxygen desaturation on exertion ([HP:0030874](https://hpo.jax.org/app/browse/term/HP:0030874))

KEGG Pathway

| **Index** | **Name** | **P-value** | **Adjusted p-value** | **Odds Ratio** | **Combined score** |
| --- | --- | --- | --- | --- | --- |
| 1 | Thyroid hormone synthesis | 0.0006123 | 0.01960 | 68.21 | 504.62 |
| 2 | Maturity onset diabetes of the young | 0.01293 | 0.09247 | 88.73 | 385.85 |
| 3 | Thyroid cancer | 0.01835 | 0.09247 | 61.59 | 246.23 |
| 4 | Autoimmune thyroid disease | 0.02619 | 0.09247 | 42.60 | 155.17 |
| 5 | Inflammatory bowel disease | 0.03204 | 0.09247 | 34.59 | 119.03 |

GO: Biological Process

| **Index** | **Name** | **P-value** | **Adjusted p-value** | **Odds Ratio** | **Combined score** |
| --- | --- | --- | --- | --- | --- |
| 1 | thyroid gland development (GO:0030878) | 5.033e-9 | 9.161e-7 | 1713.00 | 32730.63 |
| 2 | positive regulation of cell-cell adhesion mediated by cadherin (GO:2000049) | 0.000008085 | 0.0001471 | 713.68 | 8368.26 |
| 3 | endocrine system development (GO:0035270) | 1.812e-7 | 0.00001099 | 407.53 | 6326.45 |
| 4 | negative regulation of epithelial to mesenchymal transition (GO:0010719) | 3.266e-7 | 0.00001486 | 329.08 | 4914.57 |
| 5 | metanephric distal tubule development (GO:0072235) | 0.002498 | 0.01247 | 555.17 | 3326.77 |

GO: Molecular Function

| **Index** | **Name** | **P-value** | **Adjusted p-value** | **Odds Ratio** | **Combined score** |
| --- | --- | --- | --- | --- | --- |
| 1 | intronic transcription regulatory region sequence-specific DNA binding (GO:0001161) | 0.002498 | 0.009991 | 555.17 | 3326.77 |
| 2 | DEAD/H-box RNA helicase binding (GO:0017151) | 0.003495 | 0.01141 | 370.07 | 2093.27 |
| 3 | aryl hydrocarbon receptor binding (GO:0017162) | 0.003994 | 0.01141 | 317.19 | 1751.86 |
| 4 | transcription regulatory region nucleic acid binding (GO:0001067) | 3.081e-8 | 0.000001232 | 95.57 | 1652.93 |
| 5 | glucocorticoid receptor binding (GO:0035259) | 0.004492 | 0.01198 | 277.53 | 1500.17 |

GO: Cellular Component

| **Index** | **Name** | **P-value** | **Adjusted p-value** | **Odds Ratio** | **Combined score** |
| --- | --- | --- | --- | --- | --- |
| 1 | alveolar lamellar body (GO:0097208) | 0.000003371 | 0.00004719 | 1249.13 | 15739.23 |
| 2 | multivesicular body lumen (GO:0097486) | 0.000004719 | 0.00004719 | 999.25 | 12254.80 |
| 3 | late endosome lumen (GO:0031906) | 0.000008085 | 0.00005390 | 713.68 | 8368.26 |
| 4 | lamellar body (GO:0042599) | 0.00001750 | 0.00008749 | 454.07 | 4973.59 |
| 5 | multivesicular body (GO:0005771) | 0.0002403 | 0.0009613 | 110.81 | 923.40 |

Pleuritis ([HP:0002102](https://hpo.jax.org/app/browse/term/HP:0002102))

KEGG Pathway

| **Index** | **Name** | **P-value** | **Adjusted p-value** | **Odds Ratio** | **Combined score** |
| --- | --- | --- | --- | --- | --- |
| 1 | Autoimmune thyroid disease | 0.0001435 | 0.006456 | 156.41 | 1384.12 |
| 2 | African trypanosomiasis | 0.01288 | 0.08302 | 92.39 | 402.10 |
| 3 | Measles | 0.0009843 | 0.02215 | 57.97 | 401.38 |
| 4 | Allograft rejection | 0.01323 | 0.08302 | 89.89 | 388.83 |
| 5 | Graft-versus-host disease | 0.01461 | 0.08302 | 81.11 | 342.76 |

GO: Biological Process

| **Index** | **Name** | **P-value** | **Adjusted p-value** | **Odds Ratio** | **Combined score** |
| --- | --- | --- | --- | --- | --- |
| 1 | regulation of natural killer cell proliferation (GO:0032817) | 0.000002203 | 0.0002108 | 1599.04 | 20828.53 |
| 2 | negative regulation of B cell proliferation (GO:0030889) | 0.000006918 | 0.0004964 | 799.32 | 9496.99 |
| 3 | negative regulation of interleukin-10 production (GO:0032693) | 0.00001424 | 0.0008176 | 532.75 | 5945.01 |
| 4 | positive regulation of natural killer cell proliferation (GO:0032819) | 0.001749 | 0.01195 | 832.88 | 5287.72 |
| 5 | positive regulation of cell fate commitment (GO:0010455) | 0.001749 | 0.01195 | 832.88 | 5287.72 |

GO: Molecular Function

| **Index** | **Name** | **P-value** | **Adjusted p-value** | **Odds Ratio** | **Combined score** |
| --- | --- | --- | --- | --- | --- |
| 1 | IgG binding (GO:0019864) | 0.001749 | 0.01444 | 832.88 | 5287.72 |
| 2 | interleukin-12 receptor binding (GO:0005143) | 0.002098 | 0.01444 | 666.27 | 4108.59 |
| 3 | non-membrane spanning protein tyrosine phosphatase activity (GO:0004726) | 0.002098 | 0.01444 | 666.27 | 4108.59 |
| 4 | MutLalpha complex binding (GO:0032405) | 0.002448 | 0.01444 | 555.19 | 3338.15 |
| 5 | MutSalpha complex binding (GO:0032407) | 0.002448 | 0.01444 | 555.19 | 3338.15 |

GO: Cellular Component

| **Index** | **Name** | **P-value** | **Adjusted p-value** | **Odds Ratio** | **Combined score** |
| --- | --- | --- | --- | --- | --- |
| 1 | CD95 death-inducing signaling complex (GO:0031265) | 0.001749 | 0.01538 | 832.88 | 5287.72 |
| 2 | death-inducing signaling complex (GO:0031264) | 0.002797 | 0.01538 | 475.86 | 2797.66 |
| 3 | cytoplasmic side of plasma membrane (GO:0009898) | 0.01909 | 0.05251 | 61.54 | 243.60 |
| 4 | clathrin-coated vesicle (GO:0030136) | 0.02767 | 0.05386 | 42.01 | 150.72 |
| 5 | clathrin-coated endocytic vesicle (GO:0045334) | 0.02938 | 0.05386 | 39.50 | 139.34 |

Pulmonary embolism ([HP:0002204](https://hpo.jax.org/app/browse/term/HP:0002204))

KEGG Pathway

| **Index** | **Name** | **P-value** | **Adjusted p-value** | **Odds Ratio** | **Combined score** |
| --- | --- | --- | --- | --- | --- |
| 1 | GnRH secretion | 0.0004462 | 0.01795 | 80.35 | 619.92 |
| 2 | Vitamin digestion and absorption | 0.01194 | 0.07705 | 96.46 | 427.12 |
| 3 | Circadian entrainment | 0.001021 | 0.01795 | 52.36 | 360.55 |
| 4 | Aldosterone synthesis and secretion | 0.001042 | 0.01795 | 51.81 | 355.72 |
| 5 | Serotonergic synapse | 0.001382 | 0.01795 | 44.77 | 294.78 |

GO: Biological Process

| **Index** | **Name** | **P-value** | **Adjusted p-value** | **Odds Ratio** | **Combined score** |
| --- | --- | --- | --- | --- | --- |
| 1 | positive regulation of cell proliferation involved in heart morphogenesis (GO:2000138) | 0.002498 | 0.02379 | 555.17 | 3326.77 |
| 2 | atrial cardiac muscle cell membrane repolarization (GO:0099624) | 0.002498 | 0.02379 | 555.17 | 3326.77 |
| 3 | membrane repolarization during atrial cardiac muscle cell action potential (GO:0098914) | 0.002498 | 0.02379 | 555.17 | 3326.77 |
| 4 | transsulfuration (GO:0019346) | 0.002498 | 0.02379 | 555.17 | 3326.77 |
| 5 | positive regulation of histone H3-K9 acetylation (GO:2000617) | 0.002498 | 0.02379 | 555.17 | 3326.77 |

GO: Molecular Function

| **Index** | **Name** | **P-value** | **Adjusted p-value** | **Odds Ratio** | **Combined score** |
| --- | --- | --- | --- | --- | --- |
| 1 | oxidoreductase activity, acting on metal ions, NAD or NADP as acceptor (GO:0016723) | 0.002498 | 0.03991 | 555.17 | 3326.77 |
| 2 | U2 snRNA binding (GO:0030620) | 0.002498 | 0.03991 | 555.17 | 3326.77 |
| 3 | G protein-coupled serotonin receptor binding (GO:0031821) | 0.002997 | 0.03991 | 444.11 | 2580.41 |
| 4 | U1 snRNA binding (GO:0030619) | 0.003994 | 0.03991 | 317.19 | 1751.86 |
| 5 | voltage-gated potassium channel activity involved in ventricular cardiac muscle cell action potential repolarization (GO:1902282) | 0.005985 | 0.03991 | 201.81 | 1032.95 |

GO: Cellular Component

| **Index** | **Name** | **P-value** | **Adjusted p-value** | **Odds Ratio** | **Combined score** |
| --- | --- | --- | --- | --- | --- |
| 1 | heterotrimeric G-protein complex (GO:0005834) | 0.01638 | 0.1572 | 69.30 | 284.93 |
| 2 | voltage-gated potassium channel complex (GO:0008076) | 0.03591 | 0.1572 | 30.74 | 102.25 |
| 3 | potassium channel complex (GO:0034705) | 0.03930 | 0.1572 | 28.00 | 90.64 |
| 4 | lytic vacuole membrane (GO:0098852) | 0.1258 | 0.3344 | 8.24 | 17.08 |
| 5 | lysosomal membrane (GO:0005765) | 0.1533 | 0.3344 | 6.64 | 12.45 |

Reduced forced expiratory volume in one second ([HP:0032342](https://hpo.jax.org/app/browse/term/HP:0032342))

KEGG Pathway

| **Index** | **Name** | **P-value** | **Adjusted p-value** | **Odds Ratio** | **Combined score** |
| --- | --- | --- | --- | --- | --- |
| 1 | Homologous recombination | 0.0001826 | 0.0009526 | 127.89 | 1100.92 |
| 2 | Fanconi anemia pathway | 0.0003175 | 0.0009526 | 95.86 | 772.11 |
| 3 | Mismatch repair | 0.01144 | 0.02289 | 100.85 | 450.83 |
| 4 | DNA replication | 0.01786 | 0.02679 | 63.35 | 255.00 |
| 5 | Pancreatic cancer | 0.03736 | 0.04484 | 29.50 | 96.98 |

GO: Biological Process

| **Index** | **Name** | **P-value** | **Adjusted p-value** | **Odds Ratio** | **Combined score** |
| --- | --- | --- | --- | --- | --- |
| 1 | formation of extrachromosomal circular DNA (GO:0001325) | 4.360e-15 | 2.151e-13 | 3330.67 | 110132.57 |
| 2 | t-circle formation (GO:0090656) | 4.360e-15 | 2.151e-13 | 3330.67 | 110132.57 |
| 3 | telomere maintenance via telomere trimming (GO:0090737) | 4.360e-15 | 2.151e-13 | 3330.67 | 110132.57 |
| 4 | telomeric loop disassembly (GO:0090657) | 6.605e-12 | 1.086e-10 | 2220.44 | 57161.21 |
| 5 | G-quadruplex DNA unwinding (GO:0044806) | 3.147e-9 | 2.911e-8 | 2141.36 | 41921.23 |

GO: Molecular Function

| **Index** | **Name** | **P-value** | **Adjusted p-value** | **Odds Ratio** | **Combined score** |
| --- | --- | --- | --- | --- | --- |
| 1 | D-loop DNA binding (GO:0062037) | 0.000002248 | 0.00002473 | 1665.58 | 21661.60 |
| 2 | Y-form DNA binding (GO:0000403) | 0.000002248 | 0.00002473 | 1665.58 | 21661.60 |
| 3 | four-way junction helicase activity (GO:0009378) | 0.000003371 | 0.00002649 | 1249.13 | 15739.23 |
| 4 | 5'-3' DNA helicase activity (GO:0043139) | 0.000004719 | 0.00003244 | 999.25 | 12254.80 |
| 5 | 5'-flap endonuclease activity (GO:0017108) | 0.000006290 | 0.00003459 | 832.67 | 9972.50 |

GO: Cellular Component

| **Index** | **Name** | **P-value** | **Adjusted p-value** | **Odds Ratio** | **Combined score** |
| --- | --- | --- | --- | --- | --- |
| 1 | shelterin complex (GO:0070187) | 0.002997 | 0.005565 | 444.11 | 2580.41 |
| 2 | transcription factor TFIIH holo complex (GO:0005675) | 0.005985 | 0.008645 | 201.81 | 1032.95 |
| 3 | condensed nuclear chromosome (GO:0000794) | 0.009957 | 0.01294 | 116.79 | 538.34 |
| 4 | chromosome (GO:0005694) | 0.00005786 | 0.0007522 | 54.14 | 528.26 |
| 5 | nuclear chromosome (GO:0000228) | 0.0007493 | 0.001948 | 61.45 | 442.20 |

Reduced forced vital capacity ([HP:0032341](https://hpo.jax.org/app/browse/term/HP:0032341))

KEGG Pathway

| **Index** | **Name** | **P-value** | **Adjusted p-value** | **Odds Ratio** | **Combined score** |
| --- | --- | --- | --- | --- | --- |
| 1 | Terpenoid backbone biosynthesis | 0.01095 | 0.02098 | 105.66 | 476.99 |
| 2 | Mannose type O-glycan biosynthesis | 0.01144 | 0.02098 | 100.85 | 450.83 |
| 3 | Cardiac muscle contraction | 0.0008229 | 0.003669 | 58.54 | 415.82 |
| 4 | Hypertrophic cardiomyopathy | 0.0008802 | 0.003669 | 56.54 | 397.78 |
| 5 | Dilated cardiomyopathy | 0.001001 | 0.003669 | 52.91 | 365.49 |

GO: Biological Process

| **Index** | **Name** | **P-value** | **Adjusted p-value** | **Odds Ratio** | **Combined score** |
| --- | --- | --- | --- | --- | --- |
| 1 | actin-myosin filament sliding (GO:0033275) | 7.523e-7 | 0.00004251 | 244.35 | 3445.31 |
| 2 | muscle filament sliding (GO:0030049) | 7.523e-7 | 0.00004251 | 244.35 | 3445.31 |
| 3 | centriole assembly (GO:0098534) | 0.002498 | 0.03382 | 555.17 | 3326.77 |
| 4 | skeletal muscle thin filament assembly (GO:0030240) | 0.002997 | 0.03382 | 444.11 | 2580.41 |
| 5 | regulation of t-circle formation (GO:1904429) | 0.002997 | 0.03382 | 444.11 | 2580.41 |

GO: Molecular Function

| **Index** | **Name** | **P-value** | **Adjusted p-value** | **Odds Ratio** | **Combined score** |
| --- | --- | --- | --- | --- | --- |
| 1 | inositol 1,3,4,5 tetrakisphosphate binding (GO:0043533) | 0.002498 | 0.02905 | 555.17 | 3326.77 |
| 2 | prenyltransferase activity (GO:0004659) | 0.004492 | 0.02905 | 277.53 | 1500.17 |
| 3 | DNA polymerase binding (GO:0070182) | 0.008966 | 0.02905 | 130.54 | 615.43 |
| 4 | ADP binding (GO:0043531) | 0.01342 | 0.03088 | 85.32 | 367.79 |
| 5 | dicarboxylic acid transmembrane transporter activity (GO:0005310) | 0.01441 | 0.03088 | 79.21 | 335.86 |

GO: Cellular Component

| **Index** | **Name** | **P-value** | **Adjusted p-value** | **Odds Ratio** | **Combined score** |
| --- | --- | --- | --- | --- | --- |
| 1 | myosin filament (GO:0032982) | 0.002997 | 0.04064 | 444.11 | 2580.41 |
| 2 | muscle myosin complex (GO:0005859) | 0.007476 | 0.04064 | 158.54 | 776.21 |
| 3 | dense core granule (GO:0031045) | 0.007973 | 0.04064 | 147.96 | 714.91 |
| 4 | supramolecular fiber (GO:0099512) | 0.009462 | 0.04064 | 123.28 | 574.57 |
| 5 | actin filament (GO:0005884) | 0.0005645 | 0.01693 | 71.14 | 532.12 |

Reduced vital capacity ([HP:0002792](https://hpo.jax.org/app/browse/term/HP:0002792))

KEGG Pathway

| **Index** | **Name** | **P-value** | **Adjusted p-value** | **Odds Ratio** | **Combined score** |
| --- | --- | --- | --- | --- | --- |
| 1 | Neuroactive ligand-receptor interaction | 0.00001608 | 0.0001286 | 38.88 | 429.14 |
| 2 | Viral myocarditis | 0.02960 | 0.07518 | 37.53 | 132.12 |
| 3 | Cardiac muscle contraction | 0.04267 | 0.07518 | 25.72 | 81.12 |
| 4 | Hypertrophic cardiomyopathy | 0.04411 | 0.07518 | 24.85 | 77.54 |
| 5 | Dilated cardiomyopathy | 0.04699 | 0.07518 | 23.27 | 71.15 |

GO: Biological Process

| **Index** | **Name** | **P-value** | **Adjusted p-value** | **Odds Ratio** | **Combined score** |
| --- | --- | --- | --- | --- | --- |
| 1 | synaptic transmission, cholinergic (GO:0007271) | 8.685e-8 | 0.000001795 | 535.02 | 8698.92 |
| 2 | neuromuscular junction development (GO:0007528) | 1.812e-7 | 0.000002381 | 407.53 | 6326.45 |
| 3 | muscle contraction (GO:0006936) | 1.316e-11 | 8.161e-10 | 242.28 | 6070.00 |
| 4 | neuromuscular process (GO:0050905) | 0.00003830 | 0.0002638 | 293.72 | 2987.15 |
| 5 | postsynaptic membrane organization (GO:0001941) | 0.00003830 | 0.0002638 | 293.72 | 2987.15 |

GO: Molecular Function

| **Index** | **Name** | **P-value** | **Adjusted p-value** | **Odds Ratio** | **Combined score** |
| --- | --- | --- | --- | --- | --- |
| 1 | neurotransmitter receptor activity involved in regulation of postsynaptic membrane potential (GO:0099529) | 1.878e-10 | 5.070e-9 | 783.25 | 17541.62 |
| 2 | transmitter-gated ion channel activity involved in regulation of postsynaptic membrane potential (GO:1904315) | 8.580e-10 | 1.158e-8 | 511.90 | 10686.61 |
| 3 | transmitter-gated ion channel activity (GO:0022824) | 1.451e-9 | 1.305e-8 | 443.56 | 9026.96 |
| 4 | postsynaptic neurotransmitter receptor activity (GO:0098960) | 1.191e-7 | 7.447e-7 | 475.52 | 7581.26 |
| 5 | acetylcholine receptor activity (GO:0015464) | 1.379e-7 | 7.447e-7 | 450.47 | 7115.97 |

GO: Cellular Component

| **Index** | **Name** | **P-value** | **Adjusted p-value** | **Odds Ratio** | **Combined score** |
| --- | --- | --- | --- | --- | --- |
| 1 | neuromuscular junction (GO:0031594) | 4.712e-12 | 8.481e-11 | 604.76 | 15772.67 |
| 2 | acetylcholine-gated channel complex (GO:0005892) | 3.267e-8 | 2.940e-7 | 778.40 | 13417.28 |
| 3 | ion channel complex (GO:0034702) | 5.841e-7 | 0.000003505 | 267.29 | 3836.51 |
| 4 | myosin filament (GO:0032982) | 0.002997 | 0.008990 | 444.11 | 2580.41 |
| 5 | muscle myosin complex (GO:0005859) | 0.007476 | 0.01892 | 158.54 | 776.21 |

Restrictive ventilatory defect ([HP:0002091](https://hpo.jax.org/app/browse/term/HP:0002091))

KEGG Pathway

| **Index** | **Name** | **P-value** | **Adjusted p-value** | **Odds Ratio** | **Combined score** |
| --- | --- | --- | --- | --- | --- |
| 1 | Neuroactive ligand-receptor interaction | 0.00001608 | 0.0001608 | 38.88 | 429.14 |
| 2 | Primary immunodeficiency | 0.01884 | 0.07259 | 59.92 | 237.98 |
| 3 | Autoimmune thyroid disease | 0.02619 | 0.07259 | 42.60 | 155.17 |
| 4 | B cell receptor signaling pathway | 0.03978 | 0.07259 | 27.65 | 89.16 |
| 5 | Rheumatoid arthritis | 0.04555 | 0.07259 | 24.03 | 74.23 |

GO: Biological Process

| **Index** | **Name** | **P-value** | **Adjusted p-value** | **Odds Ratio** | **Combined score** |
| --- | --- | --- | --- | --- | --- |
| 1 | synaptic transmission, cholinergic (GO:0007271) | 8.685e-8 | 0.000003648 | 535.02 | 8698.92 |
| 2 | neuromuscular junction development (GO:0007528) | 1.812e-7 | 0.000004033 | 407.53 | 6326.45 |
| 3 | negative regulation of regulatory T cell differentiation (GO:0045590) | 0.002498 | 0.01614 | 555.17 | 3326.77 |
| 4 | neuromuscular process (GO:0050905) | 0.00003830 | 0.0003971 | 293.72 | 2987.15 |
| 5 | postsynaptic membrane organization (GO:0001941) | 0.00003830 | 0.0003971 | 293.72 | 2987.15 |

GO: Molecular Function

| **Index** | **Name** | **P-value** | **Adjusted p-value** | **Odds Ratio** | **Combined score** |
| --- | --- | --- | --- | --- | --- |
| 1 | neurotransmitter receptor activity involved in regulation of postsynaptic membrane potential (GO:0099529) | 1.878e-10 | 4.131e-9 | 783.25 | 17541.62 |
| 2 | transmitter-gated ion channel activity involved in regulation of postsynaptic membrane potential (GO:1904315) | 8.580e-10 | 9.438e-9 | 511.90 | 10686.61 |
| 3 | transmitter-gated ion channel activity (GO:0022824) | 1.451e-9 | 1.064e-8 | 443.56 | 9026.96 |
| 4 | postsynaptic neurotransmitter receptor activity (GO:0098960) | 1.191e-7 | 6.068e-7 | 475.52 | 7581.26 |
| 5 | acetylcholine receptor activity (GO:0015464) | 1.379e-7 | 6.068e-7 | 450.47 | 7115.97 |

GO: Cellular Component

| **Index** | **Name** | **P-value** | **Adjusted p-value** | **Odds Ratio** | **Combined score** |
| --- | --- | --- | --- | --- | --- |
| 1 | neuromuscular junction (GO:0031594) | 4.712e-12 | 5.183e-11 | 604.76 | 15772.67 |
| 2 | acetylcholine-gated channel complex (GO:0005892) | 3.267e-8 | 1.198e-7 | 778.40 | 13417.28 |
| 3 | ion channel complex (GO:0034702) | 5.841e-7 | 0.000001606 | 267.29 | 3836.51 |
| 4 | integral component of plasma membrane (GO:0005887) | 3.020e-8 | 1.198e-7 | 51.30 | 888.24 |
| 5 | voltage-gated sodium channel complex (GO:0001518) | 0.008469 | 0.01553 | 138.71 | 661.82 |

Centrilobular ground-glass opacification on pulmonary HRCT ([HP:0025180](https://hpo.jax.org/app/browse/term/HP:0025180))

KEGG Pathway

| **Index** | **Name** | **P-value** | **Adjusted p-value** | **Odds Ratio** | **Combined score** |
| --- | --- | --- | --- | --- | --- |
| 1 | TGF-beta signaling pathway | 7.834e-18 | 8.618e-17 | 925.77 | 36464.13 |
| 2 | Cytokine-cytokine receptor interaction | 2.887e-16 | 1.588e-15 | 620.06 | 22186.23 |
| 3 | Hippo signaling pathway | 5.460e-11 | 2.002e-10 | 189.49 | 4477.76 |
| 4 | Fluid shear stress and atherosclerosis | 0.00003804 | 0.00009108 | 62.57 | 636.72 |
| 5 | Basal cell carcinoma | 0.0004323 | 0.0007926 | 81.68 | 632.69 |

GO: Biological Process

| **Index** | **Name** | **P-value** | **Adjusted p-value** | **Odds Ratio** | **Combined score** |
| --- | --- | --- | --- | --- | --- |
| 1 | regulation of pathway-restricted SMAD protein phosphorylation (GO:0060393) | 1.850e-26 | 8.807e-24 | 199420.00 | 11816020.17 |
| 2 | BMP signaling pathway (GO:0030509) | 6.348e-26 | 1.511e-23 | 199350.00 | 11566105.39 |
| 3 | cellular response to BMP stimulus (GO:0071773) | 1.637e-25 | 2.598e-23 | 199290.00 | 11373779.66 |
| 4 | transmembrane receptor protein serine/threonine kinase signaling pathway (GO:0007178) | 1.197e-22 | 1.140e-20 | 198670.00 | 10028275.58 |
| 5 | positive regulation of pathway-restricted SMAD protein phosphorylation (GO:0010862) | 9.646e-24 | 1.148e-21 | 4725.47 | 250428.98 |

GO: Molecular Function

| **Index** | **Name** | **P-value** | **Adjusted p-value** | **Odds Ratio** | **Combined score** |
| --- | --- | --- | --- | --- | --- |
| 1 | BMP receptor binding (GO:0070700) | 2.247e-11 | 2.772e-10 | 1480.07 | 36289.42 |
| 2 | transmembrane receptor protein serine/threonine kinase binding (GO:0070696) | 5.717e-11 | 5.288e-10 | 1109.89 | 26176.78 |
| 3 | cytokine activity (GO:0005125) | 3.764e-13 | 1.393e-11 | 278.65 | 7971.64 |
| 4 | transforming growth factor beta-activated receptor activity (GO:0005024) | 0.00001481 | 0.00009133 | 499.50 | 5554.53 |
| 5 | BMP binding (GO:0036122) | 0.00001481 | 0.00009133 | 499.50 | 5554.53 |

GO: Cellular Component

| **Index** | **Name** | **P-value** | **Adjusted p-value** | **Odds Ratio** | **Combined score** |
| --- | --- | --- | --- | --- | --- |
| 1 | activin receptor complex (GO:0048179) | 0.003495 | 0.04214 | 370.07 | 2093.27 |
| 2 | spanning component of plasma membrane (GO:0044214) | 0.006482 | 0.04214 | 184.98 | 932.06 |
| 3 | spanning component of membrane (GO:0089717) | 0.009957 | 0.04315 | 116.79 | 538.34 |
| 4 | serine/threonine protein kinase complex (GO:1902554) | 0.01835 | 0.05964 | 61.59 | 246.23 |
| 5 | caveola (GO:0005901) | 0.02960 | 0.06771 | 37.53 | 132.12 |

Interlobular septal thickening ([HP:0030879](https://hpo.jax.org/app/browse/term/HP:0030879))

KEGG Pathway

| **Index** | **Name** | **P-value** | **Adjusted p-value** | **Odds Ratio** | **Combined score** |
| --- | --- | --- | --- | --- | --- |
| 1 | TGF-beta signaling pathway | 7.834e-18 | 8.618e-17 | 925.77 | 36464.13 |
| 2 | Cytokine-cytokine receptor interaction | 2.887e-16 | 1.588e-15 | 620.06 | 22186.23 |
| 3 | Hippo signaling pathway | 5.460e-11 | 2.002e-10 | 189.49 | 4477.76 |
| 4 | Fluid shear stress and atherosclerosis | 0.00003804 | 0.00009108 | 62.57 | 636.72 |
| 5 | Basal cell carcinoma | 0.0004323 | 0.0007926 | 81.68 | 632.69 |

GO: Biological Process

| **Index** | **Name** | **P-value** | **Adjusted p-value** | **Odds Ratio** | **Combined score** |
| --- | --- | --- | --- | --- | --- |
| 1 | regulation of pathway-restricted SMAD protein phosphorylation (GO:0060393) | 1.850e-26 | 8.807e-24 | 199420.00 | 11816020.17 |
| 2 | BMP signaling pathway (GO:0030509) | 6.348e-26 | 1.511e-23 | 199350.00 | 11566105.39 |
| 3 | cellular response to BMP stimulus (GO:0071773) | 1.637e-25 | 2.598e-23 | 199290.00 | 11373779.66 |
| 4 | transmembrane receptor protein serine/threonine kinase signaling pathway (GO:0007178) | 1.197e-22 | 1.140e-20 | 198670.00 | 10028275.58 |
| 5 | positive regulation of pathway-restricted SMAD protein phosphorylation (GO:0010862) | 9.646e-24 | 1.148e-21 | 4725.47 | 250428.98 |

GO: Molecular Function

| **Index** | **Name** | **P-value** | **Adjusted p-value** | **Odds Ratio** | **Combined score** |
| --- | --- | --- | --- | --- | --- |
| 1 | BMP receptor binding (GO:0070700) | 2.247e-11 | 2.772e-10 | 1480.07 | 36289.42 |
| 2 | transmembrane receptor protein serine/threonine kinase binding (GO:0070696) | 5.717e-11 | 5.288e-10 | 1109.89 | 26176.78 |
| 3 | cytokine activity (GO:0005125) | 3.764e-13 | 1.393e-11 | 278.65 | 7971.64 |
| 4 | transforming growth factor beta-activated receptor activity (GO:0005024) | 0.00001481 | 0.00009133 | 499.50 | 5554.53 |
| 5 | BMP binding (GO:0036122) | 0.00001481 | 0.00009133 | 499.50 | 5554.53 |

GO: Cellular Component

| **Index** | **Name** | **P-value** | **Adjusted p-value** | **Odds Ratio** | **Combined score** |
| --- | --- | --- | --- | --- | --- |
| 1 | activin receptor complex (GO:0048179) | 0.003495 | 0.04214 | 370.07 | 2093.27 |
| 2 | spanning component of plasma membrane (GO:0044214) | 0.006482 | 0.04214 | 184.98 | 932.06 |
| 3 | spanning component of membrane (GO:0089717) | 0.009957 | 0.04315 | 116.79 | 538.34 |
| 4 | serine/threonine protein kinase complex (GO:1902554) | 0.01835 | 0.05964 | 61.59 | 246.23 |
| 5 | caveola (GO:0005901) | 0.02960 | 0.06771 | 37.53 | 132.12 |

Parenchymal consolidation ([HP:0032177](https://hpo.jax.org/app/browse/term/HP:0032177))

KEGG Pathway

| **Index** | **Name** | **P-value** | **Adjusted p-value** | **Odds Ratio** | **Combined score** |
| --- | --- | --- | --- | --- | --- |
| 1 | Thyroid hormone synthesis | 0.0006123 | 0.01960 | 68.21 | 504.62 |
| 2 | Maturity onset diabetes of the young | 0.01293 | 0.09247 | 88.73 | 385.85 |
| 3 | Thyroid cancer | 0.01835 | 0.09247 | 61.59 | 246.23 |
| 4 | Autoimmune thyroid disease | 0.02619 | 0.09247 | 42.60 | 155.17 |
| 5 | Inflammatory bowel disease | 0.03204 | 0.09247 | 34.59 | 119.03 |

GO: Biological Process

| **Index** | **Name** | **P-value** | **Adjusted p-value** | **Odds Ratio** | **Combined score** |
| --- | --- | --- | --- | --- | --- |
| 1 | thyroid gland development (GO:0030878) | 5.033e-9 | 9.161e-7 | 1713.00 | 32730.63 |
| 2 | positive regulation of cell-cell adhesion mediated by cadherin (GO:2000049) | 0.000008085 | 0.0001471 | 713.68 | 8368.26 |
| 3 | endocrine system development (GO:0035270) | 1.812e-7 | 0.00001099 | 407.53 | 6326.45 |
| 4 | negative regulation of epithelial to mesenchymal transition (GO:0010719) | 3.266e-7 | 0.00001486 | 329.08 | 4914.57 |
| 5 | metanephric distal tubule development (GO:0072235) | 0.002498 | 0.01247 | 555.17 | 3326.77 |

GO: Molecular Function

| **Index** | **Name** | **P-value** | **Adjusted p-value** | **Odds Ratio** | **Combined score** |
| --- | --- | --- | --- | --- | --- |
| 1 | intronic transcription regulatory region sequence-specific DNA binding (GO:0001161) | 0.002498 | 0.009991 | 555.17 | 3326.77 |
| 2 | DEAD/H-box RNA helicase binding (GO:0017151) | 0.003495 | 0.01141 | 370.07 | 2093.27 |
| 3 | aryl hydrocarbon receptor binding (GO:0017162) | 0.003994 | 0.01141 | 317.19 | 1751.86 |
| 4 | transcription regulatory region nucleic acid binding (GO:0001067) | 3.081e-8 | 0.000001232 | 95.57 | 1652.93 |
| 5 | glucocorticoid receptor binding (GO:0035259) | 0.004492 | 0.01198 | 277.53 | 1500.17 |

GO: Cellular Component

| **Index** | **Name** | **P-value** | **Adjusted p-value** | **Odds Ratio** | **Combined score** |
| --- | --- | --- | --- | --- | --- |
| 1 | alveolar lamellar body (GO:0097208) | 0.000003371 | 0.00004719 | 1249.13 | 15739.23 |
| 2 | multivesicular body lumen (GO:0097486) | 0.000004719 | 0.00004719 | 999.25 | 12254.80 |
| 3 | late endosome lumen (GO:0031906) | 0.000008085 | 0.00005390 | 713.68 | 8368.26 |
| 4 | lamellar body (GO:0042599) | 0.00001750 | 0.00008749 | 454.07 | 4973.59 |
| 5 | multivesicular body (GO:0005771) | 0.0002403 | 0.0009613 | 110.81 | 923.40 |

Pleural thickening ([HP:0031944](https://hpo.jax.org/app/browse/term/HP:0031944))

KEGG Pathway

| **Index** | **Name** | **P-value** | **Adjusted p-value** | **Odds Ratio** | **Combined score** |
| --- | --- | --- | --- | --- | --- |
| 1 | NF-kappa B signaling pathway | 1.941e-20 | 2.212e-18 | 1884.79 | 85548.22 |
| 2 | B cell receptor signaling pathway | 2.384e-10 | 1.359e-8 | 262.03 | 5805.76 |
| 3 | T cell receptor signaling pathway | 8.515e-10 | 3.236e-8 | 200.92 | 4196.00 |
| 4 | Shigellosis | 6.478e-8 | 0.000001846 | 81.95 | 1356.40 |
| 5 | PD-L1 expression and PD-1 checkpoint pathway in cancer | 0.000009993 | 0.0002278 | 99.19 | 1142.03 |

GO: Biological Process

| **Index** | **Name** | **P-value** | **Adjusted p-value** | **Odds Ratio** | **Combined score** |
| --- | --- | --- | --- | --- | --- |
| 1 | activation of NF-kappaB-inducing kinase activity (GO:0007250) | 5.717e-11 | 1.927e-9 | 1109.89 | 26176.78 |
| 2 | positive regulation of NF-kappaB transcription factor activity (GO:0051092) | 4.814e-16 | 1.136e-13 | 539.95 | 19043.76 |
| 3 | positive regulation of antigen receptor-mediated signaling pathway (GO:0050857) | 1.878e-10 | 4.924e-9 | 783.25 | 17541.62 |
| 4 | positive regulation of T cell receptor signaling pathway (GO:0050862) | 3.267e-8 | 5.139e-7 | 778.40 | 13417.28 |
| 5 | positive regulation of DNA-binding transcription factor activity (GO:0051091) | 2.059e-14 | 2.430e-12 | 331.97 | 10461.57 |

GO: Molecular Function

| **Index** | **Name** | **P-value** | **Adjusted p-value** | **Odds Ratio** | **Combined score** |
| --- | --- | --- | --- | --- | --- |
| 1 | CARD domain binding (GO:0050700) | 5.717e-11 | 2.287e-9 | 1109.89 | 26176.78 |
| 2 | protein kinase B binding (GO:0043422) | 0.000006290 | 0.0001258 | 832.67 | 9972.50 |
| 3 | protein kinase C activity (GO:0004697) | 0.00002041 | 0.0002721 | 416.21 | 4494.85 |
| 4 | histone threonine kinase activity (GO:0035184) | 0.003495 | 0.01748 | 370.07 | 2093.27 |
| 5 | kinase activator activity (GO:0019209) | 0.0001038 | 0.001038 | 172.08 | 1578.44 |

GO: Cellular Component

| **Index** | **Name** | **P-value** | **Adjusted p-value** | **Odds Ratio** | **Combined score** |
| --- | --- | --- | --- | --- | --- |
| 1 | aggresome (GO:0016235) | 0.0001327 | 0.002123 | 151.19 | 1349.73 |
| 2 | CD40 receptor complex (GO:0035631) | 0.005488 | 0.04390 | 222.00 | 1155.57 |
| 3 | cytoplasmic side of plasma membrane (GO:0009898) | 0.02717 | 0.1449 | 41.02 | 147.91 |
| 4 | lipid droplet (GO:0005811) | 0.03785 | 0.1514 | 29.11 | 95.32 |
| 5 | mitotic spindle (GO:0072686) | 0.05842 | 0.1558 | 18.55 | 52.69 |

Pulmonary fibrosis ([HP:0002206](https://hpo.jax.org/app/browse/term/HP:0002206))

KEGG Pathway

| **Index** | **Name** | **P-value** | **Adjusted p-value** | **Odds Ratio** | **Combined score** |
| --- | --- | --- | --- | --- | --- |
| 1 | Autoimmune thyroid disease | 0.000002081 | 0.0001103 | 170.91 | 2236.02 |
| 2 | Allograft rejection | 0.0001567 | 0.002663 | 138.57 | 1214.06 |
| 3 | Graft-versus-host disease | 0.0001917 | 0.002663 | 124.69 | 1067.29 |
| 4 | Type I diabetes mellitus | 0.0002010 | 0.002663 | 121.64 | 1035.45 |
| 5 | Rheumatoid arthritis | 0.0009395 | 0.009959 | 54.67 | 381.04 |

GO: Biological Process

| **Index** | **Name** | **P-value** | **Adjusted p-value** | **Odds Ratio** | **Combined score** |
| --- | --- | --- | --- | --- | --- |
| 1 | establishment of protein localization to telomere (GO:0070200) | 0.002498 | 0.02503 | 555.17 | 3326.77 |
| 2 | positive regulation of CD4-positive, CD25-positive, alpha-beta regulatory T cell differentiation (GO:0032831) | 0.002498 | 0.02503 | 555.17 | 3326.77 |
| 3 | lung morphogenesis (GO:0060425) | 0.002498 | 0.02503 | 555.17 | 3326.77 |
| 4 | positive regulation of toll-like receptor 9 signaling pathway (GO:0034165) | 0.002498 | 0.02503 | 555.17 | 3326.77 |
| 5 | negative regulation of regulatory T cell differentiation (GO:0045590) | 0.002498 | 0.02503 | 555.17 | 3326.77 |

GO: Molecular Function

| **Index** | **Name** | **P-value** | **Adjusted p-value** | **Odds Ratio** | **Combined score** |
| --- | --- | --- | --- | --- | --- |
| 1 | intronic transcription regulatory region sequence-specific DNA binding (GO:0001161) | 0.002498 | 0.02566 | 555.17 | 3326.77 |
| 2 | RNA-directed DNA polymerase activity (GO:0003964) | 0.002997 | 0.02566 | 444.11 | 2580.41 |
| 3 | non-membrane spanning protein tyrosine phosphatase activity (GO:0004726) | 0.002997 | 0.02566 | 444.11 | 2580.41 |
| 4 | telomerase activity (GO:0003720) | 0.002997 | 0.02566 | 444.11 | 2580.41 |
| 5 | CD4 receptor binding (GO:0042609) | 0.003994 | 0.02566 | 317.19 | 1751.86 |

GO: Cellular Component

| **Index** | **Name** | **P-value** | **Adjusted p-value** | **Odds Ratio** | **Combined score** |
| --- | --- | --- | --- | --- | --- |
| 1 | transferase complex, transferring phosphorus-containing groups (GO:0061695) | 0.004492 | 0.05839 | 277.53 | 1500.17 |
| 2 | MHC class II protein complex (GO:0042613) | 0.006482 | 0.06320 | 184.98 | 932.06 |
| 3 | MHC protein complex (GO:0042611) | 0.009957 | 0.07753 | 116.79 | 538.34 |
| 4 | clathrin-coated endocytic vesicle (GO:0045334) | 0.0007857 | 0.03064 | 59.96 | 428.66 |
| 5 | lumenal side of endoplasmic reticulum membrane (GO:0098553) | 0.01392 | 0.07753 | 82.15 | 351.18 |

Pulmonary interstitial high-resolution computed tomography abnormality ([HP:0025389](https://hpo.jax.org/app/browse/term/HP:0025389))

KEGG Pathway

| **Index** | **Name** | **P-value** | **Adjusted p-value** | **Odds Ratio** | **Combined score** |
| --- | --- | --- | --- | --- | --- |
| 1 | Asthma | 0.01540 | 0.09994 | 73.93 | 308.54 |
| 2 | Allograft rejection | 0.01884 | 0.09994 | 59.92 | 237.98 |
| 3 | Graft-versus-host disease | 0.02081 | 0.09994 | 54.06 | 209.35 |
| 4 | Type I diabetes mellitus | 0.02130 | 0.09994 | 52.77 | 203.13 |
| 5 | ABC transporters | 0.02228 | 0.09994 | 50.37 | 191.61 |

GO: Biological Process

| **Index** | **Name** | **P-value** | **Adjusted p-value** | **Odds Ratio** | **Combined score** |
| --- | --- | --- | --- | --- | --- |
| 1 | positive regulation of chromosome organization (GO:2001252) | 0.00002354 | 0.003983 | 384.17 | 4094.01 |
| 2 | establishment of protein localization to telomere (GO:0070200) | 0.002498 | 0.02251 | 555.17 | 3326.77 |
| 3 | positive regulation of cardioblast differentiation (GO:0051891) | 0.002498 | 0.02251 | 555.17 | 3326.77 |
| 4 | positive regulation of CD4-positive, CD25-positive, alpha-beta regulatory T cell differentiation (GO:0032831) | 0.002498 | 0.02251 | 555.17 | 3326.77 |
| 5 | regulation of cardioblast differentiation (GO:0051890) | 0.002498 | 0.02251 | 555.17 | 3326.77 |

GO: Molecular Function

| **Index** | **Name** | **P-value** | **Adjusted p-value** | **Odds Ratio** | **Combined score** |
| --- | --- | --- | --- | --- | --- |
| 1 | intronic transcription regulatory region sequence-specific DNA binding (GO:0001161) | 0.002498 | 0.01825 | 555.17 | 3326.77 |
| 2 | phosphatidylcholine flippase activity (GO:0140345) | 0.002498 | 0.01825 | 555.17 | 3326.77 |
| 3 | phosphatidylcholine transfer activity (GO:0120019) | 0.002498 | 0.01825 | 555.17 | 3326.77 |
| 4 | cell adhesive protein binding involved in bundle of His cell-Purkinje myocyte communication (GO:0086083) | 0.002498 | 0.01825 | 555.17 | 3326.77 |
| 5 | RNA-directed DNA polymerase activity (GO:0003964) | 0.002997 | 0.01825 | 444.11 | 2580.41 |

GO: Cellular Component

| **Index** | **Name** | **P-value** | **Adjusted p-value** | **Odds Ratio** | **Combined score** |
| --- | --- | --- | --- | --- | --- |
| 1 | alveolar lamellar body (GO:0097208) | 0.002997 | 0.04423 | 444.11 | 2580.41 |
| 2 | transferase complex, transferring phosphorus-containing groups (GO:0061695) | 0.004492 | 0.04423 | 277.53 | 1500.17 |
| 3 | MHC class II protein complex (GO:0042613) | 0.006482 | 0.04423 | 184.98 | 932.06 |
| 4 | spanning component of plasma membrane (GO:0044214) | 0.006482 | 0.04423 | 184.98 | 932.06 |
| 5 | lamellar body (GO:0042599) | 0.006482 | 0.04423 | 184.98 | 932.06 |

Pulmonary opacity ([HP:0031457](https://hpo.jax.org/app/browse/term/HP:0031457))

KEGG Pathway

| **Index** | **Name** | **P-value** | **Adjusted p-value** | **Odds Ratio** | **Combined score** |
| --- | --- | --- | --- | --- | --- |
| 1 | ABC transporters | 0.02007 | 0.1097 | 56.67 | 221.48 |
| 2 | Arrhythmogenic right ventricular cardiomyopathy | 0.03413 | 0.1097 | 32.75 | 110.63 |
| 3 | RNA degradation | 0.03500 | 0.1097 | 31.91 | 106.98 |
| 4 | TGF-beta signaling pathway | 0.04152 | 0.1097 | 26.74 | 85.09 |
| 5 | AGE-RAGE signaling pathway in diabetic complications | 0.04412 | 0.1097 | 25.12 | 78.38 |

GO: Biological Process

| **Index** | **Name** | **P-value** | **Adjusted p-value** | **Odds Ratio** | **Combined score** |
| --- | --- | --- | --- | --- | --- |
| 1 | positive regulation of chromosome organization (GO:2001252) | 0.00001884 | 0.002620 | 439.08 | 4776.89 |
| 2 | establishment of protein localization to telomere (GO:0070200) | 0.002248 | 0.02051 | 624.59 | 3808.55 |
| 3 | positive regulation of cardioblast differentiation (GO:0051891) | 0.002248 | 0.02051 | 624.59 | 3808.55 |
| 4 | lung morphogenesis (GO:0060425) | 0.002248 | 0.02051 | 624.59 | 3808.55 |
| 5 | tricuspid valve development (GO:0003175) | 0.002248 | 0.02051 | 624.59 | 3808.55 |

GO: Molecular Function

| **Index** | **Name** | **P-value** | **Adjusted p-value** | **Odds Ratio** | **Combined score** |
| --- | --- | --- | --- | --- | --- |
| 1 | intronic transcription regulatory region sequence-specific DNA binding (GO:0001161) | 0.002248 | 0.01564 | 624.59 | 3808.55 |
| 2 | phosphatidylcholine flippase activity (GO:0140345) | 0.002248 | 0.01564 | 624.59 | 3808.55 |
| 3 | phosphatidylcholine transfer activity (GO:0120019) | 0.002248 | 0.01564 | 624.59 | 3808.55 |
| 4 | cell adhesive protein binding involved in bundle of His cell-Purkinje myocyte communication (GO:0086083) | 0.002248 | 0.01564 | 624.59 | 3808.55 |
| 5 | RNA-directed DNA polymerase activity (GO:0003964) | 0.002697 | 0.01564 | 499.65 | 2955.69 |

GO: Cellular Component

| **Index** | **Name** | **P-value** | **Adjusted p-value** | **Odds Ratio** | **Combined score** |
| --- | --- | --- | --- | --- | --- |
| 1 | alveolar lamellar body (GO:0097208) | 0.002697 | 0.04423 | 499.65 | 2955.69 |
| 2 | transferase complex, transferring phosphorus-containing groups (GO:0061695) | 0.004043 | 0.04423 | 312.23 | 1720.62 |
| 3 | spanning component of plasma membrane (GO:0044214) | 0.005836 | 0.04423 | 208.11 | 1070.48 |
| 4 | lamellar body (GO:0042599) | 0.005836 | 0.04423 | 208.11 | 1070.48 |
| 5 | spanning component of membrane (GO:0089717) | 0.008966 | 0.05230 | 131.39 | 619.44 |

Reticular pattern on pulmonary HRCT ([HP:0025390](https://hpo.jax.org/app/browse/term/HP:0025390))

KEGG Pathway

| **Index** | **Name** | **P-value** | **Adjusted p-value** | **Odds Ratio** | **Combined score** |
| --- | --- | --- | --- | --- | --- |
| 1 | ABC transporters | 0.01343 | 0.06259 | 90.68 | 390.89 |
| 2 | Arrhythmogenic right ventricular cardiomyopathy | 0.02288 | 0.06259 | 52.42 | 198.00 |
| 3 | RNA degradation | 0.02347 | 0.06259 | 51.07 | 191.60 |
| 4 | Gastric cancer | 0.04388 | 0.07897 | 26.82 | 83.84 |
| 5 | Hepatocellular carcinoma | 0.04936 | 0.07897 | 23.74 | 71.44 |

GO: Biological Process

| **Index** | **Name** | **P-value** | **Adjusted p-value** | **Odds Ratio** | **Combined score** |
| --- | --- | --- | --- | --- | --- |
| 1 | positive regulation of chromosome organization (GO:2001252) | 0.000007861 | 0.001019 | 768.50 | 9032.62 |
| 2 | regulation of telomere maintenance via telomere lengthening (GO:1904356) | 0.00001145 | 0.001019 | 624.31 | 7103.11 |
| 3 | establishment of protein localization to telomere (GO:0070200) | 0.001499 | 0.01519 | 999.50 | 6499.56 |
| 4 | positive regulation of cardioblast differentiation (GO:0051891) | 0.001499 | 0.01519 | 999.50 | 6499.56 |
| 5 | regulation of cardioblast differentiation (GO:0051890) | 0.001499 | 0.01519 | 999.50 | 6499.56 |

GO: Molecular Function

| **Index** | **Name** | **P-value** | **Adjusted p-value** | **Odds Ratio** | **Combined score** |
| --- | --- | --- | --- | --- | --- |
| 1 | phosphatidylcholine flippase activity (GO:0140345) | 0.001499 | 0.01079 | 999.50 | 6499.56 |
| 2 | phosphatidylcholine transfer activity (GO:0120019) | 0.001499 | 0.01079 | 999.50 | 6499.56 |
| 3 | cell adhesive protein binding involved in bundle of His cell-Purkinje myocyte communication (GO:0086083) | 0.001499 | 0.01079 | 999.50 | 6499.56 |
| 4 | telomerase RNA binding (GO:0070034) | 0.00001728 | 0.0008294 | 499.35 | 5475.89 |
| 5 | RNA-directed DNA polymerase activity (GO:0003964) | 0.001799 | 0.01079 | 799.56 | 5053.71 |

GO: Cellular Component

| **Index** | **Name** | **P-value** | **Adjusted p-value** | **Odds Ratio** | **Combined score** |
| --- | --- | --- | --- | --- | --- |
| 1 | alveolar lamellar body (GO:0097208) | 0.001799 | 0.02254 | 799.56 | 5053.71 |
| 2 | transferase complex, transferring phosphorus-containing groups (GO:0061695) | 0.002697 | 0.02254 | 499.65 | 2955.69 |
| 3 | lamellar body (GO:0042599) | 0.003894 | 0.02434 | 333.03 | 1847.77 |
| 4 | intercalated disc (GO:0014704) | 0.009265 | 0.03860 | 133.09 | 623.08 |
| 5 | cornified envelope (GO:0001533) | 0.01283 | 0.04141 | 95.01 | 413.84 |

Dyspnea ([HP:0002094](https://hpo.jax.org/app/browse/term/HP:0002094))

KEGG Pathway

| **Index** | **Name** | **P-value** | **Adjusted p-value** | **Odds Ratio** | **Combined score** |
| --- | --- | --- | --- | --- | --- |
| 1 | Neuroactive ligand-receptor interaction | 0.00001608 | 0.0001286 | 38.88 | 429.14 |
| 2 | Viral myocarditis | 0.02960 | 0.07518 | 37.53 | 132.12 |
| 3 | Cardiac muscle contraction | 0.04267 | 0.07518 | 25.72 | 81.12 |
| 4 | Hypertrophic cardiomyopathy | 0.04411 | 0.07518 | 24.85 | 77.54 |
| 5 | Dilated cardiomyopathy | 0.04699 | 0.07518 | 23.27 | 71.15 |

GO: Biological Process

| **Index** | **Name** | **P-value** | **Adjusted p-value** | **Odds Ratio** | **Combined score** |
| --- | --- | --- | --- | --- | --- |
| 1 | synaptic transmission, cholinergic (GO:0007271) | 1.217e-10 | 9.125e-9 | 887.78 | 20267.77 |
| 2 | neuromuscular junction development (GO:0007528) | 1.812e-7 | 0.000002880 | 407.53 | 6326.45 |
| 3 | atrial cardiac muscle tissue morphogenesis (GO:0055009) | 0.002498 | 0.01338 | 555.17 | 3326.77 |
| 4 | muscle contraction (GO:0006936) | 2.535e-9 | 9.507e-8 | 160.21 | 3171.03 |
| 5 | neuromuscular process (GO:0050905) | 0.00003830 | 0.0003545 | 293.72 | 2987.15 |

GO: Molecular Function

| **Index** | **Name** | **P-value** | **Adjusted p-value** | **Odds Ratio** | **Combined score** |
| --- | --- | --- | --- | --- | --- |
| 1 | neurotransmitter receptor activity involved in regulation of postsynaptic membrane potential (GO:0099529) | 1.878e-10 | 4.131e-9 | 783.25 | 17541.62 |
| 2 | transmitter-gated ion channel activity involved in regulation of postsynaptic membrane potential (GO:1904315) | 8.580e-10 | 9.438e-9 | 511.90 | 10686.61 |
| 3 | transmitter-gated ion channel activity (GO:0022824) | 1.451e-9 | 1.064e-8 | 443.56 | 9026.96 |
| 4 | postsynaptic neurotransmitter receptor activity (GO:0098960) | 1.191e-7 | 6.068e-7 | 475.52 | 7581.26 |
| 5 | acetylcholine receptor activity (GO:0015464) | 1.379e-7 | 6.068e-7 | 450.47 | 7115.97 |

GO: Cellular Component

| **Index** | **Name** | **P-value** | **Adjusted p-value** | **Odds Ratio** | **Combined score** |
| --- | --- | --- | --- | --- | --- |
| 1 | neuromuscular junction (GO:0031594) | 4.712e-12 | 5.654e-11 | 604.76 | 15772.67 |
| 2 | acetylcholine-gated channel complex (GO:0005892) | 3.267e-8 | 1.960e-7 | 778.40 | 13417.28 |
| 3 | ion channel complex (GO:0034702) | 5.841e-7 | 0.000002337 | 267.29 | 3836.51 |
| 4 | muscle myosin complex (GO:0005859) | 0.007476 | 0.01452 | 158.54 | 776.21 |
| 5 | voltage-gated sodium channel complex (GO:0001518) | 0.008469 | 0.01452 | 138.71 | 661.82 |

Exertional dyspnea ([HP:0002875](https://hpo.jax.org/app/browse/term/HP:0002875))

KEGG Pathway

| **Index** | **Name** | **P-value** | **Adjusted p-value** | **Odds Ratio** | **Combined score** |
| --- | --- | --- | --- | --- | --- |
| 1 | Neuroactive ligand-receptor interaction | 0.00001608 | 0.00001608 | 38.88 | 429.14 |

GO: Biological Process

| **Index** | **Name** | **P-value** | **Adjusted p-value** | **Odds Ratio** | **Combined score** |
| --- | --- | --- | --- | --- | --- |
| 1 | synaptic transmission, cholinergic (GO:0007271) | 1.217e-10 | 1.594e-8 | 887.78 | 20267.77 |
| 2 | neuromuscular junction development (GO:0007528) | 1.812e-7 | 0.000006289 | 407.53 | 6326.45 |
| 3 | regulation of cardioblast differentiation (GO:0051890) | 0.002498 | 0.02180 | 555.17 | 3326.77 |
| 4 | positive regulation of cardioblast differentiation (GO:0051891) | 0.002498 | 0.02180 | 555.17 | 3326.77 |
| 5 | ventricular cardiac muscle cell differentiation (GO:0055012) | 0.002498 | 0.02180 | 555.17 | 3326.77 |

GO: Molecular Function

| **Index** | **Name** | **P-value** | **Adjusted p-value** | **Odds Ratio** | **Combined score** |
| --- | --- | --- | --- | --- | --- |
| 1 | neurotransmitter receptor activity involved in regulation of postsynaptic membrane potential (GO:0099529) | 1.878e-10 | 6.009e-9 | 783.25 | 17541.62 |
| 2 | transmitter-gated ion channel activity involved in regulation of postsynaptic membrane potential (GO:1904315) | 8.580e-10 | 1.373e-8 | 511.90 | 10686.61 |
| 3 | transmitter-gated ion channel activity (GO:0022824) | 1.451e-9 | 1.547e-8 | 443.56 | 9026.96 |
| 4 | postsynaptic neurotransmitter receptor activity (GO:0098960) | 1.191e-7 | 8.827e-7 | 475.52 | 7581.26 |
| 5 | acetylcholine receptor activity (GO:0015464) | 1.379e-7 | 8.827e-7 | 450.47 | 7115.97 |

GO: Cellular Component

| **Index** | **Name** | **P-value** | **Adjusted p-value** | **Odds Ratio** | **Combined score** |
| --- | --- | --- | --- | --- | --- |
| 1 | neuromuscular junction (GO:0031594) | 4.712e-12 | 5.654e-11 | 604.76 | 15772.67 |
| 2 | acetylcholine-gated channel complex (GO:0005892) | 3.267e-8 | 1.960e-7 | 778.40 | 13417.28 |
| 3 | ion channel complex (GO:0034702) | 5.841e-7 | 0.000002337 | 267.29 | 3836.51 |
| 4 | voltage-gated sodium channel complex (GO:0001518) | 0.008469 | 0.01694 | 138.71 | 661.82 |
| 5 | neuron projection (GO:0043005) | 0.000003661 | 0.00001098 | 35.28 | 441.62 |

Hemoptysis ([HP:0002105](https://hpo.jax.org/app/browse/term/HP:0002105))

KEGG Pathway

| **Index** | **Name** | **P-value** | **Adjusted p-value** | **Odds Ratio** | **Combined score** |
| --- | --- | --- | --- | --- | --- |
| 1 | Protein digestion and absorption | 0.00001551 | 0.0004031 | 85.24 | 944.01 |
| 2 | Inflammatory bowel disease | 0.0004602 | 0.003988 | 79.08 | 607.60 |
| 3 | IL-17 signaling pathway | 0.0009597 | 0.006199 | 54.07 | 375.73 |
| 4 | Th17 cell differentiation | 0.001241 | 0.006199 | 47.35 | 316.84 |
| 5 | Platelet activation | 0.001661 | 0.006199 | 40.71 | 260.58 |

GO: Biological Process

| **Index** | **Name** | **P-value** | **Adjusted p-value** | **Odds Ratio** | **Combined score** |
| --- | --- | --- | --- | --- | --- |
| 1 | cellular response to interleukin-17 (GO:0097398) | 0.000008085 | 0.0003319 | 713.68 | 8368.26 |
| 2 | interleukin-17-mediated signaling pathway (GO:0097400) | 0.000008085 | 0.0003319 | 713.68 | 8368.26 |
| 3 | regulation of granulocyte macrophage colony-stimulating factor production (GO:0032645) | 0.00002690 | 0.0007666 | 356.71 | 3753.85 |
| 4 | positive regulation of natural killer cell proliferation (GO:0032819) | 0.002498 | 0.02134 | 555.17 | 3326.77 |
| 5 | positive regulation of cell fate commitment (GO:0010455) | 0.002498 | 0.02134 | 555.17 | 3326.77 |

GO: Molecular Function

| **Index** | **Name** | **P-value** | **Adjusted p-value** | **Odds Ratio** | **Combined score** |
| --- | --- | --- | --- | --- | --- |
| 1 | cyclic nucleotide-dependent protein kinase activity (GO:0004690) | 0.002997 | 0.01398 | 444.11 | 2580.41 |
| 2 | interleukin-12 receptor binding (GO:0005143) | 0.002997 | 0.01398 | 444.11 | 2580.41 |
| 3 | interleukin-17 receptor activity (GO:0030368) | 0.003994 | 0.01398 | 317.19 | 1751.86 |
| 4 | platelet-derived growth factor binding (GO:0048407) | 0.005488 | 0.01537 | 222.00 | 1155.57 |
| 5 | cytokine receptor activity (GO:0004896) | 0.0008418 | 0.01179 | 57.86 | 409.65 |

GO: Cellular Component

| **Index** | **Name** | **P-value** | **Adjusted p-value** | **Odds Ratio** | **Combined score** |
| --- | --- | --- | --- | --- | --- |
| 1 | elastic fiber (GO:0071953) | 0.002498 | 0.009991 | 555.17 | 3326.77 |
| 2 | supramolecular fiber (GO:0099512) | 0.00003830 | 0.0002298 | 293.72 | 2987.15 |
| 3 | microfibril (GO:0001527) | 0.005488 | 0.01646 | 222.00 | 1155.57 |
| 4 | collagen-containing extracellular matrix (GO:0062023) | 0.00002461 | 0.0002298 | 34.78 | 369.07 |
| 5 | filopodium (GO:0030175) | 0.02863 | 0.05005 | 38.86 | 138.07 |

Rhinorrhea ([HP:0031417](https://hpo.jax.org/app/browse/term/HP:0031417))

KEGG Pathway

| **Index** | **Name** | **P-value** | **Adjusted p-value** | **Odds Ratio** | **Combined score** |
| --- | --- | --- | --- | --- | --- |
| 1 | Taste transduction | 0.01709 | 0.01709 | 78.08 | 317.74 |

GO: Biological Process

| **Index** | **Name** | **P-value** | **Adjusted p-value** | **Odds Ratio** | **Combined score** |
| --- | --- | --- | --- | --- | --- |
| 1 | determination of pancreatic left/right asymmetry (GO:0035469) | 2.999e-7 | 0.000006140 | 6664.33 | 100096.56 |
| 2 | determination of liver left/right asymmetry (GO:0071910) | 4.499e-7 | 0.000006140 | 4998.00 | 73042.46 |
| 3 | determination of digestive tract left/right asymmetry (GO:0071907) | 6.298e-7 | 0.000006140 | 3998.20 | 57086.01 |
| 4 | regulation of cilium beat frequency (GO:0003356) | 6.298e-7 | 0.000006140 | 3998.20 | 57086.01 |
| 5 | epithelial cilium movement involved in determination of left/right asymmetry (GO:0060287) | 0.000001079 | 0.000008420 | 2855.57 | 39232.86 |

GO: Molecular Function

| **Index** | **Name** | **P-value** | **Adjusted p-value** | **Odds Ratio** | **Combined score** |
| --- | --- | --- | --- | --- | --- |
| 1 | voltage-gated sodium channel activity (GO:0005248) | 0.004393 | 0.007579 | 317.06 | 1720.94 |
| 2 | sodium channel activity (GO:0005272) | 0.007579 | 0.007579 | 179.81 | 877.91 |

GO: Cellular Component

| **Index** | **Name** | **P-value** | **Adjusted p-value** | **Odds Ratio** | **Combined score** |
| --- | --- | --- | --- | --- | --- |
| 1 | voltage-gated sodium channel complex (GO:0001518) | 0.003396 | 0.009982 | 416.25 | 2366.46 |
| 2 | sodium channel complex (GO:0034706) | 0.004991 | 0.009982 | 277.39 | 1470.20 |
| 3 | cilium (GO:0005929) | 0.0008259 | 0.004955 | 84.09 | 596.95 |
| 4 | axon (GO:0030424) | 0.04018 | 0.06027 | 32.50 | 104.47 |
| 5 | neuron projection (GO:0043005) | 0.1067 | 0.1280 | 11.68 | 26.13 |

Tachypnea ([HP:0002789](https://hpo.jax.org/app/browse/term/HP:0002789))

GO: Biological Process

| **Index** | **Name** | **P-value** | **Adjusted p-value** | **Odds Ratio** | **Combined score** |
| --- | --- | --- | --- | --- | --- |
| 1 | cilium assembly (GO:0060271) | 7.884e-19 | 2.129e-17 | 196860.00 | 8205971.98 |
| 2 | ciliary basal body-plasma membrane docking (GO:0097711) | 8.554e-18 | 1.155e-16 | 915.08 | 35962.78 |
| 3 | cilium organization (GO:0044782) | 2.747e-17 | 2.472e-16 | 812.51 | 30983.64 |
| 4 | plasma membrane bounded cell projection assembly (GO:0120031) | 1.681e-16 | 1.135e-15 | 659.81 | 23965.68 |
| 5 | smoothened signaling pathway (GO:0007224) | 3.049e-12 | 1.372e-11 | 665.33 | 17642.09 |

GO: Cellular Component

| **Index** | **Name** | **P-value** | **Adjusted p-value** | **Odds Ratio** | **Combined score** |
| --- | --- | --- | --- | --- | --- |
| 1 | cilium (GO:0005929) | 3.495e-12 | 2.447e-11 | 200.46 | 5288.15 |
| 2 | ciliary membrane (GO:0060170) | 4.016e-7 | 0.000001406 | 305.54 | 4499.94 |
| 3 | cell projection membrane (GO:0031253) | 0.00001104 | 0.00002576 | 95.83 | 1093.80 |
| 4 | bounding membrane of organelle (GO:0098588) | 0.005508 | 0.009639 | 10.78 | 56.10 |
| 5 | cell-cell junction (GO:0005911) | 0.1276 | 0.1786 | 8.12 | 16.71 |
