## Supplementary Table 1-3 for "Long COVID: G Protein-Coupled Receptors (GPCRs) responsible for persistent post-COVID symptoms": Supplementary File 2.10.1-Neuropsychiatric.docx

Neuropsychiatric-behavioral

Anxiety ([HP:0000739](https://hpo.jax.org/app/browse/term/HP:0000739))

GO: Biological Process

| **Index** | **Name** | **P-value** | **Adjusted p-value** | **Odds Ratio** | **Combined score** |
| --- | --- | --- | --- | --- | --- |
| 1 | equilibrioception (GO:0050957) | 5.668e-17 | 3.287e-15 | 19989.00 | 747771.16 |
| 2 | neuromuscular process controlling balance (GO:0050885) | 1.888e-14 | 2.738e-13 | 2220.11 | 70156.55 |
| 3 | sensory perception (GO:0007600) | 4.223e-14 | 4.899e-13 | 664.83 | 20473.93 |
| 4 | sensory perception of mechanical stimulus (GO:0050954) | 2.969e-15 | 7.316e-14 | 573.51 | 19184.17 |
| 5 | sensory perception of sound (GO:0007605) | 3.784e-15 | 7.316e-14 | 552.94 | 18362.12 |

GO: Molecular Function

| **Index** | **Name** | **P-value** | **Adjusted p-value** | **Odds Ratio** | **Combined score** |
| --- | --- | --- | --- | --- | --- |
| 1 | myosin binding (GO:0017022) | 0.02766 | 0.05531 | 40.27 | 144.50 |
| 2 | calcium ion binding (GO:0005509) | 0.01239 | 0.05531 | 14.19 | 62.33 |
| 3 | magnesium ion binding (GO:0000287) | 0.07066 | 0.1060 | 15.21 | 40.30 |
| 4 | metal ion binding (GO:0046872) | 0.02616 | 0.05531 | 9.45 | 34.45 |
| 5 | cadherin binding (GO:0045296) | 0.1499 | 0.1798 | 6.81 | 12.92 |

GO: Cellular Component

| **Index** | **Name** | **P-value** | **Adjusted p-value** | **Odds Ratio** | **Combined score** |
| --- | --- | --- | --- | --- | --- |
| 1 | microvillus (GO:0005902) | 0.0003539 | 0.003893 | 90.61 | 720.07 |
| 2 | actin-based cell projection (GO:0098858) | 0.0007493 | 0.004121 | 61.45 | 442.20 |
| 3 | catenin complex (GO:0016342) | 0.01540 | 0.05645 | 73.93 | 308.54 |
| 4 | basement membrane (GO:0005604) | 0.02570 | 0.07068 | 43.44 | 159.04 |
| 5 | cilium (GO:0005929) | 0.1124 | 0.2473 | 9.30 | 20.33 |

Apathy ([HP:0000741](https://hpo.jax.org/app/browse/term/HP:0000741))

KEGG Pathway

| **Index** | **Name** | **P-value** | **Adjusted p-value** | **Odds Ratio** | **Combined score** |
| --- | --- | --- | --- | --- | --- |
| 1 | Hedgehog signaling pathway | 0.000002460 | 0.00005659 | 161.22 | 2082.13 |
| 2 | Basal cell carcinoma | 0.0004323 | 0.004972 | 81.68 | 632.69 |
| 3 | Gastric cancer | 0.002386 | 0.01372 | 33.75 | 203.77 |
| 4 | Melanoma | 0.03543 | 0.1358 | 31.17 | 104.12 |
| 5 | Pathways in cancer | 0.001943 | 0.01372 | 15.80 | 98.63 |

GO: Biological Process

| **Index** | **Name** | **P-value** | **Adjusted p-value** | **Odds Ratio** | **Combined score** |
| --- | --- | --- | --- | --- | --- |
| 1 | regulation of nodal signaling pathway involved in determination of lateral mesoderm left/right asymmetry (GO:1900175) | 0.000002248 | 0.0002281 | 1665.58 | 21661.60 |
| 2 | positive regulation of T cell differentiation in thymus (GO:0033089) | 0.000003371 | 0.0002281 | 1249.13 | 15739.23 |
| 3 | dorsal/ventral pattern formation (GO:0009953) | 3.331e-10 | 1.053e-7 | 665.67 | 14526.48 |
| 4 | pituitary gland development (GO:0021983) | 0.000004719 | 0.0002281 | 999.25 | 12254.80 |
| 5 | proximal/distal pattern formation (GO:0009954) | 0.000006290 | 0.0002484 | 832.67 | 9972.50 |

GO: Molecular Function

| **Index** | **Name** | **P-value** | **Adjusted p-value** | **Odds Ratio** | **Combined score** |
| --- | --- | --- | --- | --- | --- |
| 1 | morphogen activity (GO:0016015) | 0.000004719 | 0.0001982 | 999.25 | 12254.80 |
| 2 | type 1 fibroblast growth factor receptor binding (GO:0005105) | 0.002498 | 0.02097 | 555.17 | 3326.77 |
| 3 | type 2 fibroblast growth factor receptor binding (GO:0005111) | 0.002498 | 0.02097 | 555.17 | 3326.77 |
| 4 | patched binding (GO:0005113) | 0.003495 | 0.02097 | 370.07 | 2093.27 |
| 5 | minor groove of adenine-thymine-rich DNA binding (GO:0003680) | 0.003495 | 0.02097 | 370.07 | 2093.27 |

GO: Cellular Component

| **Index** | **Name** | **P-value** | **Adjusted p-value** | **Odds Ratio** | **Combined score** |
| --- | --- | --- | --- | --- | --- |
| 1 | neurofibrillary tangle (GO:0097418) | 0.002498 | 0.02704 | 555.17 | 3326.77 |
| 2 | glial cell projection (GO:0097386) | 0.006979 | 0.03613 | 170.74 | 847.70 |
| 3 | membrane raft (GO:0045121) | 0.002846 | 0.02704 | 30.79 | 180.49 |
| 4 | dendrite (GO:0030425) | 0.007607 | 0.03613 | 18.40 | 89.75 |
| 5 | collagen-containing extracellular matrix (GO:0062023) | 0.01465 | 0.05566 | 12.97 | 54.78 |

Attention deficit hyperactivity disorder ([HP:0007018](https://hpo.jax.org/app/browse/term/HP:0007018))

KEGG Pathway

| **Index** | **Name** | **P-value** | **Adjusted p-value** | **Odds Ratio** | **Combined score** |
| --- | --- | --- | --- | --- | --- |
| 1 | Nicotine addiction | 2.854e-9 | 1.342e-7 | 369.52 | 7270.07 |
| 2 | GABAergic synapse | 7.537e-8 | 0.000001771 | 156.12 | 2560.47 |
| 3 | Morphine addiction | 0.00001068 | 0.0001674 | 96.93 | 1109.48 |
| 4 | Dopaminergic synapse | 0.00003260 | 0.0003830 | 65.98 | 681.69 |
| 5 | Retrograde endocannabinoid signaling | 0.00004587 | 0.0004312 | 58.66 | 585.95 |

GO: Biological Process

| **Index** | **Name** | **P-value** | **Adjusted p-value** | **Odds Ratio** | **Combined score** |
| --- | --- | --- | --- | --- | --- |
| 1 | neuronal action potential (GO:0019228) | 1.379e-7 | 0.000003241 | 450.47 | 7115.97 |
| 2 | synaptic transmission, GABAergic (GO:0051932) | 0.00001235 | 0.00007736 | 555.03 | 6273.06 |
| 3 | inhibitory synapse assembly (GO:1904862) | 0.00001235 | 0.00007736 | 555.03 | 6273.06 |
| 4 | membrane depolarization (GO:0051899) | 4.430e-7 | 0.000006941 | 294.99 | 4315.59 |
| 5 | regulation of postsynaptic membrane potential (GO:0060078) | 5.841e-7 | 0.000007844 | 267.29 | 3836.51 |

GO: Molecular Function

| **Index** | **Name** | **P-value** | **Adjusted p-value** | **Odds Ratio** | **Combined score** |
| --- | --- | --- | --- | --- | --- |
| 1 | benzodiazepine receptor activity (GO:0008503) | 0.00001010 | 0.0001683 | 624.44 | 7182.69 |
| 2 | extracellular ligand-gated ion channel activity (GO:0005230) | 0.00001235 | 0.0001683 | 555.03 | 6273.06 |
| 3 | GABA-gated chloride ion channel activity (GO:0022851) | 0.00001750 | 0.0001769 | 454.07 | 4973.59 |
| 4 | inhibitory extracellular ligand-gated ion channel activity (GO:0005237) | 0.00002041 | 0.0001769 | 416.21 | 4494.85 |
| 5 | glutamate-gated calcium ion channel activity (GO:0022849) | 0.002498 | 0.006494 | 555.17 | 3326.77 |

GO: Cellular Component

| **Index** | **Name** | **P-value** | **Adjusted p-value** | **Odds Ratio** | **Combined score** |
| --- | --- | --- | --- | --- | --- |
| 1 | voltage-gated sodium channel complex (GO:0001518) | 0.00003048 | 0.0001905 | 332.92 | 3461.84 |
| 2 | GABA-A receptor complex (GO:1902711) | 0.00003830 | 0.0001915 | 293.72 | 2987.15 |
| 3 | sodium channel complex (GO:0034706) | 0.00006709 | 0.0002795 | 217.03 | 2085.58 |
| 4 | neuron to neuron synapse (GO:0098984) | 0.003994 | 0.009984 | 317.19 | 1751.86 |
| 5 | NMDA selective glutamate receptor complex (GO:0017146) | 0.003994 | 0.009984 | 317.19 | 1751.86 |

Delusions ([HP:0000746](https://hpo.jax.org/app/browse/term/HP:0000746))

KEGG Pathway

| **Index** | **Name** | **P-value** | **Adjusted p-value** | **Odds Ratio** | **Combined score** |
| --- | --- | --- | --- | --- | --- |
| 1 | RNA transport | 0.00009052 | 0.0004526 | 46.39 | 431.86 |
| 2 | Tyrosine metabolism | 0.01786 | 0.02976 | 63.35 | 255.00 |
| 3 | Steroid hormone biosynthesis | 0.03009 | 0.03761 | 36.91 | 129.31 |
| 4 | Herpes simplex virus 1 infection | 0.001616 | 0.004040 | 16.88 | 108.49 |
| 5 | Dopaminergic synapse | 0.06409 | 0.06409 | 16.84 | 46.28 |

GO: Biological Process

| **Index** | **Name** | **P-value** | **Adjusted p-value** | **Odds Ratio** | **Combined score** |
| --- | --- | --- | --- | --- | --- |
| 1 | oligodendrocyte development (GO:0014003) | 5.023e-8 | 0.000005626 | 658.58 | 11068.58 |
| 2 | oligodendrocyte differentiation (GO:0048709) | 1.021e-7 | 0.000005720 | 503.52 | 8105.11 |
| 3 | response to hexose (GO:0009746) | 2.058e-7 | 0.000006513 | 388.99 | 5988.96 |
| 4 | glial cell development (GO:0021782) | 2.326e-7 | 0.000006513 | 372.06 | 5682.76 |
| 5 | response to peptide (GO:1901652) | 8.148e-7 | 0.00001825 | 237.55 | 3330.49 |

GO: Molecular Function

| **Index** | **Name** | **P-value** | **Adjusted p-value** | **Odds Ratio** | **Combined score** |
| --- | --- | --- | --- | --- | --- |
| 1 | translation initiation factor activity (GO:0003743) | 9.499e-7 | 0.00001805 | 225.02 | 3120.36 |
| 2 | O-methyltransferase activity (GO:0008171) | 0.004990 | 0.02370 | 246.68 | 1307.49 |
| 3 | guanyl-nucleotide exchange factor activity (GO:0005085) | 0.00004680 | 0.0004446 | 58.25 | 580.74 |
| 4 | S-adenosylmethionine-dependent methyltransferase activity (GO:0008757) | 0.02130 | 0.07830 | 52.77 | 203.13 |
| 5 | GTPase activator activity (GO:0005096) | 0.0005166 | 0.003272 | 25.30 | 191.47 |

GO: Cellular Component

| **Index** | **Name** | **P-value** | **Adjusted p-value** | **Odds Ratio** | **Combined score** |
| --- | --- | --- | --- | --- | --- |
| 1 | junctional sarcoplasmic reticulum membrane (GO:0014701) | 0.004492 | 0.03593 | 277.53 | 1500.17 |
| 2 | sarcoplasmic reticulum membrane (GO:0033017) | 0.01392 | 0.05566 | 82.15 | 351.18 |
| 3 | axon (GO:0030424) | 0.09746 | 0.2542 | 10.83 | 25.22 |
| 4 | dendrite (GO:0030425) | 0.1271 | 0.2542 | 8.15 | 16.80 |
| 5 | neuron projection (GO:0043005) | 0.2457 | 0.3932 | 3.89 | 5.46 |

Hallucinations ([HP:0000738](https://hpo.jax.org/app/browse/term/HP:0000738))

GO: Biological Process

| **Index** | **Name** | **P-value** | **Adjusted p-value** | **Odds Ratio** | **Combined score** |
| --- | --- | --- | --- | --- | --- |
| 1 | equilibrioception (GO:0050957) | 5.668e-17 | 3.287e-15 | 19989.00 | 747771.16 |
| 2 | neuromuscular process controlling balance (GO:0050885) | 1.888e-14 | 2.738e-13 | 2220.11 | 70156.55 |
| 3 | sensory perception (GO:0007600) | 4.223e-14 | 4.899e-13 | 664.83 | 20473.93 |
| 4 | sensory perception of mechanical stimulus (GO:0050954) | 2.969e-15 | 7.316e-14 | 573.51 | 19184.17 |
| 5 | sensory perception of sound (GO:0007605) | 3.784e-15 | 7.316e-14 | 552.94 | 18362.12 |

GO: Molecular Function

| **Index** | **Name** | **P-value** | **Adjusted p-value** | **Odds Ratio** | **Combined score** |
| --- | --- | --- | --- | --- | --- |
| 1 | myosin binding (GO:0017022) | 0.02766 | 0.05531 | 40.27 | 144.50 |
| 2 | calcium ion binding (GO:0005509) | 0.01239 | 0.05531 | 14.19 | 62.33 |
| 3 | magnesium ion binding (GO:0000287) | 0.07066 | 0.1060 | 15.21 | 40.30 |
| 4 | metal ion binding (GO:0046872) | 0.02616 | 0.05531 | 9.45 | 34.45 |
| 5 | cadherin binding (GO:0045296) | 0.1499 | 0.1798 | 6.81 | 12.92 |

GO: Cellular Component

| **Index** | **Name** | **P-value** | **Adjusted p-value** | **Odds Ratio** | **Combined score** |
| --- | --- | --- | --- | --- | --- |
| 1 | microvillus (GO:0005902) | 0.0003539 | 0.003893 | 90.61 | 720.07 |
| 2 | actin-based cell projection (GO:0098858) | 0.0007493 | 0.004121 | 61.45 | 442.20 |
| 3 | catenin complex (GO:0016342) | 0.01540 | 0.05645 | 73.93 | 308.54 |
| 4 | basement membrane (GO:0005604) | 0.02570 | 0.07068 | 43.44 | 159.04 |
| 5 | cilium (GO:0005929) | 0.1124 | 0.2473 | 9.30 | 20.33 |

Impulsivity ([HP:0100710](https://hpo.jax.org/app/browse/term/HP:0100710))

KEGG Pathway

| **Index** | **Name** | **P-value** | **Adjusted p-value** | **Odds Ratio** | **Combined score** |
| --- | --- | --- | --- | --- | --- |
| 1 | Nicotine addiction | 2.854e-9 | 8.848e-8 | 369.52 | 7270.07 |
| 2 | GABAergic synapse | 0.000009993 | 0.0001104 | 99.19 | 1142.03 |
| 3 | Morphine addiction | 0.00001068 | 0.0001104 | 96.93 | 1109.48 |
| 4 | Dopaminergic synapse | 0.00003260 | 0.0002526 | 65.98 | 681.69 |
| 5 | Retrograde endocannabinoid signaling | 0.00004587 | 0.0002844 | 58.66 | 585.95 |

GO: Biological Process

| **Index** | **Name** | **P-value** | **Adjusted p-value** | **Odds Ratio** | **Combined score** |
| --- | --- | --- | --- | --- | --- |
| 1 | membrane depolarization (GO:0051899) | 1.893e-12 | 1.703e-10 | 739.37 | 19957.87 |
| 2 | neuronal action potential (GO:0019228) | 2.294e-10 | 1.032e-8 | 739.70 | 16417.99 |
| 3 | membrane depolarization during action potential (GO:0086010) | 4.685e-10 | 1.405e-8 | 605.09 | 12998.28 |
| 4 | synaptic transmission, GABAergic (GO:0051932) | 0.00001235 | 0.00008547 | 555.03 | 6273.06 |
| 5 | inhibitory synapse assembly (GO:1904862) | 0.00001235 | 0.00008547 | 555.03 | 6273.06 |

GO: Molecular Function

| **Index** | **Name** | **P-value** | **Adjusted p-value** | **Odds Ratio** | **Combined score** |
| --- | --- | --- | --- | --- | --- |
| 1 | voltage-gated sodium channel activity (GO:0005248) | 2.294e-10 | 8.719e-9 | 739.70 | 16417.99 |
| 2 | sodium channel activity (GO:0005272) | 2.307e-9 | 4.382e-8 | 391.29 | 7781.87 |
| 3 | benzodiazepine receptor activity (GO:0008503) | 0.00001010 | 0.00009840 | 624.44 | 7182.69 |
| 4 | extracellular ligand-gated ion channel activity (GO:0005230) | 0.00001235 | 0.00009840 | 555.03 | 6273.06 |
| 5 | GABA-gated chloride ion channel activity (GO:0022851) | 0.00001750 | 0.0001108 | 454.07 | 4973.59 |

GO: Cellular Component

| **Index** | **Name** | **P-value** | **Adjusted p-value** | **Odds Ratio** | **Combined score** |
| --- | --- | --- | --- | --- | --- |
| 1 | voltage-gated sodium channel complex (GO:0001518) | 7.474e-11 | 1.570e-9 | 1024.46 | 23887.38 |
| 2 | sodium channel complex (GO:0034706) | 3.965e-10 | 3.627e-9 | 633.94 | 13723.68 |
| 3 | GABA-A receptor complex (GO:1902711) | 0.00003830 | 0.0001341 | 293.72 | 2987.15 |
| 4 | integral component of plasma membrane (GO:0005887) | 5.182e-10 | 3.627e-9 | 115.51 | 2469.57 |
| 5 | NMDA selective glutamate receptor complex (GO:0017146) | 0.003994 | 0.008387 | 317.19 | 1751.86 |

Irritability ([HP:0000737](https://hpo.jax.org/app/browse/term/HP:0000737))

KEGG Pathway

| **Index** | **Name** | **P-value** | **Adjusted p-value** | **Odds Ratio** | **Combined score** |
| --- | --- | --- | --- | --- | --- |
| 1 | Hedgehog signaling pathway | 0.0003416 | 0.005620 | 92.30 | 736.71 |
| 2 | Basal cell carcinoma | 0.0004323 | 0.005620 | 81.68 | 632.69 |
| 3 | Folate biosynthesis | 0.01293 | 0.06722 | 88.73 | 385.85 |
| 4 | Tyrosine metabolism | 0.01786 | 0.07739 | 63.35 | 255.00 |
| 5 | Gastric cancer | 0.002386 | 0.01551 | 33.75 | 203.77 |

GO: Biological Process

| **Index** | **Name** | **P-value** | **Adjusted p-value** | **Odds Ratio** | **Combined score** |
| --- | --- | --- | --- | --- | --- |
| 1 | positive regulation of T cell differentiation in thymus (GO:0033089) | 0.000003371 | 0.00007726 | 1249.13 | 15739.23 |
| 2 | metanephric collecting duct development (GO:0072205) | 0.000003371 | 0.00007726 | 1249.13 | 15739.23 |
| 3 | embryonic camera-type eye morphogenesis (GO:0048596) | 0.000003371 | 0.00007726 | 1249.13 | 15739.23 |
| 4 | dorsal/ventral pattern formation (GO:0009953) | 3.331e-10 | 9.161e-8 | 665.67 | 14526.48 |
| 5 | pituitary gland development (GO:0021983) | 0.000004719 | 0.00009926 | 999.25 | 12254.80 |

GO: Molecular Function

| **Index** | **Name** | **P-value** | **Adjusted p-value** | **Odds Ratio** | **Combined score** |
| --- | --- | --- | --- | --- | --- |
| 1 | polyol transmembrane transporter activity (GO:0015166) | 0.002498 | 0.007739 | 555.17 | 3326.77 |
| 2 | oxidoreductase activity, acting on paired donors, with incorporation or reduction of molecular oxygen, reduced pteridine as one donor, and incorporation of one atom of oxygen (GO:0016714) | 0.002498 | 0.007739 | 555.17 | 3326.77 |
| 3 | type 1 fibroblast growth factor receptor binding (GO:0005105) | 0.002498 | 0.007739 | 555.17 | 3326.77 |
| 4 | type 2 fibroblast growth factor receptor binding (GO:0005111) | 0.002498 | 0.007739 | 555.17 | 3326.77 |
| 5 | morphogen activity (GO:0016015) | 0.003495 | 0.007739 | 370.07 | 2093.27 |

GO: Cellular Component

| **Index** | **Name** | **P-value** | **Adjusted p-value** | **Odds Ratio** | **Combined score** |
| --- | --- | --- | --- | --- | --- |
| 1 | filtration diaphragm (GO:0036056) | 0.002498 | 0.02993 | 555.17 | 3326.77 |
| 2 | slit diaphragm (GO:0036057) | 0.002498 | 0.02993 | 555.17 | 3326.77 |
| 3 | melanosome membrane (GO:0033162) | 0.005985 | 0.02993 | 201.81 | 1032.95 |
| 4 | pigment granule membrane (GO:0090741) | 0.005985 | 0.02993 | 201.81 | 1032.95 |
| 5 | chitosome (GO:0045009) | 0.005985 | 0.02993 | 201.81 | 1032.95 |

Obsessive-compulsive disorder ([HP:0000722](https://hpo.jax.org/app/browse/term/HP:0000722))

KEGG Pathway

| **Index** | **Name** | **P-value** | **Adjusted p-value** | **Odds Ratio** | **Combined score** |
| --- | --- | --- | --- | --- | --- |
| 1 | Nicotine addiction | 2.854e-9 | 1.199e-7 | 369.52 | 7270.07 |
| 2 | Retrograde endocannabinoid signaling | 5.840e-7 | 0.00001226 | 91.88 | 1318.78 |
| 3 | Taste transduction | 0.000009013 | 0.00008975 | 102.79 | 1194.10 |
| 4 | GABAergic synapse | 0.000009993 | 0.00008975 | 99.19 | 1142.03 |
| 5 | Morphine addiction | 0.00001068 | 0.00008975 | 96.93 | 1109.48 |

GO: Biological Process

| **Index** | **Name** | **P-value** | **Adjusted p-value** | **Odds Ratio** | **Combined score** |
| --- | --- | --- | --- | --- | --- |
| 1 | membrane depolarization during action potential (GO:0086010) | 4.685e-10 | 7.683e-8 | 605.09 | 12998.28 |
| 2 | membrane depolarization (GO:0051899) | 1.125e-9 | 9.227e-8 | 475.29 | 9793.38 |
| 3 | neuronal action potential (GO:0019228) | 1.379e-7 | 0.000004524 | 450.47 | 7115.97 |
| 4 | synaptic transmission, GABAergic (GO:0051932) | 0.00001235 | 0.0001446 | 555.03 | 6273.06 |
| 5 | inhibitory synapse assembly (GO:1904862) | 0.00001235 | 0.0001446 | 555.03 | 6273.06 |

GO: Molecular Function

| **Index** | **Name** | **P-value** | **Adjusted p-value** | **Odds Ratio** | **Combined score** |
| --- | --- | --- | --- | --- | --- |
| 1 | voltage-gated sodium channel activity (GO:0005248) | 2.294e-10 | 9.637e-9 | 739.70 | 16417.99 |
| 2 | GABA-A receptor activity (GO:0004890) | 8.685e-8 | 0.000001216 | 535.02 | 8698.92 |
| 3 | sodium channel activity (GO:0005272) | 2.307e-9 | 4.844e-8 | 391.29 | 7781.87 |
| 4 | benzodiazepine receptor activity (GO:0008503) | 0.00001010 | 0.00005304 | 624.44 | 7182.69 |
| 5 | GABA receptor activity (GO:0016917) | 1.379e-7 | 0.000001448 | 450.47 | 7115.97 |

GO: Cellular Component

| **Index** | **Name** | **P-value** | **Adjusted p-value** | **Odds Ratio** | **Combined score** |
| --- | --- | --- | --- | --- | --- |
| 1 | voltage-gated sodium channel complex (GO:0001518) | 7.474e-11 | 1.719e-9 | 1024.46 | 23887.38 |
| 2 | sodium channel complex (GO:0034706) | 3.965e-10 | 3.973e-9 | 633.94 | 13723.68 |
| 3 | GABA-A receptor complex (GO:1902711) | 8.685e-8 | 3.995e-7 | 535.02 | 8698.92 |
| 4 | integral component of plasma membrane (GO:0005887) | 5.182e-10 | 3.973e-9 | 115.51 | 2469.57 |
| 5 | neuron to neuron synapse (GO:0098984) | 0.003994 | 0.009185 | 317.19 | 1751.86 |

Panic attack ([HP:0025269](https://hpo.jax.org/app/browse/term/HP:0025269))

KEGG Pathway

| **Index** | **Name** | **P-value** | **Adjusted p-value** | **Odds Ratio** | **Combined score** |
| --- | --- | --- | --- | --- | --- |
| 1 | Notch signaling pathway | 0.0003792 | 0.004117 | 87.43 | 688.70 |
| 2 | Long-term potentiation | 0.0004889 | 0.004117 | 76.63 | 584.21 |
| 3 | Renal cell carcinoma | 0.0005185 | 0.004117 | 74.34 | 562.35 |
| 4 | Adherens junction | 0.0005489 | 0.004117 | 72.18 | 541.87 |
| 5 | TGF-beta signaling pathway | 0.0009597 | 0.005265 | 54.07 | 375.73 |

GO: Biological Process

| **Index** | **Name** | **P-value** | **Adjusted p-value** | **Odds Ratio** | **Combined score** |
| --- | --- | --- | --- | --- | --- |
| 1 | peptidyl-lysine acetylation (GO:0018394) | 0.00001010 | 0.002869 | 624.44 | 7182.69 |
| 2 | N-terminal protein amino acid acetylation (GO:0006474) | 0.00002690 | 0.002885 | 356.71 | 3753.85 |
| 3 | positive regulation of transcription of Notch receptor target (GO:0007221) | 0.00003048 | 0.002885 | 332.92 | 3461.84 |
| 4 | glial cell-derived neurotrophic factor receptor signaling pathway (GO:0035860) | 0.002498 | 0.01650 | 555.17 | 3326.77 |
| 5 | Schwann cell differentiation (GO:0014037) | 0.002498 | 0.01650 | 555.17 | 3326.77 |

GO: Molecular Function

| **Index** | **Name** | **P-value** | **Adjusted p-value** | **Odds Ratio** | **Combined score** |
| --- | --- | --- | --- | --- | --- |
| 1 | lysine N-acetyltransferase activity, acting on acetyl phosphate as donor (GO:0004468) | 0.002498 | 0.01677 | 555.17 | 3326.77 |
| 2 | STAT family protein binding (GO:0097677) | 0.002997 | 0.01760 | 444.11 | 2580.41 |
| 3 | histone acetyltransferase activity (GO:0004402) | 0.0001038 | 0.002602 | 172.08 | 1578.44 |
| 4 | acetyltransferase activity (GO:0016407) | 0.0001107 | 0.002602 | 166.33 | 1515.06 |
| 5 | phosphatidylethanolamine binding (GO:0008429) | 0.004492 | 0.01796 | 277.53 | 1500.17 |

GO: Cellular Component

| **Index** | **Name** | **P-value** | **Adjusted p-value** | **Odds Ratio** | **Combined score** |
| --- | --- | --- | --- | --- | --- |
| 1 | NURF complex (GO:0016589) | 0.002498 | 0.01915 | 555.17 | 3326.77 |
| 2 | ISWI-type complex (GO:0031010) | 0.004990 | 0.02869 | 246.68 | 1307.49 |
| 3 | MLL1 complex (GO:0071339) | 0.01293 | 0.03717 | 88.73 | 385.85 |
| 4 | MLL1/2 complex (GO:0044665) | 0.01293 | 0.03717 | 88.73 | 385.85 |
| 5 | axon (GO:0030424) | 0.0001190 | 0.002738 | 42.19 | 381.27 |

Paranoia ([HP:0011999](https://hpo.jax.org/app/browse/term/HP:0011999))

KEGG Pathway

| **Index** | **Name** | **P-value** | **Adjusted p-value** | **Odds Ratio** | **Combined score** |
| --- | --- | --- | --- | --- | --- |
| 1 | Porphyrin and chlorophyll metabolism | 0.0002010 | 0.0006029 | 121.64 | 1035.45 |
| 2 | Inositol phosphate metabolism | 0.03591 | 0.04747 | 30.74 | 102.25 |
| 3 | Phosphatidylinositol signaling system | 0.04747 | 0.04747 | 23.03 | 70.18 |

GO: Biological Process

| **Index** | **Name** | **P-value** | **Adjusted p-value** | **Odds Ratio** | **Combined score** |
| --- | --- | --- | --- | --- | --- |
| 1 | middle ear morphogenesis (GO:0042474) | 0.002498 | 0.02532 | 555.17 | 3326.77 |
| 2 | protein retention in Golgi apparatus (GO:0045053) | 0.002498 | 0.02532 | 555.17 | 3326.77 |
| 3 | lysosomal protein catabolic process (GO:1905146) | 0.002498 | 0.02532 | 555.17 | 3326.77 |
| 4 | embryonic viscerocranium morphogenesis (GO:0048703) | 0.002498 | 0.02532 | 555.17 | 3326.77 |
| 5 | enamel mineralization (GO:0070166) | 0.002498 | 0.02532 | 555.17 | 3326.77 |

GO: Molecular Function

| **Index** | **Name** | **P-value** | **Adjusted p-value** | **Odds Ratio** | **Combined score** |
| --- | --- | --- | --- | --- | --- |
| 1 | primary miRNA binding (GO:0070878) | 0.002997 | 0.02097 | 444.11 | 2580.41 |
| 2 | oxidoreductase activity, acting on the CH-CH group of donors, oxygen as acceptor (GO:0016634) | 0.003495 | 0.02097 | 370.07 | 2093.27 |
| 3 | alkali metal ion binding (GO:0031420) | 0.006482 | 0.02917 | 184.98 | 932.06 |
| 4 | manganese ion binding (GO:0030145) | 0.02375 | 0.08549 | 47.15 | 176.34 |
| 5 | protein homodimerization activity (GO:0042803) | 0.0001825 | 0.003286 | 20.42 | 175.78 |

GO: Cellular Component

| **Index** | **Name** | **P-value** | **Adjusted p-value** | **Odds Ratio** | **Combined score** |
| --- | --- | --- | --- | --- | --- |
| 1 | neuronal dense core vesicle (GO:0098992) | 0.002498 | 0.01124 | 555.17 | 3326.77 |
| 2 | mitochondrial envelope (GO:0005740) | 0.00002904 | 0.0005228 | 68.66 | 717.28 |
| 3 | mitochondrial intermembrane space (GO:0005758) | 0.0003664 | 0.002677 | 88.99 | 704.07 |
| 4 | organelle envelope lumen (GO:0031970) | 0.0004462 | 0.002677 | 80.35 | 619.92 |
| 5 | lipid droplet (GO:0005811) | 0.03785 | 0.1363 | 29.11 | 95.32 |

Phonophobia ([HP:0002183](https://hpo.jax.org/app/browse/term/HP:0002183))

KEGG Pathway

| **Index** | **Name** | **P-value** | **Adjusted p-value** | **Odds Ratio** | **Combined score** |
| --- | --- | --- | --- | --- | --- |
| 1 | Cholesterol metabolism | 0.02473 | 0.06066 | 45.22 | 167.30 |
| 2 | Dopaminergic synapse | 0.06409 | 0.08011 | 16.84 | 46.28 |
| 3 | Endocytosis | 0.1191 | 0.1191 | 8.74 | 18.59 |
| 4 | PPAR signaling pathway | 0.03640 | 0.06066 | 30.32 | 100.44 |
| 5 | Primary bile acid biosynthesis | 0.008469 | 0.04235 | 138.71 | 661.82 |

GO: Biological Process

| **Index** | **Name** | **P-value** | **Adjusted p-value** | **Odds Ratio** | **Combined score** |
| --- | --- | --- | --- | --- | --- |
| 1 | short-term memory (GO:0007614) | 0.002498 | 0.03666 | 555.17 | 3326.77 |
| 2 | regulation of catecholamine uptake involved in synaptic transmission (GO:0051940) | 0.002498 | 0.03666 | 555.17 | 3326.77 |
| 3 | vitamin D3 metabolic process (GO:0070640) | 0.002498 | 0.03666 | 555.17 | 3326.77 |
| 4 | detection of stimulus involved in sensory perception of pain (GO:0062149) | 0.003495 | 0.03666 | 370.07 | 2093.27 |
| 5 | regulation of dopamine uptake involved in synaptic transmission (GO:0051584) | 0.003994 | 0.03666 | 317.19 | 1751.86 |

GO: Molecular Function

| **Index** | **Name** | **P-value** | **Adjusted p-value** | **Odds Ratio** | **Combined score** |
| --- | --- | --- | --- | --- | --- |
| 1 | CD4 receptor binding (GO:0042609) | 0.003994 | 0.04475 | 317.19 | 1751.86 |
| 2 | outward rectifier potassium channel activity (GO:0015271) | 0.005488 | 0.04475 | 222.00 | 1155.57 |
| 3 | voltage-gated sodium channel activity (GO:0005248) | 0.01095 | 0.04475 | 105.66 | 476.99 |
| 4 | ion gated channel activity (GO:0022839) | 0.01293 | 0.04475 | 88.73 | 385.85 |
| 5 | kinesin binding (GO:0019894) | 0.01441 | 0.04475 | 79.21 | 335.86 |

GO: Cellular Component

| **Index** | **Name** | **P-value** | **Adjusted p-value** | **Odds Ratio** | **Combined score** |
| --- | --- | --- | --- | --- | --- |
| 1 | sarcoglycan complex (GO:0016012) | 0.002498 | 0.03195 | 555.17 | 3326.77 |
| 2 | dystroglycan complex (GO:0016011) | 0.002997 | 0.03195 | 444.11 | 2580.41 |
| 3 | extrinsic component of endoplasmic reticulum membrane (GO:0042406) | 0.003994 | 0.03195 | 317.19 | 1751.86 |
| 4 | trans-Golgi network transport vesicle (GO:0030140) | 0.007476 | 0.04065 | 158.54 | 776.21 |
| 5 | voltage-gated sodium channel complex (GO:0001518) | 0.008469 | 0.04065 | 138.71 | 661.82 |

Polydipsia ([HP:0001959](https://hpo.jax.org/app/browse/term/HP:0001959))

KEGG Pathway

| **Index** | **Name** | **P-value** | **Adjusted p-value** | **Odds Ratio** | **Combined score** |
| --- | --- | --- | --- | --- | --- |
| 1 | Thyroid hormone signaling pathway | 0.000007427 | 0.0003417 | 126.32 | 1491.93 |
| 2 | Notch signaling pathway | 0.0001780 | 0.002373 | 139.90 | 1207.91 |
| 3 | Long-term potentiation | 0.0002297 | 0.002373 | 122.63 | 1027.54 |
| 4 | Renal cell carcinoma | 0.0002436 | 0.002373 | 118.96 | 989.76 |
| 5 | Adherens junction | 0.0002579 | 0.002373 | 115.50 | 954.36 |

GO: Biological Process

| **Index** | **Name** | **P-value** | **Adjusted p-value** | **Odds Ratio** | **Combined score** |
| --- | --- | --- | --- | --- | --- |
| 1 | peptidyl-lysine acetylation (GO:0018394) | 0.000004719 | 0.0004823 | 999.25 | 12254.80 |
| 2 | N-terminal protein amino acid acetylation (GO:0006474) | 0.00001257 | 0.0004823 | 570.83 | 6441.32 |
| 3 | positive regulation of transcription of Notch receptor target (GO:0007221) | 0.00001424 | 0.0004823 | 532.75 | 5945.01 |
| 4 | negative regulation of cellular respiration (GO:1901856) | 0.001749 | 0.008819 | 832.88 | 5287.72 |
| 5 | foam cell differentiation (GO:0090077) | 0.002098 | 0.009751 | 666.27 | 4108.59 |

GO: Molecular Function

| **Index** | **Name** | **P-value** | **Adjusted p-value** | **Odds Ratio** | **Combined score** |
| --- | --- | --- | --- | --- | --- |
| 1 | lysine N-acetyltransferase activity, acting on acetyl phosphate as donor (GO:0004468) | 0.001749 | 0.008538 | 832.88 | 5287.72 |
| 2 | RNA polymerase II general transcription initiation factor binding (GO:0001091) | 0.001749 | 0.008538 | 832.88 | 5287.72 |
| 3 | STAT family protein binding (GO:0097677) | 0.002098 | 0.008538 | 666.27 | 4108.59 |
| 4 | superoxide-generating NADPH oxidase activator activity (GO:0016176) | 0.002797 | 0.01002 | 475.86 | 2797.66 |
| 5 | histone acetyltransferase activity (GO:0004402) | 0.00004859 | 0.001114 | 275.37 | 2734.96 |

GO: Cellular Component

| **Index** | **Name** | **P-value** | **Adjusted p-value** | **Odds Ratio** | **Combined score** |
| --- | --- | --- | --- | --- | --- |
| 1 | phagolysosome (GO:0032010) | 0.001749 | 0.01137 | 832.88 | 5287.72 |
| 2 | elastic fiber (GO:0071953) | 0.001749 | 0.01137 | 832.88 | 5287.72 |
| 3 | NADPH oxidase complex (GO:0043020) | 0.004193 | 0.01724 | 302.76 | 1657.40 |
| 4 | secondary lysosome (GO:0005767) | 0.005587 | 0.01724 | 221.98 | 1151.45 |
| 5 | supramolecular fiber (GO:0099512) | 0.006632 | 0.01724 | 184.95 | 927.70 |

Short attention span ([HP:0000736](https://hpo.jax.org/app/browse/term/HP:0000736))

KEGG Pathway

| **Index** | **Name** | **P-value** | **Adjusted p-value** | **Odds Ratio** | **Combined score** |
| --- | --- | --- | --- | --- | --- |
| 1 | Nicotine addiction | 8.806e-7 | 0.00002114 | 231.12 | 3222.36 |
| 2 | GABAergic synapse | 0.000009993 | 0.0001199 | 99.19 | 1142.03 |
| 3 | Taste transduction | 0.0008042 | 0.004319 | 59.24 | 422.16 |
| 4 | Morphine addiction | 0.0008998 | 0.004319 | 55.90 | 392.06 |
| 5 | Dopaminergic synapse | 0.001879 | 0.007516 | 38.19 | 239.73 |

GO: Biological Process

| **Index** | **Name** | **P-value** | **Adjusted p-value** | **Odds Ratio** | **Combined score** |
| --- | --- | --- | --- | --- | --- |
| 1 | neuronal action potential (GO:0019228) | 2.294e-10 | 1.698e-8 | 739.70 | 16417.99 |
| 2 | synaptic transmission, GABAergic (GO:0051932) | 0.00001235 | 0.00005710 | 555.03 | 6273.06 |
| 3 | inhibitory synapse assembly (GO:1904862) | 0.00001235 | 0.00005710 | 555.03 | 6273.06 |
| 4 | action potential (GO:0001508) | 4.648e-9 | 1.720e-7 | 324.37 | 6223.71 |
| 5 | membrane depolarization during action potential (GO:0086010) | 2.326e-7 | 0.000002869 | 372.06 | 5682.76 |

GO: Molecular Function

| **Index** | **Name** | **P-value** | **Adjusted p-value** | **Odds Ratio** | **Combined score** |
| --- | --- | --- | --- | --- | --- |
| 1 | benzodiazepine receptor activity (GO:0008503) | 0.00001010 | 0.0001296 | 624.44 | 7182.69 |
| 2 | voltage-gated sodium channel activity (GO:0005248) | 1.379e-7 | 0.000005792 | 450.47 | 7115.97 |
| 3 | extracellular ligand-gated ion channel activity (GO:0005230) | 0.00001235 | 0.0001296 | 555.03 | 6273.06 |
| 4 | GABA-gated chloride ion channel activity (GO:0022851) | 0.00001750 | 0.0001429 | 454.07 | 4973.59 |
| 5 | inhibitory extracellular ligand-gated ion channel activity (GO:0005237) | 0.00002041 | 0.0001429 | 416.21 | 4494.85 |

GO: Cellular Component

| **Index** | **Name** | **P-value** | **Adjusted p-value** | **Odds Ratio** | **Combined score** |
| --- | --- | --- | --- | --- | --- |
| 1 | voltage-gated sodium channel complex (GO:0001518) | 6.098e-8 | 2.896e-7 | 611.51 | 10158.88 |
| 2 | sodium channel complex (GO:0034706) | 2.058e-7 | 7.821e-7 | 388.99 | 5988.96 |
| 3 | GABA-A receptor complex (GO:1902711) | 0.00003830 | 0.0001213 | 293.72 | 2987.15 |
| 4 | integral component of plasma membrane (GO:0005887) | 5.182e-10 | 9.846e-9 | 115.51 | 2469.57 |
| 5 | NMDA selective glutamate receptor complex (GO:0017146) | 0.003994 | 0.007588 | 317.19 | 1751.86 |

Visual hallucinations ([HP:0002367](https://hpo.jax.org/app/browse/term/HP:0002367))

KEGG Pathway

| **Index** | **Name** | **P-value** | **Adjusted p-value** | **Odds Ratio** | **Combined score** |
| --- | --- | --- | --- | --- | --- |
| 1 | Inositol phosphate metabolism | 0.03238 | 0.1449 | 34.58 | 118.62 |
| 2 | ECM-receptor interaction | 0.03892 | 0.1449 | 28.60 | 92.84 |
| 3 | Spinocerebellar ataxia | 0.06255 | 0.1449 | 17.47 | 48.43 |
| 4 | mTOR signaling pathway | 0.06722 | 0.1449 | 16.21 | 43.76 |
| 5 | Protein processing in endoplasmic reticulum | 0.07438 | 0.1449 | 14.57 | 37.87 |

GO: Biological Process

| **Index** | **Name** | **P-value** | **Adjusted p-value** | **Odds Ratio** | **Combined score** |
| --- | --- | --- | --- | --- | --- |
| 1 | positive regulation of AMPA receptor activity (GO:2000969) | 0.002248 | 0.03359 | 624.59 | 3808.55 |
| 2 | reelin-mediated signaling pathway (GO:0038026) | 0.002248 | 0.03359 | 624.59 | 3808.55 |
| 3 | cerebral cortex cell migration (GO:0021795) | 0.002697 | 0.03359 | 499.65 | 2955.69 |
| 4 | regulation of oxidative stress-induced neuron death (GO:1903203) | 0.003595 | 0.03359 | 356.86 | 2008.48 |
| 5 | TORC1 signaling (GO:0038202) | 0.003595 | 0.03359 | 356.86 | 2008.48 |

GO: Molecular Function

| **Index** | **Name** | **P-value** | **Adjusted p-value** | **Odds Ratio** | **Combined score** |
| --- | --- | --- | --- | --- | --- |
| 1 | cuprous ion binding (GO:1903136) | 0.003595 | 0.02845 | 356.86 | 2008.48 |
| 2 | phospholipase inhibitor activity (GO:0004859) | 0.004492 | 0.02845 | 277.53 | 1500.17 |
| 3 | lipase inhibitor activity (GO:0055102) | 0.004492 | 0.02845 | 277.53 | 1500.17 |
| 4 | lipoprotein particle receptor binding (GO:0070325) | 0.01253 | 0.05953 | 92.43 | 404.78 |
| 5 | copper ion binding (GO:0005507) | 0.02007 | 0.07628 | 56.67 | 221.48 |

GO: Cellular Component

| **Index** | **Name** | **P-value** | **Adjusted p-value** | **Odds Ratio** | **Combined score** |
| --- | --- | --- | --- | --- | --- |
| 1 | Lewy body (GO:0097413) | 0.002248 | 0.03595 | 624.59 | 3808.55 |
| 2 | clathrin-sculpted gamma-aminobutyric acid transport vesicle (GO:0061200) | 0.003595 | 0.03595 | 356.86 | 2008.48 |
| 3 | clathrin-sculpted gamma-aminobutyric acid transport vesicle membrane (GO:0061202) | 0.003595 | 0.03595 | 356.86 | 2008.48 |
| 4 | SCF ubiquitin ligase complex (GO:0019005) | 0.02580 | 0.07548 | 43.71 | 159.87 |
| 5 | azurophil granule membrane (GO:0035577) | 0.02580 | 0.07548 | 43.71 | 159.87 |

Agnosia ([HP:0010524](https://hpo.jax.org/app/browse/term/HP:0010524))

KEGG Pathway

| **Index** | **Name** | **P-value** | **Adjusted p-value** | **Odds Ratio** | **Combined score** |
| --- | --- | --- | --- | --- | --- |
| 1 | Aldosterone-regulated sodium reabsorption | 0.0001189 | 0.003782 | 162.91 | 1472.21 |
| 2 | Fanconi anemia pathway | 0.0002545 | 0.003782 | 109.55 | 906.71 |
| 3 | Endometrial cancer | 0.0002937 | 0.003782 | 101.71 | 827.20 |
| 4 | VEGF signaling pathway | 0.0003039 | 0.003782 | 99.92 | 809.23 |
| 5 | GnRH secretion | 0.0003577 | 0.003782 | 91.84 | 728.82 |

GO: Biological Process

| **Index** | **Name** | **P-value** | **Adjusted p-value** | **Odds Ratio** | **Combined score** |
| --- | --- | --- | --- | --- | --- |
| 1 | positive regulation of amyloid-beta clearance (GO:1900223) | 0.000003775 | 0.0008690 | 1142.06 | 14260.84 |
| 2 | positive regulation of apoptotic cell clearance (GO:2000427) | 0.000005033 | 0.0008690 | 951.67 | 11609.88 |
| 3 | positive regulation of phagocytosis, engulfment (GO:0060100) | 0.000006469 | 0.0008690 | 815.67 | 9746.03 |
| 4 | regulation of amyloid-beta clearance (GO:1900221) | 0.00002153 | 0.002169 | 407.69 | 4381.12 |
| 5 | regulation of astrocyte activation (GO:0061888) | 0.002248 | 0.01960 | 624.59 | 3808.55 |

GO: Molecular Function

| **Index** | **Name** | **P-value** | **Adjusted p-value** | **Odds Ratio** | **Combined score** |
| --- | --- | --- | --- | --- | --- |
| 1 | DNA insertion or deletion binding (GO:0032135) | 0.002248 | 0.01560 | 624.59 | 3808.55 |
| 2 | phosphatidylcholine floppase activity (GO:0090554) | 0.002248 | 0.01560 | 624.59 | 3808.55 |
| 3 | apolipoprotein A-I binding (GO:0034186) | 0.002248 | 0.01560 | 624.59 | 3808.55 |
| 4 | sulfatide binding (GO:0120146) | 0.002697 | 0.01560 | 499.65 | 2955.69 |
| 5 | high-density lipoprotein particle binding (GO:0008035) | 0.002697 | 0.01560 | 499.65 | 2955.69 |

GO: Cellular Component

| **Index** | **Name** | **P-value** | **Adjusted p-value** | **Odds Ratio** | **Combined score** |
| --- | --- | --- | --- | --- | --- |
| 1 | glial cell projection (GO:0097386) | 0.00001633 | 0.0006207 | 475.69 | 5243.18 |
| 2 | phosphatidylinositol 3-kinase complex, class I (GO:0097651) | 0.002248 | 0.02174 | 624.59 | 3808.55 |
| 3 | neurofibrillary tangle (GO:0097418) | 0.002248 | 0.02174 | 624.59 | 3808.55 |
| 4 | proton-transporting V-type ATPase complex (GO:0033176) | 0.006731 | 0.05116 | 178.37 | 892.01 |
| 5 | vacuolar proton-transporting V-type ATPase complex (GO:0016471) | 0.008519 | 0.05395 | 138.70 | 660.97 |

Bradykinesia ([HP:0002067](https://hpo.jax.org/app/browse/term/HP:0002067))

KEGG Pathway

| **Index** | **Name** | **P-value** | **Adjusted p-value** | **Odds Ratio** | **Combined score** |
| --- | --- | --- | --- | --- | --- |
| 1 | Nicotine addiction | 8.806e-7 | 0.00002466 | 231.12 | 3222.36 |
| 2 | Taste transduction | 0.000009013 | 0.00007479 | 102.79 | 1194.10 |
| 3 | GABAergic synapse | 0.000009993 | 0.00007479 | 99.19 | 1142.03 |
| 4 | Morphine addiction | 0.00001068 | 0.00007479 | 96.93 | 1109.48 |
| 5 | Retrograde endocannabinoid signaling | 0.00004587 | 0.0002569 | 58.66 | 585.95 |

GO: Biological Process

| **Index** | **Name** | **P-value** | **Adjusted p-value** | **Odds Ratio** | **Combined score** |
| --- | --- | --- | --- | --- | --- |
| 1 | membrane depolarization during action potential (GO:0086010) | 4.685e-10 | 2.460e-8 | 605.09 | 12998.28 |
| 2 | membrane depolarization (GO:0051899) | 1.125e-9 | 3.938e-8 | 475.29 | 9793.38 |
| 3 | neuronal action potential (GO:0019228) | 1.379e-7 | 0.000002414 | 450.47 | 7115.97 |
| 4 | synaptic transmission, GABAergic (GO:0051932) | 0.00001235 | 0.00009259 | 555.03 | 6273.06 |
| 5 | inhibitory synapse assembly (GO:1904862) | 0.00001235 | 0.00009259 | 555.03 | 6273.06 |

GO: Molecular Function

| **Index** | **Name** | **P-value** | **Adjusted p-value** | **Odds Ratio** | **Combined score** |
| --- | --- | --- | --- | --- | --- |
| 1 | voltage-gated sodium channel activity (GO:0005248) | 2.294e-10 | 7.801e-9 | 739.70 | 16417.99 |
| 2 | GABA-A receptor activity (GO:0004890) | 8.685e-8 | 9.843e-7 | 535.02 | 8698.92 |
| 3 | sodium channel activity (GO:0005272) | 2.307e-9 | 3.921e-8 | 391.29 | 7781.87 |
| 4 | benzodiazepine receptor activity (GO:0008503) | 0.00001010 | 0.00004907 | 624.44 | 7182.69 |
| 5 | GABA receptor activity (GO:0016917) | 1.379e-7 | 0.000001172 | 450.47 | 7115.97 |

GO: Cellular Component

| **Index** | **Name** | **P-value** | **Adjusted p-value** | **Odds Ratio** | **Combined score** |
| --- | --- | --- | --- | --- | --- |
| 1 | voltage-gated sodium channel complex (GO:0001518) | 7.474e-11 | 2.093e-9 | 1024.46 | 23887.38 |
| 2 | sodium channel complex (GO:0034706) | 3.965e-10 | 4.837e-9 | 633.94 | 13723.68 |
| 3 | GABA-A receptor complex (GO:1902711) | 8.685e-8 | 4.863e-7 | 535.02 | 8698.92 |
| 4 | integral component of plasma membrane (GO:0005887) | 5.182e-10 | 4.837e-9 | 115.51 | 2469.57 |
| 5 | neuron to neuron synapse (GO:0098984) | 0.003994 | 0.01398 | 317.19 | 1751.86 |

Cognitive impairment ([HP:0100543](https://hpo.jax.org/app/browse/term/HP:0100543))

KEGG Pathway

| **Index** | **Name** | **P-value** | **Adjusted p-value** | **Odds Ratio** | **Combined score** |
| --- | --- | --- | --- | --- | --- |
| 1 | Nicotine addiction | 2.854e-9 | 4.852e-8 | 369.52 | 7270.07 |
| 2 | GABAergic synapse | 7.537e-8 | 4.673e-7 | 156.12 | 2560.47 |
| 3 | Morphine addiction | 8.246e-8 | 4.673e-7 | 152.51 | 2487.64 |
| 4 | Retrograde endocannabinoid signaling | 5.840e-7 | 0.000002482 | 91.88 | 1318.78 |
| 5 | Taste transduction | 0.0008042 | 0.002278 | 59.24 | 422.16 |

GO: Biological Process

| **Index** | **Name** | **P-value** | **Adjusted p-value** | **Odds Ratio** | **Combined score** |
| --- | --- | --- | --- | --- | --- |
| 1 | neuronal action potential (GO:0019228) | 2.294e-10 | 1.170e-8 | 739.70 | 16417.99 |
| 2 | membrane depolarization (GO:0051899) | 1.125e-9 | 2.869e-8 | 475.29 | 9793.38 |
| 3 | synaptic transmission, GABAergic (GO:0051932) | 0.00001235 | 0.00004197 | 555.03 | 6273.06 |
| 4 | inhibitory synapse assembly (GO:1904862) | 0.00001235 | 0.00004197 | 555.03 | 6273.06 |
| 5 | action potential (GO:0001508) | 4.648e-9 | 7.901e-8 | 324.37 | 6223.71 |

GO: Molecular Function

| **Index** | **Name** | **P-value** | **Adjusted p-value** | **Odds Ratio** | **Combined score** |
| --- | --- | --- | --- | --- | --- |
| 1 | GABA-A receptor activity (GO:0004890) | 8.685e-8 | 0.000001195 | 535.02 | 8698.92 |
| 2 | benzodiazepine receptor activity (GO:0008503) | 0.00001010 | 0.00003060 | 624.44 | 7182.69 |
| 3 | voltage-gated sodium channel activity (GO:0005248) | 1.379e-7 | 0.000001195 | 450.47 | 7115.97 |
| 4 | GABA receptor activity (GO:0016917) | 1.379e-7 | 0.000001195 | 450.47 | 7115.97 |
| 5 | extracellular ligand-gated ion channel activity (GO:0005230) | 0.00001235 | 0.00003060 | 555.03 | 6273.06 |

GO: Cellular Component

| **Index** | **Name** | **P-value** | **Adjusted p-value** | **Odds Ratio** | **Combined score** |
| --- | --- | --- | --- | --- | --- |
| 1 | voltage-gated sodium channel complex (GO:0001518) | 6.098e-8 | 1.677e-7 | 611.51 | 10158.88 |
| 2 | GABA-A receptor complex (GO:1902711) | 8.685e-8 | 1.911e-7 | 535.02 | 8698.92 |
| 3 | sodium channel complex (GO:0034706) | 2.058e-7 | 3.773e-7 | 388.99 | 5988.96 |
| 4 | dendrite membrane (GO:0032590) | 3.266e-7 | 5.133e-7 | 329.08 | 4914.57 |
| 5 | neuron projection (GO:0043005) | 1.454e-11 | 1.600e-10 | 141.91 | 3541.27 |

Confusion ([HP:0001289](https://hpo.jax.org/app/browse/term/HP:0001289))

KEGG Pathway

| **Index** | **Name** | **P-value** | **Adjusted p-value** | **Odds Ratio** | **Combined score** |
| --- | --- | --- | --- | --- | --- |
| 1 | Nucleotide excision repair | 0.000001443 | 0.00001443 | 194.28 | 2612.86 |
| 2 | Arginine biosynthesis | 0.01095 | 0.03979 | 105.66 | 476.99 |
| 3 | Vitamin digestion and absorption | 0.01194 | 0.03979 | 96.46 | 427.12 |
| 4 | Basal transcription factors | 0.02228 | 0.05336 | 50.37 | 191.61 |
| 5 | Fanconi anemia pathway | 0.02668 | 0.05336 | 41.80 | 151.46 |

GO: Biological Process

| **Index** | **Name** | **P-value** | **Adjusted p-value** | **Odds Ratio** | **Combined score** |
| --- | --- | --- | --- | --- | --- |
| 1 | nucleotide-excision repair, DNA incision, 3'-to lesion (GO:0006295) | 1.191e-7 | 0.000009233 | 475.52 | 7581.26 |
| 2 | nucleotide-excision repair, preincision complex stabilization (GO:0006293) | 1.191e-7 | 0.000009233 | 475.52 | 7581.26 |
| 3 | negative regulation of telomerase activity (GO:0051974) | 0.00001010 | 0.0002610 | 624.44 | 7182.69 |
| 4 | urea cycle (GO:0000050) | 0.00001010 | 0.0002610 | 624.44 | 7182.69 |
| 5 | UV protection (GO:0009650) | 0.00001010 | 0.0002610 | 624.44 | 7182.69 |

GO: Molecular Function

| **Index** | **Name** | **P-value** | **Adjusted p-value** | **Odds Ratio** | **Combined score** |
| --- | --- | --- | --- | --- | --- |
| 1 | deoxyribonuclease activity (GO:0004536) | 0.00002041 | 0.0005613 | 416.21 | 4494.85 |
| 2 | oxidoreductase activity, acting on metal ions, NAD or NADP as acceptor (GO:0016723) | 0.002498 | 0.01249 | 555.17 | 3326.77 |
| 3 | L-ornithine transmembrane transporter activity (GO:0000064) | 0.002498 | 0.01249 | 555.17 | 3326.77 |
| 4 | Y-form DNA binding (GO:0000403) | 0.002498 | 0.01249 | 555.17 | 3326.77 |
| 5 | apolipoprotein A-I binding (GO:0034186) | 0.002498 | 0.01249 | 555.17 | 3326.77 |

GO: Cellular Component

| **Index** | **Name** | **P-value** | **Adjusted p-value** | **Odds Ratio** | **Combined score** |
| --- | --- | --- | --- | --- | --- |
| 1 | transcription factor TFIIH core complex (GO:0000439) | 0.004990 | 0.04870 | 246.68 | 1307.49 |
| 2 | transcription factor TFIIH holo complex (GO:0005675) | 0.005985 | 0.04870 | 201.81 | 1032.95 |
| 3 | acetylcholine-gated channel complex (GO:0005892) | 0.006979 | 0.04870 | 170.74 | 847.70 |
| 4 | voltage-gated sodium channel complex (GO:0001518) | 0.008469 | 0.04870 | 138.71 | 661.82 |
| 5 | sodium channel complex (GO:0034706) | 0.01243 | 0.05706 | 92.44 | 405.55 |

Encephalopathy ([HP:0001298](https://hpo.jax.org/app/browse/term/HP:0001298))

KEGG Pathway

| **Index** | **Name** | **P-value** | **Adjusted p-value** | **Odds Ratio** | **Combined score** |
| --- | --- | --- | --- | --- | --- |
| 1 | Nicotine addiction | 6.174e-12 | 1.667e-10 | 570.14 | 14715.76 |
| 2 | GABAergic synapse | 7.537e-8 | 0.000001017 | 156.12 | 2560.47 |
| 3 | Morphine addiction | 0.00001068 | 0.00007212 | 96.93 | 1109.48 |
| 4 | Cocaine addiction | 0.0002613 | 0.001008 | 106.08 | 875.14 |
| 5 | Neuroactive ligand-receptor interaction | 3.287e-7 | 0.000002958 | 58.49 | 873.21 |

GO: Biological Process

| **Index** | **Name** | **P-value** | **Adjusted p-value** | **Odds Ratio** | **Combined score** |
| --- | --- | --- | --- | --- | --- |
| 1 | synaptic transmission, GABAergic (GO:0051932) | 1.482e-8 | 2.964e-7 | 1070.46 | 19297.66 |
| 2 | inhibitory synapse assembly (GO:1904862) | 1.482e-8 | 2.964e-7 | 1070.46 | 19297.66 |
| 3 | cellular response to histamine (GO:0071420) | 0.000004719 | 0.00004356 | 999.25 | 12254.80 |
| 4 | excitatory chemical synaptic transmission (GO:0098976) | 0.000004719 | 0.00004356 | 999.25 | 12254.80 |
| 5 | regulation of postsynaptic membrane potential (GO:0060078) | 1.637e-9 | 6.549e-8 | 429.23 | 8683.34 |

GO: Molecular Function

| **Index** | **Name** | **P-value** | **Adjusted p-value** | **Odds Ratio** | **Combined score** |
| --- | --- | --- | --- | --- | --- |
| 1 | glutamate-gated calcium ion channel activity (GO:0022849) | 0.000002248 | 0.00001799 | 1665.58 | 21661.60 |
| 2 | GABA-gated chloride ion channel activity (GO:0022851) | 2.567e-8 | 0.000001232 | 856.29 | 14966.03 |
| 3 | NMDA glutamate receptor activity (GO:0004972) | 0.000006290 | 0.00003355 | 832.67 | 9972.50 |
| 4 | ligand-gated anion channel activity (GO:0099095) | 7.315e-8 | 0.000001390 | 570.71 | 9377.24 |
| 5 | GABA-A receptor activity (GO:0004890) | 8.685e-8 | 0.000001390 | 535.02 | 8698.92 |

GO: Cellular Component

| **Index** | **Name** | **P-value** | **Adjusted p-value** | **Odds Ratio** | **Combined score** |
| --- | --- | --- | --- | --- | --- |
| 1 | integral component of plasma membrane (GO:0005887) | 4.006e-12 | 1.082e-10 | 185460.00 | 4867070.13 |
| 2 | NMDA selective glutamate receptor complex (GO:0017146) | 0.000006290 | 0.00002830 | 832.67 | 9972.50 |
| 3 | GABA-A receptor complex (GO:1902711) | 8.685e-8 | 7.816e-7 | 535.02 | 8698.92 |
| 4 | neuron projection (GO:0043005) | 1.454e-11 | 1.963e-10 | 141.91 | 3541.27 |
| 5 | voltage-gated sodium channel complex (GO:0001518) | 0.00003048 | 0.0001176 | 332.92 | 3461.84 |

Neuropsychiatric-emotion-mood

Abnormal emotion/affect behavior ([HP:0100851](https://hpo.jax.org/app/browse/term/HP:0100851))

KEGG Pathway

| **Index** | **Name** | **P-value** | **Adjusted p-value** | **Odds Ratio** | **Combined score** |
| --- | --- | --- | --- | --- | --- |
| 1 | Nicotine addiction | 5.437e-21 | 2.882e-19 | 2494.75 | 116407.64 |
| 2 | Amphetamine addiction | 5.059e-16 | 1.341e-14 | 749.98 | 26414.43 |
| 3 | Circadian entrainment | 6.002e-15 | 1.060e-13 | 515.93 | 16894.89 |
| 4 | Long-term potentiation | 9.011e-11 | 5.970e-10 | 321.42 | 7434.41 |
| 5 | Glutamatergic synapse | 6.187e-12 | 5.466e-11 | 276.14 | 7126.73 |

GO: Biological Process

| **Index** | **Name** | **P-value** | **Adjusted p-value** | **Odds Ratio** | **Combined score** |
| --- | --- | --- | --- | --- | --- |
| 1 | regulation of NMDA receptor activity (GO:2000310) | 1.586e-18 | 1.950e-16 | 1863.40 | 76372.57 |
| 2 | excitatory chemical synaptic transmission (GO:0098976) | 3.147e-9 | 4.838e-8 | 2141.36 | 41921.23 |
| 3 | regulation of neurotransmitter receptor activity (GO:0099601) | 1.412e-16 | 5.788e-15 | 912.24 | 33293.64 |
| 4 | glutamate receptor signaling pathway (GO:0007215) | 5.464e-15 | 1.120e-13 | 965.76 | 31715.99 |
| 5 | anterograde trans-synaptic signaling (GO:0098916) | 5.108e-17 | 3.142e-15 | 756.57 | 28381.44 |

GO: Molecular Function

| **Index** | **Name** | **P-value** | **Adjusted p-value** | **Odds Ratio** | **Combined score** |
| --- | --- | --- | --- | --- | --- |
| 1 | ionotropic glutamate receptor activity (GO:0004970) | 2.918e-17 | 1.430e-15 | 2724.41 | 103726.10 |
| 2 | glutamate-gated calcium ion channel activity (GO:0022849) | 8.995e-10 | 6.296e-9 | 4283.14 | 89214.48 |
| 3 | ligand-gated channel activity (GO:0022834) | 1.397e-15 | 2.830e-14 | 1247.88 | 42682.61 |
| 4 | ligand-gated ion channel activity (GO:0015276) | 1.732e-15 | 2.830e-14 | 1197.90 | 40715.77 |
| 5 | NMDA glutamate receptor activity (GO:0004972) | 5.033e-9 | 2.740e-8 | 1713.00 | 32730.63 |

GO: Cellular Component

| **Index** | **Name** | **P-value** | **Adjusted p-value** | **Odds Ratio** | **Combined score** |
| --- | --- | --- | --- | --- | --- |
| 1 | ionotropic glutamate receptor complex (GO:0008328) | 2.132e-15 | 6.395e-14 | 1151.77 | 38908.86 |
| 2 | NMDA selective glutamate receptor complex (GO:0017146) | 5.033e-9 | 3.775e-8 | 1713.00 | 32730.63 |
| 3 | cation channel complex (GO:0034703) | 3.977e-13 | 5.966e-12 | 446.04 | 12735.72 |
| 4 | AMPA glutamate receptor complex (GO:0032281) | 1.021e-7 | 6.129e-7 | 503.52 | 8105.11 |
| 5 | synaptic membrane (GO:0097060) | 2.326e-7 | 0.000001163 | 372.06 | 5682.76 |

Abnormal fear/anxiety-related behavior ([HP:0100852](https://hpo.jax.org/app/browse/term/HP:0100852))

GO: Biological Process

| **Index** | **Name** | **P-value** | **Adjusted p-value** | **Odds Ratio** | **Combined score** |
| --- | --- | --- | --- | --- | --- |
| 1 | equilibrioception (GO:0050957) | 5.668e-17 | 3.287e-15 | 19989.00 | 747771.16 |
| 2 | neuromuscular process controlling balance (GO:0050885) | 1.888e-14 | 2.738e-13 | 2220.11 | 70156.55 |
| 3 | sensory perception (GO:0007600) | 4.223e-14 | 4.899e-13 | 664.83 | 20473.93 |
| 4 | sensory perception of mechanical stimulus (GO:0050954) | 2.969e-15 | 7.316e-14 | 573.51 | 19184.17 |
| 5 | sensory perception of sound (GO:0007605) | 3.784e-15 | 7.316e-14 | 552.94 | 18362.12 |

GO: Molecular Function

| **Index** | **Name** | **P-value** | **Adjusted p-value** | **Odds Ratio** | **Combined score** |
| --- | --- | --- | --- | --- | --- |
| 1 | myosin binding (GO:0017022) | 0.02766 | 0.05531 | 40.27 | 144.50 |
| 2 | calcium ion binding (GO:0005509) | 0.01239 | 0.05531 | 14.19 | 62.33 |
| 3 | magnesium ion binding (GO:0000287) | 0.07066 | 0.1060 | 15.21 | 40.30 |
| 4 | metal ion binding (GO:0046872) | 0.02616 | 0.05531 | 9.45 | 34.45 |
| 5 | cadherin binding (GO:0045296) | 0.1499 | 0.1798 | 6.81 | 12.92 |

GO: Cellular Component

| **Index** | **Name** | **P-value** | **Adjusted p-value** | **Odds Ratio** | **Combined score** |
| --- | --- | --- | --- | --- | --- |
| 1 | microvillus (GO:0005902) | 0.0003539 | 0.003893 | 90.61 | 720.07 |
| 2 | actin-based cell projection (GO:0098858) | 0.0007493 | 0.004121 | 61.45 | 442.20 |
| 3 | catenin complex (GO:0016342) | 0.01540 | 0.05645 | 73.93 | 308.54 |
| 4 | basement membrane (GO:0005604) | 0.02570 | 0.07068 | 43.44 | 159.04 |
| 5 | cilium (GO:0005929) | 0.1124 | 0.2473 | 9.30 | 20.33 |

Aggressive behavior ([HP:0000718](https://hpo.jax.org/app/browse/term/HP:0000718))

KEGG Pathway

| **Index** | **Name** | **P-value** | **Adjusted p-value** | **Odds Ratio** | **Combined score** |
| --- | --- | --- | --- | --- | --- |
| 1 | Thyroid hormone signaling pathway | 0.00002513 | 0.0008041 | 72.17 | 764.43 |
| 2 | Notch signaling pathway | 0.0003792 | 0.001952 | 87.43 | 688.70 |
| 3 | Long-term potentiation | 0.0004889 | 0.001952 | 76.63 | 584.21 |
| 4 | Renal cell carcinoma | 0.0005185 | 0.001952 | 74.34 | 562.35 |
| 5 | Adherens junction | 0.0005489 | 0.001952 | 72.18 | 541.87 |

GO: Biological Process

| **Index** | **Name** | **P-value** | **Adjusted p-value** | **Odds Ratio** | **Combined score** |
| --- | --- | --- | --- | --- | --- |
| 1 | peptidyl-lysine acetylation (GO:0018394) | 0.00001010 | 0.0001479 | 624.44 | 7182.69 |
| 2 | chromatin-mediated maintenance of transcription (GO:0048096) | 0.00001235 | 0.0001687 | 555.03 | 6273.06 |
| 3 | chromatin organization involved in regulation of transcription (GO:0034401) | 0.00001750 | 0.0002242 | 454.07 | 4973.59 |
| 4 | chromatin remodeling (GO:0006338) | 8.107e-10 | 1.662e-7 | 202.98 | 4248.99 |
| 5 | N-terminal protein amino acid acetylation (GO:0006474) | 0.00002690 | 0.0003064 | 356.71 | 3753.85 |

GO: Molecular Function

| **Index** | **Name** | **P-value** | **Adjusted p-value** | **Odds Ratio** | **Combined score** |
| --- | --- | --- | --- | --- | --- |
| 1 | lysine N-acetyltransferase activity, acting on acetyl phosphate as donor (GO:0004468) | 0.002498 | 0.009989 | 555.17 | 3326.77 |
| 2 | histone demethylase activity (H3-trimethyl-K4 specific) (GO:0034647) | 0.002498 | 0.009989 | 555.17 | 3326.77 |
| 3 | STAT family protein binding (GO:0097677) | 0.002997 | 0.009989 | 444.11 | 2580.41 |
| 4 | DNA-methyltransferase activity (GO:0009008) | 0.002997 | 0.009989 | 444.11 | 2580.41 |
| 5 | S-methyltransferase activity (GO:0008172) | 0.003495 | 0.01075 | 370.07 | 2093.27 |

GO: Cellular Component

| **Index** | **Name** | **P-value** | **Adjusted p-value** | **Odds Ratio** | **Combined score** |
| --- | --- | --- | --- | --- | --- |
| 1 | nBAF complex (GO:0071565) | 3.267e-8 | 3.267e-7 | 778.40 | 13417.28 |
| 2 | SWI/SNF complex (GO:0016514) | 8.685e-8 | 4.342e-7 | 535.02 | 8698.92 |
| 3 | npBAF complex (GO:0071564) | 0.00001235 | 0.00004115 | 555.03 | 6273.06 |
| 4 | NURF complex (GO:0016589) | 0.002498 | 0.004163 | 555.17 | 3326.77 |
| 5 | ISWI-type complex (GO:0031010) | 0.004990 | 0.006859 | 246.68 | 1307.49 |

Depression ([HP:0000716](https://hpo.jax.org/app/browse/term/HP:0000716))

GO: Biological Process

| **Index** | **Name** | **P-value** | **Adjusted p-value** | **Odds Ratio** | **Combined score** |
| --- | --- | --- | --- | --- | --- |
| 1 | equilibrioception (GO:0050957) | 5.668e-17 | 3.287e-15 | 19989.00 | 747771.16 |
| 2 | neuromuscular process controlling balance (GO:0050885) | 1.888e-14 | 2.738e-13 | 2220.11 | 70156.55 |
| 3 | sensory perception (GO:0007600) | 4.223e-14 | 4.899e-13 | 664.83 | 20473.93 |
| 4 | sensory perception of mechanical stimulus (GO:0050954) | 2.969e-15 | 7.316e-14 | 573.51 | 19184.17 |
| 5 | sensory perception of sound (GO:0007605) | 3.784e-15 | 7.316e-14 | 552.94 | 18362.12 |

GO: Molecular Function

| **Index** | **Name** | **P-value** | **Adjusted p-value** | **Odds Ratio** | **Combined score** |
| --- | --- | --- | --- | --- | --- |
| 1 | myosin binding (GO:0017022) | 0.02766 | 0.05531 | 40.27 | 144.50 |
| 2 | calcium ion binding (GO:0005509) | 0.01239 | 0.05531 | 14.19 | 62.33 |
| 3 | magnesium ion binding (GO:0000287) | 0.07066 | 0.1060 | 15.21 | 40.30 |
| 4 | metal ion binding (GO:0046872) | 0.02616 | 0.05531 | 9.45 | 34.45 |
| 5 | cadherin binding (GO:0045296) | 0.1499 | 0.1798 | 6.81 | 12.92 |

GO: Cellular Component

| **Index** | **Name** | **P-value** | **Adjusted p-value** | **Odds Ratio** | **Combined score** |
| --- | --- | --- | --- | --- | --- |
| 1 | microvillus (GO:0005902) | 0.0003539 | 0.003893 | 90.61 | 720.07 |
| 2 | actin-based cell projection (GO:0098858) | 0.0007493 | 0.004121 | 61.45 | 442.20 |
| 3 | catenin complex (GO:0016342) | 0.01540 | 0.05645 | 73.93 | 308.54 |
| 4 | basement membrane (GO:0005604) | 0.02570 | 0.07068 | 43.44 | 159.04 |
| 5 | cilium (GO:0005929) | 0.1124 | 0.2473 | 9.30 | 20.33 |

Emotional lability ([HP:0000712](https://hpo.jax.org/app/browse/term/HP:0000712))

KEGG Pathway

| **Index** | **Name** | **P-value** | **Adjusted p-value** | **Odds Ratio** | **Combined score** |
| --- | --- | --- | --- | --- | --- |
| 1 | Thermogenesis | 3.007e-12 | 5.412e-11 | 204.97 | 5437.89 |
| 2 | Oxidative phosphorylation | 2.958e-9 | 2.662e-8 | 155.17 | 3047.38 |
| 3 | Diabetic cardiomyopathy | 2.479e-8 | 1.488e-7 | 99.96 | 1750.56 |
| 4 | Retrograde endocannabinoid signaling | 5.840e-7 | 0.000002530 | 91.88 | 1318.78 |
| 5 | Non-alcoholic fatty liver disease | 7.027e-7 | 0.000002530 | 87.59 | 1240.99 |

GO: Biological Process

| **Index** | **Name** | **P-value** | **Adjusted p-value** | **Odds Ratio** | **Combined score** |
| --- | --- | --- | --- | --- | --- |
| 1 | NADH dehydrogenase complex assembly (GO:0010257) | 9.480e-14 | 1.659e-12 | 575.13 | 17246.54 |
| 2 | mitochondrial respiratory chain complex I assembly (GO:0032981) | 9.480e-14 | 1.659e-12 | 575.13 | 17246.54 |
| 3 | mitochondrial respiratory chain complex assembly (GO:0033108) | 1.450e-12 | 1.692e-11 | 355.46 | 9689.66 |
| 4 | mitochondrial electron transport, NADH to ubiquinone (GO:0006120) | 2.570e-9 | 1.499e-8 | 380.10 | 7518.11 |
| 5 | aerobic electron transport chain (GO:0019646) | 1.129e-10 | 8.497e-10 | 306.54 | 7021.22 |

GO: Molecular Function

| **Index** | **Name** | **P-value** | **Adjusted p-value** | **Odds Ratio** | **Combined score** |
| --- | --- | --- | --- | --- | --- |
| 1 | NADH dehydrogenase activity (GO:0003954) | 0.000004719 | 0.00001298 | 999.25 | 12254.80 |
| 2 | oxidoreduction-driven active transmembrane transporter activity (GO:0015453) | 4.283e-11 | 4.712e-10 | 376.17 | 8980.56 |
| 3 | NADH dehydrogenase (quinone) activity (GO:0050136) | 1.637e-9 | 6.003e-9 | 429.23 | 8683.34 |
| 4 | NADH dehydrogenase (ubiquinone) activity (GO:0008137) | 1.637e-9 | 6.003e-9 | 429.23 | 8683.34 |
| 5 | oxidoreductase activity, acting on the aldehyde or oxo group of donors, disulfide as acceptor (GO:0016624) | 0.003495 | 0.007322 | 370.07 | 2093.27 |

GO: Cellular Component

| **Index** | **Name** | **P-value** | **Adjusted p-value** | **Odds Ratio** | **Combined score** |
| --- | --- | --- | --- | --- | --- |
| 1 | mitochondrial respiratory chain complex I (GO:0005747) | 3.494e-9 | 2.621e-8 | 350.04 | 6815.94 |
| 2 | respiratory chain complex I (GO:0045271) | 3.494e-9 | 2.621e-8 | 350.04 | 6815.94 |
| 3 | mitochondrial envelope (GO:0005740) | 3.161e-7 | 8.832e-7 | 107.68 | 1611.67 |
| 4 | mitochondrial membrane (GO:0031966) | 3.123e-8 | 1.562e-7 | 63.26 | 1093.29 |
| 5 | mitochondrial inner membrane (GO:0005743) | 2.711e-7 | 8.832e-7 | 60.89 | 920.69 |

Mania ([HP:0100754](https://hpo.jax.org/app/browse/term/HP:0100754))

KEGG Pathway

| **Index** | **Name** | **P-value** | **Adjusted p-value** | **Odds Ratio** | **Combined score** |
| --- | --- | --- | --- | --- | --- |
| 1 | Long-term potentiation | 0.0004889 | 0.02224 | 76.63 | 584.21 |
| 2 | Morphine addiction | 0.0008998 | 0.02224 | 55.90 | 392.06 |
| 3 | Progesterone-mediated oocyte maturation | 0.001085 | 0.02224 | 50.74 | 346.39 |
| 4 | Platelet activation | 0.001661 | 0.02224 | 40.71 | 260.58 |
| 5 | Tyrosine metabolism | 0.01786 | 0.08975 | 63.35 | 255.00 |

GO: Biological Process

| **Index** | **Name** | **P-value** | **Adjusted p-value** | **Odds Ratio** | **Combined score** |
| --- | --- | --- | --- | --- | --- |
| 1 | activation of protein kinase A activity (GO:0034199) | 0.00003048 | 0.002871 | 332.92 | 3461.84 |
| 2 | negative regulation of cGMP-mediated signaling (GO:0010754) | 0.002498 | 0.02483 | 555.17 | 3326.77 |
| 3 | coronary artery morphogenesis (GO:0060982) | 0.002498 | 0.02483 | 555.17 | 3326.77 |
| 4 | middle ear morphogenesis (GO:0042474) | 0.002498 | 0.02483 | 555.17 | 3326.77 |
| 5 | embryonic viscerocranium morphogenesis (GO:0048703) | 0.002498 | 0.02483 | 555.17 | 3326.77 |

GO: Molecular Function

| **Index** | **Name** | **P-value** | **Adjusted p-value** | **Odds Ratio** | **Combined score** |
| --- | --- | --- | --- | --- | --- |
| 1 | cAMP-dependent protein kinase activity (GO:0004691) | 0.002498 | 0.02517 | 555.17 | 3326.77 |
| 2 | cyclic nucleotide-dependent protein kinase activity (GO:0004690) | 0.002997 | 0.02517 | 444.11 | 2580.41 |
| 3 | ribosomal protein S6 kinase activity (GO:0004711) | 0.003495 | 0.02517 | 370.07 | 2093.27 |
| 4 | cAMP-dependent protein kinase inhibitor activity (GO:0004862) | 0.003994 | 0.02517 | 317.19 | 1751.86 |
| 5 | L-aspartate transmembrane transporter activity (GO:0015183) | 0.004492 | 0.02517 | 277.53 | 1500.17 |

GO: Cellular Component

| **Index** | **Name** | **P-value** | **Adjusted p-value** | **Odds Ratio** | **Combined score** |
| --- | --- | --- | --- | --- | --- |
| 1 | plasma membrane raft (GO:0044853) | 0.0007315 | 0.01390 | 62.22 | 449.25 |
| 2 | calcium channel complex (GO:0034704) | 0.02228 | 0.1128 | 50.37 | 191.61 |
| 3 | membrane raft (GO:0045121) | 0.002846 | 0.02704 | 30.79 | 180.49 |
| 4 | acrosomal vesicle (GO:0001669) | 0.02375 | 0.1128 | 47.15 | 176.34 |
| 5 | cation channel complex (GO:0034703) | 0.03591 | 0.1365 | 30.74 | 102.25 |

Suicidal ideation ([HP:0031589](https://hpo.jax.org/app/browse/term/HP:0031589))

KEGG Pathway

| **Index** | **Name** | **P-value** | **Adjusted p-value** | **Odds Ratio** | **Combined score** |
| --- | --- | --- | --- | --- | --- |
| 1 | Nicotine addiction | 0.00008138 | 0.0008138 | 210.05 | 1977.93 |
| 2 | Primary bile acid biosynthesis | 0.005936 | 0.01187 | 208.09 | 1066.85 |
| 3 | Cholinergic synapse | 0.0006523 | 0.002174 | 71.65 | 525.53 |
| 4 | Neuroactive ligand-receptor interaction | 0.0001634 | 0.0008171 | 43.61 | 380.27 |
| 5 | Cholesterol metabolism | 0.01737 | 0.02895 | 67.84 | 274.94 |

GO: Biological Process

| **Index** | **Name** | **P-value** | **Adjusted p-value** | **Odds Ratio** | **Combined score** |
| --- | --- | --- | --- | --- | --- |
| 1 | synaptic transmission, cholinergic (GO:0007271) | 2.538e-8 | 0.000001954 | 936.42 | 16377.54 |
| 2 | response to acetylcholine (GO:1905144) | 0.001749 | 0.006733 | 832.88 | 5287.72 |
| 3 | response to cocaine (GO:0042220) | 0.001749 | 0.006733 | 832.88 | 5287.72 |
| 4 | vitamin D3 metabolic process (GO:0070640) | 0.001749 | 0.006733 | 832.88 | 5287.72 |
| 5 | sensory perception of pain (GO:0019233) | 0.00002198 | 0.0002821 | 420.51 | 4510.07 |

GO: Molecular Function

| **Index** | **Name** | **P-value** | **Adjusted p-value** | **Odds Ratio** | **Combined score** |
| --- | --- | --- | --- | --- | --- |
| 1 | acetylcholine-gated cation-selective channel activity (GO:0022848) | 4.326e-9 | 8.219e-8 | 1873.59 | 36082.86 |
| 2 | excitatory extracellular ligand-gated ion channel activity (GO:0005231) | 3.482e-8 | 1.915e-7 | 832.29 | 14293.04 |
| 3 | postsynaptic neurotransmitter receptor activity (GO:0098960) | 3.482e-8 | 1.915e-7 | 832.29 | 14293.04 |
| 4 | acetylcholine receptor activity (GO:0015464) | 4.031e-8 | 1.915e-7 | 788.45 | 13424.62 |
| 5 | transmitter-gated ion channel activity involved in regulation of postsynaptic membrane potential (GO:1904315) | 1.061e-7 | 4.034e-7 | 554.61 | 8906.19 |

GO: Cellular Component

| **Index** | **Name** | **P-value** | **Adjusted p-value** | **Odds Ratio** | **Combined score** |
| --- | --- | --- | --- | --- | --- |
| 1 | acetylcholine-gated channel complex (GO:0005892) | 9.539e-9 | 2.003e-7 | 1362.41 | 25160.79 |
| 2 | ion channel complex (GO:0034702) | 1.710e-7 | 0.000001795 | 467.84 | 7289.66 |
| 3 | catenin complex (GO:0016342) | 0.01080 | 0.04536 | 110.91 | 502.19 |
| 4 | neuron projection (GO:0043005) | 0.0006880 | 0.004816 | 26.37 | 191.99 |
| 5 | late endosome membrane (GO:0031902) | 0.02356 | 0.07443 | 49.57 | 185.78 |

Neuropsychiatric-headache

Headache ([HP:0002315](https://hpo.jax.org/app/browse/term/HP:0002315))

KEGG Pathway

| **Index** | **Name** | **P-value** | **Adjusted p-value** | **Odds Ratio** | **Combined score** |
| --- | --- | --- | --- | --- | --- |
| 1 | Pathways in cancer | 5.987e-14 | 7.664e-12 | 335.66 | 10219.54 |
| 2 | Gastric cancer | 3.165e-11 | 2.026e-9 | 208.19 | 5033.13 |
| 3 | Thyroid cancer | 6.931e-7 | 0.000006337 | 251.55 | 3567.44 |
| 4 | Central carbon metabolism in cancer | 2.843e-8 | 7.279e-7 | 201.25 | 3496.91 |
| 5 | Chronic myeloid leukemia | 3.973e-8 | 8.475e-7 | 184.43 | 3142.85 |

GO: Biological Process

| **Index** | **Name** | **P-value** | **Adjusted p-value** | **Odds Ratio** | **Combined score** |
| --- | --- | --- | --- | --- | --- |
| 1 | negative regulation of gene silencing by miRNA (GO:0060965) | 0.00001481 | 0.001266 | 499.50 | 5554.53 |
| 2 | negative regulation of production of miRNAs involved in gene silencing by miRNA (GO:1903799) | 0.00001481 | 0.001266 | 499.50 | 5554.53 |
| 3 | positive regulation of histone H3-K4 methylation (GO:0051571) | 0.00002354 | 0.001266 | 384.17 | 4094.01 |
| 4 | response to muscle stretch (GO:0035994) | 0.00002354 | 0.001266 | 384.17 | 4094.01 |
| 5 | regulation of phospholipase C activity (GO:1900274) | 0.00002354 | 0.001266 | 384.17 | 4094.01 |

GO: Molecular Function

| **Index** | **Name** | **P-value** | **Adjusted p-value** | **Odds Ratio** | **Combined score** |
| --- | --- | --- | --- | --- | --- |
| 1 | transcription coactivator binding (GO:0001223) | 1.021e-7 | 0.000003371 | 503.52 | 8105.11 |
| 2 | transcription coregulator binding (GO:0001221) | 9.117e-9 | 6.017e-7 | 271.31 | 5022.72 |
| 3 | I-SMAD binding (GO:0070411) | 0.00001750 | 0.0003850 | 454.07 | 4973.59 |
| 4 | RNA polymerase II general transcription initiation factor binding (GO:0001091) | 0.002498 | 0.01374 | 555.17 | 3326.77 |
| 5 | RNA-directed DNA polymerase activity (GO:0003964) | 0.002997 | 0.01413 | 444.11 | 2580.41 |

GO: Cellular Component

| **Index** | **Name** | **P-value** | **Adjusted p-value** | **Odds Ratio** | **Combined score** |
| --- | --- | --- | --- | --- | --- |
| 1 | phosphatidylinositol 3-kinase complex, class I (GO:0097651) | 0.002498 | 0.03889 | 555.17 | 3326.77 |
| 2 | transferase complex, transferring phosphorus-containing groups (GO:0061695) | 0.004492 | 0.03889 | 277.53 | 1500.17 |
| 3 | Cul4A-RING E3 ubiquitin ligase complex (GO:0031464) | 0.005488 | 0.03889 | 222.00 | 1155.57 |
| 4 | beta-catenin-TCF complex (GO:1990907) | 0.006482 | 0.03889 | 184.98 | 932.06 |
| 5 | intercalated disc (GO:0014704) | 0.01540 | 0.04602 | 73.93 | 308.54 |

Migraine ([HP:0002076](https://hpo.jax.org/app/browse/term/HP:0002076))

KEGG Pathway

| **Index** | **Name** | **P-value** | **Adjusted p-value** | **Odds Ratio** | **Combined score** |
| --- | --- | --- | --- | --- | --- |
| 1 | Prolactin signaling pathway | 2.843e-8 | 0.000001801 | 201.25 | 3496.91 |
| 2 | Chronic myeloid leukemia | 3.973e-8 | 0.000001801 | 184.43 | 3142.85 |
| 3 | Pancreatic cancer | 3.973e-8 | 0.000001801 | 184.43 | 3142.85 |
| 4 | AGE-RAGE signaling pathway in diabetic complications | 1.207e-7 | 0.000002998 | 138.15 | 2200.74 |
| 5 | Thyroid hormone signaling pathway | 2.602e-7 | 0.000005539 | 113.24 | 1716.86 |

GO: Biological Process

| **Index** | **Name** | **P-value** | **Adjusted p-value** | **Odds Ratio** | **Combined score** |
| --- | --- | --- | --- | --- | --- |
| 1 | detection of stimulus involved in sensory perception of pain (GO:0062149) | 0.000004719 | 0.001949 | 999.25 | 12254.80 |
| 2 | detection of mechanical stimulus involved in sensory perception (GO:0050974) | 0.00002354 | 0.004270 | 384.17 | 4094.01 |
| 3 | positive regulation of cell proliferation involved in heart morphogenesis (GO:2000138) | 0.002498 | 0.02406 | 555.17 | 3326.77 |
| 4 | positive regulation of histone H3-K9 acetylation (GO:2000617) | 0.002498 | 0.02406 | 555.17 | 3326.77 |
| 5 | cardiac endothelial cell differentiation (GO:0003348) | 0.002498 | 0.02406 | 555.17 | 3326.77 |

GO: Molecular Function

| **Index** | **Name** | **P-value** | **Adjusted p-value** | **Odds Ratio** | **Combined score** |
| --- | --- | --- | --- | --- | --- |
| 1 | transcription coactivator binding (GO:0001223) | 1.021e-7 | 0.000007457 | 503.52 | 8105.11 |
| 2 | RNA polymerase II general transcription initiation factor binding (GO:0001091) | 0.002498 | 0.02602 | 555.17 | 3326.77 |
| 3 | glutamate-gated calcium ion channel activity (GO:0022849) | 0.002498 | 0.02602 | 555.17 | 3326.77 |
| 4 | transcription coregulator binding (GO:0001221) | 0.000002081 | 0.00007596 | 170.91 | 2236.02 |
| 5 | 1-phosphatidylinositol-4-phosphate 3-kinase activity (GO:0035005) | 0.003495 | 0.02602 | 370.07 | 2093.27 |

GO: Cellular Component

| **Index** | **Name** | **P-value** | **Adjusted p-value** | **Odds Ratio** | **Combined score** |
| --- | --- | --- | --- | --- | --- |
| 1 | phosphatidylinositol 3-kinase complex, class I (GO:0097651) | 0.002498 | 0.03039 | 555.17 | 3326.77 |
| 2 | NMDA selective glutamate receptor complex (GO:0017146) | 0.003994 | 0.03039 | 317.19 | 1751.86 |
| 3 | voltage-gated sodium channel complex (GO:0001518) | 0.008469 | 0.03949 | 138.71 | 661.82 |
| 4 | postsynaptic density membrane (GO:0098839) | 0.009957 | 0.03949 | 116.79 | 538.34 |
| 5 | postsynaptic specialization membrane (GO:0099634) | 0.01045 | 0.03949 | 110.94 | 506.01 |

Memory impairment ([HP:0002354](https://hpo.jax.org/app/browse/term/HP:0002354))

KEGG Pathway

| **Index** | **Name** | **P-value** | **Adjusted p-value** | **Odds Ratio** | **Combined score** |
| --- | --- | --- | --- | --- | --- |
| 1 | Aldosterone-regulated sodium reabsorption | 6.931e-7 | 0.00001326 | 251.55 | 3567.44 |
| 2 | VEGF signaling pathway | 0.000002883 | 0.00003960 | 152.56 | 1946.09 |
| 3 | Cholinergic synapse | 1.976e-7 | 0.000007963 | 121.60 | 1877.08 |
| 4 | Growth hormone synthesis, secretion and action | 2.434e-7 | 0.000007963 | 115.22 | 1754.62 |
| 5 | GnRH secretion | 0.000003690 | 0.00004313 | 140.02 | 1751.57 |

GO: Biological Process

| **Index** | **Name** | **P-value** | **Adjusted p-value** | **Odds Ratio** | **Combined score** |
| --- | --- | --- | --- | --- | --- |
| 1 | cellular response to heat (GO:0034605) | 6.371e-7 | 0.0002427 | 259.18 | 3697.58 |
| 2 | activation of protein kinase A activity (GO:0034199) | 0.00003048 | 0.004313 | 332.92 | 3461.84 |
| 3 | negative regulation of tubulin deacetylation (GO:1904428) | 0.002498 | 0.02410 | 555.17 | 3326.77 |
| 4 | positive regulation of protein localization to synapse (GO:1902474) | 0.002498 | 0.02410 | 555.17 | 3326.77 |
| 5 | response to morphine (GO:0043278) | 0.002498 | 0.02410 | 555.17 | 3326.77 |

GO: Molecular Function

| **Index** | **Name** | **P-value** | **Adjusted p-value** | **Odds Ratio** | **Combined score** |
| --- | --- | --- | --- | --- | --- |
| 1 | cAMP-dependent protein kinase activity (GO:0004691) | 0.002498 | 0.02136 | 555.17 | 3326.77 |
| 2 | cyclic nucleotide-dependent protein kinase activity (GO:0004690) | 0.002997 | 0.02136 | 444.11 | 2580.41 |
| 3 | 1-phosphatidylinositol-4-phosphate 3-kinase activity (GO:0035005) | 0.003495 | 0.02136 | 370.07 | 2093.27 |
| 4 | minor groove of adenine-thymine-rich DNA binding (GO:0003680) | 0.003495 | 0.02136 | 370.07 | 2093.27 |
| 5 | protein serine/threonine/tyrosine kinase activity (GO:0004712) | 0.00008446 | 0.005068 | 191.96 | 1800.45 |

GO: Cellular Component

| **Index** | **Name** | **P-value** | **Adjusted p-value** | **Odds Ratio** | **Combined score** |
| --- | --- | --- | --- | --- | --- |
| 1 | nuclear inclusion body (GO:0042405) | 0.00001235 | 0.0004197 | 555.03 | 6273.06 |
| 2 | phosphatidylinositol 3-kinase complex, class I (GO:0097651) | 0.002498 | 0.01213 | 555.17 | 3326.77 |
| 3 | neurofibrillary tangle (GO:0097418) | 0.002498 | 0.01213 | 555.17 | 3326.77 |
| 4 | glial cell projection (GO:0097386) | 0.006979 | 0.02637 | 170.74 | 847.70 |
| 5 | membrane raft (GO:0045121) | 0.00006115 | 0.001040 | 53.12 | 515.34 |
