## Supplementary Table 1-3 for "Long COVID: G Protein-Coupled Receptors (GPCRs) responsible for persistent post-COVID symptoms": Supplementary File 2.10.2-Neuropsychiatric.docx

Neuropsychiatric-sleep

Insomnia ([HP:0100785](https://hpo.jax.org/app/browse/term/HP:0100785))

KEGG Pathway

| **Index** | **Name** | **P-value** | **Adjusted p-value** | **Odds Ratio** | **Combined score** |
| --- | --- | --- | --- | --- | --- |
| 1 | Bile secretion | 0.00001034 | 0.00008268 | 98.04 | 1125.55 |
| 2 | ABC transporters | 0.0002202 | 0.0008809 | 115.97 | 976.58 |
| 3 | Arginine biosynthesis | 0.01095 | 0.02684 | 105.66 | 476.99 |
| 4 | Collecting duct acid secretion | 0.01342 | 0.02684 | 85.32 | 367.79 |
| 5 | Porphyrin and chlorophyll metabolism | 0.02130 | 0.03297 | 52.77 | 203.13 |

GO: Biological Process

| **Index** | **Name** | **P-value** | **Adjusted p-value** | **Odds Ratio** | **Combined score** |
| --- | --- | --- | --- | --- | --- |
| 1 | regulation of bile acid metabolic process (GO:1904251) | 0.000004719 | 0.0001850 | 999.25 | 12254.80 |
| 2 | bile acid and bile salt transport (GO:0015721) | 2.929e-7 | 0.00002871 | 342.26 | 5148.70 |
| 3 | monocarboxylic acid transport (GO:0015718) | 1.229e-8 | 0.000002408 | 250.78 | 4567.88 |
| 4 | negative regulation of cellular respiration (GO:1901856) | 0.002498 | 0.02331 | 555.17 | 3326.77 |
| 5 | regulation of microvillus assembly (GO:0032534) | 0.002498 | 0.02331 | 555.17 | 3326.77 |

GO: Molecular Function

| **Index** | **Name** | **P-value** | **Adjusted p-value** | **Odds Ratio** | **Combined score** |
| --- | --- | --- | --- | --- | --- |
| 1 | glycerophospholipid flippase activity (GO:0140333) | 0.00001481 | 0.0006813 | 499.50 | 5554.53 |
| 2 | phosphatidylcholine flippase activity (GO:0140345) | 0.002498 | 0.01435 | 555.17 | 3326.77 |
| 3 | phosphatidylcholine floppase activity (GO:0090554) | 0.002498 | 0.01435 | 555.17 | 3326.77 |
| 4 | anion:sodium symporter activity (GO:0015373) | 0.002498 | 0.01435 | 555.17 | 3326.77 |
| 5 | phosphatidylserine flippase activity (GO:0140346) | 0.002498 | 0.01435 | 555.17 | 3326.77 |

GO: Cellular Component

| **Index** | **Name** | **P-value** | **Adjusted p-value** | **Odds Ratio** | **Combined score** |
| --- | --- | --- | --- | --- | --- |
| 1 | mitochondrial intermembrane space (GO:0005758) | 0.02863 | 0.1577 | 38.86 | 138.07 |
| 2 | recycling endosome membrane (GO:0055038) | 0.02863 | 0.1577 | 38.86 | 138.07 |
| 3 | organelle envelope lumen (GO:0031970) | 0.03155 | 0.1577 | 35.14 | 121.47 |
| 4 | integral component of plasma membrane (GO:0005887) | 0.0003728 | 0.007457 | 12.80 | 101.01 |
| 5 | mitochondrial envelope (GO:0005740) | 0.06173 | 0.2086 | 17.52 | 48.78 |

Restless legs ([HP:0012452](https://hpo.jax.org/app/browse/term/HP:0012452))

KEGG Pathway

| **Index** | **Name** | **P-value** | **Adjusted p-value** | **Odds Ratio** | **Combined score** |
| --- | --- | --- | --- | --- | --- |
| 1 | Pathways of neurodegeneration | 0.0001285 | 0.0007708 | 62.04 | 555.90 |
| 2 | Parkinson disease | 0.001506 | 0.004518 | 53.30 | 346.36 |
| 3 | Mitophagy | 0.01689 | 0.03377 | 74.36 | 303.47 |
| 4 | NOD-like receptor signaling pathway | 0.04444 | 0.06666 | 27.52 | 85.69 |
| 5 | Endocytosis | 0.06144 | 0.07373 | 19.67 | 54.86 |

GO: Biological Process

| **Index** | **Name** | **P-value** | **Adjusted p-value** | **Odds Ratio** | **Combined score** |
| --- | --- | --- | --- | --- | --- |
| 1 | positive regulation of vascular associated smooth muscle cell apoptotic process (GO:1905461) | 0.001249 | 0.01820 | 1249.44 | 8352.53 |
| 2 | regulation of vascular associated smooth muscle cell apoptotic process (GO:1905459) | 0.001749 | 0.01820 | 832.88 | 5287.72 |
| 3 | positive regulation of mitophagy in response to mitochondrial depolarization (GO:0098779) | 0.001749 | 0.01820 | 832.88 | 5287.72 |
| 4 | intermediate filament bundle assembly (GO:0045110) | 0.001749 | 0.01820 | 832.88 | 5287.72 |
| 5 | positive regulation of smooth muscle cell apoptotic process (GO:0034393) | 0.002248 | 0.01893 | 624.59 | 3808.55 |

GO: Molecular Function

| **Index** | **Name** | **P-value** | **Adjusted p-value** | **Odds Ratio** | **Combined score** |
| --- | --- | --- | --- | --- | --- |
| 1 | alpha-2A adrenergic receptor binding (GO:0031694) | 0.001249 | 0.008347 | 1249.44 | 8352.53 |
| 2 | adrenergic receptor binding (GO:0031690) | 0.004243 | 0.01379 | 312.17 | 1705.22 |
| 3 | ubiquitin protein ligase binding (GO:0031625) | 0.001703 | 0.008347 | 50.02 | 318.87 |
| 4 | ubiquitin-like protein ligase binding (GO:0044389) | 0.001926 | 0.008347 | 46.94 | 293.48 |
| 5 | ubiquitin binding (GO:0043130) | 0.01837 | 0.03632 | 68.23 | 272.72 |

GO: Cellular Component

| **Index** | **Name** | **P-value** | **Adjusted p-value** | **Odds Ratio** | **Combined score** |
| --- | --- | --- | --- | --- | --- |
| 1 | intrinsic component of mitochondrial membrane (GO:0098573) | 0.002997 | 0.04016 | 454.18 | 2638.91 |
| 2 | intrinsic component of mitochondrial outer membrane (GO:0031306) | 0.005737 | 0.04016 | 226.97 | 1171.32 |
| 3 | mitochondrial outer membrane (GO:0005741) | 0.03111 | 0.08167 | 39.74 | 137.91 |
| 4 | asymmetric synapse (GO:0032279) | 0.03281 | 0.08167 | 37.62 | 128.54 |
| 5 | postsynaptic density (GO:0014069) | 0.03403 | 0.08167 | 36.24 | 122.50 |

Sleep apnea ([HP:0010535](https://hpo.jax.org/app/browse/term/HP:0010535))

KEGG Pathway

| **Index** | **Name** | **P-value** | **Adjusted p-value** | **Odds Ratio** | **Combined score** |
| --- | --- | --- | --- | --- | --- |
| 1 | Thyroid cancer | 6.931e-7 | 0.00001020 | 251.55 | 3567.44 |
| 2 | Prostate cancer | 1.067e-7 | 0.000004315 | 142.63 | 2289.62 |
| 3 | Melanogenesis | 1.257e-7 | 0.000004315 | 136.72 | 2172.45 |
| 4 | Thyroid hormone signaling pathway | 2.602e-7 | 0.000005842 | 113.24 | 1716.86 |
| 5 | Long-term potentiation | 0.000004240 | 0.00003970 | 133.43 | 1650.70 |

GO: Biological Process

| **Index** | **Name** | **P-value** | **Adjusted p-value** | **Odds Ratio** | **Combined score** |
| --- | --- | --- | --- | --- | --- |
| 1 | beta-catenin-TCF complex assembly (GO:1904837) | 7.437e-10 | 3.325e-7 | 532.40 | 11190.69 |
| 2 | peptidyl-lysine acetylation (GO:0018394) | 0.00001010 | 0.0004106 | 624.44 | 7182.69 |
| 3 | negative regulation of cellular senescence (GO:2000773) | 0.00001750 | 0.0006017 | 454.07 | 4973.59 |
| 4 | N-terminal protein amino acid acetylation (GO:0006474) | 0.00002690 | 0.0007073 | 356.71 | 3753.85 |
| 5 | embryonic limb morphogenesis (GO:0030326) | 6.371e-7 | 0.00005696 | 259.18 | 3697.58 |

GO: Molecular Function

| **Index** | **Name** | **P-value** | **Adjusted p-value** | **Odds Ratio** | **Combined score** |
| --- | --- | --- | --- | --- | --- |
| 1 | lysine N-acetyltransferase activity, acting on acetyl phosphate as donor (GO:0004468) | 0.002498 | 0.01544 | 555.17 | 3326.77 |
| 2 | RNA-directed DNA polymerase activity (GO:0003964) | 0.002997 | 0.01544 | 444.11 | 2580.41 |
| 3 | STAT family protein binding (GO:0097677) | 0.002997 | 0.01544 | 444.11 | 2580.41 |
| 4 | telomerase activity (GO:0003720) | 0.002997 | 0.01544 | 444.11 | 2580.41 |
| 5 | bHLH transcription factor binding (GO:0043425) | 0.00005170 | 0.001484 | 249.63 | 2463.82 |

GO: Cellular Component

| **Index** | **Name** | **P-value** | **Adjusted p-value** | **Odds Ratio** | **Combined score** |
| --- | --- | --- | --- | --- | --- |
| 1 | transferase complex, transferring phosphorus-containing groups (GO:0061695) | 0.004492 | 0.03294 | 277.53 | 1500.17 |
| 2 | beta-catenin-TCF complex (GO:1990907) | 0.006482 | 0.03565 | 184.98 | 932.06 |
| 3 | catenin complex (GO:0016342) | 0.01540 | 0.06774 | 73.93 | 308.54 |
| 4 | nucleus (GO:0005634) | 0.0001836 | 0.004038 | 13.86 | 119.27 |
| 5 | glutamatergic synapse (GO:0098978) | 0.03398 | 0.1068 | 32.55 | 110.09 |

Sleep disturbance ([HP:0002360](https://hpo.jax.org/app/browse/term/HP:0002360))

KEGG Pathway

| **Index** | **Name** | **P-value** | **Adjusted p-value** | **Odds Ratio** | **Combined score** |
| --- | --- | --- | --- | --- | --- |
| 1 | Nicotine addiction | 9.018e-15 | 1.563e-13 | 880.41 | 28472.15 |
| 2 | Amphetamine addiction | 5.059e-16 | 2.631e-14 | 749.98 | 26414.43 |
| 3 | Circadian entrainment | 6.002e-15 | 1.560e-13 | 515.93 | 16894.89 |
| 4 | Long-term potentiation | 9.011e-11 | 7.810e-10 | 321.42 | 7434.41 |
| 5 | cAMP signaling pathway | 1.815e-12 | 2.359e-11 | 220.84 | 5970.43 |

GO: Biological Process

| **Index** | **Name** | **P-value** | **Adjusted p-value** | **Odds Ratio** | **Combined score** |
| --- | --- | --- | --- | --- | --- |
| 1 | regulation of NMDA receptor activity (GO:2000310) | 1.586e-18 | 2.394e-16 | 1863.40 | 76372.57 |
| 2 | excitatory chemical synaptic transmission (GO:0098976) | 3.147e-9 | 5.939e-8 | 2141.36 | 41921.23 |
| 3 | regulation of neurotransmitter receptor activity (GO:0099601) | 1.412e-16 | 1.066e-14 | 912.24 | 33293.64 |
| 4 | regulation of cation channel activity (GO:2001257) | 3.222e-15 | 1.622e-13 | 566.49 | 18902.93 |
| 5 | glutamate receptor signaling pathway (GO:0007215) | 4.093e-12 | 1.296e-10 | 623.69 | 16354.24 |

GO: Molecular Function

| **Index** | **Name** | **P-value** | **Adjusted p-value** | **Odds Ratio** | **Combined score** |
| --- | --- | --- | --- | --- | --- |
| 1 | glutamate-gated calcium ion channel activity (GO:0022849) | 8.995e-10 | 6.168e-9 | 4283.14 | 89214.48 |
| 2 | ionotropic glutamate receptor activity (GO:0004970) | 5.833e-14 | 2.800e-12 | 1664.83 | 50731.79 |
| 3 | NMDA glutamate receptor activity (GO:0004972) | 5.033e-9 | 3.020e-8 | 1713.00 | 32730.63 |
| 4 | ligand-gated channel activity (GO:0022834) | 1.340e-12 | 2.556e-11 | 798.60 | 21832.48 |
| 5 | ligand-gated ion channel activity (GO:0015276) | 1.597e-12 | 2.556e-11 | 767.85 | 20856.82 |

GO: Cellular Component

| **Index** | **Name** | **P-value** | **Adjusted p-value** | **Odds Ratio** | **Combined score** |
| --- | --- | --- | --- | --- | --- |
| 1 | NMDA selective glutamate receptor complex (GO:0017146) | 5.033e-9 | 4.152e-8 | 1713.00 | 32730.63 |
| 2 | ionotropic glutamate receptor complex (GO:0008328) | 1.893e-12 | 3.123e-11 | 739.37 | 19957.87 |
| 3 | cation channel complex (GO:0034703) | 3.977e-13 | 1.312e-11 | 446.04 | 12735.72 |
| 4 | synaptic membrane (GO:0097060) | 2.326e-7 | 0.000001535 | 372.06 | 5682.76 |
| 5 | AMPA glutamate receptor complex (GO:0032281) | 0.00004255 | 0.0001080 | 277.39 | 2791.90 |

Neuropsychiatric-smell-taste

Anosmia ([HP:0000458](https://hpo.jax.org/app/browse/term/HP:0000458))

KEGG Pathway

| **Index** | **Name** | **P-value** | **Adjusted p-value** | **Odds Ratio** | **Combined score** |
| --- | --- | --- | --- | --- | --- |
| 1 | GnRH secretion | 0.000003690 | 0.00005167 | 140.02 | 1751.57 |
| 2 | Neuroactive ligand-receptor interaction | 0.00001608 | 0.0001125 | 38.88 | 429.14 |
| 3 | Melanoma | 0.03543 | 0.1448 | 31.17 | 104.12 |
| 4 | GnRH signaling pathway | 0.04555 | 0.1448 | 24.03 | 74.23 |
| 5 | Breast cancer | 0.07113 | 0.1448 | 15.10 | 39.92 |

GO: Biological Process

| **Index** | **Name** | **P-value** | **Adjusted p-value** | **Odds Ratio** | **Combined score** |
| --- | --- | --- | --- | --- | --- |
| 1 | neuroepithelial cell differentiation (GO:0060563) | 0.002997 | 0.03670 | 444.11 | 2580.41 |
| 2 | negative regulation of striated muscle tissue development (GO:0045843) | 0.003495 | 0.03670 | 370.07 | 2093.27 |
| 3 | regulation of growth hormone secretion (GO:0060123) | 0.003495 | 0.03670 | 370.07 | 2093.27 |
| 4 | nose development (GO:0043584) | 0.003495 | 0.03670 | 370.07 | 2093.27 |
| 5 | limb development (GO:0060173) | 0.00009070 | 0.007618 | 184.84 | 1720.52 |

GO: Molecular Function

| **Index** | **Name** | **P-value** | **Adjusted p-value** | **Odds Ratio** | **Combined score** |
| --- | --- | --- | --- | --- | --- |
| 1 | type 1 fibroblast growth factor receptor binding (GO:0005105) | 0.002498 | 0.02123 | 555.17 | 3326.77 |
| 2 | type 2 fibroblast growth factor receptor binding (GO:0005111) | 0.002498 | 0.02123 | 555.17 | 3326.77 |
| 3 | interleukin-17 receptor activity (GO:0030368) | 0.003994 | 0.02263 | 317.19 | 1751.86 |
| 4 | peptide hormone receptor binding (GO:0051428) | 0.006979 | 0.02366 | 170.74 | 847.70 |
| 5 | neuropeptide receptor binding (GO:0071855) | 0.008966 | 0.02366 | 130.54 | 615.43 |

GO: Cellular Component

| **Index** | **Name** | **P-value** | **Adjusted p-value** | **Odds Ratio** | **Combined score** |
| --- | --- | --- | --- | --- | --- |
| 1 | cilium (GO:0005929) | 0.1124 | 0.4624 | 9.30 | 20.33 |
| 2 | Golgi membrane (GO:0000139) | 0.2125 | 0.4624 | 4.60 | 7.13 |
| 3 | integral component of plasma membrane (GO:0005887) | 0.1613 | 0.4624 | 3.19 | 5.82 |
| 4 | neuron projection (GO:0043005) | 0.2457 | 0.4624 | 3.89 | 5.46 |
| 5 | nucleolus (GO:0005730) | 0.3117 | 0.4624 | 2.92 | 3.41 |

Hyposmia ([HP:0004409](https://hpo.jax.org/app/browse/term/HP:0004409))

KEGG Pathway

| **Index** | **Name** | **P-value** | **Adjusted p-value** | **Odds Ratio** | **Combined score** |
| --- | --- | --- | --- | --- | --- |
| 1 | Melanoma | 0.0005645 | 0.006209 | 71.14 | 532.12 |
| 2 | Breast cancer | 0.002323 | 0.008327 | 34.22 | 207.51 |
| 3 | Gastric cancer | 0.002386 | 0.008327 | 33.75 | 203.77 |
| 4 | Rap1 signaling pathway | 0.004671 | 0.008327 | 23.78 | 127.59 |
| 5 | Regulation of actin cytoskeleton | 0.005024 | 0.008327 | 22.89 | 121.15 |

GO: Biological Process

| **Index** | **Name** | **P-value** | **Adjusted p-value** | **Odds Ratio** | **Combined score** |
| --- | --- | --- | --- | --- | --- |
| 1 | neuroepithelial cell differentiation (GO:0060563) | 0.002997 | 0.02194 | 444.11 | 2580.41 |
| 2 | regulation of protein localization to cilium (GO:1903564) | 0.002997 | 0.02194 | 444.11 | 2580.41 |
| 3 | negative regulation of striated muscle tissue development (GO:0045843) | 0.003495 | 0.02194 | 370.07 | 2093.27 |
| 4 | regulation of growth hormone secretion (GO:0060123) | 0.003495 | 0.02194 | 370.07 | 2093.27 |
| 5 | nose development (GO:0043584) | 0.003495 | 0.02194 | 370.07 | 2093.27 |

GO: Molecular Function

| **Index** | **Name** | **P-value** | **Adjusted p-value** | **Odds Ratio** | **Combined score** |
| --- | --- | --- | --- | --- | --- |
| 1 | type 1 fibroblast growth factor receptor binding (GO:0005105) | 0.000002248 | 0.00002361 | 1665.58 | 21661.60 |
| 2 | type 2 fibroblast growth factor receptor binding (GO:0005111) | 0.000002248 | 0.00002361 | 1665.58 | 21661.60 |
| 3 | fibroblast growth factor receptor binding (GO:0005104) | 0.00005661 | 0.0003963 | 237.73 | 2324.81 |
| 4 | interleukin-17 receptor activity (GO:0030368) | 0.003994 | 0.01398 | 317.19 | 1751.86 |
| 5 | growth factor activity (GO:0008083) | 0.0008229 | 0.004320 | 58.54 | 415.82 |

GO: Cellular Component

| **Index** | **Name** | **P-value** | **Adjusted p-value** | **Odds Ratio** | **Combined score** |
| --- | --- | --- | --- | --- | --- |
| 1 | cytoplasmic stress granule (GO:0010494) | 0.03204 | 0.3524 | 34.59 | 119.03 |
| 2 | vesicle (GO:0031982) | 0.1074 | 0.4661 | 9.76 | 21.77 |
| 3 | dendrite (GO:0030425) | 0.1271 | 0.4661 | 8.15 | 16.80 |
| 4 | Golgi membrane (GO:0000139) | 0.2125 | 0.5087 | 4.60 | 7.13 |
| 5 | nucleolus (GO:0005730) | 0.3117 | 0.5087 | 2.92 | 3.41 |

Abnormality of the sense of smell ([HP:0004408](https://hpo.jax.org/app/browse/term/HP:0004408))

KEGG Pathway

| **Index** | **Name** | **P-value** | **Adjusted p-value** | **Odds Ratio** | **Combined score** |
| --- | --- | --- | --- | --- | --- |
| 1 | GnRH secretion | 0.000003690 | 0.00005167 | 140.02 | 1751.57 |
| 2 | Neuroactive ligand-receptor interaction | 0.00001608 | 0.0001125 | 38.88 | 429.14 |
| 3 | Melanoma | 0.03543 | 0.1448 | 31.17 | 104.12 |
| 4 | GnRH signaling pathway | 0.04555 | 0.1448 | 24.03 | 74.23 |
| 5 | Breast cancer | 0.07113 | 0.1448 | 15.10 | 39.92 |

GO: Biological Process

| **Index** | **Name** | **P-value** | **Adjusted p-value** | **Odds Ratio** | **Combined score** |
| --- | --- | --- | --- | --- | --- |
| 1 | neuroepithelial cell differentiation (GO:0060563) | 0.002997 | 0.03670 | 444.11 | 2580.41 |
| 2 | negative regulation of striated muscle tissue development (GO:0045843) | 0.003495 | 0.03670 | 370.07 | 2093.27 |
| 3 | regulation of growth hormone secretion (GO:0060123) | 0.003495 | 0.03670 | 370.07 | 2093.27 |
| 4 | nose development (GO:0043584) | 0.003495 | 0.03670 | 370.07 | 2093.27 |
| 5 | limb development (GO:0060173) | 0.00009070 | 0.007618 | 184.84 | 1720.52 |

GO: Molecular Function

| **Index** | **Name** | **P-value** | **Adjusted p-value** | **Odds Ratio** | **Combined score** |
| --- | --- | --- | --- | --- | --- |
| 1 | type 1 fibroblast growth factor receptor binding (GO:0005105) | 0.002498 | 0.02123 | 555.17 | 3326.77 |
| 2 | type 2 fibroblast growth factor receptor binding (GO:0005111) | 0.002498 | 0.02123 | 555.17 | 3326.77 |
| 3 | interleukin-17 receptor activity (GO:0030368) | 0.003994 | 0.02263 | 317.19 | 1751.86 |
| 4 | peptide hormone receptor binding (GO:0051428) | 0.006979 | 0.02366 | 170.74 | 847.70 |
| 5 | neuropeptide receptor binding (GO:0071855) | 0.008966 | 0.02366 | 130.54 | 615.43 |

GO: Cellular Component

| **Index** | **Name** | **P-value** | **Adjusted p-value** | **Odds Ratio** | **Combined score** |
| --- | --- | --- | --- | --- | --- |
| 1 | cilium (GO:0005929) | 0.1124 | 0.4624 | 9.30 | 20.33 |
| 2 | Golgi membrane (GO:0000139) | 0.2125 | 0.4624 | 4.60 | 7.13 |
| 3 | integral component of plasma membrane (GO:0005887) | 0.1613 | 0.4624 | 3.19 | 5.82 |
| 4 | neuron projection (GO:0043005) | 0.2457 | 0.4624 | 3.89 | 5.46 |
| 5 | nucleolus (GO:0005730) | 0.3117 | 0.4624 | 2.92 | 3.41 |

Neuropsychiatric-speech-language

Anomic aphasia ([HP:0030784](https://hpo.jax.org/app/browse/term/HP:0030784))

KEGG Pathway

| **Index** | **Name** | **P-value** | **Adjusted p-value** | **Odds Ratio** | **Combined score** |
| --- | --- | --- | --- | --- | --- |
| 1 | Osteoclast differentiation | 0.02516 | 0.08973 | 52.57 | 193.57 |
| 2 | Parkinson disease | 0.04888 | 0.08973 | 26.54 | 80.12 |
| 3 | MAPK signaling pathway | 0.05752 | 0.08973 | 22.42 | 64.01 |
| 4 | Alzheimer disease | 0.07179 | 0.08973 | 17.78 | 46.83 |
| 5 | Pathways of neurodegeneration | 0.09168 | 0.09168 | 13.73 | 32.80 |

GO: Biological Process

| **Index** | **Name** | **P-value** | **Adjusted p-value** | **Odds Ratio** | **Combined score** |
| --- | --- | --- | --- | --- | --- |
| 1 | microglial cell activation (GO:0001774) | 0.000006921 | 0.001545 | 998.80 | 11866.74 |
| 2 | regulation of hippocampal neuron apoptotic process (GO:0110089) | 0.0009997 | 0.008525 | 1666.00 | 11508.86 |
| 3 | negative regulation of tubulin deacetylation (GO:1904428) | 0.0009997 | 0.008525 | 1666.00 | 11508.86 |
| 4 | positive regulation of protein localization to synapse (GO:1902474) | 0.0009997 | 0.008525 | 1666.00 | 11508.86 |
| 5 | chemokine (C-X-C motif) ligand 12 signaling pathway (GO:0038146) | 0.0009997 | 0.008525 | 1666.00 | 11508.86 |

GO: Molecular Function

| **Index** | **Name** | **P-value** | **Adjusted p-value** | **Odds Ratio** | **Combined score** |
| --- | --- | --- | --- | --- | --- |
| 1 | apolipoprotein A-I binding (GO:0034186) | 0.0009997 | 0.008116 | 1666.00 | 11508.86 |
| 2 | lipoprotein particle binding (GO:0071813) | 0.000008268 | 0.0002398 | 907.91 | 10625.40 |
| 3 | sulfatide binding (GO:0120146) | 0.001200 | 0.008116 | 1332.73 | 8963.74 |
| 4 | high-density lipoprotein particle binding (GO:0008035) | 0.001200 | 0.008116 | 1332.73 | 8963.74 |
| 5 | minor groove of adenine-thymine-rich DNA binding (GO:0003680) | 0.001399 | 0.008116 | 1110.56 | 7298.30 |

GO: Cellular Component

| **Index** | **Name** | **P-value** | **Adjusted p-value** | **Odds Ratio** | **Combined score** |
| --- | --- | --- | --- | --- | --- |
| 1 | neurofibrillary tangle (GO:0097418) | 0.0009997 | 0.006498 | 1666.00 | 11508.86 |
| 2 | glial cell projection (GO:0097386) | 0.002797 | 0.01212 | 512.38 | 3012.37 |
| 3 | membrane raft (GO:0045121) | 0.0003919 | 0.005094 | 123.20 | 966.45 |
| 4 | plasma membrane raft (GO:0044853) | 0.01630 | 0.05298 | 81.95 | 337.37 |
| 5 | microtubule (GO:0005874) | 0.03591 | 0.08599 | 36.49 | 121.40 |

Aphasia ([HP:0002381](https://hpo.jax.org/app/browse/term/HP:0002381))

KEGG Pathway

| **Index** | **Name** | **P-value** | **Adjusted p-value** | **Odds Ratio** | **Combined score** |
| --- | --- | --- | --- | --- | --- |
| 1 | RNA polymerase | 0.0001038 | 0.009967 | 172.08 | 1578.44 |
| 2 | Cytosolic DNA-sensing pathway | 0.0004323 | 0.01565 | 81.68 | 632.69 |
| 3 | Long-term potentiation | 0.0004889 | 0.01565 | 76.63 | 584.21 |
| 4 | Aldosterone-regulated sodium reabsorption | 0.01835 | 0.1011 | 61.59 | 246.23 |
| 5 | Thyroid cancer | 0.01835 | 0.1011 | 61.59 | 246.23 |

GO: Biological Process

| **Index** | **Name** | **P-value** | **Adjusted p-value** | **Odds Ratio** | **Combined score** |
| --- | --- | --- | --- | --- | --- |
| 1 | negative regulation of tubulin deacetylation (GO:1904428) | 0.002498 | 0.03369 | 555.17 | 3326.77 |
| 2 | positive regulation of protein localization to synapse (GO:1902474) | 0.002498 | 0.03369 | 555.17 | 3326.77 |
| 3 | cellular response to brain-derived neurotrophic factor stimulus (GO:1990416) | 0.002498 | 0.03369 | 555.17 | 3326.77 |
| 4 | intracellular distribution of mitochondria (GO:0048312) | 0.002498 | 0.03369 | 555.17 | 3326.77 |
| 5 | negative regulation of establishment of protein localization to mitochondrion (GO:1903748) | 0.002997 | 0.03369 | 444.11 | 2580.41 |

GO: Molecular Function

| **Index** | **Name** | **P-value** | **Adjusted p-value** | **Odds Ratio** | **Combined score** |
| --- | --- | --- | --- | --- | --- |
| 1 | glutamate-gated calcium ion channel activity (GO:0022849) | 0.002498 | 0.04233 | 555.17 | 3326.77 |
| 2 | minor groove of adenine-thymine-rich DNA binding (GO:0003680) | 0.003495 | 0.04233 | 370.07 | 2093.27 |
| 3 | NMDA glutamate receptor activity (GO:0004972) | 0.003994 | 0.04233 | 317.19 | 1751.86 |
| 4 | 5'-3' RNA polymerase activity (GO:0034062) | 0.00009070 | 0.002403 | 184.84 | 1720.52 |
| 5 | DNA-directed 5'-3' RNA polymerase activity (GO:0003899) | 0.00009070 | 0.002403 | 184.84 | 1720.52 |

GO: Cellular Component

| **Index** | **Name** | **P-value** | **Adjusted p-value** | **Odds Ratio** | **Combined score** |
| --- | --- | --- | --- | --- | --- |
| 1 | RNA polymerase I complex (GO:0005736) | 0.00001750 | 0.0006170 | 454.07 | 4973.59 |
| 2 | NURF complex (GO:0016589) | 0.002498 | 0.02248 | 555.17 | 3326.77 |
| 3 | neurofibrillary tangle (GO:0097418) | 0.002498 | 0.02248 | 555.17 | 3326.77 |
| 4 | RNA polymerase III complex (GO:0005666) | 0.00003428 | 0.0006170 | 312.09 | 3208.63 |
| 5 | NMDA selective glutamate receptor complex (GO:0017146) | 0.003994 | 0.02566 | 317.19 | 1751.86 |

Expressive aphasia ([HP:0002427](https://hpo.jax.org/app/browse/term/HP:0002427))

KEGG Pathway

| **Index** | **Name** | **P-value** | **Adjusted p-value** | **Odds Ratio** | **Combined score** |
| --- | --- | --- | --- | --- | --- |
| 1 | Osteoclast differentiation | 0.03135 | 0.1067 | 39.42 | 136.50 |
| 2 | mTOR signaling pathway | 0.03792 | 0.1067 | 32.42 | 106.10 |
| 3 | Parkinson disease | 0.06072 | 0.1067 | 19.91 | 55.77 |
| 4 | MAPK signaling pathway | 0.07138 | 0.1067 | 16.81 | 44.38 |
| 5 | Alzheimer disease | 0.08892 | 0.1067 | 13.33 | 32.27 |

GO: Biological Process

| **Index** | **Name** | **P-value** | **Adjusted p-value** | **Odds Ratio** | **Combined score** |
| --- | --- | --- | --- | --- | --- |
| 1 | regulation of hippocampal neuron apoptotic process (GO:0110089) | 0.001249 | 0.01102 | 1249.44 | 8352.53 |
| 2 | negative regulation of tubulin deacetylation (GO:1904428) | 0.001249 | 0.01102 | 1249.44 | 8352.53 |
| 3 | positive regulation of protein localization to synapse (GO:1902474) | 0.001249 | 0.01102 | 1249.44 | 8352.53 |
| 4 | chemokine (C-X-C motif) ligand 12 signaling pathway (GO:0038146) | 0.001249 | 0.01102 | 1249.44 | 8352.53 |
| 5 | regulation of inward rectifier potassium channel activity (GO:1901979) | 0.001249 | 0.01102 | 1249.44 | 8352.53 |

GO: Molecular Function

| **Index** | **Name** | **P-value** | **Adjusted p-value** | **Odds Ratio** | **Combined score** |
| --- | --- | --- | --- | --- | --- |
| 1 | apolipoprotein A-I binding (GO:0034186) | 0.001249 | 0.01154 | 1249.44 | 8352.53 |
| 2 | lipoprotein particle binding (GO:0071813) | 0.00001377 | 0.0004544 | 605.24 | 6774.51 |
| 3 | sulfatide binding (GO:0120146) | 0.001499 | 0.01154 | 999.50 | 6499.56 |
| 4 | high-density lipoprotein particle binding (GO:0008035) | 0.001499 | 0.01154 | 999.50 | 6499.56 |
| 5 | minor groove of adenine-thymine-rich DNA binding (GO:0003680) | 0.001749 | 0.01154 | 832.88 | 5287.72 |

GO: Cellular Component

| **Index** | **Name** | **P-value** | **Adjusted p-value** | **Odds Ratio** | **Combined score** |
| --- | --- | --- | --- | --- | --- |
| 1 | neurofibrillary tangle (GO:0097418) | 0.001249 | 0.01187 | 1249.44 | 8352.53 |
| 2 | glial cell projection (GO:0097386) | 0.003495 | 0.02214 | 384.27 | 2173.55 |
| 3 | connexin complex (GO:0005922) | 0.005239 | 0.02275 | 249.69 | 1311.24 |
| 4 | gap junction (GO:0005921) | 0.005986 | 0.02275 | 217.09 | 1111.12 |
| 5 | membrane raft (GO:0045121) | 0.0006496 | 0.01187 | 82.13 | 602.75 |

Slurred speech ([HP:0001350](https://hpo.jax.org/app/browse/term/HP:0001350))

KEGG Pathway

| **Index** | **Name** | **P-value** | **Adjusted p-value** | **Odds Ratio** | **Combined score** |
| --- | --- | --- | --- | --- | --- |
| 1 | Primary bile acid biosynthesis | 0.008469 | 0.07480 | 138.71 | 661.82 |
| 2 | Spinocerebellar ataxia | 0.00004140 | 0.001118 | 60.77 | 613.26 |
| 3 | Nicotine addiction | 0.01983 | 0.08190 | 56.84 | 222.86 |
| 4 | Cocaine addiction | 0.02424 | 0.08190 | 46.16 | 171.72 |
| 5 | Notch signaling pathway | 0.02912 | 0.08190 | 38.18 | 135.03 |

GO: Biological Process

| **Index** | **Name** | **P-value** | **Adjusted p-value** | **Odds Ratio** | **Combined score** |
| --- | --- | --- | --- | --- | --- |
| 1 | negative regulation of tubulin deacetylation (GO:1904428) | 0.002498 | 0.02880 | 555.17 | 3326.77 |
| 2 | positive regulation of protein localization to synapse (GO:1902474) | 0.002498 | 0.02880 | 555.17 | 3326.77 |
| 3 | cellular response to brain-derived neurotrophic factor stimulus (GO:1990416) | 0.002498 | 0.02880 | 555.17 | 3326.77 |
| 4 | intracellular distribution of mitochondria (GO:0048312) | 0.002498 | 0.02880 | 555.17 | 3326.77 |
| 5 | regulation of ribonucleoprotein complex localization (GO:2000197) | 0.002997 | 0.02880 | 444.11 | 2580.41 |

GO: Molecular Function

| **Index** | **Name** | **P-value** | **Adjusted p-value** | **Odds Ratio** | **Combined score** |
| --- | --- | --- | --- | --- | --- |
| 1 | glutamate-gated calcium ion channel activity (GO:0022849) | 0.002498 | 0.02618 | 555.17 | 3326.77 |
| 2 | oxidoreductase activity, acting on the CH-CH group of donors, oxygen as acceptor (GO:0016634) | 0.003495 | 0.02618 | 370.07 | 2093.27 |
| 3 | minor groove of adenine-thymine-rich DNA binding (GO:0003680) | 0.003495 | 0.02618 | 370.07 | 2093.27 |
| 4 | acyl-CoA oxidase activity (GO:0003997) | 0.003495 | 0.02618 | 370.07 | 2093.27 |
| 5 | poly(G) binding (GO:0034046) | 0.003994 | 0.02618 | 317.19 | 1751.86 |

GO: Cellular Component

| **Index** | **Name** | **P-value** | **Adjusted p-value** | **Odds Ratio** | **Combined score** |
| --- | --- | --- | --- | --- | --- |
| 1 | neurofibrillary tangle (GO:0097418) | 0.002498 | 0.01648 | 555.17 | 3326.77 |
| 2 | NMDA selective glutamate receptor complex (GO:0017146) | 0.003994 | 0.02197 | 317.19 | 1751.86 |
| 3 | nuclear inclusion body (GO:0042405) | 0.005488 | 0.02587 | 222.00 | 1155.57 |
| 4 | glial cell projection (GO:0097386) | 0.006979 | 0.02789 | 170.74 | 847.70 |
| 5 | asymmetric synapse (GO:0032279) | 0.00003334 | 0.0006142 | 65.47 | 674.94 |

Neuropsychiatric-finding

Abnormal reflex ([HP:0031826](https://hpo.jax.org/app/browse/term/HP:0031826))

KEGG Pathway

| **Index** | **Name** | **P-value** | **Adjusted p-value** | **Odds Ratio** | **Combined score** |
| --- | --- | --- | --- | --- | --- |
| 1 | Phototransduction | 2.929e-7 | 0.000001758 | 342.26 | 5148.70 |
| 2 | Retinol metabolism | 0.03349 | 0.09654 | 33.04 | 112.22 |
| 3 | Purine metabolism | 0.06267 | 0.09654 | 17.24 | 47.76 |
| 4 | Spliceosome | 0.07253 | 0.09654 | 14.80 | 38.82 |
| 5 | cGMP-PKG signaling pathway | 0.08045 | 0.09654 | 13.27 | 33.44 |

GO: Biological Process

| **Index** | **Name** | **P-value** | **Adjusted p-value** | **Odds Ratio** | **Combined score** |
| --- | --- | --- | --- | --- | --- |
| 1 | rhodopsin mediated signaling pathway (GO:0016056) | 0.00001235 | 0.0001580 | 555.03 | 6273.06 |
| 2 | regulation of rhodopsin mediated signaling pathway (GO:0022400) | 2.058e-7 | 0.000004391 | 388.99 | 5988.96 |
| 3 | phototransduction, visible light (GO:0007603) | 0.00002354 | 0.0002511 | 384.17 | 4094.01 |
| 4 | visual perception (GO:0007601) | 9.004e-8 | 0.000003140 | 149.07 | 2418.38 |
| 5 | sensory perception of light stimulus (GO:0050953) | 9.813e-8 | 0.000003140 | 145.78 | 2352.45 |

GO: Molecular Function

| **Index** | **Name** | **P-value** | **Adjusted p-value** | **Odds Ratio** | **Combined score** |
| --- | --- | --- | --- | --- | --- |
| 1 | guanylate cyclase activator activity (GO:0030250) | 0.002498 | 0.02195 | 555.17 | 3326.77 |
| 2 | NADP-retinol dehydrogenase activity (GO:0052650) | 0.002997 | 0.02195 | 444.11 | 2580.41 |
| 3 | intracellular cGMP-activated cation channel activity (GO:0005223) | 0.003495 | 0.02195 | 370.07 | 2093.27 |
| 4 | intracellular cAMP-activated cation channel activity (GO:0005222) | 0.004492 | 0.02195 | 277.53 | 1500.17 |
| 5 | leucine zipper domain binding (GO:0043522) | 0.004990 | 0.02195 | 246.68 | 1307.49 |

GO: Cellular Component

| **Index** | **Name** | **P-value** | **Adjusted p-value** | **Odds Ratio** | **Combined score** |
| --- | --- | --- | --- | --- | --- |
| 1 | spliceosomal tri-snRNP complex (GO:0097526) | 0.01589 | 0.05160 | 71.54 | 296.32 |
| 2 | U4/U6 x U5 tri-snRNP complex (GO:0046540) | 0.01589 | 0.05160 | 71.54 | 296.32 |
| 3 | U2-type precatalytic spliceosome (GO:0071005) | 0.02473 | 0.05160 | 45.22 | 167.30 |
| 4 | precatalytic spliceosome (GO:0071011) | 0.02570 | 0.05160 | 43.44 | 159.04 |
| 5 | sperm flagellum (GO:0036126) | 0.02570 | 0.05160 | 43.44 | 159.04 |

Abnormality of movement ([HP:0100022](https://hpo.jax.org/app/browse/term/HP:0100022))

KEGG Pathway

| **Index** | **Name** | **P-value** | **Adjusted p-value** | **Odds Ratio** | **Combined score** |
| --- | --- | --- | --- | --- | --- |
| 1 | Phototransduction | 0.00008446 | 0.0005068 | 191.96 | 1800.45 |
| 2 | Retinol metabolism | 0.03349 | 0.09401 | 33.04 | 112.22 |
| 3 | Drug metabolism | 0.05272 | 0.09401 | 20.65 | 60.76 |
| 4 | Purine metabolism | 0.06267 | 0.09401 | 17.24 | 47.76 |
| 5 | cGMP-PKG signaling pathway | 0.08045 | 0.09654 | 13.27 | 33.44 |

GO: Biological Process

| **Index** | **Name** | **P-value** | **Adjusted p-value** | **Odds Ratio** | **Combined score** |
| --- | --- | --- | --- | --- | --- |
| 1 | eye photoreceptor cell development (GO:0042462) | 0.00002354 | 0.0004630 | 384.17 | 4094.01 |
| 2 | visual perception (GO:0007601) | 9.004e-8 | 0.000002895 | 149.07 | 2418.38 |
| 3 | sensory perception of light stimulus (GO:0050953) | 9.813e-8 | 0.000002895 | 145.78 | 2352.45 |
| 4 | retinal rod cell development (GO:0046548) | 0.003495 | 0.02240 | 370.07 | 2093.27 |
| 5 | regulation of rhodopsin mediated signaling pathway (GO:0022400) | 0.00006709 | 0.0009895 | 217.03 | 2085.58 |

GO: Molecular Function

| **Index** | **Name** | **P-value** | **Adjusted p-value** | **Odds Ratio** | **Combined score** |
| --- | --- | --- | --- | --- | --- |
| 1 | leucine zipper domain binding (GO:0043522) | 0.00001010 | 0.0002829 | 624.44 | 7182.69 |
| 2 | LRR domain binding (GO:0030275) | 0.00002041 | 0.0002857 | 416.21 | 4494.85 |
| 3 | guanylate cyclase activator activity (GO:0030250) | 0.002498 | 0.01397 | 555.17 | 3326.77 |
| 4 | NADP-retinol dehydrogenase activity (GO:0052650) | 0.002997 | 0.01397 | 444.11 | 2580.41 |
| 5 | intracellular cGMP-activated cation channel activity (GO:0005223) | 0.003495 | 0.01397 | 370.07 | 2093.27 |

GO: Cellular Component

| **Index** | **Name** | **P-value** | **Adjusted p-value** | **Odds Ratio** | **Combined score** |
| --- | --- | --- | --- | --- | --- |
| 1 | sperm flagellum (GO:0036126) | 0.02570 | 0.1545 | 43.44 | 159.04 |
| 2 | 9+2 motile cilium (GO:0097729) | 0.02814 | 0.1545 | 39.55 | 141.22 |
| 3 | azurophil granule lumen (GO:0035578) | 0.04411 | 0.1545 | 24.85 | 77.54 |
| 4 | cytoplasmic vesicle lumen (GO:0060205) | 0.05605 | 0.1545 | 19.37 | 55.82 |
| 5 | ficolin-1-rich granule lumen (GO:1904813) | 0.05984 | 0.1545 | 18.09 | 50.96 |

Ataxia ([HP:0001251](https://hpo.jax.org/app/browse/term/HP:0001251))

GO: Biological Process

| **Index** | **Name** | **P-value** | **Adjusted p-value** | **Odds Ratio** | **Combined score** |
| --- | --- | --- | --- | --- | --- |
| 1 | cilium assembly (GO:0060271) | 7.884e-19 | 9.855e-18 | 196860.00 | 8205971.98 |
| 2 | ciliary basal body-plasma membrane docking (GO:0097711) | 8.302e-21 | 2.075e-19 | 2082.98 | 96312.24 |
| 3 | smoothened signaling pathway (GO:0007224) | 3.049e-12 | 1.270e-11 | 665.33 | 17642.09 |
| 4 | protein localization to ciliary transition zone (GO:1904491) | 0.000003371 | 0.00001204 | 1249.13 | 15739.23 |
| 5 | cilium organization (GO:0044782) | 1.113e-14 | 9.273e-14 | 359.45 | 11549.06 |

GO: Cellular Component

| **Index** | **Name** | **P-value** | **Adjusted p-value** | **Odds Ratio** | **Combined score** |
| --- | --- | --- | --- | --- | --- |
| 1 | ciliary membrane (GO:0060170) | 4.016e-7 | 0.000001606 | 305.54 | 4499.94 |
| 2 | cilium (GO:0005929) | 5.245e-10 | 4.196e-9 | 128.31 | 2741.69 |
| 3 | cell projection membrane (GO:0031253) | 0.00001104 | 0.00002944 | 95.83 | 1093.80 |
| 4 | motile cilium (GO:0031514) | 0.03349 | 0.04466 | 33.04 | 112.22 |
| 5 | cell-cell junction (GO:0005911) | 0.007662 | 0.01226 | 18.33 | 89.29 |

Babinski sign ([HP:0003487](https://hpo.jax.org/app/browse/term/HP:0003487))

KEGG Pathway

| **Index** | **Name** | **P-value** | **Adjusted p-value** | **Odds Ratio** | **Combined score** |
| --- | --- | --- | --- | --- | --- |
| 1 | Lysosome | 0.00002973 | 0.0001189 | 68.11 | 709.91 |
| 2 | Endocytosis | 0.1191 | 0.2137 | 8.74 | 18.59 |
| 3 | Amyotrophic lateral sclerosis | 0.1678 | 0.2137 | 6.01 | 10.72 |
| 4 | Pathways of neurodegeneration | 0.2137 | 0.2137 | 4.57 | 7.06 |

GO: Biological Process

| **Index** | **Name** | **P-value** | **Adjusted p-value** | **Odds Ratio** | **Combined score** |
| --- | --- | --- | --- | --- | --- |
| 1 | axo-dendritic transport (GO:0008088) | 0.002997 | 0.02686 | 444.11 | 2580.41 |
| 2 | membrane fission (GO:0090148) | 0.003495 | 0.02686 | 370.07 | 2093.27 |
| 3 | protein hexamerization (GO:0034214) | 0.004492 | 0.02686 | 277.53 | 1500.17 |
| 4 | synaptic vesicle transport (GO:0048489) | 0.005488 | 0.02686 | 222.00 | 1155.57 |
| 5 | axonal transport of mitochondrion (GO:0019896) | 0.005488 | 0.02686 | 222.00 | 1155.57 |

GO: Molecular Function

| **Index** | **Name** | **P-value** | **Adjusted p-value** | **Odds Ratio** | **Combined score** |
| --- | --- | --- | --- | --- | --- |
| 1 | olfactory receptor binding (GO:0031849) | 0.002997 | 0.01323 | 444.11 | 2580.41 |
| 2 | CD4 receptor binding (GO:0042609) | 0.003994 | 0.01323 | 317.19 | 1751.86 |
| 3 | alpha-tubulin binding (GO:0043014) | 0.01540 | 0.01854 | 73.93 | 308.54 |
| 4 | beta-tubulin binding (GO:0048487) | 0.01589 | 0.01854 | 71.54 | 296.32 |
| 5 | microtubule binding (GO:0008017) | 0.005670 | 0.01323 | 21.48 | 111.10 |

GO: Cellular Component

| **Index** | **Name** | **P-value** | **Adjusted p-value** | **Odds Ratio** | **Combined score** |
| --- | --- | --- | --- | --- | --- |
| 1 | endosome lumen (GO:0031904) | 2.326e-7 | 0.000007211 | 372.06 | 5682.76 |
| 2 | trans-Golgi network membrane (GO:0032588) | 0.00001377 | 0.0002134 | 88.81 | 994.11 |
| 3 | trans-Golgi network transport vesicle (GO:0030140) | 0.007476 | 0.03863 | 158.54 | 776.21 |
| 4 | endoplasmic reticulum tubular network (GO:0071782) | 0.01144 | 0.05068 | 100.85 | 450.83 |
| 5 | trans-Golgi network (GO:0005802) | 0.0001901 | 0.001964 | 35.87 | 307.37 |

Dysarthria ([HP:0001260](https://hpo.jax.org/app/browse/term/HP:0001260))

KEGG Pathway

| **Index** | **Name** | **P-value** | **Adjusted p-value** | **Odds Ratio** | **Combined score** |
| --- | --- | --- | --- | --- | --- |
| 1 | Amyotrophic lateral sclerosis | 4.861e-13 | 3.889e-12 | 220.61 | 6254.72 |
| 2 | Pathways of neurodegeneration | 3.368e-8 | 1.347e-7 | 62.43 | 1074.25 |
| 3 | Inositol phosphate metabolism | 0.03591 | 0.09198 | 30.74 | 102.25 |
| 4 | mRNA surveillance pathway | 0.04794 | 0.09198 | 22.79 | 69.22 |
| 5 | Spinocerebellar ataxia | 0.06926 | 0.09198 | 15.53 | 41.47 |

GO: Biological Process

| **Index** | **Name** | **P-value** | **Adjusted p-value** | **Odds Ratio** | **Combined score** |
| --- | --- | --- | --- | --- | --- |
| 1 | axo-dendritic transport (GO:0008088) | 0.002997 | 0.03670 | 444.11 | 2580.41 |
| 2 | regulation of cytoplasmic mRNA processing body assembly (GO:0010603) | 0.003495 | 0.03670 | 370.07 | 2093.27 |
| 3 | intermediate filament bundle assembly (GO:0045110) | 0.003495 | 0.03670 | 370.07 | 2093.27 |
| 4 | regulation of vesicle size (GO:0097494) | 0.003495 | 0.03670 | 370.07 | 2093.27 |
| 5 | nucleic acid transport (GO:0050657) | 0.004990 | 0.04084 | 246.68 | 1307.49 |

GO: Molecular Function

| **Index** | **Name** | **P-value** | **Adjusted p-value** | **Odds Ratio** | **Combined score** |
| --- | --- | --- | --- | --- | --- |
| 1 | epidermal growth factor receptor binding (GO:0005154) | 0.01293 | 0.06939 | 88.73 | 385.85 |
| 2 | kinesin binding (GO:0019894) | 0.01441 | 0.06939 | 79.21 | 335.86 |
| 3 | miRNA binding (GO:0035198) | 0.01490 | 0.06939 | 76.48 | 321.69 |
| 4 | protein serine/threonine kinase activator activity (GO:0043539) | 0.01835 | 0.06939 | 61.59 | 246.23 |
| 5 | regulatory RNA binding (GO:0061980) | 0.01983 | 0.06939 | 56.84 | 222.86 |

GO: Cellular Component

| **Index** | **Name** | **P-value** | **Adjusted p-value** | **Odds Ratio** | **Combined score** |
| --- | --- | --- | --- | --- | --- |
| 1 | neurofibrillary tangle (GO:0097418) | 0.002498 | 0.02498 | 555.17 | 3326.77 |
| 2 | axon (GO:0030424) | 0.0001190 | 0.001785 | 42.19 | 381.27 |
| 3 | nuclear inner membrane (GO:0005637) | 0.01392 | 0.06958 | 82.15 | 351.18 |
| 4 | neuron projection (GO:0043005) | 0.0001086 | 0.001785 | 23.48 | 214.29 |
| 5 | cytoplasmic stress granule (GO:0010494) | 0.03204 | 0.1005 | 34.59 | 119.03 |

Dysmetria ([HP:0001310](https://hpo.jax.org/app/browse/term/HP:0001310))

KEGG Pathway

| **Index** | **Name** | **P-value** | **Adjusted p-value** | **Odds Ratio** | **Combined score** |
| --- | --- | --- | --- | --- | --- |
| 1 | Spinocerebellar ataxia | 4.264e-9 | 3.113e-7 | 143.86 | 2772.51 |
| 2 | Sphingolipid signaling pathway | 0.001531 | 0.03429 | 42.46 | 275.24 |
| 3 | African trypanosomiasis | 0.01835 | 0.09771 | 61.59 | 246.23 |
| 4 | Aldosterone-regulated sodium reabsorption | 0.01835 | 0.09771 | 61.59 | 246.23 |
| 5 | Dopaminergic synapse | 0.001879 | 0.03429 | 38.19 | 239.73 |

GO: Biological Process

| **Index** | **Name** | **P-value** | **Adjusted p-value** | **Odds Ratio** | **Combined score** |
| --- | --- | --- | --- | --- | --- |
| 1 | response to morphine (GO:0043278) | 0.002498 | 0.05362 | 555.17 | 3326.77 |
| 2 | regulation of lyase activity (GO:0051339) | 0.002498 | 0.05362 | 555.17 | 3326.77 |
| 3 | regulation of cytoplasmic mRNA processing body assembly (GO:0010603) | 0.003495 | 0.05362 | 370.07 | 2093.27 |
| 4 | nucleic acid transport (GO:0050657) | 0.004990 | 0.05362 | 246.68 | 1307.49 |
| 5 | P-body assembly (GO:0033962) | 0.005985 | 0.05362 | 201.81 | 1032.95 |

GO: Molecular Function

| **Index** | **Name** | **P-value** | **Adjusted p-value** | **Odds Ratio** | **Combined score** |
| --- | --- | --- | --- | --- | --- |
| 1 | oxidoreductase activity, acting on metal ions, oxygen as acceptor (GO:0016724) | 0.003495 | 0.02156 | 370.07 | 2093.27 |
| 2 | ferric iron binding (GO:0008199) | 0.003495 | 0.02156 | 370.07 | 2093.27 |
| 3 | ferroxidase activity (GO:0004322) | 0.003495 | 0.02156 | 370.07 | 2093.27 |
| 4 | poly(G) binding (GO:0034046) | 0.003994 | 0.02156 | 317.19 | 1751.86 |
| 5 | protein-glutamine gamma-glutamyltransferase activity (GO:0003810) | 0.004492 | 0.02156 | 277.53 | 1500.17 |

GO: Cellular Component

| **Index** | **Name** | **P-value** | **Adjusted p-value** | **Odds Ratio** | **Combined score** |
| --- | --- | --- | --- | --- | --- |
| 1 | nuclear inclusion body (GO:0042405) | 0.005488 | 0.04207 | 222.00 | 1155.57 |
| 2 | protein phosphatase type 2A complex (GO:0000159) | 0.008469 | 0.04870 | 138.71 | 661.82 |
| 3 | dendrite membrane (GO:0032590) | 0.01441 | 0.06628 | 79.21 | 335.86 |
| 4 | dendrite (GO:0030425) | 0.0002722 | 0.006260 | 31.66 | 259.88 |
| 5 | cytoplasmic stress granule (GO:0010494) | 0.03204 | 0.1130 | 34.59 | 119.03 |

Dysphagia ([HP:0002015](https://hpo.jax.org/app/browse/term/HP:0002015))

KEGG Pathway

| **Index** | **Name** | **P-value** | **Adjusted p-value** | **Odds Ratio** | **Combined score** |
| --- | --- | --- | --- | --- | --- |
| 1 | Amyotrophic lateral sclerosis | 4.861e-13 | 3.889e-12 | 220.61 | 6254.72 |
| 2 | Pathways of neurodegeneration | 3.368e-8 | 1.347e-7 | 62.43 | 1074.25 |
| 3 | Inositol phosphate metabolism | 0.03591 | 0.09198 | 30.74 | 102.25 |
| 4 | mRNA surveillance pathway | 0.04794 | 0.09198 | 22.79 | 69.22 |
| 5 | Spinocerebellar ataxia | 0.06926 | 0.09198 | 15.53 | 41.47 |

GO: Biological Process

| **Index** | **Name** | **P-value** | **Adjusted p-value** | **Odds Ratio** | **Combined score** |
| --- | --- | --- | --- | --- | --- |
| 1 | axo-dendritic transport (GO:0008088) | 0.002997 | 0.03670 | 444.11 | 2580.41 |
| 2 | regulation of cytoplasmic mRNA processing body assembly (GO:0010603) | 0.003495 | 0.03670 | 370.07 | 2093.27 |
| 3 | intermediate filament bundle assembly (GO:0045110) | 0.003495 | 0.03670 | 370.07 | 2093.27 |
| 4 | regulation of vesicle size (GO:0097494) | 0.003495 | 0.03670 | 370.07 | 2093.27 |
| 5 | nucleic acid transport (GO:0050657) | 0.004990 | 0.04084 | 246.68 | 1307.49 |

GO: Molecular Function

| **Index** | **Name** | **P-value** | **Adjusted p-value** | **Odds Ratio** | **Combined score** |
| --- | --- | --- | --- | --- | --- |
| 1 | epidermal growth factor receptor binding (GO:0005154) | 0.01293 | 0.06939 | 88.73 | 385.85 |
| 2 | kinesin binding (GO:0019894) | 0.01441 | 0.06939 | 79.21 | 335.86 |
| 3 | miRNA binding (GO:0035198) | 0.01490 | 0.06939 | 76.48 | 321.69 |
| 4 | protein serine/threonine kinase activator activity (GO:0043539) | 0.01835 | 0.06939 | 61.59 | 246.23 |
| 5 | regulatory RNA binding (GO:0061980) | 0.01983 | 0.06939 | 56.84 | 222.86 |

GO: Cellular Component

| **Index** | **Name** | **P-value** | **Adjusted p-value** | **Odds Ratio** | **Combined score** |
| --- | --- | --- | --- | --- | --- |
| 1 | neurofibrillary tangle (GO:0097418) | 0.002498 | 0.02498 | 555.17 | 3326.77 |
| 2 | axon (GO:0030424) | 0.0001190 | 0.001785 | 42.19 | 381.27 |
| 3 | nuclear inner membrane (GO:0005637) | 0.01392 | 0.06958 | 82.15 | 351.18 |
| 4 | neuron projection (GO:0043005) | 0.0001086 | 0.001785 | 23.48 | 214.29 |
| 5 | cytoplasmic stress granule (GO:0010494) | 0.03204 | 0.1005 | 34.59 | 119.03 |

Dystonia ([HP:0001332](https://hpo.jax.org/app/browse/term/HP:0001332))

KEGG Pathway

| **Index** | **Name** | **P-value** | **Adjusted p-value** | **Odds Ratio** | **Combined score** |
| --- | --- | --- | --- | --- | --- |
| 1 | Nicotine addiction | 9.018e-15 | 5.050e-13 | 880.41 | 28472.15 |
| 2 | Long-term potentiation | 9.011e-11 | 1.957e-9 | 321.42 | 7434.41 |
| 3 | Amphetamine addiction | 1.048e-10 | 1.957e-9 | 311.34 | 7154.30 |
| 4 | Cocaine addiction | 6.603e-9 | 4.622e-8 | 295.48 | 5565.60 |
| 5 | Circadian entrainment | 5.976e-10 | 8.366e-9 | 216.28 | 4593.43 |

GO: Biological Process

| **Index** | **Name** | **P-value** | **Adjusted p-value** | **Odds Ratio** | **Combined score** |
| --- | --- | --- | --- | --- | --- |
| 1 | excitatory chemical synaptic transmission (GO:0098976) | 3.147e-9 | 6.383e-8 | 2141.36 | 41921.23 |
| 2 | regulation of NMDA receptor activity (GO:2000310) | 1.893e-12 | 8.958e-11 | 739.37 | 19957.87 |
| 3 | glutamate receptor signaling pathway (GO:0007215) | 4.093e-12 | 1.453e-10 | 623.69 | 16354.24 |
| 4 | response to ethanol (GO:0045471) | 4.082e-8 | 7.246e-7 | 713.50 | 12139.53 |
| 5 | anterograde trans-synaptic signaling (GO:0098916) | 1.927e-14 | 2.737e-12 | 334.81 | 10573.42 |

GO: Molecular Function

| **Index** | **Name** | **P-value** | **Adjusted p-value** | **Odds Ratio** | **Combined score** |
| --- | --- | --- | --- | --- | --- |
| 1 | glutamate-gated calcium ion channel activity (GO:0022849) | 8.995e-10 | 5.509e-9 | 4283.14 | 89214.48 |
| 2 | ionotropic glutamate receptor activity (GO:0004970) | 5.833e-14 | 2.858e-12 | 1664.83 | 50731.79 |
| 3 | NMDA glutamate receptor activity (GO:0004972) | 5.033e-9 | 2.740e-8 | 1713.00 | 32730.63 |
| 4 | ligand-gated channel activity (GO:0022834) | 1.340e-12 | 2.609e-11 | 798.60 | 21832.48 |
| 5 | ligand-gated ion channel activity (GO:0015276) | 1.597e-12 | 2.609e-11 | 767.85 | 20856.82 |

GO: Cellular Component

| **Index** | **Name** | **P-value** | **Adjusted p-value** | **Odds Ratio** | **Combined score** |
| --- | --- | --- | --- | --- | --- |
| 1 | NMDA selective glutamate receptor complex (GO:0017146) | 5.033e-9 | 3.775e-8 | 1713.00 | 32730.63 |
| 2 | ionotropic glutamate receptor complex (GO:0008328) | 1.893e-12 | 5.678e-11 | 739.37 | 19957.87 |
| 3 | cation channel complex (GO:0034703) | 1.400e-10 | 2.100e-9 | 292.97 | 6647.37 |
| 4 | synaptic membrane (GO:0097060) | 2.326e-7 | 0.000001163 | 372.06 | 5682.76 |
| 5 | postsynaptic density membrane (GO:0098839) | 0.00004255 | 0.0001410 | 277.39 | 2791.90 |

Facial paralysis ([HP:0007209](https://hpo.jax.org/app/browse/term/HP:0007209))

KEGG Pathway

| **Index** | **Name** | **P-value** | **Adjusted p-value** | **Odds Ratio** | **Combined score** |
| --- | --- | --- | --- | --- | --- |
| 1 | Rheumatoid arthritis | 0.0002120 | 0.007631 | 145.82 | 1233.48 |
| 2 | Proximal tubule bicarbonate reclamation | 0.005737 | 0.04291 | 226.97 | 1171.32 |
| 3 | Collecting duct acid secretion | 0.006732 | 0.04291 | 192.01 | 960.21 |
| 4 | Aldosterone-regulated sodium reabsorption | 0.009217 | 0.04291 | 138.60 | 649.60 |
| 5 | Carbohydrate digestion and absorption | 0.01170 | 0.04291 | 108.42 | 482.30 |

GO: Biological Process

| **Index** | **Name** | **P-value** | **Adjusted p-value** | **Odds Ratio** | **Combined score** |
| --- | --- | --- | --- | --- | --- |
| 1 | regulation of respiratory gaseous exchange by nervous system process (GO:0002087) | 0.001249 | 0.01619 | 1249.44 | 8352.53 |
| 2 | positive regulation of prostaglandin secretion (GO:0032308) | 0.001249 | 0.01619 | 1249.44 | 8352.53 |
| 3 | positive regulation of fever generation (GO:0031622) | 0.001499 | 0.01619 | 999.50 | 6499.56 |
| 4 | negative regulation of cytosolic calcium ion concentration (GO:0051481) | 0.001499 | 0.01619 | 999.50 | 6499.56 |
| 5 | membrane depolarization during action potential (GO:0086010) | 0.00001621 | 0.001377 | 554.75 | 6118.80 |

GO: Molecular Function

| **Index** | **Name** | **P-value** | **Adjusted p-value** | **Odds Ratio** | **Combined score** |
| --- | --- | --- | --- | --- | --- |
| 1 | P-type sodium transporter activity (GO:0008554) | 0.001499 | 0.01006 | 999.50 | 6499.56 |
| 2 | P-type sodium:potassium-exchanging transporter activity (GO:0005391) | 0.001499 | 0.01006 | 999.50 | 6499.56 |
| 3 | sodium ion binding (GO:0031402) | 0.001499 | 0.01006 | 999.50 | 6499.56 |
| 4 | P-type potassium transmembrane transporter activity (GO:0008556) | 0.001749 | 0.01006 | 832.88 | 5287.72 |
| 5 | potassium ion binding (GO:0030955) | 0.002248 | 0.01021 | 624.59 | 3808.55 |

GO: Cellular Component

| **Index** | **Name** | **P-value** | **Adjusted p-value** | **Odds Ratio** | **Combined score** |
| --- | --- | --- | --- | --- | --- |
| 1 | sodium:potassium-exchanging ATPase complex (GO:0005890) | 0.002498 | 0.02055 | 555.17 | 3326.77 |
| 2 | proton-transporting V-type ATPase complex (GO:0033176) | 0.003745 | 0.02055 | 356.80 | 1993.61 |
| 3 | cation-transporting ATPase complex (GO:0090533) | 0.003994 | 0.02055 | 333.00 | 1839.15 |
| 4 | voltage-gated sodium channel complex (GO:0001518) | 0.004243 | 0.02055 | 312.17 | 1705.22 |
| 5 | vacuolar proton-transporting V-type ATPase complex (GO:0016471) | 0.004741 | 0.02055 | 277.46 | 1484.80 |

Gait disturbance ([HP:0001288](https://hpo.jax.org/app/browse/term/HP:0001288))

KEGG Pathway

| **Index** | **Name** | **P-value** | **Adjusted p-value** | **Odds Ratio** | **Combined score** |
| --- | --- | --- | --- | --- | --- |
| 1 | Nicotine addiction | 9.018e-15 | 4.689e-13 | 880.41 | 28472.15 |
| 2 | GABAergic synapse | 1.354e-12 | 3.520e-11 | 359.77 | 9831.65 |
| 3 | Retrograde endocannabinoid signaling | 5.071e-9 | 6.592e-8 | 138.79 | 2650.86 |
| 4 | Morphine addiction | 8.246e-8 | 8.576e-7 | 152.51 | 2487.64 |
| 5 | Neuroactive ligand-receptor interaction | 4.661e-9 | 6.592e-8 | 88.01 | 1688.34 |

GO: Biological Process

| **Index** | **Name** | **P-value** | **Adjusted p-value** | **Odds Ratio** | **Combined score** |
| --- | --- | --- | --- | --- | --- |
| 1 | anterograde trans-synaptic signaling (GO:0098916) | 5.108e-17 | 6.896e-15 | 756.57 | 28381.44 |
| 2 | chemical synaptic transmission (GO:0007268) | 4.030e-16 | 2.720e-14 | 596.76 | 21153.59 |
| 3 | chloride transmembrane transport (GO:1902476) | 3.727e-14 | 1.677e-12 | 679.98 | 21025.29 |
| 4 | synaptic transmission, GABAergic (GO:0051932) | 1.482e-8 | 2.858e-7 | 1070.46 | 19297.66 |
| 5 | inhibitory synapse assembly (GO:1904862) | 1.482e-8 | 2.858e-7 | 1070.46 | 19297.66 |

GO: Molecular Function

| **Index** | **Name** | **P-value** | **Adjusted p-value** | **Odds Ratio** | **Combined score** |
| --- | --- | --- | --- | --- | --- |
| 1 | GABA-A receptor activity (GO:0004890) | 1.217e-10 | 6.310e-9 | 887.78 | 20267.77 |
| 2 | GABA receptor activity (GO:0016917) | 2.294e-10 | 6.310e-9 | 739.70 | 16417.99 |
| 3 | GABA-gated chloride ion channel activity (GO:0022851) | 2.567e-8 | 2.353e-7 | 856.29 | 14966.03 |
| 4 | ligand-gated anion channel activity (GO:0099095) | 7.315e-8 | 5.748e-7 | 570.71 | 9377.24 |
| 5 | transmitter-gated ion channel activity (GO:0022824) | 1.451e-9 | 2.659e-8 | 443.56 | 9026.96 |

GO: Cellular Component

| **Index** | **Name** | **P-value** | **Adjusted p-value** | **Odds Ratio** | **Combined score** |
| --- | --- | --- | --- | --- | --- |
| 1 | neuron projection (GO:0043005) | 2.546e-16 | 6.110e-15 | 194440.00 | 6981748.76 |
| 2 | GABA-A receptor complex (GO:1902711) | 1.217e-10 | 1.460e-9 | 887.78 | 20267.77 |
| 3 | excitatory synapse (GO:0060076) | 0.00006709 | 0.0003220 | 217.03 | 2085.58 |
| 4 | neuron to neuron synapse (GO:0098984) | 0.003994 | 0.006846 | 317.19 | 1751.86 |
| 5 | NMDA selective glutamate receptor complex (GO:0017146) | 0.003994 | 0.006846 | 317.19 | 1751.86 |

Hand muscle weakness ([HP:0030237](https://hpo.jax.org/app/browse/term/HP:0030237))

KEGG Pathway

| **Index** | **Name** | **P-value** | **Adjusted p-value** | **Odds Ratio** | **Combined score** |
| --- | --- | --- | --- | --- | --- |
| 1 | Neuroactive ligand-receptor interaction | 0.00001608 | 0.00008039 | 38.88 | 429.14 |
| 2 | Mitophagy | 0.03349 | 0.08373 | 33.04 | 112.22 |
| 3 | NOD-like receptor signaling pathway | 0.08692 | 0.1449 | 12.23 | 29.87 |
| 4 | Parkinson disease | 0.1178 | 0.1472 | 8.84 | 18.92 |
| 5 | Pathways of neurodegeneration | 0.2137 | 0.2137 | 4.57 | 7.06 |

GO: Biological Process

| **Index** | **Name** | **P-value** | **Adjusted p-value** | **Odds Ratio** | **Combined score** |
| --- | --- | --- | --- | --- | --- |
| 1 | synaptic transmission, cholinergic (GO:0007271) | 1.217e-10 | 9.611e-9 | 887.78 | 20267.77 |
| 2 | neuromuscular junction development (GO:0007528) | 1.812e-7 | 0.000003793 | 407.53 | 6326.45 |
| 3 | positive regulation of vascular associated smooth muscle cell apoptotic process (GO:1905461) | 0.002498 | 0.01644 | 555.17 | 3326.77 |
| 4 | neuromuscular process (GO:0050905) | 0.00003830 | 0.0003735 | 293.72 | 2987.15 |
| 5 | postsynaptic membrane organization (GO:0001941) | 0.00003830 | 0.0003735 | 293.72 | 2987.15 |

GO: Molecular Function

| **Index** | **Name** | **P-value** | **Adjusted p-value** | **Odds Ratio** | **Combined score** |
| --- | --- | --- | --- | --- | --- |
| 1 | neurotransmitter receptor activity involved in regulation of postsynaptic membrane potential (GO:0099529) | 1.878e-10 | 4.882e-9 | 783.25 | 17541.62 |
| 2 | transmitter-gated ion channel activity involved in regulation of postsynaptic membrane potential (GO:1904315) | 8.580e-10 | 1.115e-8 | 511.90 | 10686.61 |
| 3 | transmitter-gated ion channel activity (GO:0022824) | 1.451e-9 | 1.257e-8 | 443.56 | 9026.96 |
| 4 | postsynaptic neurotransmitter receptor activity (GO:0098960) | 1.191e-7 | 7.172e-7 | 475.52 | 7581.26 |
| 5 | acetylcholine receptor activity (GO:0015464) | 1.379e-7 | 7.172e-7 | 450.47 | 7115.97 |

GO: Cellular Component

| **Index** | **Name** | **P-value** | **Adjusted p-value** | **Odds Ratio** | **Combined score** |
| --- | --- | --- | --- | --- | --- |
| 1 | neuromuscular junction (GO:0031594) | 4.712e-12 | 7.067e-11 | 604.76 | 15772.67 |
| 2 | acetylcholine-gated channel complex (GO:0005892) | 3.267e-8 | 2.450e-7 | 778.40 | 13417.28 |
| 3 | ion channel complex (GO:0034702) | 5.841e-7 | 0.000002921 | 267.29 | 3836.51 |
| 4 | intrinsic component of mitochondrial membrane (GO:0098573) | 0.005985 | 0.01496 | 201.81 | 1032.95 |
| 5 | voltage-gated sodium channel complex (GO:0001518) | 0.008469 | 0.01815 | 138.71 | 661.82 |

Hyperkinetic movements ([HP:0002487](https://hpo.jax.org/app/browse/term/HP:0002487))

KEGG Pathway

| **Index** | **Name** | **P-value** | **Adjusted p-value** | **Odds Ratio** | **Combined score** |
| --- | --- | --- | --- | --- | --- |
| 1 | Nicotine addiction | 8.806e-7 | 0.00001843 | 231.12 | 3222.36 |
| 2 | Dopaminergic synapse | 2.847e-9 | 1.281e-7 | 156.40 | 3077.49 |
| 3 | Cocaine addiction | 0.000001638 | 0.00001843 | 185.81 | 2475.38 |
| 4 | Butanoate metabolism | 0.00008446 | 0.0003801 | 191.96 | 1800.45 |
| 5 | Amphetamine addiction | 0.000004635 | 0.00004171 | 129.38 | 1588.99 |

GO: Biological Process

| **Index** | **Name** | **P-value** | **Adjusted p-value** | **Odds Ratio** | **Combined score** |
| --- | --- | --- | --- | --- | --- |
| 1 | gamma-aminobutyric acid metabolic process (GO:0009448) | 0.000002248 | 0.0001158 | 1665.58 | 21661.60 |
| 2 | excitatory chemical synaptic transmission (GO:0098976) | 0.000004719 | 0.0001215 | 999.25 | 12254.80 |
| 3 | response to ethanol (GO:0045471) | 0.00002354 | 0.0004042 | 384.17 | 4094.01 |
| 4 | glutamate catabolic process (GO:0006538) | 0.002498 | 0.008039 | 555.17 | 3326.77 |
| 5 | excitatory postsynaptic potential (GO:0060079) | 0.00003830 | 0.0005636 | 293.72 | 2987.15 |

GO: Molecular Function

| **Index** | **Name** | **P-value** | **Adjusted p-value** | **Odds Ratio** | **Combined score** |
| --- | --- | --- | --- | --- | --- |
| 1 | glutamate-gated calcium ion channel activity (GO:0022849) | 0.000002248 | 0.00007644 | 1665.58 | 21661.60 |
| 2 | NMDA glutamate receptor activity (GO:0004972) | 0.000006290 | 0.00008213 | 832.67 | 9972.50 |
| 3 | ionotropic glutamate receptor activity (GO:0004970) | 0.00003048 | 0.0002072 | 332.92 | 3461.84 |
| 4 | opioid receptor binding (GO:0031628) | 0.002498 | 0.007077 | 555.17 | 3326.77 |
| 5 | ligand-gated calcium channel activity (GO:0099604) | 0.00003830 | 0.0002170 | 293.72 | 2987.15 |

GO: Cellular Component

| **Index** | **Name** | **P-value** | **Adjusted p-value** | **Odds Ratio** | **Combined score** |
| --- | --- | --- | --- | --- | --- |
| 1 | NMDA selective glutamate receptor complex (GO:0017146) | 0.000006290 | 0.0001698 | 832.67 | 9972.50 |
| 2 | flotillin complex (GO:0016600) | 0.002498 | 0.008430 | 555.17 | 3326.77 |
| 3 | synaptic membrane (GO:0097060) | 0.00007266 | 0.0009809 | 207.98 | 1981.99 |
| 4 | clathrin-sculpted gamma-aminobutyric acid transport vesicle (GO:0061200) | 0.003994 | 0.009933 | 317.19 | 1751.86 |
| 5 | clathrin-sculpted gamma-aminobutyric acid transport vesicle membrane (GO:0061202) | 0.003994 | 0.009933 | 317.19 | 1751.86 |

Hypotonia ([HP:0001252](https://hpo.jax.org/app/browse/term/HP:0001252))

KEGG Pathway

| **Index** | **Name** | **P-value** | **Adjusted p-value** | **Odds Ratio** | **Combined score** |
| --- | --- | --- | --- | --- | --- |
| 1 | Oxidative phosphorylation | 1.197e-22 | 1.317e-21 | 198670.00 | 10028275.58 |
| 2 | Retrograde endocannabinoid signaling | 3.613e-22 | 1.987e-21 | 198520.00 | 9801417.00 |
| 3 | Non-alcoholic fatty liver disease | 5.818e-22 | 2.133e-21 | 198450.00 | 9703402.49 |
| 4 | Diabetic cardiomyopathy | 9.277e-21 | 2.551e-20 | 197970.00 | 9131721.76 |
| 5 | Thermogenesis | 3.629e-20 | 7.983e-20 | 197680.00 | 8848725.20 |

GO: Biological Process

| **Index** | **Name** | **P-value** | **Adjusted p-value** | **Odds Ratio** | **Combined score** |
| --- | --- | --- | --- | --- | --- |
| 1 | mitochondrial electron transport, NADH to ubiquinone (GO:0006120) | 2.254e-28 | 4.958e-27 | 199610.00 | 12707139.91 |
| 2 | mitochondrial respiratory chain complex I assembly (GO:0032981) | 1.850e-26 | 1.357e-25 | 199420.00 | 11816020.17 |
| 3 | NADH dehydrogenase complex assembly (GO:0010257) | 1.850e-26 | 1.357e-25 | 199420.00 | 11816020.17 |
| 4 | aerobic electron transport chain (GO:0019646) | 1.407e-25 | 7.205e-25 | 199300.00 | 11404606.96 |
| 5 | mitochondrial ATP synthesis coupled electron transport (GO:0042775) | 1.637e-25 | 7.205e-25 | 199290.00 | 11373779.66 |

GO: Molecular Function

| **Index** | **Name** | **P-value** | **Adjusted p-value** | **Odds Ratio** | **Combined score** |
| --- | --- | --- | --- | --- | --- |
| 1 | NADH dehydrogenase (quinone) activity (GO:0050136) | 5.000e-25 | 1.000e-24 | 6910.62 | 386684.82 |
| 2 | NADH dehydrogenase (ubiquinone) activity (GO:0008137) | 5.000e-25 | 1.000e-24 | 6910.62 | 386684.82 |
| 3 | oxidoreduction-driven active transmembrane transporter activity (GO:0015453) | 7.535e-23 | 1.005e-22 | 3662.63 | 186574.14 |
| 4 | NADH dehydrogenase activity (GO:0003954) | 0.000004719 | 0.000004719 | 999.25 | 12254.80 |

GO: Cellular Component

| **Index** | **Name** | **P-value** | **Adjusted p-value** | **Odds Ratio** | **Combined score** |
| --- | --- | --- | --- | --- | --- |
| 1 | mitochondrial respiratory chain complex I (GO:0005747) | 5.216e-28 | 4.173e-27 | 199580.00 | 12537732.35 |
| 2 | respiratory chain complex I (GO:0045271) | 5.216e-28 | 4.173e-27 | 199580.00 | 12537732.35 |
| 3 | mitochondrial membrane (GO:0031966) | 4.573e-17 | 2.439e-16 | 195310.00 | 7348319.17 |
| 4 | mitochondrial inner membrane (GO:0005743) | 7.582e-16 | 3.033e-15 | 554.98 | 19322.03 |
| 5 | organelle inner membrane (GO:0019866) | 1.232e-15 | 3.944e-15 | 524.86 | 18018.26 |

Muscle weakness ([HP:0001324](https://hpo.jax.org/app/browse/term/HP:0001324))

KEGG Pathway

| **Index** | **Name** | **P-value** | **Adjusted p-value** | **Odds Ratio** | **Combined score** |
| --- | --- | --- | --- | --- | --- |
| 1 | Mannose type O-glycan biosynthesis | 1.586e-7 | 0.000001110 | 427.93 | 6700.13 |
| 2 | Viral myocarditis | 0.02960 | 0.05482 | 37.53 | 132.12 |
| 3 | Arrhythmogenic right ventricular cardiomyopathy | 0.03785 | 0.05482 | 29.11 | 95.32 |
| 4 | Taste transduction | 0.04219 | 0.05482 | 26.02 | 82.37 |
| 5 | Hypertrophic cardiomyopathy | 0.04411 | 0.05482 | 24.85 | 77.54 |

GO: Biological Process

| **Index** | **Name** | **P-value** | **Adjusted p-value** | **Odds Ratio** | **Combined score** |
| --- | --- | --- | --- | --- | --- |
| 1 | neuronal action potential (GO:0019228) | 2.480e-13 | 1.116e-11 | 1174.88 | 34101.37 |
| 2 | membrane depolarization during action potential (GO:0086010) | 6.190e-13 | 1.393e-11 | 950.90 | 26730.60 |
| 3 | membrane depolarization (GO:0051899) | 1.893e-12 | 2.839e-11 | 739.37 | 19957.87 |
| 4 | action potential (GO:0001508) | 1.145e-11 | 1.288e-10 | 498.75 | 12564.94 |
| 5 | sodium ion transmembrane transport (GO:0035725) | 3.433e-10 | 3.061e-9 | 242.78 | 5290.74 |

GO: Molecular Function

| **Index** | **Name** | **P-value** | **Adjusted p-value** | **Odds Ratio** | **Combined score** |
| --- | --- | --- | --- | --- | --- |
| 1 | voltage-gated sodium channel activity (GO:0005248) | 2.480e-13 | 2.232e-12 | 1174.88 | 34101.37 |
| 2 | sodium channel activity (GO:0005272) | 4.712e-12 | 2.120e-11 | 604.76 | 15772.67 |
| 3 | actinin binding (GO:0042805) | 0.01045 | 0.02686 | 110.94 | 506.01 |
| 4 | alpha-actinin binding (GO:0051393) | 0.01194 | 0.02686 | 96.46 | 427.12 |
| 5 | acetylglucosaminyltransferase activity (GO:0008375) | 0.02375 | 0.03562 | 47.15 | 176.34 |

GO: Cellular Component

| **Index** | **Name** | **P-value** | **Adjusted p-value** | **Odds Ratio** | **Combined score** |
| --- | --- | --- | --- | --- | --- |
| 1 | voltage-gated sodium channel complex (GO:0001518) | 5.833e-14 | 9.917e-13 | 1664.83 | 50731.79 |
| 2 | sodium channel complex (GO:0034706) | 5.000e-13 | 4.250e-12 | 998.50 | 28281.59 |
| 3 | axon (GO:0030424) | 2.123e-10 | 1.203e-9 | 149.94 | 3339.58 |
| 4 | neuron projection (GO:0043005) | 8.582e-8 | 3.647e-7 | 53.02 | 862.66 |
| 5 | integral component of Golgi membrane (GO:0030173) | 0.0003294 | 0.0009334 | 94.04 | 754.04 |

Orthostatic hypotension ([HP:0001278](https://hpo.jax.org/app/browse/term/HP:0001278))

KEGG Pathway

| **Index** | **Name** | **P-value** | **Adjusted p-value** | **Odds Ratio** | **Combined score** |
| --- | --- | --- | --- | --- | --- |
| 1 | JAK-STAT signaling pathway | 0.002812 | 0.04780 | 30.98 | 182.00 |
| 2 | Sphingolipid metabolism | 0.02424 | 0.09613 | 46.16 | 171.72 |
| 3 | Inflammatory bowel disease | 0.03204 | 0.09613 | 34.59 | 119.03 |
| 4 | Adipocytokine signaling pathway | 0.03398 | 0.09613 | 32.55 | 110.09 |
| 5 | Cytokine-cytokine receptor interaction | 0.009024 | 0.06752 | 16.81 | 79.12 |

GO: Biological Process

| **Index** | **Name** | **P-value** | **Adjusted p-value** | **Odds Ratio** | **Combined score** |
| --- | --- | --- | --- | --- | --- |
| 1 | negative regulation of response to food (GO:0032096) | 0.002498 | 0.03060 | 555.17 | 3326.77 |
| 2 | activation of protein kinase C activity (GO:1990051) | 0.002498 | 0.03060 | 555.17 | 3326.77 |
| 3 | positive regulation of cell fate commitment (GO:0010455) | 0.002498 | 0.03060 | 555.17 | 3326.77 |
| 4 | anion homeostasis (GO:0055081) | 0.002498 | 0.03060 | 555.17 | 3326.77 |
| 5 | response to acetylcholine (GO:1905144) | 0.002498 | 0.03060 | 555.17 | 3326.77 |

GO: Molecular Function

| **Index** | **Name** | **P-value** | **Adjusted p-value** | **Odds Ratio** | **Combined score** |
| --- | --- | --- | --- | --- | --- |
| 1 | interleukin-12 receptor binding (GO:0005143) | 0.002997 | 0.03163 | 444.11 | 2580.41 |
| 2 | acetylcholine-gated cation-selective channel activity (GO:0022848) | 0.005488 | 0.03163 | 222.00 | 1155.57 |
| 3 | arylsulfatase activity (GO:0004065) | 0.006979 | 0.03163 | 170.74 | 847.70 |
| 4 | peptide hormone receptor binding (GO:0051428) | 0.006979 | 0.03163 | 170.74 | 847.70 |
| 5 | sulfuric ester hydrolase activity (GO:0008484) | 0.008469 | 0.03163 | 138.71 | 661.82 |

GO: Cellular Component

| **Index** | **Name** | **P-value** | **Adjusted p-value** | **Odds Ratio** | **Combined score** |
| --- | --- | --- | --- | --- | --- |
| 1 | chromaffin granule (GO:0042583) | 0.002498 | 0.02997 | 555.17 | 3326.77 |
| 2 | acetylcholine-gated channel complex (GO:0005892) | 0.006979 | 0.05360 | 170.74 | 847.70 |
| 3 | ion channel complex (GO:0034702) | 0.01737 | 0.08931 | 65.22 | 264.33 |
| 4 | dendrite (GO:0030425) | 0.0002722 | 0.009798 | 31.66 | 259.88 |
| 5 | mitochondrial intermembrane space (GO:0005758) | 0.02863 | 0.1048 | 38.86 | 138.07 |

Paresthesia ([HP:0003401](https://hpo.jax.org/app/browse/term/HP:0003401))

KEGG Pathway

| **Index** | **Name** | **P-value** | **Adjusted p-value** | **Odds Ratio** | **Combined score** |
| --- | --- | --- | --- | --- | --- |
| 1 | Aldosterone-regulated sodium reabsorption | 6.931e-7 | 0.00008387 | 251.55 | 3567.44 |
| 2 | Endometrial cancer | 0.0003664 | 0.005434 | 88.99 | 704.07 |
| 3 | VEGF signaling pathway | 0.0003792 | 0.005434 | 87.43 | 688.70 |
| 4 | GnRH secretion | 0.0004462 | 0.005434 | 80.35 | 619.92 |
| 5 | Acute myeloid leukemia | 0.0004889 | 0.005434 | 76.63 | 584.21 |

GO: Biological Process

| **Index** | **Name** | **P-value** | **Adjusted p-value** | **Odds Ratio** | **Combined score** |
| --- | --- | --- | --- | --- | --- |
| 1 | chloride ion homeostasis (GO:0055064) | 0.00001010 | 0.0004991 | 624.44 | 7182.69 |
| 2 | monovalent inorganic anion homeostasis (GO:0055083) | 0.00001010 | 0.0004991 | 624.44 | 7182.69 |
| 3 | megakaryocyte differentiation (GO:0030219) | 0.00001481 | 0.0006097 | 499.50 | 5554.53 |
| 4 | regulation of glycoprotein metabolic process (GO:1903018) | 0.002498 | 0.02200 | 555.17 | 3326.77 |
| 5 | positive regulation of toll-like receptor 9 signaling pathway (GO:0034165) | 0.002498 | 0.02200 | 555.17 | 3326.77 |

GO: Molecular Function

| **Index** | **Name** | **P-value** | **Adjusted p-value** | **Odds Ratio** | **Combined score** |
| --- | --- | --- | --- | --- | --- |
| 1 | potassium:chloride symporter activity (GO:0015379) | 0.000008085 | 0.0002054 | 713.68 | 8368.26 |
| 2 | sodium:chloride symporter activity (GO:0015378) | 0.000008085 | 0.0002054 | 713.68 | 8368.26 |
| 3 | cation:chloride symporter activity (GO:0015377) | 0.00001010 | 0.0002054 | 624.44 | 7182.69 |
| 4 | anion:sodium symporter activity (GO:0015373) | 0.002498 | 0.02611 | 555.17 | 3326.77 |
| 5 | ATP-activated inward rectifier potassium channel activity (GO:0015272) | 0.002997 | 0.02611 | 444.11 | 2580.41 |

GO: Cellular Component

| **Index** | **Name** | **P-value** | **Adjusted p-value** | **Odds Ratio** | **Combined score** |
| --- | --- | --- | --- | --- | --- |
| 1 | phosphatidylinositol 3-kinase complex, class I (GO:0097651) | 0.002498 | 0.03479 | 555.17 | 3326.77 |
| 2 | intercalated disc (GO:0014704) | 0.01540 | 0.08724 | 73.93 | 308.54 |
| 3 | cell-cell contact zone (GO:0044291) | 0.02326 | 0.09237 | 48.17 | 181.19 |
| 4 | cytoplasmic side of plasma membrane (GO:0009898) | 0.02717 | 0.09237 | 41.02 | 147.91 |
| 5 | voltage-gated potassium channel complex (GO:0008076) | 0.03591 | 0.09543 | 30.74 | 102.25 |

Parkinsonism ([HP:0001300](https://hpo.jax.org/app/browse/term/HP:0001300))

KEGG Pathway

| **Index** | **Name** | **P-value** | **Adjusted p-value** | **Odds Ratio** | **Combined score** |
| --- | --- | --- | --- | --- | --- |
| 1 | Nicotine addiction | 0.0001737 | 0.006081 | 131.26 | 1136.48 |
| 2 | Proximal tubule bicarbonate reclamation | 0.01144 | 0.04634 | 100.85 | 450.83 |
| 3 | Taste transduction | 0.0008042 | 0.007873 | 59.24 | 422.16 |
| 4 | GABAergic synapse | 0.0008609 | 0.007873 | 57.19 | 403.64 |
| 5 | Morphine addiction | 0.0008998 | 0.007873 | 55.90 | 392.06 |

GO: Biological Process

| **Index** | **Name** | **P-value** | **Adjusted p-value** | **Odds Ratio** | **Combined score** |
| --- | --- | --- | --- | --- | --- |
| 1 | synaptic transmission, GABAergic (GO:0051932) | 0.00001235 | 0.0002028 | 555.03 | 6273.06 |
| 2 | inhibitory synapse assembly (GO:1904862) | 0.00001235 | 0.0002028 | 555.03 | 6273.06 |
| 3 | membrane depolarization during action potential (GO:0086010) | 2.326e-7 | 0.00001337 | 372.06 | 5682.76 |
| 4 | membrane depolarization (GO:0051899) | 4.430e-7 | 0.00001698 | 294.99 | 4315.59 |
| 5 | anterograde dendritic transport (GO:0098937) | 0.002498 | 0.01306 | 555.17 | 3326.77 |

GO: Molecular Function

| **Index** | **Name** | **P-value** | **Adjusted p-value** | **Odds Ratio** | **Combined score** |
| --- | --- | --- | --- | --- | --- |
| 1 | benzodiazepine receptor activity (GO:0008503) | 0.00001010 | 0.0001049 | 624.44 | 7182.69 |
| 2 | voltage-gated sodium channel activity (GO:0005248) | 1.379e-7 | 0.000004689 | 450.47 | 7115.97 |
| 3 | extracellular ligand-gated ion channel activity (GO:0005230) | 0.00001235 | 0.0001049 | 555.03 | 6273.06 |
| 4 | GABA-gated chloride ion channel activity (GO:0022851) | 0.00001750 | 0.0001157 | 454.07 | 4973.59 |
| 5 | inhibitory extracellular ligand-gated ion channel activity (GO:0005237) | 0.00002041 | 0.0001157 | 416.21 | 4494.85 |

GO: Cellular Component

| **Index** | **Name** | **P-value** | **Adjusted p-value** | **Odds Ratio** | **Combined score** |
| --- | --- | --- | --- | --- | --- |
| 1 | voltage-gated sodium channel complex (GO:0001518) | 6.098e-8 | 0.000001341 | 611.51 | 10158.88 |
| 2 | sodium channel complex (GO:0034706) | 2.058e-7 | 0.000002264 | 388.99 | 5988.96 |
| 3 | GABA-A receptor complex (GO:1902711) | 0.00003830 | 0.0001685 | 293.72 | 2987.15 |
| 4 | neuron to neuron synapse (GO:0098984) | 0.003994 | 0.01098 | 317.19 | 1751.86 |
| 5 | dendrite membrane (GO:0032590) | 0.00009070 | 0.0003325 | 184.84 | 1720.52 |

Polyneuropathy ([HP:0001271](https://hpo.jax.org/app/browse/term/HP:0001271))

KEGG Pathway

| **Index** | **Name** | **P-value** | **Adjusted p-value** | **Odds Ratio** | **Combined score** |
| --- | --- | --- | --- | --- | --- |
| 1 | Peroxisome | 0.000007807 | 0.0001561 | 108.02 | 1270.32 |
| 2 | Ubiquinone and other terpenoid-quinone biosynthesis | 0.005488 | 0.03658 | 222.00 | 1155.57 |
| 3 | Spinocerebellar ataxia | 0.002200 | 0.02200 | 35.19 | 215.36 |
| 4 | Nucleotide excision repair | 0.02326 | 0.06795 | 48.17 | 181.19 |
| 5 | Cocaine addiction | 0.02424 | 0.06795 | 46.16 | 171.72 |

GO: Biological Process

| **Index** | **Name** | **P-value** | **Adjusted p-value** | **Odds Ratio** | **Combined score** |
| --- | --- | --- | --- | --- | --- |
| 1 | peroxisome organization (GO:0007031) | 2.058e-7 | 0.00001405 | 388.99 | 5988.96 |
| 2 | protein import into peroxisome matrix (GO:0016558) | 0.00001481 | 0.0004776 | 499.50 | 5554.53 |
| 3 | peroxisomal membrane transport (GO:0015919) | 2.616e-7 | 0.00001405 | 356.54 | 5403.81 |
| 4 | protein targeting to peroxisome (GO:0006625) | 3.266e-7 | 0.00001405 | 329.08 | 4914.57 |
| 5 | double-strand break repair via classical nonhomologous end joining (GO:0097680) | 0.002997 | 0.02974 | 444.11 | 2580.41 |

GO: Molecular Function

| **Index** | **Name** | **P-value** | **Adjusted p-value** | **Odds Ratio** | **Combined score** |
| --- | --- | --- | --- | --- | --- |
| 1 | opioid receptor binding (GO:0031628) | 0.002498 | 0.02156 | 555.17 | 3326.77 |
| 2 | peroxisome targeting sequence binding (GO:0000268) | 0.002997 | 0.02156 | 444.11 | 2580.41 |
| 3 | alpha-actinin binding (GO:0051393) | 0.00006174 | 0.001482 | 226.91 | 2199.35 |
| 4 | adenylate cyclase inhibiting G protein-coupled glutamate receptor activity (GO:0001640) | 0.004492 | 0.02156 | 277.53 | 1500.17 |
| 5 | G protein-coupled glutamate receptor activity (GO:0098988) | 0.004492 | 0.02156 | 277.53 | 1500.17 |

GO: Cellular Component

| **Index** | **Name** | **P-value** | **Adjusted p-value** | **Odds Ratio** | **Combined score** |
| --- | --- | --- | --- | --- | --- |
| 1 | integral component of peroxisomal membrane (GO:0005779) | 0.00001750 | 0.0001531 | 454.07 | 4973.59 |
| 2 | intrinsic component of peroxisomal membrane (GO:0031231) | 0.00002041 | 0.0001531 | 416.21 | 4494.85 |
| 3 | microbody membrane (GO:0031903) | 0.000001964 | 0.00003121 | 174.41 | 2291.88 |
| 4 | peroxisomal membrane (GO:0005778) | 0.000002081 | 0.00003121 | 170.91 | 2236.02 |
| 5 | G protein-coupled receptor dimeric complex (GO:0038037) | 0.003495 | 0.01498 | 370.07 | 2093.27 |

Rigidity ([HP:0002063](https://hpo.jax.org/app/browse/term/HP:0002063))

KEGG Pathway

| **Index** | **Name** | **P-value** | **Adjusted p-value** | **Odds Ratio** | **Combined score** |
| --- | --- | --- | --- | --- | --- |
| 1 | Taste transduction | 0.0008042 | 0.01608 | 59.24 | 422.16 |
| 2 | Dopaminergic synapse | 0.001879 | 0.01879 | 38.19 | 239.73 |
| 3 | Nicotine addiction | 0.01983 | 0.1070 | 56.84 | 222.86 |
| 4 | Type II diabetes mellitus | 0.02277 | 0.1070 | 49.25 | 186.27 |
| 5 | Cortisol synthesis and secretion | 0.03204 | 0.1070 | 34.59 | 119.03 |

GO: Biological Process

| **Index** | **Name** | **P-value** | **Adjusted p-value** | **Odds Ratio** | **Combined score** |
| --- | --- | --- | --- | --- | --- |
| 1 | membrane depolarization (GO:0051899) | 2.132e-15 | 1.034e-13 | 1151.77 | 38908.86 |
| 2 | neuronal action potential (GO:0019228) | 2.480e-13 | 6.014e-12 | 1174.88 | 34101.37 |
| 3 | membrane depolarization during action potential (GO:0086010) | 6.190e-13 | 1.001e-11 | 950.90 | 26730.60 |
| 4 | inorganic cation transmembrane transport (GO:0098662) | 1.473e-16 | 1.429e-14 | 669.91 | 24420.91 |
| 5 | action potential (GO:0001508) | 1.912e-14 | 6.182e-13 | 767.35 | 24238.98 |

GO: Molecular Function

| **Index** | **Name** | **P-value** | **Adjusted p-value** | **Odds Ratio** | **Combined score** |
| --- | --- | --- | --- | --- | --- |
| 1 | voltage-gated sodium channel activity (GO:0005248) | 2.480e-13 | 4.960e-12 | 1174.88 | 34101.37 |
| 2 | sodium channel activity (GO:0005272) | 4.712e-12 | 4.712e-11 | 604.76 | 15772.67 |
| 3 | voltage-gated cation channel activity (GO:0022843) | 5.976e-10 | 3.984e-9 | 216.28 | 4593.43 |
| 4 | delayed rectifier potassium channel activity (GO:0005251) | 4.872e-7 | 0.000001624 | 285.14 | 4144.43 |
| 5 | voltage-gated sodium channel activity involved in cardiac muscle cell action potential (GO:0086006) | 0.002498 | 0.007136 | 555.17 | 3326.77 |
| 6 | voltage-gated potassium channel activity (GO:0005249) | 5.408e-8 | 2.704e-7 | 170.19 | 2847.72 |
| 7 | potassium channel activity (GO:0005267) | 8.246e-8 | 3.298e-7 | 152.51 | 2487.64 |
| 8 | sodium channel inhibitor activity (GO:0019871) | 0.003994 | 0.009984 | 317.19 | 1751.86 |
| 9 | high voltage-gated calcium channel activity (GO:0008331) | 0.004990 | 0.01088 | 246.68 | 1307.49 |
| 10 | outward rectifier potassium channel activity (GO:0015271) | 0.005488 | 0.01088 | 222.00 | 1155.57 |
| 11 | voltage-gated potassium channel activity involved in ventricular cardiac muscle cell action potential repolarization (GO:1902282) | 0.005985 | 0.01088 | 201.81 | 1032.95 |
| 12 | voltage-gated potassium channel activity involved in cardiac muscle cell action potential repolarization (GO:0086008) | 0.007476 | 0.01246 | 158.54 | 776.21 |
| 13 | potassium ion transmembrane transporter activity (GO:0015079) | 0.01589 | 0.02294 | 71.54 | 296.32 |
| 14 | ion channel inhibitor activity (GO:0008200) | 0.01835 | 0.02294 | 61.59 | 246.23 |
| 15 | voltage-gated calcium channel activity (GO:0005245) | 0.01835 | 0.02294 | 61.59 | 246.23 |
| 16 | sodium channel regulator activity (GO:0017080) | 0.01835 | 0.02294 | 61.59 | 246.23 |
| 17 | amyloid-beta binding (GO:0001540) | 0.03930 | 0.04580 | 28.00 | 90.64 |
| 18 | calcium channel activity (GO:0005262) | 0.04122 | 0.04580 | 26.65 | 84.98 |
| 19 | cation channel activity (GO:0005261) | 0.04794 | 0.05047 | 22.79 | 69.22 |
| 20 | protein heterodimerization activity (GO:0046982) | 0.09014 | 0.09014 | 11.77 | 28.31 |

GO: Cellular Component

| **Index** | **Name** | **P-value** | **Adjusted p-value** | **Odds Ratio** | **Combined score** |
| --- | --- | --- | --- | --- | --- |
| 1 | voltage-gated sodium channel complex (GO:0001518) | 5.833e-14 | 8.167e-13 | 1664.83 | 50731.79 |
| 2 | sodium channel complex (GO:0034706) | 5.000e-13 | 3.500e-12 | 998.50 | 28281.59 |
| 3 | axon (GO:0030424) | 1.211e-12 | 5.652e-12 | 234.43 | 6432.76 |
| 4 | voltage-gated potassium channel complex (GO:0008076) | 3.373e-8 | 7.870e-8 | 192.47 | 3311.50 |
| 5 | potassium channel complex (GO:0034705) | 4.893e-8 | 9.785e-8 | 174.68 | 2940.45 |
| 6 | integral component of plasma membrane (GO:0005887) | 5.182e-10 | 1.814e-9 | 115.51 | 2469.57 |
| 7 | neuron projection (GO:0043005) | 1.379e-9 | 3.863e-9 | 82.63 | 1685.72 |
| 8 | dendrite membrane (GO:0032590) | 0.01441 | 0.02155 | 79.21 | 335.86 |
| 9 | intercalated disc (GO:0014704) | 0.01540 | 0.02155 | 73.93 | 308.54 |
| 10 | cell-cell contact zone (GO:0044291) | 0.02326 | 0.02960 | 48.17 | 181.19 |
| 11 | cation channel complex (GO:0034703) | 0.03591 | 0.04190 | 30.74 | 102.25 |
| 12 | dendrite (GO:0030425) | 0.007607 | 0.01331 | 18.40 | 89.75 |
| 13 | asymmetric synapse (GO:0032279) | 0.06456 | 0.06691 | 16.72 | 45.80 |
| 14 | postsynaptic density (GO:0014069) | 0.06691 | 0.06691 | 16.10 | 43.54 |

Seizure ([HP:0001250](https://hpo.jax.org/app/browse/term/HP:0001250))

GO: Biological Process

| **Index** | **Name** | **P-value** | **Adjusted p-value** | **Odds Ratio** | **Combined score** |
| --- | --- | --- | --- | --- | --- |
| 1 | cilium assembly (GO:0060271) | 7.884e-19 | 1.419e-17 | 196860.00 | 8205971.98 |
| 2 | ciliary basal body-plasma membrane docking (GO:0097711) | 8.554e-18 | 7.699e-17 | 915.08 | 35962.78 |
| 3 | cilium organization (GO:0044782) | 2.747e-17 | 1.648e-16 | 812.51 | 30983.64 |
| 4 | plasma membrane bounded cell projection assembly (GO:0120031) | 1.681e-16 | 7.564e-16 | 659.81 | 23965.68 |
| 5 | protein localization to ciliary transition zone (GO:1904491) | 0.000003371 | 0.000008669 | 1249.13 | 15739.23 |
| 6 | organelle assembly (GO:0070925) | 7.972e-15 | 2.870e-14 | 423.48 | 13747.25 |
| 7 | smoothened signaling pathway (GO:0007224) | 1.637e-9 | 4.912e-9 | 429.23 | 8683.34 |
| 8 | retinal rod cell development (GO:0046548) | 0.003495 | 0.006291 | 370.07 | 2093.27 |
| 9 | non-motile cilium assembly (GO:1905515) | 0.00009715 | 0.0002186 | 178.23 | 1646.74 |
| 10 | retinal rod cell differentiation (GO:0060221) | 0.004492 | 0.007350 | 277.53 | 1500.17 |
| 11 | protein localization to cilium (GO:0061512) | 0.0001327 | 0.0002654 | 151.19 | 1349.73 |
| 12 | eye photoreceptor cell development (GO:0042462) | 0.007476 | 0.01121 | 158.54 | 776.21 |
| 13 | negative regulation of G protein-coupled receptor signaling pathway (GO:0045744) | 0.01144 | 0.01584 | 100.85 | 450.83 |
| 14 | regulation of protein localization (GO:0032880) | 0.03252 | 0.04181 | 34.06 | 116.68 |
| 15 | regulation of G protein-coupled receptor signaling pathway (GO:0008277) | 0.04026 | 0.04831 | 27.31 | 87.73 |
| 16 | microtubule cytoskeleton organization involved in mitosis (GO:1902850) | 0.06220 | 0.06998 | 17.38 | 48.27 |
| 17 | mitotic spindle organization (GO:0007052) | 0.07580 | 0.08026 | 14.13 | 36.44 |
| 18 | negative regulation of signal transduction (GO:0009968) | 0.1258 | 0.1258 | 8.24 | 17.08 |

GO: Molecular Function

| **Index** | **Name** | **P-value** | **Adjusted p-value** | **Odds Ratio** | **Combined score** |
| --- | --- | --- | --- | --- | --- |
| 1 | gamma-tubulin binding (GO:0043015) | 0.01144 | 0.02289 | 100.85 | 450.83 |
| 2 | tubulin binding (GO:0015631) | 0.1433 | 0.1433 | 7.15 | 13.88 |

GO: Cellular Component

| **Index** | **Name** | **P-value** | **Adjusted p-value** | **Odds Ratio** | **Combined score** |
| --- | --- | --- | --- | --- | --- |
| 1 | ciliary membrane (GO:0060170) | 0.0001038 | 0.0002595 | 172.08 | 1578.44 |
| 2 | cilium (GO:0005929) | 5.378e-8 | 2.689e-7 | 85.16 | 1425.50 |
| 3 | cell projection membrane (GO:0031253) | 0.0009195 | 0.001533 | 55.28 | 386.48 |
| 4 | bounding membrane of organelle (GO:0098588) | 0.05389 | 0.06736 | 6.28 | 18.35 |
| 5 | cell-cell junction (GO:0005911) | 0.1276 | 0.1276 | 8.12 | 16.71 |

Skeletal muscle atrophy ([HP:0003202](https://hpo.jax.org/app/browse/term/HP:0003202))

GO: Biological Process

| **Index** | **Name** | **P-value** | **Adjusted p-value** | **Odds Ratio** | **Combined score** |
| --- | --- | --- | --- | --- | --- |
| 1 | cilium assembly (GO:0060271) | 5.098e-16 | 3.772e-14 | 580.87 | 20453.89 |
| 2 | pigment granule transport (GO:0051904) | 7.315e-8 | 9.022e-7 | 570.71 | 9377.24 |
| 3 | melanosome transport (GO:0032402) | 7.315e-8 | 9.022e-7 | 570.71 | 9377.24 |
| 4 | establishment of melanosome localization (GO:0032401) | 8.685e-8 | 9.181e-7 | 535.02 | 8698.92 |
| 5 | melanosome localization (GO:0032400) | 1.191e-7 | 0.000001102 | 475.52 | 7581.26 |

GO: Molecular Function

| **Index** | **Name** | **P-value** | **Adjusted p-value** | **Odds Ratio** | **Combined score** |
| --- | --- | --- | --- | --- | --- |
| 1 | patched binding (GO:0005113) | 0.003495 | 0.006990 | 370.07 | 2093.27 |
| 2 | RNA polymerase II-specific DNA-binding transcription factor binding (GO:0061629) | 1.780e-8 | 8.402e-8 | 107.05 | 1910.30 |
| 3 | DNA-binding transcription factor binding (GO:0140297) | 2.801e-8 | 8.402e-8 | 97.47 | 1695.14 |
| 4 | phosphatidylinositol-3-phosphate binding (GO:0032266) | 0.01933 | 0.02900 | 58.34 | 230.20 |
| 5 | phosphatidylinositol phosphate binding (GO:1901981) | 0.04986 | 0.05983 | 21.88 | 65.61 |

GO: Cellular Component

| **Index** | **Name** | **P-value** | **Adjusted p-value** | **Odds Ratio** | **Combined score** |
| --- | --- | --- | --- | --- | --- |
| 1 | motile cilium (GO:0031514) | 0.03349 | 0.06698 | 33.04 | 112.22 |
| 2 | neuron projection (GO:0043005) | 0.02994 | 0.06698 | 8.77 | 30.77 |
| 3 | cilium (GO:0005929) | 0.1124 | 0.1276 | 9.30 | 20.33 |
| 4 | cell-cell junction (GO:0005911) | 0.1276 | 0.1276 | 8.12 | 16.71 |

Somatic sensory dysfunction ([HP:0003474](https://hpo.jax.org/app/browse/term/HP:0003474))

KEGG Pathway

| **Index** | **Name** | **P-value** | **Adjusted p-value** | **Odds Ratio** | **Combined score** |
| --- | --- | --- | --- | --- | --- |
| 1 | Non-small cell lung cancer | 0.03543 | 0.1299 | 31.17 | 104.12 |
| 2 | Amyotrophic lateral sclerosis | 0.01350 | 0.1228 | 13.56 | 58.36 |
| 3 | Dopaminergic synapse | 0.06409 | 0.1570 | 16.84 | 46.28 |
| 4 | Cell adhesion molecules | 0.07160 | 0.1570 | 15.00 | 39.55 |
| 5 | Pathways of neurodegeneration | 0.02233 | 0.1228 | 10.32 | 39.22 |

GO: Biological Process

| **Index** | **Name** | **P-value** | **Adjusted p-value** | **Odds Ratio** | **Combined score** |
| --- | --- | --- | --- | --- | --- |
| 1 | synaptic vesicle transport (GO:0048489) | 0.00001235 | 0.0006828 | 555.03 | 6273.06 |
| 2 | synaptic vesicle localization (GO:0097479) | 0.00001750 | 0.0006828 | 454.07 | 4973.59 |
| 3 | endoplasmic reticulum tubular network organization (GO:0071786) | 0.00002354 | 0.0006828 | 384.17 | 4094.01 |
| 4 | anterograde dendritic transport (GO:0098937) | 0.002498 | 0.01672 | 555.17 | 3326.77 |
| 5 | anterograde dendritic transport of neurotransmitter receptor complex (GO:0098971) | 0.002498 | 0.01672 | 555.17 | 3326.77 |

GO: Molecular Function

| **Index** | **Name** | **P-value** | **Adjusted p-value** | **Odds Ratio** | **Combined score** |
| --- | --- | --- | --- | --- | --- |
| 1 | olfactory receptor binding (GO:0031849) | 0.002997 | 0.01299 | 444.11 | 2580.41 |
| 2 | microtubule binding (GO:0008017) | 0.0001741 | 0.002263 | 36.98 | 320.12 |
| 3 | alpha-tubulin binding (GO:0043014) | 0.01540 | 0.03443 | 73.93 | 308.54 |
| 4 | beta-tubulin binding (GO:0048487) | 0.01589 | 0.03443 | 71.54 | 296.32 |
| 5 | tubulin binding (GO:0015631) | 0.0003968 | 0.002579 | 27.75 | 217.37 |

GO: Cellular Component

| **Index** | **Name** | **P-value** | **Adjusted p-value** | **Odds Ratio** | **Combined score** |
| --- | --- | --- | --- | --- | --- |
| 1 | endoplasmic reticulum tubular network membrane (GO:0098826) | 0.002498 | 0.01748 | 555.17 | 3326.77 |
| 2 | endoplasmic reticulum tubular network (GO:0071782) | 0.00005661 | 0.001585 | 237.73 | 2324.81 |
| 3 | Golgi cis cisterna (GO:0000137) | 0.01243 | 0.03868 | 92.44 | 405.55 |
| 4 | intrinsic component of endoplasmic reticulum membrane (GO:0031227) | 0.001583 | 0.01748 | 41.75 | 269.21 |
| 5 | integral component of endoplasmic reticulum membrane (GO:0030176) | 0.002170 | 0.01748 | 35.45 | 217.39 |

Spasticity ([HP:0001257](https://hpo.jax.org/app/browse/term/HP:0001257))

KEGG Pathway

| **Index** | **Name** | **P-value** | **Adjusted p-value** | **Odds Ratio** | **Combined score** |
| --- | --- | --- | --- | --- | --- |
| 1 | Lysosome | 0.00002973 | 0.0003271 | 68.11 | 709.91 |
| 2 | Non-small cell lung cancer | 0.03543 | 0.07795 | 31.17 | 104.12 |
| 3 | Endocytosis | 0.006657 | 0.03661 | 19.74 | 98.94 |
| 4 | Amyotrophic lateral sclerosis | 0.01350 | 0.04949 | 13.56 | 58.36 |
| 5 | Dopaminergic synapse | 0.06409 | 0.1175 | 16.84 | 46.28 |

GO: Biological Process

| **Index** | **Name** | **P-value** | **Adjusted p-value** | **Odds Ratio** | **Combined score** |
| --- | --- | --- | --- | --- | --- |
| 1 | synaptic vesicle transport (GO:0048489) | 0.00001235 | 0.0002012 | 555.03 | 6273.06 |
| 2 | synaptic vesicle localization (GO:0097479) | 0.00001750 | 0.0002012 | 454.07 | 4973.59 |
| 3 | anterograde dendritic transport (GO:0098937) | 0.002498 | 0.01413 | 555.17 | 3326.77 |
| 4 | anterograde dendritic transport of neurotransmitter receptor complex (GO:0098971) | 0.002498 | 0.01413 | 555.17 | 3326.77 |
| 5 | establishment of vesicle localization (GO:0051650) | 0.00003428 | 0.0002628 | 312.09 | 3208.63 |

GO: Molecular Function

| **Index** | **Name** | **P-value** | **Adjusted p-value** | **Odds Ratio** | **Combined score** |
| --- | --- | --- | --- | --- | --- |
| 1 | olfactory receptor binding (GO:0031849) | 0.002997 | 0.01512 | 444.11 | 2580.41 |
| 2 | CD4 receptor binding (GO:0042609) | 0.003994 | 0.01512 | 317.19 | 1751.86 |
| 3 | microtubule motor activity (GO:0003777) | 0.02766 | 0.04336 | 40.27 | 144.50 |
| 4 | motor activity (GO:0003774) | 0.03252 | 0.04336 | 34.06 | 116.68 |
| 5 | microtubule binding (GO:0008017) | 0.005670 | 0.01512 | 21.48 | 111.10 |

GO: Cellular Component

| **Index** | **Name** | **P-value** | **Adjusted p-value** | **Odds Ratio** | **Combined score** |
| --- | --- | --- | --- | --- | --- |
| 1 | endosome lumen (GO:0031904) | 2.326e-7 | 0.000006745 | 372.06 | 5682.76 |
| 2 | trans-Golgi network membrane (GO:0032588) | 0.00001377 | 0.0001996 | 88.81 | 994.11 |
| 3 | trans-Golgi network transport vesicle (GO:0030140) | 0.007476 | 0.03097 | 158.54 | 776.21 |
| 4 | endoplasmic reticulum tubular network (GO:0071782) | 0.01144 | 0.04148 | 100.85 | 450.83 |
| 5 | trans-Golgi network (GO:0005802) | 0.0001901 | 0.001837 | 35.87 | 307.37 |

Tremor ([HP:0001337](https://hpo.jax.org/app/browse/term/HP:0001337))

GO: Biological Process

| **Index** | **Name** | **P-value** | **Adjusted p-value** | **Odds Ratio** | **Combined score** |
| --- | --- | --- | --- | --- | --- |
| 1 | ciliary basal body-plasma membrane docking (GO:0097711) | 3.584e-24 | 1.147e-22 | 199050.00 | 10745825.19 |
| 2 | cilium assembly (GO:0060271) | 7.884e-19 | 1.261e-17 | 196860.00 | 8205971.98 |
| 3 | protein localization to ciliary transition zone (GO:1904491) | 0.000003371 | 0.00001541 | 1249.13 | 15739.23 |
| 4 | cilium organization (GO:0044782) | 1.113e-14 | 1.187e-13 | 359.45 | 11549.06 |
| 5 | plasma membrane bounded cell projection assembly (GO:0120031) | 5.535e-14 | 4.428e-13 | 292.15 | 8917.85 |

GO: Cellular Component

| **Index** | **Name** | **P-value** | **Adjusted p-value** | **Odds Ratio** | **Combined score** |
| --- | --- | --- | --- | --- | --- |
| 1 | cilium (GO:0005929) | 5.245e-10 | 4.196e-9 | 128.31 | 2741.69 |
| 2 | ciliary membrane (GO:0060170) | 0.0001038 | 0.0004153 | 172.08 | 1578.44 |
| 3 | cell projection membrane (GO:0031253) | 0.0009195 | 0.002452 | 55.28 | 386.48 |
| 4 | motile cilium (GO:0031514) | 0.03349 | 0.05359 | 33.04 | 112.22 |
| 5 | cell-cell junction (GO:0005911) | 0.007662 | 0.01532 | 18.33 | 89.29 |
