## Supplementary Table 1-3 for "Long COVID: G Protein-Coupled Receptors (GPCRs) responsible for persistent post-COVID symptoms": Supplementary File 3- hub genes.docx

**Immunology-autoimmunity**

| **Symptom (HPO ID)** | **Hub Genes (ranked)** |
| --- | --- |
| Anaphylactic shock ([HP:0100845](https://hpo.jax.org/app/browse/term/HP:0100845)) | CBL |
|  | KIT |
|  | KITLG |
|  | GRB2 |
|  | EPOR |
|  | PTPN11 |
|  | NRAS |
|  | PIK3R1 |
|  | HRAS |
|  | PTPN6 |
| Antinuclear antibody positivity ([HP:0003493](https://hpo.jax.org/app/browse/term/HP:0003493)) | CTLA4 |
|  | IL2RA |
|  | FAS |
|  | FASLG |
|  | FCGR2B |
|  | RIPK1 |
|  | IL12RB1 |
|  | MS4A1 |
|  | PTPN22 |
|  | ACP5 |
| Anti-thyroid peroxidase antibody positivity ([HP:0025379](https://hpo.jax.org/app/browse/term/HP:0025379)) | RASGRP1 |
|  | HRAS |
|  | KRAS |
|  | NRAS |
|  | RAP1B |
|  | RAP1A |
|  | PLCG1 |
|  | PRKCQ |
|  | SKAP1 |
|  | DGKZ |
| Anti-thyroglobulin antibody positivity ([HP:0032069](https://hpo.jax.org/app/browse/term/HP:0032069)) | - |
| Lymphadenopathy ([HP:0002716](https://hpo.jax.org/app/browse/term/HP:0002716)) | CTLA4 |
|  | CD19 |
|  | IL2RA |
|  | STAT5B |
|  | PRF1 |
|  | FAS |
|  | ZAP70 |
|  | RAG1 |
|  | FASLG |
|  | SYK |
| Lymphopenia ([HP:0001888](https://hpo.jax.org/app/browse/term/HP:0001888)) | CD19 |
|  | CTLA4 |
|  | ZAP70 |
|  | CD3E |
|  | RAG1 |
|  | RAG2 |
|  | CD3D |
|  | IL2RA |
|  | CD3G |
|  | IKZF1 |

**Dermatographical-finding**

| **Symptom (HPO ID)** | **Hub Genes (ranked)** |
| --- | --- |
| Alopecia ([HP:0001596](https://hpo.jax.org/app/browse/term/HP:0001596)) | HRAS |
|  | KRAS |
|  | NRAS |
|  | GNA11 |
|  | PRKAR1A |
|  | PRKACA |
|  | CTLA4 |
|  | BTK |
|  | ABCA12 |
|  | CERS3 |
| Dermatographic urticaria ([HP:0011971](https://hpo.jax.org/app/browse/term/HP:0011971)) | - |
| Flushing ([HP:0031284](https://hpo.jax.org/app/browse/term/HP:0031284)) | NF1 |
|  | SDHC |
|  | RET |
|  | MAX |
|  | KIF1B |
|  | SDHAF2 |
|  | KIT |
|  | DNMT3A |
|  | SETD2 |
|  | NSD1 |
| Fragile nails ([HP:0001808](https://hpo.jax.org/app/browse/term/HP:0001808)) | COL17A1 |
|  | HRAS |
|  | ITGB4 |
|  | DSP |
|  | EFNB1 |
|  | DLX3 |
|  | ERCC2 |
|  | GTF2H5 |
|  | CAMK2G |
|  | MSX1 |
| Hyperhidrosis ([HP:0000975](https://hpo.jax.org/app/browse/term/HP:0000975)) | NF1 |
|  | SDHC |
|  | RET |
|  | SDHAF2 |
|  | MAX |
|  | KIF1B |
|  | MEN1 |
|  | TERT |
|  | HNF4A |
|  | YY1 |
| Inflammatory abnormality of the skin ([HP:0011123](https://hpo.jax.org/app/browse/term/HP:0011123)) | ZAP70 |
|  | CD3E |
|  | CTLA4 |
|  | CD3D |
|  | CD3G |
|  | RAG1 |
|  | CD79B |
|  | RAG2 |
|  | CD79A |
|  | IL2RA |
| Petechiae ([HP:0000967](https://hpo.jax.org/app/browse/term/HP:0000967)) | ITGA2B |
|  | GATA1 |
|  | GP9 |
|  | STAT5B |
|  | PRF1 |
|  | STXBP2 |
|  | GP1BB |
|  | GFI1B |
|  | RAB27A |
|  | FERMT3 |
| Pruritus ([HP:0000989](https://hpo.jax.org/app/browse/term/HP:0000989)) | ABCA12 |
|  | PNPLA1 |
|  | CYP4F22 |
|  | SDR9C7 |
|  | LIPN |
|  | CERS3 |
|  | ALOXE3 |
|  | ALOX12B |
|  | RAG1 |
|  | KIT |
| Scaling skin ([HP:0040189](https://hpo.jax.org/app/browse/term/HP:0040189)) | CDSN |
|  | KIT |
|  | IL2RA |
|  | ZMPSTE24 |
|  | ADAM17 |
|  | RASA1 |
|  | KRT74 |
|  | CHST8 |
|  | CARD14 |
|  | CASP14 |
| Skin rash ([HP:0000988](https://hpo.jax.org/app/browse/term/HP:0000988)) | CTLA4 |
|  | ZAP70 |
|  | CD79B |
|  | CD79A |
|  | BTK |
|  | SYK |
|  | FCGR2B |
|  | RAG2 |
|  | RAG1 |
|  | PTPN22 |

**Reproductive-Genitourinary-Endocrine-Metabolism**

| **Symptom (HPO ID)** | **Hub Genes (ranked)** |
| --- | --- |
| Abnormal female reproductive system physiology ([HP:0030012](https://hpo.jax.org/app/browse/term/HP:0030012)) | GNRH1 |
|  | GNRHR |
|  | KISS1R |
|  | KAL1 |
|  | TAC3 |
|  | FGF8 |
|  | FEZF1 |
|  | CHD7 |
|  | KISS1 |
|  | IL17RD |
| Decreased glomerular filtration rate ([HP:0012213](https://hpo.jax.org/app/browse/term/HP:0012213)) | BSND |
|  | CLCNKB |
|  | CLCNKA |
|  | CASR |
|  | SLC22A12 |
|  | SLC2A9 |
|  | G6PC |
|  | ITGA3 |
|  | SLC7A7 |
|  | HGD |
| Diabetes mellitus ([HP:0000819](https://hpo.jax.org/app/browse/term/HP:0000819)) | CRX |
|  | GUCA1B |
|  | PRPH2 |
|  | RDH12 |
|  | RPGR |
|  | NR2E3 |
|  | TULP1 |
|  | SPATA7 |
|  | PRPF3 |
|  | PDE6G |
| Abnormality of the digestive system ([HP:0025031](https://hpo.jax.org/app/browse/term/HP:0025031)) | DRC1 |
|  | CCDC65 |
|  | HYDIN |
|  | RSPH4A |
|  | RSPH1 |
|  | CCDC103 |
|  | HEATR2 |
|  | DYX1C1 |
|  | CCDC40 |
|  | CCDC39 |
| Edema ([HP:0000969](https://hpo.jax.org/app/browse/term/HP:0000969)) | NR2E3 |
|  | RHO |
|  | NRL |
|  | RLBP1 |
|  | PRPF8 |
|  | PRPH2 |
|  | PDE6G |
|  | RP9 |
|  | KRAS |
|  | NRAS |
| Female sexual dysfunction ([HP:0030014](https://hpo.jax.org/app/browse/term/HP:0030014)) | KAL1 |
|  | FGF8 |
|  | GNRH1 |
|  | TAC3 |
|  | KISS1R |
|  | GNRHR |
|  | FEZF1 |
|  | CHD7 |
|  | IL17RD |
|  | KISS1 |
| Fever ([HP:0001945](https://hpo.jax.org/app/browse/term/HP:0001945)) | CTLA4 |
|  | CD3E |
|  | CD79A |
|  | IKZF1 |
|  | CD79B |
|  | RAG1 |
|  | RAG2 |
|  | CD3D |
|  | SYK |
|  | KIT |
| Heat intolerance ([HP:0002046](https://hpo.jax.org/app/browse/term/HP:0002046)) | EDAR |
|  | OCA2 |
|  | EDA |
|  | EDARADD |
|  | UBE3A |
|  | POFUT1 |
|  | ITPR2 |
|  | CLDN10 |
|  | HNRNPK |
|  | LBX1 |
| Hypothermia ([HP:0002045](https://hpo.jax.org/app/browse/term/HP:0002045)) | TSHB |
|  | TG |
|  | TPO |
|  | TSHR |
|  | POU1F1 |
|  | PROP1 |
|  | LHX4 |
|  | SLC5A5 |
|  | LHX3 |
|  | SUCLG1 |
| Irregular menstruation ([HP:0000858](https://hpo.jax.org/app/browse/term/HP:0000858)) | PRKAR1A |
|  | PRKACA |
|  | PHKB |
|  | PDE11A |
|  | PDE4D |
|  | PHKA2 |
|  | AIP |
|  | MEN1 |
|  | PHKG2 |
|  | GPR101 |
| Low-grade fever ([HP:0011134](https://hpo.jax.org/app/browse/term/HP:0011134)) | - |
| Male sexual dysfunction ([HP:0040307](https://hpo.jax.org/app/browse/term/HP:0040307)) | KAL1 |
|  | FGF8 |
|  | GNRH1 |
|  | TAC3 |
|  | KISS1R |
|  | GNRHR |
|  | FEZF1 |
|  | CHD7 |
|  | IL17RD |
|  | KISS1 |
| Menorrhagia ([HP:0000132](https://hpo.jax.org/app/browse/term/HP:0000132)) | GP9 |
|  | GP1BB |
|  | ITGA2B |
|  | F8 |
|  | BLOC1S5 |
|  | SLFN14 |
|  | BLOC1S3 |
|  | PRKACG |
|  | PRLR |
| Pancreatitis ([HP:0001733](https://hpo.jax.org/app/browse/term/HP:0001733)) | CASR |
|  | CTRC |
|  | PRSS1 |
|  | PRSS3P2 |
|  | CTLA4 |
|  | BCKDHB |
|  | PTPN22 |
|  | BCKDHA |
|  | G6PC |
|  | NFS1 |
| Recurrent fever ([HP:0001954](https://hpo.jax.org/app/browse/term/HP:0001954)) | NLRC4 |
|  | MEFV |
|  | NLRP3 |
|  | NLRP12 |
|  | SYK |
|  | XIAP |
|  | LPIN2 |
|  | PRF1 |
|  | PRKCD |
|  | TMEM173 |
| Renal insufficiency ([HP:0000083](https://hpo.jax.org/app/browse/term/HP:0000083)) | RAD51 |
|  | FANCM |
|  | FANCD2 |
|  | FANCA |
|  | FANCB |
|  | SLX4 |
|  | FANCE |
|  | FANCC |
|  | FANCF |
|  | FANCG |
| Temperature instability ([HP:0005968](https://hpo.jax.org/app/browse/term/HP:0005968)) | ZIC2 |
|  | SIX3 |
|  | SHH |
|  | GLI2 |
|  | FGF8 |
|  | CDON |
|  | NODAL |
|  | SNRPN |
|  | NPAP1 |
|  | DDC |
| Urinary incontinence ([HP:0000020](https://hpo.jax.org/app/browse/term/HP:0000020)) | KIF5A |
|  | NIPA1 |
|  | KIAA0196 |
|  | SPG11 |
|  | RTN2 |
|  | SPAST |
|  | ATL1 |
|  | GJC2 |
|  | ALDH18A1 |
|  | KCNT1 |

**HEENT-ear**

| **Symptom (HPO ID)** | **Hub Genes (ranked)** |
| --- | --- |
| Ear pain ([HP:0030766](https://hpo.jax.org/app/browse/term/HP:0030766)) | SMO |
|  | PIK3CA |
|  | SUFU |
|  | TERT |
|  | KRT6A |
|  | TRAF7 |
| Hearing impairment ([HP:0000365](https://hpo.jax.org/app/browse/term/HP:0000365)) | RPGR |
|  | IMPDH1 |
|  | RDH12 |
|  | SPATA7 |
|  | CRX |
|  | PRPH2 |
|  | TULP1 |
|  | CNGA1 |
|  | NR2E3 |
|  | GUCA1B |
| Hyperacusis ([HP:0010780](https://hpo.jax.org/app/browse/term/HP:0010780)) | - |
| Pulsatile tinnitus ([HP:0008629](https://hpo.jax.org/app/browse/term/HP:0008629)) | MAX |
|  | NF1 |
|  | SDHC |
|  | RET |
|  | SDHAF2 |
|  | KIF1B |
|  | DNMT3A |
|  | DLST |
| Tinnitus ([HP:0000360](https://hpo.jax.org/app/browse/term/HP:0000360)) | NF1 |
|  | MAX |
|  | RET |
|  | KIF1B |
|  | SDHC |
|  | SDHAF2 |
|  | TERT |
|  | CACNA1D |
|  | CYP11B2 |
|  | KCNJ5 |
| Vertigo ([HP:0002321](https://hpo.jax.org/app/browse/term/HP:0002321)) | SCN5A |
|  | SCN2A |
|  | SCN1A |
|  | CACNA1G |
|  | RYR2 |
|  | SCN1B |
|  | CASQ2 |
|  | NF1 |
|  | RET |
|  | SDHC |

**HEENT-ENT**

| **Symptom (HPO ID)** | **Hub Genes (ranked)** |
| --- | --- |
| Abnormality of the pharynx ([HP:0000600](https://hpo.jax.org/app/browse/term/HP:0000600)) | RSPH4A |
|  | HEATR2 |
|  | RSPH1 |
|  | SPAG1 |
|  | DYX1C1 |
|  | C21orf59 |
|  | CCDC65 |
|  | HYDIN |
|  | DNAI2 |
|  | DNAI1 |
| Dysphonia ([HP:0001618](https://hpo.jax.org/app/browse/term/HP:0001618)) | MAX |
|  | SDHC |
|  | NF1 |
|  | KIF1B |
|  | RET |
|  | SDHAF2 |
|  | SYT2 |
|  | CHAT |
|  | SLC18A3 |
|  | MFN2 |
| Nasal congestion ([HP:0001742](https://hpo.jax.org/app/browse/term/HP:0001742)) | CCDC65 |
|  | DRC1 |
|  | HYDIN |
|  | RSPH4A |
|  | RSPH1 |
|  | CCDC103 |
|  | HEATR2 |
|  | DYX1C1 |
|  | DNAI2 |
|  | CCDC40 |
| Rhinitis ([HP:0012384](https://hpo.jax.org/app/browse/term/HP:0012384)) | CCDC65 |
|  | DRC1 |
|  | HYDIN |
|  | RSPH4A |
|  | RSPH1 |
|  | CCDC103 |
|  | HEATR2 |
|  | DYX1C1 |
|  | DNAI2 |
|  | CCDC40 |

**HEENT-eye**

| **Symptom (HPO ID)** | **Hub Genes (ranked)** |
| --- | --- |
| Blindness ([HP:0000618](https://hpo.jax.org/app/browse/term/HP:0000618)) | RPGR |
|  | CRX |
|  | CNGA1 |
|  | GUCA1B |
|  | NR2E3 |
|  | PRPH2 |
|  | RDH12 |
|  | TULP1 |
|  | SPATA7 |
|  | IMPDH1 |
| Blurred vision ([HP:0000622](https://hpo.jax.org/app/browse/term/HP:0000622)) | KCNA1 |
|  | SCN9A |
|  | SCN4A |
|  | ATP1A2 |
|  | SPATA7 |
|  | COL8A2 |
|  | VSX1 |
|  | CLCNKB |
|  | GRHL2 |
|  | RPE65 |
| Conjunctivitis ([HP:0000509](https://hpo.jax.org/app/browse/term/HP:0000509)) | CD19 |
|  | CD79B |
|  | CD79A |
|  | CR2 |
|  | BTK |
|  | TNFRSF13C |
|  | TNFRSF13B |
|  | RAG1 |
|  | RAG2 |
|  | IKZF1 |
| Diplopia ([HP:0000651](https://hpo.jax.org/app/browse/term/HP:0000651)) | DOK7 |
|  | MUSK |
|  | RAPSN |
|  | SCN4A |
|  | CHRNE |
|  | CHRNA1 |
|  | CHRNB1 |
|  | CHRND |
|  | CHAT |
|  | KCNA1 |
| Gaze-evoked nystagmus ([HP:0000640](https://hpo.jax.org/app/browse/term/HP:0000640)) | ATXN10 |
|  | PRKCG |
|  | ATXN1 |
|  | ATXN2 |
|  | ATXN3 |
|  | CACNA1G |
|  | SCN8A |
|  | SCN1A |
|  | KCND3 |
|  | ATP1A2 |
| Keratoconjunctivitis sicca ([HP:0001097](https://hpo.jax.org/app/browse/term/HP:0001097)) | ERCC6 |
|  | GTF2H5 |
|  | ERCC2 |
|  | HLA-DRB1 |
|  | BTNL2 |
|  | COL17A1 |
|  | MEFV |
|  | RNF125 |
|  | GJB6 |
|  | SCN9A |
| Ocular pain ([HP:0200026](https://hpo.jax.org/app/browse/term/HP:0200026)) | COL8A2 |
|  | GNA11 |
|  | GRHL2 |
|  | VSX1 |
|  | GNAQ |
|  | OVOL2 |
|  | CHST6 |
|  | AGBL1 |
|  | SCN9A |
|  | COL17A1 |
| Peripheral visual field loss ([HP:0007994](https://hpo.jax.org/app/browse/term/HP:0007994)) | ABCA4 |
|  | RHO |
|  | CRX |
|  | CNGA1 |
|  | PRPH2 |
|  | RLBP1 |
|  | RPE65 |
|  | SPATA7 |
|  | GUCY2D |
|  | PDE6A |
| Photophobia ([HP:0000613](https://hpo.jax.org/app/browse/term/HP:0000613)) | GUCA1B |
|  | NR2E3 |
|  | PRPH2 |
|  | CRX |
|  | CNGA1 |
|  | NRL |
|  | PDE6G |
|  | RPGR |
|  | RDH12 |
|  | TULP1 |
| Red eye ([HP:0025337](https://hpo.jax.org/app/browse/term/HP:0025337)) | CD19 |
|  | CD79A |
|  | CD79B |
|  | CR2 |
|  | BTK |
|  | TNFRSF13C |
|  | TNFRSF13B |
|  | RAG1 |
|  | RAG2 |
|  | IKZF1 |
| Visual loss ([HP:0000572](https://hpo.jax.org/app/browse/term/HP:0000572)) | USH2A |
|  | CLRN1 |
|  | DFNB31 |
|  | CDH23 |
|  | PCDH15 |
|  | CIB2 |
|  | USH1C |
|  | PDZD7 |
|  | USH1G |
|  | GPR98 |
| Vitreous floaters ([HP:0100832](https://hpo.jax.org/app/browse/term/HP:0100832)) | NDP |
|  | TSPAN12 |
|  | ZNF408 |
|  | CTNNB1 |

**Lab**

| **Symptom (HPO ID)** | **Hub Genes (ranked)** |
| --- | --- |
| Abnormal calcium-phosphate regulating hormone level ([HP:0100530](https://hpo.jax.org/app/browse/term/HP:0100530)) | PTH |
|  | VDR |
|  | KL |
|  | SLC34A3 |
|  | CASR |
|  | SLC34A1 |
|  | RET |
|  | NF1 |
|  | TRPV6 |
|  | SDHC |
| Abnormal circulating protein concentration ([HP:0010876](https://hpo.jax.org/app/browse/term/HP:0010876)) | TCAP |
|  | LDB3 |
|  | CSRP3 |
|  | MYH6 |
|  | MYL2 |
|  | DMD |
|  | ACTN2 |
|  | MYBPC3 |
|  | TNNI3 |
|  | MYH7 |
| Abnormality of fibrinolysis ([HP:0040224](https://hpo.jax.org/app/browse/term/HP:0040224)) | - |
| Decreased circulating calcifediol concentration ([HP:0012053](https://hpo.jax.org/app/browse/term/HP:0012053)) | CYP3A4 |
|  | CYP2C9 |
|  | CYP2C19 |
|  | ABCB1 |
|  | CYP3A7 |
|  | UGT2B7 |
|  | UGT1A8 |
|  | POR |
|  | CYB5A |
|  | CES1 |
| Elevated erythrocyte sedimentation rate ([HP:0003565](https://hpo.jax.org/app/browse/term/HP:0003565)) | PTPN22 |
|  | CTLA4 |
|  | HLA-DRB1 |
|  | CIITA |
|  | MEFV |
|  | IL2RA |
|  | LPIN2 |
|  | NLRP3 |
|  | TMEM173 |
|  | NLRP12 |
| Elevated circulating alkaline phosphatase concentration ([HP:0003155](https://hpo.jax.org/app/browse/term/HP:0003155)) | PIGV |
|  | PIGA |
|  | PIGL |
|  | PIGB |
|  | PIGQ |
|  | PIGY |
|  | PGAP2 |
|  | PIGW |
|  | FGF23 |
|  | PHEX |
| Elevated circulating alanine aminotransferase concentration ([HP:0031964](https://hpo.jax.org/app/browse/term/HP:0031964)) | NR1H4 |
|  | BAAT |
|  | SLC51B |
|  | SLC2A2 |
|  | SLC25A13 |
|  | SCO1 |
|  | DAK |
|  | TTC26 |
|  | KIF12 |
|  | PKHD1 |
| Elevated circulating aspartate aminotransferase concentration ([HP:0031956](https://hpo.jax.org/app/browse/term/HP:0031956)) | NR1H4 |
|  | BAAT |
|  | SLC51B |
|  | SLC2A2 |
|  | COG8 |
|  | TTC26 |
|  | KIF12 |
|  | NFS1 |
|  | PGM1 |
|  | MARS |
| Elevated circulating creatine kinase concentration ([HP:0003236](https://hpo.jax.org/app/browse/term/HP:0003236)) | TCAP |
|  | LDB3 |
|  | CSRP3 |
|  | MYH6 |
|  | DMD |
|  | MYL2 |
|  | ACTN2 |
|  | MYBPC3 |
|  | TNNI3 |
|  | MYH7 |
| Elevated circulating creatinine concentration ([HP:0003259](https://hpo.jax.org/app/browse/term/HP:0003259)) | WDR19 |
|  | CC2D2A |
|  | TTC26 |
|  | INVS |
|  | ADAMTS13 |
|  | CFHR3 |
|  | TREX1 |
|  | FAN1 |
|  | IKBKAP |
|  | HNF1B |
| Elevated circulating C-reactive protein concentration ([HP:0011227](https://hpo.jax.org/app/browse/term/HP:0011227)) | NLRC4 |
|  | MEFV |
|  | NLRP3 |
|  | PSTPIP1 |
|  | PTPN22 |
|  | CTLA4 |
|  | CIITA |
|  | SYK |
|  | PRSS1 |
|  | HLA-DRB1 |
| Increased circulating ferritin concentration ([HP:0003281](https://hpo.jax.org/app/browse/term/HP:0003281)) | XIAP |
|  | SLC25A38 |
|  | STXBP2 |
|  | PRF1 |
|  | GLRX5 |
|  | CPOX |
|  | NLRC4 |
|  | SLC19A1 |
|  | TTC26 |
|  | SLC7A7 |
| Elevated gamma-glutamyltransferase level ([HP:0030948](https://hpo.jax.org/app/browse/term/HP:0030948)) | - |
| Increased circulating interleukin 6 ([HP:0030783](https://hpo.jax.org/app/browse/term/HP:0030783)) | - |
| Increased circulating lactate dehydrogenase concentration ([HP:0025435](https://hpo.jax.org/app/browse/term/HP:0025435)) | ACAD9 |
|  | SLC7A7 |
|  | CPT2 |
|  | SLC25A13 |
|  | RHAG |
|  | SLC19A1 |
|  | RB1 |
|  | HLA-DRB1 |
|  | RHCE |
|  | PIGA |
| Increased circulating NT-proBNP concentration ([HP:0031185](https://hpo.jax.org/app/browse/term/HP:0031185)) | - |
| Elevated circulating thyroid-stimulating hormone concentration ([HP:0002925](https://hpo.jax.org/app/browse/term/HP:0002925)) | SLC5A5 |
|  | TSHR |
|  | TG |
|  | NKX2-1 |
|  | TPO |
|  | SLC16A2 |
|  | SECISBP2 |
|  | SLC35A2 |
|  | PRKAR1A |
|  | CDH23 |
| Hyperglycemia ([HP:0003074](https://hpo.jax.org/app/browse/term/HP:0003074)) | ABCC8 |
|  | INS |
|  | KCNJ11 |
|  | GCK |
|  | PAX4 |
|  | PDX1 |
|  | HNF4A |
|  | HNF1A |
|  | SLC2A2 |
|  | NEUROD1 |
| Hypocalcemia ([HP:0002901](https://hpo.jax.org/app/browse/term/HP:0002901)) | PTH |
|  | CASR |
|  | VDR |
|  | FGF23 |
|  | TNFSF11 |
|  | CYP2R1 |
|  | TBX1 |
|  | DGCR2 |
|  | TNFRSF11A |
|  | GCM2 |
| Hypofibrinogenemia ([HP:0011900](https://hpo.jax.org/app/browse/term/HP:0011900)) | XIAP |
|  | PRF1 |
|  | EPB42 |
|  | ANK1 |
|  | STXBP2 |
|  | SPTA1 |
|  | STAT5B |
|  | AHCY |
|  | NLRC4 |
|  | PRKAR1A |
| Hypoglycemia ([HP:0001943](https://hpo.jax.org/app/browse/term/HP:0001943)) | INS |
|  | GCK |
|  | ABCC8 |
|  | KCNJ11 |
|  | PAX4 |
|  | HNF4A |
|  | HNF1A |
|  | PDX1 |
|  | SLC2A2 |
|  | NEUROD1 |
| Hypophosphatemia ([HP:0002148](https://hpo.jax.org/app/browse/term/HP:0002148)) | FGF23 |
|  | SLC34A3 |
|  | PHEX |
|  | SLC34A1 |
|  | VDR |
|  | DMP1 |
|  | CASR |
|  | ENPP1 |
|  | TNFSF11 |
|  | CLCN5 |
| Thrombocytopenia ([HP:0001873](https://hpo.jax.org/app/browse/term/HP:0001873)) | BRCA1 |
|  | RAD51 |
|  | FANCM |
|  | PALB2 |
|  | SLX4 |
|  | FANCA |
|  | FANCD2 |
|  | FANCC |
|  | FANCE |
|  | FANCL |

**General-pain**

| **Symptom (HPO ID)** | **Hub Genes (ranked)** |
| --- | --- |
| Arthralgia ([HP:0002829](https://hpo.jax.org/app/browse/term/HP:0002829)) | CTLA4 |
|  | CD19 |
|  | CR2 |
|  | TNFRSF13B |
|  | TNFRSF13C |
|  | FAS |
|  | MS4A1 |
|  | IL2RA |
|  | COL2A1 |
|  | COL10A1 |
| Back pain ([HP:0003418](https://hpo.jax.org/app/browse/term/HP:0003418)) | PIK3CA |
|  | KRAS |
|  | SMAD4 |
|  | BRCA1 |
|  | PALB2 |
|  | TERT |
|  | TBX6 |
|  | ALDH18A1 |
|  | RASA1 |
|  | MESP2 |
| Bone pain ([HP:0002653](https://hpo.jax.org/app/browse/term/HP:0002653)) | PHEX |
|  | FGF23 |
|  | SLC34A3 |
|  | DMP1 |
|  | SLC34A1 |
|  | VDR |
|  | ENPP1 |
|  | TNFSF11 |
|  | CLCN5 |
|  | STAT5B |
| Chest pain ([HP:0100749](https://hpo.jax.org/app/browse/term/HP:0100749)) | NF1 |
|  | RET |
|  | SDHC |
|  | MAX |
|  | SDHAF2 |
|  | KIF1B |
|  | SMAD4 |
|  | CTNNB1 |
|  | ACTA2 |
|  | COL3A1 |
| Limb pain ([HP:0009763](https://hpo.jax.org/app/browse/term/HP:0009763)) | COL9A1 |
|  | COL9A3 |
|  | COL2A1 |
|  | COL5A2 |
|  | MATN3 |
|  | SPTLC1 |
|  | BSCL2 |
|  | MFN2 |
|  | CPOX |
|  | ATL1 |
| Myalgia ([HP:0003326](https://hpo.jax.org/app/browse/term/HP:0003326)) | CAV3 |
|  | MYOT |
|  | FKTN |
|  | FKRP |
|  | CAPN3 |
|  | DMD |
|  | TRIM32 |
|  | PYGM |
|  | PHKA1 |
|  | PHKB |
| Pain ([HP:0012531](https://hpo.jax.org/app/browse/term/HP:0012531)) | KRAS |
|  | CTNNB1 |
|  | ESR1 |
|  | PIK3CA |
|  | SMAD4 |
|  | NF1 |
|  | KIT |
|  | RET |
|  | ERBB4 |
|  | GNAQ |
| Pedal edema ([HP:0010741](https://hpo.jax.org/app/browse/term/HP:0010741)) | NKX2-5 |
|  | TBX20 |
|  | MYH6 |
|  | GATA4 |
|  | GATA6 |
|  | SNRPN |
|  | SETD2 |
|  | MAGEL2 |
|  | OCA2 |
|  | RBM8A |

**General-symptom**

| **Symptom (HPO ID)** | **Hub Genes (ranked)** |
| --- | --- |
| Arthritis ([HP:0001369](https://hpo.jax.org/app/browse/term/HP:0001369)) | COL2A1 |
|  | CTLA4 |
|  | SYK |
|  | BTK |
|  | CD79B |
|  | ACAN |
|  | FCGR2B |
|  | COL9A3 |
|  | MATN3 |
|  | ASPN |
| Asthenia ([HP:0025406](https://hpo.jax.org/app/browse/term/HP:0025406)) | - |
| Chills ([HP:0025143](https://hpo.jax.org/app/browse/term/HP:0025143)) | EPB42 |
|  | ANK1 |
|  | SPTA1 |
|  | EPB41 |
|  | NLRP3 |
|  | PKHD1 |
| Constitutional symptom ([HP:0025142](https://hpo.jax.org/app/browse/term/HP:0025142)) | MYH6 |
|  | MYL2 |
|  | MYBPC3 |
|  | TNNI3 |
|  | MYH7 |
|  | SCN5A |
|  | RYR2 |
|  | MYL3 |
|  | LDB3 |
|  | CASQ2 |
| Difficulty walking ([HP:0002355](https://hpo.jax.org/app/browse/term/HP:0002355)) | REEP1 |
|  | RTN2 |
|  | SPG21 |
|  | AP4E1 |
|  | AP4B1 |
|  | SPG11 |
|  | AP4S1 |
|  | KIAA0196 |
|  | BSCL2 |
|  | GJC2 |
| Exercise intolerance ([HP:0003546](https://hpo.jax.org/app/browse/term/HP:0003546)) | PYGM |
|  | PFKM |
|  | PGAM2 |
|  | PHKA1 |
|  | MYH6 |
|  | ENO3 |
|  | PHKA2 |
|  | ACTA1 |
|  | PHKB |
|  | PHKG2 |
| Fatigue ([HP:0012378)](https://hpo.jax.org/app/browse/term/HP:0012378) | CTNNB1 |
|  | NF1 |
|  | SMAD4 |
|  | KRAS |
|  | RET |
|  | PIK3CA |
|  | BRCA1 |
|  | KIT |
|  | ERBB4 |
|  | PMS2 |
| Impaired ability to dress oneself ([HP:0031060](https://hpo.jax.org/app/browse/term/HP:0031060)) | - |
| Impairment of activities of daily living ([HP:0031058](https://hpo.jax.org/app/browse/term/HP:0031058)) | KIF5A |
|  | KIAA0196 |
|  | RTN2 |
|  | NIPA1 |
|  | SPG11 |
|  | SPAST |
|  | ATL1 |
|  | GJC2 |
|  | PLP1 |
|  | GABRA1 |
| Night sweats ([HP:0030166](https://hpo.jax.org/app/browse/term/HP:0030166)) | - |
| Postexertional malaise ([HP:0030973](https://hpo.jax.org/app/browse/term/HP:0030973)) | SLC2A9 |
|  | COL9A1 |
|  | SCN5A |
|  | NPPA |
|  | SLC22A12 |
|  | COL9A3 |
|  | PYGM |
|  | SVIL |
| Shivering ([HP:0025144](https://hpo.jax.org/app/browse/term/HP:0025144)) | CHRNA2 |
|  | CHRNB2 |
|  | CHRNA4 |
|  | CHRNB4 |
|  | CHRNA7 |
|  | CHRNA9 |
|  | CHRNA3 |
|  | CHRNB3 |
|  | CHRNB1 |
|  | KCNT1 |
| Stiff neck ([HP:0025258](https://hpo.jax.org/app/browse/term/HP:0025258)) | - |
| Weight loss ([HP:0001824](https://hpo.jax.org/app/browse/term/HP:0001824)) | FANCD2 |
|  | RAD51 |
|  | FANCM |
|  | FANCA |
|  | FANCC |
|  | FANCL |
|  | FANCG |
|  | UBE2T |
|  | FANCB |
|  | FANCF |
| Xerostomia ([HP:0000217](https://hpo.jax.org/app/browse/term/HP:0000217)) | C9orf72 |
|  | FUS |
|  | ATXN2 |
|  | MATR3 |
|  | NEK1 |
|  | NEFH |
|  | FIG4 |
|  | TAF15 |
|  | CCNF |
|  | GLE1 |

**Cardiovascular-finding**

| **Symptom (HPO ID)** | **Hub Genes (ranked)** |
| --- | --- |
| Abnormal heart morphology ([HP:0001627](https://hpo.jax.org/app/browse/term/HP:0001627)) | DRC1 |
|  | CCDC65 |
|  | HYDIN |
|  | RSPH4A |
|  | RSPH1 |
|  | CCDC103 |
|  | HEATR2 |
|  | DYX1C1 |
|  | DNAI2 |
|  | CCDC40 |
| Abnormal heart rate variability ([HP:0031860](https://hpo.jax.org/app/browse/term/HP:0031860)) | ZIC2 |
|  | SIX3 |
|  | SHH |
|  | GLI2 |
|  | CDON |
|  | FGF8 |
|  | NODAL |
|  | TGIF1 |
|  | PHOX2B |
|  | SMC1A |
| Abnormal left ventricular function ([HP:0005162](https://hpo.jax.org/app/browse/term/HP:0005162)) | MYH6 |
|  | TNNI3 |
|  | NKX2-5 |
|  | GATA4 |
|  | SCN5A |
|  | NPPA |
|  | MYH7 |
|  | TBX20 |
|  | ACTA2 |
|  | LDB3 |
| Abnormal pericardium morphology ([HP:0001697](https://hpo.jax.org/app/browse/term/HP:0001697)) | CTLA4 |
|  | PTPN22 |
|  | HLA-DRB1 |
|  | IL23R |
|  | MYBPC3 |
|  | CCBE1 |
|  | FAS |
|  | FCGR2B |
|  | ABCC9 |
|  | PTPN14 |
| Bradycardia ([HP:0001662](https://hpo.jax.org/app/browse/term/HP:0001662)) | SCN5A |
|  | ANK2 |
|  | CAV3 |
|  | CASQ2 |
|  | KCNE2 |
|  | HCN4 |
|  | KCNJ5 |
|  | SCN10A |
|  | TRDN |
|  | CACNA1D |
| Increased circulating troponin I concentration ([HP:0410173](https://hpo.jax.org/app/browse/term/HP:0410173)) | SVIL |
|  | ACTB |
|  | KDM1A |
|  | ACTG1 |
|  | FLOT2 |
|  | FLOT1 |
|  | AR |
|  | NEB |
|  | POTEF |
|  | KIF14 |
| Increased circulating troponin T concentration ([HP:0410174](https://hpo.jax.org/app/browse/term/HP:0410174)) | - |
| Hypertension ([HP:0000822](https://hpo.jax.org/app/browse/term/HP:0000822)) | BBS1 |
|  | BBS2 |
|  | MKS1 |
|  | SDCCAG8 |
|  | BBS5 |
|  | BBS7 |
|  | BBS12 |
|  | BBS10 |
|  | WDPCP |
|  | CCDC28B |
| Hypotension ([HP:0002615](https://hpo.jax.org/app/browse/term/HP:0002615)) | REN |
|  | KCNJ1 |
|  | SCNN1A |
|  | NR3C2 |
|  | SLC12A3 |
|  | CLCNKB |
|  | SLC12A1 |
|  | ACE |
|  | LHX4 |
|  | POU1F1 |
| Myocarditis ([HP:0012819](https://hpo.jax.org/app/browse/term/HP:0012819)) | - |
| Pericardial effusion ([HP:0001698](https://hpo.jax.org/app/browse/term/HP:0001698)) | MYBPC3 |
|  | CCBE1 |
|  | HLA-DRB1 |
|  | CLCNKB |
|  | ABCC9 |
|  | ADAMTS3 |
|  | PTPN14 |
|  | ENPP1 |
|  | SLC12A3 |
|  | ABCC6 |
| Reduced ejection fraction ([HP:0012664](https://hpo.jax.org/app/browse/term/HP:0012664)) | GTPBP3 |
|  | POLG2 |
|  | NPPA |
|  | C10orf2 |
|  | RPL3L |
|  | SCN5A |
|  | FHOD3 |
|  | PPCS |
|  | COA6 |
| Tachycardia ([HP:0001649](https://hpo.jax.org/app/browse/term/HP:0001649)) | SCN5A |
|  | HCN4 |
|  | KCND3 |
|  | CACNB2 |
|  | KCNE3 |
|  | SCN1B |
|  | ABCC9 |
|  | SCN3B |
|  | RYR2 |
|  | KCNJ5 |
| Venous thrombosis ([HP:0004936](https://hpo.jax.org/app/browse/term/HP:0004936)) | CTNNB1 |
|  | CASR |
|  | PRSS3P2 |
|  | SMAD4 |
|  | CTLA4 |
|  | PRSS1 |
|  | CTRC |
|  | GNAQ |
|  | MET |
|  | PTPN22 |

**Cardiovascular-symptom**

| **Symptom (HPO ID)** | **Hub Genes (ranked)** |
| --- | --- |
| Angina pectoris ([HP:0001681](https://hpo.jax.org/app/browse/term/HP:0001681)) | ABCG5 |
|  | ABCG8 |
|  | CYP27A1 |
|  | ABCC6 |
|  | LIPC |
|  | ENPP1 |
|  | PTPN22 |
|  | CTLA4 |
|  | ZMPSTE24 |
|  | GLA |
| Arrhythmia ([HP:0011675](https://hpo.jax.org/app/browse/term/HP:0011675)) | MYH6 |
|  | ACTN2 |
|  | TNNI3 |
|  | MYL2 |
|  | MYBPC3 |
|  | CSRP3 |
|  | MYH7 |
|  | RYR2 |
|  | LDB3 |
|  | MYL3 |
| Palpitations ([HP:0001962](https://hpo.jax.org/app/browse/term/HP:0001962)) | SCN5A |
|  | MYH6 |
|  | MYL2 |
|  | NKX2-5 |
|  | NPPA |
|  | TBX20 |
|  | GATA4 |
|  | MYL3 |
|  | CAV3 |
|  | KCND3 |
| Stroke ([HP:0001297](https://hpo.jax.org/app/browse/term/HP:0001297)) | NKX2-5 |
|  | MYH6 |
|  | TNNI3 |
|  | SCN5A |
|  | NPPA |
|  | GATA4 |
|  | TBX20 |
|  | MYBPC3 |
|  | ACTA2 |
|  | COL3A1 |
| Syncope ([HP:0001279](https://hpo.jax.org/app/browse/term/HP:0001279)) | SCN5A |
|  | HCN4 |
|  | KCND3 |
|  | CACNB2 |
|  | KCNE2 |
|  | KCNJ5 |
|  | SCN1B |
|  | ANK2 |
|  | KCNE3 |
|  | RYR2 |

**gi-finding**

| **Symptom (HPO ID)** | **Hub Genes (ranked)** |
| --- | --- |
| Abnormal pancreas morphology ([HP:0012090](https://hpo.jax.org/app/browse/term/HP:0012090)) | RPGRIP1L |
|  | TMEM216 |
|  | MKS1 |
|  | TCTN2 |
|  | TCTN3 |
|  | CC2D2A |
|  | B9D2 |
|  | B9D1 |
|  | TMEM231 |
|  | NPHP3 |
| Gastric ulcer ([HP:0002592](https://hpo.jax.org/app/browse/term/HP:0002592)) | ARID1B |
|  | SMARCA4 |
|  | SMARCC1 |
|  | SMARCA2 |
|  | SMARCB1 |
|  | SMARCC2 |
|  | SMARCD1 |
|  | SMARCE1 |
|  | ACTL6A |
|  | SS18 |
| Gastroesophageal reflux ([HP:0002020](https://hpo.jax.org/app/browse/term/HP:0002020)) | GRIN1 |
|  | GRIN2B |
|  | CAMK2A |
|  | GABRG2 |
|  | GABRB2 |
|  | NRXN1 |
|  | CACNA1B |
|  | CAMK2B |
|  | ERBB4 |
|  | HRAS |
| Gastroparesis ([HP:0002578](https://hpo.jax.org/app/browse/term/HP:0002578)) | SNRPN |
|  | MAGEL2 |
|  | OCA2 |
|  | NDN |
|  | C10orf2 |
|  | POLG2 |
|  | DNAJC6 |
|  | ACTG2 |
|  | MGAT2 |
|  | TMEM70 |
| Hepatic steatosis ([HP:0001397](https://hpo.jax.org/app/browse/term/HP:0001397)) | ACADS |
|  | ETFDH |
|  | CPT2 |
|  | HADH |
|  | SLC25A20 |
|  | ACOX1 |
|  | ACAD9 |
|  | SLC22A5 |
|  | PLIN1 |
|  | CIDEC |
| Hepatitis ([HP:0012115](https://hpo.jax.org/app/browse/term/HP:0012115)) | SYK |
|  | PIK3CA |
|  | CD40LG |
|  | CD79B |
|  | CD79A |
|  | CD3D |
|  | CD3E |
|  | BTK |
|  | FASLG |
|  | FAS |
| Hepatomegaly ([HP:0002240](https://hpo.jax.org/app/browse/term/HP:0002240)) | CTLA4 |
|  | CD40LG |
|  | CD3E |
|  | PRF1 |
|  | ZAP70 |
|  | CD3D |
|  | CD19 |
|  | IL2RA |
|  | RAG1 |
|  | RAG2 |
| Malnutrition ([HP:0004395](https://hpo.jax.org/app/browse/term/HP:0004395)) | MSX1 |
|  | PVRL1 |
|  | IRF6 |
|  | SBDS |
|  | SLC5A1 |
|  | SLC7A7 |
|  | EFTUD1 |
|  | HLA-DQA1 |
|  | ACTG2 |
|  | CRLF1 |
| Splenomegaly ([HP:0001744](https://hpo.jax.org/app/browse/term/HP:0001744)) | CD40LG |
|  | CD19 |
|  | CTLA4 |
|  | IL2RA |
|  | ZAP70 |
|  | CD3E |
|  | CD3D |
|  | PRF1 |
|  | RAG1 |
|  | RAG2 |

**gi-symptom**

| **Symptom (HPO ID)** | **Hub Genes (ranked)** |
| --- | --- |
| Abdominal pain ([HP:0002027](https://hpo.jax.org/app/browse/term/HP:0002027)) | KRAS |
|  | SMAD4 |
|  | CTNNB1 |
|  | ESR1 |
|  | PIK3CA |
|  | RET |
|  | KIT |
|  | NF1 |
|  | BRCA1 |
|  | CDKN1B |
| Abdominal symptom ([HP:0011458](https://hpo.jax.org/app/browse/term/HP:0011458)) | SCN2A |
|  | SCN1A |
|  | SCN8A |
|  | SCN9A |
|  | SCN5A |
|  | SCN11A |
|  | SCN10A |
|  | SCN1B |
|  | SCN4A |
|  | SCN3A |
| Anorexia ([HP:0002039](https://hpo.jax.org/app/browse/term/HP:0002039)) | BRCA1 |
|  | SMAD4 |
|  | KRAS |
|  | CDKN1B |
|  | PTPN22 |
|  | CD3D |
|  | CD3E |
|  | SYK |
|  | MEN1 |
|  | PALB2 |
| Bowel incontinence ([HP:0002607](https://hpo.jax.org/app/browse/term/HP:0002607)) | TBX1 |
|  | VANGL1 |
|  | RTN2 |
|  | FUZ |
|  | LRIG2 |
|  | PLP1 |
|  | HPSE2 |
|  | GP1BB |
|  | C19orf12 |
|  | COMT |
| Constipation ([HP:0002019](https://hpo.jax.org/app/browse/term/HP:0002019)) | SHH |
|  | FGF8 |
|  | OTX2 |
|  | SIX3 |
|  | SOX3 |
|  | ZIC2 |
|  | GLI2 |
|  | FOXG1 |
|  | SOX10 |
|  | LHX3 |
| Diarrhea ([HP:0002014](https://hpo.jax.org/app/browse/term/HP:0002014)) | CTLA4 |
|  | CD19 |
|  | CD79A |
|  | CD79B |
|  | CD3E |
|  | ZAP70 |
|  | CD3D |
|  | RAG2 |
|  | RAG1 |
|  | CD40LG |
| Nausea ([HP:0002018](https://hpo.jax.org/app/browse/term/HP:0002018)) | RET |
|  | NF1 |
|  | SDHC |
|  | SDHAF2 |
|  | MAX |
|  | KIF1B |
|  | ESR1 |
|  | MEN1 |
|  | KIT |
|  | KCNJ5 |
| Poor appetite ([HP:0004396](https://hpo.jax.org/app/browse/term/HP:0004396)) | PALB2 |
|  | BRCA1 |
|  | SMAD4 |
|  | KRAS |
|  | ACTA1 |
|  | PALLD |
|  | TPM3 |
|  | MYL2 |
|  | NAGS |
|  | SLC25A13 |
| Vomiting ([HP:0002013](https://hpo.jax.org/app/browse/term/HP:0002013)) | NDUFS3 |
|  | NDUFAF4 |
|  | NDUFA1 |
|  | NDUFA6 |
|  | NDUFB10 |
|  | NDUFB11 |
|  | NDUFS1 |
|  | TIMMDC1 |
|  | CTNNB1 |
|  | SHH |

**Pulmonary-finding**

| **Symptom (HPO ID)** | **Hub Genes (ranked)** |
| --- | --- |
| Abnormality on pulmonary function testing ([HP:0030878](https://hpo.jax.org/app/browse/term/HP:0030878)) | RAPSN |
|  | CHRNA1 |
|  | DOK7 |
|  | MUSK |
|  | SCN4A |
|  | CHRND |
|  | CHRNB1 |
|  | CHRNE |
|  | CTLA4 |
|  | CD19 |
| Airway obstruction ([HP:0006536](https://hpo.jax.org/app/browse/term/HP:0006536)) | CCDC65 |
|  | DRC1 |
|  | HYDIN |
|  | RSPH4A |
|  | RSPH9 |
|  | RSPH1 |
|  | CCDC103 |
|  | HEATR2 |
|  | DYX1C1 |
|  | DNAI2 |
| Decreased DLCO ([HP:0045051](https://hpo.jax.org/app/browse/term/HP:0045051)) | - |
| Ground-glass opacification ([HP:0025179](https://hpo.jax.org/app/browse/term/HP:0025179)) | TERT |
|  | RTEL1 |
|  | OBFC1 |
|  | ABCA3 |
|  | DHX36 |
|  | NKX2-1 |
|  | DSP |
|  | STON1 |
|  | BMPR2 |
| Hypoxemia ([HP:0012418](https://hpo.jax.org/app/browse/term/HP:0012418)) | NKX2-1 |
|  | GATA6 |
|  | TPM3 |
|  | MYL2 |
|  | ACTA1 |
|  | HLA-DRB1 |
|  | ZFPM2 |
|  | ABCA3 |
|  | BTNL2 |
|  | RHAG |
| Oxygen desaturation on exertion ([HP:0030874](https://hpo.jax.org/app/browse/term/HP:0030874)) | NKX2-1 |
|  | PAX8 |
|  | NAPSA |
|  | KRT7 |
|  | TG |
|  | SMAD3 |
|  | NCOA1 |
|  | FOXA2 |
|  | FOXA1 |
|  | SFTPB |
| Pleuritis ([HP:0002102](https://hpo.jax.org/app/browse/term/HP:0002102)) | CTLA4 |
|  | PTPN22 |
|  | FCGR2B |
|  | IL23R |
|  | FAS |
|  | MEFV |
|  | TREX1 |
| Pulmonary embolism ([HP:0002204](https://hpo.jax.org/app/browse/term/HP:0002204)) | SMAD4 |
|  | GDF2 |
|  | GNAQ |
|  | CBSL |
|  | ENSP00000381231 |
|  | COIL |
|  | TGS1 |
|  | MMACHC |
|  | MEFV |
|  | KCNJ5 |
| Reduced forced expiratory volume in one second ([HP:0032342](https://hpo.jax.org/app/browse/term/HP:0032342)) | RTEL1 |
|  | DNA2 |
|  | WRN |
|  | RAD51 |
|  | PIF1 |
|  | BLM |
|  | EXO1 |
|  | TERF1 |
|  | MMS19 |
|  | FAM96B |
| Reduced FEV1/FVC ratio ([HP:0030877](https://hpo.jax.org/app/browse/term/HP:0030877)) | - |
| Reduced forced vital capacity ([HP:0032341](https://hpo.jax.org/app/browse/term/HP:0032341)) | MYH7 |
|  | ACTA1 |
|  | UNC45B |
|  | TPM3 |
|  | FKRP |
|  | SLC25A21 |
|  | MCIDAS |
|  | GGPS1 |
|  | RTEL1 |
|  | SYT2 |
| Reduced vital capacity ([HP:0002792](https://hpo.jax.org/app/browse/term/HP:0002792)) | RAPSN |
|  | CHRNA1 |
|  | CHRND |
|  | CHRNE |
|  | CHRNB1 |
|  | DOK7 |
|  | MUSK |
|  | SCN4A |
|  | ACTA1 |
|  | MYH7 |
| Restrictive ventilatory defect ([HP:0002091](https://hpo.jax.org/app/browse/term/HP:0002091)) | SCN4A |
|  | CHRNE |
|  | RAPSN |
|  | CHRND |
|  | CHRNA1 |
|  | CHRNB1 |
|  | DOK7 |
|  | MUSK |
|  | CTLA4 |
|  | CD19 |

**Pulmonary-imaging**

| **Symptom (HPO ID)** | **Hub Genes (ranked)** |
| --- | --- |
| Abnormal pulmonary thoracic imaging finding ([HP:0031983](https://hpo.jax.org/app/browse/term/HP:0031983)) | DRC1 |
|  | CCDC65 |
|  | HYDIN |
|  | RSPH4A |
|  | RSPH1 |
|  | CCDC103 |
|  | HEATR2 |
|  | DYX1C1 |
|  | DNAI2 |
|  | CCDC40 |
| Atelectasis ([HP:0100750](https://hpo.jax.org/app/browse/term/HP:0100750)) | CCDC65 |
|  | DRC1 |
|  | HYDIN |
|  | RSPH4A |
|  | RSPH1 |
|  | CCDC103 |
|  | HEATR2 |
|  | DYX1C1 |
|  | DNAI2 |
|  | CCDC40 |
| Bronchiectasis ([HP:0002110](https://hpo.jax.org/app/browse/term/HP:0002110)) | DRC1 |
|  | CCDC65 |
|  | HYDIN |
|  | RSPH4A |
|  | RSPH1 |
|  | CCDC103 |
|  | HEATR2 |
|  | DYX1C1 |
|  | DNAI2 |
|  | CCDC40 |
| Centrilobular ground-glass opacification on pulmonary HRCT ([HP:0025180](https://hpo.jax.org/app/browse/term/HP:0025180)) | BMPR2 |
|  | ACVR1 |
|  | SMAD6 |
|  | BMP4 |
|  | BMP2 |
|  | BMP10 |
|  | BMP7 |
|  | GDF5 |
|  | BMP6 |
|  | GDF2 |
| Interlobular septal thickening ([HP:0030879](https://hpo.jax.org/app/browse/term/HP:0030879)) | BMPR2 |
|  | ACVR1 |
|  | SMAD6 |
|  | BMP4 |
|  | BMP2 |
|  | BMP10 |
|  | BMP7 |
|  | GDF5 |
|  | BMP6 |
|  | GDF2 |
| Parenchymal consolidation ([HP:0032177](https://hpo.jax.org/app/browse/term/HP:0032177)) | NKX2-1 |
|  | NAPSA |
|  | PAX8 |
|  | KRT7 |
|  | TG |
|  | SMAD3 |
|  | NCOA1 |
|  | FOXA2 |
|  | FOXA1 |
|  | SFTPB |
| Pleural thickening ([HP:0031944](https://hpo.jax.org/app/browse/term/HP:0031944)) | CARD10 |
|  | PRKCQ |
|  | BCL10 |
|  | MALT1 |
|  | CARD11 |
|  | PRKCB |
|  | TRAF6 |
|  | IKBKG |
|  | CARD14 |
|  | RRNAD1 |
| Pulmonary bulla ([HP:0032446](https://hpo.jax.org/app/browse/term/HP:0032446)) | - |
| Pulmonary fibrosis ([HP:0002206](https://hpo.jax.org/app/browse/term/HP:0002206)) | CTLA4 |
|  | PTPN22 |
|  | HLA-DRB1 |
|  | NR5A1 |
|  | TERT |
|  | BMP15 |
|  | NKX2-1 |
|  | BTNL2 |
|  | OBFC1 |
|  | FASLG |
| Pulmonary interstitial high-resolution computed tomography abnormality ([HP:0025389](https://hpo.jax.org/app/browse/term/HP:0025389)) | TERT |
|  | OBFC1 |
|  | RTEL1 |
|  | NKX2-1 |
|  | ABCA3 |
|  | DHX36 |
|  | COL3A1 |
|  | HLA-DRB1 |
|  | BMPR2 |
|  | DSP |
| Pulmonary opacity ([HP:0031457](https://hpo.jax.org/app/browse/term/HP:0031457)) | TERT |
|  | OBFC1 |
|  | RTEL1 |
|  | NKX2-1 |
|  | ABCA3 |
|  | DHX36 |
|  | DSP |
|  | COL3A1 |
|  | BMPR2 |
| Reticular pattern on pulmonary HRCT ([HP:0025390](https://hpo.jax.org/app/browse/term/HP:0025390)) | TERT |
|  | OBFC1 |
|  | RTEL1 |
|  | DHX36 |
|  | ABCA3 |
|  | DSP |

**Neuropsychiatric-behavioural**

| **Symptom (HPO ID)** | **Hub Genes (ranked)** |
| --- | --- |
| Anxiety ([HP:0000739](https://hpo.jax.org/app/browse/term/HP:0000739)) | GPR98 |
|  | USH1C |
|  | USH2A |
|  | USH1G |
|  | PCDH15 |
|  | DFNB31 |
|  | CIB2 |
|  | CDH23 |
|  | CLRN1 |
|  | PDZD7 |
| Apathy ([HP:0000741](https://hpo.jax.org/app/browse/term/HP:0000741)) | ZIC2 |
|  | SIX3 |
|  | SHH |
|  | GLI2 |
|  | FGF8 |
|  | CDON |
|  | MAPT |
|  | NODAL |
|  | C9orf72 |
|  | ATXN10 |
| Attention deficit hyperactivity disorder ([HP:0007018](https://hpo.jax.org/app/browse/term/HP:0007018)) | GABRG2 |
|  | GABRA1 |
|  | SCN1A |
|  | KCNA2 |
|  | KCNT1 |
|  | SCN8A |
|  | GRIN2A |
|  | SLC6A1 |
|  | CACNA1B |
|  | ATP1A3 |
| Auditory hallucinations ([HP:0008765](https://hpo.jax.org/app/browse/term/HP:0008765)) | - |
| Behavioral abnormalities ([HP:0000708](https://hpo.jax.org/app/browse/term/HP:0000708)) | GUCA1B |
|  | PRPH2 |
|  | NR2E3 |
|  | CRX |
|  | CNGA1 |
|  | NRL |
|  | PDE6G |
|  | RPGR |
|  | RDH12 |
|  | TULP1 |
| Delusions ([HP:0000746](https://hpo.jax.org/app/browse/term/HP:0000746)) | COMT |
|  | DGCR2 |
|  | ENSP00000331681 |
|  | TBX1 |
|  | EIF2B2 |
|  | DAOA |
|  | EIF2B3 |
|  | JPH3 |
|  | EIF2B1 |
|  | DGCR14 |
| Hallucinations ([HP:0000738](https://hpo.jax.org/app/browse/term/HP:0000738)) | GPR98 |
|  | USH1C |
|  | USH2A |
|  | DFNB31 |
|  | CIB2 |
|  | CDH23 |
|  | USH1G |
|  | PCDH15 |
|  | CLRN1 |
|  | PDZD7 |
| Impulsivity ([HP:0100710](https://hpo.jax.org/app/browse/term/HP:0100710)) | SCN1A |
|  | SCN2A |
|  | KCNA2 |
|  | GABRG2 |
|  | GABRA1 |
|  | SCN8A |
|  | PCDH19 |
|  | GRIN2A |
|  | SCN1B |
|  | CACNA1B |
| Irritability ([HP:0000737](https://hpo.jax.org/app/browse/term/HP:0000737)) | SHH |
|  | SIX3 |
|  | FGF8 |
|  | ZIC2 |
|  | PAX2 |
|  | GLI2 |
|  | FOXG1 |
|  | NPHS1 |
|  | TH |
|  | AQP2 |
| Obsessive-compulsive disorder ([HP:0000722](https://hpo.jax.org/app/browse/term/HP:0000722)) | GABRG2 |
|  | SCN1A |
|  | SCN2A |
|  | GABRA1 |
|  | GABRD |
|  | SCN1B |
|  | GRIA2 |
|  | SCN9A |
|  | PCDH19 |
|  | EP300 |
| Panic attack ([HP:0025269](https://hpo.jax.org/app/browse/term/HP:0025269)) | MAX |
|  | NF1 |
|  | SDHC |
|  | KIF1B |
|  | SDHAF2 |
|  | RET |
|  | EP300 |
|  | BPTF |
|  | CREBBP |
|  | TRIP12 |
| Paranoia ([HP:0011999](https://hpo.jax.org/app/browse/term/HP:0011999)) | ENSP00000331681 |
|  | DGCR2 |
|  | TBX1 |
|  | DGCR14 |
|  | CPOX |
|  | DGCR8 |
|  | HMBS |
|  | TIMM8A |
|  | VPS13A |
|  | IMPA1 |
| Personality disorders ([HP:0012075](https://hpo.jax.org/app/browse/term/HP:0012075)) | - |
| Phonophobia ([HP:0002183](https://hpo.jax.org/app/browse/term/HP:0002183)) | SGCE |
|  | SCN1A |
|  | KCNT1 |
|  | KCTD17 |
|  | TOR1A |
|  | ENSP00000381231 |
|  | CYP27A1 |
|  | CUX2 |
|  | XK |
|  | SPG21 |
| Polydipsia ([HP:0001959](https://hpo.jax.org/app/browse/term/HP:0001959)) | EP300 |
|  | CREBBP |
|  | ESR1 |
|  | MLXIPL |
|  | ELN |
|  | NCF1 |
|  | DNAJC30 |
| Short attention span ([HP:0000736](https://hpo.jax.org/app/browse/term/HP:0000736)) | SCN1A |
|  | SCN2A |
|  | GABRG2 |
|  | KCNT1 |
|  | GABRA1 |
|  | SCN8A |
|  | KCNA2 |
|  | PCDH19 |
|  | SLC6A1 |
|  | GRIN2A |
| Visual hallucinations ([HP:0002367](https://hpo.jax.org/app/browse/term/HP:0002367)) | DNAJC5 |
|  | FBXO7 |
|  | GIGYF2 |
|  | SNCB |
|  | NPRL3 |
|  | FIG4 |
|  | LGI1 |
|  | RELN |
|  | EPM2A |

**Neuropsychiatric-cognitivedysfunction**

| **Symptom (HPO ID)** | **Hub Genes (ranked)** |
| --- | --- |
| Abnormality of higher mental function ([HP:0011446](https://hpo.jax.org/app/browse/term/HP:0011446)) | RPGR |
|  | IMPDH1 |
|  | RDH12 |
|  | CRX |
|  | SPATA7 |
|  | PRPH2 |
|  | TULP1 |
|  | CNGA1 |
|  | NR2E3 |
|  | GUCA1B |
| Agnosia ([HP:0010524](https://hpo.jax.org/app/browse/term/HP:0010524)) | TREM2 |
|  | MAPT |
|  | PIK3CA |
|  | PMS2 |
|  | ABCA7 |
|  | KRAS |
|  | C9orf72 |
|  | FAN1 |
|  | ATP6AP2 |
| Bradykinesia ([HP:0002067](https://hpo.jax.org/app/browse/term/HP:0002067)) | SCN1A |
|  | GABRG2 |
|  | SCN2A |
|  | GABRA1 |
|  | SCN1B |
|  | GABRD |
|  | ATP1A3 |
|  | PCDH19 |
|  | SCN9A |
|  | ATXN3 |
| Bradyphrenia ([HP:0031843](https://hpo.jax.org/app/browse/term/HP:0031843)) | - |
| Cognitive impairment ([HP:0100543](https://hpo.jax.org/app/browse/term/HP:0100543)) | SCN2A |
|  | SCN1A |
|  | KCNA2 |
|  | KCNT1 |
|  | GABRG2 |
|  | KCNC1 |
|  | GABRA1 |
|  | CACNA1B |
|  | SCN8A |
|  | GABRD |
| Confusion ([HP:0001289](https://hpo.jax.org/app/browse/term/HP:0001289)) | ERCC4 |
|  | ERCC2 |
|  | NAGS |
|  | SCN1A |
|  | SLC25A15 |
|  | ERCC5 |
|  | MEN1 |
|  | MMACHC |
|  | CHRNA2 |
|  | TREM2 |
| Encephalopathy ([HP:0001298](https://hpo.jax.org/app/browse/term/HP:0001298)) | GABRA1 |
|  | GABRG2 |
|  | SCN2A |
|  | SLC6A1 |
|  | GRIN2A |
|  | SCN1A |
|  | GRIN2B |
|  | KCNA2 |
|  | NRXN1 |
|  | GABRB2 |
| Diminished ability to concentrate ([HP:0031987](https://hpo.jax.org/app/browse/term/HP:0031987)) | - |

**Neuropsychiatric-emotion-mood**

| **Symptom (HPO ID)** | **Hub Genes (ranked)** |
| --- | --- |
| Abnormal emotion/affect behavior ([HP:0100851](https://hpo.jax.org/app/browse/term/HP:0100851)) | GRIA2 |
|  | SLC6A1 |
|  | GRIN2A |
|  | GRIN1 |
|  | GRIA3 |
|  | CAMK2A |
|  | GABRG2 |
|  | GABRA1 |
|  | GRIA4 |
|  | GRIN2B |
| Abnormal fear/anxiety-related behavior ([HP:0100852](https://hpo.jax.org/app/browse/term/HP:0100852)) | GPR98 |
|  | USH1C |
|  | CLRN1 |
|  | PDZD7 |
|  | USH2A |
|  | USH1G |
|  | PCDH15 |
|  | DFNB31 |
|  | CIB2 |
|  | CDH23 |
| Aggressive behavior ([HP:0000718](https://hpo.jax.org/app/browse/term/HP:0000718)) | EP300 |
|  | CREBBP |
|  | ARID1A |
|  | DNMT3A |
|  | KDM5C |
|  | SIN3A |
|  | SETD2 |
|  | SMARCC2 |
|  | BPTF |
|  | ARID1B |
| Depression ([HP:0000716](https://hpo.jax.org/app/browse/term/HP:0000716)) | GPR98 |
|  | USH1G |
|  | PCDH15 |
|  | DFNB31 |
|  | CIB2 |
|  | CDH23 |
|  | USH1C |
|  | CLRN1 |
|  | PDZD7 |
|  | USH2A |
| Emotional lability ([HP:0000712](https://hpo.jax.org/app/browse/term/HP:0000712)) | NDUFS3 |
|  | NDUFS1 |
|  | NDUFA2 |
|  | NDUFA9 |
|  | NDUFAF2 |
|  | NDUFAF6 |
|  | COX15 |
|  | TACO1 |
|  | PDHA1 |
|  | C9orf72 |
| Euphoria ([HP:0031844](https://hpo.jax.org/app/browse/term/HP:0031844)) | - |
| Mania ([HP:0100754](https://hpo.jax.org/app/browse/term/HP:0100754)) | PRKACA |
|  | TBX1 |
|  | PRKAR1A |
|  | PDE11A |
|  | C10orf2 |
|  | GP1BB |
|  | POLG2 |
|  | COMT |
|  | RPS6KA3 |
|  | SLC25A13 |
| Suicidal ideation ([HP:0031589](https://hpo.jax.org/app/browse/term/HP:0031589)) | KCNT1 |
|  | CHRNB2 |
|  | CHRNA4 |
|  | CHRNA2 |
|  | CYP27A1 |
|  | FIG4 |
|  | CDH23 |

**Neuropsychiatric-headache**

| **Symptom (HPO ID)** | **Hub Genes (ranked)** |
| --- | --- |
| Headache ([HP:0002315](https://hpo.jax.org/app/browse/term/HP:0002315)) | CTNNB1 |
|  | KRAS |
|  | ESR1 |
|  | KIT |
|  | RET |
|  | SMAD4 |
|  | TERT |
|  | PIK3CA |
|  | NF1 |
|  | CDKN1B |
| Migraine ([HP:0002076](https://hpo.jax.org/app/browse/term/HP:0002076)) | KRAS |
|  | PIK3CA |
|  | ESR1 |
|  | SMAD4 |
|  | NF1 |
|  | NOTCH3 |
|  | RELA |
|  | SCN1A |
|  | GRIN2A |
|  | KCNA1 |

**Neuropsychiatric-memory**

| **Symptom (HPO ID)** | **Hub Genes (ranked)** |
| --- | --- |
| Memory impairment ([HP:0002354](https://hpo.jax.org/app/browse/term/HP:0002354)) | MAPT |
|  | NF1 |
|  | KRAS |
|  | ATXN3 |
|  | PIK3CA |
|  | PRKACA |
|  | ATXN1 |
|  | C9orf72 |
|  | PRKAR1A |
|  | PRKCG |

**Neuropsychiatric-sleep**

| **Symptom (HPO ID)** | **Hub Genes (ranked)** |
| --- | --- |
| Insomnia ([HP:0100785](https://hpo.jax.org/app/browse/term/HP:0100785)) | NR1H4 |
|  | ATP8B1 |
|  | ABCB4 |
|  | ABCB11 |
|  | NAGS |
|  | CPOX |
|  | CLCNKB |
|  | MLXIPL |
|  | SLC25A13 |
|  | SLC12A3 |
| Maintenance insomnia ([HP:0031355](https://hpo.jax.org/app/browse/term/HP:0031355)) | - |
| Restless legs ([HP:0012452](https://hpo.jax.org/app/browse/term/HP:0012452)) | NEFL |
|  | UCHL1 |
|  | MFN2 |
|  | DNAJC6 |
|  | ATXN7 |
| Sleep apnea ([HP:0010535](https://hpo.jax.org/app/browse/term/HP:0010535)) | CTNNB1 |
|  | HRAS |
|  | EP300 |
|  | CREBBP |
|  | TERT |
|  | RUNX2 |
|  | TWIST1 |
|  | RET |
|  | USP7 |
|  | CHAT |
| Sleep disturbance ([HP:0002360](https://hpo.jax.org/app/browse/term/HP:0002360)) | GRIN2B |
|  | GRIN2A |
|  | GRIN1 |
|  | GRIA4 |
|  | CAMK2A |
|  | GRIA3 |
|  | GABRG2 |
|  | KCNA1 |
|  | SCN2A |
|  | CAMK2B |
| Sleep onset insomnia ([HP:0031354](https://hpo.jax.org/app/browse/term/HP:0031354)) | - |

**Neuropsychiatric-smell-taste**

| **Symptom (HPO ID)** | **Hub Genes (ranked)** |
| --- | --- |
| Abnormality of taste sensation ([HP:0000223](https://hpo.jax.org/app/browse/term/HP:0000223)) | - |
| Anosmia ([HP:0000458](https://hpo.jax.org/app/browse/term/HP:0000458)) | FGF8 |
|  | KAL1 |
|  | GNRH1 |
|  | TAC3 |
|  | FEZF1 |
|  | CHD7 |
|  | PROKR2 |
|  | KISS1R |
|  | IL17RD |
|  | KISS1 |
| Hypogeusia ([HP:0000224](https://hpo.jax.org/app/browse/term/HP:0000224)) | - |
| Hyposmia ([HP:0004409](https://hpo.jax.org/app/browse/term/HP:0004409)) | KAL1 |
|  | PROKR2 |
|  | FEZF1 |
|  | SOX10 |
|  | FGF8 |
|  | CHD7 |
|  | FGF17 |
|  | IL17RD |
|  | GIGYF2 |
|  | LZTFL1 |
| Abnormality of the sense of smell ([HP:0004408](https://hpo.jax.org/app/browse/term/HP:0004408)) | FGF8 |
|  | KAL1 |
|  | GNRH1 |
|  | TAC3 |
|  | FEZF1 |
|  | CHD7 |
|  | PROKR2 |
|  | KISS1R |
|  | IL17RD |
|  | KISS1 |

**Neuropsychiatric-speech-language**

| **Symptom (HPO ID)** | **Hub Genes (ranked)** |
| --- | --- |
| Anomic aphasia ([HP:0030784](https://hpo.jax.org/app/browse/term/HP:0030784)) | MAPT |
|  | TREM2 |
|  | C9orf72 |
|  | PLP1 |
| Aphasia ([HP:0002381](https://hpo.jax.org/app/browse/term/HP:0002381)) | MAPT |
|  | BPTF |
|  | GRIN2A |
|  | TRIP12 |
|  | SCN1A |
|  | PLP1 |
|  | C9orf72 |
|  | POLR1D |
|  | POLR1C |
|  | KRAS |
| Expressive aphasia ([HP:0002427](https://hpo.jax.org/app/browse/term/HP:0002427)) | C9orf72 |
|  | MAPT |
|  | TREM2 |
|  | NPRL3 |
|  | GJB1 |
| Neurological speech impairment ([HP:0002167](https://hpo.jax.org/app/browse/term/HP:0002167)) | DRC1 |
|  | CCDC65 |
|  | HYDIN |
|  | RSPH4A |
|  | RSPH1 |
|  | CCDC103 |
|  | HEATR2 |
|  | DYX1C1 |
|  | DNAI2 |
|  | CCDC40 |
| Slurred speech ([HP:0001350](https://hpo.jax.org/app/browse/term/HP:0001350)) | MAPT |
|  | KCNA1 |
|  | MOG |
|  | GRIN2A |
|  | DNAJC6 |
|  | KCND3 |
|  | ATXN1 |
|  | SETD2 |
|  | ACOX2 |
|  | HCRT |

**Neuropsychiatric-finding**

| **Symptom (HPO ID)** | **Hub Genes (ranked)** |
| --- | --- |
| Abnormal reflex ([HP:0031826](https://hpo.jax.org/app/browse/term/HP:0031826)) | RDH12 |
|  | NR2E3 |
|  | CRX |
|  | GUCA1B |
|  | PRPH2 |
|  | RPGR |
|  | TULP1 |
|  | PRPF3 |
|  | PDE6G |
|  | CNGA1 |
| Abnormality of movement ([HP:0100022](https://hpo.jax.org/app/browse/term/HP:0100022)) | RDH12 |
|  | NR2E3 |
|  | CRX |
|  | GUCA1B |
|  | PRPH2 |
|  | RPGR |
|  | TULP1 |
|  | CNGA1 |
|  | IMPDH1 |
|  | NRL |
| Ataxia ([HP:0001251](https://hpo.jax.org/app/browse/term/HP:0001251)) | B9D1 |
|  | CC2D2A |
|  | MKS1 |
|  | TMEM216 |
|  | RPGRIP1L |
|  | TCTN2 |
|  | NPHP1 |
|  | TMEM231 |
|  | TCTN3 |
|  | TCTN1 |
| Babinski sign ([HP:0003487](https://hpo.jax.org/app/browse/term/HP:0003487)) | REEP1 |
|  | RTN2 |
|  | SPG21 |
|  | SPG11 |
|  | KIAA0196 |
|  | NIPA1 |
|  | AP4B1 |
|  | AP4S1 |
|  | AP4E1 |
|  | SPAST |
| Dysarthria ([HP:0001260](https://hpo.jax.org/app/browse/term/HP:0001260)) | ATXN2 |
|  | SPG11 |
|  | C9orf72 |
|  | FUS |
|  | MATR3 |
|  | NEFH |
|  | SETX |
|  | NEK1 |
|  | FIG4 |
|  | ALS2 |
| Dysmetria ([HP:0001310](https://hpo.jax.org/app/browse/term/HP:0001310)) | ATXN7 |
|  | ATXN10 |
|  | PPP2R2B |
|  | ATXN2 |
|  | KCNC3 |
|  | ATXN1 |
|  | ATN1 |
|  | TGM6 |
|  | PRKCG |
|  | FXN |
| Dysphagia ([HP:0002015](https://hpo.jax.org/app/browse/term/HP:0002015)) | ATXN2 |
|  | C9orf72 |
|  | NEFH |
|  | FUS |
|  | SPG11 |
|  | ALS2 |
|  | NEK1 |
|  | MATR3 |
|  | SETX |
|  | FIG4 |
| Dystonia ([HP:0001332](https://hpo.jax.org/app/browse/term/HP:0001332)) | GRIN2B |
|  | GRIA2 |
|  | SCN2A |
|  | CACNA1B |
|  | KCNA1 |
|  | GRIN1 |
|  | GRIN2A |
|  | CAMK2A |
|  | GABRB2 |
|  | GRIK2 |
| Facial paralysis ([HP:0007209](https://hpo.jax.org/app/browse/term/HP:0007209)) | ATP1A2 |
|  | TCIRG1 |
|  | SCN1A |
|  | TNFSF11 |
|  | SH3TC2 |
| Frontal release signs ([HP:0000743](https://hpo.jax.org/app/browse/term/HP:0000743)) | - |
| Gait disturbance ([HP:0001288](https://hpo.jax.org/app/browse/term/HP:0001288)) | GRIA2 |
|  | GABRA1 |
|  | GABRG2 |
|  | GABRB2 |
|  | GRIN1 |
|  | CAMK2A |
|  | KCNA1 |
|  | GABRD |
|  | SLC12A5 |
|  | SLC6A1 |
| Hand muscle weakness ([HP:0030237](https://hpo.jax.org/app/browse/term/HP:0030237)) | RAPSN |
|  | CHRNA1 |
|  | CHRNE |
|  | CHRNB1 |
|  | CHRND |
|  | DOK7 |
|  | MUSK |
|  | SCN4A |
|  | COLQ |
|  | MFN2 |
| Hyperesthesia ([HP:0100963](https://hpo.jax.org/app/browse/term/HP:0100963)) | - |
| Hyperkinetic movements ([HP:0002487](https://hpo.jax.org/app/browse/term/HP:0002487)) | GRIN2A |
|  | GAD1 |
|  | CACNA1B |
|  | GRIN1 |
|  | SCN1A |
|  | GNAO1 |
|  | SLC6A3 |
|  | IQSEC2 |
|  | ALDH5A1 |
|  | PNKD |
| Hypotonia ([HP:0001252](https://hpo.jax.org/app/browse/term/HP:0001252)) | NDUFS3 |
|  | NDUFA9 |
|  | NDUFA2 |
|  | NDUFC2 |
|  | NDUFA6 |
|  | NDUFA12 |
|  | NDUFA1 |
|  | NDUFA8 |
|  | NDUFB10 |
|  | NDUFB11 |
| Muscle spasm ([HP:0003394](https://hpo.jax.org/app/browse/term/HP:0003394)) | C9orf72 |
|  | FUS |
|  | ATXN2 |
|  | MATR3 |
|  | NEK1 |
|  | FIG4 |
|  | NEFH |
|  | TAF15 |
|  | CCNF |
|  | GLE1 |
| Muscle weakness ([HP:0001324](https://hpo.jax.org/app/browse/term/HP:0001324)) | FKTN |
|  | FKRP |
|  | POMGNT1 |
|  | SCN4A |
|  | SCN1A |
|  | SCN2A |
|  | SCN8A |
|  | MYOT |
|  | SCN11A |
|  | SGCB |
| Orthostatic hypotension ([HP:0001278](https://hpo.jax.org/app/browse/term/HP:0001278)) | GIGYF2 |
|  | CHCHD2 |
|  | SPG11 |
|  | IL12RB1 |
|  | LEP |
|  | CYB561 |
|  | AAAS |
|  | IKBKAP |
|  | ARSA |
|  | CHRNA3 |
| Paresthesia ([HP:0003401](https://hpo.jax.org/app/browse/term/HP:0003401)) | CASR |
|  | CLCNKB |
|  | KCNJ1 |
|  | PIK3CA |
|  | KRAS |
|  | SLC12A3 |
|  | THPO |
|  | SLC12A1 |
|  | PTPN22 |
|  | GATA1 |
| Parkinsonism ([HP:0001300](https://hpo.jax.org/app/browse/term/HP:0001300)) | GABRG2 |
|  | SCN2A |
|  | GABRA1 |
|  | SCN1A |
|  | PCDH19 |
|  | ATP1A3 |
|  | SCN1B |
|  | KIF5A |
|  | C9orf72 |
|  | ATXN2 |
| Polyneuropathy ([HP:0001271](https://hpo.jax.org/app/browse/term/HP:0001271)) | PEX11B |
|  | PEX7 |
|  | PEX12 |
|  | SETX |
|  | GRM1 |
|  | MYOT |
|  | COQ7 |
|  | PDYN |
|  | ERCC6 |
|  | LDB3 |
| Rigidity ([HP:0002063](https://hpo.jax.org/app/browse/term/HP:0002063)) | SCN1A |
|  | SCN2A |
|  | KCNA2 |
|  | SCN8A |
|  | SCN9A |
|  | KCND3 |
|  | KCNA4 |
|  | SCN1B |
|  | KCNB1 |
|  | CACNA1B |
| Seizure ([HP:0001250](https://hpo.jax.org/app/browse/term/HP:0001250)) | CC2D2A |
|  | B9D1 |
|  | TCTN2 |
|  | TMEM231 |
|  | TMEM216 |
|  | B9D2 |
|  | MKS1 |
|  | RPGRIP1L |
|  | TCTN1 |
|  | TMEM138 |
| Skeletal muscle atrophy ([HP:0003202](https://hpo.jax.org/app/browse/term/HP:0003202)) | BBS5 |
|  | BBS7 |
|  | BBS1 |
|  | LZTFL1 |
|  | BBS2 |
|  | BBS10 |
|  | WDPCP |
|  | BBS12 |
|  | MKS1 |
|  | SDCCAG8 |
| Somatic sensory dysfunction ([HP:0003474](https://hpo.jax.org/app/browse/term/HP:0003474)) | BSCL2 |
|  | REEP1 |
|  | KIF5A |
|  | SPG11 |
|  | NIPA1 |
|  | KIAA0196 |
|  | RTN2 |
|  | SPAST |
|  | ATL1 |
|  | MPZ |
| Spasticity ([HP:0001257](https://hpo.jax.org/app/browse/term/HP:0001257)) | SPG11 |
|  | REEP1 |
|  | RTN2 |
|  | SPG21 |
|  | KIAA0196 |
|  | NIPA1 |
|  | AP4E1 |
|  | AP4B1 |
|  | AP4S1 |
|  | KIF5A |
| Tremor ([HP:0001337](https://hpo.jax.org/app/browse/term/HP:0001337)) | B9D1 |
|  | CC2D2A |
|  | TCTN2 |
|  | CEP41 |
|  | TCTN3 |
|  | MKS1 |
|  | TMEM216 |
|  | RPGRIP1L |
|  | TCTN1 |
|  | NPHP1 |
| Unilateral facial palsy ([HP:0012799](https://hpo.jax.org/app/browse/term/HP:0012799)) | - |

**All symptoms**

**Hub Genes (ranked)**

GNGT1

GNG12

GNB3

GNB4

GNG13

GNG8

GNG3

GNG7

GNG10

GNAI1
